# Ultrafast Sample Placement on Existing Trees (UShER) Empowers Real-Time Phylogenetics for the SARS-CoV-2 Pandemic

**DOI:** 10.1101/2020.09.26.314971

**Authors:** Yatish Turakhia, Bryan Thornlow, Angie S. Hinrichs, Nicola De Maio, Landen Gozashti, Robert Lanfear, David Haussler, Russell Corbett-Detig

## Abstract

As the SARS-CoV-2 virus spreads through human populations, the unprecedented accumulation of viral genome sequences is ushering a new era of “genomic contact tracing” – that is, using viral genome sequences to trace local transmission dynamics. However, because the viral phylogeny is already so large – and will undoubtedly grow many fold – placing new sequences onto the tree has emerged as a barrier to real-time genomic contact tracing. Here, we resolve this challenge by building an efficient, tree-based data structure encoding the inferred evolutionary history of the virus. We demonstrate that our approach improves the speed of phylogenetic placement of new samples and data visualization by orders of magnitude, making it possible to complete the placements under real-time constraints. Our method also provides the key ingredient for maintaining a fully-updated reference phylogeny. We make these tools available to the research community through the UCSC SARS-CoV-2 Genome Browser to enable rapid cross-referencing of information in new virus sequences with an ever-expanding array of molecular and structural biology data. The methods described here will empower research and genomic contact tracing for laboratories worldwide.

**Software Availability:** USHER is available to users through the UCSC Genome Browser at https://genome.ucsc.edu/cgi-bin/hgPhyloPlace. The source code and detailed instructions on how to compile and run UShER are available from https://github.com/yatisht/usher.

## Introduction

In the past year, the SARS-CoV-2 virus emerged from a presumably zoonotic source to spread across human populations worldwide^1–3^. Recent technological advances have enabled rapid and cost-efficient sequencing of the viral genome – over 1560 groups worldwide have generated 97,733 high coverage whole-genome SARS-CoV-2 sequences that are available on GISAID^4^ as of Sep. 24, 2020. These vast datasets and rapid sequencing turnaround times are enabling a type of “genomic contact tracing” where genetic similarities (or dissimilarity) between viral genomes isolated for different hosts carries important information about the transmission dynamics of the virus. For example, these data can be used to infer the number of unique introductions of the viral genome in a given area^5–11^ and to identify “transmission chains” among otherwise seemingly unrelated infections^12–16^.

Despite great potential, this unprecedented and ongoing accumulation of sequencing data is overwhelming existing systems for analysis and interpretation of viral transmission and evolutionary dynamics. In part, this is because typical phylogenetics applications accumulate all of the relevant sequence data before beginning phylogenetic inference. For genomic contact tracing to work effectively, each new viral genome sequence must be contextualized within the entire evolutionary history of the virus rapidly and accurately as it is collected. This could be accomplished by re-inferring the full phylogeny, but with current SARS-CoV-2 datasets, this takes more than a day even using powerful computational resources. Alternatively, new genome sequences could be contextualized by placing samples onto an existing “reference phylogeny” and several methods have been developed for this purpose^17–20^. These methods have been used to place new samples onto a phylogeny created from a small subset of available SARS-CoV-2 isolates^21^, and to provide regular updates to a global phylogeny of SARS-CoV-2^22^. Nonetheless, existing algorithms for placing sequences onto reference phylogenies are far too slow to enable real-time genomic contact tracing.

Quantification of uncertainty is a fundamental aspect of interpreting phylogenetic inferences^23^ and sample placements onto a reference phylogeny. Non-parametric bootstrapping^24^ has been a cornerstone of phylogenetic inference for decades, but this is impractical for the extremely large sample sizes and the limited phylogenetic information in SARS-CoV-2 genome isolates. More recently developed methods such as Ultrafast Bootstrapping^25,26^ are fast, but not applicable to the problem of placing individual samples onto a reference phylogeny. An alternative to these approaches is the approximate likelihood ratio test^27^, but its computation is prohibitively slow and interpretation challenging. Quantifying uncertainty in sample placement on reference phylogenies is therefore an important unsolved problem and particularly relevant during this pandemic.

In this work, we describe an efficient method that facilitates rapid, maximum parsimony placement of samples onto an existing phylogeny. We show that our method for placing new genome sequences onto a SARS-CoV-2 phylogeny is orders of magnitude faster than existing approaches and produces highly accurate results. Additionally, we introduce the Branch Parsimony Score (BPS) which is the minimum number of additional mutations (the parsimony score) required to accommodate a sample placement at a given branch. This offers an intuitive means of quantifying uncertainty in sample placements for SARS-CoV-2 phylogenies. Our placement approach and related data visualization tools are available from the UCSC SARS-CoV-2 Genome Browser^28^ and will empower genomic contact tracing applications worldwide.

## Results and Discussion

### Existing Placement Methods are Inadequate for the SARS-CoV-2 Global Phylogeny

Genomic contact tracing during this global pandemic necessitates algorithms that efficiently place samples onto the vast global tree. With this requirement in mind, we evaluated the performance of several existing approaches^17–20^ and compared their runtime and memory usage by adding just one additional sequence to a SARS-CoV-2 global phylogeny containing 38,342 leaves, our “reference phylogeny”, which comes from the 11/7/2020 release of^22^. We found that the time required to place a single sample is unacceptably large. For example, EPA-NG^18^ takes approximately 28 CPU minutes to place one sample and requires 791 GB of memory (Table 1).

**Table 1:** Average and range of time required to place one sample and peak memory usage across 20 replicate runs of each placement algorithm. A typical use case for placing SARS-CoV-2 samples onto the global phylogeny will often require placing 10-100 sequences. We do not evaluate that here because we found that several other algorithms could not be run on larger sample sets due to exceptionally high memory usage and runtimes. Note that while the remaining tools use a multiple sequence alignment (MSA) as input, UShER accepts a VCF for new samples, which can be generated very quickly (compared to adding sequences to an existing MSA) using pairwise alignments (in *e.g*., minimap2^29^) and whose overhead we ignore. We also note that TreeBeST is not developed explicitly for this purpose, but we include it here because it has tree placement capabilities.

| Program | Average time to place one sample | Time range over 20 replicates | Average peak memory used (GB) | Memory range (GB) over 20 replicates |
| --- | --- | --- | --- | --- |
| IQ-TREE 2.1.0 | 46m 31s | 29m 56s - 68m 52s | 12.85 | 12.82 - 12.89 |
| EPA-NG | 27m 38s | 25m 19s - 31m 13s | 791.82 | 781.80 - 800.85 |
| PAGAN2 | 120m 32s | 102m 5s - 156m 15s | 470.74 | 468.10 - 473.84 |
| TreeBeST | 48+ hours | N/A | N/A | N/A |
| **UShER (with pre-processed mutation-annotated tree)** | **0.5s** | **0.40s - 0.65s** | **0.28** | **0.17 - 0.32** |
| **UShER (without pre-processed mutation-annotated tree)** | **1m 43s** | **1m 40s - 1m 46s** | **1.02** | **0.99 - 1.04** |

To address the challenge of real-time sample placement, we developed a new tool called <u>U</u>Jtrafast <u>S</u>ample placement on <u>E</u>xisting t<u>R</u>ees (UShER). UShER can place a SARS-CoV-2 sample onto our reference phylogeny in just 0.5 seconds – several orders of magnitude improvement over the next fastest tool. A part of the increased efficiency of UShER stems from its heavily optimized encoding of mutations compared to a multiple sequence alignment (MSA) and from its pre-computed data object storing the inferred histories of mutation events on the tree before placing samples during each execution (details below).

### UShER Uses Efficient Tree-Based Data Objects

Existing approaches to sample placement use a Multiple Sequence Alignment (MSA) of genomes that requires storing a whole-genome sequence for each sample (see Figure 1, Methods). UShER’s primary data structure is substantially more efficient. It starts with a list of variants with respect to a reference sequence for each sample and represents genotype data based on the inferred phylogeny of the viral population itself. UShER uses the Fitch-Sankoff algorithm to infer the placement of mutations on a given tree and on the variant list^30,31^. Besides the phylogeny itself, UShER records only the nodes for which mutations are inferred to have occurred on the branches leading to them in a representation that we call mutation-annotated tree (Figure 1). This representation is particularly favorable for the SARS-CoV-2 phylogeny in which the mutations are relatively rare and often shared across several samples. This approach has parallels to efficient tree-based representations used recently in population genetics^32,33^. For our SARS-CoV-2 reference phylogeny, UShER’s mutation-annotated tree uses only 3.4MB of memory (that fits easily in a last-level cache^34^) to encode virtually the same information as the full MSA which requires 1.14GB (>300x improvement).

**Figure 1:**
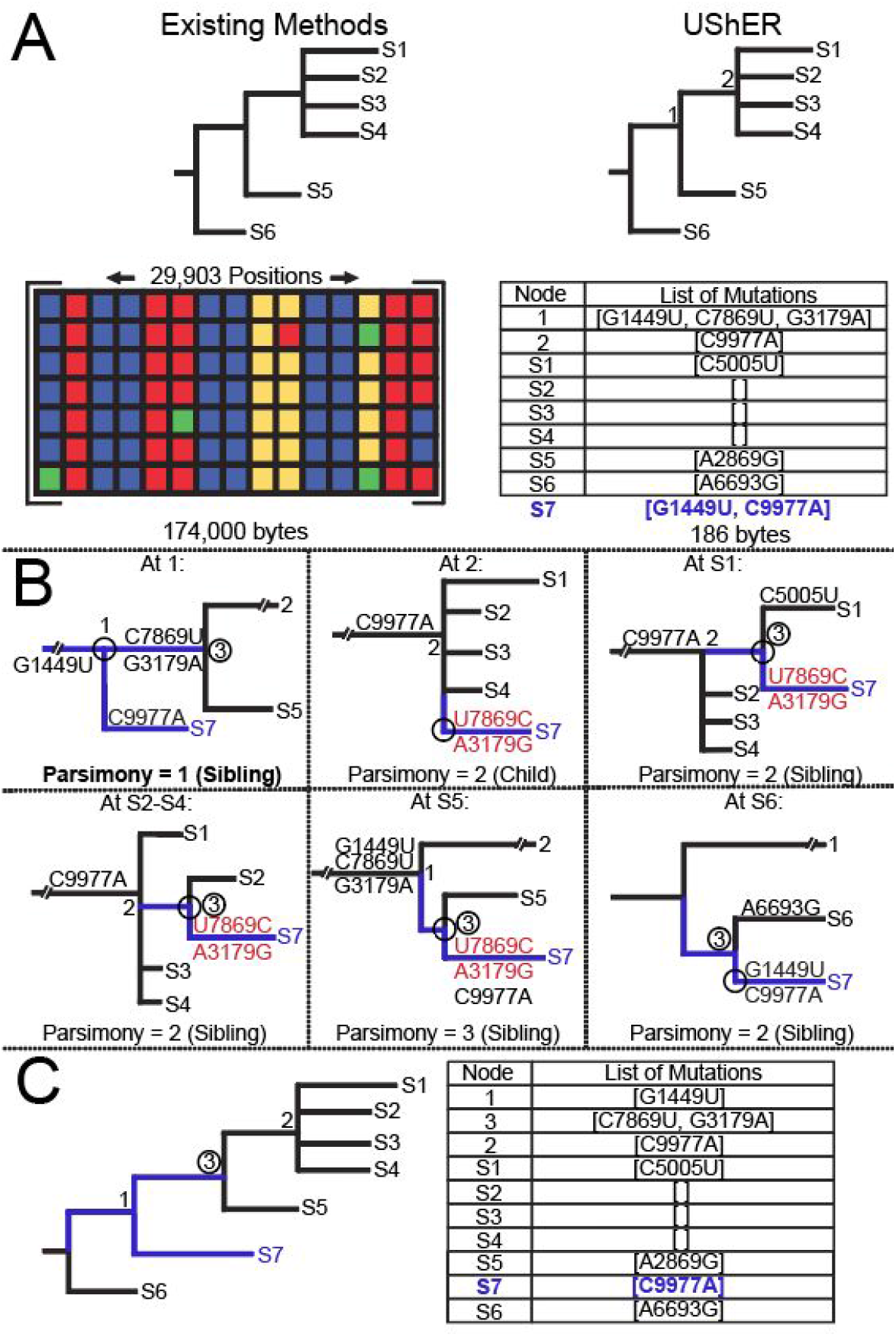
Overview of UShER’s placement algorithm and data object. (A) Prior methods rely on a full multiple sequence alignment (MSA) to inform phylogenetic structure (left), while UShER uses a mutation-annotated tree (right). (B) UShER evaluations of the parsimony score for placing the sample S7 (blue) at each possible position (see Methods) of our example phylogeny (shown in panel A). The branches that need to be modified or added to the phylogeny to accommodate S7 are shown in blue; back mutations (if present) are colored in red and new nodes are circled. Placing S7 at node 1 is optimal by parsimony. Note that UShER ultimately discards placement configurations in which branches leading to internal nodes have no assigned mutations, since they are equivalent to placement at parent (e.g. at S1, S2-S4). (C) The final tree with S7 added, in which an additional internal node 3 is added to support S7 (left), and the mutation annotations for the final tree with S7 colored in blue (right). Note that the memory efficiency of the mutation-annotated tree can vary depending on the dataset.

UShER can generate a mutation-annotated tree for our reference tree with 38,342 leaves and 15,129 variant positions in just 2:24 minutes:seconds using four threads (Table S1). This data structure is then stored as a pre-processed protocol buffer (https://developers.google.com/protocol-buffers), which is a customizable binary file format that can be rapidly loaded (~150 milliseconds) during sample placement and data visualization (Figure 1) and obviates the need to recompute the assignments for each execution.

### Sample Placement using Mutation-Annotated Tree

UShER uses this mutation-annotated tree to rapidly place newly acquired samples onto the tree of all known SARS-CoV-2 variation. More specifically, UShER uses a maximum-parsimony approach where it searches the entire reference tree (see Figure 1, Methods) for a placement that requires the fewest additional mutations to accommodate the added sample (*i.e*., the maximum-parsimony placement of a sample). UShER breaks ties based on the number of descendant leaves at the placement nodes when multiple placements are parsimony-optimal (Methods). When a pre-processed mutation-annotated tree is already available, this procedure takes approximately 0.5 seconds to place a single sample onto the SARS-CoV-2 reference tree (Table 1) and is even more efficient when placing larger sets of samples since the time to load the mutation-annotated tree gets amortized. For example, it only takes ~18 seconds to place 1000 samples onto our reference tree using 16 threads (Table S1). This means that our implementation is fast enough to facilitate real-time placement of SARS-CoV-2 sequences and sufficiently memory efficient (Table 1, Table S1–S2) that everything we present could be run on a basic laptop, which should facilitate widespread adoption of this approach.

### UShER Accurately Places Simulated SARS-CoV-2 Samples

To evaluate the accuracy of UShER’s maximum parsimony-based placement algorithm when the viral evolutionary history is known, we generated a SARS-CoV-2 simulated dataset using a fixed tree that we supplied (see Methods). UShER places samples with the correct sister node in 97.2% of cases. For samples with just one parsimony-optimal placement, UShER achieves 98.5% accurate local placements. When incorrect, UShER’s placements tend to be quite close to the correct node on the SARS-CoV-2 global phylogeny – separated by just 1.1 edges from the correct position on the tree, on average (Figure 2, Methods). We therefore conclude that UShER is capable of accurately placing new samples onto a fixed SARS-CoV-2 global reference phylogeny in practice and could indeed facilitate the ongoing genomic contact tracing efforts. Although UShER works well for SARS-CoV-2, it will not be as accurate for phylogenetic analyses in which maximum parsimony algorithms are known to perform poorly (*e.g*., cases of long branch attraction^35^).

**Figure 2:**
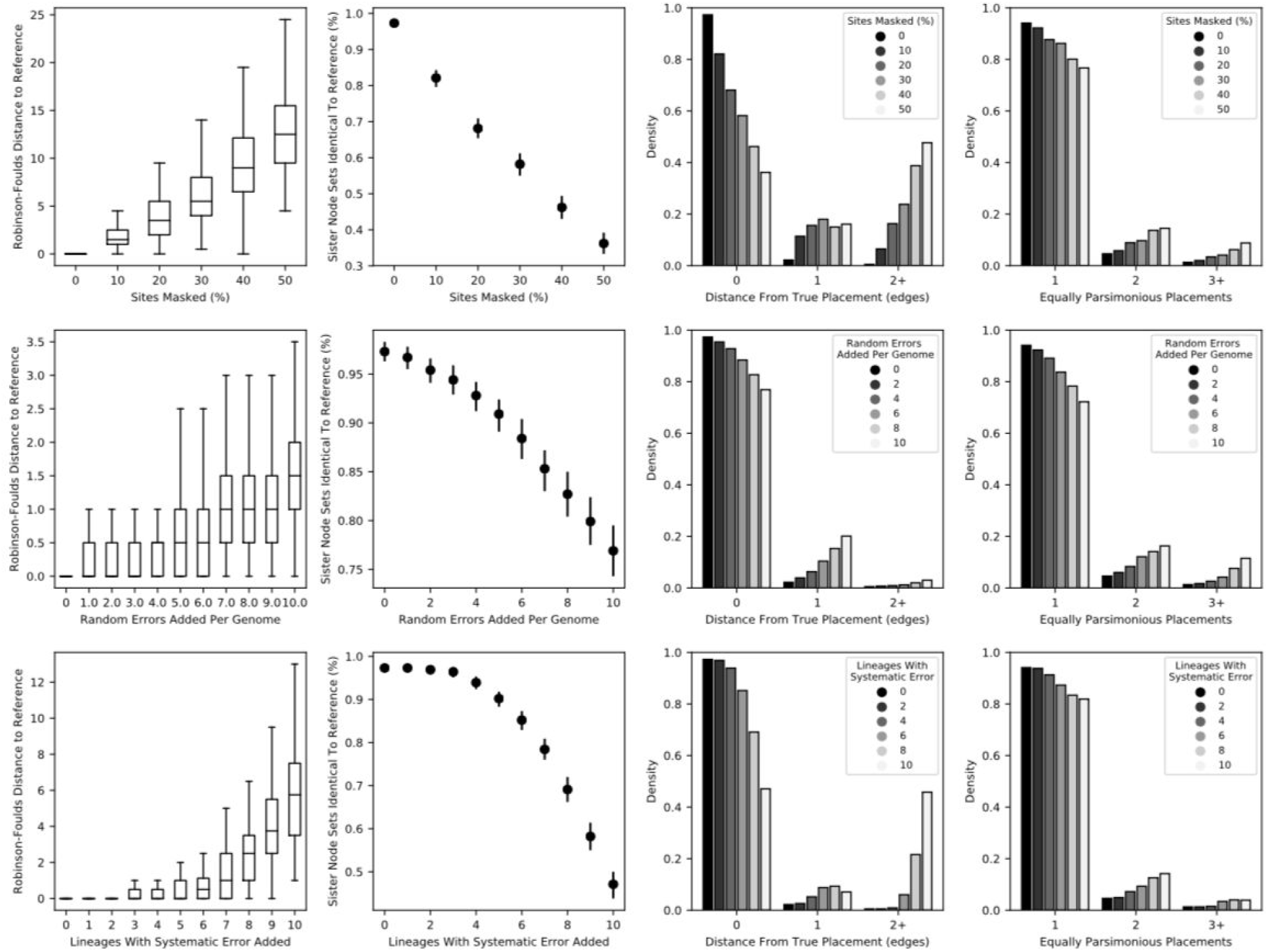
The maximum-parsimony algorithm used by UShER is robust to moderate rates of missing data and simulated errors in SARS-CoV-2 genomes. Top: We independently masked sites at 10, 20, 30, 40, and 50 percent of sites for each of 10 simulated genomes to be added to the phylogeny, and computed the Robinson Foulds distance^39^, the average number of lineages added that had identical sister node sets to those in the simulated reference tree, the distance from true placement for each lineage added (see Methods), and the number of equally parsimonious sites per placement for each lineage added. Middle: We introduced random nucleotide substitutions to the genomes of the 10 lineages added to the tree by UShER at a rate of 1, 2,… 10 sites per genome, drawn independently, and computed the same measures of coherence to the reference tree, with error bars representing 95% confidence intervals. Bottom: We introduced one systematic error to 1, 2,… 10 of the genomes added to the tree by UShER and computed the same metrics as above.

### Missing Data Has A Large Effect on Sample Placement of SARS-CoV-2 Genomes

Given the low mutation rate and therefore low phylogenetic signal in SARS-CoV-2 viral genomes, missing data has a large impact on phylogenetic placement, as expected (Figure 2). When we randomly masked between 0 and 50% of positions in samples to be placed by UShER, all measures of placement accuracy were negatively impacted. With 50% of all sites masked, we find that only 41.9% of samples are assigned identical sister nodes as their true placement on the reference tree. However, the mean distance between UShER and correct placements on the tree remained relatively small – just 1.61 edges– and 81.0% of sister nodes contain leaves that overlap those of the correct sister node (see Methods). This suggests that missing data affects the precision of UShER’s placements (and is expected to affect other placement algorithms similarly), but that the placement is sufficiently accurate with low rates of missing data that it can still be valuable for epidemiological studies and for genomic contact tracing.

### UShER is Robust to Low Rates of Sequencing Error in SARS-CoV-2 Genomes

Two types of errors in SARS-CoV-2 consensus sequences also affect the accuracy of sample placements. First, stochastic errors are likely present in many available SARS-CoV-2 sequences^36^. When we simulated independent errors, we found the effects on UShER’s accuracy are modest (Figure 2). With 10 errors on average, the placement is approximately 20% less likely to select the correct sister node, and other distance metrics are similarly impacted (Figure 2). Our results indicate that especially low quality samples should be rigorously identified and excluded from analyses using UShER. Additionally, poor quality samples can be easily flagged because they will tend to appear as unrealistically long terminal branches in UShER’s placements. UShER reports all newly added samples with a parsimony score greater than 3 along with a list of parsimony-increasing sites.

Second, systematic error, where the same apparent variant is introduced into many sequences, are present in some SARS-CoV-2 sequences and have the potential to affect phylogenetic inference because they appear as inherited mutations^37,38^. Whereas UShER appears to be robust to a single systematic error present in fewer than five samples (Figure 2), a single systematic error present in all 10 samples had a similar overall effect on placement accuracy as 50% missing data in error-free sequences. Consistent with our previous work^37,38^, addition of two perfectly correlated systematic errors can drastically affect UShER performance (Figure S1). Systematic errors should be rigorously identified and removed before sample placements are performed. We refer readers to methods that we developed previously to detect and eliminate such errors^37,38^ and the UShER package includes a tool to remove known problematic positions when preparing input data.

We emphasize that sequencing errors are likely to pose similar challenges for other placement tools and our analysis is meant to serve as a guideline to the user rather than highlight the limitations of UShER.

### Quantifying Uncertainty in Sample Placement

Quantifying uncertainty in phylogenetic placement is critical for accurately interpreting SARS-CoV-2 phylogenies where true phylogenetic signal is limited and sometimes even contradictory^36,37^. We developed functionality within UShER to report the number of equally parsimonious placements by default. Additionally, UShER can output the minimum number of additional mutations required to accommodate a single sample placed on each branch of the reference tree, a measure which we call the Branch Parsimony Score (BPS). We limit this function to single sample placements because it would be challenging to quantify and to represent the uncertainty imposed by the sequential incorporation of additional samples. As would be expected given the typically unambiguous sample placements for high quality sequences on the global phylogeny, BPS typically increases rapidly with increasing distance along the tree (*e.g*. Figure 3).

**Figure 3.**
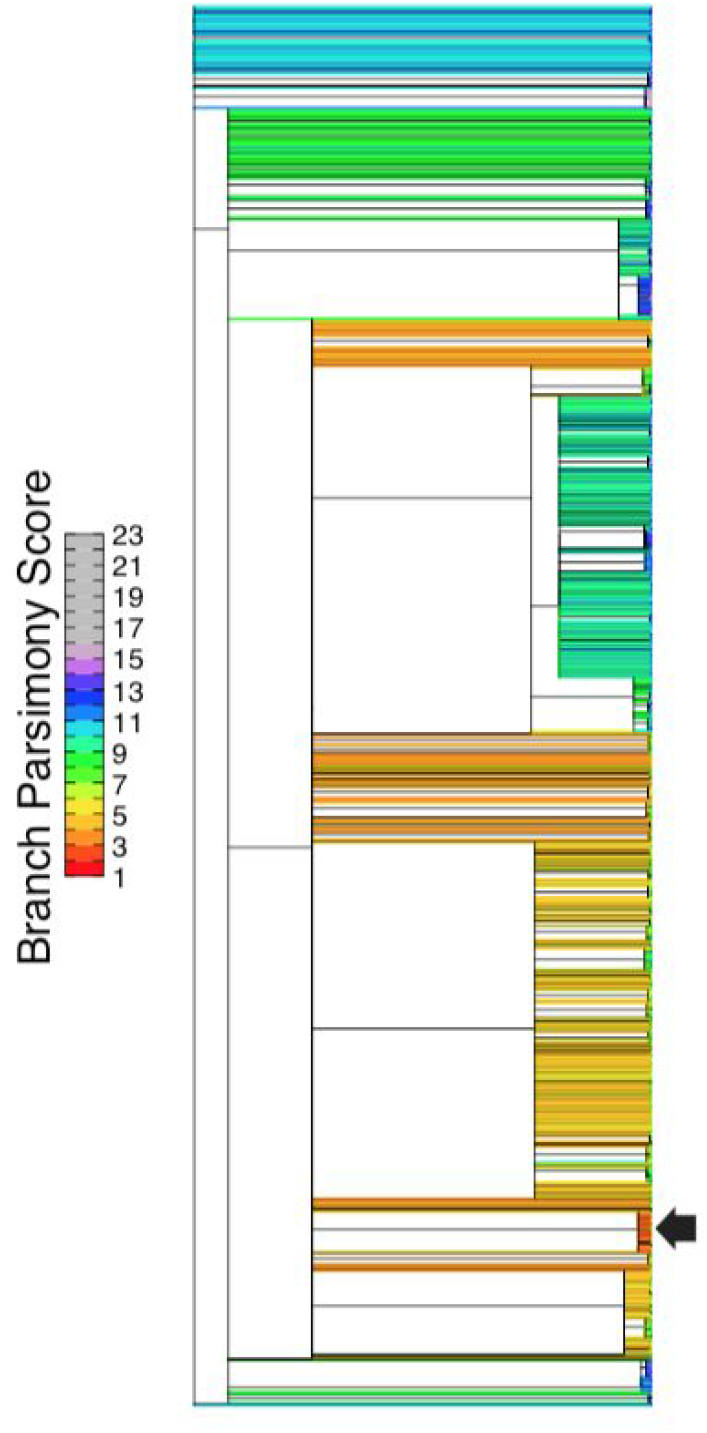
The branch parsimony score (BPS) statistic for a single sample across the global SARS-CoV-2 phylogeny. The correct sample placement which corresponds to the maximally parsimonious placement is shown with an arrow and each branch is colored by the BPS for that sample on that branch.

### UShER is Consistent with Standard Phylogenetics Methods Using Real SARS-CoV-2 Data

To evaluate the performance of our approach under realistic conditions with genuine SARS-CoV-2 data, we used UShER to place real samples onto a global reference phylogeny. Because the phylogeny was necessarily inferred from real data (see Methods), this approach measures the consistency of placement between more typical tree-building approaches and UShER placement algorithm rather than placement accuracy *per se*. To evaluate the consistency, we randomly pruned and replaced 100 sets of 10 samples each using the reference tree (see Methods). We found that UShER placed each with an identical sister node as in the reference tree in 90.0% of cases (Figure 4). Additionally, the placements tend to be quite close to correct and the mean number of edges between the reference position and UShER’s placement is just 0.159 and the mean Robinson-Foulds distance for trees with 10 samples added is 1.27 (Figure 4A). When we mimicked a plausible use case by removing larger sets of related sequences, we found that UShER is also able to accurately reconstruct larger subtrees for the added samples (Figure 4D-G). Collectively, our metrics are not far from those we obtained when analyzing the simulated datasets, and indicate that missing data, errors and other features of real sequences occasionally impact UShER’s placements.

**Figure 4.**
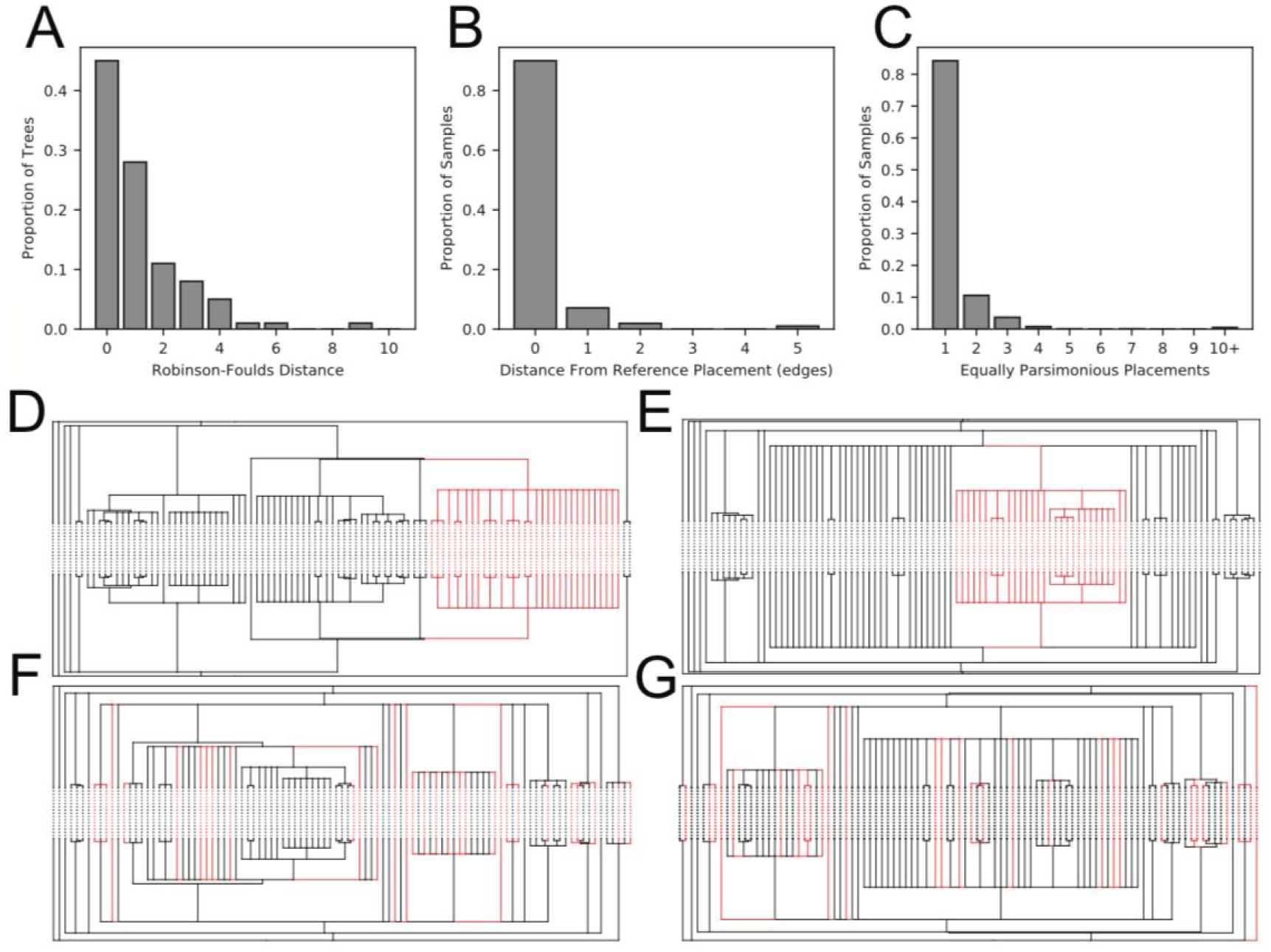
UShER is accurate using real data. The Robinson-Foulds distance between 100 reference and UShER-generated trees produced by removing and re-adding 10 samples in each (A), the distance from reference placement for each of 1,000 placed samples (B), and the number of equally parsimonious placements for each of the 1,000 placed samples (C). Comparisons of subsets of the global phylogeny released on 11/7 with reconstruction of this phylogeny using UShER (D-G). In each case, we pruned lineages colored in red from the phylogeny and added them back using UShER. UShER accurately places subtrees lineages collected in the Western United States in March/April (D) and in Europe in March (E), as well as more distantly related lineages whose times and places of collection differed more widely (F, G).

We found that samples causing inconsistent placements between the reference tree and UShER were mostly challenging cases. In particular, six of the 1000 sequences that we attempted to place using UShER have large numbers of equally parsimonious placements (5-65) and were placed inconsistently relative to the reference tree. Each of these consensus sequences has a large number of ambiguous nucleotide positions (8-15) that overlap many phylogenetically informative sites in the reference tree. This may suggest a mixture of two genetically divergent samples–either a true mixed infection or laboratory induced. Regardless of the source, we believe future versions of the reference tree should rigorously filter sequences containing ambiguities at phylogenetically informative positions.

Additionally increasing genetic distance and sequencing errors are expected to affect placement accuracy. We found that samples are more likely to be placed inconsistently when the parsimony score is higher (P = 2.98E-5, Wilcoxon Rank Sum Test). There is also a strong correlation between incorrectly placed samples and the number of equally parsimonious placements (P < 2.2E-16). In fact, 15% of real samples have more than one equally parsimonious placement on the reference phylogeny and many distinct nodes are identical in the reference tree. However, if we restrict the analysis to samples with only a single most parsimonious placement, we find that 97% of UShER’s placements are consistent with the maximum-likelihood reference tree. We suggest that the placements of samples that are unusually genetically distant or that have many equally parsimonious placements on a reference tree should be regarded with caution. Both statistics are reported by UShER.

### UShER can Empower Real-Time Phylogenetics by Maintaining a Global Phylogeny

We propose that UShER could form the basis for “real time” phylogenetics platforms in periodically updating the reference tree itself or be used in conjunction with maximum-likelihood updates. To investigate this, we used UShER to add all of the 9437 additional sequences in the 31/7/2020 release of the global tree to our 11/7/2020 reference tree. We also extensively optimized both trees using a maximum likelihood approach in fasttree (^40^ see Methods, File S1). The Robinson-Foulds distance between all trees is similar suggesting that the UShER updated topologies are close to *de novo* phylogenies (Table S3). Additionally, the optimized version of the phylogeny produced by UShER resulted in a substantially increased likelihood over the 31/7/2020 tree inferred *de novo* with similarly extensive optimization (Table S4). We obtained the highest likelihood topology from a heavily optimized 11/7/2020 tree, sample addition with UShER, and then another round of tree optimization (Table S4). This indicates that UShER, combined with additional rounds of optimization, does not result in unrecoverable local-minima but rather may help avoid them. In combination with periodic maximum-likelihood updates to the global phylogeny, UShER can offer an appealing combination of real-time phylogenetic methods and model-based practices. This combination can be used to maintain an updated phylogeny for the SARS-CoV-2 pandemic.

### Web Interface with Rapid Subtree Access for Data Visualization And Genomic Contact Tracing

Interpretation of UShER’s placements often involves scrutinizing the relationships and genotypes among closely related samples already present in the reference tree. In addition to providing the complete phylogenetic tree with new samples added, UShER can optionally provide local subtree outputs of a specified size (number of sample leaf nodes) so that the relationship of added samples to their nearest neighbors can be visualized and examined in detail. If all added samples fit within the specified size then one subtree is created; otherwise, multiple subtrees are created as necessary to provide local subtrees for all samples.

To make genomic contact tracing using UShER widely available, we developed a web interface integrated with the SARS-CoV-2 UCSC Genome Browser (https://genome.ucsc.edu/cgi-bin/hgPhyloPlace). Users may upload new sample sequences in a FASTA file, or alternatively, new sample variants relative to the reference sequence (NC_045512.2 / MN908947.3 / Wuhan/Hu-1) in a VCF file. The web server runs UShER on the new sequences and presents a summary of the sample placements to the user, with a link to download the phylogenetic tree including the newly placed sample(s). The user can click a button to view custom tracks in the Genome Browser that show subtree(s) with a configurable number of (default 50) leaves including the new sample(s) and related sequences from the initial phylogenetic tree (Figure 5). The web server uses UShER’s mutation-annotated tree to provide almost instant visualization. Additionally, to facilitate tree exploration and cross-referencing sequences against privately-maintained personal health information, our web platform also exports a JSON formatted subtree file that can be viewed behind a firewall using auspice (Figure S1, https://auspice.us, ^41^).

**Figure 5.**
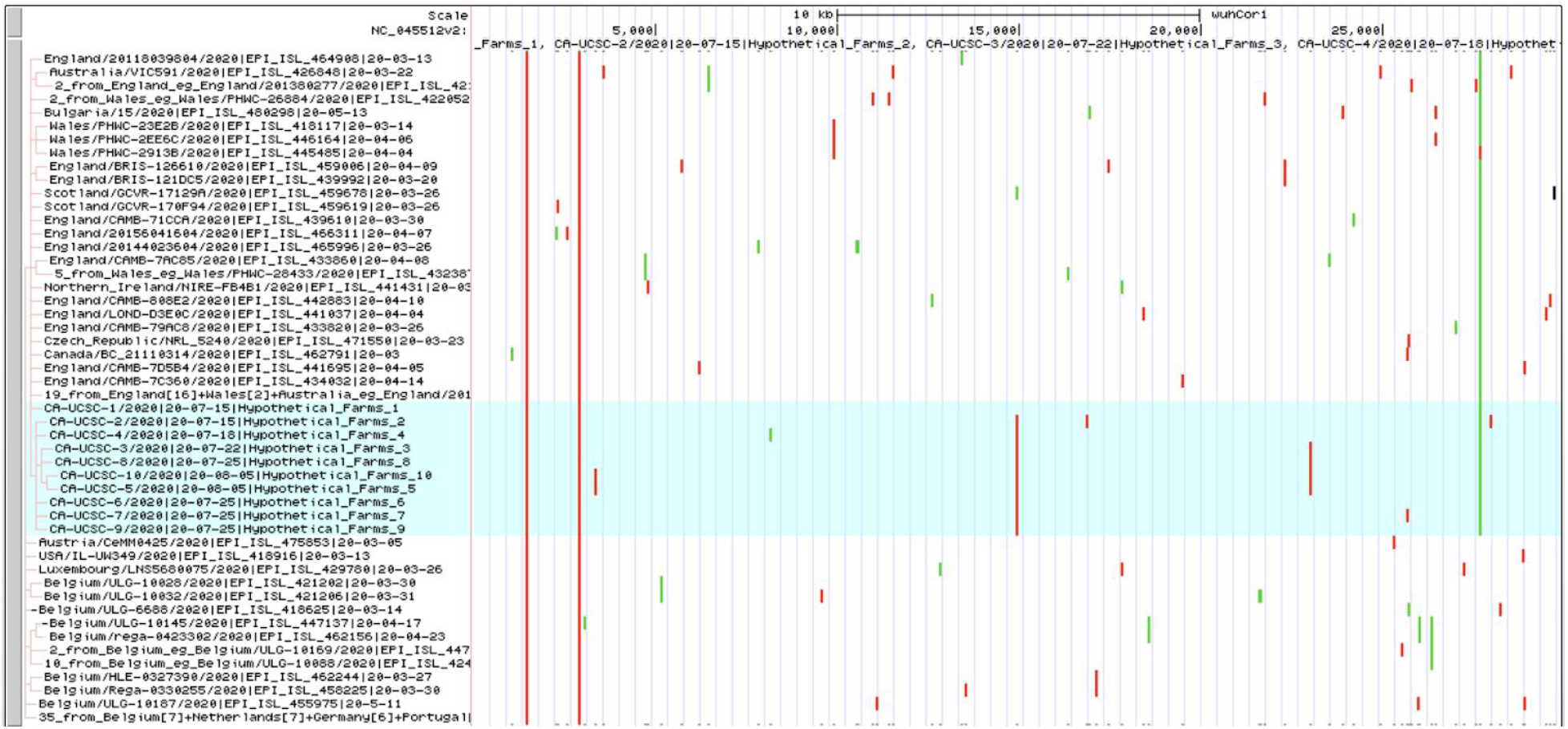
UCSC Genome Browser display of subtree where hypothetical example sequences have been placed by UShER. Newly added samples are highlighted in blue and the tree displaying their relationships and placement on the global tree is shown to the left. Interactive view: https://genome.ucsc.edu/s/AngieHinrichs/UShER_example

Widespread usage of genomic contact tracing has the potential to enable public health practitioners to link apparently independent incidences of infection even across disparate sequencing centers, as well as to disprove false links inferred from circumstantial evidence. This provides important and actionable information for suppressing transmission and for refining public health practices. The user-uploaded viral genome sequences are discarded shortly after use and not shared or stored on the UCSC Genome Browser servers, unless the user saves the subtree custom tracks in a Genome Browser Session. But we stress that for global contact tracing to be maximally effective, most users must upload their viral genome sequences to public sequence repositories so that their data can also be incorporated into the reference tree. We therefore echo the call from the INSDC (https://ncbiinsights.ncbi.nlm.nih.gov/2020/08/17/insdc-covid-data-sharing/) for all SARS-CoV-2 sequencing datasets to be made publicly available as soon as is practical.

## Conclusion

The SARS-CoV-2 pandemic has been accompanied by unprecedented levels of pathogen genomic sequencing which has truly empowered near real-time monitoring of viral transmission and evolution. This seemingly endless flood of genome sequence data has also pushed phylogenetic analysis frameworks over the edge of their capabilities, requiring new approaches to rapidly incorporate and contextualize newly sequenced viral genomes. UShER is an extremely efficient software package inspired by the ongoing evolution of the virus itself, that provides a method to immediately incorporate viral genome isolates into a global phylogenetic tree. Compared to its closest counterpart, UShER is over 3,000x faster and orders of magnitude more memory efficient. It is currently the only tool with actual real-time capabilities. UShER is also available to the worldwide research community through a user-friendly web interface in the SARS-CoV-2 UCSC Genome Browser. Though several challenges still remain, UShER can significantly decrease the turnaround time from sample to analysis and empower real-time genomic contact tracing efforts during the SARS-CoV-2 pandemic and beyond.

## Methods

### Implementation and Optimization of Algorithms in UShER

Given existing samples, whose genotypes and phylogenetic tree is known, and the genotypes of new samples, UShER aims to incorporate new samples into the phylogenetic tree while preserving the topology of existing samples and maximizing parsimony. UShER’s algorithm consists of two phases: (i) the pre-processing phase and (ii) the placement phase.

In the pre-processing phase, UShER accepts the phylogenetic tree of existing samples in a Newick format and their genotypes, specified as a set of single-nucleotide variants with respect to a reference sequence (UShER currently ignores indels), in a VCF format. For each site in the VCF, UShER uses Fitch-Sankoff algorithm^30,31^ to find the most parsimonious nucleotide assignment for every node of the tree. When a sample contains ambiguous genotypes, multiple nucleotides may be most parsimonious at a node. To resolve these, UShER assigns it any one of the most parsimonious nucleotides with preference, when possible, given to the reference base. UShER also allows the VCF to specify ambiguous bases in samples using IUPAC format (https://www.bioinformatics.org/sms/iupac.html), which are also resolved to a unique base using the above strategy. When a branch leading to a node is found to carry a mutation, i.e. the base assigned to the node differs from its parent, the mutation (e.g. G3179A at node 1, Figure 1A) gets added to a list of mutations corresponding to the branches leading to that node. Finally, UShER uses protocol buffers (https://developers.google.com/protocol-buffers) to store in a file, the Newick string corresponding to the input tree and a list of lists of node mutations (which we also refer to as mutation-annotated tree object), as shown in Figure 1. The outer list is ordered according to the depth-first traversal of nodes. UShER also parallelizes the independent Fitch-Sankoff computations for multiple VCF sites efficiently using multiple threads (Table S1–S2).

In the placement phase, UShER loads the pre-processed mutation-annotated tree and the genotypes of new samples in a VCF format and sequentially adds the new samples to the tree. For each new sample, UShER computes the additional parsimony score required for placing it at every node in the current tree while considering the full path of mutations on the branches from the root of the tree to that node (Figure 1B). For internal nodes, the parsimonious placement can be as a sibling to the node (*e.g*. node 1, Figure 1B), when there are mutations on the branch leading to that node that are not shared by the new sample, or as a child to that node (*e.g*. node 2, Figure 1B), when all mutations on the branch leading to that node are shared by the new sample. For leaf nodes, only the sibling placement is considered (*e.g*. S1-S6 in Figure 1B) to ensure that samples are always maintained as leaves of the tree. Next, UShER places the new sample at the node that results in the smallest additional parsimony score. When multiple node placements are equally parsimonious, UShER picks the node with a greater number of descendant leaves for placement. All else being equal, the sample is most likely to come from the node with the largest number of descendants. If the choice is between a parent and its child node, the parent node would always be selected by this rule because the set of leaves descendant from the parent node necessarily contains the full set descendant from the child node. However, a more accurate placement can be obtained by evaluating the number of leaves uniquely attributable to the child versus parent node. Therefore, in these cases, UShER picks the parent node if the number of descendant leaves of the parent that are not shared with the child node exceed the number of descendant leaves of the child. While placing a new sample, UShER parallelizes the parsimony score computation over different nodes of the tree using multiple threads.

At the end of the placement phase, UShER allows the user to create another protocol-buffer file containing the mutation-annotated tree object for the newly generated tree including added samples (Figure 1C). This allows another round of placements to be carried out over and above the newly added samples. While UShER’s sequential placement of new samples helps it achieve high speed, the placements could potentially be worse than a collective strategy; however, in practice, we have found UShER’s accuracy over iterated placements to be reasonably high.

UShER implements a few additional optimizations to speed up the placement phase. For example, the parsimony score for a node requires computing the symmetric set difference between the set of new sample variants and the set of mutations on the branches from the root of the tree to that node. UShER maintains mutations sorted by positions to speed up this computation. UShER also maintains the minimum parsimony score of previously traversed nodes in a shared variable and terminates the computation of the set difference in a new node as soon as the parsimony score corresponding to it exceeds the value of this shared variable. Finally, UShER also allows an option during the pre-processing phase to collapse nodes (i.e. delete the node after moving its child nodes to its parent node), branches leading to which, are not inferred to contain a mutation through the Fitch-Sankoff algorithm as well as to condense nodes in a polytomy that contain identical sequences into a single representative node, both of which help in significantly reducing the search space for the placement phase.

### Simulating Realistic SARS-CoV-2 Genome Evolution

To evaluate the accuracy of our placement algorithm on a known phylogeny, we produced a set of simulated samples for which we knew the correct placement on the tree. To do this, we first needed to obtain a mutation rate matrix. Each position of the reference genome (NC_045512.2 / MN908947.3 / Wuhan/Hu-1, see https://www.ncbi.nlm.nih.gov/nuccore/MN908947) was classified as coding or non-coding. Start and Stop codons were not considered and were simulated as constant. First and last 100bp of the genome and sites marked as problematic in https://github.com/W-L/ProblematicSites_SARS-CoV2/blob/master/problematic_sites_sarsCov2.vcf were also not considered here (for estimating substitution rates), but they were simulated like the other “normal” sites. We counted “opportunities” of mutations based on the reference genome: for example a non-coding C allele in the reference genome represents 3 opportunities for non-coding mutations (C->A, C->G and C->U). For coding sites, we split synonymous and non-synonymous mutation opportunities into two separate counts. Then, we counted the number of observed mutation events of each type using the inferred mutational history of the viral population based on the phylogeny and global alignment from 7/11 and using our software that we have described previously (available from https://github.com/yatisht/strain_phylogenetics,^37^). As above, we masked the ends of the genome, previously identified problematic sites, and start/stop codons.

To further avoid potential biases resulting from sequencing errors, RNA degradation, contamination, or other possible sources of error, we used a threshold for minor allele counts for variants/mutations to be included in our analysis. All the results presented here assume that only variants/mutations with at least a minor allele count of 7 can be used reliably. Results vary a bit by varying this threshold, in particular, as one would expect, the nonsynonymous/synonymous mutation ratio decreases with increasing this threshold, however the effect is overall limited. Also, we use two different ways to count mutation events. In the first approach, we use the reconstructed mutation histories from the strain_phylogenetics package and only count substitution events with at least 7 descendant lineages. The other approach only uses the alignment and ignores the reconstructed mutation history: we count variant alleles in the alignment that have a minor allele count of at least 7. These two approaches give comparable results, and here we use results from the first approach.

For each class of mutations (synonymous or nonsynonymous, and from any nucleotide to any other nucleotide) we calculate a rate by dividing its total opportunity count by its mutation count. Given the small number of noncoding sites, we merge the noncoding and synonymous counts. Our final rate matrix is available from Table S5.

### Simulating sequence evolution along the tree

We assume a root genome equal to the reference genome and we simulate its evolution along a phylogeny with 38,342 leaves using pyvolve^42^. We simulate evolution of each site with a nucleotide substitution process, but still take into account the genetic code and a non-neutral nonsynonymous/synonymous rate ratio omega in doing so. We achieve this by assuming that the codon context of each position is constant - this is sufficiently realistic because it is very rare that two positions in the same codon are both mutated (with respect to the reference) in the same SARS-CoV-2 sequence. So, considering each genome position in turn, we evolve noncoding sites under the synonymous matrix above, while for coding sites, we use the same rates for synonymous mutations, while we multiply the rate of nonsynonymous mutations by omega. The reason for using this approach instead of simulating unde a codon model is that a codon model would have been prohibitively computationally intensive.

We assume that omega is variable across sites, and that there is a probability of 20% that omega=0.01, of 30% that omega=0.1, of 30% that omega=0.3, and of 20% that omega=1. Pyvolve automatically rescales input rate matrices, so we rescale the tree at each site so that the synonymous rates are the same across sites. In addition, we rescaled the overall tree so that the observed numbers of mutation events in the simulated alignment is similar to the number in the real alignment.

### Accuracy Evaluation

We measured UShER’s accuracy in placing samples onto a reference phylogeny using simulated (described above) as well as real data. We did not evaluate the accuracy of other placement algorithms because their runtimes are too large for it to be practical to evaluate their accuracy. For simulated data, both reference phylogeny and sequences were simulated, while for real data, we used the global phylogeny dated 11/7/20 (https://github.com/roblanf/sarscov2phylo, File S1) as reference and its corresponding sequences were obtained from GISAID^4^ In each case, we first randomly pruned out 10 samples from the global phylogeny, which was then used as the input phylogeny while adding back the pruned samples using UShER. UShER’s accuracy in placing back the samples was computed using average values of three different statistics (described below) over 100 such replicates.

First, we used TreeCmp^43^ to compute the Robinson-Foulds distance between the reference phylogeny and the tree constructed by samples using UShER. Second, we recorded whether the sister node for each placed sample is identical to the true sister clade (i.e., the sister clade in the reference phylogeny). Finally, we computed the distance between the UShER placement and the correct placement in terms of the minimum number of edges separating them. Ordinarily, distance between two nodes in a tree can be computed using their lowest common ancestor (LCA)^44^ i.e. by taking the sum of the number of edges to each node from the LCA. To determine the distance between the node placement in two trees (reference phylogeny and the one resulting from UShER placement), we developed a utility that reports all descendant lineages from N-th generation ancestor of any given node in a tree, with N provided as input (i.e. when N=1, it reports unpruned lineages in the sister clade). For each pruned lineage, we found the descendants varying the number of generations, N1 and N2, in global and UShER phylogenies, respectively, and reported the distance between the UShER placement and the correct placement as the smallest (N1+N2-2) that resulted in the same set of descendant lineages in both phylogenies. Note that the second statistic records cases for which the sister clades in the two trees to be identical, which would always have 0-distance in our third statistic (with N1=N2=1).

We also measured UShER’s accuracy in a realistic scenario of placing closely related samples that form their own subtree. In this case, we required during pruning that the pruned samples together form a subtree (that is not a trivial polytomy) in the reference phylogeny.

### Benchmarking Placement Algorithms

We compared UShER to four other lineage placement algorithms: IQ-TREE 2, EPA-ng PAGAN2, and TreeBEsT^17–20^ We initially attempted to add 1,000 lineages to our simulated phylogeny, however except UShER, which required 18 seconds to finish using 64 threads, none of the placement programs finished within 24 hours. Due to time and memory constraints, we instead added only one lineage to the tree in 20 replicates, recording the average and range of time and peak memory usage across these 20 replicates in Table 1. A full list of commands used to run each test can be found in Supplementary Table S6. We installed and ran each program on a server with 160 processors (Intel(R) Xeon(R) CPU E7-8870 v4 @ 2.10GHz), each with 20 cpu cores.

### Tree Construction for SARS-CoV-2 Samples

Full details and reproducible code for the construction of the global tree of SARS-CoV-2 samples are available in the 31/7/20 release and at^22^. To summarise, this code creates a global phylogeny of all available samples from the GISAID data repository as follows. First, all sequences marked as ‘complete’ and ‘high coverage’ submitted up to 31/7/20 were downloaded from GISAID. Sequences with known issues from previous analyses were then removed from this database (details are in the excluded_sequences.tsv file at the above DOI). Second, a global alignment was created by aligning every sequence to the NC_045512.2 accession from NCBI, using mafft^45^, faSplit (http://hgdownload.soe.ucsc.edu/admin/exe/), faSomeRecords (https://github.com/ENCODE-DCC/kentUtils), and GNU parallel^46^. Third, sites that are likely to be dominated by sequencing error^37^ are masked from the alignment using faSplit, seqmagick (https://seqmagick.readthedocs.io/en/latest/), and GNU parallel, sequences shorter than 28KB and/or with >1000 ambiguities are removed from the alignment using esl-alimanip (hmmer.org), and subsequently sites that are >50% gaps are removed (after converting N’s to gaps) esl-alimask. Fourth, the global phylogeny was estimated using fasttree^47^ in two stages: (i) an initial analysis that produces a Neighbour Joining tree which is optimised with 5 rounds of SPR moves of length 500; and (ii) a second analysis which uses the tree from the first analysis as a starting tree with 5 rounds of SPR moves of length 200 and otherwise default fasttree settings. Finally, goalign (https://github.com/evolbioinfo/goalign) was used to create 100 bootstrap alignments followed by re-estimating all the ML trees with fasttree with the -fastest setting, using GNU parallel to manage parallelisation. FBP and TBE values were calculated with gotree (https://github.com/evolbioinfo/gotree), and SH values are calculated with fasttree. The resulting trees were rooted with our reference (NC_045512.2 / MN908947.3 / Wuhan/Hu-1) sequence using nw_reroot^48^.

From the resulting tree, we removed sequences on very long branches using TreeShrink^49^. These sequences are likely to be either of poor quality and/or poorly aligned, so rather unreliable to interpret in a phylogeny with such limited variation.

### Tree Optimization for Inferred SARS-CoV-2 Phylogenies

For the 11/7-20 and 31/7/20 reference trees, we created “optimized” versions of each using FastTree^40^, using ten iterations of the command *“fasttree -nt -nni 0 -spr 1 -sprlength 1000 -nosupport -intree <initial tree> global.fa > <new tree>“*, replacing the initial tree with the new tree from the previous iteration each time, followed by the command *“fasttree -nt -nni 0 -spr 1 -sprlength 1000 -nosupport -gamma -intree <initial tree> global.fa > <new tree>”*. Because FastTree requires binary trees, we randomly resolved all polytomies prior to optimization. We also generated two other trees using UShER, by taking the original and “optimized” 11/7 trees, pruning out all lineages in the 11/7/20 tree that are not present in the 31/7/20 tree, and using UShER to add in all lineages present in the 31/7/20 that were not present in the 11/7 tree. We then optimized these two new trees using ten iterations of FastTree, followed by another round of optimization using the *-gamma* flag as described above.

### Parsing the Mutation Annotated Tree Object for Rapid Subtree Visualization

When we were developing the web-application for UShER we discovered that parsing genotype data from a VCF file containing 40,000+ samples was the most time consuming step for displaying the resulting subtrees. We therefore developed an approach for parsing genotype data from subtrees from the mutation annotated tree object. Briefly, our approach descends from the root of the phylogeny to the focal subtree, accumulates the relevant mutations along the path, and then extracts the variation within the subtree that will be used for visualization. This heavily reduced dataset can then be visualized using the existing code-base of the UCSC Genome Browser and we output a JSON-formatted file that can be viewed using auspice (https://nextstrain.github.io/auspice/). With the current dataset sizes, this procedure takes approximately 0.03 seconds in total to extract genotype data for a subtree of 50 sequences. Software for rapid subtree VCF extraction from our mutation annotated tree object is available from https://github.com/ucscGenomeBrowser/kent/tree/master/src/hg/hgPhyloPlace/phyloPlace.c.

## Acknowledgements

We thank all of the Nextstrain team members whose platform we have built on as a part of this work. We also thank Sidney Bell and Minh Bui for providing valuable feedback on UShER features.

Until about the 24th of June, 2020, GISAID acknowledgements were provided in tabulated form on their website so could be combined into a single file. This is File S2. Subsequent to this, GISAID shifted to only allow PDF downloads of this table. It is not possible though to download the entire table at once, only small subsections of it. As a result, acknowledgements after the 24th of June are provided in .pdf files (File S3-S6). This statement applies to all formats of acknowledgement tables: We gratefully acknowledge the following Authors from the Originating laboratories responsible for obtaining the specimens, as well as the Submitting laboratories where the genome data were generated and shared via GISAID, on which this research is based.

## Supplemental Files

**File S1.** Newick-formatted trees created in this work. In order, the phylogenies in this file are (1) the reference 31/7/2020 tree, (2) the optimized 31/7/2020 tree, (3) the 11/7/2020 reference tree with missing samples added by UShER, (4) the 11/7/2020 optimized tree with samples added by UShER, (5) the 11/7/2020 reference tree with samples added by UShER and then optimized, (6) the 11/7/2020 optimized tree with samples added by UShER then optimized again, (7) the 11/7/2020 reference tree with missing samples added at random, (8) then 11/7/2020 optimized tree with missing samples added at random.

**Table S1:**
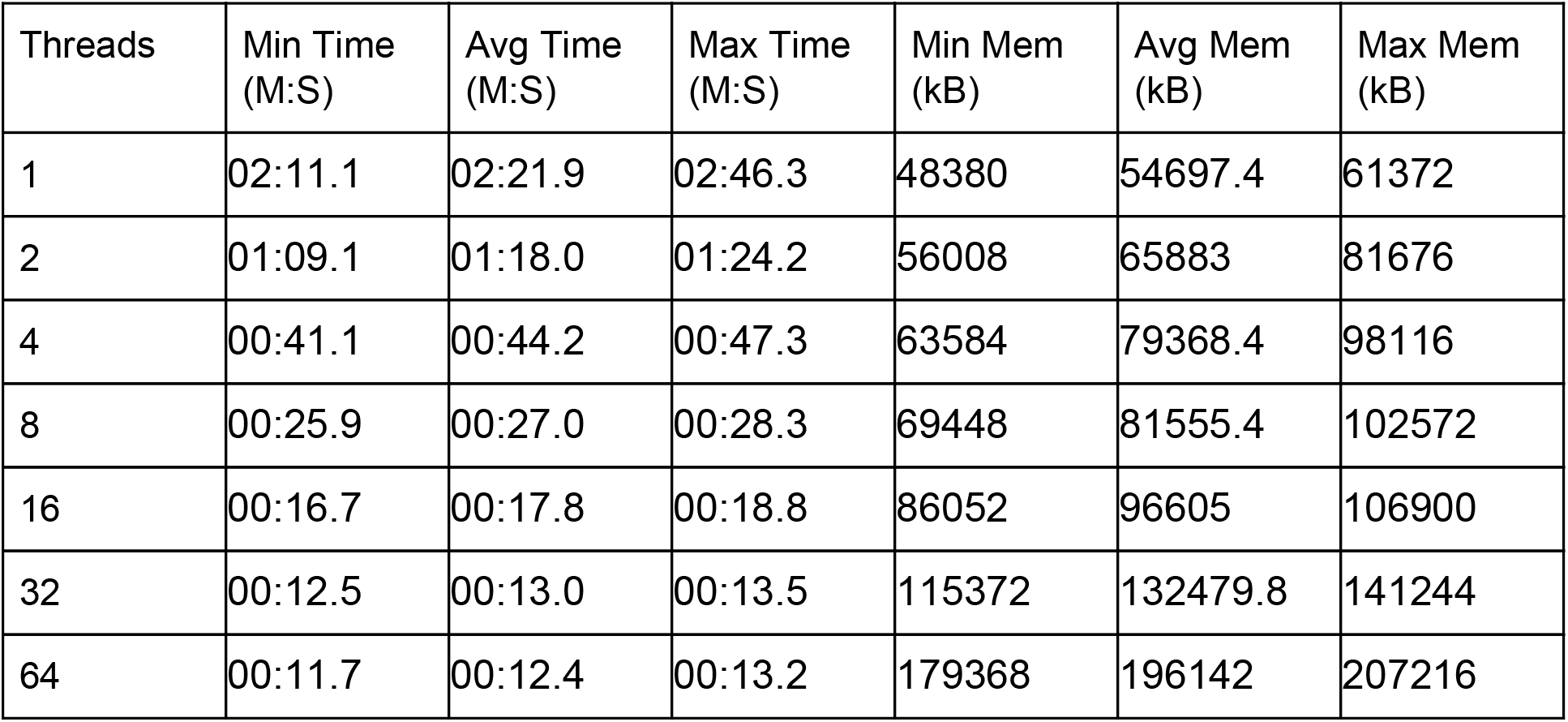
The minimum, average, and maximum total time used and peak memory usage across 20 replicates used by UShER to place 1,000 samples with varied thread arguments. This table shows data for only the lineage placement step, as protobuf files were previously generated for each replicate.

| Threads | Min Time (M:S) | Avg Time (M:S) | Max Time (M:S) | Min Mem (kB) | Avg Mem (kB) | Max Mem (kB) |
| --- | --- | --- | --- | --- | --- | --- |
| 1 | 02:11.1 | 02:21.9 | 02:46.3 | 48380 | 54697.4 | 61372 |
| 2 | 01:09.1 | 01:18.0 | 01:24.2 | 56008 | 65883 | 81676 |
| 4 | 00:41.1 | 00:44.2 | 00:47.3 | 63584 | 79368.4 | 98116 |
| 8 | 00:25.9 | 00:27.0 | 00:28.3 | 69448 | 81555.4 | 102572 |
| 16 | 00:16.7 | 00:17.8 | 00:18.8 | 86052 | 96605 | 106900 |
| 32 | 00:12.5 | 00:13.0 | 00:13.5 | 115372 | 132479.8 | 141244 |
| 64 | 00:11.7 | 00:12.4 | 00:13.2 | 179368 | 196142 | 207216 |

**Table S2:**
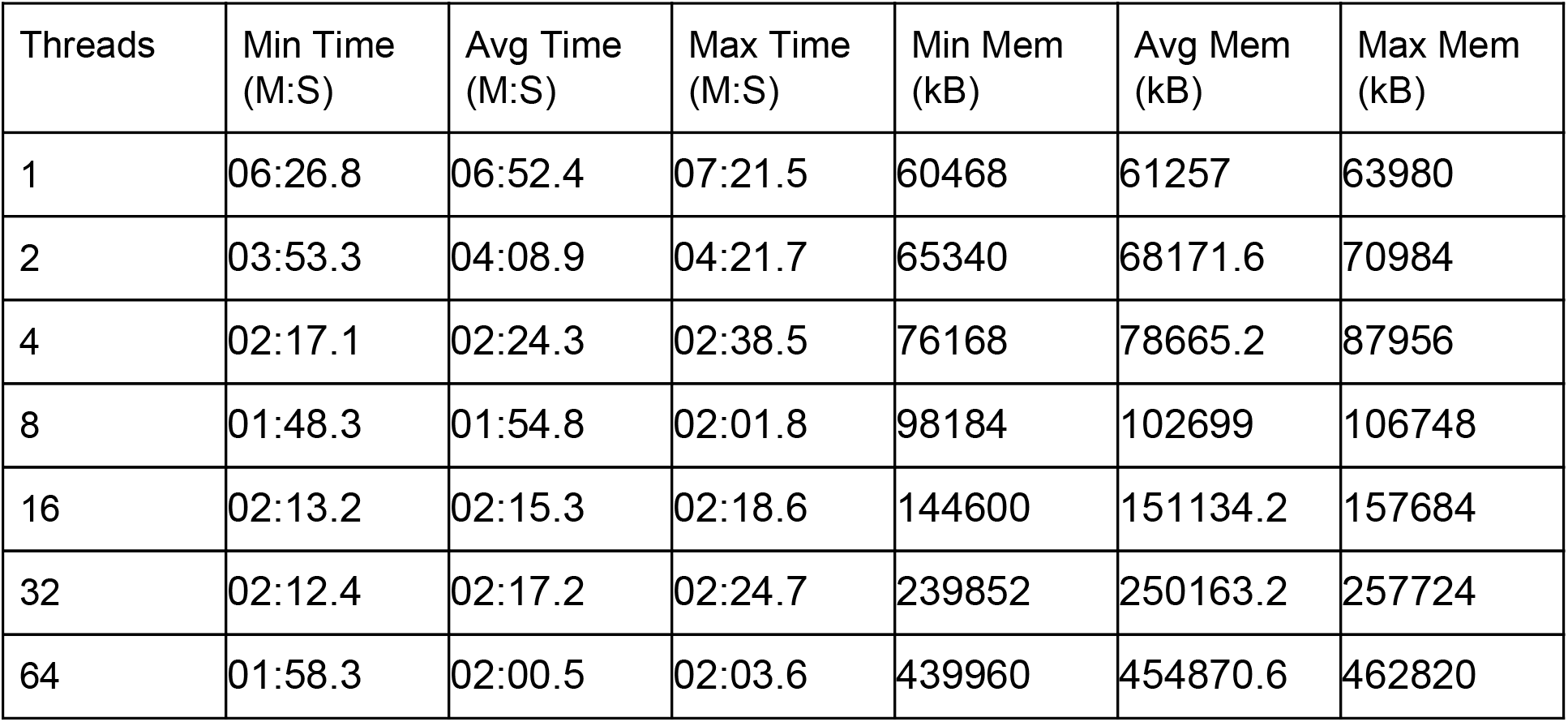
The minimum, average, and maximum total time used and peak memory usage across 20 replicates used by UShER to create a protobuf file containing the mutation-annotated tree for a 38,342 lineage SARS-CoV-2 reference tree with varied thread arguments.

| Threads | Min Time (M:S) | Avg Time (M:S) | Max Time (M:S) | Min Mem (kB) | Avg Mem (kB) | Max Mem (kB) |
| --- | --- | --- | --- | --- | --- | --- |
| 1 | 06:26.8 | 06:52.4 | 07:21.5 | 60468 | 61257 | 63980 |
| 2 | 03:53.3 | 04:08.9 | 04:21.7 | 65340 | 68171.6 | 70984 |
| 4 | 02:17.1 | 02:24.3 | 02:38.5 | 76168 | 78665.2 | 87956 |
| 8 | 01:48.3 | 01:54.8 | 02:01.8 | 98184 | 102699 | 106748 |
| 16 | 02:13.2 | 02:15.3 | 02:18.6 | 144600 | 151134.2 | 157684 |
| 32 | 02:12.4 | 02:17.2 | 02:24.7 | 239852 | 250163.2 | 257724 |
| 64 | 01:58.3 | 02:00.5 | 02:03.6 | 439960 | 454870.6 | 462820 |

**Table S3.** Robinson-Foulds distances between 11/7 and 31/7 trees and modifications by UShER. Most tree topologies, using fasttree or UShER, are similar.

|  | Reference<br>31/7 | Optimized<br>31/7 | 11/7<br>Reference<br>+USHER | 11/7<br>Optimized +<br>USHER | 11/7<br>USHER<br>then<br>Optimized | 11/7<br>Optimized+U<br>ShER<br>then<br>Optimized | 11/7+<br>Randomly<br>Placed<br>Samples | 11/7_<br>Optimized+R<br>andomly<br>Placed<br>Samples |
| --- | --- | --- | --- | --- | --- | --- | --- | --- |
| Reference 31/7 | 0 | 1522.5 | 1768.5 | 2112 | 1929.5 | 2014 | 5882.5 | 6178 |
| Optimized 31/7 |  | 0 | 2104 | 2103.5 | 1838 | 1876.5 | 6140 | 6276.5 |
| 11/7 Reference<br>+USHER |  |  | 0 | 1378.5 | 1708 | 1915.5 | 5414 | 5965 |
| 11/7<br>Optimized+USHER |  |  |  | 0 | 1812.5 | 1550 | 5868.5 | 5699 |
| 11/7 USHER then<br>Optimized |  |  |  |  | 0 | 1646.5 | 6030 | 6211.5 |
| 11/7<br>Optimized+USHER<br>then Optimized |  |  |  |  |  | 0 | 6079.5 | 6137 |
| 11/7+Randomly<br>Placed Samples |  |  |  |  |  |  | 0 | 6171.5 |
| 11/7_Optimized+Ra<br>ndomly Placed<br>Samples |  |  |  |  |  |  |  | 0 |

**Table S4.** Log likelihoods of each phylogeny after ten rounds of optimization by using FastTree2. Higher (closer to zero) log-likelihoods imply a better tree.

| **Tree** | **Log-Likelihood** |
| --- | --- |
| Optimized 31/7 Reference Tree | **-455888.368** |
| 11/7 Reference Tree + UShER then Optimized | **-455680.894** |
| Optimized 11/7 Reference Tree + UShER then Optimized | **-455241.894** |

**Table S5.**
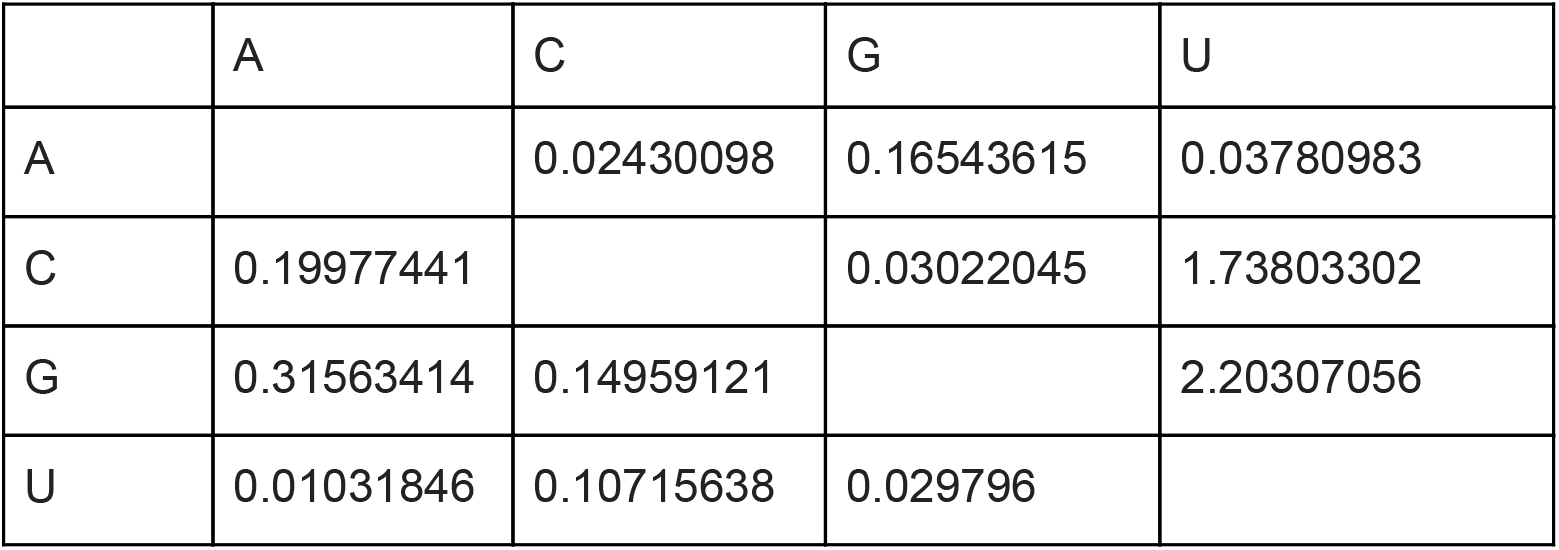
Inferred mutation rate matrix for SARS-CoV-2 evolution at synonymous and untranscribed sites.

**Table S6.**
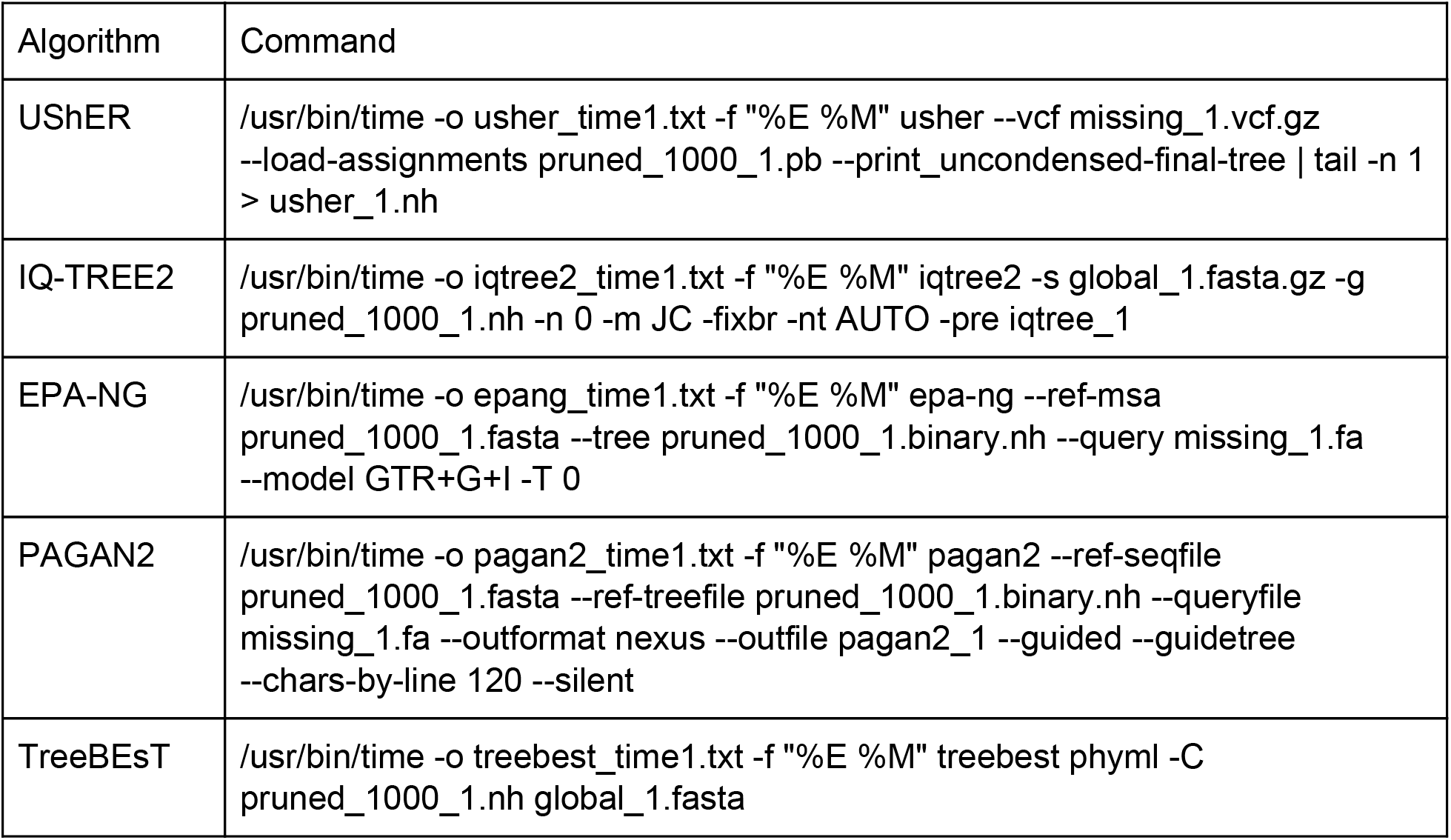
Exact commands used to profile time and memory usage of each tree placement algorithm. “Pruned” files contain data related only to lineages present in the tree prior to addition. “Missing” files contain data related only to lineages to be added to the tree. “Global” files contain data related to all lineages. For algorithms that require binary trees, we used the multi2di command in the ape package in R^50^.

**Figure S1.**
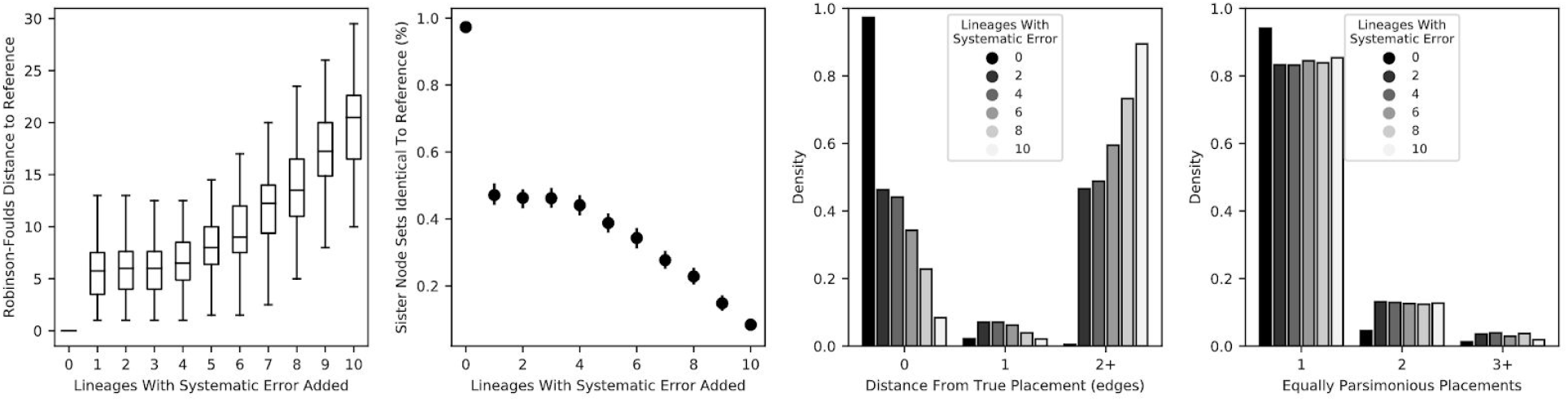
Addition of two perfectly correlated errors significantly reduces UShER accuracy. As in Figure 2, the Robinson-Foulds distances, proportion of sister nodes identical to the reference tree, distance from true placement and equally parsimonious placements, respecitvely, are shown for UShER experiments in placing 10 lineages, with two perfectly correlated errors added to 1, 2… 10 of the lineages to be placed.

**Figure S2.**
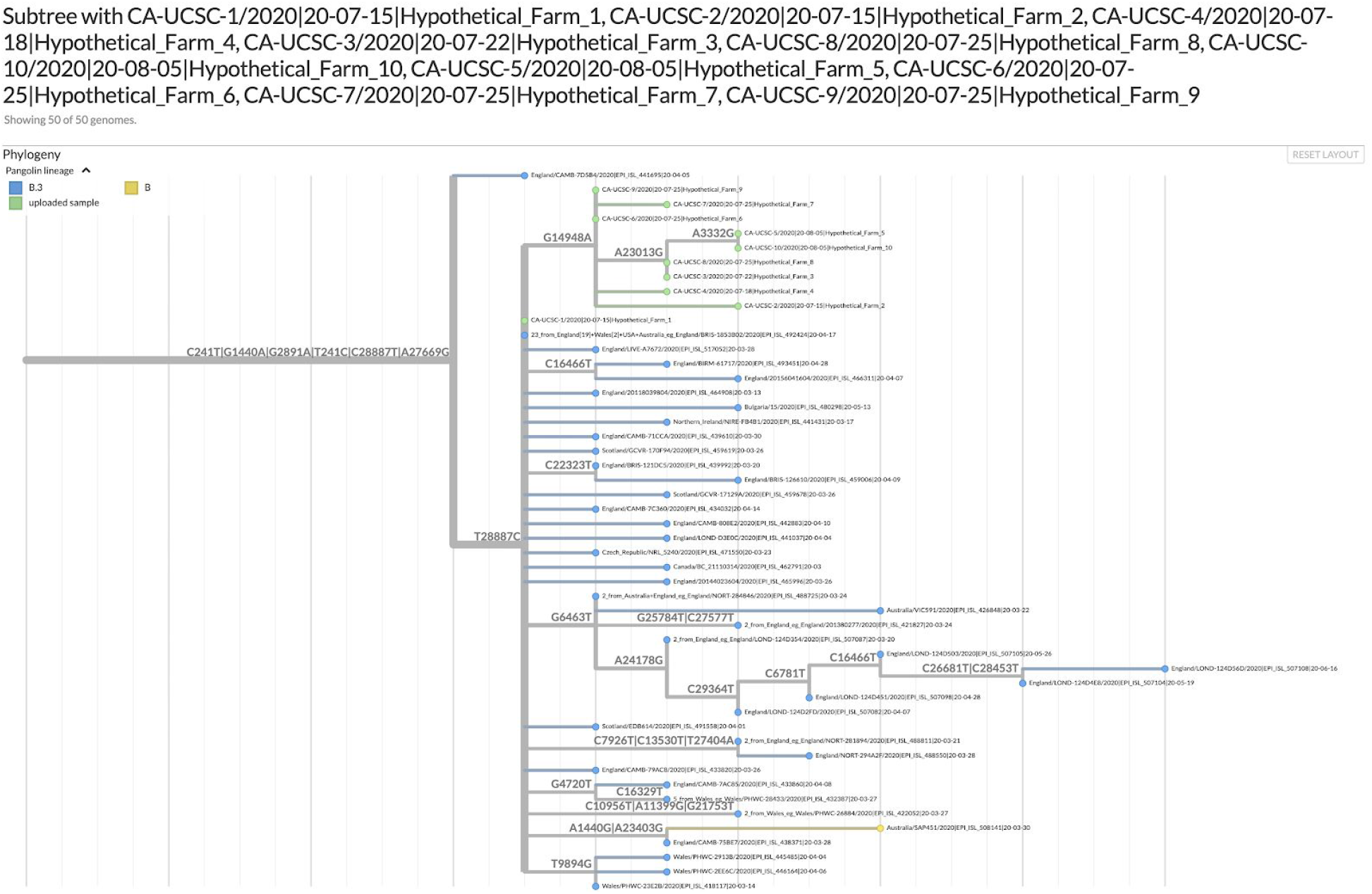
Auspice view of subtree created by UShER placing the same hypothetical example samples as in Figure 5. Temporary link (herokuapp.com temporary host will be nextstrain.org when the new /fetch/ feature is released, https://github.com/nextstrain/nextstrain.org/pull/216): https://nextstrain-s-fetch-sour-8nfiar.herokuapp.com/fetch/hgwdev.gi.ucsc.edu/~angie/usher_example.json

