## Supplementary material for "Ultrafast Sample Placement on Existing Trees (UShER) Empowers Real-Time Phylogenetics for the SARS-CoV-2 Pandemic": File_S6

We gratefully acknowledge the following Authors from the Originating laboratories responsible for obtaining the specimens, as well as the Submitting laboratories where the genome data were generated and shared via GISAID, on which this research is based.

All Submitters of data may be contacted directly via www.gisaid.org

| Accession ID | Originating Laboratory | Submitting Laboratory | Authors |
|---|---|---|---|
| EPI_ISL_495917, EPI_ISL_495918, EPI_ISL_495919, EPI_ISL_495920, EPI_ISL_495921, EPI_ISL_495922, EPI_ISL_495923, EPI_ISL_495924, EPI_ISL_495925, EPI_ISL_495926, EPI_ISL_495927, EPI_ISL_495928, EPI_ISL_495929, EPI_ISL_495930, EPI_ISL_495931, EPI_ISL_495932, EPI_ISL_495933, EPI_ISL_495934, EPI_ISL_495935, EPI_ISL_495936, EPI_ISL_495937, EPI_ISL_495938, EPI_ISL_495939, EPI_ISL_495940, EPI_ISL_495941, EPI_ISL_495942, EPI_ISL_495943, EPI_ISL_495944, EPI_ISL_495945, EPI_ISL_495946, EPI_ISL_495947, EPI_ISL_495948, EPI_ISL_495949, EPI_ISL_495950, EPI_ISL_495951, EPI_ISL_495952, EPI_ISL_495953, EPI_ISL_495954, EPI_ISL_495955, EPI_ISL_495956, EPI_ISL_495957, EPI_ISL_495958, EPI_ISL_495959, EPI_ISL_495960, EPI_ISL_495961, EPI_ISL_495962, EPI_ISL_495963, EPI_ISL_495964, EPI_ISL_495965, EPI_ISL_495966, EPI_ISL_495967, EPI_ISL_495968, EPI_ISL_495969, EPI_ISL_495970, EPI_ISL_495971, EPI_ISL_495972, EPI_ISL_495973, EPI_ISL_495974, EPI_ISL_495975, EPI_ISL_495976, EPI_ISL_495977, EPI_ISL_495978, EPI_ISL_495979, EPI_ISL_495980, EPI_ISL_495981, EPI_ISL_495982, EPI_ISL_495983, EPI_ISL_495984, EPI_ISL_495985, EPI_ISL_495986, EPI_ISL_495987, EPI_ISL_495988, EPI_ISL_495989, EPI_ISL_495990, EPI_ISL_495991, EPI_ISL_495992, EPI_ISL_495993, EPI_ISL_495994, EPI_ISL_495995, EPI_ISL_495996, EPI_ISL_495997, EPI_ISL_495998, EPI_ISL_495999, EPI_ISL_496000, EPI_ISL_496001, EPI_ISL_496002, EPI_ISL_496003, EPI_ISL_496004, EPI_ISL_496005, EPI_ISL_496006, EPI_ISL_496007, EPI_ISL_496008, EPI_ISL_496009, EPI_ISL_496010, EPI_ISL_496011, EPI_ISL_496012, EPI_ISL_496013, EPI_ISL_496014, EPI_ISL_496015, EPI_ISL_496016, EPI_ISL_496017, EPI_ISL_496018, EPI_ISL_496019, EPI_ISL_496020, EPI_ISL_496021, EPI_ISL_496022, EPI_ISL_496023, EPI_ISL_496024, EPI_ISL_496025, EPI_ISL_496026, EPI_ISL_496027, EPI_ISL_496028, EPI_ISL_496029, EPI_ISL_496030, EPI_ISL_496031, EPI_ISL_496032, EPI_ISL_496033, EPI_ISL_496034, EPI_ISL_496035, EPI_ISL_496036, EPI_ISL_496037, EPI_ISL_496038, EPI_ISL_496039, EPI_ISL_496040, EPI_ISL_496041, EPI_ISL_496042, EPI_ISL_496043, EPI_ISL_496044, EPI_ISL_496045, EPI_ISL_496046, EPI_ISL_496047, EPI_ISL_496048, EPI_ISL_496049, EPI_ISL_496050, EPI_ISL_496051, EPI_ISL_496052, EPI_ISL_496053, EPI_ISL_496054, EPI_ISL_496055, EPI_ISL_496056, EPI_ISL_496057, EPI_ISL_496058, EPI_ISL_496059, EPI_ISL_496060, EPI_ISL_496061, EPI_ISL_496062, EPI_ISL_496063, EPI_ISL_496064, EPI_ISL_496065, EPI_ISL_496066, EPI_ISL_496067, EPI_ISL_496068, EPI_ISL_496069, EPI_ISL_496070, EPI_ISL_496071, EPI_ISL_496072, EPI_ISL_496073, EPI_ISL_496074, EPI_ISL_496075, EPI_ISL_496076, EPI_ISL_496077, EPI_ISL_496078, EPI_ISL_496079, EPI_ISL_496080, EPI_ISL_496081, EPI_ISL_496082, EPI_ISL_496083, EPI_ISL_496084, EPI_ISL_496085, EPI_ISL_496086, EPI_ISL_496087, EPI_ISL_496088, EPI_ISL_496089, EPI_ISL_496090, EPI_ISL_496091, EPI_ISL_496092, EPI_ISL_496093, EPI_ISL_496094, EPI_ISL_496095, EPI_ISL_496096, EPI_ISL_496097, EPI_ISL_496098, EPI_ISL_496099, EPI_ISL_496100, EPI_ISL_496101, EPI_ISL_496102, EPI_ISL_496103, EPI_ISL_496104, EPI_ISL_496105, EPI_ISL_496106, EPI_ISL_496107, EPI_ISL_496108, EPI_ISL_496109, EPI_ISL_496110, EPI_ISL_496111, EPI_ISL_496112, EPI_ISL_496113, EPI_ISL_496114, EPI_ISL_496115, EPI_ISL_496116, EPI_ISL_496117, EPI_ISL_496118, EPI_ISL_496119, EPI_ISL_496120, EPI_ISL_496121, EPI_ISL_496122, EPI_ISL_496123, EPI_ISL_496124, EPI_ISL_496125, EPI_ISL_496126, EPI_ISL_496127, EPI_ISL_496128, EPI_ISL_496129, EPI_ISL_496130, EPI_ISL_496131, EPI_ISL_496132, EPI_ISL_496133, EPI_ISL_496134, EPI_ISL_496135, EPI_ISL_496136, EPI_ISL_496137, EPI_ISL_496138, EPI_ISL_496139, EPI_ISL_496140, EPI_ISL_496141, EPI_ISL_496142, EPI_ISL_496143, EPI_ISL_496144, EPI_ISL_496145, EPI_ISL_496146, EPI_ISL_496147, EPI_ISL_496148, EPI_ISL_496149, EPI_ISL_496150, EPI_ISL_496151, EPI_ISL_496152, EPI_ISL_496153, EPI_ISL_496154, EPI_ISL_496155, EPI_ISL_496156, EPI_ISL_496157, EPI_ISL_496158, EPI_ISL_496159, EPI_ISL_496160, EPI_ISL_496161, EPI_ISL_496162, EPI_ISL_496163, EPI_ISL_496164, EPI_ISL_496165, EPI_ISL_496166, EPI_ISL_496167, EPI_ISL_496168, EPI_ISL_496169, EPI_ISL_496170, EPI_ISL_496171, EPI_ISL_496172, EPI_ISL_496173, EPI_ISL_496174, EPI_ISL_496175, EPI_ISL_496176, EPI_ISL_496177, EPI_ISL_496178, EPI_ISL_496179, EPI_ISL_496180, EPI_ISL_496181, EPI_ISL_496182, EPI_ISL_496183, EPI_ISL_496184, EPI_ISL_496185, EPI_ISL_496186, EPI_ISL_496187, EPI_ISL_496188, EPI_ISL_496189, EPI_ISL_496190, EPI_ISL_496191, EPI_ISL_496192, EPI_ISL_496193, EPI_ISL_496194, EPI_ISL_496195, EPI_ISL_496196, EPI_ISL_496197, EPI_ISL_496198, EPI_ISL_496199, EPI_ISL_496200, EPI_ISL_496201, EPI_ISL_496202, EPI_ISL_496203, EPI_ISL_496204, EPI_ISL_496205, EPI_ISL_496206, EPI_ISL_496207, EPI_ISL_496208, EPI_ISL_496209, EPI_ISL_496210, EPI_ISL_496211, EPI_ISL_496212, EPI_ISL_496213, EPI_ISL_496214, EPI_ISL_496215, EPI_ISL_496216, EPI_ISL_496217, EPI_ISL_496218, EPI_ISL_496219, EPI_ISL_496220, EPI_ISL_496221, EPI_ISL_496222, EPI_ISL_496223, EPI_ISL_496224, EPI_ISL_496225, EPI_ISL_496226, EPI_ISL_496227, EPI_ISL_496228, EPI_ISL_496229, EPI_ISL_496230, EPI_ISL_496231, EPI_ISL_496232, EPI_ISL_496233, EPI_ISL_496234, EPI_ISL_496235, EPI_ISL_496236, EPI_ISL_496237, EPI_ISL_496238, EPI_ISL_496239, EPI_ISL_496240, EPI_ISL_496241, EPI_ISL_496242, EPI_ISL_496243, EPI_ISL_496244, EPI_ISL_496245, EPI_ISL_496246, EPI_ISL_496247, EPI_ISL_496248, EPI_ISL_496249, EPI_ISL_496250, EPI_ISL_496251, EPI_ISL_496252, EPI_ISL_496253, EPI_ISL_496254, EPI_ISL_496255, EPI_ISL_496256, EPI_ISL_496257, EPI_ISL_496258, EPI_ISL_496259, EPI_ISL_496260, EPI_ISL_496261, EPI_ISL_496262, EPI_ISL_496263, EPI_ISL_496264, EPI_ISL_496265, EPI_ISL_496266, EPI_ISL_496267, EPI_ISL_496268, EPI_ISL_496269, EPI_ISL_496270, EPI_ISL_496271, EPI_ISL_496272, EPI_ISL_496273, EPI_ISL_496274, EPI_ISL_496275, EPI_ISL_496276, EPI_ISL_496277, EPI_ISL_496278, EPI_ISL_496279, EPI_ISL_496280, EPI_ISL_496281, EPI_ISL_496282, EPI_ISL_496283, EPI_ISL_496284, EPI_ISL_496285, EPI_ISL_496286, EPI_ISL_496287, EPI_ISL_496288, EPI_ISL_496289, EPI_ISL_496290, EPI_ISL_496291, EPI_ISL_496292, EPI_ISL_496293, EPI_ISL_496294, EPI_ISL_496295, EPI_ISL_496296, EPI_ISL_496297, EPI_ISL_496298, EPI_ISL_496299, EPI_ISL_496300, EPI_ISL_496301, EPI_ISL_496302, EPI_ISL_496303, EPI_ISL_496304, EPI_ISL_496305, EPI_ISL_496306, EPI_ISL_496307, EPI_ISL_496308, EPI_ISL_496309, EPI_ISL_496310, EPI_ISL_496311, EPI_ISL_496312, EPI_ISL_496313, EPI_ISL_496314, EPI_ISL_496315, EPI_ISL_496316, EPI_ISL_496317, EPI_ISL_496318, EPI_ISL_496319, EPI_ISL_496320, EPI_ISL_496321, EPI_ISL_496322, EPI_ISL_496323, EPI_ISL_496324, EPI_ISL_496325, EPI_ISL_496326, EPI_ISL_496327, EPI_ISL_496328, EPI_ISL_496329, EPI_ISL_496330, EPI_ISL_496331, EPI_ISL_496332, EPI_ISL_496333, EPI_ISL_496334, EPI_ISL_496335, EPI_ISL_496336, EPI_ISL_496337, EPI_ISL_496338 | | | |
| see above | Washington State Department of Health | Seattle Flu Study | Deborah A. Nickerson, Chris D. Frazar, Jover Lee, Benjamin Pelle, Matthew Richardson, Amanda Adler, Elisabeth Brandstetter, Peter D. Han, Kairsten Fay, Misja Ilcisin, Kirsten Lacombe, Thomas R. Sibley, Melissa Truong, Caitlin R. Wolf, Romesh Gautom, Geoff Melly, Brian Hiatt, Philip Dykema, Scott Lindquist, Michael Boeckh, Janet A. Englund, Michael Famulare, Barry R. Lutz, Mark J. Rieder, Lea M. Starita, Matthew Thompson, Helen Y. Chu, Jay Shendure, Trevor Bedford |
| EPI_ISL_496339, EPI_ISL_496340, EPI_ISL_496341, EPI_ISL_496342, EPI_ISL_496343, EPI_ISL_496344, EPI_ISL_496345, EPI_ISL_496346, EPI_ISL_496347, EPI_ISL_496348, EPI_ISL_496349, EPI_ISL_496350, EPI_ISL_496351, EPI_ISL_496352, EPI_ISL_496353, EPI_ISL_496354, EPI_ISL_496355, EPI_ISL_496356, EPI_ISL_496357, EPI_ISL_496358, EPI_ISL_496359, EPI_ISL_496360, EPI_ISL_496361, EPI_ISL_496362, EPI_ISL_496363, EPI_ISL_496364, EPI_ISL_496365, EPI_ISL_496366, EPI_ISL_496367, EPI_ISL_496368, EPI_ISL_496369, EPI_ISL_496370, EPI_ISL_496371, EPI_ISL_496372, EPI_ISL_496373, EPI_ISL_496374, EPI_ISL_496375, EPI_ISL_496376 | | | |
| see above | Infectolab | Andersen lab at Scripps Research | SEARCH Alliance San Diego with Samuel Navarro Alvarez, Carlos A. Cota Haros, Octavio Renteria Pacheco |
| EPI_ISL_496482 | Dept. Infectious, Tropical Diseases & Microbiology, IRCCS Sacro Cuore Don Calabria Hospital | 1) Dept. Infectious, Tropical Diseases & Microbiology, IRCCS Sacro Cuore Don Calabria Hospital; 2) Centro Piattaforme Tecnologiche, University of Verona; 3) Dept. Neurosciences, Biomedicine and Movement Sciences, University of Verona. | 1) Antonio Mori, Michela Deiana, Elena Pomari, Chiara Piubelli; 2) Monica Castellucci and Francesca Griggio; 3) Giovanni Malerba |
| EPI_ISL_496491, EPI_ISL_496492, EPI_ISL_496493, EPI_ISL_496494, EPI_ISL_496495, EPI_ISL_496496, EPI_ISL_496497, EPI_ISL_496498, EPI_ISL_496499, EPI_ISL_496500, EPI_ISL_496501, EPI_ISL_496502, EPI_ISL_496503, EPI_ISL_496504, EPI_ISL_496505, EPI_ISL_496506, EPI_ISL_496507, EPI_ISL_496508, EPI_ISL_496509, EPI_ISL_496510, EPI_ISL_496511, EPI_ISL_496512, EPI_ISL_496513, EPI_ISL_496514, EPI_ISL_496515, EPI_ISL_496516, EPI_ISL_496517 | | | |
| see above | Viral Respiratory Lab, National Institute for Biomedical Research (INRB) | Pathogen Sequencing Lab, National Institute for Biomedical Research (INRB) | Placide Mbala-Kingebeni, Edith Nkwembe, Eddy Kinganda-Lusamaki, Amuri Aziza, Francisca Muyembe Mawete, Emmanuel Lokilo Lofiko, Catherine Pratt, Matthias Pauthner, Josh Quick, Allison Black, James Hadfield, Trevor Bedford, Ian Goodfellow, Andrew Rambaut, Nick Loman, Kristian Andersen, Michael Wiley, Steve Ahuka-Mundeke, Jean-Jacques Muyembe Tamfum |
| EPI_ISL_496518, EPI_ISL_496519, EPI_ISL_496520 | Armed Forces Medical College | National Centre For Cell Science | Dhiraj Paul, Kunal Jani, Radha Chauhan, Janesh Kumar, Vasudevan Seshadri, Girdhari Lal, Rajesh Karyakarte, Suvarna Joshi, Murlidhar Tambe, Sourav Sen, Santosh Karade, Kavita Bala Anand, Shelinder Pal Singh Shergill, Rajiv Mohan Gupta, Manoj Kumar Bhat, Arvind Sahu, Maharashtra COVID-19 Study Group, DBT's PAN-INDIA 1000 SARS-CoV2 RNA genome sequencing consortium, Yogesh S Shouche |
| EPI_ISL_496521, EPI_ISL_496522, EPI_ISL_496523 | B.J. Govt. Medical College | National Centre For Cell Science | Dhiraj Paul, Kunal Jani, Radha Chauhan, Janesh Kumar, Vasudevan Seshadri, Girdhari Lal, Rajesh Karyakarte, Suvarna Joshi, Murlidhar Tambe, Sourav Sen, Santosh Karade, Kavita Bala Anand, Shelinder Pal Singh Shergill, Rajiv Mohan Gupta, Manoj Kumar Bhat, Arvind Sahu, Maharashtra COVID-19 Study Group, DBT's PAN-INDIA 1000 SARS-CoV2 RNA genome sequencing consortium, Yogesh S Shouche |
| EPI_ISL_496524, EPI_ISL_496525, EPI_ISL_496526 | National Centre For Cell Science | National Centre For Cell Science | Dhiraj Paul, Kunal Jani, Radha Chauhan, Janesh Kumar, Vasudevan Seshadri, Girdhari Lal, Rajesh Karyakarte, Suvarna Joshi, Murlidhar Tambe, Sourav Sen, Santosh Karade, Kavita Bala Anand, Shelinder Pal Singh Shergill, Rajiv Mohan Gupta, Manoj Kumar Bhat, Arvind Sahu, Maharashtra COVID-19 Study Group, DBT's PAN-INDIA 1000 SARS-CoV2 RNA genome sequencing consortium, Yogesh S Shouche |
| EPI_ISL_496527, EPI_ISL_496528 | B.J. Govt. Medical College | National Centre For Cell Science | Dhiraj Paul, Kunal Jani, Radha Chauhan, Janesh Kumar, Vasudevan Seshadri, Girdhari Lal, Rajesh Karyakarte, Suvarna Joshi, Murlidhar Tambe, Sourav Sen, Santosh Karade, Kavita Bala Anand, Shelinder Pal Singh Shergill, Rajiv Mohan Gupta, Manoj Kumar Bhat, Arvind Sahu, Maharashtra COVID-19 Study Group, DBT's PAN-INDIA 1000 SARS-CoV2 RNA genome sequencing consortium, Yogesh S Shouche |
| EPI_ISL_496529 | Armed Forces Medical College | National Centre For Cell Science | Dhiraj Paul, Kunal Jani, Radha Chauhan, Janesh Kumar, Vasudevan Seshadri, Girdhari Lal, Rajesh Karyakarte, Suvarna Joshi, Murlidhar Tambe, Sourav Sen, Santosh Karade, Kavita Bala Anand, Shelinder Pal Singh Shergill, Rajiv Mohan Gupta, Manoj Kumar Bhat, Arvind Sahu, Maharashtra COVID-19 Study Group, DBT's PAN-INDIA 1000 SARS-CoV2 RNA genome sequencing consortium, Yogesh S Shouche |
| EPI_ISL_496530 | B.J. Govt. Medical College | National Centre For Cell Science | Dhiraj Paul, Kunal Jani, Radha Chauhan, Janesh Kumar, Vasudevan Seshadri, Girdhari Lal, Rajesh Karyakarte, Suvarna Joshi, Murlidhar Tambe, Sourav Sen, Santosh Karade, Kavita Bala Anand, Shelinder Pal Singh Shergill, Rajiv Mohan Gupta, Manoj Kumar Bhat, Arvind Sahu, Maharashtra COVID-19 Study Group, DBT's PAN-INDIA 1000 SARS-CoV2 RNA genome sequencing consortium, Yogesh S Shouche |
| EPI_ISL_496531, EPI_ISL_496532 | National Centre For Cell Science | National Centre For Cell Science | Dhiraj Paul, Kunal Jani, Radha Chauhan, Janesh Kumar, Vasudevan Seshadri, Girdhari Lal, Rajesh Karyakarte, Suvarna Joshi, Murlidhar Tambe, Sourav Sen, Santosh Karade, Kavita Bala Anand, Shelinder Pal Singh Shergill, Rajiv Mohan Gupta, Manoj Kumar Bhat, Arvind Sahu, Maharashtra COVID-19 Study Group, DBT's PAN-INDIA 1000 SARS-CoV2 RNA genome sequencing consortium, Yogesh S Shouche |
| EPI_ISL_496533 | Armed Forces Medical College | National Centre For Cell Science | Dhiraj Paul, Kunal Jani, Radha Chauhan, Janesh Kumar, Vasudevan Seshadri, Girdhari Lal, Rajesh Karyakarte, Suvarna Joshi, Murlidhar Tambe, Sourav Sen, Santosh Karade, Kavita Bala Anand, Shelinder Pal Singh Shergill, Rajiv Mohan Gupta, Manoj Kumar Bhat, Arvind Sahu, Maharashtra COVID-19 Study Group, DBT's PAN-INDIA 1000 SARS-CoV2 RNA genome sequencing consortium, Yogesh S Shouche |
| EPI_ISL_496534 | B.J. Govt. Medical College | National Centre For Cell Science | Dhiraj Paul, Kunal Jani, Radha Chauhan, Janesh Kumar, Vasudevan Seshadri, Girdhari Lal, Rajesh Karyakarte, Suvarna Joshi, Murlidhar Tambe, Sourav Sen, Santosh Karade, Kavita Bala Anand, Shelinder Pal Singh Shergill, Rajiv Mohan Gupta, Manoj Kumar Bhat, Arvind Sahu, Maharashtra COVID-19 Study Group, DBT's PAN-INDIA 1000 SARS-CoV2 RNA genome sequencing consortium, Yogesh S Shouche |
| EPI_ISL_496535, EPI_ISL_496536 | National Centre For Cell Science | National Centre For Cell Science | Dhiraj Paul, Kunal Jani, Radha Chauhan, Janesh Kumar, Vasudevan Seshadri, Girdhari Lal, Rajesh Karyakarte, Suvarna Joshi, Murlidhar Tambe, Sourav Sen, Santosh Karade, Kavita Bala Anand, Shelinder Pal Singh Shergill, Rajiv Mohan Gupta, Manoj Kumar Bhat, Arvind Sahu, Maharashtra COVID-19 Study Group, DBT's PAN-INDIA 1000 SARS-CoV2 RNA genome sequencing consortium, Yogesh S Shouche |
| EPI_ISL_496537, EPI_ISL_496538, EPI_ISL_496539, EPI_ISL_496540, EPI_ISL_496541, EPI_ISL_496542, EPI_ISL_496543, EPI_ISL_496544, EPI_ISL_496545 | Armed Forces Medical College | National Centre For Cell Science | Dhiraj Paul, Kunal Jani, Radha Chauhan, Janesh Kumar, Vasudevan Seshadri, Girdhari Lal, Rajesh Karyakarte, Suvarna Joshi, Murlidhar Tambe, Sourav Sen, Santosh Karade, Kavita Bala Anand, Shelinder Pal Singh Shergill, Rajiv Mohan Gupta, Manoj Kumar Bhat, Arvind Sahu, Maharashtra COVID-19 Study Group, DBT's PAN-INDIA 1000 SARS-CoV2 RNA genome sequencing consortium, Yogesh S Shouche |
| EPI_ISL_496546, EPI_ISL_496547, EPI_ISL_496548, EPI_ISL_496549, EPI_ISL_496550, EPI_ISL_496551, EPI_ISL_496552, EPI_ISL_496553, EPI_ISL_496554 | B.J. Govt. Medical College | National Centre For Cell Science | Dhiraj Paul, Kunal Jani, Radha Chauhan, Janesh Kumar, Vasudevan Seshadri, Girdhari Lal, Rajesh Karyakarte, Suvarna Joshi, Murlidhar Tambe, Sourav Sen, Santosh Karade, Kavita Bala Anand, Shelinder Pal Singh Shergill, Rajiv Mohan Gupta, Manoj Kumar Bhat, Arvind Sahu, Maharashtra COVID-19 Study Group, DBT's PAN-INDIA 1000 SARS-CoV2 RNA genome sequencing consortium, Yogesh S Shouche |
| EPI_ISL_496555, EPI_ISL_496556, EPI_ISL_496557, EPI_ISL_496558, EPI_ISL_496559, EPI_ISL_496560, EPI_ISL_496561, EPI_ISL_496562, EPI_ISL_496563, EPI_ISL_496564, EPI_ISL_496565, EPI_ISL_496566, EPI_ISL_496567, EPI_ISL_496568, EPI_ISL_496569, EPI_ISL_496570, EPI_ISL_496571, EPI_ISL_496572, EPI_ISL_496573, EPI_ISL_496574, EPI_ISL_496575, EPI_ISL_496576, EPI_ISL_496577, EPI_ISL_496578, EPI_ISL_496579, EPI_ISL_496580, EPI_ISL_496581, EPI_ISL_496582, EPI_ISL_496583, EPI_ISL_496584, EPI_ISL_496585, EPI_ISL_496586, EPI_ISL_496587 | | | |
| see above | National Centre For Cell Science | National Centre For Cell Science | Dhiraj Paul, Kunal Jani, Radha Chauhan, Janesh Kumar, Vasudevan Seshadri, Girdhari Lal, Rajesh Karyakarte, Suvarna Joshi, Murlidhar Tambe, Sourav Sen, Santosh Karade, Kavita Bala Anand, Shelinder Pal Singh Shergill, Rajiv Mohan Gupta, Manoj Kumar Bhat, Arvind Sahu, Maharashtra COVID-19 Study Group, DBT's PAN-INDIA 1000 SARS-CoV2 RNA genome sequencing consortium, Yogesh S Shouche |
| EPI_ISL_496602 | Armed Forces Medical College | National Centre For Cell Science | Dhiraj Paul, Kunal Jani, Radha Chauhan, Janesh Kumar, Vasudevan Seshadri, Girdhari Lal, Rajesh Karyakarte, Suvarna Joshi, Murlidhar Tambe, Sourav Sen, Santosh Karade, Kavita Bala Anand, Shelinder Pal Singh Shergill, Rajiv Mohan Gupta, Manoj Kumar Bhat, Arvind Sahu, Maharashtra COVID-19 Study Group, DBT's PAN-INDIA 1000 SARS-CoV2 RNA genome sequencing consortium, Yogesh S Shouche |
| EPI_ISL_496603, EPI_ISL_496604, EPI_ISL_496605, EPI_ISL_496606, EPI_ISL_496607, EPI_ISL_496608, EPI_ISL_496609, EPI_ISL_496610, EPI_ISL_496611, EPI_ISL_496612, EPI_ISL_496613, EPI_ISL_496614, EPI_ISL_496615, EPI_ISL_496616, EPI_ISL_496617, EPI_ISL_496618, EPI_ISL_496619, EPI_ISL_496620, EPI_ISL_496621, EPI_ISL_496622, EPI_ISL_496623, EPI_ISL_496624, EPI_ISL_496625, EPI_ISL_496626, EPI_ISL_496627, EPI_ISL_496628, EPI_ISL_496629, EPI_ISL_496630, EPI_ISL_496631, EPI_ISL_496632, EPI_ISL_496633, EPI_ISL_496634, EPI_ISL_496635, EPI_ISL_496636, EPI_ISL_496637, EPI_ISL_496638, EPI_ISL_496639, EPI_ISL_496640, EPI_ISL_496641, EPI_ISL_496642, EPI_ISL_496643, EPI_ISL_496644, EPI_ISL_496645, EPI_ISL_496646, EPI_ISL_496647, EPI_ISL_496648, EPI_ISL_496649, EPI_ISL_496650, EPI_ISL_496651, EPI_ISL_496652, EPI_ISL_496653, EPI_ISL_496654, EPI_ISL_496655, EPI_ISL_496656, EPI_ISL_496657, EPI_ISL_496658, EPI_ISL_496659, EPI_ISL_496660, EPI_ISL_496661, EPI_ISL_496662, EPI_ISL_496663, EPI_ISL_496664, EPI_ISL_496665, EPI_ISL_496666, EPI_ISL_496667, EPI_ISL_496668, EPI_ISL_496669, EPI_ISL_496670, EPI_ISL_496671, EPI_ISL_496672, EPI_ISL_496673, EPI_ISL_496674, EPI_ISL_496675, EPI_ISL_496676, EPI_ISL_496677, EPI_ISL_496678, EPI_ISL_496679, EPI_ISL_496680, EPI_ISL_496681, EPI_ISL_496682, EPI_ISL_496683, EPI_ISL_496684, EPI_ISL_496685, EPI_ISL_496686, EPI_ISL_496687, EPI_ISL_496688, EPI_ISL_496689, EPI_ISL_496690, EPI_ISL_496691, EPI_ISL_496692, EPI_ISL_496693, EPI_ISL_496694, EPI_ISL_496695, EPI_ISL_496696, EPI_ISL_496697, EPI_ISL_496698, EPI_ISL_496699, EPI_ISL_496700, EPI_ISL_496701, EPI_ISL_496702, EPI_ISL_496703, EPI_ISL_496704, EPI_ISL_496705, EPI_ISL_496706, EPI_ISL_496707, EPI_ISL_496708, EPI_ISL_496709, EPI_ISL_496710, EPI_ISL_496711, EPI_ISL_496712, EPI_ISL_496713, EPI_ISL_496714, EPI_ISL_496715, EPI_ISL_496716, EPI_ISL_496717, EPI_ISL_496718, EPI_ISL_496719, EPI_ISL_496720, EPI_ISL_496721, EPI_ISL_496722, EPI_ISL_496723, EPI_ISL_496724, EPI_ISL_496725, EPI_ISL_496726, EPI_ISL_496727, EPI_ISL_496728, EPI_ISL_496729, EPI_ISL_496730, EPI_ISL_496731, EPI_ISL_496732, EPI_ISL_496733, EPI_ISL_496734, EPI_ISL_496735, EPI_ISL_496736, EPI_ISL_496737, EPI_ISL_496738, EPI_ISL_496739, EPI_ISL_496740, EPI_ISL_496741, EPI_ISL_496742, EPI_ISL_496743, EPI_ISL_496744, EPI_ISL_496745, EPI_ISL_496746, EPI_ISL_496747, EPI_ISL_496748, EPI_ISL_496749, EPI_ISL_496750, EPI_ISL_496751, EPI_ISL_496752, EPI_ISL_496753, EPI_ISL_496754, EPI_ISL_496755, EPI_ISL_496756, EPI_ISL_496757, EPI_ISL_496758, EPI_ISL_496759, EPI_ISL_496760, EPI_ISL_496761, EPI_ISL_496762, EPI_ISL_496763, EPI_ISL_496764, EPI_ISL_496765, EPI_ISL_496766, EPI_ISL_496767, EPI_ISL_496768, EPI_ISL_496769, EPI_ISL_496770, EPI_ISL_496771, EPI_ISL_496772, EPI_ISL_496773, EPI_ISL_496774, EPI_ISL_496775, EPI_ISL_496776, EPI_ISL_496777, EPI_ISL_496778, EPI_ISL_496779, EPI_ISL_496780, EPI_ISL_496781, EPI_ISL_496782, EPI_ISL_496783, EPI_ISL_496784, EPI_ISL_496785, EPI_ISL_496786, EPI_ISL_496787, EPI_ISL_496788, EPI_ISL_496789, EPI_ISL_496790, EPI_ISL_496791, EPI_ISL_496792, EPI_ISL_496793, EPI_ISL_496794, EPI_ISL_496795, EPI_ISL_496796, EPI_ISL_496797, EPI_ISL_496798, EPI_ISL_496799, EPI_ISL_496800, EPI_ISL_496801, EPI_ISL_496802, EPI_ISL_496803, EPI_ISL_496804, EPI_ISL_496805, EPI_ISL_496806, EPI_ISL_496807, EPI_ISL_496808, EPI_ISL_496809, EPI_ISL_496810, EPI_ISL_496811, EPI_ISL_496812, EPI_ISL_496813, EPI_ISL_496814, EPI_ISL_496815, EPI_ISL_496816, EPI_ISL_496817, EPI_ISL_496818, EPI_ISL_496819, EPI_ISL_496820, EPI_ISL_496821, EPI_ISL_496822, EPI_ISL_496823, EPI_ISL_496824, EPI_ISL_496825, EPI_ISL_496826, EPI_ISL_496827, EPI_ISL_496828, EPI_ISL_496829, EPI_ISL_496830, EPI_ISL_496831, EPI_ISL_496832, EPI_ISL_496833, EPI_ISL_496834, EPI_ISL_496835, EPI_ISL_496836, EPI_ISL_496837, EPI_ISL_496838, EPI_ISL_496839, EPI_ISL_496840, EPI_ISL_496841, EPI_ISL_496842, EPI_ISL_496843, EPI_ISL_496844, EPI_ISL_496845, EPI_ISL_496846, EPI_ISL_496847, EPI_ISL_496848, EPI_ISL_496849, EPI_ISL_496850, EPI_ISL_496851, EPI_ISL_496852, EPI_ISL_496853, EPI_ISL_496854, EPI_ISL_496855, EPI_ISL_496856, EPI_ISL_496857, EPI_ISL_496858, EPI_ISL_496859, EPI_ISL_496860, EPI_ISL_496861, EPI_ISL_496862, EPI_ISL_496863, EPI_ISL_496864, EPI_ISL_496865, EPI_ISL_496866, EPI_ISL_496867, EPI_ISL_496868, EPI_ISL_496869, EPI_ISL_496870, EPI_ISL_496871, EPI_ISL_496872, EPI_ISL_496873, EPI_ISL_496874, EPI_ISL_496875, EPI_ISL_496876, EPI_ISL_496877, EPI_ISL_496878, EPI_ISL_496879, EPI_ISL_496880, EPI_ISL_496881, EPI_ISL_496882, EPI_ISL_496883, EPI_ISL_496884, EPI_ISL_496885, EPI_ISL_496886, EPI_ISL_496887, EPI_ISL_496888, EPI_ISL_496889, EPI_ISL_496890, EPI_ISL_496891, EPI_ISL_496892, EPI_ISL_496893, EPI_ISL_496894, EPI_ISL_496895, EPI_ISL_496896, EPI_ISL_496897, EPI_ISL_496898, EPI_ISL_496899, EPI_ISL_496900, EPI_ISL_496901, EPI_ISL_496902, EPI_ISL_496903, EPI_ISL_496904, EPI_ISL_496905, EPI_ISL_496906, EPI_ISL_496907, EPI_ISL_496908, EPI_ISL_496909, EPI_ISL_496910, EPI_ISL_496911, EPI_ISL_496912, EPI_ISL_496913, EPI_ISL_496914, EPI_ISL_496915 | | | |
| see above | Gorgas Memorial Laboratory of Health Studies | Gorgas Memorial Laboratory of Health Studies | Danilo Franco, Claudia Gonzalez Sandra Lopez-Verges, Alexander A Martinez |
| EPI_ISL_496916, EPI_ISL_496917 | Minnesota Department of Health, Public Health Laboratory | Minnesota Department of Health, Public Health Laboratory | Matt Plumb, Jacob Garfin, and Xiong Wang |
| EPI_ISL_496918 | Mayo Clinic & Mayo Clinic Laboratories | Minnesota Department of Health, Public Health Laboratory | Matt Plumb, Jacob Garfin, and Xiong Wang |
| EPI_ISL_496919, EPI_ISL_496920, EPI_ISL_496921, EPI_ISL_496922, EPI_ISL_496923, EPI_ISL_496924, EPI_ISL_496925, EPI_ISL_496926, EPI_ISL_496927, EPI_ISL_496928, EPI_ISL_496929, EPI_ISL_496930 | | | |
| see above | Minnesota Department of Health, Public Health Laboratory | Minnesota Department of Health, Public Health Laboratory | Matt Plumb, Jacob Garfin, and Xiong Wang |
| EPI_ISL_496931, EPI_ISL_496932, EPI_ISL_496933, EPI_ISL_496934, EPI_ISL_496935, EPI_ISL_496936, EPI_ISL_496937, EPI_ISL_496938, EPI_ISL_496939, EPI_ISL_496940, EPI_ISL_496941, EPI_ISL_496942, EPI_ISL_496943, EPI_ISL_496944, EPI_ISL_496945, EPI_ISL_496946, EPI_ISL_496947, EPI_ISL_496948, EPI_ISL_496949, EPI_ISL_496950, EPI_ISL_496951, EPI_ISL_496952, EPI_ISL_496953, EPI_ISL_496954, | | | |

| | | | |
|---|---|---|---|
| EPI_ISL_496955, EPI_ISL_496956, EPI_ISL_496957, EPI_ISL_496958, EPI_ISL_496959, EPI_ISL_496960, EPI_ISL_496961, EPI_ISL_496962, EPI_ISL_496963, EPI_ISL_496964, EPI_ISL_496965, EPI_ISL_496966, EPI_ISL_496967, EPI_ISL_496968, EPI_ISL_496969, EPI_ISL_496970, EPI_ISL_496971, EPI_ISL_496972, EPI_ISL_496973, EPI_ISL_496974, EPI_ISL_496975, EPI_ISL_496976, EPI_ISL_496977, EPI_ISL_496978, EPI_ISL_496979, EPI_ISL_496980, EPI_ISL_496981, EPI_ISL_496982, EPI_ISL_496983, EPI_ISL_496984, EPI_ISL_496985, EPI_ISL_496986, EPI_ISL_496987, EPI_ISL_496988, EPI_ISL_496989, EPI_ISL_496990, EPI_ISL_496991, EPI_ISL_496992, EPI_ISL_496993, EPI_ISL_496994, EPI_ISL_496995, EPI_ISL_496996, EPI_ISL_496997, EPI_ISL_496998, EPI_ISL_496999, EPI_ISL_497000, EPI_ISL_497001, EPI_ISL_497002, EPI_ISL_497003, EPI_ISL_497004, EPI_ISL_497005, EPI_ISL_497006, EPI_ISL_497007, EPI_ISL_497008, EPI_ISL_497009, EPI_ISL_497010, EPI_ISL_497011, EPI_ISL_497012, EPI_ISL_497013, EPI_ISL_497014, EPI_ISL_497015, EPI_ISL_497016, EPI_ISL_497017, EPI_ISL_497018, EPI_ISL_497019, EPI_ISL_497020, EPI_ISL_497021, EPI_ISL_497022, EPI_ISL_497023, EPI_ISL_497024, EPI_ISL_497025, EPI_ISL_497026, EPI_ISL_497027, EPI_ISL_497028, EPI_ISL_497029, EPI_ISL_497030, EPI_ISL_497031, EPI_ISL_497032, EPI_ISL_497033, EPI_ISL_497034, EPI_ISL_497035, EPI_ISL_497036, EPI_ISL_497037, EPI_ISL_497038, EPI_ISL_497039, EPI_ISL_497040, EPI_ISL_497041, EPI_ISL_497042, EPI_ISL_497043, EPI_ISL_497044, EPI_ISL_497045, EPI_ISL_497046, EPI_ISL_497047, EPI_ISL_497048, EPI_ISL_497049, EPI_ISL_497050, EPI_ISL_497051, EPI_ISL_497052, EPI_ISL_497053, EPI_ISL_497054, EPI_ISL_497055, EPI_ISL_497056, EPI_ISL_497057, EPI_ISL_497058, EPI_ISL_497059, EPI_ISL_497060, EPI_ISL_497061, EPI_ISL_497062, EPI_ISL_497063, EPI_ISL_497064, EPI_ISL_497065, EPI_ISL_497066, EPI_ISL_497067, EPI_ISL_497068, EPI_ISL_497069, EPI_ISL_497070, EPI_ISL_497071, EPI_ISL_497072, EPI_ISL_497073, EPI_ISL_497074, EPI_ISL_497075, EPI_ISL_497076, EPI_ISL_497077, EPI_ISL_497078, EPI_ISL_497079, EPI_ISL_497080, EPI_ISL_497081, EPI_ISL_497082, EPI_ISL_497083, EPI_ISL_497084, EPI_ISL_497085, EPI_ISL_497086, EPI_ISL_497087, EPI_ISL_497088, EPI_ISL_497089, EPI_ISL_497090, EPI_ISL_497091, EPI_ISL_497092, EPI_ISL_497093, EPI_ISL_497094, EPI_ISL_497095, EPI_ISL_497096, EPI_ISL_497097, EPI_ISL_497098, EPI_ISL_497099, EPI_ISL_497100, EPI_ISL_497101, EPI_ISL_497102, EPI_ISL_497103, EPI_ISL_497104, EPI_ISL_497105, EPI_ISL_497106, EPI_ISL_497107, EPI_ISL_497108, EPI_ISL_497109, EPI_ISL_497110, EPI_ISL_497111, EPI_ISL_497112, EPI_ISL_497113, EPI_ISL_497114, EPI_ISL_497115, EPI_ISL_497116, EPI_ISL_497117, EPI_ISL_497118, EPI_ISL_497119, EPI_ISL_497120, EPI_ISL_497121, EPI_ISL_497122, EPI_ISL_497123, EPI_ISL_497124, EPI_ISL_497125, EPI_ISL_497126, EPI_ISL_497127, EPI_ISL_497128, EPI_ISL_497129, EPI_ISL_497130, EPI_ISL_497131, EPI_ISL_497132, EPI_ISL_497133, EPI_ISL_497134, EPI_ISL_497135, EPI_ISL_497136, EPI_ISL_497137, EPI_ISL_497138, EPI_ISL_497139, EPI_ISL_497140, EPI_ISL_497141, EPI_ISL_497142, EPI_ISL_497143, EPI_ISL_497144, EPI_ISL_497145, EPI_ISL_497146, EPI_ISL_497147, EPI_ISL_497148, EPI_ISL_497149, EPI_ISL_497150, EPI_ISL_497151, EPI_ISL_497152, EPI_ISL_497153, EPI_ISL_497154, EPI_ISL_497155, EPI_ISL_497156, EPI_ISL_497157, EPI_ISL_497158, EPI_ISL_497159, EPI_ISL_497160, EPI_ISL_497161, EPI_ISL_497162, EPI_ISL_497163, EPI_ISL_497164, EPI_ISL_497165, EPI_ISL_497166, EPI_ISL_497167, EPI_ISL_497168, EPI_ISL_497169, EPI_ISL_497170, EPI_ISL_497171, EPI_ISL_497172, EPI_ISL_497173, EPI_ISL_497174, EPI_ISL_497175, EPI_ISL_497176, EPI_ISL_497177, EPI_ISL_497178, EPI_ISL_497179, EPI_ISL_497180, EPI_ISL_497181, EPI_ISL_497182, EPI_ISL_497183, EPI_ISL_497184, EPI_ISL_497185, EPI_ISL_497186, EPI_ISL_497187, EPI_ISL_497188, EPI_ISL_497189, EPI_ISL_497190, EPI_ISL_497191, EPI_ISL_497192, EPI_ISL_497193, EPI_ISL_497194, EPI_ISL_497195, EPI_ISL_497196, EPI_ISL_497197, EPI_ISL_497198, EPI_ISL_497199, EPI_ISL_497200, EPI_ISL_497201, EPI_ISL_497202, EPI_ISL_497203, EPI_ISL_497204, EPI_ISL_497205, EPI_ISL_497206, EPI_ISL_497207, EPI_ISL_497208, EPI_ISL_497209, EPI_ISL_497210, EPI_ISL_497211, EPI_ISL_497212, EPI_ISL_497213, EPI_ISL_497214, EPI_ISL_497215, EPI_ISL_497216, EPI_ISL_497217, EPI_ISL_497218, EPI_ISL_497219, EPI_ISL_497220, EPI_ISL_497221, EPI_ISL_497222, EPI_ISL_497223, EPI_ISL_497224, EPI_ISL_497225, EPI_ISL_497226, EPI_ISL_497227, EPI_ISL_497228, EPI_ISL_497229, EPI_ISL_497230, EPI_ISL_497231, EPI_ISL_497232, EPI_ISL_497233, EPI_ISL_497234, EPI_ISL_497235, EPI_ISL_497236, EPI_ISL_497237, EPI_ISL_497238, EPI_ISL_497239, EPI_ISL_497240, EPI_ISL_497241, EPI_ISL_497242, EPI_ISL_497243, EPI_ISL_497244, EPI_ISL_497245, EPI_ISL_497246, EPI_ISL_497247, EPI_ISL_497248, EPI_ISL_497249, EPI_ISL_497250, EPI_ISL_497251, EPI_ISL_497252, EPI_ISL_497253, EPI_ISL_497254, EPI_ISL_497255, EPI_ISL_497256, EPI_ISL_497257, EPI_ISL_497258, EPI_ISL_497259, EPI_ISL_497260, EPI_ISL_497261, EPI_ISL_497262, EPI_ISL_497263, EPI_ISL_497264, EPI_ISL_497265, EPI_ISL_497266, EPI_ISL_497267, EPI_ISL_497268, EPI_ISL_497269, EPI_ISL_497270, EPI_ISL_497271, EPI_ISL_497272, EPI_ISL_497273, EPI_ISL_497274, EPI_ISL_497275, EPI_ISL_497276, EPI_ISL_497277, EPI_ISL_497278, EPI_ISL_497279, EPI_ISL_497280, EPI_ISL_497281, EPI_ISL_497282, EPI_ISL_497283, EPI_ISL_497284, EPI_ISL_497285, EPI_ISL_497286, EPI_ISL_497287, EPI_ISL_497288, EPI_ISL_497289, EPI_ISL_497290, EPI_ISL_497291, EPI_ISL_497292, EPI_ISL_497293, EPI_ISL_497294, EPI_ISL_497295, EPI_ISL_497296, EPI_ISL_497297, EPI_ISL_497298, EPI_ISL_497299, EPI_ISL_497300, EPI_ISL_497301, EPI_ISL_497302, EPI_ISL_497303, EPI_ISL_497304, EPI_ISL_497305, EPI_ISL_497306, EPI_ISL_497307, EPI_ISL_497308, EPI_ISL_497309, EPI_ISL_497310, EPI_ISL_497311, EPI_ISL_497312, EPI_ISL_497313, EPI_ISL_497314, EPI_ISL_497315, EPI_ISL_497316, EPI_ISL_497317, EPI_ISL_497318, EPI_ISL_497319, EPI_ISL_497320, EPI_ISL_497321, EPI_ISL_497322, EPI_ISL_497323, EPI_ISL_497324, EPI_ISL_497325, EPI_ISL_497326, EPI_ISL_497327, EPI_ISL_497328, EPI_ISL_497329, EPI_ISL_497330, EPI_ISL_497331, EPI_ISL_497332, EPI_ISL_497333, EPI_ISL_497334, EPI_ISL_497335, EPI_ISL_497336, EPI_ISL_497337, EPI_ISL_497338, EPI_ISL_497339, EPI_ISL_497340, EPI_ISL_497341, EPI_ISL_497342, EPI_ISL_497343, EPI_ISL_497344, EPI_ISL_497345, EPI_ISL_497346, EPI_ISL_497347, EPI_ISL_497348, EPI_ISL_497349, EPI_ISL_497350, EPI_ISL_497351, EPI_ISL_497352, EPI_ISL_497353, EPI_ISL_497354, EPI_ISL_497355, EPI_ISL_497356, EPI_ISL_497357, EPI_ISL_497358, EPI_ISL_497359, EPI_ISL_497360, EPI_ISL_497361, EPI_ISL_497362, EPI_ISL_497363, EPI_ISL_497364, EPI_ISL_497365, EPI_ISL_497366, EPI_ISL_497367, EPI_ISL_497368, EPI_ISL_497369, EPI_ISL_497370, EPI_ISL_497371, EPI_ISL_497372, EPI_ISL_497373, EPI_ISL_497374, EPI_ISL_497375, EPI_ISL_497376, EPI_ISL_497377, EPI_ISL_497378, EPI_ISL_497379, EPI_ISL_497380, EPI_ISL_497381, EPI_ISL_497382, EPI_ISL_497383, EPI_ISL_497384, EPI_ISL_497385, EPI_ISL_497386, EPI_ISL_497387, EPI_ISL_497388, EPI_ISL_497389, EPI_ISL_497390, EPI_ISL_497391, EPI_ISL_497392, EPI_ISL_497393, EPI_ISL_497394, EPI_ISL_497395, EPI_ISL_497396, EPI_ISL_497397, EPI_ISL_497398, EPI_ISL_497399, EPI_ISL_497400, EPI_ISL_497401, EPI_ISL_497402, EPI_ISL_497403, EPI_ISL_497404, EPI_ISL_497405, EPI_ISL_497406, EPI_ISL_497407, EPI_ISL_497408, EPI_ISL_497409, EPI_ISL_497410, EPI_ISL_497411, EPI_ISL_497412, EPI_ISL_497413, EPI_ISL_497414, EPI_ISL_497415, EPI_ISL_497416, EPI_ISL_497417, EPI_ISL_497418, EPI_ISL_497419, EPI_ISL_497420, EPI_ISL_497421, EPI_ISL_497422, EPI_ISL_497423, EPI_ISL_497424, EPI_ISL_497425, EPI_ISL_497426, EPI_ISL_497427, EPI_ISL_497428, EPI_ISL_497429, EPI_ISL_497430, EPI_ISL_497431, EPI_ISL_497432, EPI_ISL_497433, EPI_ISL_497434, EPI_ISL_497435, EPI_ISL_497436, EPI_ISL_497437, EPI_ISL_497438, EPI_ISL_497439, EPI_ISL_497440, EPI_ISL_497441, EPI_ISL_497442, EPI_ISL_497443, EPI_ISL_497444, EPI_ISL_497445, EPI_ISL_497446, EPI_ISL_497447, EPI_ISL_497448, EPI_ISL_497449, EPI_ISL_497450, EPI_ISL_497451, EPI_ISL_497452, EPI_ISL_497453, EPI_ISL_497454, EPI_ISL_497455, EPI_ISL_497456, EPI_ISL_497457, EPI_ISL_497458, EPI_ISL_497459, EPI_ISL_497460, EPI_ISL_497461, EPI_ISL_497462, EPI_ISL_497463, EPI_ISL_497464, EPI_ISL_497465, EPI_ISL_497466, EPI_ISL_497467, EPI_ISL_497468, EPI_ISL_497469, EPI_ISL_497470, EPI_ISL_497471, EPI_ISL_497472, EPI_ISL_497473, EPI_ISL_497474, EPI_ISL_497475, EPI_ISL_497476, EPI_ISL_497477, EPI_ISL_497478, EPI_ISL_497479, EPI_ISL_497480, EPI_ISL_497481, EPI_ISL_497482, EPI_ISL_497483, EPI_ISL_497484, EPI_ISL_497485, EPI_ISL_497486, EPI_ISL_497487, EPI_ISL_497488, EPI_ISL_497489, EPI_ISL_497490, EPI_ISL_497491, EPI_ISL_497492, EPI_ISL_497493, EPI_ISL_497494, EPI_ISL_497495, EPI_ISL_497496, EPI_ISL_497497, EPI_ISL_497498, EPI_ISL_497499, EPI_ISL_497500, EPI_ISL_497501, EPI_ISL_497502, EPI_ISL_497503, EPI_ISL_497504, EPI_ISL_497505, EPI_ISL_497506, EPI_ISL_497507, EPI_ISL_497508, EPI_ISL_497509, EPI_ISL_497510, EPI_ISL_497511, EPI_ISL_497512, EPI_ISL_497513, EPI_ISL_497514, EPI_ISL_497515, EPI_ISL_497516, EPI_ISL_497517, EPI_ISL_497518, EPI_ISL_497519, EPI_ISL_497520, EPI_ISL_497521, EPI_ISL_497522, EPI_ISL_497523, EPI_ISL_497524, EPI_ISL_497525, EPI_ISL_497526, EPI_ISL_497527, EPI_ISL_497528, EPI_ISL_497529, EPI_ISL_497530, EPI_ISL_497531, EPI_ISL_497532, EPI_ISL_497533, EPI_ISL_497534, EPI_ISL_497535, EPI_ISL_497536, EPI_ISL_497537, EPI_ISL_497538, EPI_ISL_497539, EPI_ISL_497540, EPI_ISL_497541, EPI_ISL_497542, EPI_ISL_497543, EPI_ISL_497544, EPI_ISL_497545, EPI_ISL_497546, EPI_ISL_497547, EPI_ISL_497548, EPI_ISL_497549, EPI_ISL_497550, EPI_ISL_497551, EPI_ISL_497552, EPI_ISL_497553, EPI_ISL_497554, EPI_ISL_497555, EPI_ISL_497556, EPI_ISL_497557, EPI_ISL_497558, EPI_ISL_497559, EPI_ISL_497560, EPI_ISL_497561, EPI_ISL_497562, EPI_ISL_497563, EPI_ISL_497564, EPI_ISL_497565, EPI_ISL_497566, EPI_ISL_497567, EPI_ISL_497568, EPI_ISL_497569, EPI_ISL_497570, EPI_ISL_497571, EPI_ISL_497572, EPI_ISL_497573, EPI_ISL_497574, EPI_ISL_497575, EPI_ISL_497576, EPI_ISL_497577, EPI_ISL_497578, EPI_ISL_497579, EPI_ISL_497580, EPI_ISL_497581, EPI_ISL_497582, EPI_ISL_497583, EPI_ISL_497584, EPI_ISL_497585, EPI_ISL_497586, EPI_ISL_497587, EPI_ISL_497588, EPI_ISL_497589, EPI_ISL_497590, EPI_ISL_497591, EPI_ISL_497592, EPI_ISL_497593, EPI_ISL_497594, EPI_ISL_497595, EPI_ISL_497596, EPI_ISL_497597, EPI_ISL_497598, EPI_ISL_497599, EPI_ISL_497600, EPI_ISL_497601, EPI_ISL_497602, EPI_ISL_497603, EPI_ISL_497604, EPI_ISL_497605, EPI_ISL_497606, EPI_ISL_497607, EPI_ISL_497608, EPI_ISL_497609, EPI_ISL_497610, EPI_ISL_497611, EPI_ISL_497612, EPI_ISL_497613, EPI_ISL_497614, EPI_ISL_497615, EPI_ISL_497616, EPI_ISL_497617, EPI_ISL_497618, EPI_ISL_497619, EPI_ISL_497620, EPI_ISL_497621, EPI_ISL_497622, EPI_ISL_497623, EPI_ISL_497624, EPI_ISL_497625, EPI_ISL_497626, EPI_ISL_497627, EPI_ISL_497628, EPI_ISL_497629, EPI_ISL_497630, EPI_ISL_497631, EPI_ISL_497632, EPI_ISL_497633, EPI_ISL_497634, EPI_ISL_497635, EPI_ISL_497636, EPI_ISL_497637, EPI_ISL_497638, EPI_ISL_497639, EPI_ISL_497640, EPI_ISL_497641, EPI_ISL_497642, EPI_ISL_497643, EPI_ISL_497644, EPI_ISL_497645, EPI_ISL_497646, EPI_ISL_497647, EPI_ISL_497648, EPI_ISL_497649, EPI_ISL_497650, EPI_ISL_497651, EPI_ISL_497652, EPI_ISL_497653, EPI_ISL_497654, EPI_ISL_497655, EPI_ISL_497656, EPI_ISL_497657, EPI_ISL_497658, EPI_ISL_497659, EPI_ISL_497660, EPI_ISL_497661, EPI_ISL_497662, EPI_ISL_497663, EPI_ISL_497664, EPI_ISL_497665, EPI_ISL_497666, EPI_ISL_497667, EPI_ISL_497668, EPI_ISL_497669, EPI_ISL_497670, EPI_ISL_497671, EPI_ISL_497672, EPI_ISL_497673, EPI_ISL_497674, EPI_ISL_497675, EPI_ISL_497676, EPI_ISL_497677, EPI_ISL_497678, EPI_ISL_497679, EPI_ISL_497680, EPI_ISL_497681, EPI_ISL_497682, EPI_ISL_497683, EPI_ISL_497684, EPI_ISL_497685, EPI_ISL_497686, EPI_ISL_497687, EPI_ISL_497688, EPI_ISL_497689, EPI_ISL_497690, EPI_ISL_497691, EPI_ISL_497692, EPI_ISL_497693, EPI_ISL_497694, EPI_ISL_497695, EPI_ISL_497696, EPI_ISL_497697, EPI_ISL_497698, EPI_ISL_497699, EPI_ISL_497700, EPI_ISL_497701, EPI_ISL_497702, EPI_ISL_497703, EPI_ISL_497704, EPI_ISL_497705, EPI_ISL_497706, EPI_ISL_497707, EPI_ISL_497708, EPI_ISL_497709, EPI_ISL_497710, EPI_ISL_497711, EPI_ISL_497712, EPI_ISL_497713, EPI_ISL_497714, EPI_ISL_497715, EPI_ISL_497716, EPI_ISL_497717, EPI_ISL_497718, EPI_ISL_497719, EPI_ISL_497720, EPI_ISL_497721, EPI_ISL_497722, EPI_ISL_497723, EPI_ISL_497724, EPI_ISL_497725, EPI_ISL_497726 | | | |
| see above | Washington State Department of Health | Seattle Flu Study | Deborah A. Nickerson, Chris D. Frazar, Jover Lee, Benjamin Pelle, Matthew Richardson, Amanda Adler, Elisabeth Brandstetter, Peter D. Han, Kairsten Fay, Misja Ilcisin, Kirsten Lacombe, Thomas R. Sibley, Melissa Truong, Caitlin R. Wolf, Romesh Gautom, Geoff Melly, Brian Hiatt, Philip Dykema, Scott Lindquist, Michael Boeckh, Janet A. Englund, Michael Famulare, Barry R. Lutz, Mark J. Rieder, Lea M. Starita, Matthew Thompson, Helen Y. Chu, Jay Shendure, Trevor Bedford |
| EPI_ISL_497736, EPI_ISL_497738, EPI_ISL_497744, EPI_ISL_497745 | Instituto Nacional de Salud, Bogotá, Colombia | Instituto Nacional de Salud, Bogotá, Colombia | Katherine Laiton-Donato, Diego A. Álvarez-Díaz, Carlos Franco-Muñoz, Jonathan Reales, Diego Andrés Prada, Jose A. Usme-Ciro, Nicolas D. Franco-Sierra, Zulma M. Cucunubá, Christian Julian Villabona-Arenas, Liz Villabona-Arenas, Sussy Echeverria, Astrid C. Flórez, Carolina Ferro, Diana Marcela Walteros-Acero, Franklin Prieto, Carlos Andrés Durán, Martha Lucia Ospina Martinez, Marcela Mercado-Reyes |
| EPI_ISL_497758, EPI_ISL_497760 | CSIR-CDRI/SGPGI, Lucknow | CSIr-CDRI/SGPGI, Lucknow | Saumya Sarkar, Dharam Veer Singh, Rahul Vishvkarma, Ujjala Ghoshal, Uday Ghoshal, Ravishankar Ramachandran, Tapas Kumar Kundu, Rajender Singh |
| EPI_ISL_497762, EPI_ISL_497763 | CSIR-CDRI/SGPGI, Lucknow | CSIR-CDRI/SGPGI, Lucknow | Saumya Sarkar, Dharam Veer Singh, Rahul Vishvkarma, Ujjala Ghoshal, Uday Ghoshal, Ravishankar Ramachandran, Tapas Kumar Kundu, Rajender Singh |
| EPI_ISL_497764 | CSIR-CDRi/SGPGI, Lucknow | CSIR-CDRi/SGPGI, Lucknow | Saumya Sarkar, Dharam Veer Singh, Rahul Vishvkarma, Ujjala Ghoshal, Uday Ghoshal, Ravishankar Ramachandran, Tapas Kumar Kundu, Rajender Singh |
| EPI_ISL_497765 | CSIR-CDRI/SGPGI, Lucknow | CSIR-CDRI/SGPGI, Lucknow | Saumya Sarkar, Dharam Veer Singh, Rahul Vishvkarma, Ujjala Ghoshal, Uday Ghoshal, Ravishankar Ramachandran, Tapas Kumar Kundu, Rajender Singh |
| EPI_ISL_497766 | CSIR-CDRI/SGPGI, Lucknow | CSIR-CDRi/SGPGI, Lucknow | Saumya Sarkar, Dharam Veer Singh, Rahul Vishvkarma, Ujjala Ghoshal, Uday Ghoshal, Ravishankar Ramachandran, Tapas Kumar Kundu, Rajender Singh |
| EPI_ISL_497767 | CSIR-CDRI/SGPGI, Lucknow | CSIR-CDRI/SGPGI, Lucknow | Saumya Sarkar, Dharam Veer Singh, Rahul Vishvkarma, Ujjala Ghoshal, Uday Ghoshal, Ravishankar Ramachandran, Tapas Kumar Kundu, Rajender Singh |
| EPI_ISL_497768, EPI_ISL_497769, EPI_ISL_497770, EPI_ISL_497771, EPI_ISL_497772, EPI_ISL_497773, EPI_ISL_497774, EPI_ISL_497775, EPI_ISL_497776, EPI_ISL_497777, EPI_ISL_497778, EPI_ISL_497779, EPI_ISL_497780, EPI_ISL_497781, EPI_ISL_497782, EPI_ISL_497783, EPI_ISL_497784, EPI_ISL_497785, EPI_ISL_497786, EPI_ISL_497787, EPI_ISL_497788, EPI_ISL_497789, EPI_ISL_497790, EPI_ISL_497791, EPI_ISL_497792, EPI_ISL_497793, EPI_ISL_497794, EPI_ISL_497795, EPI_ISL_497796, EPI_ISL_497797, EPI_ISL_497798, EPI_ISL_497799, EPI_ISL_497800, EPI_ISL_497801, EPI_ISL_497802, EPI_ISL_497803, EPI_ISL_497804, EPI_ISL_497805, EPI_ISL_497806, EPI_ISL_497807, EPI_ISL_497808, EPI_ISL_497809, EPI_ISL_497810, EPI_ISL_497811, EPI_ISL_497812, EPI_ISL_497813, EPI_ISL_497814, EPI_ISL_497815, EPI_ISL_497816, EPI_ISL_497817, EPI_ISL_497818, EPI_ISL_497819, EPI_ISL_497820, EPI_ISL_497821, EPI_ISL_497822, EPI_ISL_497823, EPI_ISL_497824, EPI_ISL_497825, EPI_ISL_497826, EPI_ISL_497827, EPI_ISL_497828, EPI_ISL_497829, EPI_ISL_497830, EPI_ISL_497831, EPI_ISL_497832, EPI_ISL_497833, EPI_ISL_497834, EPI_ISL_497835, EPI_ISL_497836, EPI_ISL_497837, EPI_ISL_497838, EPI_ISL_497839, EPI_ISL_497840, EPI_ISL_497841, EPI_ISL_497842, EPI_ISL_497843, EPI_ISL_497844, EPI_ISL_497845, EPI_ISL_497846, EPI_ISL_497847, EPI_ISL_497848, EPI_ISL_497849, EPI_ISL_497850, EPI_ISL_497851, EPI_ISL_497852, EPI_ISL_497853, EPI_ISL_497854, EPI_ISL_497855, EPI_ISL_497856, EPI_ISL_497857, EPI_ISL_497858, EPI_ISL_497859, EPI_ISL_497860, EPI_ISL_497861, EPI_ISL_497862, EPI_ISL_497863, EPI_ISL_497864, EPI_ISL_497865, EPI_ISL_497866, EPI_ISL_497867, EPI_ISL_497868, EPI_ISL_497869, EPI_ISL_497870 | | | |
| see above | Department of Microbiology, The University of Hong Kong | Department of Microbiology, The University of Hong Kong | Kelvin K.W. To, Kwok-Yung Yuen |
| EPI_ISL_497872 | UW Virology Lab | UW Virology Lab | Pavitra Roychoudhury, Amin Addetia, Hong Xie, Lasata Shrestha, Truong Nguyen, Meei-Li Huang, Keith Jerome, Alexander Greninger |
| EPI_ISL_497873, EPI_ISL_497874, EPI_ISL_497875, EPI_ISL_497876, EPI_ISL_497877, EPI_ISL_497878, EPI_ISL_497879 | National Centre For Cell Science | National Centre For Cell Science | Dhiraj Paul, Kunal Jani, Radha Chauhan, Janesh Kumar, Vasudevan Seshadri, Girdhari Lal, Rajesh Karyakarte, Suvarna Joshi, Murlidhar Tambe, Sourav Sen, Santosh Karade, Kavita Bala Anand, Shelinder Pal Singh Shergill, Rajiv Mohan Gupta, Manoj Kumar Bhat, Arvind Sahu, Maharashtra COVID-19 Study Group, DBT's PAN-INDIA 1000 SARS-CoV2 RNA genome sequencing consortium, Yogesh S Shouche |
| EPI_ISL_497880, EPI_ISL_497881, EPI_ISL_497882, EPI_ISL_497883, EPI_ISL_497884, EPI_ISL_497885, EPI_ISL_497886, EPI_ISL_497887 | Armed Forces Medical College | National Centre For Cell Science | Dhiraj Paul, Kunal Jani, Radha Chauhan, Janesh Kumar, Vasudevan Seshadri, Girdhari Lal, Rajesh Karyakarte, Suvarna Joshi, Murlidhar Tambe, Sourav Sen, Santosh Karade, Kavita Bala Anand, Shelinder Pal Singh Shergill, Rajiv Mohan Gupta, Manoj Kumar Bhat, Arvind Sahu, Maharashtra COVID-19 Study Group, DBT's PAN-INDIA 1000 SARS-CoV2 RNA genome sequencing consortium, Yogesh S Shouche |
| EPI_ISL_497888, EPI_ISL_497889, EPI_ISL_497890, EPI_ISL_497891 | B.J. Govt. Medical College | National Centre For Cell Science | Dhiraj Paul, Kunal Jani, Radha Chauhan, Janesh Kumar, Vasudevan Seshadri, Girdhari Lal, Rajesh Karyakarte, Suvarna Joshi, Murlidhar Tambe, Sourav Sen, Santosh Karade, Kavita Bala Anand, Shelinder Pal Singh Shergill, Rajiv Mohan Gupta, Manoj Kumar Bhat, Arvind Sahu, Maharashtra COVID-19 Study Group, DBT's PAN-INDIA 1000 SARS-CoV2 RNA genome sequencing consortium, Yogesh S Shouche |
| EPI_ISL_497950 | Shaoxing CDC | Zhejiang Provincial Center for Disease Control and Prevention | Yin Chen, Yanjun Zhang, Haiyan Mao, Junhang Pan, Xiuyu Lou, Yi Sun, Hao Yan, Zhen Li, Wen Shi |
| EPI_ISL_497951 | Division of Viral Diseases, Center for Laboratory Control of Infectious Diseases, Korea Centers for Diseases Control and Prevention | Division of Viral Diseases, Center for Laboratory Control of Infectious Diseases, Korea Centers for Diseases Control and Prevention | Jeong-Min Kim, Yoon-Seok Chung, Namjoo Lee, Sang Hee Woo, Hye-Jun Jo, Heui Man Kim, Jun-Sub Kim, Myung Guk Han |
| EPI_ISL_497953, EPI_ISL_497954, EPI_ISL_497955, EPI_ISL_497956, EPI_ISL_497957, EPI_ISL_497958, EPI_ISL_497959, EPI_ISL_497960 | Division of Viral Diseases, Center for Laboratory Control of Infectious Diseases, Korea Centers for Diseases Control and Prevention | Division of Viral Diseases, Center for Laboratory Control of Infectious Diseases, Korea Centers for Diseases Control and Prevention | Jeong-Min Kim, Yoon-Seok Chung, Namjoo Lee, Sang Hee Woo, Hye-Jun Jo, Heui Man Kim, Jun-Sub Kim, Dong Hyun Song, Daesang Lee, Seong Tae Jeong, Myung Guk Han |
| EPI_ISL_497961, EPI_ISL_497962, EPI_ISL_497963, EPI_ISL_497964, EPI_ISL_497965, EPI_ISL_497966, EPI_ISL_497967 | Division of Viral Diseases, Center for Laboratory Control of Infectious Diseases, Korea Centers for Diseases Control and Prevention | Division of Viral Diseases, Center for Laboratory Control of Infectious Diseases, Korea Centers for Diseases Control and Prevention | Jeong-Min Kim, Yoon-Seok Chung, Namjoo Lee, Sang Hee Woo, Hye-Jun Jo, Heui Man Kim, Jun-Sub Kim, Myung Guk Han |
| EPI_ISL_497968, EPI_ISL_497969, EPI_ISL_497970, EPI_ISL_497971 | Division of Viral Diseases, Center for Laboratory Control of Infectious Diseases, Korea Centers for Diseases Control and Prevention | Division of Viral Diseases, Center for Laboratory Control of Infectious Diseases, Korea Centers for Diseases Control and Prevention | Jeong-Min Kim, Yoon-Seok Chung, Namjoo Lee, Sang Hee Woo, Hye-Jun Jo, Heui Man Kim, Jun-Sub Kim, Dong Hyun Song, Daesang Lee, Seong Tae Jeong, Myung Guk Han |
| EPI_ISL_497972, EPI_ISL_497973 | Division of Viral Diseases, Center for Laboratory Control of Infectious Diseases, Korea Centers for Diseases Control and Prevention | Division of Viral Diseases, Center for Laboratory Control of Infectious Diseases, Korea Centers for Diseases Control and Prevention | Jeong-Min Kim, Yoon-Seok Chung, Namjoo Lee, Sang Hee Woo, Hye-Jun Jo, Heui Man Kim, Jun-Sub Kim, Myung Guk Han |
| EPI_ISL_497974 | Division of Viral Diseases, Center for Laboratory Control of Infectious Diseases, Korea Centers for Diseases Control and Prevention | Division of Viral Diseases, Center for Laboratory Control of Infectious Diseases, Korea Centers for Diseases Control and Prevention | Jeong-Min Kim, Yoon-Seok Chung, Namjoo Lee, Sang Hee Woo, Hye-Jun Jo, Heui Man Kim, Jun-Sub Kim, Dong Hyun Song, Daesang Lee, Seong Tae Jeong, Myung Guk Han |
| EPI_ISL_497975, EPI_ISL_497976, EPI_ISL_497977, EPI_ISL_497978, EPI_ISL_497979, EPI_ISL_497980, EPI_ISL_497981, EPI_ISL_497982, EPI_ISL_497983, EPI_ISL_497984 | Division of Viral Diseases, Center for Laboratory Control of Infectious Diseases, Korea Centers for Diseases Control and Prevention | Division of Viral Diseases, Center for Laboratory Control of Infectious Diseases, Korea Centers for Diseases Control and Prevention | Jeong-Min Kim, Yoon-Seok Chung, Namjoo Lee, Sang Hee Woo, Hye-Jun Jo, Heui Man Kim, Jun-Sub Kim, Myung Guk Han |
| EPI_ISL_497985 | Division of Viral Diseases, Center for Laboratory Control of Infectious Diseases, Korea Centers for Diseases Control and Prevention | Division of Viral Diseases, Center for Laboratory Control of Infectious Diseases, Korea Centers for Diseases Control and Prevention | Jeong-Min Kim, Yoon-Seok Chung, Namjoo Lee, Sang Hee Woo, Hye-Jun Jo, Heui Man Kim, Jun-Sub Kim, Dong Hyun Song, Daesang Lee, Seong Tae Jeong, Myung Guk Han |
| EPI_ISL_497986 | Division of Viral Diseases, Center for Laboratory Control of Infectious Diseases, Korea Centers for Diseases Control and Prevention | Division of Viral Diseases, Center for Laboratory Control of Infectious Diseases, Korea Centers for Diseases Control and Prevention | Jeong-Min Kim, Yoon-Seok Chung, Namjoo Lee, Sang Hee Woo, Hye-Jun Jo, Heui Man Kim, Jun-Sub Kim, Myung Guk Han |
| EPI_ISL_497987, EPI_ISL_497988, EPI_ISL_497989, EPI_ISL_497990 | Division of Viral Diseases, Center for Laboratory Control of Infectious Diseases, Korea Centers for Diseases Control and Prevention | Division of Viral Diseases, Center for Laboratory Control of Infectious Diseases, Korea Centers for Diseases Control and Prevention | Jeong-Min Kim, Yoon-Seok Chung, Namjoo Lee, Sang Hee Woo, Hye-Jun Jo, Heui Man Kim, Jun-Sub Kim, Dong Hyun Song, Daesang Lee, Seong Tae Jeong, Myung Guk Han |
| EPI_ISL_497991, EPI_ISL_497992, EPI_ISL_497993 | Division of Viral Diseases, Center for Laboratory Control of Infectious Diseases, Korea Centers for Diseases Control and Prevention | Division of Viral Diseases, Center for Laboratory Control of Infectious Diseases, Korea Centers for Diseases Control and Prevention | Jeong-Min Kim, Yoon-Seok Chung, Namjoo Lee, Sang Hee Woo, Hye-Jun Jo, Heui Man Kim, Jun-Sub Kim, Myung Guk Han |
| EPI_ISL_497994, EPI_ISL_497995 | Division of Viral Diseases, Center for Laboratory Control of Infectious Diseases, Korea Centers for Diseases Control and Prevention | Division of Viral Diseases, Center for Laboratory Control of Infectious Diseases, Korea Centers for Diseases Control and Prevention | Jeong-Min Kim, Yoon-Seok Chung, Namjoo Lee, Sang Hee Woo, Hye-Jun Jo, Heui Man Kim, Jun-Sub Kim, Dong Hyun Song, Daesang Lee, Seong Tae Jeong, Myung Guk Han |
| EPI_ISL_497996 | Division of Viral Diseases, Center for Laboratory Control of | Division of Viral Diseases, Center for Laboratory Control of Infectious | Jeong-Min Kim, Yoon-Seok Chung, Namjoo Lee, Sang Hee Woo, Hye-Jun Jo, Heui Man Kim, Jun-Sub Kim, Myung Guk Han |

| | | | |
|---|---|---|---|
| | Infectious Diseases, Korea Centers for Diseases Control and Prevention | Diseases, Korea Centers for Diseases Control and Prevention | |
| EPI_ISL_497997 | Division of Viral Diseases, Center for Laboratory Control of Infectious Diseases, Korea Centers for Diseases Control and Prevention | Division of Viral Diseases, Center for Laboratory Control of Infectious Diseases, Korea Centers for Diseases Control and Prevention | Jeong-Min Kim, Yoon-Seok Chung, Namjoo Lee, Sang Hee Woo, Hye-Jun Jo, Heui Man Kim, Jun-Sub Kim, Dong Hyun Song, Daesang Lee, Seong Tae Jeong, Myung Guk Han |
| EPI_ISL_497998, EPI_ISL_497999, EPI_ISL_498000, EPI_ISL_498001, EPI_ISL_498002, EPI_ISL_498003 | Division of Viral Diseases, Center for Laboratory Control of Infectious Diseases, Korea Centers for Diseases Control and Prevention | Division of Viral Diseases, Center for Laboratory Control of Infectious Diseases, Korea Centers for Diseases Control and Prevention | Jeong-Min Kim, Yoon-Seok Chung, Namjoo Lee, Sang Hee Woo, Hye-Jun Jo, Heui Man Kim, Jun-Sub Kim, Myung Guk Han |
| EPI_ISL_498004 | Division of Viral Diseases, Center for Laboratory Control of Infectious Diseases, Korea Centers for Diseases Control and Prevention | Division of Viral Diseases, Center for Laboratory Control of Infectious Diseases, Korea Centers for Diseases Control and Prevention | Jeong-Min Kim, Yoon-Seok Chung, Namjoo Lee, Sang Hee Woo, Hye-Jun Jo, Heui Man Kim, Jun-Sub Kim, Dong Hyun Song, Daesang Lee, Seong Tae Jeong, Myung Guk Han |
| EPI_ISL_498005 | Division of Viral Diseases, Center for Laboratory Control of Infectious Diseases, Korea Centers for Diseases Control and Prevention | Division of Viral Diseases, Center for Laboratory Control of Infectious Diseases, Korea Centers for Diseases Control and Prevention | Jeong-Min Kim, Yoon-Seok Chung, Namjoo Lee, Sang Hee Woo, Hye-Jun Jo, Heui Man Kim, Jun-Sub Kim, Myung Guk Han |
| EPI_ISL_498006, EPI_ISL_498007, EPI_ISL_498008, EPI_ISL_498009, EPI_ISL_498010, EPI_ISL_498011, EPI_ISL_498012, EPI_ISL_498013, EPI_ISL_498014, EPI_ISL_498015, EPI_ISL_498016 | | | |
| see above | Division of Viral Diseases, Center for Laboratory Control of Infectious Diseases, Korea Centers for Diseases Control and Prevention | Division of Viral Diseases, Center for Laboratory Control of Infectious Diseases, Korea Centers for Diseases Control and Prevention | Jeong-Min Kim, Yoon-Seok Chung, Namjoo Lee, Sang Hee Woo, Hye-Jun Jo, Heui Man Kim, Jun-Sub Kim, Dong Hyun Song, Daesang Lee, Seong Tae Jeong, Myung Guk Han |
| EPI_ISL_498017 | Division of Viral Diseases, Center for Laboratory Control of Infectious Diseases, Korea Centers for Diseases Control and Prevention | Division of Viral Diseases, Center for Laboratory Control of Infectious Diseases, Korea Centers for Diseases Control and Prevention | Jeong-Min Kim, Yoon-Seok Chung, Namjoo Lee, Sang Hee Woo, Hye-Jun Jo, Heui Man Kim, Jun-Sub Kim, Myung Guk Han |
| EPI_ISL_498018, EPI_ISL_498019, EPI_ISL_498020, EPI_ISL_498021, EPI_ISL_498022, EPI_ISL_498023, EPI_ISL_498024, EPI_ISL_498025, EPI_ISL_498026, EPI_ISL_498027, EPI_ISL_498028, EPI_ISL_498029, EPI_ISL_498030, EPI_ISL_498031, EPI_ISL_498032, EPI_ISL_498033, EPI_ISL_498034, EPI_ISL_498035 | | | |
| see above | Division of Viral Diseases, Center for Laboratory Control of Infectious Diseases, Korea Centers for Diseases Control and Prevention | Division of Viral Diseases, Center for Laboratory Control of Infectious Diseases, Korea Centers for Diseases Control and Prevention | Jeong-Min Kim, Yoon-Seok Chung, Namjoo Lee, Sang Hee Woo, Hye-Jun Jo, Heui Man Kim, Jun-Sub Kim, Dong Hyun Song, Daesang Lee, Seong Tae Jeong, Myung Guk Han |
| EPI_ISL_498036 | Division of Viral Diseases, Center for Laboratory Control of Infectious Diseases, Korea Centers for Diseases Control and Prevention | Division of Viral Diseases, Center for Laboratory Control of Infectious Diseases, Korea Centers for Diseases Control and Prevention | Jeong-Min Kim, Yoon-Seok Chung, Namjoo Lee, Sang Hee Woo, Hye-Jun Jo, Heui Man Kim, Jun-Sub Kim, Myung Guk Han |
| EPI_ISL_498037, EPI_ISL_498038, EPI_ISL_498039, EPI_ISL_498040, EPI_ISL_498041, EPI_ISL_498042, EPI_ISL_498043, EPI_ISL_498044, EPI_ISL_498045, EPI_ISL_498046, EPI_ISL_498047, EPI_ISL_498048, EPI_ISL_498049, EPI_ISL_498050, EPI_ISL_498051, EPI_ISL_498052, EPI_ISL_498053 | | | |
| see above | Division of Viral Diseases, Center for Laboratory Control of Infectious Diseases, Korea Centers for Diseases Control and Prevention | Division of Viral Diseases, Center for Laboratory Control of Infectious Diseases, Korea Centers for Diseases Control and Prevention | Jeong-Min Kim, Yoon-Seok Chung, Namjoo Lee, Sang Hee Woo, Hye-Jun Jo, Heui Man Kim, Jun-Sub Kim, Dong Hyun Song, Daesang Lee, Seong Tae Jeong, Myung Guk Han |
| EPI_ISL_498054, EPI_ISL_498055, EPI_ISL_498056, EPI_ISL_498057, EPI_ISL_498058, EPI_ISL_498059, EPI_ISL_498060, EPI_ISL_498061, EPI_ISL_498062, EPI_ISL_498063, EPI_ISL_498064, EPI_ISL_498065, EPI_ISL_498066, EPI_ISL_498067, EPI_ISL_498068, EPI_ISL_498069, EPI_ISL_498070, EPI_ISL_498071, EPI_ISL_498072, EPI_ISL_498073, EPI_ISL_498074, EPI_ISL_498075, EPI_ISL_498076, EPI_ISL_498077, EPI_ISL_498078, EPI_ISL_498079, EPI_ISL_498080, EPI_ISL_498081, EPI_ISL_498082, EPI_ISL_498083, EPI_ISL_498084, EPI_ISL_498085, EPI_ISL_498086, EPI_ISL_498087, EPI_ISL_498088, EPI_ISL_498089, EPI_ISL_498090, EPI_ISL_498091, EPI_ISL_498092, EPI_ISL_498093, EPI_ISL_498094, EPI_ISL_498095, EPI_ISL_498096, EPI_ISL_498097, EPI_ISL_498098, EPI_ISL_498099, EPI_ISL_498100, EPI_ISL_498101, EPI_ISL_498102, EPI_ISL_498103, EPI_ISL_498104, EPI_ISL_498105, EPI_ISL_498106, EPI_ISL_498107, EPI_ISL_498108, EPI_ISL_498109, EPI_ISL_498110, EPI_ISL_498111, EPI_ISL_498112, EPI_ISL_498113, EPI_ISL_498114, EPI_ISL_498115, EPI_ISL_498116, EPI_ISL_498117, EPI_ISL_498118, EPI_ISL_498119, EPI_ISL_498120, EPI_ISL_498121, EPI_ISL_498122, EPI_ISL_498123, EPI_ISL_498124, EPI_ISL_498125, EPI_ISL_498126 | | | |
| see above | NHLS-IALCH | KRISP, KZN Research Innovation and Sequencing Platform | Giandhari J, Pillay S, Lessells R, Chimukangara B, Mdlalose K, York D, Khan S, Tegally H, Wilkinson E, de Oliveira T |
| EPI_ISL_498127, EPI_ISL_498128, EPI_ISL_498129, EPI_ISL_498130, EPI_ISL_498131, EPI_ISL_498132, EPI_ISL_498133, EPI_ISL_498134, EPI_ISL_498135, EPI_ISL_498136, EPI_ISL_498137, EPI_ISL_498138, EPI_ISL_498139, EPI_ISL_498140, EPI_ISL_498141, EPI_ISL_498142, EPI_ISL_498143, EPI_ISL_498144, EPI_ISL_498145, EPI_ISL_498146, EPI_ISL_498147, EPI_ISL_498148, EPI_ISL_498149, EPI_ISL_498150, EPI_ISL_498151 | | | |
| see above | Department of Clinical Microbiology | GIGA Medical Genomics | Keith Durkin, Maria Artesi, Sébastien Bontems, Raphaël Boreux, Cécile Meex, Axelle Chaslain, Céline Fombellida-Lopez, Pierrette Melin, Marie-Pierre Hayette, Vincent Bours. |
| EPI_ISL_498152, EPI_ISL_498153, EPI_ISL_498154, EPI_ISL_498155, EPI_ISL_498156, EPI_ISL_498157, EPI_ISL_498158, EPI_ISL_498159, EPI_ISL_498160, EPI_ISL_498161, EPI_ISL_498162, EPI_ISL_498163, EPI_ISL_498164, EPI_ISL_498165, EPI_ISL_498166, EPI_ISL_498167, EPI_ISL_498168, EPI_ISL_498169, EPI_ISL_498170 | | | |
| see above | Instituto Nacional de Salud, Bogotá, Colombia | Instituto Nacional de Salud, Bogotá, Colombia | Katherine Laiton-Donato, Diego A. Álvarez-Díaz, Carlos Franco-Muñoz, Jonathan Reales, Diego Andrés Prada, Jose A. Usme-Ciro, Nicolas D. Franco-Sierra, Zulma M. Cucunubá, Christian Julian Villabona-Arenas, Liz Villabona-Arenas, Sussy Echeverria, Astrid C. Flórez, Carolina Ferro, Diana Marcela Walteros-Acero, Franklin Prieto, Carlos Andrés Durán, Martha Lucia Ospina Martinez, Marcela Mercado-Reyes |
| EPI_ISL_498171, EPI_ISL_498172, EPI_ISL_498173, EPI_ISL_498174, EPI_ISL_498175, EPI_ISL_498176, EPI_ISL_498177, EPI_ISL_498178, EPI_ISL_498179, EPI_ISL_498180, EPI_ISL_498181, EPI_ISL_498182, EPI_ISL_498183, EPI_ISL_498184, EPI_ISL_498185, EPI_ISL_498186, EPI_ISL_498187, EPI_ISL_498188, EPI_ISL_498189, EPI_ISL_498190, EPI_ISL_498191, EPI_ISL_498192 | | | |
| see above | OUCRU | OUCRU | Nguyen Van Vinh Chau, Nguyen Thi Thu Hong, Nguyen Thi Han Ny, Le Nguyen Truc Nhu, Nghiem My Ngoc, Vo Thanh Lam, Nguyen Thanh Dung, Lam Minh Yen, Ngo Ngoc Quang Minh, Le Manh Hung, Nguyen Tri Dung, Dinh Nguyen Huy Man, Lam Anh Nguyet, Tran Chanh Xuan, Tran Tinh Hien, Nguyen Thanh Phong, Tran Nguyen Hoang Tu, Tran Tan Thanh, Nguyen Thanh Truong, Nguyen Tan Binh, Tang Chi Thuong, Guy Thwaites, and Le Van Tan, for OUCRU COVID-19 research group* |
| EPI_ISL_498193 | Molecular Pathology Division, Department of Pathology, Hong Kong Sanatorium & Hospital | Molecular Pathology Division, Department of Pathology, Hong Kong Sanatorium & Hospital | Chun Hang AU, Wai Sing CHAN, Ho Yin LAM, Dona N. HO, Simon Y.M. LAM, Jonpaul S.T. ZEE, Tsun Leung CHAN, Edmond S.K. MA |
| EPI_ISL_498226 | LIC | LIC | LIC |
| EPI_ISL_498227, EPI_ISL_498228 | National Institute of Laboratory Medicine and Referral Center | Genomic Research Lab, BCSIR | Shahina Akter, Abu Sayeed Mohammad Mahmud, Mohammad Samir Uzzaman, Eshrar Osman, Md. Ahasan Habib, Tanjina Akhter Banu, Md. Murshed Hasan Sarkar, Barna Goswami, Iffat Jahan, Md. Saddam Hossain, Tasnim Nafisa, Md. Maruf Ahmed Molla, Mahmuda Yeasmin, Asish Kumar Ghosh, A. K. M. Shamsuzzaman, Sheikh Md. Selim Al Din, Utpal Chandra Ray, Salek Ahmed Sajib, Md. Salim Khan |
| EPI_ISL_498229, EPI_ISL_498230, EPI_ISL_498231, EPI_ISL_498232, EPI_ISL_498233, EPI_ISL_498234, EPI_ISL_498235, EPI_ISL_498236, EPI_ISL_498237, EPI_ISL_498238, EPI_ISL_498239, EPI_ISL_498240, EPI_ISL_498241, EPI_ISL_498242, EPI_ISL_498243, EPI_ISL_498244, EPI_ISL_498245, EPI_ISL_498246, EPI_ISL_498247, EPI_ISL_498248, EPI_ISL_498249, EPI_ISL_498250, EPI_ISL_498251, EPI_ISL_498252 | | | |
| see above | Institut Pasteur de Dakar | Institut Pasteur de Dakar | Ndongo Dia, Moussa Moise Diagne, Mamadou Diop, Marie Henriette Dior Ndione, Mamadou Malado Jallow, Safietou Sankhe Mbengue, Ousmane Faye, Amadou Alpha Sall. |
| EPI_ISL_498253, EPI_ISL_498254 | National Institute of Laboratory Medicine and Referral Center | Genomic Research Lab, BCSIR | Tanjina Akhter Banu, Abu Sayeed Mohammad Mahmud, Mohammad Samir Uzzaman, Eshrar Osman, Md. Ahasan Habib, Shahina Akter, Md. Murshed Hasan Sarkar, Barna Goswami, Iffat Jahan, Md. Saddam Hossain, Tasnim Nafisa, Md. Maruf Ahmed Molla, Mahmuda Yeasmin, Asish Kumar Ghosh, A. K. M. Shamsuzzaman, Sheikh Md. Selim Al Din, Utpal Chandra Ray, Salek Ahmed Sajib, Md. Salim Khan |
| EPI_ISL_498255, EPI_ISL_498256, EPI_ISL_498257, EPI_ISL_498258, EPI_ISL_498259, EPI_ISL_498260, EPI_ISL_498261 | Hospital for Tropical Diseases | COVID-19 Network Investigations (CONI) Alliance | Elizabeth Batty, Nantarat Chantawat, Wasun Chantratita, Thanat Chookajorn, Stefan Fernandez, Angkana Huang, Weena Janwitthayanan, Akanitt Jittmittraphap, Anthony R. Jones, Khajohn Joonsalak, Chonticha Klungtong, Theerarat Kochakarn, Namfon Kotanan, Krittikorn Kumpornsin, Pornsawan Leaungwutiwong, Wudtichai Manasatienkij, Bhakbhoom Panthan, Ekawat Pasomsub, Kingkan Rakmanee, Insee Sensorn, Janjira Thaipadungpanit, Arporn Wangwiwatsin,Treewat Watthanachockchai |
| EPI_ISL_498262, EPI_ISL_498263, EPI_ISL_498264, EPI_ISL_498265, EPI_ISL_498266 | Ramathibodi Hospital | COVID-19 Network Investigations (CONI) Alliance | Elizabeth Batty, Wasun Chantratita, Thanat Chookajorn, Stefan Fernandez, Angkana Huang, Anthony R. Jones, Khajohn Joonsalak, Chonticha Klungtong, Theerarat Kochakarn, Namfon Kotanan, Krittikorn Kumpornsin, Wudtichai Manasatienkij, Bhakbhoom Panthan, Ekawat Pasomsub, Kingkan Rakmanee, Insee Sensorn, Janjira Thaipadungpanit, Arporn Wangwiwatsin,Treewat Watthanachockchai |
| EPI_ISL_498267, EPI_ISL_498268 | National Institute of Laboratory Medicine and Referral Center | Genomic Research Lab, BCSIR | Barna Goswami, Abu Sayeed Mohammad Mahmud, Mohammad Samir Uzzaman, Eshrar Osman, Md. Ahasan Habib, Shahina Akter, Tanjina Akhter Banu, Md. Murshed Hasan Sarkar, Iffat Jahan, Md. Saddam Hossain, Tasnim Nafisa, Md. Maruf Ahmed Molla, Mahmuda Yeasmin, Asish Kumar Ghosh, A. K. M. Shamsuzzaman, Sheikh Md. Selim Al Din, Utpal Chandra Ray, Salek Ahmed Sajib, Md. Salim Khan |
| EPI_ISL_498269, EPI_ISL_498270, EPI_ISL_498271 | Department of Microbiology, The University of Hong Kong | Department of Microbiology, The University of Hong Kong | Kelvin K.W. To, Kwok-Yung Yuen |
| EPI_ISL_498272, EPI_ISL_498273 | National Institute of Laboratory Medicine and Referral Center | Genomic Research Lab, BCSIR | Iffat Jahan, Abu Sayeed Mohammad Mahmud, Mohammad Samir Uzzaman, Eshrar Osman, Md. Ahasan Habib, Shahina Akter, Tanjina Akhter Banu, Md. Murshed Hasan Sarkar, Barna Goswami, Md. Saddam Hossain, Tasnim Nafisa, Md. Maruf Ahmed Molla, Mahmuda Yeasmin, Asish Kumar Ghosh, A. K. M. Shamsuzzaman, Sheikh Md. Selim Al Din, Utpal Chandra Ray, Salek Ahmed Sajib, Md. Salim Khan |
| EPI_ISL_498274, EPI_ISL_498417 | National Institute of Laboratory Medicine and Referral Center | Genomic Research Lab, BCSIR | Tasnim Nafisa, Abu Sayeed Mohammad Mahmud, Mohammad Samir Uzzaman, Eshrar Osman, Md. Ahasan Habib, Shahina Akter, Tanjina Akhter Banu, Md. Murshed Hasan Sarkar, Barna Goswami, Iffat Jahan, Md. Saddam Hossain, Md. Maruf Ahmed Molla, Mahmuda Yeasmin, Asish Kumar Ghosh, A. K. M. Shamsuzzaman, Sheikh Md. Selim Al Din, Utpal Chandra Ray, Salek Ahmed Sajib, Md. Salim Khan |
| EPI_ISL_498418, EPI_ISL_498419 | National Institute of Laboratory Medicine and Referral Center | Genomic Research Lab, BCSIR | Md. Maruf Ahmed Molla, Abu Sayeed Mohammad Mahmud, Mohammad Samir Uzzaman, Eshrar Osman, Md. Ahasan Habib, Shahina Akter, Tanjina Akhter Banu, Md. Murshed Hasan Sarkar, Barna Goswami, Iffat Jahan, Md. Saddam Hossain, Tasnim Nafisa, Mahmuda Yeasmin, Asish Kumar Ghosh, A. K. M. Shamsuzzaman, Sheikh Md. Selim Al Din, Utpal Chandra Ray, Salek Ahmed Sajib, Md. Salim Khan |
| EPI_ISL_498466, EPI_ISL_498467 | National Institute of Laboratory Medicine and Referral Center | Genomic Research Lab, BCSIR | Mahmuda Yeasmin, Abu Sayeed Mohammad Mahmud, Mohammad Samir Uzzaman, Eshrar Osman, Md. Ahasan Habib, Shahina Akter, Tanjina Akhter Banu, Md. Murshed Hasan Sarkar, Barna Goswami, Iffat Jahan, Md. Saddam Hossain, Tasnim Nafisa, Md. Maruf Ahmed Molla, Asish Kumar Ghosh, A. K. M. Shamsuzzaman, Sheikh Md. Selim Al Din, Utpal Chandra Ray, Salek Ahmed Sajib, Md. Salim Khan |
| EPI_ISL_498468, EPI_ISL_498469, EPI_ISL_498470, EPI_ISL_498471, EPI_ISL_498472, EPI_ISL_498473, EPI_ISL_498474, EPI_ISL_498475, EPI_ISL_498476, EPI_ISL_498477, EPI_ISL_498478, EPI_ISL_498479, EPI_ISL_498480, EPI_ISL_498481, EPI_ISL_498482, EPI_ISL_498483, EPI_ISL_498484, EPI_ISL_498485, EPI_ISL_498486, EPI_ISL_498487, EPI_ISL_498488, EPI_ISL_498489, EPI_ISL_498490, EPI_ISL_498491, EPI_ISL_498492, EPI_ISL_498493, EPI_ISL_498494, EPI_ISL_498495, EPI_ISL_498496, EPI_ISL_498497, EPI_ISL_498498, EPI_ISL_498499, EPI_ISL_498500, EPI_ISL_498501, EPI_ISL_498502, EPI_ISL_498503, EPI_ISL_498504, EPI_ISL_498505, EPI_ISL_498506, EPI_ISL_498507, EPI_ISL_498508, EPI_ISL_498509, EPI_ISL_498510, EPI_ISL_498511, EPI_ISL_498512, EPI_ISL_498513, EPI_ISL_498514, EPI_ISL_498515, EPI_ISL_498516, EPI_ISL_498517, EPI_ISL_498518, EPI_ISL_498519, EPI_ISL_498520, EPI_ISL_498521, EPI_ISL_498522, EPI_ISL_498523, EPI_ISL_498524, EPI_ISL_498525, EPI_ISL_498526, EPI_ISL_498527, EPI_ISL_498528, EPI_ISL_498529, EPI_ISL_498530, EPI_ISL_498531, EPI_ISL_498532, EPI_ISL_498533, EPI_ISL_498534, EPI_ISL_498535, EPI_ISL_498536, EPI_ISL_498537, EPI_ISL_498538, EPI_ISL_498539, EPI_ISL_498540, EPI_ISL_498541, EPI_ISL_498542, EPI_ISL_498543, EPI_ISL_498544, EPI_ISL_498545, EPI_ISL_498546, EPI_ISL_498547, EPI_ISL_498548 | | | |
| see above | ACT Pathology | Schwessinger Lab | Ashley Jones, Benjamin Schwessinger, Robert Lanfear, Robyn N Hall, Megan McDonald, Ming-Dao Chia, Kevin Murray, Craig Kennedy, Karina Kennedy |
| EPI_ISL_498549, EPI_ISL_498550 | National Institute of Laboratory Medicine and Referral Center | Genomic Research Lab, BCSIR | Asish Kumar Ghosh, Abu Sayeed Mohammad Mahmud, Mohammad Samir Uzzaman, Eshrar Osman, Md. Ahasan Habib, Shahina Akter, Tanjina Akhter Banu, Md. Murshed Hasan Sarkar, Barna Goswami, Iffat Jahan, Md. Saddam Hossain, Tasnim Nafisa, Md. Maruf Ahmed Molla, Mahmuda Yeasmin, A. K. M. Shamsuzzaman, Sheikh Md. Selim Al Din, Utpal Chandra Ray, Salek Ahmed Sajib, Md. Salim Khan |
| EPI_ISL_498551, EPI_ISL_498552, EPI_ISL_498554, EPI_ISL_498556 | Lebanese American University | Lebanese American University | Abi Habib,W., Abdallah,J., El Shesheny,R., Mokhbat,J., Webby,R.J., Goldstein,J. and Kayali,G. |
| EPI_ISL_498558, EPI_ISL_498559, EPI_ISL_498560, EPI_ISL_498561 | Laboratory of Molecular Virology International Center for Genetic Engineering and Biotechnology (ICGEB) | ARGO Open Lab Platform for Genome Sequencing | Licastro D, Rajasekharan S, Dal Monego S, Segat L, D'Agaro P, Marcello A |
| EPI_ISL_498562 | Laboratory of Molecular Virology International Center for Genetic Engineering and Biotechnology (ICGEB) | ARGO Open Lab Platform for Genome Sequencing | Licastro D, Rajasekharan S, Dal Monego S, Segat L, D'Agaro P, Marcello A |
| EPI_ISL_498563 | Laboratory of Molecular Virology International Center for Genetic Engineering and Biotechnology (ICGEB) | ARGO Open Lab Platform for Genome Sequencing | Licastro D, Rajasekharan S, Dal Monego S, Segat L, D'Agaro P, Marcello A |
| EPI_ISL_498564, EPI_ISL_498565, EPI_ISL_498566, EPI_ISL_498567, EPI_ISL_498568, EPI_ISL_498569, EPI_ISL_498570, EPI_ISL_498571, EPI_ISL_498572, EPI_ISL_498573, EPI_ISL_498574, EPI_ISL_498575, EPI_ISL_498576, EPI_ISL_498577, EPI_ISL_498578, EPI_ISL_498579, EPI_ISL_498580, EPI_ISL_498581, EPI_ISL_498582, EPI_ISL_498583, EPI_ISL_498584, EPI_ISL_498585, EPI_ISL_498586, EPI_ISL_498587, EPI_ISL_498588, EPI_ISL_498589, EPI_ISL_498590, EPI_ISL_498591, EPI_ISL_498592, EPI_ISL_498593, EPI_ISL_498594, EPI_ISL_498595, EPI_ISL_498596, EPI_ISL_498597, EPI_ISL_498598, EPI_ISL_498599, EPI_ISL_498600, EPI_ISL_498601, EPI_ISL_498602, EPI_ISL_498603, EPI_ISL_498604, EPI_ISL_498605, EPI_ISL_498606, EPI_ISL_498607, EPI_ISL_498608, EPI_ISL_498609, EPI_ISL_498610, EPI_ISL_498611, EPI_ISL_498612, EPI_ISL_498613, EPI_ISL_498614, EPI_ISL_498615, EPI_ISL_498616, EPI_ISL_498617, EPI_ISL_498618 | | | |

| | | | |
|---|---|---|---|
| see above | National Public Health Laboratory, National Centre for Infectious Diseases | National Public Health Laboratory, National Centre for Infectious Diseases | Mak TM, Octavia S, Zhou Z, Chavatte JM, Cui L, Lin RTP |
| EPI_ISL_498619, EPI_ISL_498620, EPI_ISL_498621, EPI_ISL_498622, EPI_ISL_498623, EPI_ISL_498624, EPI_ISL_498625, EPI_ISL_498626, EPI_ISL_498627 | Viollier AG | Department of Biosystems Science and Engineering, ETH Zürich | Christian Beisel, Sarah Nadeau, Ivan Topolsky, Pedro Ferreira, Philipp Jablonski, Susana Posada-Céspedes, Tobias Schär, Ina Nissen, Natascha Santacroce, Elodie Burcklen, Christiane Beckmann, Maurice Redondo, Olivier Kobel, Christoph Noppen, Sophie Seidel, Noemie Santamaria de Souza, Niko Beerenwinkel, Tanja Stadler |
| EPI_ISL_498628, EPI_ISL_498629 | Department of Clinical Microbiology | GIGA Medical Genomics | Keith Durkin, Maria Artesi, Sébastien Bontems, Raphaël Boreux, Cécile Meex, Axelle Chaslain, Céline Fombellida-Lopez, Pierrette Melin, Marie-Pierre Hayette, Vincent Bours. |
| EPI_ISL_498630, EPI_ISL_498631, EPI_ISL_498632, EPI_ISL_498633, EPI_ISL_498634, EPI_ISL_498635, EPI_ISL_498637, EPI_ISL_498638, EPI_ISL_498639, EPI_ISL_498640, EPI_ISL_498641, EPI_ISL_498642, EPI_ISL_498643, EPI_ISL_498644, EPI_ISL_498645, EPI_ISL_498646, EPI_ISL_498647, EPI_ISL_498648, EPI_ISL_498649, EPI_ISL_498650, EPI_ISL_498651, EPI_ISL_498652, EPI_ISL_498653, EPI_ISL_498654, EPI_ISL_498655, EPI_ISL_498656, EPI_ISL_498657, EPI_ISL_498658, EPI_ISL_498659, EPI_ISL_498660, EPI_ISL_498661, EPI_ISL_498662, EPI_ISL_498663, EPI_ISL_498664, EPI_ISL_498665, EPI_ISL_498666, EPI_ISL_498667, EPI_ISL_498668, EPI_ISL_498669, EPI_ISL_498670, EPI_ISL_498671, EPI_ISL_498672, EPI_ISL_498673, EPI_ISL_498674, EPI_ISL_498675, EPI_ISL_498676, EPI_ISL_498677, EPI_ISL_498678, EPI_ISL_498679, EPI_ISL_498680, EPI_ISL_498681, EPI_ISL_498682, EPI_ISL_498683, EPI_ISL_498684, EPI_ISL_498685, EPI_ISL_498686, EPI_ISL_498687, EPI_ISL_498688, EPI_ISL_498689, EPI_ISL_498690 | | | |
| see above | Utah Public Health Laboratory | Utah Public Health Laboratory | Heidi Butz, Erin Young, Kelly Oakeson |
| EPI_ISL_498691, EPI_ISL_498692, EPI_ISL_498693, EPI_ISL_498694 | National Institute for Viral Disease Control and Prevention, China CDC | National Institute for Viral Disease Control and Prevention, China CDC | Xiang Zhao,LingLing Mao,Yao Meng,Zhixiao Chen,Yuchao Wu,Yong ZhangBo ZhijianJianqun Zhang,Yang Song,Dayan Wang,WenQing YaoWenbo Xu |
| EPI_ISL_498695, EPI_ISL_498696, EPI_ISL_498697, EPI_ISL_498698, EPI_ISL_498699, EPI_ISL_498700, EPI_ISL_498701, EPI_ISL_498702, EPI_ISL_498703, EPI_ISL_498704, EPI_ISL_498705, EPI_ISL_498706, EPI_ISL_498707, EPI_ISL_498708, EPI_ISL_498709, EPI_ISL_498710, EPI_ISL_498711, EPI_ISL_498712, EPI_ISL_498713, EPI_ISL_498714, EPI_ISL_498715, EPI_ISL_498716, EPI_ISL_498717, EPI_ISL_498718, EPI_ISL_498719, EPI_ISL_498720, EPI_ISL_498721, EPI_ISL_498722, EPI_ISL_498723, EPI_ISL_498724, EPI_ISL_498725, EPI_ISL_498726, EPI_ISL_498727, EPI_ISL_498728, EPI_ISL_498729, EPI_ISL_498730, EPI_ISL_498731, EPI_ISL_498732, EPI_ISL_498733, EPI_ISL_498734, EPI_ISL_498735, EPI_ISL_498736, EPI_ISL_498737, EPI_ISL_498738, EPI_ISL_498739, EPI_ISL_498740, EPI_ISL_498741, EPI_ISL_498742, EPI_ISL_498743, EPI_ISL_498744, EPI_ISL_498745, EPI_ISL_498746, EPI_ISL_498747 | | | |
| see above | Quest Diagnostics | Quest Diagnostics | Rosenthal,S.H., Gerasimova,A., Kagan,R.M. and Owen, R. |
| EPI_ISL_498748, EPI_ISL_498749, EPI_ISL_498750 | Pathology West - NSW Health Pathology | NSW Health Pathology - Institute of Clinical Pathology and Medical Research; Westmead Hospital; University of Sydney | CIDM-PH et al. |
| EPI_ISL_498751 | South Eastern Area Laboratory Services (SEALS) | NSW Health Pathology - Institute of Clinical Pathology and Medical Research; Westmead Hospital; University of Sydney | CIDM-PH et al. |
| EPI_ISL_498761, EPI_ISL_498762 | Pathology West - NSW Health Pathology | NSW Health Pathology - Institute of Clinical Pathology and Medical Research; Westmead Hospital; University of Sydney | CIDM-PH et al. |
| EPI_ISL_498763, EPI_ISL_498764 | Sydney South West Pathology Service (SSWPS) - Liverpool Hospital - NSW Health Pathology | NSW Health Pathology - Institute of Clinical Pathology and Medical Research; Westmead Hospital; University of Sydney | CIDM-PH et al. |
| EPI_ISL_498765 | Sydney South West Pathology Service (SSWPS) - Concord Repatriation General Hospital - NSW Health Pathology | NSW Health Pathology - Institute of Clinical Pathology and Medical Research; Westmead Hospital; University of Sydney | CIDM-PH et al. |
| EPI_ISL_498766, EPI_ISL_498767 | Pathology West - NSW Health Pathology | NSW Health Pathology - Institute of Clinical Pathology and Medical Research; Westmead Hospital; University of Sydney | CIDM-PH et al. |
| EPI_ISL_498768 | South Eastern Area Laboratory Services (SEALS) | NSW Health Pathology - Institute of Clinical Pathology and Medical Research; Westmead Hospital; University of Sydney | CIDM-PH et al. |
| EPI_ISL_498769, EPI_ISL_498770, EPI_ISL_498771, EPI_ISL_498772 | Laverty Pathology | NSW Health Pathology - Institute of Clinical Pathology and Medical Research; Westmead Hospital; University of Sydney | CIDM-PH et al. |
| EPI_ISL_498773 | Douglass Hanly Moir Pathology | NSW Health Pathology - Institute of Clinical Pathology and Medical Research; Westmead Hospital; University of Sydney | CIDM-PH et al. |
| EPI_ISL_498774, EPI_ISL_498775 | Histopath | NSW Health Pathology - Institute of Clinical Pathology and Medical Research; Westmead Hospital; University of Sydney | CIDM-PH et al. |
| EPI_ISL_498776, EPI_ISL_498777 | Pathology West - NSW Health Pathology | NSW Health Pathology - Institute of Clinical Pathology and Medical Research; Westmead Hospital; University of Sydney | CIDM-PH et al. |
| EPI_ISL_498778 | Sydney South West Pathology Service (SSWPS) - Liverpool Hospital - NSW Health Pathology | NSW Health Pathology - Institute of Clinical Pathology and Medical Research; Westmead Hospital; University of Sydney | CIDM-PH et al. |
| EPI_ISL_498779, EPI_ISL_498780 | Pathology West - NSW Health Pathology | NSW Health Pathology - Institute of Clinical Pathology and Medical Research; Westmead Hospital; University of Sydney | CIDM-PH et al. |
| EPI_ISL_498781, EPI_ISL_498782 | Sydney South West Pathology Service (SSWPS) - Liverpool Hospital - NSW Health Pathology | NSW Health Pathology - Institute of Clinical Pathology and Medical Research; Westmead Hospital; University of Sydney | CIDM-PH et al. |
| EPI_ISL_498783, EPI_ISL_498784, EPI_ISL_498785, EPI_ISL_498786, EPI_ISL_498787, EPI_ISL_498788, EPI_ISL_498789, EPI_ISL_498790, EPI_ISL_498791, EPI_ISL_498792 | National Institute of Laboratory Medicine and Referral Center | Genomic Research Lab, BCSIR | Md. Saddam Hossain, Abu Sayeed Mohammad Mahmud, Mohammad Samir Uzzaman, Eshrar Osman, Md. Ahasan Habib, Shahina Akter, Tanjina Akhter Banu, Md. Murshed Hasan Sarkar, Barna Goswami, Iffat Jahan, Tasnim Nafisa, Md. Maruf Ahmed Molla, Mahmuda Yeasmin, Asish Kumar Ghosh, A. K. M. Shamsuzzaman, Sheikh Md. Selim Al Din, Utpal Chandra Ray, Salek Ahmed Sajib, Md. Salim Khan |
| EPI_ISL_498793, EPI_ISL_498794, EPI_ISL_498795, EPI_ISL_498796, EPI_ISL_498797, EPI_ISL_498800, EPI_ISL_498801, EPI_ISL_498802, EPI_ISL_498803, EPI_ISL_498804, EPI_ISL_498805, EPI_ISL_498806 | | | |
| see above | National Institute of Laboratory Medicine and Referral Center | Genomic Research Lab, BCSIR | Md. Murshed Hasan Sarkar, Abu Sayeed Mohammad Mahmud, Mohammad Samir Uzzaman, Eshrar Osman, Md. Ahasan Habib, Shahina Akter, Tanjina Akhter Banu, Barna Goswami, Iffat Jahan, Md. Saddam Hossain, Tasnim Nafisa, Md. Maruf Ahmed Molla, Mahmuda Yeasmin, Asish Kumar Ghosh, A. K. M. Shamsuzzaman, Sheikh Md. Selim Al Din, Utpal Chandra Ray, Salek Ahmed Sajib, Md. Salim Khan |
| EPI_ISL_498808, EPI_ISL_498809, EPI_ISL_498811, EPI_ISL_498814, EPI_ISL_498815, EPI_ISL_498816, EPI_ISL_498817, EPI_ISL_498818, EPI_ISL_498830, EPI_ISL_498892, EPI_ISL_498932, EPI_ISL_498933 | | | |
| see above | National Institute of Laboratory Medicine and Referral Center | Genomic Research Lab, BCSIR | Abu Sayeed Mohammad Mahmud, Mohammad Samir Uzzaman, Eshrar Osman, Md. Ahasan Habib, Shahina Akter, Tanjina Akhter Banu, Md. Murshed Hasan Sarkar, Barna Goswami, Iffat Jahan, Md. Saddam Hossain, Tasnim Nafisa, Md. Maruf Ahmed Molla, Mahmuda Yeasmin, Asish Kumar Ghosh, A. K. M. Shamsuzzaman, Sheikh Md. Selim Al Din, Utpal Chandra Ray, Salek Ahmed Sajib, Md. Salim Khan |
| EPI_ISL_499042 | National Institute for Biological Standards and Control (NIBSC) | National Institute for Biological Standards and Control (NIBSC) | Javier Martin, Dimitra Klapsa, Thomas Wilton |
| EPI_ISL_499083 | Instituto de Virologia "Dr. J. M. Vanella", Facultad de Ciencias Medicas, Universidad Nacional de Cordoba. Laboratorio Central de la Provincia de Cordoba, Argentina. Ministerio de Salud de la provincia de Cordoba, Argentina. | Laboratorio de Virología, Hospital de Niños Ricardo Gutiérrez, CABA, Argentina. | Sandra Gallego, Brenda Konigheim, Sebastian Blanco, Lorena Spinsanti, Javier Aguilar, Adrian Diaz, Gonzalo Castro, Gabriela Barbas, Mercedes Nabaes, Stephanie Goya, Monica Natale, Silvina Lusso, Mariana Viegas. |
| EPI_ISL_499266, EPI_ISL_499267, EPI_ISL_499268, EPI_ISL_499269 | Queens Medical Centre, Clinical Microbiology Department / DeepSeq Nottingham | COVID-19 Genomics UK (COG-UK) Consortium | Gemma Clark, Wendy Smith, Manjinder Khakh, Vicki M Fleming, Michelle M Lister, Hannah Howson-Wells, Jonathan Ball, Patrick McClure, Joseph Chappell, Theocharis Tsoleridis, Nadine Holmes, Matthew Carlisle, Christopher Moore, Fei Sang, Johnny Debebe, Victoria Wright, Matthew Loose |
| EPI_ISL_499270, EPI_ISL_499271, EPI_ISL_499272, EPI_ISL_499273, EPI_ISL_499274, EPI_ISL_499275, EPI_ISL_499276, EPI_ISL_499277, EPI_ISL_499278, EPI_ISL_499279, EPI_ISL_499280, EPI_ISL_499281, EPI_ISL_499282, EPI_ISL_499283, EPI_ISL_499284, EPI_ISL_499285, EPI_ISL_499286, EPI_ISL_499287, EPI_ISL_499288, EPI_ISL_499289, EPI_ISL_499290, EPI_ISL_499291, EPI_ISL_499292, EPI_ISL_499293, EPI_ISL_499294, EPI_ISL_499295, EPI_ISL_499296, EPI_ISL_499297, EPI_ISL_499298, EPI_ISL_499299, EPI_ISL_499300, EPI_ISL_499301, EPI_ISL_499302, EPI_ISL_499303, EPI_ISL_499304, EPI_ISL_499305, EPI_ISL_499306, EPI_ISL_499307, EPI_ISL_499308, EPI_ISL_499309, EPI_ISL_499310, EPI_ISL_499311, EPI_ISL_499312, EPI_ISL_499313, EPI_ISL_499314, EPI_ISL_499315, EPI_ISL_499316, EPI_ISL_499317, EPI_ISL_499318, EPI_ISL_499319, EPI_ISL_499320, EPI_ISL_499321, EPI_ISL_499322, EPI_ISL_499323, EPI_ISL_499324, EPI_ISL_499325, EPI_ISL_499326, EPI_ISL_499327, EPI_ISL_499328, EPI_ISL_499329 | | | |
| see above | Centre for Enzyme Innovation, University of Portsmouth / Translational Research Laboratory, Portsmouth Hospitals NHS Trust | COVID-19 Genomics UK (COG-UK) Consortium | Angela Beckett,Yann Bourgeois,Garry Scarlett,Sharon Glaysher,Scott Elliott,Kelly Bicknell,Robert Impey,Allyson Lloyd,Sarah Wyllie,Ethan Butcher,Anoop Chauhan,Samuel Robson |
| EPI_ISL_499330, EPI_ISL_499331, EPI_ISL_499332, EPI_ISL_499333, EPI_ISL_499334, EPI_ISL_499335, EPI_ISL_499336, EPI_ISL_499337, EPI_ISL_499338, EPI_ISL_499339, EPI_ISL_499340, EPI_ISL_499341, EPI_ISL_499342, EPI_ISL_499343, EPI_ISL_499344, EPI_ISL_499345, EPI_ISL_499346, EPI_ISL_499347, EPI_ISL_499348, EPI_ISL_499349, EPI_ISL_499350, EPI_ISL_499351 | | | |
| see above | Virology Department, Sheffield Teaching Hospitals NHS Foundation Trust/Department of Infection, Immunity and Cardiovascular Disease, The Medical School, University of Sheffield | COVID-19 Genomics UK (COG-UK) Consortium | Thushan de Silva, Matthew Parker, Nikki Smith, Adri Angyal, Rebecca Brown, Luke Green, Rachel Tucker, Paul Parsons, Danielle Groves, Katie Johnson, Laura Carrilero, Alex Keeley, Dave Partridge, Matthew Wyles, Benjamin Lindsey, Mehmet Yavuz, Mohammad Raza, Cariad Evans |
| EPI_ISL_499352, EPI_ISL_499353, EPI_ISL_499354 | West of Scotland Specialist Virology Centre, NHSGGC / MRC-University of Glasgow Centre for Virus Research | COVID-19 Genomics UK (COG-UK) Consortium | Ana da Silva Filipe, Natasha Johnson, Kathy Smollett, Daniel Mair, Stephen Carmichael, Lily Tong, Jenna Nichols, Elihu Aranday-Cortes, Kirstyn Brunker, Yasmin Parr, Alice Broos, Kyriaki Nomikou; Sarah McDonald, Marc Niebel, Patawee Asamaphan; Richard Orton, Joseph Hughes, Sreenu Vattipally, David L Robertson; Alasdair MacLean, Rory Gunson; Kathy Li, Natasha Jesudason, Rajiv Shah, James Shepherd, Antonia Ho, Emma Thomson |
| EPI_ISL_499355, EPI_ISL_499356, EPI_ISL_499357, EPI_ISL_499358, EPI_ISL_499359, EPI_ISL_499360, EPI_ISL_499361, EPI_ISL_499362, EPI_ISL_499363, EPI_ISL_499364, EPI_ISL_499365, EPI_ISL_499366, EPI_ISL_499367, EPI_ISL_499368, EPI_ISL_499369, EPI_ISL_499370, EPI_ISL_499371, EPI_ISL_499372, EPI_ISL_499373, EPI_ISL_499374, EPI_ISL_499375, EPI_ISL_499376, EPI_ISL_499377, EPI_ISL_499378, EPI_ISL_499379, EPI_ISL_499380, EPI_ISL_499381, EPI_ISL_499382, EPI_ISL_499383, EPI_ISL_499384, EPI_ISL_499385, EPI_ISL_499386, EPI_ISL_499387, EPI_ISL_499388, EPI_ISL_499389, EPI_ISL_499390, EPI_ISL_499391, EPI_ISL_499392, EPI_ISL_499393, EPI_ISL_499394, EPI_ISL_499395, EPI_ISL_499396, EPI_ISL_499397 | | | |
| see above | Originating lab: Wales Specialist Virology Centre Sequencing lab: Pathogen Genomics Unit | COVID-19 Genomics UK (COG-UK) Consortium | Catherine Moore, Johnathan Evans, Laura Gifford, Malorie Perry, Simon Cottrell, Angela Marchbank, Alec Birchley, Alexander Adams, Amy Gaskin, Bree Gatica-Wilcox, Jason Coombes, Joel Southgate, Lauren Gilbert, Lee Graham, Nicole Pacchiarini, Sara Kumziene-Summerhayes, Sarah Taylor, Sophie Jones, Sara Rey, Matthew Bull, Joanne Watkins, Sally Corden, Tom Connor |
| EPI_ISL_499398, EPI_ISL_499399, EPI_ISL_499400, EPI_ISL_499401, EPI_ISL_499402, EPI_ISL_499403, EPI_ISL_499404, EPI_ISL_499405, EPI_ISL_499406, EPI_ISL_499407, EPI_ISL_499408, EPI_ISL_499409, EPI_ISL_499410, EPI_ISL_499411, EPI_ISL_499412, EPI_ISL_499413, EPI_ISL_499414, EPI_ISL_499415, EPI_ISL_499416, EPI_ISL_499417, EPI_ISL_499418, EPI_ISL_499419, EPI_ISL_499420, EPI_ISL_499421, EPI_ISL_499422, EPI_ISL_499423, EPI_ISL_499424, EPI_ISL_499425, EPI_ISL_499426, EPI_ISL_499427, EPI_ISL_499428, EPI_ISL_499429, EPI_ISL_499430, EPI_ISL_499431, EPI_ISL_499432, EPI_ISL_499433, EPI_ISL_499434, EPI_ISL_499435, EPI_ISL_499436, EPI_ISL_499437, EPI_ISL_499438, EPI_ISL_499439, EPI_ISL_499440, EPI_ISL_499441, EPI_ISL_499442, EPI_ISL_499443, EPI_ISL_499444, EPI_ISL_499445, EPI_ISL_499446, EPI_ISL_499447, EPI_ISL_499448, EPI_ISL_499449, EPI_ISL_499450, EPI_ISL_499451, EPI_ISL_499452, EPI_ISL_499453, EPI_ISL_499454, EPI_ISL_499455, EPI_ISL_499456, EPI_ISL_499457, EPI_ISL_499458, EPI_ISL_499459 | | | |
| see above | Wales Specialist Virology Centre Sequencing lab: Pathogen Genomics Unit | COVID-19 Genomics UK (COG-UK) Consortium | Catherine Moore, Johnathan Evans, Laura Gifford, Malorie Perry, Simon Cottrell, Angela Marchbank, Alec Birchley, Alexander Adams, Amy Gaskin, Bree Gatica-Wilcox, Jason Coombes, Joel Southgate, Lauren Gilbert, Lee Graham, Nicole Pacchiarini, Sara Kumziene-Summerhayes, Sarah Taylor, Sophie Jones, Sara Rey, Matthew Bull, Joanne Watkins, Sally Corden, Tom Connor |
| EPI_ISL_499460, EPI_ISL_499461, EPI_ISL_499462, EPI_ISL_499463, EPI_ISL_499464, EPI_ISL_499465, EPI_ISL_499466, EPI_ISL_499467, EPI_ISL_499468, EPI_ISL_499469, EPI_ISL_499470, EPI_ISL_499471 | | | |
| see above | Liverpool Clinical Laboratories | COVID-19 Genomics UK (COG-UK) Consortium | Sam Haldenby, Anita Lucaci, Steve Paterson, Julian Hiscox, Alistair Darby, M Almsaud, A Alrezaihi, Muhannad Alruwaili, Stuart D Armstrong, Jones Benjamin, Eleanor G Bentley, Anu Chawla, Jordan J Clark, Angela Cowell, Richard Eccles, Isabel García-Dorival, Matthew Gemmell, Alessandro Gerada, PKF Gilmore, Richard Gregory, Ximeng Han, Catherine Hartley, Margaret Hughes, Miren Iturriza-Gomara, James Johnson, L Luu, Jenifer Manson, Charlotte Nelson, Elaine O'Toole, Cassie Olateju, Rebekah Penrice-Randal , Lucille Rainbow, N.P Randle, Trevor Ian Robinson, Parul Sharma, Ghada T Shawli, James P Stewart, Neil Swainston, Ecaterina Vamos, Joanne Watts, Mark Whitehead |
| EPI_ISL_499472 | Northumbria University / South Tees Hospitals NHS Foundation Trust / North Cumbria Integrated Care NHS Foundation Trust / North Tees and Hartlepool NHS Foundation Trust / Newcastle Hospitals NHS Foundation Trust | COVID-19 Genomics UK (COG-UK) Consortium | Darren L Smith,Andrew Nelson,Matthew Bashton,Greg R Young,Joshua Loh,John Allan,Mohammad A Tariq,Giles S Holt,Gary Black,Wen C Yew,Lynn Dover,Paul Baker,Steve Liggett,Sarah Essex,Jane Greenaway,Debra Padgett,Clive Graham,Garren Scott,Edward Barton,Emma Swindells,Brendan Payne,Jennifer Collins,Yusri Taha,Gary Eltringham |

| | | | |
|---|---|---|---|
| EPI_ISL_499473 | Liverpool Clinical Laboratories | COVID-19 Genomics UK (COG-UK) Consortium | Sam Haldenby, Anita Lucaci, Steve Paterson, Julian Hiscox, Alistair Darby, M Almsaud, A Alrezaihi, Muhannad Alruwaili, Stuart D Armstrong, Jones Benjamin, Eleanor G Bentley, Anu Chawla, Jordan J Clark, Angela Cowell, Richard Eccles, Isabel García-Dorival, Matthew Gemmell, Alessandro Gerada, PKF Gilmore, Richard Gregory, Ximeng Han, Catherine Hartley, Margaret Hughes, Miren Iturriza-Gomara, James Johnson, L Luu, Jenifer Manson, Charlotte Nelson, Elaine O’Toole, Cassie Olateju, Rebekah Penrice-Randal , Lucille Rainbow, N.P Randle, Trevor Ian Robinson, Parul Sharma, Ghada T Shawli, James P Stewart, Neil Swainston, Ecaterina Vamos, Joanne Watts, Mark Whitehead |
| EPI_ISL_499474 | Northumbria University / South Tees Hospitals NHS Foundation Trust / North Cumbria Integrated Care NHS Foundation Trust / North Tees and Hartlepool NHS Foundation Trust / Newcastle Hospitals NHS Foundation Trust | COVID-19 Genomics UK (COG-UK) Consortium | Darren L Smith,Andrew Nelson,Matthew Bashton,Greg R Young,Joshua Loh,John Allan,Mohammad A Tariq,Giles S Holt,Gary Black,Wen C Yew,Lynn Dover,Paul Baker,Steve Liggett,Sarah Essex,Jane Greenaway,Debra Padgett,Clive Graham,Garren Scott,Edward Barton,Emma Swindells,Brendan Payne,Jennifer Collins,Yusri Taha,Gary Eltringham |
| EPI_ISL_499475, EPI_ISL_499476, EPI_ISL_499477 | Liverpool Clinical Laboratories | COVID-19 Genomics UK (COG-UK) Consortium | Sam Haldenby, Anita Lucaci, Steve Paterson, Julian Hiscox, Alistair Darby, M Almsaud, A Alrezaihi, Muhannad Alruwaili, Stuart D Armstrong, Jones Benjamin, Eleanor G Bentley, Anu Chawla, Jordan J Clark, Angela Cowell, Richard Eccles, Isabel García-Dorival, Matthew Gemmell, Alessandro Gerada, PKF Gilmore, Richard Gregory, Ximeng Han, Catherine Hartley, Margaret Hughes, Miren Iturriza-Gomara, James Johnson, L Luu, Jenifer Manson, Charlotte Nelson, Elaine O’Toole, Cassie Olateju, Rebekah Penrice-Randal , Lucille Rainbow, N.P Randle, Trevor Ian Robinson, Parul Sharma, Ghada T Shawli, James P Stewart, Neil Swainston, Ecaterina Vamos, Joanne Watts, Mark Whitehead |
| EPI_ISL_499478 | Northumbria University / South Tees Hospitals NHS Foundation Trust / North Cumbria Integrated Care NHS Foundation Trust / North Tees and Hartlepool NHS Foundation Trust / Newcastle Hospitals NHS Foundation Trust | COVID-19 Genomics UK (COG-UK) Consortium | Darren L Smith,Andrew Nelson,Matthew Bashton,Greg R Young,Joshua Loh,John Allan,Mohammad A Tariq,Giles S Holt,Gary Black,Wen C Yew,Lynn Dover,Paul Baker,Steve Liggett,Sarah Essex,Jane Greenaway,Debra Padgett,Clive Graham,Garren Scott,Edward Barton,Emma Swindells,Brendan Payne,Jennifer Collins,Yusri Taha,Gary Eltringham |
| EPI_ISL_499479 | Quadram Institute Bioscience | COVID-19 Genomics UK (COG-UK) Consortium | Dave J. Baker, Gemma L. Kay, Alp Aydin, Thanh Le-Viet, Steven Rudder, Ana P. Tedim, Anastasia Kolyva, Maria Diaz, Leonardo de Oliveira Martins, Nabil-Fareed Alikhan, Lizzie Meadows, Rachael Stanley, Ngozi Elumogo, Muhammed Yasir, Nicholas M. Thomson, Alexander J Trotter, Rachel Gilroy, Samuel Bloomfield, Claire Stuart, Andrew Bell, Reenesh Prakash, Samir Dervisevic, Alison E. Mather, John Wain, Mark Webber, Andrew J. Page, Justin O’Grady |
| EPI_ISL_499480, EPI_ISL_499481 | Liverpool Clinical Laboratories | COVID-19 Genomics UK (COG-UK) Consortium | Sam Haldenby, Anita Lucaci, Steve Paterson, Julian Hiscox, Alistair Darby, M Almsaud, A Alrezaihi, Muhannad Alruwaili, Stuart D Armstrong, Jones Benjamin, Eleanor G Bentley, Anu Chawla, Jordan J Clark, Angela Cowell, Richard Eccles, Isabel García-Dorival, Matthew Gemmell, Alessandro Gerada, PKF Gilmore, Richard Gregory, Ximeng Han, Catherine Hartley, Margaret Hughes, Miren Iturriza-Gomara, James Johnson, L Luu, Jenifer Manson, Charlotte Nelson, Elaine O’Toole, Cassie Olateju, Rebekah Penrice-Randal , Lucille Rainbow, N.P Randle, Trevor Ian Robinson, Parul Sharma, Ghada T Shawli, James P Stewart, Neil Swainston, Ecaterina Vamos, Joanne Watts, Mark Whitehead |
| EPI_ISL_499482 | Northumbria University / South Tees Hospitals NHS Foundation Trust / North Cumbria Integrated Care NHS Foundation Trust / North Tees and Hartlepool NHS Foundation Trust / Newcastle Hospitals NHS Foundation Trust | COVID-19 Genomics UK (COG-UK) Consortium | Darren L Smith,Andrew Nelson,Matthew Bashton,Greg R Young,Joshua Loh,John Allan,Mohammad A Tariq,Giles S Holt,Gary Black,Wen C Yew,Lynn Dover,Paul Baker,Steve Liggett,Sarah Essex,Jane Greenaway,Debra Padgett,Clive Graham,Garren Scott,Edward Barton,Emma Swindells,Brendan Payne,Jennifer Collins,Yusri Taha,Gary Eltringham |
| EPI_ISL_499483, EPI_ISL_499484 | Liverpool Clinical Laboratories | COVID-19 Genomics UK (COG-UK) Consortium | Sam Haldenby, Anita Lucaci, Steve Paterson, Julian Hiscox, Alistair Darby, M Almsaud, A Alrezaihi, Muhannad Alruwaili, Stuart D Armstrong, Jones Benjamin, Eleanor G Bentley, Anu Chawla, Jordan J Clark, Angela Cowell, Richard Eccles, Isabel García-Dorival, Matthew Gemmell, Alessandro Gerada, PKF Gilmore, Richard Gregory, Ximeng Han, Catherine Hartley, Margaret Hughes, Miren Iturriza-Gomara, James Johnson, L Luu, Jenifer Manson, Charlotte Nelson, Elaine O’Toole, Cassie Olateju, Rebekah Penrice-Randal , Lucille Rainbow, N.P Randle, Trevor Ian Robinson, Parul Sharma, Ghada T Shawli, James P Stewart, Neil Swainston, Ecaterina Vamos, Joanne Watts, Mark Whitehead |
| EPI_ISL_499485 | Northumbria University / South Tees Hospitals NHS Foundation Trust / North Cumbria Integrated Care NHS Foundation Trust / North Tees and Hartlepool NHS Foundation Trust / Newcastle Hospitals NHS Foundation Trust | COVID-19 Genomics UK (COG-UK) Consortium | Darren L Smith,Andrew Nelson,Matthew Bashton,Greg R Young,Joshua Loh,John Allan,Mohammad A Tariq,Giles S Holt,Gary Black,Wen C Yew,Lynn Dover,Paul Baker,Steve Liggett,Sarah Essex,Jane Greenaway,Debra Padgett,Clive Graham,Garren Scott,Edward Barton,Emma Swindells,Brendan Payne,Jennifer Collins,Yusri Taha,Gary Eltringham |
| EPI_ISL_499486, EPI_ISL_499487, EPI_ISL_499488, EPI_ISL_499489 | Liverpool Clinical Laboratories | COVID-19 Genomics UK (COG-UK) Consortium | Sam Haldenby, Anita Lucaci, Steve Paterson, Julian Hiscox, Alistair Darby, M Almsaud, A Alrezaihi, Muhannad Alruwaili, Stuart D Armstrong, Jones Benjamin, Eleanor G Bentley, Anu Chawla, Jordan J Clark, Angela Cowell, Richard Eccles, Isabel García-Dorival, Matthew Gemmell, Alessandro Gerada, PKF Gilmore, Richard Gregory, Ximeng Han, Catherine Hartley, Margaret Hughes, Miren Iturriza-Gomara, James Johnson, L Luu, Jenifer Manson, Charlotte Nelson, Elaine O’Toole, Cassie Olateju, Rebekah Penrice-Randal , Lucille Rainbow, N.P Randle, Trevor Ian Robinson, Parul Sharma, Ghada T Shawli, James P Stewart, Neil Swainston, Ecaterina Vamos, Joanne Watts, Mark Whitehead |
| EPI_ISL_499490 | Quadram Institute Bioscience | COVID-19 Genomics UK (COG-UK) Consortium | Dave J. Baker, Gemma L. Kay, Alp Aydin, Thanh Le-Viet, Steven Rudder, Ana P. Tedim, Anastasia Kolyva, Maria Diaz, Leonardo de Oliveira Martins, Nabil-Fareed Alikhan, Lizzie Meadows, Rachael Stanley, Ngozi Elumogo, Muhammed Yasir, Nicholas M. Thomson, Alexander J Trotter, Rachel Gilroy, Samuel Bloomfield, Claire Stuart, Andrew Bell, Reenesh Prakash, Samir Dervisevic, Alison E. Mather, John Wain, Mark Webber, Andrew J. Page, Justin O’Grady |
| EPI_ISL_499491, EPI_ISL_499492 | Liverpool Clinical Laboratories | COVID-19 Genomics UK (COG-UK) Consortium | Sam Haldenby, Anita Lucaci, Steve Paterson, Julian Hiscox, Alistair Darby, M Almsaud, A Alrezaihi, Muhannad Alruwaili, Stuart D Armstrong, Jones Benjamin, Eleanor G Bentley, Anu Chawla, Jordan J Clark, Angela Cowell, Richard Eccles, Isabel García-Dorival, Matthew Gemmell, Alessandro Gerada, PKF Gilmore, Richard Gregory, Ximeng Han, Catherine Hartley, Margaret Hughes, Miren Iturriza-Gomara, James Johnson, L Luu, Jenifer Manson, Charlotte Nelson, Elaine O’Toole, Cassie Olateju, Rebekah Penrice-Randal , Lucille Rainbow, N.P Randle, Trevor Ian Robinson, Parul Sharma, Ghada T Shawli, James P Stewart, Neil Swainston, Ecaterina Vamos, Joanne Watts, Mark Whitehead |
| EPI_ISL_499493, EPI_ISL_499494 | Northumbria University / South Tees Hospitals NHS Foundation Trust / North Cumbria Integrated Care NHS Foundation Trust / North Tees and Hartlepool NHS Foundation Trust / Newcastle Hospitals NHS Foundation Trust | COVID-19 Genomics UK (COG-UK) Consortium | Darren L Smith,Andrew Nelson,Matthew Bashton,Greg R Young,Joshua Loh,John Allan,Mohammad A Tariq,Giles S Holt,Gary Black,Wen C Yew,Lynn Dover,Paul Baker,Steve Liggett,Sarah Essex,Jane Greenaway,Debra Padgett,Clive Graham,Garren Scott,Edward Barton,Emma Swindells,Brendan Payne,Jennifer Collins,Yusri Taha,Gary Eltringham |
| EPI_ISL_499495 | Quadram Institute Bioscience | COVID-19 Genomics UK (COG-UK) Consortium | Dave J. Baker, Gemma L. Kay, Alp Aydin, Thanh Le-Viet, Steven Rudder, Ana P. Tedim, Anastasia Kolyva, Maria Diaz, Leonardo de Oliveira Martins, Nabil-Fareed Alikhan, Lizzie Meadows, Rachael Stanley, Ngozi Elumogo, Muhammed Yasir, Nicholas M. Thomson, Alexander J Trotter, Rachel Gilroy, Samuel Bloomfield, Claire Stuart, Andrew Bell, Reenesh Prakash, Samir Dervisevic, Alison E. Mather, John Wain, Mark Webber, Andrew J. Page, Justin O’Grady |
| EPI_ISL_499496, EPI_ISL_499497, EPI_ISL_499498, EPI_ISL_499499, EPI_ISL_499500, EPI_ISL_499501 | Liverpool Clinical Laboratories | COVID-19 Genomics UK (COG-UK) Consortium | Sam Haldenby, Anita Lucaci, Steve Paterson, Julian Hiscox, Alistair Darby, M Almsaud, A Alrezaihi, Muhannad Alruwaili, Stuart D Armstrong, Jones Benjamin, Eleanor G Bentley, Anu Chawla, Jordan J Clark, Angela Cowell, Richard Eccles, Isabel García-Dorival, Matthew Gemmell, Alessandro Gerada, PKF Gilmore, Richard Gregory, Ximeng Han, Catherine Hartley, Margaret Hughes, Miren Iturriza-Gomara, James Johnson, L Luu, Jenifer Manson, Charlotte Nelson, Elaine O’Toole, Cassie Olateju, Rebekah Penrice-Randal , Lucille Rainbow, N.P Randle, Trevor Ian Robinson, Parul Sharma, Ghada T Shawli, James P Stewart, Neil Swainston, Ecaterina Vamos, Joanne Watts, Mark Whitehead |
| EPI_ISL_499502, EPI_ISL_499503, EPI_ISL_499504, EPI_ISL_499505, EPI_ISL_499506, EPI_ISL_499507, EPI_ISL_499508 | Northumbria University / South Tees Hospitals NHS Foundation Trust / North Cumbria Integrated Care NHS Foundation Trust / North Tees and Hartlepool NHS Foundation Trust / Newcastle Hospitals NHS Foundation Trust | COVID-19 Genomics UK (COG-UK) Consortium | Darren L Smith,Andrew Nelson,Matthew Bashton,Greg R Young,Joshua Loh,John Allan,Mohammad A Tariq,Giles S Holt,Gary Black,Wen C Yew,Lynn Dover,Paul Baker,Steve Liggett,Sarah Essex,Jane Greenaway,Debra Padgett,Clive Graham,Garren Scott,Edward Barton,Emma Swindells,Brendan Payne,Jennifer Collins,Yusri Taha,Gary Eltringham |
| EPI_ISL_499509 | Liverpool Clinical Laboratories | COVID-19 Genomics UK (COG-UK) Consortium | Sam Haldenby, Anita Lucaci, Steve Paterson, Julian Hiscox, Alistair Darby, M Almsaud, A Alrezaihi, Muhannad Alruwaili, Stuart D Armstrong, Jones Benjamin, Eleanor G Bentley, Anu Chawla, Jordan J Clark, Angela Cowell, Richard Eccles, Isabel García-Dorival, Matthew Gemmell, Alessandro Gerada, PKF Gilmore, Richard Gregory, Ximeng Han, Catherine Hartley, Margaret Hughes, Miren Iturriza-Gomara, James Johnson, L Luu, Jenifer Manson, Charlotte Nelson, Elaine O’Toole, Cassie Olateju, Rebekah Penrice-Randal , Lucille Rainbow, N.P Randle, Trevor Ian Robinson, Parul Sharma, Ghada T Shawli, James P Stewart, Neil Swainston, Ecaterina Vamos, Joanne Watts, Mark Whitehead |
| EPI_ISL_499510 | Northumbria University / South Tees Hospitals NHS Foundation Trust / North Cumbria Integrated Care NHS Foundation Trust / North Tees and Hartlepool NHS Foundation Trust / Newcastle Hospitals NHS Foundation Trust | COVID-19 Genomics UK (COG-UK) Consortium | Darren L Smith,Andrew Nelson,Matthew Bashton,Greg R Young,Joshua Loh,John Allan,Mohammad A Tariq,Giles S Holt,Gary Black,Wen C Yew,Lynn Dover,Paul Baker,Steve Liggett,Sarah Essex,Jane Greenaway,Debra Padgett,Clive Graham,Garren Scott,Edward Barton,Emma Swindells,Brendan Payne,Jennifer Collins,Yusri Taha,Gary Eltringham |
| EPI_ISL_499511, EPI_ISL_499512, EPI_ISL_499513, EPI_ISL_499514 | Liverpool Clinical Laboratories | COVID-19 Genomics UK (COG-UK) Consortium | Sam Haldenby, Anita Lucaci, Steve Paterson, Julian Hiscox, Alistair Darby, M Almsaud, A Alrezaihi, Muhannad Alruwaili, Stuart D Armstrong, Jones Benjamin, Eleanor G Bentley, Anu Chawla, Jordan J Clark, Angela Cowell, Richard Eccles, Isabel García-Dorival, Matthew Gemmell, Alessandro Gerada, PKF Gilmore, Richard Gregory, Ximeng Han, Catherine Hartley, Margaret Hughes, Miren Iturriza-Gomara, James Johnson, L Luu, Jenifer Manson, Charlotte Nelson, Elaine O’Toole, Cassie Olateju, Rebekah Penrice-Randal , Lucille Rainbow, N.P Randle, Trevor Ian Robinson, Parul Sharma, Ghada T Shawli, James P Stewart, Neil Swainston, Ecaterina Vamos, Joanne Watts, Mark Whitehead |
| EPI_ISL_499515 | Northumbria University / South Tees Hospitals NHS Foundation Trust / North Cumbria Integrated Care NHS Foundation Trust / North Tees and Hartlepool NHS Foundation Trust / Newcastle Hospitals NHS Foundation Trust | COVID-19 Genomics UK (COG-UK) Consortium | Darren L Smith,Andrew Nelson,Matthew Bashton,Greg R Young,Joshua Loh,John Allan,Mohammad A Tariq,Giles S Holt,Gary Black,Wen C Yew,Lynn Dover,Paul Baker,Steve Liggett,Sarah Essex,Jane Greenaway,Debra Padgett,Clive Graham,Garren Scott,Edward Barton,Emma Swindells,Brendan Payne,Jennifer Collins,Yusri Taha,Gary Eltringham |
| EPI_ISL_499516, EPI_ISL_499517, EPI_ISL_499518, EPI_ISL_499519, EPI_ISL_499520, EPI_ISL_499521, EPI_ISL_499522, EPI_ISL_499523, EPI_ISL_499524, EPI_ISL_499525, EPI_ISL_499526, EPI_ISL_499527, EPI_ISL_499528, EPI_ISL_499529, EPI_ISL_499530, EPI_ISL_499531, EPI_ISL_499532, EPI_ISL_499533, EPI_ISL_499534, EPI_ISL_499535, EPI_ISL_499536, EPI_ISL_499537, EPI_ISL_499538, EPI_ISL_499539, EPI_ISL_499540, EPI_ISL_499541, EPI_ISL_499542, EPI_ISL_499543, EPI_ISL_499544, EPI_ISL_499545 | | | |
| see above | Liverpool Clinical Laboratories | COVID-19 Genomics UK (COG-UK) Consortium | Sam Haldenby, Anita Lucaci, Steve Paterson, Julian Hiscox, Alistair Darby, M Almsaud, A Alrezaihi, Muhannad Alruwaili, Stuart D Armstrong, Jones Benjamin, Eleanor G Bentley, Anu Chawla, Jordan J Clark, Angela Cowell, Richard Eccles, Isabel García-Dorival, Matthew Gemmell, Alessandro Gerada, PKF Gilmore, Richard Gregory, Ximeng Han, Catherine Hartley, Margaret Hughes, Miren Iturriza-Gomara, James Johnson, L Luu, Jenifer Manson, Charlotte Nelson, Elaine O’Toole, Cassie Olateju, Rebekah Penrice-Randal , Lucille Rainbow, N.P Randle, Trevor Ian Robinson, Parul Sharma, Ghada T Shawli, James P Stewart, Neil Swainston, Ecaterina Vamos, Joanne Watts, Mark Whitehead |
| EPI_ISL_499546, EPI_ISL_499547, EPI_ISL_499548, EPI_ISL_499549, EPI_ISL_499550, EPI_ISL_499551, EPI_ISL_499552, EPI_ISL_499553, EPI_ISL_499554, EPI_ISL_499555, EPI_ISL_499556 | | | |
| see above | Northumbria University / South Tees Hospitals NHS Foundation Trust / North Cumbria Integrated Care NHS Foundation Trust / North Tees and Hartlepool NHS Foundation Trust / Newcastle Hospitals NHS Foundation Trust | COVID-19 Genomics UK (COG-UK) Consortium | Darren L Smith,Andrew Nelson,Matthew Bashton,Greg R Young,Joshua Loh,John Allan,Mohammad A Tariq,Giles S Holt,Gary Black,Wen C Yew,Lynn Dover,Paul Baker,Steve Liggett,Sarah Essex,Jane Greenaway,Debra Padgett,Clive Graham,Garren Scott,Edward Barton,Emma Swindells,Brendan Payne,Jennifer Collins,Yusri Taha,Gary Eltringham |
| EPI_ISL_499557 | Liverpool Clinical Laboratories | COVID-19 Genomics UK (COG-UK) Consortium | Sam Haldenby, Anita Lucaci, Steve Paterson, Julian Hiscox, Alistair Darby, M Almsaud, A Alrezaihi, Muhannad Alruwaili, Stuart D Armstrong, Jones Benjamin, Eleanor G Bentley, Anu Chawla, Jordan J Clark, Angela Cowell, Richard Eccles, Isabel García-Dorival, Matthew Gemmell, Alessandro Gerada, PKF Gilmore, Richard Gregory, Ximeng Han, Catherine Hartley, Margaret Hughes, Miren Iturriza-Gomara, James Johnson, L Luu, Jenifer Manson, Charlotte Nelson, Elaine O’Toole, Cassie Olateju, Rebekah Penrice-Randal , Lucille Rainbow, N.P Randle, Trevor Ian Robinson, Parul Sharma, Ghada T Shawli, James P Stewart, Neil Swainston, Ecaterina Vamos, Joanne Watts, Mark Whitehead |
| EPI_ISL_499558, EPI_ISL_499559, EPI_ISL_499560, EPI_ISL_499561 | Northumbria University / South Tees Hospitals NHS Foundation Trust / North Cumbria Integrated Care NHS Foundation Trust / North Tees and Hartlepool NHS Foundation Trust / Newcastle Hospitals NHS Foundation Trust | COVID-19 Genomics UK (COG-UK) Consortium | Darren L Smith,Andrew Nelson,Matthew Bashton,Greg R Young,Joshua Loh,John Allan,Mohammad A Tariq,Giles S Holt,Gary Black,Wen C Yew,Lynn Dover,Paul Baker,Steve Liggett,Sarah Essex,Jane Greenaway,Debra Padgett,Clive Graham,Garren Scott,Edward Barton,Emma Swindells,Brendan Payne,Jennifer Collins,Yusri Taha,Gary Eltringham |
| EPI_ISL_499562 | Quadram Institute Bioscience | COVID-19 Genomics UK (COG-UK) Consortium | Dave J. Baker, Gemma L. Kay, Alp Aydin, Thanh Le-Viet, Steven Rudder, Ana P. Tedim, Anastasia Kolyva, Maria Diaz, Leonardo de Oliveira Martins, Nabil-Fareed Alikhan, Lizzie Meadows, Rachael Stanley, Ngozi Elumogo, Muhammed Yasir, Nicholas M. Thomson, Alexander J Trotter, Rachel Gilroy, Samuel Bloomfield, Claire Stuart, Andrew Bell, Reenesh Prakash, Samir Dervisevic, Alison E. Mather, John Wain, Mark Webber, Andrew J. Page, Justin O’Grady |

| | | | |
|---|---|---|---|
| EPI_ISL_499563 | Northumbria University / South Tees Hospitals NHS Foundation Trust / North Cumbria Integrated Care NHS Foundation Trust / North Tees and Hartlepool NHS Foundation Trust / Newcastle Hospitals NHS Foundation Trust | COVID-19 Genomics UK (COG-UK) Consortium | Darren L Smith,Andrew Nelson,Matthew Bashton,Greg R Young,Joshua Loh,John Allan,Mohammad A Tariq,Giles S Holt,Gary Black,Wen C Yew,Lynn Dover,Paul Baker,Steve Liggett,Sarah Essex,Jane Greenaway,Debra Padgett,Clive Graham,Garren Scott,Edward Barton,Emma Swindells,Brendan Payne,Jennifer Collins,Yusri Taha,Gary Eltringham |
| EPI_ISL_499564, EPI_ISL_499565, EPI_ISL_499566, EPI_ISL_499567 | Liverpool Clinical Laboratories | COVID-19 Genomics UK (COG-UK) Consortium | Sam Haldenby, Anita Lucaci, Steve Paterson, Julian Hiscox, Alistair Darby, M Almsaud, A Alrezaihi, Muhannad Alruwaili, Stuart D Armstrong, Jones Benjamin, Eleanor G Bentley, Anu Chawla, Jordan J Clark, Angela Cowell, Richard Eccles, Isabel García-Dorival, Matthew Gemmell, Alessandro Gerada, PKF Gilmore, Richard Gregory, Ximeng Han, Catherine Hartley, Margaret Hughes, Miren Iturriza-Gomara, James Johnson, L Luu, Jenifer Manson, Charlotte Nelson, Elaine O'Toole, Cassie Olateju, Rebekah Penrice-Randal , Lucille Rainbow, N.P Randle, Trevor Ian Robinson, Parul Sharma, Ghada T Shawli, James P Stewart, Neil Swainston, Ecaterina Vamos, Joanne Watts, Mark Whitehead |
| EPI_ISL_499568 | Northumbria University / South Tees Hospitals NHS Foundation Trust / North Cumbria Integrated Care NHS Foundation Trust / North Tees and Hartlepool NHS Foundation Trust / Newcastle Hospitals NHS Foundation Trust | COVID-19 Genomics UK (COG-UK) Consortium | Darren L Smith,Andrew Nelson,Matthew Bashton,Greg R Young,Joshua Loh,John Allan,Mohammad A Tariq,Giles S Holt,Gary Black,Wen C Yew,Lynn Dover,Paul Baker,Steve Liggett,Sarah Essex,Jane Greenaway,Debra Padgett,Clive Graham,Garren Scott,Edward Barton,Emma Swindells,Brendan Payne,Jennifer Collins,Yusri Taha,Gary Eltringham |
| EPI_ISL_499569, EPI_ISL_499570, EPI_ISL_499571 | Liverpool Clinical Laboratories | COVID-19 Genomics UK (COG-UK) Consortium | Sam Haldenby, Anita Lucaci, Steve Paterson, Julian Hiscox, Alistair Darby, M Almsaud, A Alrezaihi, Muhannad Alruwaili, Stuart D Armstrong, Jones Benjamin, Eleanor G Bentley, Anu Chawla, Jordan J Clark, Angela Cowell, Richard Eccles, Isabel García-Dorival, Matthew Gemmell, Alessandro Gerada, PKF Gilmore, Richard Gregory, Ximeng Han, Catherine Hartley, Margaret Hughes, Miren Iturriza-Gomara, James Johnson, L Luu, Jenifer Manson, Charlotte Nelson, Elaine O'Toole, Cassie Olateju, Rebekah Penrice-Randal , Lucille Rainbow, N.P Randle, Trevor Ian Robinson, Parul Sharma, Ghada T Shawli, James P Stewart, Neil Swainston, Ecaterina Vamos, Joanne Watts, Mark Whitehead |
| EPI_ISL_499572 | Northumbria University / South Tees Hospitals NHS Foundation Trust / North Cumbria Integrated Care NHS Foundation Trust / North Tees and Hartlepool NHS Foundation Trust / Newcastle Hospitals NHS Foundation Trust | COVID-19 Genomics UK (COG-UK) Consortium | Darren L Smith,Andrew Nelson,Matthew Bashton,Greg R Young,Joshua Loh,John Allan,Mohammad A Tariq,Giles S Holt,Gary Black,Wen C Yew,Lynn Dover,Paul Baker,Steve Liggett,Sarah Essex,Jane Greenaway,Debra Padgett,Clive Graham,Garren Scott,Edward Barton,Emma Swindells,Brendan Payne,Jennifer Collins,Yusri Taha,Gary Eltringham |
| EPI_ISL_499573, EPI_ISL_499574, EPI_ISL_499575, EPI_ISL_499576, EPI_ISL_499577, EPI_ISL_499578, EPI_ISL_499579, EPI_ISL_499580, EPI_ISL_499581, EPI_ISL_499582, EPI_ISL_499583, EPI_ISL_499584, EPI_ISL_499585, EPI_ISL_499586 | | | |
| see above | Liverpool Clinical Laboratories | COVID-19 Genomics UK (COG-UK) Consortium | Sam Haldenby, Anita Lucaci, Steve Paterson, Julian Hiscox, Alistair Darby, M Almsaud, A Alrezaihi, Muhannad Alruwaili, Stuart D Armstrong, Jones Benjamin, Eleanor G Bentley, Anu Chawla, Jordan J Clark, Angela Cowell, Richard Eccles, Isabel García-Dorival, Matthew Gemmell, Alessandro Gerada, PKF Gilmore, Richard Gregory, Ximeng Han, Catherine Hartley, Margaret Hughes, Miren Iturriza-Gomara, James Johnson, L Luu, Jenifer Manson, Charlotte Nelson, Elaine O'Toole, Cassie Olateju, Rebekah Penrice-Randal , Lucille Rainbow, N.P Randle, Trevor Ian Robinson, Parul Sharma, Ghada T Shawli, James P Stewart, Neil Swainston, Ecaterina Vamos, Joanne Watts, Mark Whitehead |
| EPI_ISL_499587 | Northumbria University / South Tees Hospitals NHS Foundation Trust / North Cumbria Integrated Care NHS Foundation Trust / North Tees and Hartlepool NHS Foundation Trust / Newcastle Hospitals NHS Foundation Trust | COVID-19 Genomics UK (COG-UK) Consortium | Darren L Smith,Andrew Nelson,Matthew Bashton,Greg R Young,Joshua Loh,John Allan,Mohammad A Tariq,Giles S Holt,Gary Black,Wen C Yew,Lynn Dover,Paul Baker,Steve Liggett,Sarah Essex,Jane Greenaway,Debra Padgett,Clive Graham,Garren Scott,Edward Barton,Emma Swindells,Brendan Payne,Jennifer Collins,Yusri Taha,Gary Eltringham |
| EPI_ISL_499588 | Quadram Institute Bioscience | COVID-19 Genomics UK (COG-UK) Consortium | Dave J. Baker, Gemma L. Kay, Alp Aydin, Thanh Le-Viet, Steven Rudder, Ana P. Tedim, Anastasia Kolyva, Maria Diaz, Leonardo de Oliveira Martins, Nabil-Fareed Alikhan, Lizzie Meadows, Rachael Stanley, Ngozi Elumogo, Muhammed Yasir, Nicholas M. Thomson, Alexander J Trotter, Rachel Gilroy, Samuel Bloomfield, Claire Stuart, Andrew Bell, Reenesh Prakash, Samir Dervisevic, Alison E. Mather, John Wain, Mark Webber, Andrew J. Page, Justin O'Grady |
| EPI_ISL_499589, EPI_ISL_499590 | Liverpool Clinical Laboratories | COVID-19 Genomics UK (COG-UK) Consortium | Sam Haldenby, Anita Lucaci, Steve Paterson, Julian Hiscox, Alistair Darby, M Almsaud, A Alrezaihi, Muhannad Alruwaili, Stuart D Armstrong, Jones Benjamin, Eleanor G Bentley, Anu Chawla, Jordan J Clark, Angela Cowell, Richard Eccles, Isabel García-Dorival, Matthew Gemmell, Alessandro Gerada, PKF Gilmore, Richard Gregory, Ximeng Han, Catherine Hartley, Margaret Hughes, Miren Iturriza-Gomara, James Johnson, L Luu, Jenifer Manson, Charlotte Nelson, Elaine O'Toole, Cassie Olateju, Rebekah Penrice-Randal , Lucille Rainbow, N.P Randle, Trevor Ian Robinson, Parul Sharma, Ghada T Shawli, James P Stewart, Neil Swainston, Ecaterina Vamos, Joanne Watts, Mark Whitehead |
| EPI_ISL_499591, EPI_ISL_499592 | Quadram Institute Bioscience | COVID-19 Genomics UK (COG-UK) Consortium | Dave J. Baker, Gemma L. Kay, Alp Aydin, Thanh Le-Viet, Steven Rudder, Ana P. Tedim, Anastasia Kolyva, Maria Diaz, Leonardo de Oliveira Martins, Nabil-Fareed Alikhan, Lizzie Meadows, Rachael Stanley, Ngozi Elumogo, Muhammed Yasir, Nicholas M. Thomson, Alexander J Trotter, Rachel Gilroy, Samuel Bloomfield, Claire Stuart, Andrew Bell, Reenesh Prakash, Samir Dervisevic, Alison E. Mather, John Wain, Mark Webber, Andrew J. Page, Justin O'Grady |
| EPI_ISL_499593, EPI_ISL_499594 | Liverpool Clinical Laboratories | COVID-19 Genomics UK (COG-UK) Consortium | Sam Haldenby, Anita Lucaci, Steve Paterson, Julian Hiscox, Alistair Darby, M Almsaud, A Alrezaihi, Muhannad Alruwaili, Stuart D Armstrong, Jones Benjamin, Eleanor G Bentley, Anu Chawla, Jordan J Clark, Angela Cowell, Richard Eccles, Isabel García-Dorival, Matthew Gemmell, Alessandro Gerada, PKF Gilmore, Richard Gregory, Ximeng Han, Catherine Hartley, Margaret Hughes, Miren Iturriza-Gomara, James Johnson, L Luu, Jenifer Manson, Charlotte Nelson, Elaine O'Toole, Cassie Olateju, Rebekah Penrice-Randal , Lucille Rainbow, N.P Randle, Trevor Ian Robinson, Parul Sharma, Ghada T Shawli, James P Stewart, Neil Swainston, Ecaterina Vamos, Joanne Watts, Mark Whitehead |
| EPI_ISL_499595, EPI_ISL_499596, EPI_ISL_499597, EPI_ISL_499598 | Northumbria University / South Tees Hospitals NHS Foundation Trust / North Cumbria Integrated Care NHS Foundation Trust / North Tees and Hartlepool NHS Foundation Trust / Newcastle Hospitals NHS Foundation Trust | COVID-19 Genomics UK (COG-UK) Consortium | Darren L Smith,Andrew Nelson,Matthew Bashton,Greg R Young,Joshua Loh,John Allan,Mohammad A Tariq,Giles S Holt,Gary Black,Wen C Yew,Lynn Dover,Paul Baker,Steve Liggett,Sarah Essex,Jane Greenaway,Debra Padgett,Clive Graham,Garren Scott,Edward Barton,Emma Swindells,Brendan Payne,Jennifer Collins,Yusri Taha,Gary Eltringham |
| EPI_ISL_499599 | Quadram Institute Bioscience | COVID-19 Genomics UK (COG-UK) Consortium | Dave J. Baker, Gemma L. Kay, Alp Aydin, Thanh Le-Viet, Steven Rudder, Ana P. Tedim, Anastasia Kolyva, Maria Diaz, Leonardo de Oliveira Martins, Nabil-Fareed Alikhan, Lizzie Meadows, Rachael Stanley, Ngozi Elumogo, Muhammed Yasir, Nicholas M. Thomson, Alexander J Trotter, Rachel Gilroy, Samuel Bloomfield, Claire Stuart, Andrew Bell, Reenesh Prakash, Samir Dervisevic, Alison E. Mather, John Wain, Mark Webber, Andrew J. Page, Justin O'Grady |
| EPI_ISL_499600 | Liverpool Clinical Laboratories | COVID-19 Genomics UK (COG-UK) Consortium | Sam Haldenby, Anita Lucaci, Steve Paterson, Julian Hiscox, Alistair Darby, M Almsaud, A Alrezaihi, Muhannad Alruwaili, Stuart D Armstrong, Jones Benjamin, Eleanor G Bentley, Anu Chawla, Jordan J Clark, Angela Cowell, Richard Eccles, Isabel García-Dorival, Matthew Gemmell, Alessandro Gerada, PKF Gilmore, Richard Gregory, Ximeng Han, Catherine Hartley, Margaret Hughes, Miren Iturriza-Gomara, James Johnson, L Luu, Jenifer Manson, Charlotte Nelson, Elaine O'Toole, Cassie Olateju, Rebekah Penrice-Randal , Lucille Rainbow, N.P Randle, Trevor Ian Robinson, Parul Sharma, Ghada T Shawli, James P Stewart, Neil Swainston, Ecaterina Vamos, Joanne Watts, Mark Whitehead |
| EPI_ISL_499601 | Quadram Institute Bioscience | COVID-19 Genomics UK (COG-UK) Consortium | Dave J. Baker, Gemma L. Kay, Alp Aydin, Thanh Le-Viet, Steven Rudder, Ana P. Tedim, Anastasia Kolyva, Maria Diaz, Leonardo de Oliveira Martins, Nabil-Fareed Alikhan, Lizzie Meadows, Rachael Stanley, Ngozi Elumogo, Muhammed Yasir, Nicholas M. Thomson, Alexander J Trotter, Rachel Gilroy, Samuel Bloomfield, Claire Stuart, Andrew Bell, Reenesh Prakash, Samir Dervisevic, Alison E. Mather, John Wain, Mark Webber, Andrew J. Page, Justin O'Grady |
| EPI_ISL_499602 | Northumbria University / South Tees Hospitals NHS Foundation Trust / North Cumbria Integrated Care NHS Foundation Trust / North Tees and Hartlepool NHS Foundation Trust / Newcastle Hospitals NHS Foundation Trust | COVID-19 Genomics UK (COG-UK) Consortium | Darren L Smith,Andrew Nelson,Matthew Bashton,Greg R Young,Joshua Loh,John Allan,Mohammad A Tariq,Giles S Holt,Gary Black,Wen C Yew,Lynn Dover,Paul Baker,Steve Liggett,Sarah Essex,Jane Greenaway,Debra Padgett,Clive Graham,Garren Scott,Edward Barton,Emma Swindells,Brendan Payne,Jennifer Collins,Yusri Taha,Gary Eltringham |
| EPI_ISL_499603 | Quadram Institute Bioscience | COVID-19 Genomics UK (COG-UK) Consortium | Dave J. Baker, Gemma L. Kay, Alp Aydin, Thanh Le-Viet, Steven Rudder, Ana P. Tedim, Anastasia Kolyva, Maria Diaz, Leonardo de Oliveira Martins, Nabil-Fareed Alikhan, Lizzie Meadows, Rachael Stanley, Ngozi Elumogo, Muhammed Yasir, Nicholas M. Thomson, Alexander J Trotter, Rachel Gilroy, Samuel Bloomfield, Claire Stuart, Andrew Bell, Reenesh Prakash, Samir Dervisevic, Alison E. Mather, John Wain, Mark Webber, Andrew J. Page, Justin O'Grady |
| EPI_ISL_499604, EPI_ISL_499605 | Liverpool Clinical Laboratories | COVID-19 Genomics UK (COG-UK) Consortium | Sam Haldenby, Anita Lucaci, Steve Paterson, Julian Hiscox, Alistair Darby, M Almsaud, A Alrezaihi, Muhannad Alruwaili, Stuart D Armstrong, Jones Benjamin, Eleanor G Bentley, Anu Chawla, Jordan J Clark, Angela Cowell, Richard Eccles, Isabel García-Dorival, Matthew Gemmell, Alessandro Gerada, PKF Gilmore, Richard Gregory, Ximeng Han, Catherine Hartley, Margaret Hughes, Miren Iturriza-Gomara, James Johnson, L Luu, Jenifer Manson, Charlotte Nelson, Elaine O'Toole, Cassie Olateju, Rebekah Penrice-Randal , Lucille Rainbow, N.P Randle, Trevor Ian Robinson, Parul Sharma, Ghada T Shawli, James P Stewart, Neil Swainston, Ecaterina Vamos, Joanne Watts, Mark Whitehead |
| EPI_ISL_499606 | Northumbria University / South Tees Hospitals NHS Foundation Trust / North Cumbria Integrated Care NHS Foundation Trust / North Tees and Hartlepool NHS Foundation Trust / Newcastle Hospitals NHS Foundation Trust | COVID-19 Genomics UK (COG-UK) Consortium | Darren L Smith,Andrew Nelson,Matthew Bashton,Greg R Young,Joshua Loh,John Allan,Mohammad A Tariq,Giles S Holt,Gary Black,Wen C Yew,Lynn Dover,Paul Baker,Steve Liggett,Sarah Essex,Jane Greenaway,Debra Padgett,Clive Graham,Garren Scott,Edward Barton,Emma Swindells,Brendan Payne,Jennifer Collins,Yusri Taha,Gary Eltringham |
| EPI_ISL_499607, EPI_ISL_499608 | Liverpool Clinical Laboratories | COVID-19 Genomics UK (COG-UK) Consortium | Sam Haldenby, Anita Lucaci, Steve Paterson, Julian Hiscox, Alistair Darby, M Almsaud, A Alrezaihi, Muhannad Alruwaili, Stuart D Armstrong, Jones Benjamin, Eleanor G Bentley, Anu Chawla, Jordan J Clark, Angela Cowell, Richard Eccles, Isabel García-Dorival, Matthew Gemmell, Alessandro Gerada, PKF Gilmore, Richard Gregory, Ximeng Han, Catherine Hartley, Margaret Hughes, Miren Iturriza-Gomara, James Johnson, L Luu, Jenifer Manson, Charlotte Nelson, Elaine O'Toole, Cassie Olateju, Rebekah Penrice-Randal , Lucille Rainbow, N.P Randle, Trevor Ian Robinson, Parul Sharma, Ghada T Shawli, James P Stewart, Neil Swainston, Ecaterina Vamos, Joanne Watts, Mark Whitehead |
| EPI_ISL_499609, EPI_ISL_499610 | Northumbria University / South Tees Hospitals NHS Foundation Trust / North Cumbria Integrated Care NHS Foundation Trust / North Tees and Hartlepool NHS Foundation Trust / Newcastle Hospitals NHS Foundation Trust | COVID-19 Genomics UK (COG-UK) Consortium | Darren L Smith,Andrew Nelson,Matthew Bashton,Greg R Young,Joshua Loh,John Allan,Mohammad A Tariq,Giles S Holt,Gary Black,Wen C Yew,Lynn Dover,Paul Baker,Steve Liggett,Sarah Essex,Jane Greenaway,Debra Padgett,Clive Graham,Garren Scott,Edward Barton,Emma Swindells,Brendan Payne,Jennifer Collins,Yusri Taha,Gary Eltringham |
| EPI_ISL_499611, EPI_ISL_499612, EPI_ISL_499613, EPI_ISL_499614, EPI_ISL_499615, EPI_ISL_499616, EPI_ISL_499617, EPI_ISL_499618, EPI_ISL_499619, EPI_ISL_499620, EPI_ISL_499621, EPI_ISL_499622 | | | |
| see above | Liverpool Clinical Laboratories | COVID-19 Genomics UK (COG-UK) Consortium | Sam Haldenby, Anita Lucaci, Steve Paterson, Julian Hiscox, Alistair Darby, M Almsaud, A Alrezaihi, Muhannad Alruwaili, Stuart D Armstrong, Jones Benjamin, Eleanor G Bentley, Anu Chawla, Jordan J Clark, Angela Cowell, Richard Eccles, Isabel García-Dorival, Matthew Gemmell, Alessandro Gerada, PKF Gilmore, Richard Gregory, Ximeng Han, Catherine Hartley, Margaret Hughes, Miren Iturriza-Gomara, James Johnson, L Luu, Jenifer Manson, Charlotte Nelson, Elaine O'Toole, Cassie Olateju, Rebekah Penrice-Randal , Lucille Rainbow, N.P Randle, Trevor Ian Robinson, Parul Sharma, Ghada T Shawli, James P Stewart, Neil Swainston, Ecaterina Vamos, Joanne Watts, Mark Whitehead |
| EPI_ISL_499623 | Quadram Institute Bioscience | COVID-19 Genomics UK (COG-UK) Consortium | Dave J. Baker, Gemma L. Kay, Alp Aydin, Thanh Le-Viet, Steven Rudder, Ana P. Tedim, Anastasia Kolyva, Maria Diaz, Leonardo de Oliveira Martins, Nabil-Fareed Alikhan, Lizzie Meadows, Rachael Stanley, Ngozi Elumogo, Muhammed Yasir, Nicholas M. Thomson, Alexander J Trotter, Rachel Gilroy, Samuel Bloomfield, Claire Stuart, Andrew Bell, Reenesh Prakash, Samir Dervisevic, Alison E. Mather, John Wain, Mark Webber, Andrew J. Page, Justin O'Grady |
| EPI_ISL_499624, EPI_ISL_499625 | Liverpool Clinical Laboratories | COVID-19 Genomics UK (COG-UK) Consortium | Sam Haldenby, Anita Lucaci, Steve Paterson, Julian Hiscox, Alistair Darby, M Almsaud, A Alrezaihi, Muhannad Alruwaili, Stuart D Armstrong, Jones Benjamin, Eleanor G Bentley, Anu Chawla, Jordan J Clark, Angela Cowell, Richard Eccles, Isabel García-Dorival, Matthew Gemmell, Alessandro Gerada, PKF Gilmore, Richard Gregory, Ximeng Han, Catherine Hartley, Margaret Hughes, Miren Iturriza-Gomara, James Johnson, L Luu, Jenifer Manson, Charlotte Nelson, Elaine O'Toole, Cassie Olateju, Rebekah Penrice-Randal , Lucille Rainbow, N.P Randle, Trevor Ian Robinson, Parul Sharma, Ghada T Shawli, James P Stewart, Neil Swainston, Ecaterina Vamos, Joanne Watts, Mark Whitehead |
| EPI_ISL_499626 | Northumbria University / South Tees Hospitals NHS Foundation Trust / North Cumbria Integrated Care NHS Foundation Trust / North Tees and Hartlepool NHS Foundation Trust / Newcastle Hospitals NHS Foundation Trust | COVID-19 Genomics UK (COG-UK) Consortium | Darren L Smith,Andrew Nelson,Matthew Bashton,Greg R Young,Joshua Loh,John Allan,Mohammad A Tariq,Giles S Holt,Gary Black,Wen C Yew,Lynn Dover,Paul Baker,Steve Liggett,Sarah Essex,Jane Greenaway,Debra Padgett,Clive Graham,Garren Scott,Edward Barton,Emma Swindells,Brendan Payne,Jennifer Collins,Yusri Taha,Gary Eltringham |
| EPI_ISL_499627, EPI_ISL_499628, EPI_ISL_499629, | Liverpool Clinical Laboratories | COVID-19 Genomics UK (COG-UK) Consortium | Sam Haldenby, Anita Lucaci, Steve Paterson, Julian Hiscox, Alistair Darby, M Almsaud, A Alrezaihi, Muhannad Alruwaili, Stuart D Armstrong, Jones Benjamin, Eleanor G Bentley, Anu Chawla, Jordan J Clark, Angela |

| | | | |
|---|---|---|---|
| EPI_ISL_499630 | | | Cowell, Richard Eccles, Isabel García-Dorival, Matthew Gemmell, Alessandro Gerada, PKF Gilmore, Richard Gregory, Ximeng Han, Catherine Hartley, Margaret Hughes, Miren Iturriza-Gomara, James Johnson, L Luu, Jenifer Manson, Charlotte Nelson, Elaine O'Toole, Cassie Olateju, Rebekah Penrice-Randal , Lucille Rainbow, N.P Randle, Trevor Ian Robinson, Parul Sharma, Ghada T Shawli, James P Stewart, Neil Swainston, Ecaterina Vamos, Joanne Watts, Mark Whitehead |
| EPI_ISL_499631 | Quadram Institute Bioscience | COVID-19 Genomics UK (COG-UK) Consortium | Dave J. Baker, Gemma L. Kay, Alp Aydin, Thanh Le-Viet, Steven Rudder, Ana P. Tedim, Anastasia Kolyva, Maria Diaz, Leonardo de Oliveira Martins, Nabil-Fareed Alikhan, Lizzie Meadows, Rachael Stanley, Ngozi Elumogo, Muhammed Yasir, Nicholas M. Thomson, Alexander J Trotter, Rachel Gilroy, Samuel Bloomfield, Claire Stuart, Andrew Bell, Reenesh Prakash, Samir Dervisevic, Alison E. Mather, John Wain, Mark Webber, Andrew J. Page, Justin O'Grady |
| EPI_ISL_499632, EPI_ISL_499633, EPI_ISL_499634, EPI_ISL_499635, EPI_ISL_499636, EPI_ISL_499637, EPI_ISL_499638, EPI_ISL_499639, EPI_ISL_499640, EPI_ISL_499641, EPI_ISL_499642, EPI_ISL_499643, EPI_ISL_499644, EPI_ISL_499645, EPI_ISL_499646, EPI_ISL_499647, EPI_ISL_499648, EPI_ISL_499649, EPI_ISL_499650, EPI_ISL_499651, EPI_ISL_499652, EPI_ISL_499653, EPI_ISL_499654, EPI_ISL_499655, EPI_ISL_499656, EPI_ISL_499657, EPI_ISL_499658, EPI_ISL_499659, EPI_ISL_499660, EPI_ISL_499661, EPI_ISL_499662, EPI_ISL_499663, EPI_ISL_499664, EPI_ISL_499665, EPI_ISL_499666, EPI_ISL_499667, EPI_ISL_499668, EPI_ISL_499669, EPI_ISL_499670, EPI_ISL_499671, EPI_ISL_499672, EPI_ISL_499673, EPI_ISL_499674, EPI_ISL_499675, EPI_ISL_499676, EPI_ISL_499677, EPI_ISL_499678, EPI_ISL_499679, EPI_ISL_499680, EPI_ISL_499681, EPI_ISL_499682, EPI_ISL_499683, EPI_ISL_499684, EPI_ISL_499685, EPI_ISL_499686, EPI_ISL_499687, EPI_ISL_499688, EPI_ISL_499689, EPI_ISL_499690, EPI_ISL_499691, EPI_ISL_499692, EPI_ISL_499693, EPI_ISL_499694, EPI_ISL_499695, EPI_ISL_499696, EPI_ISL_499697, EPI_ISL_499698, EPI_ISL_499699, EPI_ISL_499700, EPI_ISL_499701, EPI_ISL_499702, EPI_ISL_499703, EPI_ISL_499704, EPI_ISL_499705, EPI_ISL_499706, EPI_ISL_499707, EPI_ISL_499708, EPI_ISL_499709, EPI_ISL_499710, EPI_ISL_499711, EPI_ISL_499712, EPI_ISL_499713, EPI_ISL_499714, EPI_ISL_499715, EPI_ISL_499716, EPI_ISL_499717, EPI_ISL_499718, EPI_ISL_499719, EPI_ISL_499720, EPI_ISL_499721, EPI_ISL_499722, EPI_ISL_499723, EPI_ISL_499724, EPI_ISL_499725, EPI_ISL_499726, EPI_ISL_499727, EPI_ISL_499728, EPI_ISL_499729, EPI_ISL_499730, EPI_ISL_499731, EPI_ISL_499732, EPI_ISL_499733, EPI_ISL_499734, EPI_ISL_499735, EPI_ISL_499736, EPI_ISL_499737, EPI_ISL_499738, EPI_ISL_499739, EPI_ISL_499740, EPI_ISL_499741, EPI_ISL_499742, EPI_ISL_499743, EPI_ISL_499744, EPI_ISL_499745, EPI_ISL_499746, EPI_ISL_499747, EPI_ISL_499748, EPI_ISL_499749, EPI_ISL_499750, EPI_ISL_499751, EPI_ISL_499752, EPI_ISL_499753, EPI_ISL_499754, EPI_ISL_499755, EPI_ISL_499756, EPI_ISL_499757, EPI_ISL_499758, EPI_ISL_499759, EPI_ISL_499760, EPI_ISL_499761, EPI_ISL_499762, EPI_ISL_499763, EPI_ISL_499764, EPI_ISL_499765, EPI_ISL_499766, EPI_ISL_499767, EPI_ISL_499768, EPI_ISL_499769, EPI_ISL_499770, EPI_ISL_499771, EPI_ISL_499772, EPI_ISL_499773, EPI_ISL_499774, EPI_ISL_499775, EPI_ISL_499776 | | | |
| see above | Liverpool Clinical Laboratories | COVID-19 Genomics UK (COG-UK) Consortium | Sam Haldenby, Anita Lucaci, Steve Paterson, Julian Hiscox, Alistair Darby, M Almsaud, A Alrezaihi, Muhannad Alruwaili, Stuart D Armstrong, Jones Benjamin, Eleanor G Bentley, Anu Chawla, Jordan J Clark, Angela Cowell, Richard Eccles, Isabel García-Dorival, Matthew Gemmell, Alessandro Gerada, PKF Gilmore, Richard Gregory, Ximeng Han, Catherine Hartley, Margaret Hughes, Miren Iturriza-Gomara, James Johnson, L Luu, Jenifer Manson, Charlotte Nelson, Elaine O'Toole, Cassie Olateju, Rebekah Penrice-Randal , Lucille Rainbow, N.P Randle, Trevor Ian Robinson, Parul Sharma, Ghada T Shawli, James P Stewart, Neil Swainston, Ecaterina Vamos, Joanne Watts, Mark Whitehead |
| EPI_ISL_499777, EPI_ISL_499778, EPI_ISL_499779, EPI_ISL_499780, EPI_ISL_499781, EPI_ISL_499782, EPI_ISL_499783, EPI_ISL_499784, EPI_ISL_499785, EPI_ISL_499786, EPI_ISL_499787, EPI_ISL_499788, EPI_ISL_499789, EPI_ISL_499790, EPI_ISL_499791, EPI_ISL_499792, EPI_ISL_499793, EPI_ISL_499794, EPI_ISL_499795, EPI_ISL_499796, EPI_ISL_499797, EPI_ISL_499798, EPI_ISL_499799, EPI_ISL_499800, EPI_ISL_499801, EPI_ISL_499802, EPI_ISL_499803, EPI_ISL_499804, EPI_ISL_499805, EPI_ISL_499806, EPI_ISL_499807 | | | |
| see above | Northumbria University / South Tees Hospitals NHS Foundation Trust / North Cumbria Integrated Care NHS Foundation Trust / North Tees and Hartlepool NHS Foundation Trust / Newcastle Hospitals NHS Foundation Trust | COVID-19 Genomics UK (COG-UK) Consortium | Darren L Smith,Andrew Nelson,Matthew Bashton,Greg R Young,Joshua Loh,John Allan,Mohammad A Tariq,Giles S Holt,Gary Black,Wen C Yew,Lynn Dover,Paul Baker,Steve Liggett,Sarah Essex,Jane Greenaway,Debra Padgett,Clive Graham,Garren Scott,Edward Barton,Emma Swindells,Brendan Payne,Jennifer Collins,Yusri Taha,Gary Eltringham |
| EPI_ISL_499808, EPI_ISL_499809 | Queens Medical Centre, Clinical Microbiology Department / DeepSeq Nottingham | COVID-19 Genomics UK (COG-UK) Consortium | Gemma Clark, Wendy Smith, Manjinder Khakh, Vicki M Fleming, Michelle M Lister, Hannah Howson-Wells, Jonathan Ball, Patrick McClure, Joseph Chappell, Theocharis Tsoleridis, Nadine Holmes, Matthew Carlisle, Christopher Moore, Fei Sang, Johnny Debebe, Victoria Wright, Matthew Loose |
| EPI_ISL_499810 | University Hospitals Of Leicester NHS Trust and DeepSeq Nottingham | COVID-19 Genomics UK (COG-UK) Consortium | Christopher Holmes, Paul Bird, Thomas Helmer, Karlie Fallon, Julian Tang, Jonathan Ball, Patrick McClure, Joeseph Chappell, Nadine Holmes, Matthew Carlisle, Christopher Moore, Fei Sang, Johnny Debebe, Victoria Wright, Matthew Loose |
| EPI_ISL_499811, EPI_ISL_499812, EPI_ISL_499813, EPI_ISL_499814, EPI_ISL_499815, EPI_ISL_499816, EPI_ISL_499817, EPI_ISL_499818, EPI_ISL_499819, EPI_ISL_499820, EPI_ISL_499821, EPI_ISL_499822, EPI_ISL_499823, EPI_ISL_499824, EPI_ISL_499825, EPI_ISL_499826, EPI_ISL_499827 | | | |
| see above | Liverpool Clinical Laboratories | COVID-19 Genomics UK (COG-UK) Consortium | Sam Haldenby, Anita Lucaci, Steve Paterson, Julian Hiscox, Alistair Darby, M Almsaud, A Alrezaihi, Muhannad Alruwaili, Stuart D Armstrong, Jones Benjamin, Eleanor G Bentley, Anu Chawla, Jordan J Clark, Angela Cowell, Richard Eccles, Isabel García-Dorival, Matthew Gemmell, Alessandro Gerada, PKF Gilmore, Richard Gregory, Ximeng Han, Catherine Hartley, Margaret Hughes, Miren Iturriza-Gomara, James Johnson, L Luu, Jenifer Manson, Charlotte Nelson, Elaine O'Toole, Cassie Olateju, Rebekah Penrice-Randal , Lucille Rainbow, N.P Randle, Trevor Ian Robinson, Parul Sharma, Ghada T Shawli, James P Stewart, Neil Swainston, Ecaterina Vamos, Joanne Watts, Mark Whitehead |
| EPI_ISL_499828, EPI_ISL_499829 | University Hospitals Of Leicester NHS Trust and DeepSeq Nottingham | COVID-19 Genomics UK (COG-UK) Consortium | Christopher Holmes, Paul Bird, Thomas Helmer, Karlie Fallon, Julian Tang, Jonathan Ball, Patrick McClure, Joeseph Chappell, Nadine Holmes, Matthew Carlisle, Christopher Moore, Fei Sang, Johnny Debebe, Victoria Wright, Matthew Loose |
| EPI_ISL_499830 | University of Birmingham | COVID-19 Genomics UK (COG-UK) Consortium | Institute of Microbiology, University of Birmingham: Claire McMurray, Joanne Stockton, Samuel Nicholls, Radoslaw Poplawski, Will Rowe, Josh Quick, Nicholas Loman. University of Birmingham Testing Laboratory: Celina M Whalley, Andrew Bosworth, Charlotte Poxon, Kasun Wanigasooriya, Oliver Pickles, Mike Kidd, Alex Richter, Andrew D Beggs PHE Heartlands Lab: Husam Osman, Andrew Bosworth. Queen Elizabeth Hospital: Anna Casey |
| EPI_ISL_499831, EPI_ISL_499832, EPI_ISL_499833, EPI_ISL_499834, EPI_ISL_499835, EPI_ISL_499836, EPI_ISL_499837, EPI_ISL_499838 | Liverpool Clinical Laboratories | COVID-19 Genomics UK (COG-UK) Consortium | Sam Haldenby, Anita Lucaci, Steve Paterson, Julian Hiscox, Alistair Darby, M Almsaud, A Alrezaihi, Muhannad Alruwaili, Stuart D Armstrong, Jones Benjamin, Eleanor G Bentley, Anu Chawla, Jordan J Clark, Angela Cowell, Richard Eccles, Isabel García-Dorival, Matthew Gemmell, Alessandro Gerada, PKF Gilmore, Richard Gregory, Ximeng Han, Catherine Hartley, Margaret Hughes, Miren Iturriza-Gomara, James Johnson, L Luu, Jenifer Manson, Charlotte Nelson, Elaine O'Toole, Cassie Olateju, Rebekah Penrice-Randal , Lucille Rainbow, N.P Randle, Trevor Ian Robinson, Parul Sharma, Ghada T Shawli, James P Stewart, Neil Swainston, Ecaterina Vamos, Joanne Watts, Mark Whitehead |
| EPI_ISL_499839 | University of Birmingham | COVID-19 Genomics UK (COG-UK) Consortium | Institute of Microbiology, University of Birmingham: Claire McMurray, Joanne Stockton, Samuel Nicholls, Radoslaw Poplawski, Will Rowe, Josh Quick, Nicholas Loman. University of Birmingham Testing Laboratory: Celina M Whalley, Andrew Bosworth, Charlotte Poxon, Kasun Wanigasooriya, Oliver Pickles, Mike Kidd, Alex Richter, Andrew D Beggs PHE Heartlands Lab: Husam Osman, Andrew Bosworth. Queen Elizabeth Hospital: Anna Casey |
| EPI_ISL_499840, EPI_ISL_499841, EPI_ISL_499842, EPI_ISL_499843, EPI_ISL_499844, EPI_ISL_499845, EPI_ISL_499846, EPI_ISL_499847, EPI_ISL_499848, EPI_ISL_499849, EPI_ISL_499850 | | | |
| see above | Liverpool Clinical Laboratories | COVID-19 Genomics UK (COG-UK) Consortium | Sam Haldenby, Anita Lucaci, Steve Paterson, Julian Hiscox, Alistair Darby, M Almsaud, A Alrezaihi, Muhannad Alruwaili, Stuart D Armstrong, Jones Benjamin, Eleanor G Bentley, Anu Chawla, Jordan J Clark, Angela Cowell, Richard Eccles, Isabel García-Dorival, Matthew Gemmell, Alessandro Gerada, PKF Gilmore, Richard Gregory, Ximeng Han, Catherine Hartley, Margaret Hughes, Miren Iturriza-Gomara, James Johnson, L Luu, Jenifer Manson, Charlotte Nelson, Elaine O'Toole, Cassie Olateju, Rebekah Penrice-Randal , Lucille Rainbow, N.P Randle, Trevor Ian Robinson, Parul Sharma, Ghada T Shawli, James P Stewart, Neil Swainston, Ecaterina Vamos, Joanne Watts, Mark Whitehead |
| EPI_ISL_499851, EPI_ISL_499852 | University of Birmingham | COVID-19 Genomics UK (COG-UK) Consortium | Institute of Microbiology, University of Birmingham: Claire McMurray, Joanne Stockton, Samuel Nicholls, Radoslaw Poplawski, Will Rowe, Josh Quick, Nicholas Loman. University of Birmingham Testing Laboratory: Celina M Whalley, Andrew Bosworth, Charlotte Poxon, Kasun Wanigasooriya, Oliver Pickles, Mike Kidd, Alex Richter, Andrew D Beggs PHE Heartlands Lab: Husam Osman, Andrew Bosworth. Queen Elizabeth Hospital: Anna Casey |
| EPI_ISL_499853 | Department of Pathology, University of Cambridge | COVID-19 Genomics UK (COG-UK) Consortium | Luke W Meredith, M. Estée Török, Myra Hosmillo, William L. Hamilton, Martin D. Curran, Theresa Feltwell, Grant Hall, Anna Yakovleva, Fahad A Khokhar, Charlotte J. Houldcroft, Laura G Caller, Aminu S. Jahun, Sarah L. Caddy, Yasmin Chaudhry, Malte Pinckert, Ian Goodfellow |
| EPI_ISL_499854 | Liverpool Clinical Laboratories | COVID-19 Genomics UK (COG-UK) Consortium | Sam Haldenby, Anita Lucaci, Steve Paterson, Julian Hiscox, Alistair Darby, M Almsaud, A Alrezaihi, Muhannad Alruwaili, Stuart D Armstrong, Jones Benjamin, Eleanor G Bentley, Anu Chawla, Jordan J Clark, Angela Cowell, Richard Eccles, Isabel García-Dorival, Matthew Gemmell, Alessandro Gerada, PKF Gilmore, Richard Gregory, Ximeng Han, Catherine Hartley, Margaret Hughes, Miren Iturriza-Gomara, James Johnson, L Luu, Jenifer Manson, Charlotte Nelson, Elaine O'Toole, Cassie Olateju, Rebekah Penrice-Randal , Lucille Rainbow, N.P Randle, Trevor Ian Robinson, Parul Sharma, Ghada T Shawli, James P Stewart, Neil Swainston, Ecaterina Vamos, Joanne Watts, Mark Whitehead |
| EPI_ISL_499855 | Department of Pathology, University of Cambridge | COVID-19 Genomics UK (COG-UK) Consortium | Luke W Meredith, M. Estée Török, Myra Hosmillo, William L. Hamilton, Martin D. Curran, Theresa Feltwell, Grant Hall, Anna Yakovleva, Fahad A Khokhar, Charlotte J. Houldcroft, Laura G Caller, Aminu S. Jahun, Sarah L. Caddy, Yasmin Chaudhry, Malte Pinckert, Ian Goodfellow |
| EPI_ISL_499856, EPI_ISL_499857 | Liverpool Clinical Laboratories | COVID-19 Genomics UK (COG-UK) Consortium | Sam Haldenby, Anita Lucaci, Steve Paterson, Julian Hiscox, Alistair Darby, M Almsaud, A Alrezaihi, Muhannad Alruwaili, Stuart D Armstrong, Jones Benjamin, Eleanor G Bentley, Anu Chawla, Jordan J Clark, Angela Cowell, Richard Eccles, Isabel García-Dorival, Matthew Gemmell, Alessandro Gerada, PKF Gilmore, Richard Gregory, Ximeng Han, Catherine Hartley, Margaret Hughes, Miren Iturriza-Gomara, James Johnson, L Luu, Jenifer Manson, Charlotte Nelson, Elaine O'Toole, Cassie Olateju, Rebekah Penrice-Randal , Lucille Rainbow, N.P Randle, Trevor Ian Robinson, Parul Sharma, Ghada T Shawli, James P Stewart, Neil Swainston, Ecaterina Vamos, Joanne Watts, Mark Whitehead |
| EPI_ISL_499858 | University Hospitals Of Leicester NHS Trust and DeepSeq Nottingham | COVID-19 Genomics UK (COG-UK) Consortium | Christopher Holmes, Paul Bird, Thomas Helmer, Karlie Fallon, Julian Tang, Jonathan Ball, Patrick McClure, Joeseph Chappell, Nadine Holmes, Matthew Carlisle, Christopher Moore, Fei Sang, Johnny Debebe, Victoria Wright, Matthew Loose |
| EPI_ISL_499859, EPI_ISL_499860 | Liverpool Clinical Laboratories | COVID-19 Genomics UK (COG-UK) Consortium | Sam Haldenby, Anita Lucaci, Steve Paterson, Julian Hiscox, Alistair Darby, M Almsaud, A Alrezaihi, Muhannad Alruwaili, Stuart D Armstrong, Jones Benjamin, Eleanor G Bentley, Anu Chawla, Jordan J Clark, Angela Cowell, Richard Eccles, Isabel García-Dorival, Matthew Gemmell, Alessandro Gerada, PKF Gilmore, Richard Gregory, Ximeng Han, Catherine Hartley, Margaret Hughes, Miren Iturriza-Gomara, James Johnson, L Luu, Jenifer Manson, Charlotte Nelson, Elaine O'Toole, Cassie Olateju, Rebekah Penrice-Randal , Lucille Rainbow, N.P Randle, Trevor Ian Robinson, Parul Sharma, Ghada T Shawli, James P Stewart, Neil Swainston, Ecaterina Vamos, Joanne Watts, Mark Whitehead |
| EPI_ISL_499861 | University Hospitals Of Leicester NHS Trust and DeepSeq Nottingham | COVID-19 Genomics UK (COG-UK) Consortium | Christopher Holmes, Paul Bird, Thomas Helmer, Karlie Fallon, Julian Tang, Jonathan Ball, Patrick McClure, Joeseph Chappell, Nadine Holmes, Matthew Carlisle, Christopher Moore, Fei Sang, Johnny Debebe, Victoria Wright, Matthew Loose |
| EPI_ISL_499862, EPI_ISL_499863, EPI_ISL_499864, EPI_ISL_499865, EPI_ISL_499866, EPI_ISL_499867, EPI_ISL_499868 | Liverpool Clinical Laboratories | COVID-19 Genomics UK (COG-UK) Consortium | Sam Haldenby, Anita Lucaci, Steve Paterson, Julian Hiscox, Alistair Darby, M Almsaud, A Alrezaihi, Muhannad Alruwaili, Stuart D Armstrong, Jones Benjamin, Eleanor G Bentley, Anu Chawla, Jordan J Clark, Angela Cowell, Richard Eccles, Isabel García-Dorival, Matthew Gemmell, Alessandro Gerada, PKF Gilmore, Richard Gregory, Ximeng Han, Catherine Hartley, Margaret Hughes, Miren Iturriza-Gomara, James Johnson, L Luu, Jenifer Manson, Charlotte Nelson, Elaine O'Toole, Cassie Olateju, Rebekah Penrice-Randal , Lucille Rainbow, N.P Randle, Trevor Ian Robinson, Parul Sharma, Ghada T Shawli, James P Stewart, Neil Swainston, Ecaterina Vamos, Joanne Watts, Mark Whitehead |
| EPI_ISL_499869 | University Hospitals Of Leicester NHS Trust and DeepSeq Nottingham | COVID-19 Genomics UK (COG-UK) Consortium | Christopher Holmes, Paul Bird, Thomas Helmer, Karlie Fallon, Julian Tang, Jonathan Ball, Patrick McClure, Joeseph Chappell, Nadine Holmes, Matthew Carlisle, Christopher Moore, Fei Sang, Johnny Debebe, Victoria Wright, Matthew Loose |
| EPI_ISL_499870, EPI_ISL_499871, EPI_ISL_499872, EPI_ISL_499873, EPI_ISL_499874, EPI_ISL_499875, EPI_ISL_499876, EPI_ISL_499877, EPI_ISL_499878, EPI_ISL_499879, EPI_ISL_499880, EPI_ISL_499881, EPI_ISL_499882, EPI_ISL_499883, EPI_ISL_499884, EPI_ISL_499885, EPI_ISL_499886, EPI_ISL_499887, EPI_ISL_499888, EPI_ISL_499889, EPI_ISL_499890, EPI_ISL_499891, EPI_ISL_499892, EPI_ISL_499893 | | | |
| see above | Liverpool Clinical Laboratories | COVID-19 Genomics UK (COG-UK) Consortium | Sam Haldenby, Anita Lucaci, Steve Paterson, Julian Hiscox, Alistair Darby, M Almsaud, A Alrezaihi, Muhannad Alruwaili, Stuart D Armstrong, Jones Benjamin, Eleanor G Bentley, Anu Chawla, Jordan J Clark, Angela Cowell, Richard Eccles, Isabel García-Dorival, Matthew Gemmell, Alessandro Gerada, PKF Gilmore, Richard Gregory, Ximeng Han, Catherine Hartley, Margaret Hughes, Miren Iturriza-Gomara, James Johnson, L Luu, Jenifer Manson, Charlotte Nelson, Elaine O'Toole, Cassie Olateju, Rebekah Penrice-Randal , Lucille Rainbow, N.P Randle, Trevor Ian Robinson, Parul Sharma, Ghada T Shawli, James P Stewart, Neil Swainston, Ecaterina Vamos, Joanne Watts, Mark Whitehead |
| EPI_ISL_499894, EPI_ISL_499895 | University Hospitals Of Leicester NHS Trust and DeepSeq Nottingham | COVID-19 Genomics UK (COG-UK) Consortium | Christopher Holmes, Paul Bird, Thomas Helmer, Karlie Fallon, Julian Tang, Jonathan Ball, Patrick McClure, Joeseph Chappell, Nadine Holmes, Matthew Carlisle, Christopher Moore, Fei Sang, Johnny Debebe, Victoria Wright, Matthew Loose |
| EPI_ISL_499896 | Liverpool Clinical Laboratories | COVID-19 Genomics UK (COG-UK) Consortium | Sam Haldenby, Anita Lucaci, Steve Paterson, Julian Hiscox, Alistair Darby, M Almsaud, A Alrezaihi, Muhannad Alruwaili, Stuart D Armstrong, Jones Benjamin, Eleanor G Bentley, Anu Chawla, Jordan J Clark, Angela Cowell, Richard Eccles, Isabel García-Dorival, Matthew Gemmell, Alessandro Gerada, PKF Gilmore, Richard Gregory, Ximeng Han, Catherine Hartley, Margaret Hughes, Miren Iturriza-Gomara, James Johnson, L Luu, Jenifer Manson, Charlotte Nelson, Elaine O'Toole, Cassie Olateju, Rebekah Penrice-Randal , Lucille Rainbow, N.P Randle, Trevor Ian Robinson, Parul Sharma, Ghada T Shawli, James P Stewart, Neil Swainston, Ecaterina Vamos, Joanne Watts, Mark Whitehead |
| EPI_ISL_499897 | University Hospitals Of Leicester NHS Trust and DeepSeq Nottingham | COVID-19 Genomics UK (COG-UK) Consortium | Christopher Holmes, Paul Bird, Thomas Helmer, Karlie Fallon, Julian Tang, Jonathan Ball, Patrick McClure, Joeseph Chappell, Nadine Holmes, Matthew Carlisle, Christopher Moore, Fei Sang, Johnny Debebe, Victoria Wright, Matthew Loose |
| EPI_ISL_499898, EPI_ISL_499899, EPI_ISL_499900, EPI_ISL_499901, EPI_ISL_499902, EPI_ISL_499903, EPI_ISL_499904 | Liverpool Clinical Laboratories | COVID-19 Genomics UK (COG-UK) Consortium | Sam Haldenby, Anita Lucaci, Steve Paterson, Julian Hiscox, Alistair Darby, M Almsaud, A Alrezaihi, Muhannad Alruwaili, Stuart D Armstrong, Jones Benjamin, Eleanor G Bentley, Anu Chawla, Jordan J Clark, Angela Cowell, Richard Eccles, Isabel García-Dorival, Matthew Gemmell, Alessandro Gerada, PKF Gilmore, Richard Gregory, Ximeng Han, Catherine Hartley, Margaret Hughes, Miren Iturriza-Gomara, James Johnson, L Luu, Jenifer Manson, Charlotte Nelson, Elaine O'Toole, Cassie Olateju, Rebekah Penrice-Randal , Lucille Rainbow, N.P Randle, Trevor Ian Robinson, Parul Sharma, Ghada T Shawli, James P Stewart, Neil Swainston, Ecaterina Vamos, Joanne Watts, Mark Whitehead |
| EPI_ISL_499905 | Department of Pathology, University of Cambridge | COVID-19 Genomics UK (COG-UK) Consortium | Luke W Meredith, M. Estée Török, Myra Hosmillo, William L. Hamilton, Martin D. Curran, Theresa Feltwell, Grant Hall, Anna Yakovleva, Fahad A Khokhar, Charlotte J. Houldcroft, Laura G Caller, Aminu S. Jahun, |

| | | | |
|---|---|---|---|
| | | | Sarah L. Caddy, Yasmin Chaudhry, Malte Pinckert, Ian Goodfellow |
| EPI_ISL_499906, EPI_ISL_499907 | Liverpool Clinical Laboratories | COVID-19 Genomics UK (COG-UK) Consortium | Sam Haldenby, Anita Lucaci, Steve Paterson, Julian Hiscox, Alistair Darby, M Almsaud, A Alrezaihi, Muhannad Alruwaili, Stuart D Armstrong, Jones Benjamin, Eleanor G Bentley, Anu Chawla, Jordan J Clark, Angela Cowell, Richard Eccles, Isabel García-Dorival, Matthew Gemmell, Alessandro Gerada, PKF Gilmore, Richard Gregory, Ximeng Han, Catherine Hartley, Margaret Hughes, Miren Iturriza-Gomara, James Johnson, L Luu, Jenifer Manson, Charlotte Nelson, Elaine O’Toole, Cassie Olateju, Rebekah Penrice-Randal , Lucille Rainbow, N.P Randle, Trevor Ian Robinson, Parul Sharma, Ghada T Shawli, James P Stewart, Neil Swainston, Ecaterina Vamos, Joanne Watts, Mark Whitehead |
| EPI_ISL_499908 | Department of Pathology, University of Cambridge | COVID-19 Genomics UK (COG-UK) Consortium | Luke W Meredith, M. Estée Török, Myra Hosmillo, William L. Hamilton, Martin D. Curran, Theresa Feltwell, Grant Hall, Anna Yakovleva, Fahad A Khokhar, Charlotte J. Houldcroft, Laura G Caller, Aminu S. Jahun, Sarah L. Caddy, Yasmin Chaudhry, Malte Pinckert, Ian Goodfellow |
| EPI_ISL_499909 | Liverpool Clinical Laboratories | COVID-19 Genomics UK (COG-UK) Consortium | Sam Haldenby, Anita Lucaci, Steve Paterson, Julian Hiscox, Alistair Darby, M Almsaud, A Alrezaihi, Muhannad Alruwaili, Stuart D Armstrong, Jones Benjamin, Eleanor G Bentley, Anu Chawla, Jordan J Clark, Angela Cowell, Richard Eccles, Isabel García-Dorival, Matthew Gemmell, Alessandro Gerada, PKF Gilmore, Richard Gregory, Ximeng Han, Catherine Hartley, Margaret Hughes, Miren Iturriza-Gomara, James Johnson, L Luu, Jenifer Manson, Charlotte Nelson, Elaine O’Toole, Cassie Olateju, Rebekah Penrice-Randal , Lucille Rainbow, N.P Randle, Trevor Ian Robinson, Parul Sharma, Ghada T Shawli, James P Stewart, Neil Swainston, Ecaterina Vamos, Joanne Watts, Mark Whitehead |
| EPI_ISL_499910 | University Hospitals Of Leicester NHS Trust and DeepSeq Nottingham | COVID-19 Genomics UK (COG-UK) Consortium | Christopher Holmes, Paul Bird, Thomas Helmer, Karlie Fallon, Julian Tang, Jonathan Ball, Patrick McClure, Joeseph Chappell, Nadine Holmes, Matthew Carlisle, Christopher Moore, Fei Sang, Johnny Debebe, Victoria Wright, Matthew Loose |
| EPI_ISL_499911 | Liverpool Clinical Laboratories | COVID-19 Genomics UK (COG-UK) Consortium | Sam Haldenby, Anita Lucaci, Steve Paterson, Julian Hiscox, Alistair Darby, M Almsaud, A Alrezaihi, Muhannad Alruwaili, Stuart D Armstrong, Jones Benjamin, Eleanor G Bentley, Anu Chawla, Jordan J Clark, Angela Cowell, Richard Eccles, Isabel García-Dorival, Matthew Gemmell, Alessandro Gerada, PKF Gilmore, Richard Gregory, Ximeng Han, Catherine Hartley, Margaret Hughes, Miren Iturriza-Gomara, James Johnson, L Luu, Jenifer Manson, Charlotte Nelson, Elaine O’Toole, Cassie Olateju, Rebekah Penrice-Randal , Lucille Rainbow, N.P Randle, Trevor Ian Robinson, Parul Sharma, Ghada T Shawli, James P Stewart, Neil Swainston, Ecaterina Vamos, Joanne Watts, Mark Whitehead |
| EPI_ISL_499912, EPI_ISL_499913 | University Hospitals Of Leicester NHS Trust and DeepSeq Nottingham | COVID-19 Genomics UK (COG-UK) Consortium | Christopher Holmes, Paul Bird, Thomas Helmer, Karlie Fallon, Julian Tang, Jonathan Ball, Patrick McClure, Joeseph Chappell, Nadine Holmes, Matthew Carlisle, Christopher Moore, Fei Sang, Johnny Debebe, Victoria Wright, Matthew Loose |
| EPI_ISL_499914, EPI_ISL_499915, EPI_ISL_499916, EPI_ISL_499917, EPI_ISL_499918, EPI_ISL_499919, EPI_ISL_499920, EPI_ISL_499921, EPI_ISL_499922, EPI_ISL_499923, EPI_ISL_499924, EPI_ISL_499925, EPI_ISL_499926, EPI_ISL_499927, EPI_ISL_499928, EPI_ISL_499929, EPI_ISL_499930, EPI_ISL_499931, EPI_ISL_499932 | | | |
| see above | Liverpool Clinical Laboratories | COVID-19 Genomics UK (COG-UK) Consortium | Sam Haldenby, Anita Lucaci, Steve Paterson, Julian Hiscox, Alistair Darby, M Almsaud, A Alrezaihi, Muhannad Alruwaili, Stuart D Armstrong, Jones Benjamin, Eleanor G Bentley, Anu Chawla, Jordan J Clark, Angela Cowell, Richard Eccles, Isabel García-Dorival, Matthew Gemmell, Alessandro Gerada, PKF Gilmore, Richard Gregory, Ximeng Han, Catherine Hartley, Margaret Hughes, Miren Iturriza-Gomara, James Johnson, L Luu, Jenifer Manson, Charlotte Nelson, Elaine O’Toole, Cassie Olateju, Rebekah Penrice-Randal , Lucille Rainbow, N.P Randle, Trevor Ian Robinson, Parul Sharma, Ghada T Shawli, James P Stewart, Neil Swainston, Ecaterina Vamos, Joanne Watts, Mark Whitehead |
| EPI_ISL_499933 | University Hospitals Of Leicester NHS Trust and DeepSeq Nottingham | COVID-19 Genomics UK (COG-UK) Consortium | Christopher Holmes, Paul Bird, Thomas Helmer, Karlie Fallon, Julian Tang, Jonathan Ball, Patrick McClure, Joeseph Chappell, Nadine Holmes, Matthew Carlisle, Christopher Moore, Fei Sang, Johnny Debebe, Victoria Wright, Matthew Loose |
| EPI_ISL_499934, EPI_ISL_499935, EPI_ISL_499936, EPI_ISL_499937, EPI_ISL_499938, EPI_ISL_499939, EPI_ISL_499940, EPI_ISL_499941, EPI_ISL_499942, EPI_ISL_499943, EPI_ISL_499944, EPI_ISL_499945, EPI_ISL_499946 | | | |
| see above | Liverpool Clinical Laboratories | COVID-19 Genomics UK (COG-UK) Consortium | Sam Haldenby, Anita Lucaci, Steve Paterson, Julian Hiscox, Alistair Darby, M Almsaud, A Alrezaihi, Muhannad Alruwaili, Stuart D Armstrong, Jones Benjamin, Eleanor G Bentley, Anu Chawla, Jordan J Clark, Angela Cowell, Richard Eccles, Isabel García-Dorival, Matthew Gemmell, Alessandro Gerada, PKF Gilmore, Richard Gregory, Ximeng Han, Catherine Hartley, Margaret Hughes, Miren Iturriza-Gomara, James Johnson, L Luu, Jenifer Manson, Charlotte Nelson, Elaine O’Toole, Cassie Olateju, Rebekah Penrice-Randal , Lucille Rainbow, N.P Randle, Trevor Ian Robinson, Parul Sharma, Ghada T Shawli, James P Stewart, Neil Swainston, Ecaterina Vamos, Joanne Watts, Mark Whitehead |
| EPI_ISL_499947 | University of Birmingham | COVID-19 Genomics UK (COG-UK) Consortium | Institute of Microbiology, University of Birmingham: Claire McMurray, Joanne Stockton, Samuel Nicholls, Radoslaw Poplawski, Will Rowe, Josh Quick, Nicholas Loman. University of Birmingham Testing Laboratory: Celina M Whalley, Andrew Bosworth, Charlotte Poxon, Kasun Wanigasooriya, Oliver Pickles, Mike Kidd, Alex Richter, Andrew D Beggs PHE Heartlands Lab: Husam Osman, Andrew Bosworth. Queen Elizabeth Hospital: Anna Casey |
| EPI_ISL_499948, EPI_ISL_499949 | Liverpool Clinical Laboratories | COVID-19 Genomics UK (COG-UK) Consortium | Sam Haldenby, Anita Lucaci, Steve Paterson, Julian Hiscox, Alistair Darby, M Almsaud, A Alrezaihi, Muhannad Alruwaili, Stuart D Armstrong, Jones Benjamin, Eleanor G Bentley, Anu Chawla, Jordan J Clark, Angela Cowell, Richard Eccles, Isabel García-Dorival, Matthew Gemmell, Alessandro Gerada, PKF Gilmore, Richard Gregory, Ximeng Han, Catherine Hartley, Margaret Hughes, Miren Iturriza-Gomara, James Johnson, L Luu, Jenifer Manson, Charlotte Nelson, Elaine O’Toole, Cassie Olateju, Rebekah Penrice-Randal , Lucille Rainbow, N.P Randle, Trevor Ian Robinson, Parul Sharma, Ghada T Shawli, James P Stewart, Neil Swainston, Ecaterina Vamos, Joanne Watts, Mark Whitehead |
| EPI_ISL_499950 | University Hospitals Of Leicester NHS Trust and DeepSeq Nottingham | COVID-19 Genomics UK (COG-UK) Consortium | Christopher Holmes, Paul Bird, Thomas Helmer, Karlie Fallon, Julian Tang, Jonathan Ball, Patrick McClure, Joeseph Chappell, Nadine Holmes, Matthew Carlisle, Christopher Moore, Fei Sang, Johnny Debebe, Victoria Wright, Matthew Loose |
| EPI_ISL_499951, EPI_ISL_499952, EPI_ISL_499953, EPI_ISL_499954, EPI_ISL_499955, EPI_ISL_499956, EPI_ISL_499957, EPI_ISL_499958, EPI_ISL_499959, EPI_ISL_499960, EPI_ISL_499961, EPI_ISL_499962, EPI_ISL_499963, EPI_ISL_499964, EPI_ISL_499965 | | | |
| see above | Liverpool Clinical Laboratories | COVID-19 Genomics UK (COG-UK) Consortium | Sam Haldenby, Anita Lucaci, Steve Paterson, Julian Hiscox, Alistair Darby, M Almsaud, A Alrezaihi, Muhannad Alruwaili, Stuart D Armstrong, Jones Benjamin, Eleanor G Bentley, Anu Chawla, Jordan J Clark, Angela Cowell, Richard Eccles, Isabel García-Dorival, Matthew Gemmell, Alessandro Gerada, PKF Gilmore, Richard Gregory, Ximeng Han, Catherine Hartley, Margaret Hughes, Miren Iturriza-Gomara, James Johnson, L Luu, Jenifer Manson, Charlotte Nelson, Elaine O’Toole, Cassie Olateju, Rebekah Penrice-Randal , Lucille Rainbow, N.P Randle, Trevor Ian Robinson, Parul Sharma, Ghada T Shawli, James P Stewart, Neil Swainston, Ecaterina Vamos, Joanne Watts, Mark Whitehead |
| EPI_ISL_499966 | Department of Pathology, University of Cambridge | COVID-19 Genomics UK (COG-UK) Consortium | Luke W Meredith, M. Estée Török, Myra Hosmillo, William L. Hamilton, Martin D. Curran, Theresa Feltwell, Grant Hall, Anna Yakovleva, Fahad A Khokhar, Charlotte J. Houldcroft, Laura G Caller, Aminu S. Jahun, Sarah L. Caddy, Yasmin Chaudhry, Malte Pinckert, Ian Goodfellow |
| EPI_ISL_499967 | Liverpool Clinical Laboratories | COVID-19 Genomics UK (COG-UK) Consortium | Sam Haldenby, Anita Lucaci, Steve Paterson, Julian Hiscox, Alistair Darby, M Almsaud, A Alrezaihi, Muhannad Alruwaili, Stuart D Armstrong, Jones Benjamin, Eleanor G Bentley, Anu Chawla, Jordan J Clark, Angela Cowell, Richard Eccles, Isabel García-Dorival, Matthew Gemmell, Alessandro Gerada, PKF Gilmore, Richard Gregory, Ximeng Han, Catherine Hartley, Margaret Hughes, Miren Iturriza-Gomara, James Johnson, L Luu, Jenifer Manson, Charlotte Nelson, Elaine O’Toole, Cassie Olateju, Rebekah Penrice-Randal , Lucille Rainbow, N.P Randle, Trevor Ian Robinson, Parul Sharma, Ghada T Shawli, James P Stewart, Neil Swainston, Ecaterina Vamos, Joanne Watts, Mark Whitehead |
| EPI_ISL_499968, EPI_ISL_499969, EPI_ISL_499970, EPI_ISL_499971, EPI_ISL_499972, EPI_ISL_499973 | Department of Pathology, University of Cambridge | COVID-19 Genomics UK (COG-UK) Consortium | Luke W Meredith, M. Estée Török, Myra Hosmillo, William L. Hamilton, Martin D. Curran, Theresa Feltwell, Grant Hall, Anna Yakovleva, Fahad A Khokhar, Charlotte J. Houldcroft, Laura G Caller, Aminu S. Jahun, Sarah L. Caddy, Yasmin Chaudhry, Malte Pinckert, Ian Goodfellow |
| EPI_ISL_499974, EPI_ISL_499975, EPI_ISL_499976, EPI_ISL_499977, EPI_ISL_499978, EPI_ISL_499979, EPI_ISL_499980, EPI_ISL_499981, EPI_ISL_499982, EPI_ISL_499983 | University Hospitals Of Leicester NHS Trust and DeepSeq Nottingham | COVID-19 Genomics UK (COG-UK) Consortium | Christopher Holmes, Paul Bird, Thomas Helmer, Karlie Fallon, Julian Tang, Jonathan Ball, Patrick McClure, Joeseph Chappell, Nadine Holmes, Matthew Carlisle, Christopher Moore, Fei Sang, Johnny Debebe, Victoria Wright, Matthew Loose |
| EPI_ISL_499984, EPI_ISL_499985, EPI_ISL_499986, EPI_ISL_499987, EPI_ISL_499988, EPI_ISL_499989, EPI_ISL_499990, EPI_ISL_499991, EPI_ISL_499992, EPI_ISL_499993, EPI_ISL_499994, EPI_ISL_499995, EPI_ISL_499996, EPI_ISL_499997, EPI_ISL_499998, EPI_ISL_499999, EPI_ISL_500000, EPI_ISL_500001, EPI_ISL_500002, EPI_ISL_500003, EPI_ISL_500004, EPI_ISL_500005, EPI_ISL_500006, EPI_ISL_500007, EPI_ISL_500008, EPI_ISL_500009, EPI_ISL_500010, EPI_ISL_500011, EPI_ISL_500012, EPI_ISL_500013, EPI_ISL_500014, EPI_ISL_500015, EPI_ISL_500016, EPI_ISL_500017, EPI_ISL_500018, EPI_ISL_500019, EPI_ISL_500020, EPI_ISL_500021, EPI_ISL_500022, EPI_ISL_500023, EPI_ISL_500024, EPI_ISL_500025, EPI_ISL_500026, EPI_ISL_500027, EPI_ISL_500028, EPI_ISL_500029, EPI_ISL_500030, EPI_ISL_500031, EPI_ISL_500032, EPI_ISL_500033, EPI_ISL_500034, EPI_ISL_500035, EPI_ISL_500036, EPI_ISL_500037, EPI_ISL_500038, EPI_ISL_500039, EPI_ISL_500040, EPI_ISL_500041, EPI_ISL_500042, EPI_ISL_500043, EPI_ISL_500044, EPI_ISL_500045, EPI_ISL_500046, EPI_ISL_500047, EPI_ISL_500048, EPI_ISL_500049, EPI_ISL_500050, EPI_ISL_500051, EPI_ISL_500052, EPI_ISL_500053, EPI_ISL_500054, EPI_ISL_500055, EPI_ISL_500056, EPI_ISL_500057, EPI_ISL_500058, EPI_ISL_500059, EPI_ISL_500060, EPI_ISL_500061, EPI_ISL_500062, EPI_ISL_500063, EPI_ISL_500064, EPI_ISL_500065, EPI_ISL_500066, EPI_ISL_500067, EPI_ISL_500068, EPI_ISL_500069, EPI_ISL_500070, EPI_ISL_500071, EPI_ISL_500072, EPI_ISL_500073, EPI_ISL_500074, EPI_ISL_500075, EPI_ISL_500076, EPI_ISL_500077, EPI_ISL_500078, EPI_ISL_500079, EPI_ISL_500080, EPI_ISL_500081, EPI_ISL_500082, EPI_ISL_500083, EPI_ISL_500084, EPI_ISL_500085, EPI_ISL_500086, EPI_ISL_500087, EPI_ISL_500088, EPI_ISL_500089, EPI_ISL_500090, EPI_ISL_500091, EPI_ISL_500092, EPI_ISL_500093, EPI_ISL_500094, EPI_ISL_500095, EPI_ISL_500096, EPI_ISL_500097, EPI_ISL_500098, EPI_ISL_500099, EPI_ISL_500100, EPI_ISL_500101, EPI_ISL_500102, EPI_ISL_500103, EPI_ISL_500104, EPI_ISL_500105, EPI_ISL_500106, EPI_ISL_500107, EPI_ISL_500108, EPI_ISL_500109, EPI_ISL_500110, EPI_ISL_500111, EPI_ISL_500112, EPI_ISL_500113, EPI_ISL_500114, EPI_ISL_500115, EPI_ISL_500116, EPI_ISL_500117, EPI_ISL_500118, EPI_ISL_500119, EPI_ISL_500120, EPI_ISL_500121, EPI_ISL_500122, EPI_ISL_500123, EPI_ISL_500124, EPI_ISL_500125, EPI_ISL_500126, EPI_ISL_500127, EPI_ISL_500128, EPI_ISL_500129, EPI_ISL_500130, EPI_ISL_500131, EPI_ISL_500132, EPI_ISL_500133, EPI_ISL_500134, EPI_ISL_500135, EPI_ISL_500136, EPI_ISL_500137, EPI_ISL_500138, EPI_ISL_500139, EPI_ISL_500140, EPI_ISL_500141, EPI_ISL_500142, EPI_ISL_500143, EPI_ISL_500144, EPI_ISL_500145, EPI_ISL_500146, EPI_ISL_500147, EPI_ISL_500148, EPI_ISL_500149, EPI_ISL_500150, EPI_ISL_500151, EPI_ISL_500152, EPI_ISL_500153, EPI_ISL_500154, EPI_ISL_500155, EPI_ISL_500156 | | | |
| see above | Liverpool Clinical Laboratories | COVID-19 Genomics UK (COG-UK) Consortium | Sam Haldenby, Anita Lucaci, Steve Paterson, Julian Hiscox, Alistair Darby, M Almsaud, A Alrezaihi, Muhannad Alruwaili, Stuart D Armstrong, Jones Benjamin, Eleanor G Bentley, Anu Chawla, Jordan J Clark, Angela Cowell, Richard Eccles, Isabel García-Dorival, Matthew Gemmell, Alessandro Gerada, PKF Gilmore, Richard Gregory, Ximeng Han, Catherine Hartley, Margaret Hughes, Miren Iturriza-Gomara, James Johnson, L Luu, Jenifer Manson, Charlotte Nelson, Elaine O’Toole, Cassie Olateju, Rebekah Penrice-Randal , Lucille Rainbow, N.P Randle, Trevor Ian Robinson, Parul Sharma, Ghada T Shawli, James P Stewart, Neil Swainston, Ecaterina Vamos, Joanne Watts, Mark Whitehead |
| EPI_ISL_500157, EPI_ISL_500158, EPI_ISL_500159 | Complejo Hospitalario Universitario de Albacete | SeqCOVID-SPAIN consortium/IBV(CSIC) | Encarnacion Simarro Córdoba, Julia Lozano Serra, Lorena Robles Fonseca , Monica Parra Grandes, Caridad Sainz de Baranda Camino and SeqCOVID-SPAIN consortium |
| EPI_ISL_500160, EPI_ISL_500161 | Hospital Universitario Virgen de las Nieves de Granada-SAS | SeqCOVID-SPAIN consortium/IBV(CSIC) | Mercedes Pérez Ruiz, Sara Sanbonmatsu Gámez, Irene Pedrosa Corral, José M. Navarro-Marí and SeqCOVID-SPAIN consortium |
| EPI_ISL_500162 | Hospital Clínico Universitario de Santiago de Compostela | SeqCOVID-SPAIN consortium/IBV(CSIC) | José Javier Costa Alcalde, Antonio Aguilera Guirao, Mª Luisa Pérez del Molino Bernal, Amparo Coira Nieto, Gema Barbeito Castiñeiras, Rocio Trastoy Pena and SeqCOVID-SPAIN consortium |
| EPI_ISL_500163, EPI_ISL_500164, EPI_ISL_500165 | Hospital Universitario Virgen de las Nieves de Granada-SAS | SeqCOVID-SPAIN consortium/IBV(CSIC) | Mercedes Pérez Ruiz, Sara Sanbonmatsu Gámez, Irene Pedrosa Corral, José M. Navarro-Marí and SeqCOVID-SPAIN consortium |
| EPI_ISL_500166 | Complejo Hospitalario Universitario de Albacete | SeqCOVID-SPAIN consortium/IBV(CSIC) | Encarnacion Simarro Córdoba, Julia Lozano Serra, Lorena Robles Fonseca , Monica Parra Grandes, Caridad Sainz de Baranda Camino and SeqCOVID-SPAIN consortium |
| EPI_ISL_500167 | Hospital Clínico Universitario de Santiago de Compostela | SeqCOVID-SPAIN consortium/IBV(CSIC) | José Javier Costa Alcalde, Antonio Aguilera Guirao, Mª Luisa Pérez del Molino Bernal, Amparo Coira Nieto, Gema Barbeito Castiñeiras, Rocio Trastoy Pena and SeqCOVID-SPAIN consortium |
| EPI_ISL_500168, EPI_ISL_500169, EPI_ISL_500170, EPI_ISL_500171, EPI_ISL_500172, EPI_ISL_500173, EPI_ISL_500174, EPI_ISL_500175, EPI_ISL_500176, EPI_ISL_500177, EPI_ISL_500178, EPI_ISL_500179, EPI_ISL_500180, EPI_ISL_500181, EPI_ISL_500182, EPI_ISL_500183, EPI_ISL_500184, EPI_ISL_500185, EPI_ISL_500186, EPI_ISL_500187, EPI_ISL_500188, EPI_ISL_500189, EPI_ISL_500190, EPI_ISL_500191, EPI_ISL_500192, EPI_ISL_500193, EPI_ISL_500194, EPI_ISL_500195, EPI_ISL_500196, EPI_ISL_500197, EPI_ISL_500198, EPI_ISL_500199, EPI_ISL_500200, EPI_ISL_500201, EPI_ISL_500202, EPI_ISL_500203, EPI_ISL_500204, EPI_ISL_500205, EPI_ISL_500206 | | | |
| see above | Hospital Universitario Virgen de las Nieves de Granada-SAS | SeqCOVID-SPAIN consortium/IBV(CSIC) | Mercedes Pérez Ruiz, Sara Sanbonmatsu Gámez, Irene Pedrosa Corral, José M. Navarro-Marí and SeqCOVID-SPAIN consortium |
| EPI_ISL_500207 | Complejo Hospitalario Universitario de Albacete | SeqCOVID-SPAIN consortium/IBV(CSIC) | Encarnacion Simarro Córdoba, Julia Lozano Serra, Lorena Robles Fonseca , Monica Parra Grandes, Caridad Sainz de Baranda Camino and SeqCOVID-SPAIN consortium |
| EPI_ISL_500208 | Hospital Universitario Virgen de las Nieves de Granada-SAS | SeqCOVID-SPAIN consortium/IBV(CSIC) | Mercedes Pérez Ruiz, Sara Sanbonmatsu Gámez, Irene Pedrosa Corral, José M. Navarro-Marí and SeqCOVID-SPAIN consortium |
| EPI_ISL_500209 | Hospital Clínico Universitario de Santiago de Compostela | SeqCOVID-SPAIN consortium/IBV(CSIC) | José Javier Costa Alcalde, Antonio Aguilera Guirao, Mª Luisa Pérez del Molino Bernal, Amparo Coira Nieto, Gema Barbeito Castiñeiras, Rocio Trastoy Pena and SeqCOVID-SPAIN consortium |
| EPI_ISL_500210, EPI_ISL_500211, EPI_ISL_500212, EPI_ISL_500213, EPI_ISL_500214, EPI_ISL_500215, EPI_ISL_500216, EPI_ISL_500217 | Hospital Universitario Virgen de las Nieves de Granada-SAS | SeqCOVID-SPAIN consortium/IBV(CSIC) | Mercedes Pérez Ruiz, Sara Sanbonmatsu Gámez, Irene Pedrosa Corral, José M. Navarro-Marí and SeqCOVID-SPAIN consortium |
| EPI_ISL_500218 | Complejo Hospitalario Universitario de Albacete | SeqCOVID-SPAIN consortium/IBV(CSIC) | Encarnacion Simarro Córdoba, Julia Lozano Serra, Lorena Robles Fonseca , Monica Parra Grandes, Caridad Sainz de Baranda Camino and SeqCOVID-SPAIN consortium |
| EPI_ISL_500219 | Hospital Clínico Universitario de Santiago de Compostela | SeqCOVID-SPAIN consortium/IBV(CSIC) | José Javier Costa Alcalde, Antonio Aguilera Guirao, Mª Luisa Pérez del Molino Bernal, Amparo Coira Nieto, Gema Barbeito Castiñeiras, Rocio Trastoy Pena and SeqCOVID-SPAIN consortium |
| EPI_ISL_500220 | Hospital Universitario Virgen de las Nieves de Granada-SAS | SeqCOVID-SPAIN consortium/IBV(CSIC) | Mercedes Pérez Ruiz, Sara Sanbonmatsu Gámez, Irene Pedrosa Corral, José M. Navarro-Marí and SeqCOVID-SPAIN consortium |
| EPI_ISL_500221 | Hospital Clínico Universitario de Santiago de Compostela | SeqCOVID-SPAIN consortium/IBV(CSIC) | José Javier Costa Alcalde, Antonio Aguilera Guirao, Mª Luisa Pérez del Molino Bernal, Amparo Coira Nieto, Gema Barbeito Castiñeiras, Rocio Trastoy Pena and SeqCOVID-SPAIN consortium |
| EPI_ISL_500222, EPI_ISL_500223, EPI_ISL_500224, EPI_ISL_500225, EPI_ISL_500226, EPI_ISL_500227 | Hospital Universitario Virgen de las Nieves de Granada-SAS | SeqCOVID-SPAIN consortium/IBV(CSIC) | Mercedes Pérez Ruiz, Sara Sanbonmatsu Gámez, Irene Pedrosa Corral, José M. Navarro-Marí and SeqCOVID-SPAIN consortium |
| EPI_ISL_500228, EPI_ISL_500229, EPI_ISL_500230, EPI_ISL_500231, EPI_ISL_500232, EPI_ISL_500233, EPI_ISL_500234, EPI_ISL_500235, EPI_ISL_500236, EPI_ISL_500237, EPI_ISL_500238, EPI_ISL_500239, EPI_ISL_500240, EPI_ISL_500241, EPI_ISL_500242, EPI_ISL_500243, EPI_ISL_500244, EPI_ISL_500245, EPI_ISL_500246, EPI_ISL_500247, EPI_ISL_500248, EPI_ISL_500249, EPI_ISL_500250, EPI_ISL_500251, EPI_ISL_500252, EPI_ISL_500253, EPI_ISL_500254, EPI_ISL_500255, EPI_ISL_500256, EPI_ISL_500257, EPI_ISL_500258, EPI_ISL_500259, EPI_ISL_500260, EPI_ISL_500261, EPI_ISL_500262, EPI_ISL_500263, EPI_ISL_500264, EPI_ISL_500265, EPI_ISL_500266, EPI_ISL_500267, EPI_ISL_500268, EPI_ISL_500269, EPI_ISL_500270, EPI_ISL_500271, EPI_ISL_500272, EPI_ISL_500273, EPI_ISL_500274, EPI_ISL_500275, | | | |

| | | | |
|---|---|---|---|
| EPI_ISL_500276, EPI_ISL_500277, EPI_ISL_500278, EPI_ISL_500279, EPI_ISL_500280, EPI_ISL_500281, EPI_ISL_500282, EPI_ISL_500283, EPI_ISL_500284, EPI_ISL_500285, EPI_ISL_500286 | | | |
| see above | Hospital Universitario Araba. Vitoria-Gasteiz | SeqCOVID-SPAIN consortium/IBV(CSIC) | Silvia Hernáez Crespo, Carmen Gómez González, Amaia Aguirre Quiñonero, Marina Fernández Torres, Mª Rosario Almela Ferrer, Mª Concepción Lecaroz Agara, Andrés Canut Blasco and SeqCOVID-SPAIN consortium |
| EPI_ISL_500287, EPI_ISL_500288, EPI_ISL_500290, EPI_ISL_500291 | Servicio de Microbiología, Hospital Miguel Servet, Zaragoza | SeqCOVID-SPAIN consortium/IBV(CSIC) | Antonio Rezusta López, Alexander Tristancho Baró, Ana Milagro, Yolanda Gracia Grataloup, Nieves Martínez Cameo and SeqCOVID-SPAIN consortium |
| EPI_ISL_500292, EPI_ISL_500293, EPI_ISL_500294, EPI_ISL_500295, EPI_ISL_500296, EPI_ISL_500297, EPI_ISL_500298, EPI_ISL_500299, EPI_ISL_500300, EPI_ISL_500301, EPI_ISL_500302, EPI_ISL_500303, EPI_ISL_500304, EPI_ISL_500305, EPI_ISL_500306, EPI_ISL_500307, EPI_ISL_500308, EPI_ISL_500309, EPI_ISL_500310, EPI_ISL_500311, EPI_ISL_500312, EPI_ISL_500313, EPI_ISL_500314, EPI_ISL_500315, EPI_ISL_500316, EPI_ISL_500317, EPI_ISL_500318, EPI_ISL_500319, EPI_ISL_500320, EPI_ISL_500321, EPI_ISL_500322, EPI_ISL_500323, EPI_ISL_500324, EPI_ISL_500325, EPI_ISL_500326, EPI_ISL_500327, EPI_ISL_500328, EPI_ISL_500329, EPI_ISL_500330, EPI_ISL_500331, EPI_ISL_500332, EPI_ISL_500333, EPI_ISL_500334, EPI_ISL_500335 | | | |
| see above | Servicio de Microbiología. Hospital Universitario Donostia. OSI Donostialdea. Área de Enfermedades Infecciosas, Grupo de Infección Respiratoria y Resistencia Antimicrobiana. Instituto de Investigación Sanitaria Biodonostia | SeqCOVID-SPAIN consortium/IBV(CSIC) | Gustavo Cilla, Milagrosa Montes, Luis Piñeiro, Jose Maria Marimón and SeqCOVID-SPAIN consortium |
| EPI_ISL_500336, EPI_ISL_500337, EPI_ISL_500338, EPI_ISL_500339, EPI_ISL_500340, EPI_ISL_500341, EPI_ISL_500343, EPI_ISL_500344, EPI_ISL_500345, EPI_ISL_500346, EPI_ISL_500347, EPI_ISL_500349, EPI_ISL_500350, EPI_ISL_500351, EPI_ISL_500352, EPI_ISL_500353, EPI_ISL_500354, EPI_ISL_500355, EPI_ISL_500356, EPI_ISL_500357, EPI_ISL_500359, EPI_ISL_500360, EPI_ISL_500361, EPI_ISL_500362, EPI_ISL_500363, EPI_ISL_500364, EPI_ISL_500365, EPI_ISL_500366, EPI_ISL_500367, EPI_ISL_500368 | | | |
| see above | Servicio de Microbiología, Hospital Miguel Servet, Zaragoza | SeqCOVID-SPAIN consortium/IBV(CSIC) | Antonio Rezusta López, Alexander Tristancho Baró, Ana Milagro, Yolanda Gracia Grataloup, Nieves Martínez Cameo and SeqCOVID-SPAIN consortium |
| EPI_ISL_500369, EPI_ISL_500370, EPI_ISL_500371, EPI_ISL_500372, EPI_ISL_500374, EPI_ISL_500375, EPI_ISL_500376, EPI_ISL_500377, EPI_ISL_500378, EPI_ISL_500379, EPI_ISL_500380, EPI_ISL_500381, EPI_ISL_500383, EPI_ISL_500385, EPI_ISL_500386, EPI_ISL_500387, EPI_ISL_500388, EPI_ISL_500389, EPI_ISL_500390, EPI_ISL_500391, EPI_ISL_500392, EPI_ISL_500393, EPI_ISL_500394, EPI_ISL_500395, EPI_ISL_500396, EPI_ISL_500397, EPI_ISL_500398, EPI_ISL_500399, EPI_ISL_500400, EPI_ISL_500401, EPI_ISL_500402, EPI_ISL_500403, EPI_ISL_500404, EPI_ISL_500405, EPI_ISL_500406, EPI_ISL_500407, EPI_ISL_500408, EPI_ISL_500409, EPI_ISL_500410, EPI_ISL_500411, EPI_ISL_500412, EPI_ISL_500413, EPI_ISL_500414, EPI_ISL_500415, EPI_ISL_500416, EPI_ISL_500417, EPI_ISL_500418, EPI_ISL_500419, EPI_ISL_500420, EPI_ISL_500421, EPI_ISL_500422, EPI_ISL_500423, EPI_ISL_500424, EPI_ISL_500425, EPI_ISL_500426, EPI_ISL_500427, EPI_ISL_500428, EPI_ISL_500429, EPI_ISL_500430, EPI_ISL_500431, EPI_ISL_500432, EPI_ISL_500433, EPI_ISL_500434, EPI_ISL_500435, EPI_ISL_500436, EPI_ISL_500437, EPI_ISL_500438, EPI_ISL_500439, EPI_ISL_500440, EPI_ISL_500441, EPI_ISL_500442, EPI_ISL_500443, EPI_ISL_500444, EPI_ISL_500445, EPI_ISL_500446, EPI_ISL_500447, EPI_ISL_500448, EPI_ISL_500449, EPI_ISL_500450, EPI_ISL_500451, EPI_ISL_500452, EPI_ISL_500453, EPI_ISL_500454, EPI_ISL_500455, EPI_ISL_500456, EPI_ISL_500457, EPI_ISL_500458 | | | |
| see above | Centro de Investigación Biomédica de La Rioja - Hospital San Pedro Logroño | SeqCOVID-SPAIN consortium/IBV(CSIC) | María de Toro, José Manuel Azcona Gutiérrez, María Pilar Bea Escudero, Miriam Blasco Alberdi and SeqCOVID-SPAIN consortium |
| EPI_ISL_500459 | Quest Diagnostics | Quest Diagnostics | Rosenthal,S.H., Gerasimova,A., Kagan,R.M., Owen, R. AND Lacbawan, F. |
| EPI_ISL_500460, EPI_ISL_500461, EPI_ISL_500462, EPI_ISL_500463, EPI_ISL_500464, EPI_ISL_500465, EPI_ISL_500466, EPI_ISL_500467, EPI_ISL_500468, EPI_ISL_500469, EPI_ISL_500470, EPI_ISL_500471, EPI_ISL_500472, EPI_ISL_500473, EPI_ISL_500474, EPI_ISL_500475, EPI_ISL_500476, EPI_ISL_500477, EPI_ISL_500478, EPI_ISL_500479, EPI_ISL_500480, EPI_ISL_500481, EPI_ISL_500482, EPI_ISL_500483, EPI_ISL_500484, EPI_ISL_500485, EPI_ISL_500486 | | | |
| see above | LACEN/PE | WallauLab, Aggeu Magalhaes Institute | Marcelo Henrique dos Santos Paiva, Duschinka Ribeiro Duarte Guedes, Cássia Docena, Matheus Filgueira Bezerra, Filipe Zimmer Dezordi, Laís Ceschini Machado, Larissa Krokovsky, Elisama Helvecio, Alexandre Freitas da Silva, Antonio Mauro Rezende5, Sinval Pinto Brandão Filho7, Constância Flávia Junqueira Ayres, Gabriel Luz Wallau |
| EPI_ISL_500500, EPI_ISL_500501, EPI_ISL_500502 | University of Washington Virology Lab | University of Washington Virology Lab | Pavitra Roychoudhury, Hong Xie, Lasata Shrestha, Amin Addetia, Truong Nguyen, Victoria M Rachleff, Meei-Li Huang, Keith R Jerome, Alexander Greninger |
| EPI_ISL_500503, EPI_ISL_500504, EPI_ISL_500505, EPI_ISL_500506, EPI_ISL_500507, EPI_ISL_500508, EPI_ISL_500509, EPI_ISL_500510, EPI_ISL_500511, EPI_ISL_500512, EPI_ISL_500513, EPI_ISL_500514, EPI_ISL_500515, EPI_ISL_500516, EPI_ISL_500517, EPI_ISL_500518, EPI_ISL_500519, EPI_ISL_500520, EPI_ISL_500521, EPI_ISL_500522, EPI_ISL_500523, EPI_ISL_500524, EPI_ISL_500525, EPI_ISL_500526, EPI_ISL_500527, EPI_ISL_500528, EPI_ISL_500529, EPI_ISL_500530, EPI_ISL_500531, EPI_ISL_500532, EPI_ISL_500533, EPI_ISL_500534, EPI_ISL_500535, EPI_ISL_500536, EPI_ISL_500537, EPI_ISL_500538 | | | |
| see above | Mayo Clinic Laboratories | University of Washington Virology Lab | Pavitra Roychoudhury, Hong Xie, Lasata Shrestha, Amin Addetia, Truong Nguyen, Victoria M Rachleff, Meei-Li Huang, Keith R Jerome, Alexander Greninger |
| EPI_ISL_500539, EPI_ISL_500540, EPI_ISL_500541, EPI_ISL_500542, EPI_ISL_500543, EPI_ISL_500544, EPI_ISL_500545, EPI_ISL_500546, EPI_ISL_500547, EPI_ISL_500548, EPI_ISL_500549, EPI_ISL_500550, EPI_ISL_500551, EPI_ISL_500552, EPI_ISL_500553, EPI_ISL_500554, EPI_ISL_500555, EPI_ISL_500556, EPI_ISL_500557, EPI_ISL_500558, EPI_ISL_500559, EPI_ISL_500560, EPI_ISL_500561, EPI_ISL_500562, EPI_ISL_500563, EPI_ISL_500564, EPI_ISL_500565, EPI_ISL_500566, EPI_ISL_500567, EPI_ISL_500568, EPI_ISL_500569, EPI_ISL_500570, EPI_ISL_500571, EPI_ISL_500572 | | | |
| see above | Singapore General Hospital | Department of Microbiology | Nurdyana Abdul Rahman, Kun Lee Lim, Chenhao Li, Kian Sing Chan, Lynette Oon, Kern Rei Chng, Niranjan Nagarajan, Karrie Ko |
| EPI_ISL_500573, EPI_ISL_500574, EPI_ISL_500575, EPI_ISL_500576, EPI_ISL_500577, EPI_ISL_500578, EPI_ISL_500579, EPI_ISL_500580, EPI_ISL_500581, EPI_ISL_500582, EPI_ISL_500583, EPI_ISL_500584, EPI_ISL_500585, EPI_ISL_500586, EPI_ISL_500587, EPI_ISL_500588, EPI_ISL_500589, EPI_ISL_500590, EPI_ISL_500591, EPI_ISL_500592, EPI_ISL_500593, EPI_ISL_500594, EPI_ISL_500595 | | | |
| see above | National Virus Reference Laboratory | National Virus Reference Laboratory | Michael Carr, Gabriel Gonzalez, Jonathan Dean, Suzie Coughlan, Cillian F De Gascun |
| EPI_ISL_500596, EPI_ISL_500597, EPI_ISL_500598, EPI_ISL_500599, EPI_ISL_500600, EPI_ISL_500601, EPI_ISL_500602, EPI_ISL_500603, EPI_ISL_500604, EPI_ISL_500605, EPI_ISL_500606, EPI_ISL_500607, EPI_ISL_500608, EPI_ISL_500609, EPI_ISL_500610, EPI_ISL_500611, EPI_ISL_500612, EPI_ISL_500613, EPI_ISL_500614, EPI_ISL_500615, EPI_ISL_500616, EPI_ISL_500617, EPI_ISL_500618, EPI_ISL_500619, EPI_ISL_500620, EPI_ISL_500621, EPI_ISL_500622, EPI_ISL_500623, EPI_ISL_500624, EPI_ISL_500625, EPI_ISL_500626, EPI_ISL_500627, EPI_ISL_500628, EPI_ISL_500629, EPI_ISL_500630, EPI_ISL_500631, EPI_ISL_500632, EPI_ISL_500633, EPI_ISL_500634, EPI_ISL_500635, EPI_ISL_500636, EPI_ISL_500637, EPI_ISL_500638, EPI_ISL_500639, EPI_ISL_500640, EPI_ISL_500641, EPI_ISL_500642, EPI_ISL_500643, EPI_ISL_500644, EPI_ISL_500645, EPI_ISL_500646, EPI_ISL_500647, EPI_ISL_500648, EPI_ISL_500649, EPI_ISL_500650, EPI_ISL_500651, EPI_ISL_500652, EPI_ISL_500653, EPI_ISL_500654, EPI_ISL_500655, EPI_ISL_500656, EPI_ISL_500657, EPI_ISL_500658, EPI_ISL_500659, EPI_ISL_500660, EPI_ISL_500661, EPI_ISL_500662, EPI_ISL_500663, EPI_ISL_500664, EPI_ISL_500665, EPI_ISL_500666, EPI_ISL_500667, EPI_ISL_500668, EPI_ISL_500669, EPI_ISL_500670, EPI_ISL_500671, EPI_ISL_500672, EPI_ISL_500673, EPI_ISL_500674, EPI_ISL_500675, EPI_ISL_500676, EPI_ISL_500677, EPI_ISL_500678, EPI_ISL_500679, EPI_ISL_500680, EPI_ISL_500681, EPI_ISL_500682, EPI_ISL_500683, EPI_ISL_500684, EPI_ISL_500685, EPI_ISL_500686, EPI_ISL_500687, EPI_ISL_500688, EPI_ISL_500689, EPI_ISL_500690, EPI_ISL_500691, EPI_ISL_500692, EPI_ISL_500693, EPI_ISL_500694, EPI_ISL_500695, EPI_ISL_500696, EPI_ISL_500697, EPI_ISL_500698, EPI_ISL_500699, EPI_ISL_500700, EPI_ISL_500701, EPI_ISL_500702, EPI_ISL_500703, EPI_ISL_500704, EPI_ISL_500705, EPI_ISL_500706 | | | |
| see above | Area of Virology, Serology and Virology Division (SAViD), New South Wales Health Pathology Randwick | Area of Virology, Serology and Virology Division (SAViD), New South Wales Health Pathology Randwick | Rawlinson, W. |
| EPI_ISL_500707, EPI_ISL_500708, EPI_ISL_500709, EPI_ISL_500710, EPI_ISL_500711, EPI_ISL_500712, EPI_ISL_500713, EPI_ISL_500714, EPI_ISL_500715 | Respiratory Virus Unit, Microbiology Services Colindale, Public Health England | Respiratory Virus Unit, Microbiology Services Colindale, Public Health England | PHE Covid Sequencing Team |
| EPI_ISL_500716 | BSL3 Lab Pendik Veterinary Control Institute | Department of Medicinal Genetics, Bursa Uludağ University, Faculty of medicine By Sehime Gülsün Temel, Adem Alemdar, Kadir Yeşilbağ | Mustafa HASOKSUZ, Fahriye SARAC, Osman ERGANIS, Serdar UZAR, Hakan ENUL, Cumhur ADIAY, Ahmet SAIT, Orbay SAYI, Kadir YESILBAG, Oguz KARABEY |
| EPI_ISL_500717 | Area of Virology, Serology and Virology Division (SAViD), New South Wales Health Pathology Randwick | Area of Virology, Serology and Virology Division (SAViD), New South Wales Health Pathology Randwick | Rawlinson, W |
| EPI_ISL_500719 | Respiratory Virus Unit, Microbiology Services Colindale, Public Health England | Respiratory Virus Unit, Microbiology Services Colindale, Public Health England | PHE Covid Sequencing Team |
| EPI_ISL_500768, EPI_ISL_500769, EPI_ISL_500770, EPI_ISL_500771, EPI_ISL_500772, EPI_ISL_500773, EPI_ISL_500774 | Furst Medical Laboratory | Norwegian Institute of Public Health, Department of Virology | Kathrine Stene-Johansen, Kamilla Heddeland Instefjord, Hilde Elshaug, Rasmus Riis Kopperud, Karoline Bragstad, Olav Hungnes |
| EPI_ISL_500775 | Hospital of Southern Norway - Kristiansand, Department of Medical Microbiology | Norwegian Institute of Public Health, Department of Virology | Kathrine Stene-Johansen, Kamilla Heddeland Instefjord, Hilde Elshaug, Rasmus Riis Kopperud, Karoline Bragstad, Olav Hungnes |
| EPI_ISL_500776, EPI_ISL_500777, EPI_ISL_500778 | Foerde Hospital, Department of Microbiology | Norwegian Institute of Public Health, Department of Virology | Kathrine Stene-Johansen, Kamilla Heddeland Instefjord, Hilde Elshaug, Rasmus Riis Kopperud, Karoline Bragstad, Olav Hungnes |
| EPI_ISL_500779, EPI_ISL_500780, EPI_ISL_500781, EPI_ISL_500782, EPI_ISL_500783 | Akershus University Hospital, Department for Microbiology and Infectious Disease Control | Norwegian Institute of Public Health, Department of Virology | Kathrine Stene-Johansen, Kamilla Heddeland Instefjord, Hilde Elshaug, Rasmus Riis Kopperud, Karoline Bragstad, Olav Hungnes |
| EPI_ISL_500784, EPI_ISL_500785, EPI_ISL_500786, EPI_ISL_500787, EPI_ISL_500788, EPI_ISL_500789, EPI_ISL_500790 | Furst Medical Laboratory | Norwegian Institute of Public Health, Department of Virology | Kathrine Stene-Johansen, Kamilla Heddeland Instefjord, Hilde Elshaug, Rasmus Riis Kopperud, Karoline Bragstad, Olav Hungnes |
| EPI_ISL_500791, EPI_ISL_500792 | Hospital of Southern Norway - Kristiansand, Department of Medical Microbiology | Norwegian Institute of Public Health, Department of Virology | Kathrine Stene-Johansen, Kamilla Heddeland Instefjord, Hilde Elshaug, Rasmus Riis Kopperud, Karoline Bragstad, Olav Hungnes |
| EPI_ISL_500793, EPI_ISL_500794, EPI_ISL_500795, EPI_ISL_500796 | Akershus University Hospital, Department for Microbiology and Infectious Disease Control | Norwegian Institute of Public Health, Department of Virology | Kathrine Stene-Johansen, Kamilla Heddeland Instefjord, Hilde Elshaug, Rasmus Riis Kopperud, Karoline Bragstad, Olav Hungnes |
| EPI_ISL_500797, EPI_ISL_500798, EPI_ISL_500799 | Hospital of Southern Norway - Kristiansand, Department of Medical Microbiology | Norwegian Institute of Public Health, Department of Virology | Kathrine Stene-Johansen, Kamilla Heddeland Instefjord, Hilde Elshaug, Rasmus Riis Kopperud, Karoline Bragstad, Olav Hungnes |
| EPI_ISL_500800 | Oslo University Hospital, Department of Medical Microbiology | Norwegian Institute of Public Health, Department of Virology | Kathrine Stene-Johansen, Kamilla Heddeland Instefjord, Hilde Elshaug, Rasmus Riis Kopperud, Karoline Bragstad, Olav Hungnes |
| EPI_ISL_500801, EPI_ISL_500802, EPI_ISL_500803, EPI_ISL_500804, EPI_ISL_500805, EPI_ISL_500806, EPI_ISL_500807, EPI_ISL_500808, EPI_ISL_500809, EPI_ISL_500810, EPI_ISL_500811, EPI_ISL_500812, EPI_ISL_500813, EPI_ISL_500814, EPI_ISL_500815, EPI_ISL_500816, EPI_ISL_500817, EPI_ISL_500818, EPI_ISL_500819, EPI_ISL_500820, EPI_ISL_500821, EPI_ISL_500822, EPI_ISL_500823, EPI_ISL_500824, EPI_ISL_500825, EPI_ISL_500826, EPI_ISL_500827, EPI_ISL_500828, EPI_ISL_500829, EPI_ISL_500830 | | | |
| see above | National Institute for Biological Standards and Control | National Institute for Biological Standards and Control | Javier Martin, Dimitra Klapsa, Thomas Wilton |
| EPI_ISL_500831, EPI_ISL_500832, EPI_ISL_500833, EPI_ISL_500834, EPI_ISL_500835, EPI_ISL_500836, EPI_ISL_500837, EPI_ISL_500838, EPI_ISL_500839, EPI_ISL_500840, EPI_ISL_500841, EPI_ISL_500842, EPI_ISL_500843, EPI_ISL_500844, EPI_ISL_500845, EPI_ISL_500846, EPI_ISL_500847, EPI_ISL_500848, EPI_ISL_500849, EPI_ISL_500850, EPI_ISL_500851, EPI_ISL_500852, EPI_ISL_500853, EPI_ISL_500854, EPI_ISL_500855, EPI_ISL_500856, EPI_ISL_500857, EPI_ISL_500858, EPI_ISL_500859, EPI_ISL_500860, EPI_ISL_500861, EPI_ISL_500862, EPI_ISL_500863 | | | |
| see above | Virginia DCLS | Virginia DCLS | Virginia DCLS |
| EPI_ISL_500864 | Instituto de Pesquisas Biomédicas, Hospital Naval Marcílio Dias | Bioinformatics Laboratory / LNCC | Andressa Rangel de Oliveira Lima, Luiz Gonzaga Paula de Almeida, Alexandra Lehmkuhl Gerber, Marlon Daniel Lima Tonin, Mônica Gomes Monteiro da R. Santos, Claudio Alberto Mule Monteiro, Átila Duque Rossi, Isabel de Medeiros Magalhaes de As, Luana Ferreira Martins de Toledo, Amilcar Tanuri, Albina Luciana da Silva Freitas, Fernanda Conceição Silva do Amorim, Shana Priscila Coutinho Barroso, Cynthia Chester Cardoso, Carolina Moreira Voloch, Ana Tereza Vasconcelos |
| EPI_ISL_500865, EPI_ISL_500866, EPI_ISL_500867, EPI_ISL_500868, EPI_ISL_500869, EPI_ISL_500870, EPI_ISL_500871, EPI_ISL_500872, EPI_ISL_500873, EPI_ISL_500874, EPI_ISL_500875 | | | |
| see above | LACEN/PE | WallauLab, Aggeu Magalhaes Institute | Marcelo Henrique dos Santos Paiva, Duschinka Ribeiro Duarte Guedes, Cássia Docena, Matheus Filgueira Bezerra, Filipe Zimmer Dezordi, Laís Ceschini Machado, Larissa Krokovsky, Elisama Helvecio, Alexandre Freitas da Silva, Antonio Mauro Rezende5, Sinval Pinto Brandão Filho7, Constância Flávia Junqueira Ayres, Gabriel Luz Wallau |
| EPI_ISL_500877, EPI_ISL_500878, EPI_ISL_500879, EPI_ISL_500880, EPI_ISL_500881, EPI_ISL_500882, EPI_ISL_500883, EPI_ISL_500884, EPI_ISL_500885, EPI_ISL_500886, EPI_ISL_500887, EPI_ISL_500888, EPI_ISL_500889, EPI_ISL_500890, EPI_ISL_500891, EPI_ISL_500892, EPI_ISL_500893, EPI_ISL_500894, EPI_ISL_500895, EPI_ISL_500896, EPI_ISL_500897, EPI_ISL_500898, EPI_ISL_500899, EPI_ISL_500900, EPI_ISL_500901, EPI_ISL_500902, EPI_ISL_500903, EPI_ISL_500904, EPI_ISL_500905, EPI_ISL_500906, EPI_ISL_500907, EPI_ISL_500908, EPI_ISL_500909, EPI_ISL_500910, EPI_ISL_500911, EPI_ISL_500912, EPI_ISL_500913, EPI_ISL_500914, EPI_ISL_500915, EPI_ISL_500916, EPI_ISL_500917, EPI_ISL_500918, EPI_ISL_500919, EPI_ISL_500920, EPI_ISL_500921, EPI_ISL_500922, EPI_ISL_500923, EPI_ISL_500924, EPI_ISL_500925, EPI_ISL_500926, EPI_ISL_500927, EPI_ISL_500928, EPI_ISL_500929, EPI_ISL_500930, EPI_ISL_500931, EPI_ISL_500932, EPI_ISL_500933, EPI_ISL_500934, EPI_ISL_500935, EPI_ISL_500936, EPI_ISL_500937, EPI_ISL_500938, EPI_ISL_500939, EPI_ISL_500940, EPI_ISL_500941, EPI_ISL_500942, EPI_ISL_500943, EPI_ISL_500944, EPI_ISL_500945 | | | |
| see above | Viollier AG | Department of Biosystems Science and Engineering, ETH Zürich | Christian Beisel, Sarah Nadeau, Ivan Topolsky, Pedro Ferreira, Philipp Jablonski, Susana Posada-Céspedes, Tobias Schär, Ina Nissen, Natascha Santacroce, Elodie Burcklen, Christiane Beckmann, Maurice Redondo, Olivier Kobel, Christoph Noppen, Sophie Seidel, Noemie Santamaria de Souza, Niko Beerenwinkel, Tanja Stadler |
| EPI_ISL_500946 | GMERS Medical College & Hospital, Gotri, Vadodara | Gujarat Biotechnology Research Centre | Zuber Saiyed, Komal Patel, Labdhi Pandya, Afzal Ansari, Nikha Trivedi, Meenakshi Shah, Neena Doshi, Varsha Godbole, Apurvasinh Puvar, Janvi Raval, Zarna Patel, Monika Gandhi, Pinal Trivedi, Maharshi Pandya, Nidhi Patel, Nitin Savaliya, Raghawendra Kumar, Dinesh Kumar, R D Dixit, A M Kadri, Harsh Bakshi, Chaitanya Joshi, Madhvi Joshi |
| EPI_ISL_500947 | Department of MicroBiology, Government Medical College, Surat | Gujarat Biotechnology Research Centre | Maharshi Pandya, Nidhi Patel, Nitin Savaliya, Raghawendra Kumar, Dinesh Kumar, Zuber Saiyed, Komal Patel, Labdhi Pandya, Afzal Ansari, Nikha Trivedi, Naresh Chauhan, Summaiya Mullan, Amit gamit, Apurvasinh Puvar, Janvi Raval, Zarna Patel, Monika Gandhi, Pinal Trivedi, R D Dixit, A M Kadri, Harsh Bakshi, Chaitanya Joshi, Madhvi Joshi |

| | | | |
|---|---|---|---|
| EPI_ISL_500948 | Department of MicroBiology, Government Medical College, Surat | Gujarat Biotechnology Research Centre | Zuber Saiyed, Komal Patel, Labdhi Pandya, Afzal Ansari, Nikha Trivedi, Naresh Chauhan, Summaiya Mullan, Amit gamit, Apurvasinh Puvar, Janvi Raval, Zarna Patel, Monika Gandhi, Pinal Trivedi, Maharshi Pandya, Nidhi Patel, Nitin Savaliya, Raghawendra Kumar, Dinesh Kumar, R D Dixit, A M Kadri, Harsh Bakshi, Chaitanya Joshi, Madhvi Joshi |
| EPI_ISL_500949 | Department of MicroBiology, Government Medical College, Surat | Gujarat Biotechnology Research Centre | Komal Patel, Labdhi Pandya, Afzal Ansari, Nikha Trivedi, Naresh Chauhan, Summaiya Mullan, Amit gamit, Apurvasinh Puvar, Janvi Raval, Zarna Patel, Monika Gandhi, Pinal Trivedi, Maharshi Pandya, Nidhi Patel, Nitin Savaliya, Raghawendra Kumar, Dinesh Kumar, Zuber Saiyed, R D Dixit, A M Kadri, Harsh Bakshi, Chaitanya Joshi, Madhvi Joshi |
| EPI_ISL_500950 | Department of MicroBiology, Government Medical College, Surat | Gujarat Biotechnology Research Centre | Labdhi Pandya, Afzal Ansari, Nikha Trivedi, Naresh Chauhan, Summaiya Mullan, Amit gamit, Apurvasinh Puvar, Janvi Raval, Zarna Patel, Monika Gandhi, Pinal Trivedi, Maharshi Pandya, Nidhi Patel, Nitin Savaliya, Raghawendra Kumar, Dinesh Kumar, Zuber Saiyed, Komal Patel, R D Dixit, A M Kadri, Harsh Bakshi, Chaitanya Joshi, Madhvi Joshi |
| EPI_ISL_500951, EPI_ISL_500952, EPI_ISL_500953 | Respiratory Virus Unit, Microbiology Services Colindale, Public Health England | Respiratory Virus Unit, Microbiology Services Colindale, Public Health England | PHE Covid Sequencing Team |
| EPI_ISL_500954, EPI_ISL_500955, EPI_ISL_500956, EPI_ISL_500957, EPI_ISL_500958, EPI_ISL_500959, EPI_ISL_500960, EPI_ISL_500961, EPI_ISL_500962, EPI_ISL_500963, EPI_ISL_500964, EPI_ISL_500965, EPI_ISL_500966, EPI_ISL_500967, EPI_ISL_500968, EPI_ISL_500969, EPI_ISL_500970, EPI_ISL_500971, EPI_ISL_500972, EPI_ISL_500973, EPI_ISL_500974, EPI_ISL_500975, EPI_ISL_500976, EPI_ISL_500977, EPI_ISL_500978, EPI_ISL_500979, EPI_ISL_500980, EPI_ISL_500981, EPI_ISL_500982, EPI_ISL_500983, EPI_ISL_500984, EPI_ISL_500985, EPI_ISL_500986, EPI_ISL_500987, EPI_ISL_500988, EPI_ISL_500989, EPI_ISL_500990, EPI_ISL_500991, EPI_ISL_500992, EPI_ISL_500993, EPI_ISL_500994, EPI_ISL_500995, EPI_ISL_500996, EPI_ISL_500997, EPI_ISL_500998, EPI_ISL_500999, EPI_ISL_501000, EPI_ISL_501001, EPI_ISL_501002, EPI_ISL_501003, EPI_ISL_501004, EPI_ISL_501005, EPI_ISL_501006, EPI_ISL_501007, EPI_ISL_501008, EPI_ISL_501009, EPI_ISL_501010, EPI_ISL_501011, EPI_ISL_501012, EPI_ISL_501013, EPI_ISL_501014, EPI_ISL_501015, EPI_ISL_501016, EPI_ISL_501017, EPI_ISL_501018, EPI_ISL_501019, EPI_ISL_501020, EPI_ISL_501021, EPI_ISL_501022, EPI_ISL_501023, EPI_ISL_501024, EPI_ISL_501025, EPI_ISL_501026, EPI_ISL_501027, EPI_ISL_501028, EPI_ISL_501029, EPI_ISL_501030, EPI_ISL_501031, EPI_ISL_501032, EPI_ISL_501033, EPI_ISL_501034, EPI_ISL_501035, EPI_ISL_501036, EPI_ISL_501037, EPI_ISL_501038, EPI_ISL_501039, EPI_ISL_501040, EPI_ISL_501041, EPI_ISL_501042, EPI_ISL_501043, EPI_ISL_501044, EPI_ISL_501045, EPI_ISL_501046, EPI_ISL_501047, EPI_ISL_501048, EPI_ISL_501049, EPI_ISL_501050, EPI_ISL_501051, EPI_ISL_501052, EPI_ISL_501053, EPI_ISL_501054, EPI_ISL_501055, EPI_ISL_501056, EPI_ISL_501057, EPI_ISL_501058, EPI_ISL_501059, EPI_ISL_501060, EPI_ISL_501061, EPI_ISL_501062, EPI_ISL_501063, EPI_ISL_501064, EPI_ISL_501065, EPI_ISL_501066, EPI_ISL_501067, EPI_ISL_501068, EPI_ISL_501069, EPI_ISL_501070, EPI_ISL_501071 | | | |
| see above | Regional Virus Laboratory, Belfast Health and Social Care Trust | Wellcome Sanger Institute for the COVID-19 Genomics UK Consortium | Conall McCaughey, James McKenna, Tanya Curran, Susan Feeney, Alison Watt, Ciara Cox, Mairead Connor, Zoltan Molnar, David Simpson, Derek Fairley; and Alex Alderton, Roberto Amato, Sonia Goncalves, Ewan Harrison, David K. Jackson, Ian Johnston, Dominic Kwiatkowski, Cordelia Langford, John Sillitoe on behalf of the Wellcome Sanger Institute COVID-19 Surveillance Team (http://www.sanger.ac.uk/covid-team) |
| EPI_ISL_501072, EPI_ISL_501073, EPI_ISL_501074, EPI_ISL_501075, EPI_ISL_501076, EPI_ISL_501077, EPI_ISL_501078, EPI_ISL_501079, EPI_ISL_501080, EPI_ISL_501081, EPI_ISL_501082 | | | |
| see above | Mayo Clinic Laboratories | University of Washington Virology Lab | Pavitra Roychoudhury, Hong Xie, Lasata Shrestha, Amin Addetia, Truong Nguyen, Victoria M Rachleff, Meei-Li Huang, Keith R Jerome, Alexander Greninger |
| EPI_ISL_501083, EPI_ISL_501084, EPI_ISL_501085, EPI_ISL_501086, EPI_ISL_501087, EPI_ISL_501088, EPI_ISL_501089, EPI_ISL_501090, EPI_ISL_501091, EPI_ISL_501092, EPI_ISL_501093, EPI_ISL_501094, EPI_ISL_501095, EPI_ISL_501096, EPI_ISL_501097, EPI_ISL_501098, EPI_ISL_501099, EPI_ISL_501100, EPI_ISL_501101, EPI_ISL_501102, EPI_ISL_501103, EPI_ISL_501104, EPI_ISL_501105, EPI_ISL_501106, EPI_ISL_501107, EPI_ISL_501108, EPI_ISL_501109, EPI_ISL_501110, EPI_ISL_501111, EPI_ISL_501112, EPI_ISL_501113, EPI_ISL_501114, EPI_ISL_501115, EPI_ISL_501116, EPI_ISL_501117, EPI_ISL_501118, EPI_ISL_501119, EPI_ISL_501120, EPI_ISL_501121, EPI_ISL_501122, EPI_ISL_501123, EPI_ISL_501124, EPI_ISL_501125, EPI_ISL_501126, EPI_ISL_501127, EPI_ISL_501128, EPI_ISL_501129, EPI_ISL_501130, EPI_ISL_501131, EPI_ISL_501132, EPI_ISL_501133, EPI_ISL_501134, EPI_ISL_501135, EPI_ISL_501136, EPI_ISL_501137, EPI_ISL_501138, EPI_ISL_501139, EPI_ISL_501140, EPI_ISL_501141, EPI_ISL_501142, EPI_ISL_501143, EPI_ISL_501144, EPI_ISL_501145, EPI_ISL_501146, EPI_ISL_501147, EPI_ISL_501148, EPI_ISL_501149, EPI_ISL_501150, EPI_ISL_501151, EPI_ISL_501152, EPI_ISL_501153, EPI_ISL_501154, EPI_ISL_501155, EPI_ISL_501156, EPI_ISL_501157, EPI_ISL_501158, EPI_ISL_501159, EPI_ISL_501160, EPI_ISL_501161, EPI_ISL_501162, EPI_ISL_501163, EPI_ISL_501164 | | | |
| see above | University of Washington Virology Lab | University of Washington Virology Lab | Pavitra Roychoudhury, Hong Xie, Lasata Shrestha, Amin Addetia, Truong Nguyen, Victoria M Rachleff, Meei-Li Huang, Keith R Jerome, Alexander Greninger |
| EPI_ISL_501165 | Mayo Clinic Laboratories | University of Washington Virology Lab | Pavitra Roychoudhury, Hong Xie, Lasata Shrestha, Amin Addetia, Truong Nguyen, Victoria M Rachleff, Meei-Li Huang, Keith R Jerome, Alexander Greninger |
| EPI_ISL_501167, EPI_ISL_501168, EPI_ISL_501169, EPI_ISL_501170, EPI_ISL_501171, EPI_ISL_501172, EPI_ISL_501173, EPI_ISL_501174 | Baylor College of Medicine | Baylor College of Medicine: HGSC | Vasanthi Avadhanula, Erin Nicholson, David Henke, Pedro Piedra, Harsha Doddapaneni, Donna Muzny, Qingchang Meng, Hsu Chao, Zeineen Momin, Hua Shen, George Weissenberger, Kavya Kottapalli, Yimiti Meiheerguli, Sejal Salvi, Ginger Metcalf, Vipin Menon, Sara J.J. Cregeen, Matthew C. Ross, Tulin Ayvaz, Richard Sucgang, Kristi L. Hoffman, Matthew Wong, Joseph F. Petrosino |
| EPI_ISL_501176, EPI_ISL_501177, EPI_ISL_501178, EPI_ISL_501179, EPI_ISL_501180, EPI_ISL_501181, EPI_ISL_501182, EPI_ISL_501183, EPI_ISL_501184, EPI_ISL_501185, EPI_ISL_501186, EPI_ISL_501187, EPI_ISL_501188, EPI_ISL_501189, EPI_ISL_501190, EPI_ISL_501191, EPI_ISL_501192, EPI_ISL_501193, EPI_ISL_501194, EPI_ISL_501195, EPI_ISL_501196, EPI_ISL_501197, EPI_ISL_501198, EPI_ISL_501199, EPI_ISL_501200, EPI_ISL_501201, EPI_ISL_501202, EPI_ISL_501203, EPI_ISL_501204, EPI_ISL_501205, EPI_ISL_501206, EPI_ISL_501207, EPI_ISL_501208, EPI_ISL_501209, EPI_ISL_501210, EPI_ISL_501211, EPI_ISL_501212, EPI_ISL_501213, EPI_ISL_501214, EPI_ISL_501215, EPI_ISL_501216, EPI_ISL_501217, EPI_ISL_501218, EPI_ISL_501219, EPI_ISL_501220, EPI_ISL_501221, EPI_ISL_501222, EPI_ISL_501223, EPI_ISL_501224, EPI_ISL_501225, EPI_ISL_501226, EPI_ISL_501227, EPI_ISL_501228 | | | |
| see above | Department of Medical Microbiology, University Malaya Medical Centre | Department of Medical Microbiology, Faculty of Medicine, University of Malaya | Yoong Min CHONG, Jennifer Chong, I-Ching SAM, Yoke Fun CHAN, University Malaya Medical Centre COVID Team |
| EPI_ISL_501230 | Hellenic Pasteur Institute, Public Health Laboratories | Hellenic Pasteur Institute, National Influenza Reference laboratory of Southern Greece & Unit of Bioinformatics and Applied Genomics | Vasiliki Pogka, Timokratis Karamitros, Athanasios Kossyvakis, Antonios Kalliaropoulos, Horefti Elina, Evangelidou Maria, Androniki Voulgari-Kokota, Aspasia Kontou, Andreas Mentis |
| EPI_ISL_501231, EPI_ISL_501232, EPI_ISL_501233, EPI_ISL_501234, EPI_ISL_501235, EPI_ISL_501236, EPI_ISL_501237, EPI_ISL_501238, EPI_ISL_501241, EPI_ISL_501242, EPI_ISL_501246, EPI_ISL_501248, EPI_ISL_501249, EPI_ISL_501250, EPI_ISL_501251 | | | |
| see above | Hellenic Pasteur Institute, National Influenza Reference laboratory of Southern Greece & Unit of Bioinformatics and Applied Genomics | Hellenic Pasteur Institute, National Influenza Reference laboratory of Southern Greece & Unit of Bioinformatics and Applied Genomics | Vasiliki Pogka, Timokratis Karamitros, Athanasios Kossyvakis, Antonios Kalliaropoulos, Horefti Elina, Evangelidou Maria, Androniki Voulgari-Kokota, Aspasia Kontou, Andreas Mentis |
| EPI_ISL_501252, EPI_ISL_501253, EPI_ISL_501254, EPI_ISL_501255, EPI_ISL_501256, EPI_ISL_501257, EPI_ISL_501258, EPI_ISL_501259, EPI_ISL_501260, EPI_ISL_501261, EPI_ISL_501262, EPI_ISL_501263, EPI_ISL_501264, EPI_ISL_501265, EPI_ISL_501266, EPI_ISL_501267, EPI_ISL_501268, EPI_ISL_501269, EPI_ISL_501270, EPI_ISL_501271, EPI_ISL_501272, EPI_ISL_501273, EPI_ISL_501274 | | | |
| see above | National Virus Reference Laboratory | National Virus Reference Laboratory | Michael Carr, Gabriel Gonzalez, Jonathan Dean, Suzie Coughlan, Cillian F De Gascun |
| EPI_ISL_501275 | E. Gulbja Laboratorija | Latvian Biomedical Research and Study Centre | Ivars Silamiķelis, Kaspars Megnis, Monta Ustinova, Ņikita Zrelovs, Vita Rovīte, Mikus Gavars, Dmitrijs Perminovs, Uga Dumpis, Jānis Kloviņš |
| EPI_ISL_501284, EPI_ISL_501285 | E. Gulbja Laboratorija | Latvian Biomedical Research and Study Centre | Ivars Silamiķelis, Kaspars Megnis, Monta Ustinova, Ņikita Zrelovs, Vita Rovīte, Mikus Gavars, Dmitrijs Perminovs, Uga Dumpis, Jānis Kloviņš |
| EPI_ISL_501286, EPI_ISL_501287, EPI_ISL_501288, EPI_ISL_501289 | Centrālā laboratorija | Latvian Biomedical Research and Study Centre | Ivars Silamiķelis, Kaspars Megnis, Monta Ustinova, Ņikita Zrelovs, Vita Rovīte, Stella Lapiņa, Jana Osīte, Marta Priedīte, Uga Dumpis, Jānis Kloviņš |
| EPI_ISL_501555, EPI_ISL_501556 | PHE South West Regional Laboratory, National Infection Service | Wellcome Sanger Institute for the COVID-19 Genomics UK Consortium | Stephanie Hutchings, Hannah Pymont, Dr Peter Muir, Barry Vipond, Rich Hopes; and Alex Alderton, Roberto Amato, Sonia Goncalves, Ewan Harrison, David K. Jackson, Ian Johnston, Dominic Kwiatkowski, Cordelia Langford, John Sillitoe on behalf of the Wellcome Sanger Institute COVID-19 Surveillance Team (http://www.sanger.ac.uk/covid-team) |
| EPI_ISL_501557 | Department of Medical Microbiology, Western Sussex Hospitals NHS Foundation Trust, St Richard's Hospital | Wellcome Sanger Institute for the COVID-19 Genomics UK Consortium | Manasa Mutingwende, Sarah Lowdon, Olga Podplomyk, Michelle Erkiert, Jonathan Lewis, Paul Randell and Alex Alderton, Roberto Amato, Sonia Goncalves, Ewan Harrison, David K. Jackson, Ian Johnston, Dominic Kwiatkowski, Cordelia Langford, John Sillitoe on behalf of the Wellcome Sanger Institute COVID-19 Surveillance Team (http://www.sanger.ac.uk/covid-team) |
| EPI_ISL_501558, EPI_ISL_501559, EPI_ISL_501560, EPI_ISL_501561 | PHE South West Regional Laboratory, National Infection Service | Wellcome Sanger Institute for the COVID-19 Genomics UK Consortium | Stephanie Hutchings, Hannah Pymont, Dr Peter Muir, Barry Vipond, Rich Hopes; and Alex Alderton, Roberto Amato, Sonia Goncalves, Ewan Harrison, David K. Jackson, Ian Johnston, Dominic Kwiatkowski, Cordelia Langford, John Sillitoe on behalf of the Wellcome Sanger Institute COVID-19 Surveillance Team (http://www.sanger.ac.uk/covid-team) |
| EPI_ISL_501562 | Department of Medical Microbiology, Western Sussex Hospitals NHS Foundation Trust, St Richard's Hospital | Wellcome Sanger Institute for the COVID-19 Genomics UK Consortium | Manasa Mutingwende, Sarah Lowdon, Olga Podplomyk, Michelle Erkiert, Jonathan Lewis, Paul Randell and Alex Alderton, Roberto Amato, Sonia Goncalves, Ewan Harrison, David K. Jackson, Ian Johnston, Dominic Kwiatkowski, Cordelia Langford, John Sillitoe on behalf of the Wellcome Sanger Institute COVID-19 Surveillance Team (http://www.sanger.ac.uk/covid-team) |
| EPI_ISL_501563, EPI_ISL_501564, EPI_ISL_501565, EPI_ISL_501566, EPI_ISL_501567, EPI_ISL_501568, EPI_ISL_501569, EPI_ISL_501570, EPI_ISL_501571, EPI_ISL_501572, EPI_ISL_501573, EPI_ISL_501574, EPI_ISL_501575, EPI_ISL_501576, EPI_ISL_501577, EPI_ISL_501578, EPI_ISL_501579, EPI_ISL_501580, EPI_ISL_501581, EPI_ISL_501582, EPI_ISL_501583, EPI_ISL_501584, EPI_ISL_501585, EPI_ISL_501586, EPI_ISL_501587, EPI_ISL_501588, EPI_ISL_501589, EPI_ISL_501590, EPI_ISL_501591, EPI_ISL_501592, EPI_ISL_501593, EPI_ISL_501594, EPI_ISL_501595, EPI_ISL_501596, EPI_ISL_501597, EPI_ISL_501598, EPI_ISL_501599, EPI_ISL_501601 | | | |
| see above | PHE South West Regional Laboratory, National Infection Service | Wellcome Sanger Institute for the COVID-19 Genomics UK Consortium | Stephanie Hutchings, Hannah Pymont, Dr Peter Muir, Barry Vipond, Rich Hopes; and Alex Alderton, Roberto Amato, Sonia Goncalves, Ewan Harrison, David K. Jackson, Ian Johnston, Dominic Kwiatkowski, Cordelia Langford, John Sillitoe on behalf of the Wellcome Sanger Institute COVID-19 Surveillance Team (http://www.sanger.ac.uk/covid-team) |
| EPI_ISL_501602 | Department of Medical Microbiology, Western Sussex Hospitals NHS Foundation Trust, St Richard's Hospital | Wellcome Sanger Institute for the COVID-19 Genomics UK Consortium | Manasa Mutingwende, Sarah Lowdon, Olga Podplomyk, Michelle Erkiert, Jonathan Lewis, Paul Randell and Alex Alderton, Roberto Amato, Sonia Goncalves, Ewan Harrison, David K. Jackson, Ian Johnston, Dominic Kwiatkowski, Cordelia Langford, John Sillitoe on behalf of the Wellcome Sanger Institute COVID-19 Surveillance Team (http://www.sanger.ac.uk/covid-team) |
| EPI_ISL_501603, EPI_ISL_501604, EPI_ISL_501605, EPI_ISL_501606, EPI_ISL_501607, EPI_ISL_501608, EPI_ISL_501609, EPI_ISL_501610, EPI_ISL_501611, EPI_ISL_501612 | PHE South West Regional Laboratory, National Infection Service | Wellcome Sanger Institute for the COVID-19 Genomics UK Consortium | Stephanie Hutchings, Hannah Pymont, Dr Peter Muir, Barry Vipond, Rich Hopes; and Alex Alderton, Roberto Amato, Sonia Goncalves, Ewan Harrison, David K. Jackson, Ian Johnston, Dominic Kwiatkowski, Cordelia Langford, John Sillitoe on behalf of the Wellcome Sanger Institute COVID-19 Surveillance Team (http://www.sanger.ac.uk/covid-team) |
| EPI_ISL_501613 | Department of Medical Microbiology, Western Sussex Hospitals NHS Foundation Trust, St Richard's Hospital | Wellcome Sanger Institute for the COVID-19 Genomics UK Consortium | Manasa Mutingwende, Sarah Lowdon, Olga Podplomyk, Michelle Erkiert, Jonathan Lewis, Paul Randell and Alex Alderton, Roberto Amato, Sonia Goncalves, Ewan Harrison, David K. Jackson, Ian Johnston, Dominic Kwiatkowski, Cordelia Langford, John Sillitoe on behalf of the Wellcome Sanger Institute COVID-19 Surveillance Team (http://www.sanger.ac.uk/covid-team) |
| EPI_ISL_501614, EPI_ISL_501615, EPI_ISL_501616, EPI_ISL_501617, EPI_ISL_501618, EPI_ISL_501619, EPI_ISL_501620, EPI_ISL_501621, EPI_ISL_501622, EPI_ISL_501623, EPI_ISL_501624, EPI_ISL_501625, EPI_ISL_501626, EPI_ISL_501627, EPI_ISL_501628, EPI_ISL_501629, EPI_ISL_501630, EPI_ISL_501631 | | | |
| see above | Lab Microbiology, Pathology Department, William Harvey Hospital | Wellcome Sanger Institute for the COVID-19 Genomics UK Consortium | Samuel Moses, Hannah Lowe, Felicity Ryan and Alex Alderton, Roberto Amato, Sonia Goncalves, Ewan Harrison, David K. Jackson, Ian Johnston, Dominic Kwiatkowski, Cordelia Langford, John Sillitoe on behalf of the Wellcome Sanger Institute COVID-19 Surveillance Team (http://www.sanger.ac.uk/covid-team) |
| EPI_ISL_501632 | Virology Department, Royal Infirmary of Edinburgh, NHS Lothian / School of Biological Sciences, University of Edinburgh | Wellcome Sanger Institute for the COVID-19 Genomics UK Consortium | McHugh M, Dewar R, Rooke S, O'Toole Á, Scher E, Hill V, McCrone JT, Colquhoun R, Yu X, Jackson B, Rambaut A, Templeton K and Alex Alderton, Roberto Amato, Sonia Goncalves, Ewan Harrison, David K. Jackson, Ian Johnston, Dominic Kwiatkowski, Cordelia Langford, John Sillitoe on behalf of the Wellcome Sanger Institute COVID-19 Surveillance Team (http://www.sanger.ac.uk/covid-team) |
| EPI_ISL_501633, EPI_ISL_501634, EPI_ISL_501635 | NHSGGC West of Scotland Specialist Virology Centre / MRC-University of Glasgow Centre for Virus Research | Wellcome Sanger Institute for the COVID-19 Genomics UK Consortium | Ana da Silva Filipe, Natasha Johnson, Kathy Smollett, Daniel Mair, Stephen Carmichael, Lily Tong, Jenna Nichols, Elihu Aranday-Cortes, Kirstyn Brunker, Yasmin Parr, Kyriaki Nomikou; Sarah McDonald, Marc Niebel, Patawee Asamaphan; Richard Orton, Joseph Hughes, Sreenu Vattipally, David L Robertson; Alasdair MacLean, Rory Gunson; Kathy Li, Natasha Jesudason, Rajiv Shah, James Shepherd, Antonia Ho, Alice Broos, Emma Thomson and Alex Alderton, Roberto Amato, Sonia Goncalves, Ewan Harrison, David K. Jackson, Ian Johnston, Dominic Kwiatkowski, Cordelia Langford, John Sillitoe on behalf of the Wellcome Sanger Institute COVID-19 Surveillance Team (http://www.sanger.ac.uk/covid-team) |
| EPI_ISL_501808, EPI_ISL_501817 | Centrālā laboratorija | Latvian Biomedical Research and Study Centre | Ivars Silamiķelis, Kaspars Megnis, Monta Ustinova, Ņikita Zrelovs, Vita Rovīte, Stella Lapiņa, Jana Osīte, Marta Priedīte, Uga Dumpis, Jānis Kloviņš |
| EPI_ISL_501823, EPI_ISL_501829, EPI_ISL_501833, EPI_ISL_501839, EPI_ISL_501849, EPI_ISL_501894 | E. Gulbja Laboratorija | Latvian Biomedical Research and Study Centre | Ivars Silamiķelis, Kaspars Megnis, Monta Ustinova, Ņikita Zrelovs, Vita Rovīte, Mikus Gavars, Dmitrijs Perminovs, Uga Dumpis, Jānis Kloviņš |
| EPI_ISL_501895, EPI_ISL_501896, EPI_ISL_501915, EPI_ISL_501922 | Centrālā laboratorija | Latvian Biomedical Research and Study Centre | Ivars Silamiķelis, Kaspars Megnis, Monta Ustinova, Ņikita Zrelovs, Vita Rovīte, Stella Lapiņa, Jana Osīte, Marta Priedīte, Uga Dumpis, Jānis Kloviņš |
| EPI_ISL_501929, EPI_ISL_501936 | E. Gulbja Laboratorija | Latvian Biomedical Research and Study Centre | Ivars Silamiķelis, Kaspars Megnis, Monta Ustinova, Ņikita Zrelovs, Vita Rovīte, Mikus Gavars, Dmitrijs Perminovs, Uga Dumpis, Jānis Kloviņš |
| EPI_ISL_502779 | LACEN/PE | LABBE, Federal University of Pernambuco | WILSON JOSE DA SILVA JUNIOR, HEIDI LACERDA ALVES DA CRUZ, MARCOS DA SILVEIRA REGUEIRA NETO, BRUNO SAMPAIO, SERGIO DE SA LEITAO PAIVA JUNIOR, ZILDENE DE SOUSA SILVEIRA, MAIRA GALDINO DA ROCHA PITTA, MICHELLY CRISTINY PEREIRA, MARCOS ANTONIO DE MORAIS JUNIOR, ANTONIO CARLOS DE FREITAS, VALDIR DE QUEIROZ BALBINO. |
| EPI_ISL_502875 | LACEN/PE | LABBE, Federal University of Pernambuco | WILSON JOSE DA SILVA JUNIOR, HEIDI LACERDA ALVES DA CRUZ, MARCOS DA SILVEIRA REGUEIRA NETO, BRUNO SAMPAIO, SERGIO DE SA LEITAO PAIVA JUNIOR, ZILDENE DE SOUSA SILVEIRA, MAIRA GALDINO DA ROCHA PITTA, MICHELLY CRISTINY PEREIRA, REGINALDO GONCALVES DE LIMA NETO, MARCOS ANTONIO DE MORAIS JUNIOR, ANTONIO CARLOS DE FREITAS, VALDIR DE QUEIROZ BALBINO. |
