## Supplementary material for "Ultrafast Sample Placement on Existing Trees (UShER) Empowers Real-Time Phylogenetics for the SARS-CoV-2 Pandemic": File_S4

We gratefully acknowledge the following Authors from the Originating laboratories responsible for obtaining the specimens, as well as the Submitting laboratories where the genome data were generated and shared via GISAID, on which this research is based.

All Submitters of data may be contacted directly via www.gisaid.org

| Accession ID | Originating Laboratory | Submitting Laboratory | Authors |
|---|---|---|---|
| EPI_ISL_480414, EPI_ISL_480415, EPI_ISL_480416, EPI_ISL_480417, EPI_ISL_480418 | National Institute of Laboratory Medicine and Referral Center | Bangladesh Council of Scientific and Industrial Research | Md. Saddam Hossain, Abu Sayeed Mohammad Mahmud, Mohammad Samir Uzzaman, Eshrar Osman, Md. Ahasan Habib, Shahina Akter, Tanjina Akhter Banu, Md. Murshed Hasan Sarkar, Barna Goswami, Iffat Jahan, Tasnim Nafisa, Md. Maruf Ahmed Molla, Mahmuda Yeasmin, Asish Kumar Ghosh, Shahjahan Siddike, A. K. M. Shamsuzzaman, Sheikh Md. Selim Al Din, Utpal Chandra Ray, Salek Ahmed Sajib, Md. Salim Khan |
| EPI_ISL_480419, EPI_ISL_480420, EPI_ISL_480421, EPI_ISL_480424, EPI_ISL_480425 | National Institute of Laboratory Medicine and Referral Center | Bangladesh Council of Scientific and Industrial Research | Md. Murshed Hasan Sarkar, Abu Sayeed Mohammad Mahmud, Mohammad Samir Uzzaman, Eshrar Osman, Md. Ahasan Habib, Shahina Akter, Tanjina Akhter Banu, Barna Goswami, Iffat Jahan, Md. Saddam Hossain, Tasnim Nafisa, Md. Maruf Ahmed Molla, Mahmuda Yeasmin, Asish Kumar Ghosh, Shahjahan Siddike, A. K. M. Shamsuzzaman, Sheikh Md. Selim Al Din, Utpal Chandra Ray, Salek Ahmed Sajib, Md. Salim Khan |
| EPI_ISL_480426, EPI_ISL_480427 | National Institute of Laboratory Medicine and Referral Center | Bangladesh Council of Scientific and Industrial Research | Shahina Akter, Abu Sayeed Mohammad Mahmud, Mohammad Samir Uzzaman, Eshrar Osman, Md. Ahasan Habib, Tanjina Akhter Banu, Md. Murshed Hasan Sarkar, Barna Goswami, Iffat Jahan, Md. Saddam Hossain, Tasnim Nafisa, Md. Maruf Ahmed Molla, Mahmuda Yeasmin, Asish Kumar Ghosh, Shahjahan Siddike, A. K. M. Shamsuzzaman, Sheikh Md. Selim Al Din, Utpal Chandra Ray, Salek Ahmed Sajib, Md. Salim Khan |
| EPI_ISL_480428, EPI_ISL_480429, EPI_ISL_480430, EPI_ISL_480431, EPI_ISL_480432, EPI_ISL_480433, EPI_ISL_480434, EPI_ISL_480435, EPI_ISL_480436, EPI_ISL_480437, EPI_ISL_480438 | | | |
| see above | Laboratorio de Biología Molecular Asociación Española Primera en Salud | Departments of Pathology and Medicine, New York University School of Medicine | Maria Victoria Elizondo, Maria Noel Zubillaga, Gonzalo Manrique, Paul Zappile, Gael Westby, Matthew T Maurano, Christian Marier, Adriana Heguy |
| EPI_ISL_480439, EPI_ISL_480440 | National Institute of Laboratory Medicine and Referral Center | Bangladesh Council of Scientific and Industrial Research | Tanjina Akhter Banu, Abu Sayeed Mohammad Mahmud, Mohammad Samir Uzzaman, Eshrar Osman, Md. Ahasan Habib, Shahina Akter, Md. Murshed Hasan Sarkar, Barna Goswami, Iffat Jahan, Md. Saddam Hossain, Tasnim Nafisa, Md. Maruf Ahmed Molla, Mahmuda Yeasmin, Asish Kumar Ghosh, Shahjahan Siddike, A. K. M. Shamsuzzaman, Sheikh Md. Selim Al Din, Utpal Chandra Ray, Salek Ahmed Sajib, Md. Salim Khan |
| EPI_ISL_480441, EPI_ISL_480442 | National Institute of Laboratory Medicine and Referral Center | Bangladesh Council of Scientific and Industrial Research | Barna Goswami, Abu Sayeed Mohammad Mahmud, Mohammad Samir Uzzaman, Eshrar Osman, Md. Ahasan Habib, Shahina Akter, Tanjina Akhter Banu, Md. Murshed Hasan Sarkar, Iffat Jahan, Md. Saddam Hossain, Tasnim Nafisa, Md. Maruf Ahmed Molla, Mahmuda Yeasmin, Asish Kumar Ghosh, Shahjahan Siddike, A. K. M. Shamsuzzaman, Sheikh Md. Selim Al Din, Utpal Chandra Ray, Salek Ahmed Sajib, Md. Salim Khan |
| EPI_ISL_480443, EPI_ISL_480444 | National Institute of Laboratory Medicine and Referral Center | Bangladesh Council of Scientific and Industrial Research | Iffat Jahan, Abu Sayeed Mohammad Mahmud, Mohammad Samir Uzzaman, Eshrar Osman, Md. Ahasan Habib, Shahina Akter, Tanjina Akhter Banu, Md. Murshed Hasan Sarkar, Barna Goswami, Md. Saddam Hossain, Tasnim Nafisa, Md. Maruf Ahmed Molla, Mahmuda Yeasmin, Asish Kumar Ghosh, Shahjahan Siddike, A. K. M. Shamsuzzaman, Sheikh Md. Selim Al Din, Utpal Chandra Ray, Salek Ahmed Sajib, Md. Salim Khan |
| EPI_ISL_480445 | National Institute of Laboratory Medicine and Referral Center | Genomic Research Lab, BCSIR | Md. Ahasan Habib, Abu Sayeed Mohammad Mahmud, Mohammad Samir Uzzaman, Eshrar Osman, , Shahina Akter, Tanjina Akhter Banu, Md. Murshed Hasan Sarkar, Barna Goswami, Iffat Jahan, Md. Saddam Hossain, Tasnim Nafisa, Md. Maruf Ahmed Molla, Mahmuda Yeasmin, Asish Kumar Ghosh, Shahjahan Siddike, A. K. M. Shamsuzzaman, Sheikh Md. Selim Al Din, Utpal Chandra Ray, Salek Ahmed Sajib, Md. Salim Khan |
| EPI_ISL_480446, EPI_ISL_480447, EPI_ISL_480448, EPI_ISL_480449, EPI_ISL_480450 | National Institute of Laboratory Medicine and Referral Center | Genomic Research Lab, BCSIR | Abu Sayeed Mohammad Mahmud, Mohammad Samir Uzzaman, Eshrar Osman, Md. Ahasan Habib, Shahina Akter, Tanjina Akhter Banu, Md. Murshed Hasan Sarkar, Barna Goswami, Iffat Jahan, Md. Saddam Hossain, Tasnim Nafisa, Md. Maruf Ahmed Molla, Mahmuda Yeasmin, Asish Kumar Ghosh, Shahjahan Siddike, A. K. M. Shamsuzzaman, Sheikh Md. Selim Al Din, Utpal Chandra Ray, Salek Ahmed Sajib, Md. Salim Khan |
| EPI_ISL_480554, EPI_ISL_480556 | Institut Pasteur Dakar | Institut Pasteur de Dakar | Ndongo Dia, Moussa Moise Diagne, Mamadou Diop, Marie Henriette Dior Ndione, Mamadou Malado Jallow, Safietou Sanke, Ousmane Faye, Amadou Alpha Sall. |
| EPI_ISL_480557 | Victorian Infectious Diseases Reference Laboratory (VIDRL) | VIDRL and MDU-PHL | Caly L., Seemann T., Sait, M., Schultz M., Druce J., Sherry, N. |
| EPI_ISL_480558, EPI_ISL_480559, EPI_ISL_480560, EPI_ISL_480561, EPI_ISL_480562 | Microbiological Diagnostic Unit - Public Health Laboratory (MDU-PHL) | MDU-PHL | Seemann T., Schultz M., Sait, M., Sherry, N. |
| EPI_ISL_480563, EPI_ISL_480564, EPI_ISL_480565, EPI_ISL_480566, EPI_ISL_480567, EPI_ISL_480568, EPI_ISL_480569, EPI_ISL_480570, EPI_ISL_480571, EPI_ISL_480572, EPI_ISL_480573, EPI_ISL_480574, EPI_ISL_480576, EPI_ISL_480577, EPI_ISL_480579, EPI_ISL_480581, EPI_ISL_480582, EPI_ISL_480583, EPI_ISL_480584, EPI_ISL_480585, EPI_ISL_480586 | | | |
| see above | Victorian Infectious Diseases Reference Laboratory (VIDRL) | VIDRL and MDU-PHL | Caly L., Seemann T., Sait, M., Schultz M., Druce J., Sherry, N. |
| EPI_ISL_480587 | Microbiological Diagnostic Unit - Public Health Laboratory (MDU-PHL) | MDU-PHL | Seemann T., Schultz M., Sait, M., Sherry, N. |
| EPI_ISL_480588, EPI_ISL_480589, EPI_ISL_480590, EPI_ISL_480591, EPI_ISL_480592, EPI_ISL_480593, EPI_ISL_480594, EPI_ISL_480595, EPI_ISL_480597, EPI_ISL_480598, EPI_ISL_480599, EPI_ISL_480600 | | | |
| see above | Victorian Infectious Diseases Reference Laboratory (VIDRL) | VIDRL and MDU-PHL | Caly L., Seemann T., Sait, M., Schultz M., Druce J., Sherry, N. |
| EPI_ISL_480601, EPI_ISL_480602 | National Health Laboratory, Timor-Leste | MDU-PHL | Soares da Silva, E., Dolores de Jesus da Costa, M., Salles de Sousa, A., Jayanti Pereira Tilman, A., Antonia da Costa, E., Barreto, I., Marr, I., Wapling, J., Francis, J., Ximenes, J., Canisia, D., Freeman, K., Dakh, F., Douglas, N., Baird, R., Caly, L., Seemann, T., Sait, M., Schultz, M., Sherry, N. |
| EPI_ISL_480603, EPI_ISL_480604, EPI_ISL_480605, EPI_ISL_480606, EPI_ISL_480607, EPI_ISL_480608, EPI_ISL_480609, EPI_ISL_480610, EPI_ISL_480611 | Victorian Infectious Diseases Reference Laboratory (VIDRL) | VIDRL and MDU-PHL | Caly L., Seemann T., Sait, M., Schultz M., Druce J., Sherry, N. |
| EPI_ISL_480612, EPI_ISL_480613, EPI_ISL_480614, EPI_ISL_480615, EPI_ISL_480616, EPI_ISL_480617, EPI_ISL_480618, EPI_ISL_480620 | Microbiological Diagnostic Unit - Public Health Laboratory (MDU-PHL) | MDU-PHL | Seemann T., Schultz M., Sait, M., Sherry, N. |
| EPI_ISL_480621, EPI_ISL_480622, EPI_ISL_480624, EPI_ISL_480625, EPI_ISL_480626, EPI_ISL_480627, EPI_ISL_480628, EPI_ISL_480629, EPI_ISL_480630, EPI_ISL_480631, EPI_ISL_480632, EPI_ISL_480633, EPI_ISL_480634, EPI_ISL_480635, EPI_ISL_480636, EPI_ISL_480637, EPI_ISL_480638, EPI_ISL_480639, EPI_ISL_480640, EPI_ISL_480641, EPI_ISL_480642, EPI_ISL_480643, EPI_ISL_480644, EPI_ISL_480645, EPI_ISL_480646, EPI_ISL_480647, EPI_ISL_480648, EPI_ISL_480649, EPI_ISL_480650, EPI_ISL_480651, EPI_ISL_480652, EPI_ISL_480653, EPI_ISL_480654, EPI_ISL_480655, EPI_ISL_480656, EPI_ISL_480657, EPI_ISL_480658, EPI_ISL_480659, EPI_ISL_480660, EPI_ISL_480661, EPI_ISL_480662, EPI_ISL_480663, EPI_ISL_480664, EPI_ISL_480665, EPI_ISL_480666, EPI_ISL_480667, EPI_ISL_480668, EPI_ISL_480669, EPI_ISL_480670, EPI_ISL_480671, EPI_ISL_480672, EPI_ISL_480673, EPI_ISL_480674, EPI_ISL_480675, EPI_ISL_480676, EPI_ISL_480677, EPI_ISL_480678, EPI_ISL_480679, EPI_ISL_480680, EPI_ISL_480681, EPI_ISL_480682, EPI_ISL_480683, EPI_ISL_480684, EPI_ISL_480685, EPI_ISL_480686 | | | |
| see above | Victorian Infectious Diseases Reference Laboratory (VIDRL) | VIDRL and MDU-PHL | Caly L., Seemann T., Sait, M., Schultz M., Druce J., Sherry, N. |
| EPI_ISL_480687 | Microbiological Diagnostic Unit - Public Health Laboratory (MDU-PHL) | MDU-PHL | Seemann T., Schultz M., Sait, M., Sherry, N. |
| EPI_ISL_480688, EPI_ISL_480689, EPI_ISL_480690 | Victorian Infectious Diseases Reference Laboratory (VIDRL) | VIDRL and MDU-PHL | Caly L., Seemann T., Sait, M., Schultz M., Druce J., Sherry, N. |
| EPI_ISL_480691, EPI_ISL_480692, EPI_ISL_480693, EPI_ISL_480694, EPI_ISL_480695, EPI_ISL_480696, EPI_ISL_480697 | Royal Darwin Hospital Pathology | MDU-PHL | Meumann, E., Caly L., Seemann T., Sait, M., Schultz M., Druce J., Sherry, N. |
| EPI_ISL_480698, EPI_ISL_480699, EPI_ISL_480700, EPI_ISL_480701, EPI_ISL_480702, EPI_ISL_480703, EPI_ISL_480704, EPI_ISL_480705, EPI_ISL_480706, EPI_ISL_480707, EPI_ISL_480708, EPI_ISL_480709, EPI_ISL_480710, EPI_ISL_480711, EPI_ISL_480712, EPI_ISL_480713, EPI_ISL_480714, EPI_ISL_480715, EPI_ISL_480716, EPI_ISL_480717, EPI_ISL_480718, EPI_ISL_480719, EPI_ISL_480720, EPI_ISL_480721, EPI_ISL_480722, EPI_ISL_480723, EPI_ISL_480724, EPI_ISL_480725, EPI_ISL_480726, EPI_ISL_480727, EPI_ISL_480728, EPI_ISL_480729, EPI_ISL_480730, EPI_ISL_480731, EPI_ISL_480732, EPI_ISL_480733, EPI_ISL_480734, EPI_ISL_480735, EPI_ISL_480736, EPI_ISL_480737, EPI_ISL_480738, EPI_ISL_480739, EPI_ISL_480740, EPI_ISL_480741, EPI_ISL_480742, EPI_ISL_480743, EPI_ISL_480744 | | | |
| see above | Victorian Infectious Diseases Reference Laboratory (VIDRL) | VIDRL and MDU-PHL | Caly L., Seemann T., Sait, M., Schultz M., Druce J., Sherry, N. |
| EPI_ISL_480745, EPI_ISL_480746, EPI_ISL_480747, EPI_ISL_480748, EPI_ISL_480749, EPI_ISL_480750, EPI_ISL_480751, EPI_ISL_480752, EPI_ISL_480753, EPI_ISL_480754, EPI_ISL_480755, EPI_ISL_480756, EPI_ISL_480757, EPI_ISL_480758, EPI_ISL_480759, EPI_ISL_480760, EPI_ISL_480761, EPI_ISL_480762, EPI_ISL_480763 | | | |
| see above | Microbiological Diagnostic Unit - Public Health Laboratory (MDU-PHL) | MDU-PHL | Seemann T., Schultz M., Sait, M., Sherry, N. |
| EPI_ISL_480764, EPI_ISL_480765, EPI_ISL_480766 | Victorian Infectious Diseases Reference Laboratory (VIDRL) | VIDRL and MDU-PHL | Caly L., Seemann T., Sait, M., Schultz M., Druce J., Sherry, N. |
| EPI_ISL_480767, EPI_ISL_480768, EPI_ISL_480769, EPI_ISL_480770, EPI_ISL_480771, EPI_ISL_480772, EPI_ISL_480773, EPI_ISL_480774, EPI_ISL_480775, EPI_ISL_480776, EPI_ISL_480777 | | | |
| see above | Microbiological Diagnostic Unit - Public Health Laboratory (MDU-PHL) | MDU-PHL | Seemann T., Schultz M., Sait, M., Sherry, N. |
| EPI_ISL_480778, EPI_ISL_480779, EPI_ISL_480780, EPI_ISL_480781 | Victorian Infectious Diseases Reference Laboratory (VIDRL) | VIDRL and MDU-PHL | Caly L., Seemann T., Sait, M., Schultz M., Druce J., Sherry, N. |
| EPI_ISL_480782, EPI_ISL_480783, EPI_ISL_480786, EPI_ISL_480787, EPI_ISL_480788, EPI_ISL_480789 | Institut Pasteur Dakar | Institut Pasteur de Dakar | Ndongo Dia, Moussa Moise Diagne, Mamadou Diop, Marie Henriette Dior Ndione, Mamadou Malado Jallow, Safietou Sanke, Ousmane Faye, Amadou Alpha Sall. |
| EPI_ISL_480790, EPI_ISL_480791, EPI_ISL_480792, EPI_ISL_480793, EPI_ISL_480794, EPI_ISL_480795, EPI_ISL_480796, EPI_ISL_480797, EPI_ISL_480798, EPI_ISL_480799, EPI_ISL_480800, EPI_ISL_480801, EPI_ISL_480802, EPI_ISL_480803, EPI_ISL_480804, EPI_ISL_480805, EPI_ISL_480806, EPI_ISL_480807, EPI_ISL_480808, EPI_ISL_480809, EPI_ISL_480810, EPI_ISL_480811, EPI_ISL_480812, EPI_ISL_480813, EPI_ISL_480814, EPI_ISL_480815, EPI_ISL_480816, EPI_ISL_480817, EPI_ISL_480818, EPI_ISL_480819, EPI_ISL_480820, EPI_ISL_480821, EPI_ISL_480822, EPI_ISL_480823, EPI_ISL_480824, EPI_ISL_480825, EPI_ISL_480826, EPI_ISL_480827, EPI_ISL_480828, EPI_ISL_480829, EPI_ISL_480830, EPI_ISL_480831, EPI_ISL_480832, EPI_ISL_480833, EPI_ISL_480834, EPI_ISL_480835, EPI_ISL_480836, EPI_ISL_480837, EPI_ISL_480838, EPI_ISL_480839, EPI_ISL_480840, EPI_ISL_480841, EPI_ISL_480842, EPI_ISL_480843, EPI_ISL_480844, EPI_ISL_480845, EPI_ISL_480846, EPI_ISL_480847, EPI_ISL_480848, EPI_ISL_480849, EPI_ISL_480850, EPI_ISL_480851, EPI_ISL_480852, EPI_ISL_480853, EPI_ISL_480854, EPI_ISL_480855, EPI_ISL_480856, EPI_ISL_480857, EPI_ISL_480858, EPI_ISL_480859, EPI_ISL_480860, EPI_ISL_480861, EPI_ISL_480862, EPI_ISL_480863, EPI_ISL_480864, EPI_ISL_480865, EPI_ISL_480866, EPI_ISL_480867, EPI_ISL_480868, EPI_ISL_480869, EPI_ISL_480870, EPI_ISL_480871, EPI_ISL_480872, EPI_ISL_480873, EPI_ISL_480874, EPI_ISL_480875, EPI_ISL_480876, EPI_ISL_480877, EPI_ISL_480878, EPI_ISL_480879, EPI_ISL_480880, EPI_ISL_480881, EPI_ISL_480882, EPI_ISL_480883, EPI_ISL_480884, EPI_ISL_480885, | | | |

| | | | |
|---|---|---|---|
| EPI_ISL_480886, EPI_ISL_480887, EPI_ISL_480888, EPI_ISL_480889, EPI_ISL_480890, EPI_ISL_480891, EPI_ISL_480892, EPI_ISL_480893, EPI_ISL_480894, EPI_ISL_480895, EPI_ISL_480896, EPI_ISL_480897, EPI_ISL_480898, EPI_ISL_480899, EPI_ISL_480900, EPI_ISL_480901, EPI_ISL_480902, EPI_ISL_480903, EPI_ISL_480904, EPI_ISL_480905, EPI_ISL_480906, EPI_ISL_480907, EPI_ISL_480908, EPI_ISL_480909, EPI_ISL_480910, EPI_ISL_480911, EPI_ISL_480912, EPI_ISL_480913, EPI_ISL_480914, EPI_ISL_480915, EPI_ISL_480916, EPI_ISL_480917, EPI_ISL_480918, EPI_ISL_480919, EPI_ISL_480920, EPI_ISL_480921, EPI_ISL_480922, EPI_ISL_480923, EPI_ISL_480924, EPI_ISL_480925, EPI_ISL_480926, EPI_ISL_480927, EPI_ISL_480928, EPI_ISL_480929, EPI_ISL_480930, EPI_ISL_480931, EPI_ISL_480932, EPI_ISL_480933, EPI_ISL_480934, EPI_ISL_480935, EPI_ISL_480936, EPI_ISL_480937, EPI_ISL_480938, EPI_ISL_480939, EPI_ISL_480940, EPI_ISL_480941, EPI_ISL_480942, EPI_ISL_480943, EPI_ISL_480944, EPI_ISL_480945, EPI_ISL_480946, EPI_ISL_480947, EPI_ISL_480948, EPI_ISL_480949, EPI_ISL_480950, EPI_ISL_480951 | | | |
| see above | Florida Bureau of Public Health Laboratories | Florida Bureau of Public Health Laboratories | Sarah Schmedes, Jason Blanton |
| EPI_ISL_480952, EPI_ISL_480953, EPI_ISL_480954, EPI_ISL_480955, EPI_ISL_480956, EPI_ISL_480957, EPI_ISL_480958, EPI_ISL_480959, EPI_ISL_480960 | Servicio de Microbiología. Hospital Universitario Donostia. OSI Donostialdea. Área de Enfermedades Infecciosas, Grupo de Infección Respiratoria y Resistencia Antimicrobiana. Instituto de Investigación Sanitaria Biodonostia | SeqCOVID-SPAIN consortium/IBV(CSIC) | Gustavo Cilla, Milagrosa Montes, Luis Piñeiro, Jose Maria Marimón and SeqCOVID-SPAIN consortium |
| EPI_ISL_480961 | ISGlobal, Institut de Salut Global de Barcelona | SeqCOVID-SPAIN consortium/IBV(CSIC) | Alfredo Mayor, Alberto L Garcia-Basteiro, Carlota Dobaño, Gemma Moncunill, Pau Cisteró and SeqCOVID-SPAIN consortium |
| EPI_ISL_480962, EPI_ISL_480963, EPI_ISL_480964, EPI_ISL_480965, EPI_ISL_480966, EPI_ISL_480967, EPI_ISL_480968, EPI_ISL_480969, EPI_ISL_480970, EPI_ISL_480971, EPI_ISL_480972, EPI_ISL_480973, EPI_ISL_480974 | | | |
| see above | Servicio de Microbiología. Hospital Universitario Donostia. OSI Donostialdea. Área de Enfermedades Infecciosas, Grupo de Infección Respiratoria y Resistencia Antimicrobiana. Instituto de Investigación Sanitaria Biodonostia | SeqCOVID-SPAIN consortium/IBV(CSIC) | Gustavo Cilla, Milagrosa Montes, Luis Piñeiro, Jose Maria Marimón and SeqCOVID-SPAIN consortium |
| EPI_ISL_480975 | ISGlobal, Institut de Salut Global de Barcelona | SeqCOVID-SPAIN consortium/IBV(CSIC) | Alfredo Mayor, Alberto L Garcia-Basteiro, Carlota Dobaño, Gemma Moncunill, Pau Cisteró and SeqCOVID-SPAIN consortium |
| EPI_ISL_480976, EPI_ISL_480977, EPI_ISL_480978, EPI_ISL_480979, EPI_ISL_480980 | Servicio de Microbiología. Hospital Universitario Donostia. OSI Donostialdea. Área de Enfermedades Infecciosas, Grupo de Infección Respiratoria y Resistencia Antimicrobiana. Instituto de Investigación Sanitaria Biodonostia | SeqCOVID-SPAIN consortium/IBV(CSIC) | Gustavo Cilla, Milagrosa Montes, Luis Piñeiro, Jose Maria Marimón and SeqCOVID-SPAIN consortium |
| EPI_ISL_480981 | ISGlobal, Institut de Salut Global de Barcelona | SeqCOVID-SPAIN consortium/IBV(CSIC) | Alfredo Mayor, Alberto L Garcia-Basteiro, Carlota Dobaño, Gemma Moncunill, Pau Cisteró and SeqCOVID-SPAIN consortium |
| EPI_ISL_480982, EPI_ISL_480983, EPI_ISL_480984, EPI_ISL_480985, EPI_ISL_480986, EPI_ISL_480987, EPI_ISL_480988 | Servicio de Microbiología. Hospital Universitario Donostia. OSI Donostialdea. Área de Enfermedades Infecciosas, Grupo de Infección Respiratoria y Resistencia Antimicrobiana. Instituto de Investigación Sanitaria Biodonostia | SeqCOVID-SPAIN consortium/IBV(CSIC) | Gustavo Cilla, Milagrosa Montes, Luis Piñeiro, Jose Maria Marimón and SeqCOVID-SPAIN consortium |
| EPI_ISL_480989 | ISGlobal, Institut de Salut Global de Barcelona | SeqCOVID-SPAIN consortium/IBV(CSIC) | Alfredo Mayor, Alberto L Garcia-Basteiro, Carlota Dobaño, Gemma Moncunill, Pau Cisteró and SeqCOVID-SPAIN consortium |
| EPI_ISL_480990, EPI_ISL_480991, EPI_ISL_480992, EPI_ISL_480993, EPI_ISL_480994 | Servicio de Microbiología. Hospital Universitario Donostia. OSI Donostialdea. Área de Enfermedades Infecciosas, Grupo de Infección Respiratoria y Resistencia Antimicrobiana. Instituto de Investigación Sanitaria Biodonostia | SeqCOVID-SPAIN consortium/IBV(CSIC) | Gustavo Cilla, Milagrosa Montes, Luis Piñeiro, Jose Maria Marimón and SeqCOVID-SPAIN consortium |
| EPI_ISL_480995 | ISGlobal, Institut de Salut Global de Barcelona | SeqCOVID-SPAIN consortium/IBV(CSIC) | Alfredo Mayor, Alberto L Garcia-Basteiro, Carlota Dobaño, Gemma Moncunill, Pau Cisteró and SeqCOVID-SPAIN consortium |
| EPI_ISL_480996, EPI_ISL_480997, EPI_ISL_480998, EPI_ISL_480999, EPI_ISL_481000, EPI_ISL_481001, EPI_ISL_481002 | Servicio de Microbiología. Hospital Universitario Donostia. OSI Donostialdea. Área de Enfermedades Infecciosas, Grupo de Infección Respiratoria y Resistencia Antimicrobiana. Instituto de Investigación Sanitaria Biodonostia | SeqCOVID-SPAIN consortium/IBV(CSIC) | Gustavo Cilla, Milagrosa Montes, Luis Piñeiro, Jose Maria Marimón and SeqCOVID-SPAIN consortium |
| EPI_ISL_481003 | ISGlobal, Institut de Salut Global de Barcelona | SeqCOVID-SPAIN consortium/IBV(CSIC) | Alfredo Mayor, Alberto L Garcia-Basteiro, Carlota Dobaño, Gemma Moncunill, Pau Cisteró and SeqCOVID-SPAIN consortium |
| EPI_ISL_481004, EPI_ISL_481005, EPI_ISL_481006, EPI_ISL_481007, EPI_ISL_481008, EPI_ISL_481009, EPI_ISL_481010, EPI_ISL_481011, EPI_ISL_481012, EPI_ISL_481013, EPI_ISL_481014, EPI_ISL_481015, EPI_ISL_481016 | | | |
| see above | Servicio de Microbiología. Hospital Universitario Donostia. OSI Donostialdea. Área de Enfermedades Infecciosas, Grupo de Infección Respiratoria y Resistencia Antimicrobiana. Instituto de Investigación Sanitaria Biodonostia | SeqCOVID-SPAIN consortium/IBV(CSIC) | Gustavo Cilla, Milagrosa Montes, Luis Piñeiro, Jose Maria Marimón and SeqCOVID-SPAIN consortium |
| EPI_ISL_481017 | ISGlobal, Institut de Salut Global de Barcelona | SeqCOVID-SPAIN consortium/IBV(CSIC) | Alfredo Mayor, Alberto L Garcia-Basteiro, Carlota Dobaño, Gemma Moncunill, Pau Cisteró and SeqCOVID-SPAIN consortium |
| EPI_ISL_481018, EPI_ISL_481019, EPI_ISL_481020, EPI_ISL_481021, EPI_ISL_481022, EPI_ISL_481023, EPI_ISL_481024 | Servicio de Microbiología. Hospital Universitario Donostia. OSI Donostialdea. Área de Enfermedades Infecciosas, Grupo de Infección Respiratoria y Resistencia Antimicrobiana. Instituto de Investigación Sanitaria Biodonostia | SeqCOVID-SPAIN consortium/IBV(CSIC) | Gustavo Cilla, Milagrosa Montes, Luis Piñeiro, Jose Maria Marimón and SeqCOVID-SPAIN consortium |
| EPI_ISL_481025 | ISGlobal, Institut de Salut Global de Barcelona | SeqCOVID-SPAIN consortium/IBV(CSIC) | Alfredo Mayor, Alberto L Garcia-Basteiro, Carlota Dobaño, Gemma Moncunill, Pau Cisteró and SeqCOVID-SPAIN consortium |
| EPI_ISL_481026, EPI_ISL_481027, EPI_ISL_481028 | Servicio de Microbiología. Hospital Universitario Donostia. OSI Donostialdea. Área de Enfermedades Infecciosas, Grupo de Infección Respiratoria y Resistencia Antimicrobiana. Instituto de Investigación Sanitaria Biodonostia | SeqCOVID-SPAIN consortium/IBV(CSIC) | Gustavo Cilla, Milagrosa Montes, Luis Piñeiro, Jose Maria Marimón and SeqCOVID-SPAIN consortium |
| EPI_ISL_481029 | ISGlobal, Institut de Salut Global de Barcelona | SeqCOVID-SPAIN consortium/IBV(CSIC) | Alfredo Mayor, Alberto L Garcia-Basteiro, Carlota Dobaño, Gemma Moncunill, Pau Cisteró and SeqCOVID-SPAIN consortium |
| EPI_ISL_481030, EPI_ISL_481031, EPI_ISL_481032, EPI_ISL_481033 | Servicio de Microbiología. Hospital Universitario Donostia. OSI Donostialdea. Área de Enfermedades Infecciosas, Grupo de Infección Respiratoria y Resistencia Antimicrobiana. Instituto de Investigación Sanitaria Biodonostia | SeqCOVID-SPAIN consortium/IBV(CSIC) | Gustavo Cilla, Milagrosa Montes, Luis Piñeiro, Jose Maria Marimón and SeqCOVID-SPAIN consortium |
| EPI_ISL_481034, EPI_ISL_481035 | ISGlobal, Institut de Salut Global de Barcelona | SeqCOVID-SPAIN consortium/IBV(CSIC) | Alfredo Mayor, Alberto L Garcia-Basteiro, Carlota Dobaño, Gemma Moncunill, Pau Cisteró and SeqCOVID-SPAIN consortium |
| EPI_ISL_481036, EPI_ISL_481037, EPI_ISL_481038, EPI_ISL_481039, EPI_ISL_481040 | Servicio de Microbiología. Hospital Universitario Donostia. OSI Donostialdea. Área de Enfermedades Infecciosas, Grupo de Infección Respiratoria y Resistencia Antimicrobiana. Instituto de Investigación Sanitaria Biodonostia | SeqCOVID-SPAIN consortium/IBV(CSIC) | Gustavo Cilla, Milagrosa Montes, Luis Piñeiro, Jose Maria Marimón and SeqCOVID-SPAIN consortium |
| EPI_ISL_481041, EPI_ISL_481042, EPI_ISL_481043, EPI_ISL_481044, EPI_ISL_481045, EPI_ISL_481046, EPI_ISL_481047, EPI_ISL_481048, EPI_ISL_481049, EPI_ISL_481050, EPI_ISL_481051, EPI_ISL_481052, EPI_ISL_481053, EPI_ISL_481054, EPI_ISL_481055, EPI_ISL_481056, EPI_ISL_481057, EPI_ISL_481058, EPI_ISL_481059, EPI_ISL_481060, EPI_ISL_481061, EPI_ISL_481062, EPI_ISL_481063, EPI_ISL_481064, EPI_ISL_481065, EPI_ISL_481066, EPI_ISL_481067, EPI_ISL_481068, EPI_ISL_481069, EPI_ISL_481070, EPI_ISL_481071, EPI_ISL_481072, EPI_ISL_481073, EPI_ISL_481074, EPI_ISL_481075, EPI_ISL_481076, EPI_ISL_481077, EPI_ISL_481078, EPI_ISL_481079, EPI_ISL_481080, EPI_ISL_481081, EPI_ISL_481082, EPI_ISL_481083, EPI_ISL_481084, EPI_ISL_481085, EPI_ISL_481086, EPI_ISL_481087, EPI_ISL_481088, EPI_ISL_481089, EPI_ISL_481090, EPI_ISL_481091, EPI_ISL_481092, EPI_ISL_481093, EPI_ISL_481094, EPI_ISL_481095, EPI_ISL_481096, EPI_ISL_481097, EPI_ISL_481098, EPI_ISL_481099, EPI_ISL_481100, EPI_ISL_481101, EPI_ISL_481102, EPI_ISL_481103, EPI_ISL_481104, EPI_ISL_481105, EPI_ISL_481106, EPI_ISL_481107, EPI_ISL_481108, EPI_ISL_481109 | | | |
| see above | Hospital General Universitario Gregorio Marañón | SeqCOVID-SPAIN consortium/IBV(CSIC) | Laura Pérez-Lago, Marta Herranz, Jon Sicilia, Julia Suárez, Pilar Catalán, Patricia Muñoz, Darío García de Viedma and SeqCOVID-SPAIN consortium |
| EPI_ISL_481110, EPI_ISL_481111, EPI_ISL_481112, EPI_ISL_481113, EPI_ISL_481114, EPI_ISL_481115, EPI_ISL_481116, EPI_ISL_481117, EPI_ISL_481118, EPI_ISL_481119, EPI_ISL_481120, EPI_ISL_481121, EPI_ISL_481122, EPI_ISL_481123, EPI_ISL_481124, EPI_ISL_481125, EPI_ISL_481126, EPI_ISL_481127, EPI_ISL_481128, EPI_ISL_481129, EPI_ISL_481130, EPI_ISL_481131, EPI_ISL_481132, EPI_ISL_481133 | | | |
| see above | Immunogenomics lab, Institute of Life Sciences, Bhubaneswar | Immunogenomics lab, Institute of Life Sciences, Bhubaneswar | Sunil Raghav, Arup Ghosh, Deepika Singh, Ankita Datey, P. Sushree Shyamli, Bharati Singh, Neha Singh, Atimukta Jha, Viplov K. Biswas, Swati Madhulika, Manasi Priyadarshini, Sneha Dutta, Auromira Khuntia, Rupesh Dash, Soma Chattopadhyay, Ghulam Hussain Syed, Shanti Senapati, Tushar K. Beuria, Rajeeb Swain, Punit Prasad, Orissa COVID-19 Study Group, DBT's PAN-INDIA 1000 SARS-CoV2 RNA genome sequencing consortium, Ajay Parida |
| EPI_ISL_481134, EPI_ISL_481135, EPI_ISL_481136, EPI_ISL_481137, EPI_ISL_481138, EPI_ISL_481139, EPI_ISL_481140, EPI_ISL_481141, EPI_ISL_481142, EPI_ISL_481143, EPI_ISL_481144, EPI_ISL_481145, EPI_ISL_481146, EPI_ISL_481147, EPI_ISL_481148, EPI_ISL_481149, EPI_ISL_481150, EPI_ISL_481151, EPI_ISL_481152, EPI_ISL_481153, EPI_ISL_481154, EPI_ISL_481155, EPI_ISL_481156, EPI_ISL_481157 | | | |
| see above | Immunogenomics lab, Institute of Life Sciences, Bhubaneswar | Immunogenomics lab, Institute of Life Sciences, Bhubaneswar | Sunil Raghav, Arup Ghosh, Ankita Datey, P. Sushree Shyamli, Bharati Singh, Neha Singh, Deepika Singh, Atimukta Jha, Viplov K. Biswas, Swati Madhulika, Manasi Priyadarshini, Aditi Chatterjee, Rahul Das, Soumyajit Ghosh, Rupesh Dash, Soma Chattopadhyay, Ghulam Hussain Syed, Shanti Senapati, Tushar K. Beuria, Rajeeb Swain, Punit Prasad, Amol Ratnakar Suryawanshi, Dileep Vasudeva, Orissa COVID-19 Study Group, DBT's PAN-INDIA 1000 SARS-CoV2 RNA genome sequencing consortium, Ajay Parida |
| EPI_ISL_481158, EPI_ISL_481159, EPI_ISL_481160, EPI_ISL_481161, EPI_ISL_481162, EPI_ISL_481163, EPI_ISL_481164, EPI_ISL_481165, EPI_ISL_481166, EPI_ISL_481167, EPI_ISL_481168, EPI_ISL_481169, EPI_ISL_481170, EPI_ISL_481171, EPI_ISL_481172, EPI_ISL_481173, EPI_ISL_481174, EPI_ISL_481175, EPI_ISL_481176, EPI_ISL_481177, EPI_ISL_481178, EPI_ISL_481179, EPI_ISL_481180, EPI_ISL_481181 | | | |
| see above | Immunogenomics lab, Institute of Life Sciences, Bhubaneswar | Immunogenomics lab, Institute of Life Sciences, Bhubaneswar | Sunil Raghav, Arup Ghosh, P. Sushree Shyamli, Bharati Singh, Neha Singh, Ankita Datey, Deepika Singh, Atimukta Jha, Viplov K. Biswas, Swati Madhulika, Manasi Priyadarshini, Tsheten Sheroa, Auromira Khuntia, Rupesh Dash, Soma Chattopadhyay, Ghulam Hussain Syed, Shanti Senapati, Tushar K. Beuria, Rajeeb Swain, Punit Prasad, Amol Ratnakar Suryawanshi, Dileep Vasudevan, Orissa COVID-19 Study Group, DBT's PAN-INDIA 1000 SARS-CoV2 RNA genome sequencing consortium, Ajay Parida |
| EPI_ISL_481182, EPI_ISL_481183, EPI_ISL_481184, EPI_ISL_481185, EPI_ISL_481186, EPI_ISL_481187, EPI_ISL_481188, EPI_ISL_481189, EPI_ISL_481190, EPI_ISL_481191, EPI_ISL_481192, EPI_ISL_481193, EPI_ISL_481194, EPI_ISL_481195, EPI_ISL_481196, EPI_ISL_481197, EPI_ISL_481198, EPI_ISL_481199, EPI_ISL_481200, EPI_ISL_481201, EPI_ISL_481202, EPI_ISL_481203, EPI_ISL_481204, EPI_ISL_481205 | | | |
| see above | Immunogenomics lab, Institute of Life Sciences, Bhubaneswar | Immunogenomics lab, Institute of Life Sciences, Bhubaneswar | Sunil Raghav, Arup Ghosh, Atimukta Jha, Viplov K. Biswas, Swati Madhulika, Manasi Priyadarshini, Ajit Singh, Sivaram Krishna, Naga Jogayya Kothakota, Rupesh Dash, Soma Chattopadhyay, Ghulam Hussain Syed, Shanti Senapati, Tushar K. Beuria, Rajeeb Swain, Punit Prasad, Amol Ratnakar Suryawanshi, Dileep Vasudevan, Orissa COVID-19 Study Group, DBT's PAN-INDIA 1000 SARS-CoV2 RNA genome sequencing consortium, Ajay Parida |
| EPI_ISL_481206 | Hospital of Southern Norway - Kristiansand, Department of Medical Microbiology | Norwegian Institute of Public Health, Department of Virology | Kathrine Stene-Johansen, Kamilla Heddeland Instefjord, Hilde Elshaug, Rasmus Riis Kopperud, Karoline Bragstad, Olav Hungnes |
| EPI_ISL_481207 | Ostfold Hospital Trust - Kalnes, Centre for Laboratory Medicine, Section for gene technology and infection serology | Norwegian Institute of Public Health, Department of Virology | Kathrine Stene-Johansen, Kamilla Heddeland Instefjord, Hilde Elshaug, Rasmus Riis Kopperud, Karoline Bragstad, Olav Hungnes |
| EPI_ISL_481208 | Furst Medical Laboratory | Norwegian Institute of Public Health, | Kathrine Stene-Johansen, Kamilla Heddeland Instefjord, Hilde Elshaug, Rasmus Riis Kopperud, Karoline Bragstad, Olav Hungnes |

| | | | |
|---|---|---|---|
| | | Department of Virology | |
| EPI_ISL_481209, EPI_ISL_481210, EPI_ISL_481211, EPI_ISL_481212, EPI_ISL_481213 | Ostfold Hospital Trust - Kalnes, Centre for Laboratory Medicine, Section for gene technology and infection serology | Norwegian Institute of Public Health, Department of Virology | Kathrine Stene-Johansen, Kamilla Heddeland Instefjord, Hilde Elshaug, Rasmus Riis Kopperud, Karoline Bragstad, Olav Hungnes |
| EPI_ISL_481214, EPI_ISL_481215 | Oslo University Hospital, Department of Medical Microbiology | Norwegian Institute of Public Health, Department of Virology | Kathrine Stene-Johansen, Kamilla Heddeland Instefjord, Hilde Elshaug, Rasmus Riis Kopperud, Karoline Bragstad, Olav Hungnes |
| EPI_ISL_481216 | Medical Microbiology Unit, Department for Laboratory Medicine, Drammen Hospital, Vestre Viken Health Trust, | Norwegian Institute of Public Health, Department of Virology | Kathrine Stene-Johansen, Kamilla Heddeland Instefjord, Hilde Elshaug, Rasmus Riis Kopperud, Karoline Bragstad, Olav Hungnes |
| EPI_ISL_481217, EPI_ISL_481218, EPI_ISL_481219 | Oslo University Hospital, Department of Medical Microbiology | Norwegian Institute of Public Health, Department of Virology | Kathrine Stene-Johansen, Kamilla Heddeland Instefjord, Hilde Elshaug, Rasmus Riis Kopperud, Karoline Bragstad, Olav Hungnes |
| EPI_ISL_481220 | Institut Pasteur Dakar | Institut Pasteur de Dakar | Ndongo Dia, Moussa Moise Diagne, Mamadou Diop, Marie Henriette Dior Ndione, Mamadou Malado Jallow, Safietou Sanke, Ousmane Faye, Amadou Alpha Sall. |
| EPI_ISL_481221, EPI_ISL_481222, EPI_ISL_481223, EPI_ISL_481224, EPI_ISL_481225, EPI_ISL_481226 | Lab voor klinische biologie | Onderzoeksgroep Virologie | Laurens Lambrechts, Nick Vereecke, Marthe Pauwels, Bruno Verhasselt, Linos Vandekerckhove, Hans Nauwynck, Sebastiaan Theuns |
| EPI_ISL_481227, EPI_ISL_481228, EPI_ISL_481229, EPI_ISL_481230, EPI_ISL_481231, EPI_ISL_481232, EPI_ISL_481233 | Lab voor klinische biologie | Onderzoeksgroep Virologie | Nick Vereecke, Laurens Lambrechts, Marthe Pauwels, Bruno Verhasselt, Linos Vandekerckhove, Hans Nauwynck, Sebastiaan Theuns |
| EPI_ISL_481234, EPI_ISL_481235, EPI_ISL_481236, EPI_ISL_481237, EPI_ISL_481238, EPI_ISL_481239, EPI_ISL_481240 | Institut Pasteur Dakar | Institut Pasteur de Dakar | Ndongo Dia, Moussa Moise Diagne, Mamadou Diop, Marie Henriette Dior Ndione, Mamadou Malado Jallow, Safietou Sanke, Ousmane Faye, Amadou Alpha Sall. |
| EPI_ISL_481241 | M Health Fairview | Minnesota Department of Health, Public Health Laboratory | Matt Plumb, Jacob Garfin, Kelly Pung, and Xiong Wang |
| EPI_ISL_481242 | Mayo Clinic & Mayo Clinic Laboratories | Minnesota Department of Health, Public Health Laboratory | Matt Plumb, Jacob Garfin, Kelly Pung, and Xiong Wang |
| EPI_ISL_481243 | Institut Pasteur Dakar | Institut Pasteur de Dakar | Ndongo Dia, Moussa Moise Diagne, Mamadou Diop, Marie Henriette Dior Ndione, Mamadou Malado Jallow, Safietou Sanke, Ousmane Faye, Amadou Alpha Sall. |
| EPI_ISL_481244, EPI_ISL_481245, EPI_ISL_481246, EPI_ISL_481247, EPI_ISL_481248 | Hospital IESS Babahoyo | Institute of Microbiology, Universidad San Francisco de Quito | Belén Prado-Vivar, Sully Márquez, Juan José Guadalupe, Monica Becerra-Wong, Carla Torres, Bernardo Gutiérrez, Francisco Cordova, Ninfa Henriquez, Killen Briones-Zamora, Killen Briones-Claudette, Verónica Barragán, Patricio Rojas-Silva, Gabriel Trueba, Michelle Grunauer, Paúl Cárdenas |
| EPI_ISL_481251, EPI_ISL_481252 | Department of Emerging Infectious Diseases, Institute of Tropical Medicine, Nagasaki University | Department of Emerging Infectious Diseases, Institute of Tropical Medicine, Nagasaki University | Jiro Yasuda, Rokusuke Yoshikawa, Yuichiro Furusato, Haruka Abe |
| EPI_ISL_481253 | Robert Koch Institute, National Reference center for Influenza, Berlin, Germany | Robert Koch Institute, Bioinformatics MF1, Berlin, Germany | Marianne Wedde, Oliver Drechsel, Andrea Thuermer, Rene Kmiecinski, Ralf Duerrwald, Thorsten Wolff, Stephan Fuchs, Max v. Kleist |
| EPI_ISL_481254, EPI_ISL_481255 | Department of Emerging Infectious Diseases, Institute of Tropical Medicine, Nagasaki University | Department of Emerging Infectious Diseases, Institute of Tropical Medicine, Nagasaki University | Jiro Yasuda, Rokusuke Yoshikawa, Yuichiro Furusato, Haruka Abe |
| EPI_ISL_481256 | Robert Koch Institute, National Reference center for Influenza, Berlin, Germany | Robert Koch Institute, Bioinformatics MF1, Berlin, Germany | Marianne Wedde, Oliver Drechsel, Andrea Thuermer, Rene Kmiecinski, Ralf Duerrwald, Thorsten Wolff, Stephan Fuchs, Max v. Kleist |
| EPI_ISL_481257, EPI_ISL_481258, EPI_ISL_481259, EPI_ISL_481260, EPI_ISL_481261 | Department of Emerging Infectious Diseases, Institute of Tropical Medicine, Nagasaki University | Department of Emerging Infectious Diseases, Institute of Tropical Medicine, Nagasaki University | Jiro Yasuda, Rokusuke Yoshikawa, Yuichiro Furusato, Haruka Abe |
| EPI_ISL_481262 | Robert Koch Institute, National Reference center for Influenza, Berlin, Germany | Robert Koch Institute, Bioinformatics MF1, Berlin, Germany | Marianne Wedde, Oliver Drechsel, Andrea Thuermer, Rene Kmiecinski, Ralf Duerrwald, Thorsten Wolff, Stephan Fuchs, Max v. Kleist |
| EPI_ISL_481263 | Department of Emerging Infectious Diseases, Institute of Tropical Medicine, Nagasaki University | Department of Emerging Infectious Diseases, Institute of Tropical Medicine, Nagasaki University | Jiro Yasuda, Rokusuke Yoshikawa, Yuichiro Furusato, Haruka Abe |
| EPI_ISL_481264 | Robert Koch Institute, National Reference center for Influenza, Berlin, Germany | Robert Koch Institute, Bioinformatics MF1, Berlin, Germany | Marianne Wedde, Oliver Drechsel, Andrea Thuermer, Rene Kmiecinski, Ralf Duerrwald, Thorsten Wolff, Stephan Fuchs, Max v. Kleist |
| EPI_ISL_481265, EPI_ISL_481266, EPI_ISL_481267, EPI_ISL_481268, EPI_ISL_481269, EPI_ISL_481270, EPI_ISL_481271, EPI_ISL_481272, EPI_ISL_481273, EPI_ISL_481274, EPI_ISL_481275, EPI_ISL_481276, EPI_ISL_481277, EPI_ISL_481278, EPI_ISL_481279, EPI_ISL_481280, EPI_ISL_481281, EPI_ISL_481282 | | | |
| see above | unknown | Core Sequencing, Maryland Department of Health | Keller,E. |
| EPI_ISL_481283 | unknown | Center for Genomics and System Biology | Roder,A., Banakis,S., Johnson,K., Khalfan,M., Borenstein,E.S., Samanovic,M., Cornelius,A., Herati,R., Ulrich,R., Fleming,A., Kottkamp,A., Raabe,V., Mulligan,M.J., Gresham,D. and Ghedin,E. |
| EPI_ISL_481284 | unknown | Department of Experimental Modeling and Pathogenesis of Infectious Diseases | Sobolev,I.A., Shanshin,D.V., Chepurnov,A.A., Kononova,J.V., Bondar,A.A., Alekseev,A.Y. and Shestopalov,A.M. |
| EPI_ISL_481370 | Division of Viral Diseases, Center for Laboratory Control of Infectious Diseases, Korea Centers for Diseases Control and Prevention | Division of Viral Diseases, Center for Laboratory Control of Infectious Diseases, Korea Centers for Diseases Control and Prevention | Jeong-Min Kim, Yoon-Seok Chung, Namjoo Lee, Sang Hee Woo, Hye-Jun Jo, Heui Man Kim, Jun-Sub Kim, Myung Guk Han |
| EPI_ISL_481371, EPI_ISL_481372, EPI_ISL_481373, EPI_ISL_481374, EPI_ISL_481375, EPI_ISL_481376, EPI_ISL_481377, EPI_ISL_481378, EPI_ISL_481379 | Division of Viral Diseases, Center for Laboratory Control of Infectious Diseases, Korea Centers for Diseases Control and Prevention | Division of Viral Diseases, Center for Laboratory Control of Infectious Diseases, Korea Centers for Diseases Control and Prevention | Jeong-Min Kim, Yoon-Seok Chung, Namjoo Lee, Sang Hee Woo, Hye-Jun Jo, Heui Man Kim, Jun-Sub Kim, Dong Hyun Song, Daesang Lee, Seong Tae Jeong, Myung Guk Han |
| EPI_ISL_481380 | Department for Virology, Molecular Biology and Genome Research, R. G. Lugar Center for Public Health Research, National Center for Disease Control and Public Health (NCDC) of Georgia. | Department for Virology, Molecular Biology and Genome Research, R. G. Lugar Center for Public Health Research, National Center for Disease Control and Public Health (NCDC) of Georgia. | Ana Papkiauri, Tata Imnadze, Giorgi Tomashvili, Meri Pantsulaia, Gvantsa Brachveli, Gvantsa Chanturia, Ann Machablishvili, Nato Kotaria, Marine Murtskhvaladze, Lela Sabadze, Mari Gavashelidze, Tamar Jashiashvili, Tea Tevdoradze, Ketevan Sidamonidze, Ekaterine Khmaladze, Ekaterine Zhghenti, Roena Sukhiashvili, Mariam Zakalashvili, Lela Urushadze, Magda Dgebuadze, Davit Tsaguria, Ekaterine Zangaladze, Nino Berishvili, Adam Kotorashvili, Maia Alkhazashvili, Irma Burjanadze, Anna Kasradze, Khatuna Zakhashvili, Paata Imnadze, Amiran Gamkrelidze. |
| EPI_ISL_481483 | Department for Virology, Molecular Biology and Genome Research, R. G. Lugar Center for Public Health Research, National Center for Disease Control and Public Health (NCDC) of Georgia. | Department for Virology, Molecular Biology and Genome Research, R. G. Lugar Center for Public Health Research, National Center for Disease Control and Public Health (NCDC) of Georgia. | Nino Berishvili, Tata Imnadze, Giorgi Tomashvili, Ana Papkiauri, Meri Pantsulaia, Gvantsa Brachveli, Gvantsa Chanturia, Ann Machablishvili, Nato Kotaria, Marine Murtskhvaladze, Lela Sabadze, Mari Gavashelidze, Tamar Jashiashvili, Tea Tevdoradze, Ketevan Sidamonidze, Ekaterine Khmaladze, Ekaterine Zhghenti, Roena Sukhiashvili, Mariam Zakalashvili, Lela Urushadze, Magda Dgebuadze, Davit Tsaguria, Ekaterine Zangaladze, Adam Kotorashvili, Maia Alkhazashvili, Irma Burjanadze, Anna Kasradze, Khatuna Zakhashvili, Paata Imnadze, Amiran Gamkrelidze. |
| EPI_ISL_481510, EPI_ISL_481511 | Prof. Massimo Zollo CEINGE TASK-FORCE COVID19 - Regione Campania | Prof. Massimo Zollo CEINGE TASK-FORCE COVID19 - Regione Campania | Veronica Ferrucci1,2, Dae young Kong8, Fatemeh asadzadeh1,2, Laura Marrone1,2, Roberto Siciliano1,2, Rino Cerino3, Giovanna Fusco3, Marika Comegna1,2, Angelo Boccia2, Maurizio Viscardi3, Giorgia Borriello3, Sergio Brandi3, Claudia Tiberio4, Luigi Atripaldi4, Giovanni Paolella1,2, Giuseppe Castaldo1,2, Stefano Pascarella4, Martina Bianchi4, Lorenzo Chiariotti1,2, Jae Myun Lee5, Jae Ho Jung6, Kyong Seop Yun7, Hong Yeoul Kim 7,8* and Massimo Zollo1,2* 1 CEINGE Biotecnologie Avanzate, Naples, Italia 2 Dipartimento di Medicina Molecolare e Biotecnologie Mediche DMMBM University of Naples Federico II, Italia 3 Istituto Zooprofilattico Sperimentale del Mezzogiorno, Naples, Italia 4 -U.O.C. di Patologia Clinica Ospedale D. Cotugno, Azienda Sanitaria Ospedali dei Colli, Naples, Italy. 5 Università La Sapienza di Roma, Italia 6 Department of Microbiology, Yonsei University College of Medicine, Seoul, Korea 7 Department of Surgery, Yonsei University College of Medicine, Seoul, Korea 8 Haim bio co., Ltd, , Indust |
| EPI_ISL_481512 | Prof. Massimo Zollo CEINGE TASK-FORCE COVID19 - Regione Campania | Prof. Massimo Zollo CEINGE TASK-FORCE COVID19 - Regione Campania | Veronica Ferrucci1,2, Dae young Kong8, Fatemeh asadzadeh1,2, Laura Marrone1,2, Roberto Siciliano1,2, Rino Cerino3, Giovanna Fusco3, Marika Comegna1,2, Angelo Boccia2, Maurizio Viscardi3, Giorgia Borriello3, Sergio Brandi3, Claudia Tiberio4, Luigi Atripaldi4, Giovanni Paolella1,2, Giuseppe Castaldo1,2, Stefano Pascarella4, Martina Bianchi4, Lorenzo Chiariotti1,2, Jae Myun Lee5, Jae Ho Jung6, Kyong Seop Yun7, Hong Yeoul Kim 7,8* and Massimo Zollo1,2* 1 CEINGE Biotecnologie Avanzate, Naples, Italia 2 Dipartimento di Medicina Molecolare e Biotecnologie Mediche DMMBM University of Naples Federico II, Italia 3 Istituto Zooprofilattico Sperimentale del Mezzogiorno, Naples, Italia 4 -U.O.C. di Patologia Clinica Ospedale D. Cotugno, Azienda Sanitaria Ospedali dei Colli, Naples, Italy. 5 Università La Sapienza di Roma, Italia 6 Department of Microbiology, Yonsei University College of Medicine, Seoul, Korea 7 Department of Surgery, Yonsei University College of Medicine, Seoul, Korea 8 Haim bio co., Ltd, , Indust |
| EPI_ISL_481513, EPI_ISL_481514, EPI_ISL_481515, EPI_ISL_481516, EPI_ISL_481517, EPI_ISL_481518, EPI_ISL_481519, EPI_ISL_481520, EPI_ISL_481521, EPI_ISL_481522, EPI_ISL_481523, EPI_ISL_481524, EPI_ISL_481525, EPI_ISL_481526, EPI_ISL_481527, EPI_ISL_481528, EPI_ISL_481529, EPI_ISL_481530, EPI_ISL_481531, EPI_ISL_481532, EPI_ISL_481533, EPI_ISL_481534, EPI_ISL_481535, EPI_ISL_481536, EPI_ISL_481537, EPI_ISL_481538, EPI_ISL_481539, EPI_ISL_481540, EPI_ISL_481541, EPI_ISL_481542, EPI_ISL_481543, EPI_ISL_481544, EPI_ISL_481545, EPI_ISL_481546, EPI_ISL_481547, EPI_ISL_481548, EPI_ISL_481549, EPI_ISL_481550, EPI_ISL_481551, EPI_ISL_481552, EPI_ISL_481553, EPI_ISL_481554, EPI_ISL_481555, EPI_ISL_481556, EPI_ISL_481557, EPI_ISL_481558, EPI_ISL_481559, EPI_ISL_481560, EPI_ISL_481561, EPI_ISL_481562, EPI_ISL_481563, EPI_ISL_481564, EPI_ISL_481565, EPI_ISL_481566, EPI_ISL_481567, EPI_ISL_481568, EPI_ISL_481569, EPI_ISL_481570, EPI_ISL_481571, EPI_ISL_481572, EPI_ISL_481573, EPI_ISL_481574, EPI_ISL_481575, EPI_ISL_481576, EPI_ISL_481577, EPI_ISL_481578, EPI_ISL_481579, EPI_ISL_481580, EPI_ISL_481581, EPI_ISL_481582, EPI_ISL_481583, EPI_ISL_481584, EPI_ISL_481585, EPI_ISL_481586, EPI_ISL_481587, EPI_ISL_481588, EPI_ISL_481589, EPI_ISL_481590, EPI_ISL_481591, EPI_ISL_481592, EPI_ISL_481593, EPI_ISL_481594, EPI_ISL_481595, EPI_ISL_481596, EPI_ISL_481597, EPI_ISL_481598, EPI_ISL_481599, EPI_ISL_481600, EPI_ISL_481601, EPI_ISL_481602, EPI_ISL_481603, EPI_ISL_481604, EPI_ISL_481605, EPI_ISL_481606, EPI_ISL_481607, EPI_ISL_481608, EPI_ISL_481609, EPI_ISL_481610, EPI_ISL_481611, EPI_ISL_481612, EPI_ISL_481613, EPI_ISL_481614, EPI_ISL_481615, EPI_ISL_481616, EPI_ISL_481617, EPI_ISL_481618, EPI_ISL_481619, EPI_ISL_481620, EPI_ISL_481621, EPI_ISL_481622, EPI_ISL_481623, EPI_ISL_481624, EPI_ISL_481625, EPI_ISL_481626, EPI_ISL_481627, EPI_ISL_481628, EPI_ISL_481629, EPI_ISL_481630, EPI_ISL_481631, EPI_ISL_481632, EPI_ISL_481633, EPI_ISL_481634, EPI_ISL_481635, EPI_ISL_481636, EPI_ISL_481637, EPI_ISL_481638, EPI_ISL_481639, EPI_ISL_481640, EPI_ISL_481641, EPI_ISL_481642, EPI_ISL_481643, EPI_ISL_481644, EPI_ISL_481645, EPI_ISL_481646, EPI_ISL_481647, EPI_ISL_481648, EPI_ISL_481649, EPI_ISL_481650, EPI_ISL_481651, EPI_ISL_481652, EPI_ISL_481653, EPI_ISL_481654, EPI_ISL_481655, EPI_ISL_481656, EPI_ISL_481657, EPI_ISL_481658, EPI_ISL_481659, EPI_ISL_481660, EPI_ISL_481661, EPI_ISL_481662, EPI_ISL_481663, EPI_ISL_481664, EPI_ISL_481665, EPI_ISL_481666, EPI_ISL_481667, EPI_ISL_481668, EPI_ISL_481669, EPI_ISL_481670, EPI_ISL_481671, EPI_ISL_481672, EPI_ISL_481673, EPI_ISL_481674, EPI_ISL_481675, EPI_ISL_481676, EPI_ISL_481677, EPI_ISL_481678, EPI_ISL_481679, EPI_ISL_481680, EPI_ISL_481681, EPI_ISL_481682, EPI_ISL_481683, EPI_ISL_481684, EPI_ISL_481685, EPI_ISL_481686, EPI_ISL_481687, EPI_ISL_481688, EPI_ISL_481689, EPI_ISL_481690, EPI_ISL_481691, EPI_ISL_481692, EPI_ISL_481693, EPI_ISL_481694, EPI_ISL_481695, EPI_ISL_481696, EPI_ISL_481697, EPI_ISL_481698, EPI_ISL_481699, EPI_ISL_481700, EPI_ISL_481701, EPI_ISL_481702, EPI_ISL_481703, EPI_ISL_481704, EPI_ISL_481705, EPI_ISL_481706, EPI_ISL_481707, EPI_ISL_481708, EPI_ISL_481709, EPI_ISL_481710, EPI_ISL_481711, EPI_ISL_481712, EPI_ISL_481713, EPI_ISL_481714, EPI_ISL_481715 | | | |

| | | | |
|---|---|---|---|
| see above | Department of Virology and Immunology, University of Helsinki and Helsinki University Hospital, Huslab Finland | Department of Virology, Faculty of Medicine, University of Helsinki, Helsinki, Finland | Teemu Smura, Hannimari Kallio-Kokko, Jenni Virtanen, Maija Suvanto, Sari Hannula, Harri Kangas, Pekka Ellonen, Olli Vapalahti |
| EPI_ISL_481716 | Prof. Massimo Zollo CEINGE TASK-FORCE COVID19 - Regione Campania | Prof. Massimo Zollo CEINGE TASK-FORCE COVID19 - Regione Campania | Veronica Ferrucci1,2, Dae young Kong8, Fatemeh asadzadeh1,2, Laura Marrone1,2, Roberto Siciliano1,2, Rino Cerino3, Giovanna Fusco3, Marika Comegna1,2, Angelo Boccia2, Maurizio Viscardi3, Giorgia Borriello3, Sergio Brandi3, Claudia Tiberio4, Luigi Atripaldi4, Giovanni Paolella1,2, Giuseppe Castaldo1,2, Stefano Pascarella4, Martina Bianchi4, Lorenzo Chiariotti1,2, Jae Myun Lee5, Jae Ho Jung6, Kyong Seop Yun7, Hong Yeoul Kim 7,8* and Massimo Zollo1,2* 1 CEINGE Biotecnologie Avanzate, Naples, Italia 2 Dipartimento di Medicina Molecolare e Biotecnologie Mediche DMMBM University of Naples Federico II, Italia 3 Istituto Zooprofilattico Sperimentale del Mezzogiorno, Naples, Italia 4 -U.O.C. di Patologia Clinica Ospedale D. Cotugno, Azienda Sanitaria Ospedali dei Colli, Naples, Italy. 5 Università La Sapienza di Roma, Italia 6 Department of Microbiology, Yonsei University College of Medicine, Seoul, Korea 7 Department of Surgery, Yonsei University College of Medicine, Seoul, Korea 8 Haim bio co., Ltd., , Indust |
| EPI_ISL_481717, EPI_ISL_481718, EPI_ISL_481719, EPI_ISL_481720, EPI_ISL_481721, EPI_ISL_481722, EPI_ISL_481723, EPI_ISL_481724, EPI_ISL_481725, EPI_ISL_481726, EPI_ISL_481727, EPI_ISL_481728, EPI_ISL_481729, EPI_ISL_481730, EPI_ISL_481731, EPI_ISL_481732, EPI_ISL_481733, EPI_ISL_481734, EPI_ISL_481735, EPI_ISL_481736, EPI_ISL_481737, EPI_ISL_481738, EPI_ISL_481739, EPI_ISL_481740 | | | |
| see above | Department of Virology and Immunology, University of Helsinki and Helsinki University Hospital, Huslab Finland | Department of Virology, Faculty of Medicine, University of Helsinki, Helsinki, Finland | Teemu Smura, Hannimari Kallio-Kokko, Jenni Virtanen, Maija Suvanto, Sari Hannula, Harri Kangas, Pekka Ellonen, Olli Vapalahti |
| EPI_ISL_481741 | Prof. Massimo Zollo CEINGE TASK-FORCE COVID19 - Regione Campania | Prof. Massimo Zollo CEINGE TASK-FORCE COVID19 - Regione Campania | Veronica Ferrucci1,2, Dae young Kong8, Fatemeh asadzadeh1,2, Laura Marrone1,2, Roberto Siciliano1,2, Rino Cerino3, Giovanna Fusco3, Marika Comegna1,2, Angelo Boccia2, Maurizio Viscardi3, Giorgia Borriello3, Sergio Brandi3, Claudia Tiberio4, Luigi Atripaldi4, Giovanni Paolella1,2, Giuseppe Castaldo1,2, Stefano Pascarella4, Martina Bianchi4, Lorenzo Chiariotti1,2, Jae Myun Lee5, Jae Ho Jung6, Kyong Seop Yun7, Hong Yeoul Kim 7,8* and Massimo Zollo1,2* 1 CEINGE Biotecnologie Avanzate, Naples, Italia 2 Dipartimento di Medicina Molecolare e Biotecnologie Mediche DMMBM University of Naples Federico II, Italia 3 Istituto Zooprofilattico Sperimentale del Mezzogiorno, Naples, Italia 4 -U.O.C. di Patologia Clinica Ospedale D. Cotugno, Azienda Sanitaria Ospedali dei Colli, Naples, Italy. 5 Università La Sapienza di Roma, Italia 6 Department of Microbiology, Yonsei University College of Medicine, Seoul, Korea 7 Department of Surgery, Yonsei University College of Medicine, Seoul, Korea 8 Haim bio co., Ltd., , Indust |
| EPI_ISL_481742, EPI_ISL_481743, EPI_ISL_481744, EPI_ISL_481745, EPI_ISL_481746, EPI_ISL_481747, EPI_ISL_481748, EPI_ISL_481749, EPI_ISL_481750, EPI_ISL_481751, EPI_ISL_481752, EPI_ISL_481753, EPI_ISL_481754, EPI_ISL_481755, EPI_ISL_481756, EPI_ISL_481757, EPI_ISL_481758 | | | |
| see above | Dr. Georges-L.-Dumont University Hospital Centre | National Microbiology Laboratory | Anna Majer, Shari Tyson, Grace Seo, Kristyn Burak, Philip Mabon, Elsie Grudeski, Rhiannon Huzarewich, Russell Mandes, Jennifer Tanner, Natalie Knox, Morag Graham, Gary Van Domselaar, Richard Garceau, Guillaume Desnoyers, Nathalie Bastien, Yan Li, Timothy Booth |
| EPI_ISL_481759 | Prof. Massimo Zollo CEINGE TASK-FORCE COVID19 - Regione Campania | Prof. Massimo Zollo CEINGE TASK-FORCE COVID19 - Regione Campania | Veronica Ferrucci1,2, Dae young Kong8, Fatemeh asadzadeh1,2, Laura Marrone1,2, Roberto Siciliano1,2, Rino Cerino3, Giovanna Fusco3, Marika Comegna1,2, Angelo Boccia2, Maurizio Viscardi3, Giorgia Borriello3, Sergio Brandi3, Claudia Tiberio4, Luigi Atripaldi4, Giovanni Paolella1,2, Giuseppe Castaldo1,2, Stefano Pascarella4, Martina Bianchi4, Lorenzo Chiariotti1,2, Jae Myun Lee5, Jae Ho Jung6, Kyong Seop Yun7, Hong Yeoul Kim 7,8* and Massimo Zollo1,2* 1 CEINGE Biotecnologie Avanzate, Naples, Italia 2 Dipartimento di Medicina Molecolare e Biotecnologie Mediche DMMBM University of Naples Federico II, Italia 3 Istituto Zooprofilattico Sperimentale del Mezzogiorno, Naples, Italia 4 -U.O.C. di Patologia Clinica Ospedale D. Cotugno, Azienda Sanitaria Ospedali dei Colli, Naples, Italy. 5 Università La Sapienza di Roma, Italia 6 Department of Microbiology, Yonsei University College of Medicine, Seoul, Korea 7 Department of Surgery, Yonsei University College of Medicine, Seoul, Korea 8 Haim bio co., Ltd., , Indust |
| EPI_ISL_481760, EPI_ISL_481761, EPI_ISL_481762 | Prof. Massimo Zollo CEINGE TASK-FORCE COVID19 - Regione Campania | Prof. Massimo Zollo CEINGE TASK-FORCE COVID19 - Regione Campania | Veronica Ferrucci1,2, Dae young Kong8, Fatemeh asadzadeh1,2, Laura Marrone1,2, Roberto Siciliano1,2, Rino Cerino3, Giovanna Fusco3, Marika Comegna1,2, Angelo Boccia2, Maurizio Viscardi3, Giorgia Borriello3, Sergio Brandi3, Claudia Tiberio4, Luigi Atripaldi4, Giovanni Paolella1,2, Giuseppe Castaldo1,2, Stefano Pascarella4, Martina Bianchi4, Lorenzo Chiariotti1,2, Jae Myun Lee5, Jae Ho Jung6, Kyong Seop Yun7, Hong Yeoul Kim 7,8* and Massimo Zollo1,2* 1 CEINGE Biotecnologie Avanzate, Naples, Italia 2 Dipartimento di Medicina Molecolare e Biotecnologie Mediche DMMBM University of Naples Federico II, Italia 3 Istituto Zooprofilattico Sperimentale del Mezzogiorno, Naples, Italia 4 -U.O.C. di Patologia Clinica Ospedale D. Cotugno, Azienda Sanitaria Ospedali dei Colli, Naples, Italy. 5 Università La Sapienza di Roma, Italia 6 Department of Microbiology, Yonsei University College of Medicine, Seoul, Korea 7 Department of Surgery, Yonsei University College of Medicine, Seoul, Korea 8 Haim bio co., Ltd., , Indust |
| EPI_ISL_481763, EPI_ISL_481764, EPI_ISL_481765, EPI_ISL_481766, EPI_ISL_481767, EPI_ISL_481768, EPI_ISL_481769, EPI_ISL_481770, EPI_ISL_481771, EPI_ISL_481772, EPI_ISL_481773, EPI_ISL_481774, EPI_ISL_481775, EPI_ISL_481776, EPI_ISL_481777, EPI_ISL_481778, EPI_ISL_481779, EPI_ISL_481780, EPI_ISL_481781, EPI_ISL_481782, EPI_ISL_481783, EPI_ISL_481784, EPI_ISL_481785, EPI_ISL_481786, EPI_ISL_481787, EPI_ISL_481788, EPI_ISL_481789, EPI_ISL_481790, EPI_ISL_481791, EPI_ISL_481792, EPI_ISL_481793, EPI_ISL_481794, EPI_ISL_481795, EPI_ISL_481796, EPI_ISL_481797, EPI_ISL_481798, EPI_ISL_481799, EPI_ISL_481800, EPI_ISL_481801, EPI_ISL_481802, EPI_ISL_481803, EPI_ISL_481804, EPI_ISL_481805, EPI_ISL_481806, EPI_ISL_481807, EPI_ISL_481808, EPI_ISL_481809, EPI_ISL_481810, EPI_ISL_481811, EPI_ISL_481812, EPI_ISL_481813, EPI_ISL_481814, EPI_ISL_481815, EPI_ISL_481816, EPI_ISL_481817, EPI_ISL_481818, EPI_ISL_481819, EPI_ISL_481820, EPI_ISL_481821, EPI_ISL_481822, EPI_ISL_481823, EPI_ISL_481824, EPI_ISL_481825, EPI_ISL_481826, EPI_ISL_481827, EPI_ISL_481828, EPI_ISL_481829, EPI_ISL_481830, EPI_ISL_481831, EPI_ISL_481832, EPI_ISL_481833, EPI_ISL_481834, EPI_ISL_481835, EPI_ISL_481836, EPI_ISL_481837, EPI_ISL_481838, EPI_ISL_481839, EPI_ISL_481840, EPI_ISL_481841, EPI_ISL_481842, EPI_ISL_481843, EPI_ISL_481844, EPI_ISL_481845, EPI_ISL_481846, EPI_ISL_481847, EPI_ISL_481848, EPI_ISL_481849, EPI_ISL_481850, EPI_ISL_481851, EPI_ISL_481852, EPI_ISL_481853, EPI_ISL_481854, EPI_ISL_481855, EPI_ISL_481856, EPI_ISL_481857, EPI_ISL_481858, EPI_ISL_481859, EPI_ISL_481860, EPI_ISL_481861, EPI_ISL_481862, EPI_ISL_481863, EPI_ISL_481864, EPI_ISL_481865, EPI_ISL_481866, EPI_ISL_481867, EPI_ISL_481868, EPI_ISL_481869, EPI_ISL_481870, EPI_ISL_481871, EPI_ISL_481872, EPI_ISL_481873, EPI_ISL_481874, EPI_ISL_481875, EPI_ISL_481876, EPI_ISL_481877, EPI_ISL_481878, EPI_ISL_481879, EPI_ISL_481880, EPI_ISL_481881, EPI_ISL_481882, EPI_ISL_481883, EPI_ISL_481884, EPI_ISL_481885, EPI_ISL_481886, EPI_ISL_481887, EPI_ISL_481888, EPI_ISL_481889, EPI_ISL_481890, EPI_ISL_481891, EPI_ISL_481892, EPI_ISL_481893, EPI_ISL_481894, EPI_ISL_481895, EPI_ISL_481896, EPI_ISL_481897, EPI_ISL_481898, EPI_ISL_481899, EPI_ISL_481900, EPI_ISL_481901, EPI_ISL_481902, EPI_ISL_481903, EPI_ISL_481904, EPI_ISL_481905, EPI_ISL_481906, EPI_ISL_481907, EPI_ISL_481908, EPI_ISL_481909, EPI_ISL_481910, EPI_ISL_481911, EPI_ISL_481912, EPI_ISL_481913, EPI_ISL_481914, EPI_ISL_481915, EPI_ISL_481916, EPI_ISL_481917, EPI_ISL_481918, EPI_ISL_481919, EPI_ISL_481920, EPI_ISL_481921, EPI_ISL_481922, EPI_ISL_481923, EPI_ISL_481924, EPI_ISL_481925, EPI_ISL_481926, EPI_ISL_481927, EPI_ISL_481928, EPI_ISL_481929, EPI_ISL_481930, EPI_ISL_481931, EPI_ISL_481932, EPI_ISL_481933, EPI_ISL_481934, EPI_ISL_481935, EPI_ISL_481936, EPI_ISL_481937, EPI_ISL_481938, EPI_ISL_481939, EPI_ISL_481940, EPI_ISL_481941, EPI_ISL_481942, EPI_ISL_481943, EPI_ISL_481944, EPI_ISL_481945, EPI_ISL_481946, EPI_ISL_481947, EPI_ISL_481948, EPI_ISL_481949, EPI_ISL_481950, EPI_ISL_481951, EPI_ISL_481952, EPI_ISL_481953, EPI_ISL_481954, EPI_ISL_481955, EPI_ISL_481956, EPI_ISL_481957, EPI_ISL_481958, EPI_ISL_481959, EPI_ISL_481960, EPI_ISL_481961, EPI_ISL_481962, EPI_ISL_481963, EPI_ISL_481964, EPI_ISL_481965, EPI_ISL_481966, EPI_ISL_481967, EPI_ISL_481968, EPI_ISL_481969, EPI_ISL_481970, EPI_ISL_481971, EPI_ISL_481972, EPI_ISL_481973, EPI_ISL_481974, EPI_ISL_481975, EPI_ISL_481976, EPI_ISL_481977, EPI_ISL_481978, EPI_ISL_481979, EPI_ISL_481980, EPI_ISL_481981, EPI_ISL_481982, EPI_ISL_481983, EPI_ISL_481984, EPI_ISL_481985, EPI_ISL_481986, EPI_ISL_481987, EPI_ISL_481988, EPI_ISL_481989, EPI_ISL_481990, EPI_ISL_481991, EPI_ISL_481992, EPI_ISL_481993, EPI_ISL_481994, EPI_ISL_481995, EPI_ISL_481996, EPI_ISL_481997, EPI_ISL_481998, EPI_ISL_481999, EPI_ISL_482000, EPI_ISL_482001, EPI_ISL_482002, EPI_ISL_482003, EPI_ISL_482004, EPI_ISL_482005, EPI_ISL_482006, EPI_ISL_482007, EPI_ISL_482008, EPI_ISL_482009, EPI_ISL_482010, EPI_ISL_482011, EPI_ISL_482012, EPI_ISL_482013, EPI_ISL_482014, EPI_ISL_482015, EPI_ISL_482016, EPI_ISL_482017, EPI_ISL_482018, EPI_ISL_482019, EPI_ISL_482020, EPI_ISL_482021, EPI_ISL_482022, EPI_ISL_482023, EPI_ISL_482024, EPI_ISL_482025, EPI_ISL_482026, EPI_ISL_482027, EPI_ISL_482029, EPI_ISL_482030, EPI_ISL_482031 | | | |
| see above | PHE South West Regional Laboratory, National Infection Service | Wellcome Sanger Institute for the COVID-19 Genomics UK Consortium | Stephanie Hutchings, Hannah Pymont, Dr Peter Muir, Barry Vipond, Rich Hopes; and Alex Alderton, Roberto Amato, Sonia Goncalves, Ewan Harrison, David K. Jackson, Ian Johnston, Dominic Kwiatkowski, Cordelia Langford, John Sillitoe on behalf of the Wellcome Sanger Institute COVID-19 Surveillance Team (http://www.sanger.ac.uk/covid-team) |
| EPI_ISL_482032, EPI_ISL_482033, EPI_ISL_482034, EPI_ISL_482035, EPI_ISL_482036, EPI_ISL_482037, EPI_ISL_482038, EPI_ISL_482039, EPI_ISL_482040, EPI_ISL_482041, EPI_ISL_482042, EPI_ISL_482043, EPI_ISL_482044, EPI_ISL_482045, EPI_ISL_482046, EPI_ISL_482047, EPI_ISL_482048, EPI_ISL_482049, EPI_ISL_482050, EPI_ISL_482051, EPI_ISL_482052, EPI_ISL_482053, EPI_ISL_482054, EPI_ISL_482055, EPI_ISL_482056 | | | |
| see above | Regional Virus Laboratory, Belfast Health and Social Care Trust | Wellcome Sanger Institute for the COVID-19 Genomics UK Consortium | Conall McCaughey, James McKenna, Tanya Curran, Susan Feeney, Alison Watt, Ciara Cox, Mairead Connor, Zoltan Molnar, David Simpson, Derek Fairley; and Alex Alderton, Roberto Amato, Sonia Goncalves, Ewan Harrison, David K. Jackson, Ian Johnston, Dominic Kwiatkowski, Cordelia Langford, John Sillitoe on behalf of the Wellcome Sanger Institute COVID-19 Surveillance Team (http://www.sanger.ac.uk/covid-team) |
| EPI_ISL_482057, EPI_ISL_482058, EPI_ISL_482059, EPI_ISL_482060, EPI_ISL_482061, EPI_ISL_482062, EPI_ISL_482063, EPI_ISL_482064, EPI_ISL_482065, EPI_ISL_482066, EPI_ISL_482067, EPI_ISL_482068 | | | |
| see above | The Department of Microbiology, Torbay and South Devon NHS Foundation Trust | Wellcome Sanger Institute for the COVID-19 Genomics UK Consortium | Amy Hurd, Sophie Lloyd, Anthony Mogridge, Jack Howe, Helen Brown, Gary Booth, Mel Brown, Cheryl Bailiss, Michelle Harrison and Alex Alderton, Roberto Amato, Sonia Goncalves, Ewan Harrison, David K. Jackson, Ian Johnston, Dominic Kwiatkowski, Cordelia Langford, John Sillitoe on behalf of the Wellcome Sanger Institute COVID-19 Surveillance Team (http://www.sanger.ac.uk/covid-team) |
| EPI_ISL_482069 | Microbiology Department, Hereford County Hospital | Wellcome Sanger Institute for the COVID-19 Genomics UK Consortium | Alison Johnson, Venkat Sivaprakasam, Fenella Halstead, Jane Thomas, Wendy Hogsden, Samantha Lamb and Alex Alderton, Roberto Amato, Sonia Goncalves, Ewan Harrison, David K. Jackson, Ian Johnston, Dominic Kwiatkowski, Cordelia Langford, John Sillitoe on behalf of the Wellcome Sanger Institute COVID-19 Surveillance Team (http://www.sanger.ac.uk/covid-team) |
| EPI_ISL_482070 | Regional Virus Laboratory, Belfast Health and Social Care Trust | Wellcome Sanger Institute for the COVID-19 Genomics UK Consortium | Conall McCaughey, James McKenna, Tanya Curran, Susan Feeney, Alison Watt, Ciara Cox, Mairead Connor, Zoltan Molnar, David Simpson, Derek Fairley; and Alex Alderton, Roberto Amato, Sonia Goncalves, Ewan Harrison, David K. Jackson, Ian Johnston, Dominic Kwiatkowski, Cordelia Langford, John Sillitoe on behalf of the Wellcome Sanger Institute COVID-19 Surveillance Team (http://www.sanger.ac.uk/covid-team) |
| EPI_ISL_482071, EPI_ISL_482072, EPI_ISL_482073, EPI_ISL_482074 | Microbiology Department, Hereford County Hospital | Wellcome Sanger Institute for the COVID-19 Genomics UK Consortium | Alison Johnson, Venkat Sivaprakasam, Fenella Halstead, Jane Thomas, Wendy Hogsden, Samantha Lamb and Alex Alderton, Roberto Amato, Sonia Goncalves, Ewan Harrison, David K. Jackson, Ian Johnston, Dominic Kwiatkowski, Cordelia Langford, John Sillitoe on behalf of the Wellcome Sanger Institute COVID-19 Surveillance Team (http://www.sanger.ac.uk/covid-team) |
| EPI_ISL_482075 | Regional Virus Laboratory, Belfast Health and Social Care Trust | Wellcome Sanger Institute for the COVID-19 Genomics UK Consortium | Conall McCaughey, James McKenna, Tanya Curran, Susan Feeney, Alison Watt, Ciara Cox, Mairead Connor, Zoltan Molnar, David Simpson, Derek Fairley; and Alex Alderton, Roberto Amato, Sonia Goncalves, Ewan Harrison, David K. Jackson, Ian Johnston, Dominic Kwiatkowski, Cordelia Langford, John Sillitoe on behalf of the Wellcome Sanger Institute COVID-19 Surveillance Team (http://www.sanger.ac.uk/covid-team) |
| EPI_ISL_482076 | Microbiology Department, Hereford County Hospital | Wellcome Sanger Institute for the COVID-19 Genomics UK Consortium | Alison Johnson, Venkat Sivaprakasam, Fenella Halstead, Jane Thomas, Wendy Hogsden, Samantha Lamb and Alex Alderton, Roberto Amato, Sonia Goncalves, Ewan Harrison, David K. Jackson, Ian Johnston, Dominic Kwiatkowski, Cordelia Langford, John Sillitoe on behalf of the Wellcome Sanger Institute COVID-19 Surveillance Team (http://www.sanger.ac.uk/covid-team) |
| EPI_ISL_482077 | Regional Virus Laboratory, Belfast Health and Social Care Trust | Wellcome Sanger Institute for the COVID-19 Genomics UK Consortium | Conall McCaughey, James McKenna, Tanya Curran, Susan Feeney, Alison Watt, Ciara Cox, Mairead Connor, Zoltan Molnar, David Simpson, Derek Fairley; and Alex Alderton, Roberto Amato, Sonia Goncalves, Ewan Harrison, David K. Jackson, Ian Johnston, Dominic Kwiatkowski, Cordelia Langford, John Sillitoe on behalf of the Wellcome Sanger Institute COVID-19 Surveillance Team (http://www.sanger.ac.uk/covid-team) |
| EPI_ISL_482078, EPI_ISL_482079, EPI_ISL_482080, EPI_ISL_482081 | Microbiology Department, Hereford County Hospital | Wellcome Sanger Institute for the COVID-19 Genomics UK Consortium | Alison Johnson, Venkat Sivaprakasam, Fenella Halstead, Jane Thomas, Wendy Hogsden, Samantha Lamb and Alex Alderton, Roberto Amato, Sonia Goncalves, Ewan Harrison, David K. Jackson, Ian Johnston, Dominic Kwiatkowski, Cordelia Langford, John Sillitoe on behalf of the Wellcome Sanger Institute COVID-19 Surveillance Team (http://www.sanger.ac.uk/covid-team) |
| EPI_ISL_482082 | Regional Virus Laboratory, Belfast Health and Social Care Trust | Wellcome Sanger Institute for the COVID-19 Genomics UK Consortium | Conall McCaughey, James McKenna, Tanya Curran, Susan Feeney, Alison Watt, Ciara Cox, Mairead Connor, Zoltan Molnar, David Simpson, Derek Fairley; and Alex Alderton, Roberto Amato, Sonia Goncalves, Ewan Harrison, David K. Jackson, Ian Johnston, Dominic Kwiatkowski, Cordelia Langford, John Sillitoe on behalf of the Wellcome Sanger Institute COVID-19 Surveillance Team (http://www.sanger.ac.uk/covid-team) |
| EPI_ISL_482083, EPI_ISL_482084, EPI_ISL_482085, EPI_ISL_482086, EPI_ISL_482087, EPI_ISL_482088, EPI_ISL_482089, EPI_ISL_482090 | Microbiology Department, Hereford County Hospital | Wellcome Sanger Institute for the COVID-19 Genomics UK Consortium | Alison Johnson, Venkat Sivaprakasam, Fenella Halstead, Jane Thomas, Wendy Hogsden, Samantha Lamb and Alex Alderton, Roberto Amato, Sonia Goncalves, Ewan Harrison, David K. Jackson, Ian Johnston, Dominic Kwiatkowski, Cordelia Langford, John Sillitoe on behalf of the Wellcome Sanger Institute COVID-19 Surveillance Team (http://www.sanger.ac.uk/covid-team) |
| EPI_ISL_482091 | Regional Virus Laboratory, Belfast Health and Social Care Trust | Wellcome Sanger Institute for the COVID-19 Genomics UK Consortium | Conall McCaughey, James McKenna, Tanya Curran, Susan Feeney, Alison Watt, Ciara Cox, Mairead Connor, Zoltan Molnar, David Simpson, Derek Fairley; and Alex Alderton, Roberto Amato, Sonia Goncalves, Ewan Harrison, David K. Jackson, Ian Johnston, Dominic Kwiatkowski, Cordelia Langford, John Sillitoe on behalf of the Wellcome Sanger Institute COVID-19 Surveillance Team (http://www.sanger.ac.uk/covid-team) |
| EPI_ISL_482092, EPI_ISL_482093, EPI_ISL_482094, EPI_ISL_482095 | Microbiology Department, Hereford County Hospital | Wellcome Sanger Institute for the COVID-19 Genomics UK Consortium | Alison Johnson, Venkat Sivaprakasam, Fenella Halstead, Jane Thomas, Wendy Hogsden, Samantha Lamb and Alex Alderton, Roberto Amato, Sonia Goncalves, Ewan Harrison, David K. Jackson, Ian Johnston, Dominic Kwiatkowski, Cordelia Langford, John Sillitoe on behalf of the Wellcome Sanger Institute COVID-19 Surveillance Team (http://www.sanger.ac.uk/covid-team) |
| EPI_ISL_482096, EPI_ISL_482097 | Regional Virus Laboratory, Belfast Health and Social Care Trust | Wellcome Sanger Institute for the COVID-19 Genomics UK Consortium | Conall McCaughey, James McKenna, Tanya Curran, Susan Feeney, Alison Watt, Ciara Cox, Mairead Connor, Zoltan Molnar, David Simpson, Derek Fairley; and Alex Alderton, Roberto Amato, Sonia Goncalves, Ewan Harrison, David K. Jackson, Ian Johnston, Dominic Kwiatkowski, Cordelia Langford, John Sillitoe on behalf of the Wellcome Sanger Institute COVID-19 Surveillance Team (http://www.sanger.ac.uk/covid-team) |
| EPI_ISL_482098, EPI_ISL_482099, EPI_ISL_482100 | Microbiology Department, Hereford County Hospital | Wellcome Sanger Institute for the COVID-19 Genomics UK Consortium | Alison Johnson, Venkat Sivaprakasam, Fenella Halstead, Jane Thomas, Wendy Hogsden, Samantha Lamb and Alex Alderton, Roberto Amato, Sonia Goncalves, Ewan Harrison, David K. Jackson, Ian Johnston, Dominic Kwiatkowski, Cordelia Langford, John Sillitoe on behalf of the Wellcome Sanger Institute COVID-19 Surveillance Team (http://www.sanger.ac.uk/covid-team) |
| EPI_ISL_482101, EPI_ISL_482102 | Regional Virus Laboratory, Belfast Health and Social Care Trust | Wellcome Sanger Institute for the COVID-19 Genomics UK Consortium | Conall McCaughey, James McKenna, Tanya Curran, Susan Feeney, Alison Watt, Ciara Cox, Mairead Connor, Zoltan Molnar, David Simpson, Derek Fairley; and Alex Alderton, Roberto Amato, Sonia Goncalves, Ewan Harrison, David K. Jackson, Ian Johnston, Dominic Kwiatkowski, Cordelia Langford, John Sillitoe on behalf of the Wellcome Sanger Institute COVID-19 Surveillance Team (http://www.sanger.ac.uk/covid-team) |
| EPI_ISL_482103, EPI_ISL_482104 | Microbiology Department, Hereford County Hospital | Wellcome Sanger Institute for the COVID-19 Genomics UK Consortium | Alison Johnson, Venkat Sivaprakasam, Fenella Halstead, Jane Thomas, Wendy Hogsden, Samantha Lamb and Alex Alderton, Roberto Amato, Sonia Goncalves, Ewan Harrison, David K. Jackson, Ian Johnston, Dominic Kwiatkowski, Cordelia Langford, John Sillitoe on behalf of the Wellcome Sanger Institute COVID-19 Surveillance Team (http://www.sanger.ac.uk/covid-team) |
| EPI_ISL_482105 | Regional Virus Laboratory, Belfast Health and Social Care Trust | Wellcome Sanger Institute for the COVID-19 Genomics UK Consortium | Conall McCaughey, James McKenna, Tanya Curran, Susan Feeney, Alison Watt, Ciara Cox, Mairead Connor, Zoltan Molnar, David Simpson, Derek Fairley; and Alex Alderton, Roberto Amato, Sonia Goncalves, Ewan Harrison, David K. Jackson, Ian Johnston, Dominic Kwiatkowski, Cordelia Langford, John Sillitoe on behalf of the Wellcome Sanger Institute COVID-19 Surveillance Team (http://www.sanger.ac.uk/covid-team) |
| EPI_ISL_482106, EPI_ISL_482107, EPI_ISL_482108, EPI_ISL_482109, EPI_ISL_482110, EPI_ISL_482111, EPI_ISL_482112, EPI_ISL_482113, EPI_ISL_482114 | Microbiology Department, Hereford County Hospital | Wellcome Sanger Institute for the COVID-19 Genomics UK Consortium | Alison Johnson, Venkat Sivaprakasam, Fenella Halstead, Jane Thomas, Wendy Hogsden, Samantha Lamb and Alex Alderton, Roberto Amato, Sonia Goncalves, Ewan Harrison, David K. Jackson, Ian Johnston, Dominic Kwiatkowski, Cordelia Langford, John Sillitoe on behalf of the Wellcome Sanger Institute COVID-19 Surveillance Team (http://www.sanger.ac.uk/covid-team) |
| EPI_ISL_482115 | Regional Virus Laboratory, Belfast Health and Social Care Trust | Wellcome Sanger Institute for the COVID-19 Genomics UK Consortium | Conall McCaughey, James McKenna, Tanya Curran, Susan Feeney, Alison Watt, Ciara Cox, Mairead Connor, Zoltan Molnar, David Simpson, Derek Fairley; and Alex Alderton, Roberto Amato, Sonia Goncalves, Ewan Harrison, David K. Jackson, Ian Johnston, Dominic Kwiatkowski, Cordelia Langford, John Sillitoe on behalf of the Wellcome Sanger Institute COVID-19 Surveillance Team (http://www.sanger.ac.uk/covid-team) |
| EPI_ISL_482116, EPI_ISL_482117, EPI_ISL_482118, EPI_ISL_482119 | Microbiology Department, Hereford County Hospital | Wellcome Sanger Institute for the COVID-19 Genomics UK Consortium | Alison Johnson, Venkat Sivaprakasam, Fenella Halstead, Jane Thomas, Wendy Hogsden, Samantha Lamb and Alex Alderton, Roberto Amato, Sonia Goncalves, Ewan Harrison, David K. Jackson, Ian Johnston, Dominic Kwiatkowski, Cordelia Langford, John Sillitoe on behalf of the Wellcome Sanger Institute COVID-19 Surveillance Team (http://www.sanger.ac.uk/covid-team) |
| EPI_ISL_482120, EPI_ISL_482121, | The Department of Microbiology, Torbay and South | Wellcome Sanger Institute for the COVID-19 | Amy Hurd, Sophie Lloyd, Anthony Mogridge, Jack Howe, Helen Brown, Gary Booth, Mel Brown, Cheryl Bailiss, Michelle Harrison and Alex Alderton, Roberto Amato, Sonia Goncalves, Ewan Harrison, David K. Jackson, Ian Johnston, Dominic Kwiatkowski, |

| | | | |
|---|---|---|---|
| EPI_ISL_482122, EPI_ISL_482123 | Devon NHS Foundation Trust | Genomics UK Consortium | Cordelia Langford, John Sillitoe on behalf of the Wellcome Sanger Institute COVID-19 Surveillance Team (http://www.sanger.ac.uk/covid-team) |
| EPI_ISL_482124 | North West London Pathology, Imperial College Healthcare NHS Trust | Wellcome Sanger Institute for the COVID-19 Genomics UK Consortium | Ling Li, Paul Randell, David Muir, Frankie Bolt, Alison Holmes, James Price, Aileen Rowan, Graham Taylor, Anjna Badhan, Carolina Herrera and Alex Alderton, Roberto Amato, Sonia Goncalves, Ewan Harrison, David K. Jackson, Ian Johnston, Dominic Kwiatkowski, Cordelia Langford, John Sillitoe on behalf of the Wellcome Sanger Institute COVID-19 Surveillance Team (http://www.sanger.ac.uk/covid-team) |
| EPI_ISL_482125, EPI_ISL_482126, EPI_ISL_482127, EPI_ISL_482128, EPI_ISL_482129, EPI_ISL_482130 | The Department of Microbiology, Torbay and South Devon NHS Foundation Trust | Wellcome Sanger Institute for the COVID-19 Genomics UK Consortium | Amy Hurd, Sophie Lloyd, Anthony Mogridge, Jack Howe, Helen Brown, Gary Booth, Mel Brown, Cheryl Bailiss, Michelle Harrison and Alex Alderton, Roberto Amato, Sonia Goncalves, Ewan Harrison, David K. Jackson, Ian Johnston, Dominic Kwiatkowski, Cordelia Langford, John Sillitoe on behalf of the Wellcome Sanger Institute COVID-19 Surveillance Team (http://www.sanger.ac.uk/covid-team) |
| EPI_ISL_482131 | North West London Pathology, Imperial College Healthcare NHS Trust | Wellcome Sanger Institute for the COVID-19 Genomics UK Consortium | Ling Li, Paul Randell, David Muir, Frankie Bolt, Alison Holmes, James Price, Aileen Rowan, Graham Taylor, Anjna Badhan, Carolina Herrera and Alex Alderton, Roberto Amato, Sonia Goncalves, Ewan Harrison, David K. Jackson, Ian Johnston, Dominic Kwiatkowski, Cordelia Langford, John Sillitoe on behalf of the Wellcome Sanger Institute COVID-19 Surveillance Team (http://www.sanger.ac.uk/covid-team) |
| EPI_ISL_482132, EPI_ISL_482133, EPI_ISL_482134, EPI_ISL_482135, EPI_ISL_482136, EPI_ISL_482137, EPI_ISL_482139, EPI_ISL_482140, EPI_ISL_482141, EPI_ISL_482142, EPI_ISL_482143, EPI_ISL_482144, EPI_ISL_482145, EPI_ISL_482146, EPI_ISL_482147, EPI_ISL_482148, EPI_ISL_482149, EPI_ISL_482150, EPI_ISL_482151, EPI_ISL_482152, EPI_ISL_482153, EPI_ISL_482154, EPI_ISL_482155, EPI_ISL_482156, EPI_ISL_482157, EPI_ISL_482158, EPI_ISL_482159 | | | |
| see above | Regional Virus Laboratory, Belfast Health and Social Care Trust | Wellcome Sanger Institute for the COVID-19 Genomics UK Consortium | Conall McCaughey, James McKenna, Tanya Curran, Susan Feeney, Alison Watt, Ciara Cox, Mairead Connor, Zoltan Molnar, David Simpson, Derek Fairley; and Alex Alderton, Roberto Amato, Sonia Goncalves, Ewan Harrison, David K. Jackson, Ian Johnston, Dominic Kwiatkowski, Cordelia Langford, John Sillitoe on behalf of the Wellcome Sanger Institute COVID-19 Surveillance Team (http://www.sanger.ac.uk/covid-team) |
| EPI_ISL_482160, EPI_ISL_482161, EPI_ISL_482162, EPI_ISL_482163, EPI_ISL_482164, EPI_ISL_482165, EPI_ISL_482166, EPI_ISL_482167, EPI_ISL_482168, EPI_ISL_482169, EPI_ISL_482170, EPI_ISL_482171, EPI_ISL_482172, EPI_ISL_482173, EPI_ISL_482174, EPI_ISL_482175, EPI_ISL_482176, EPI_ISL_482177, EPI_ISL_482178, EPI_ISL_482179, EPI_ISL_482180, EPI_ISL_482181, EPI_ISL_482182, EPI_ISL_482183, EPI_ISL_482184, EPI_ISL_482185, EPI_ISL_482186, EPI_ISL_482187, EPI_ISL_482188, EPI_ISL_482189, EPI_ISL_482190, EPI_ISL_482191, EPI_ISL_482192, EPI_ISL_482193, EPI_ISL_482194, EPI_ISL_482195, EPI_ISL_482196, EPI_ISL_482197, EPI_ISL_482198, EPI_ISL_482199, EPI_ISL_482200, EPI_ISL_482201, EPI_ISL_482202, EPI_ISL_482203, EPI_ISL_482204, EPI_ISL_482205, EPI_ISL_482206, EPI_ISL_482207, EPI_ISL_482208, EPI_ISL_482209, EPI_ISL_482210, EPI_ISL_482211, EPI_ISL_482212, EPI_ISL_482213, EPI_ISL_482214, EPI_ISL_482215, EPI_ISL_482216, EPI_ISL_482217, EPI_ISL_482218 | | | |
| see above | University College London, Great Ormond Street Hospital for Children NHS Foundation Trust, Imperial College Healthcare NHS Trust | Wellcome Sanger Institute for the COVID-19 Genomics UK Consortium | Sergi Castellano, Rachel Williams, Mark Kristiansen, Paola Resende Silva, Sunando Roy, Tony Brooks, Helena Tutill, Paola Niola, Patricia Dyal, Charlotte Williams, Leysa Forrest, Yasmin Panchbhaya, Jacqueline Findlay, Sam Weeks, Julianne Brown, Kathryn Harris, Paul Randell, James Price, Alison Holmes, Judith Breuer and Alex Alderton, Roberto Amato, Sonia Goncalves, Ewan Harrison, David K. Jackson, Ian Johnston, Dominic Kwiatkowski, Cordelia Langford, John Sillitoe on behalf of the Wellcome Sanger Institute COVID-19 Surveillance Team (http://www.sanger.ac.uk/covid-team) |
| EPI_ISL_482292, EPI_ISL_482293, EPI_ISL_482294, EPI_ISL_482295, EPI_ISL_482296, EPI_ISL_482297, EPI_ISL_482298, EPI_ISL_482299, EPI_ISL_482300, EPI_ISL_482301, EPI_ISL_482302, EPI_ISL_482303, EPI_ISL_482304, EPI_ISL_482305, EPI_ISL_482306, EPI_ISL_482307, EPI_ISL_482308, EPI_ISL_482309, EPI_ISL_482310, EPI_ISL_482311, EPI_ISL_482312, EPI_ISL_482313, EPI_ISL_482314, EPI_ISL_482315, EPI_ISL_482316, EPI_ISL_482317, EPI_ISL_482318, EPI_ISL_482319, EPI_ISL_482320, EPI_ISL_482321, EPI_ISL_482322, EPI_ISL_482323, EPI_ISL_482324, EPI_ISL_482325, EPI_ISL_482326, EPI_ISL_482327, EPI_ISL_482328, EPI_ISL_482329, EPI_ISL_482330, EPI_ISL_482331, EPI_ISL_482332, EPI_ISL_482333, EPI_ISL_482334, EPI_ISL_482335, EPI_ISL_482336, EPI_ISL_482337, EPI_ISL_482338, EPI_ISL_482339, EPI_ISL_482340, EPI_ISL_482341, EPI_ISL_482342, EPI_ISL_482343, EPI_ISL_482344, EPI_ISL_482345, EPI_ISL_482346, EPI_ISL_482347, EPI_ISL_482348, EPI_ISL_482349, EPI_ISL_482350, EPI_ISL_482351, EPI_ISL_482352, EPI_ISL_482353, EPI_ISL_482354, EPI_ISL_482355, EPI_ISL_482356, EPI_ISL_482357, EPI_ISL_482358, EPI_ISL_482359, EPI_ISL_482360, EPI_ISL_482361, EPI_ISL_482362, EPI_ISL_482363, EPI_ISL_482364, EPI_ISL_482365, EPI_ISL_482366, EPI_ISL_482367, EPI_ISL_482368, EPI_ISL_482369, EPI_ISL_482370, EPI_ISL_482371, EPI_ISL_482372, EPI_ISL_482373, EPI_ISL_482374, EPI_ISL_482375, EPI_ISL_482376, EPI_ISL_482377, EPI_ISL_482378, EPI_ISL_482379, EPI_ISL_482380, EPI_ISL_482381, EPI_ISL_482382, EPI_ISL_482383, EPI_ISL_482384, EPI_ISL_482385, EPI_ISL_482386, EPI_ISL_482387, EPI_ISL_482388, EPI_ISL_482389, EPI_ISL_482390, EPI_ISL_482391, EPI_ISL_482392, EPI_ISL_482393, EPI_ISL_482394, EPI_ISL_482395, EPI_ISL_482396, EPI_ISL_482397, EPI_ISL_482398, EPI_ISL_482399, EPI_ISL_482400, EPI_ISL_482401, EPI_ISL_482402, EPI_ISL_482403, EPI_ISL_482404, EPI_ISL_482405, EPI_ISL_482406, EPI_ISL_482407, EPI_ISL_482408, EPI_ISL_482409, EPI_ISL_482410, EPI_ISL_482411, EPI_ISL_482412, EPI_ISL_482413, EPI_ISL_482414, EPI_ISL_482415, EPI_ISL_482416, EPI_ISL_482417, EPI_ISL_482418, EPI_ISL_482419, EPI_ISL_482420, EPI_ISL_482421, EPI_ISL_482422, EPI_ISL_482423, EPI_ISL_482424, EPI_ISL_482425, EPI_ISL_482426, EPI_ISL_482427, EPI_ISL_482428, EPI_ISL_482429, EPI_ISL_482430, EPI_ISL_482431, EPI_ISL_482432, EPI_ISL_482433, EPI_ISL_482434, EPI_ISL_482435, EPI_ISL_482436, EPI_ISL_482437, EPI_ISL_482438, EPI_ISL_482439, EPI_ISL_482440, EPI_ISL_482441, EPI_ISL_482442, EPI_ISL_482443, EPI_ISL_482444, EPI_ISL_482445, EPI_ISL_482446, EPI_ISL_482447, EPI_ISL_482448, EPI_ISL_482449, EPI_ISL_482450, EPI_ISL_482451, EPI_ISL_482452, EPI_ISL_482453, EPI_ISL_482454, EPI_ISL_482455, EPI_ISL_482456, EPI_ISL_482457, EPI_ISL_482458, EPI_ISL_482459, EPI_ISL_482460, EPI_ISL_482461, EPI_ISL_482462, EPI_ISL_482463, EPI_ISL_482464, EPI_ISL_482465, EPI_ISL_482466, EPI_ISL_482467 | | | |
| see above | Providence St. Joseph Health Molecular Genomics Laboratory | Providence St. Joseph Health Molecular Genomics Laboratory | Alexa K Dowdell, Brian D Piening, Fred L Robinson, Carlo B Bifulco, Mary Campbell |
| EPI_ISL_482468 | Laboratorio de Referencia Nacional de Virus Respiratorios. Instituto Nacional de Salud Peru | Laboratorio de Referencia Nacional de Biotecnologia y Biologia Molecular. Instituto Nacional de Salud Peru | Carlos Padilla Rojas, Priscila Lope Pari, Karolyn Vega Chozo, Johanna Balbuena Torres, Omar Caceres Rey, Henri Bailon Calderon, Maribel Huaringa Nuñez, Nancy Rojas Serrano |
| EPI_ISL_482469, EPI_ISL_482470 | Queen Elizabeth II Health Science Centre | National Microbiology Laboratory | Anna Majer, Shari Tyson, Grace Seo, Kristyn Burak, Philip Mabon, Elsie Grudeski, Rhiannon Huzarewich, Russell Mandes, Jennifer Tanner, Natalie Knox, Morag Graham, Gary Van Domselaar, Todd Hatchette, Jason LeBlanc, Nathalie Bastien, Yan Li, Timothy Booth |
| EPI_ISL_482471, EPI_ISL_482472, EPI_ISL_482473 | Dr. Georges-L.-Dumont University Hospital Centre | National Microbiology Laboratory | Anna Majer, Shari Tyson, Grace Seo, Kristyn Burak, Philip Mabon, Elsie Grudeski, Rhiannon Huzarewich, Russell Mandes, Jennifer Tanner, Natalie Knox, Morag Graham, Gary Van Domselaar, Richard Garceau, Guillaume Desnoyers, Nathalie Bastien, Yan Li, Timothy Booth |
| EPI_ISL_482474, EPI_ISL_482475 | Cadham Provincial Laboratory | National Microbiology Laboratory | Anna Majer, Shari Tyson, Grace Seo, Kristyn Burak, Philip Mabon, Elsie Grudeski, Rhiannon Huzarewich, Russell Mandes, Jennifer Tanner, Natalie Knox, Morag Graham, Gary Van Domselaar, Paul Van Caeseele, Jared Bullard, David Alexander, Kerry Dust, Nathalie Bastien, Yan Li, Timothy Booth, |
| EPI_ISL_482476, EPI_ISL_482477 | Queen Elizabeth II Health Science Centre | National Microbiology Laboratory | Anna Majer, Shari Tyson, Grace Seo, Kristyn Burak, Philip Mabon, Elsie Grudeski, Rhiannon Huzarewich, Russell Mandes, Jennifer Tanner, Natalie Knox, Morag Graham, Gary Van Domselaar, Todd Hatchette, Jason LeBlanc, Nathalie Bastien, Yan Li, Timothy Booth |
| EPI_ISL_482478 | Cadham Provincial Laboratory | National Microbiology Laboratory | Anna Majer, Shari Tyson, Grace Seo, Kristyn Burak, Philip Mabon, Elsie Grudeski, Rhiannon Huzarewich, Russell Mandes, Jennifer Tanner, Natalie Knox, Morag Graham, Gary Van Domselaar, Paul Van Caeseele, Jared Bullard, David Alexander, Kerry Dust, Nathalie Bastien, Yan Li, Timothy Booth, |
| EPI_ISL_482479 | Public Health Laboratory | National Microbiology Laboratory | Anna Majer, Shari Tyson, Grace Seo, Kristyn Burak, Philip Mabon, Elsie Grudeski, Rhiannon Huzarewich, Russell Mandes, Jennifer Tanner, Natalie Knox, Morag Graham, Gary Van Domselaar, Robert Needle, Yang Yu, Adel Malek, Laura Gilbert, George Zahariadis, Nathalie Bastien, Yan Li, Timothy Booth |
| EPI_ISL_482480, EPI_ISL_482481, EPI_ISL_482482, EPI_ISL_482483, EPI_ISL_482484 | Cadham Provincial Laboratory | National Microbiology Laboratory | Anna Majer, Shari Tyson, Grace Seo, Kristyn Burak, Philip Mabon, Elsie Grudeski, Rhiannon Huzarewich, Russell Mandes, Jennifer Tanner, Natalie Knox, Morag Graham, Gary Van Domselaar, Paul Van Caeseele, Jared Bullard, David Alexander, Kerry Dust, Nathalie Bastien, Yan Li, Timothy Booth, |
| EPI_ISL_482485, EPI_ISL_482486, EPI_ISL_482487 | National Institute of Laboratory Medicine and Referral Center | Genomic Research Lab, BCSIR | Abu Sayeed Mohammad Mahmud, Mohammad Samir Uzzaman, Eshrar Osman, Md. Ahasan Habib, Shahina Akter, Tanjina Akhter Banu, Md. Murshed Hasan Sarkar, Barna Goswami, Iffat Jahan, Md. Saddam Hossain, Tasnim Nafisa, Md. Maruf Ahmed Molla, Mahmuda Yeasmin, Asish Kumar Ghosh, Shahjahan Siddike, A. K. M. Shamsuzzaman, Sheikh Md. Selim Al Din, Utpal Chandra Ray, Salek Ahmed Sajib, Md. Salim Khan |
| EPI_ISL_482488 | National Institute of Laboratory Medicine and Referral Center | Genomic Research Lab, BCSIR | Md. Murshed Hasan Sarkar, Abu Sayeed Mohammad Mahmud, Mohammad Samir Uzzaman, Eshrar Osman, Md. Ahasan Habib, Shahina Akter, Tanjina Akhter Banu, Barna Goswami, Iffat Jahan, Md. Saddam Hossain, Tasnim Nafisa, Md. Maruf Ahmed Molla, Mahmuda Yeasmin, Asish Kumar Ghosh, Shahjahan Siddike, A. K. M. Shamsuzzaman, Sheikh Md. Selim Al Din, Utpal Chandra Ray, Salek Ahmed Sajib, Md. Salim Khan |
| EPI_ISL_482489 | National Institute of Laboratory Medicine and Referral Center | Genomic Research Lab, BCSIR | Md. Ahasan Habib, Abu Sayeed Mohammad Mahmud, Mohammad Samir Uzzaman, Eshrar Osman, Shahina Akter, Tanjina Akhter Banu, Md. Murshed Hasan Sarkar, Barna Goswami, Iffat Jahan, Md. Saddam Hossain, Tasnim Nafisa, Md. Maruf Ahmed Molla, Mahmuda Yeasmin, Asish Kumar Ghosh, Shahjahan Siddike, A. K. M. Shamsuzzaman, Sheikh Md. Selim Al Din, Utpal Chandra Ray, Salek Ahmed Sajib, Md. Salim Khan |
| EPI_ISL_482491, EPI_ISL_482492, EPI_ISL_482493, EPI_ISL_482494, EPI_ISL_482495, EPI_ISL_482496, EPI_ISL_482497, EPI_ISL_482498, EPI_ISL_482499, EPI_ISL_482500, EPI_ISL_482501, EPI_ISL_482502, EPI_ISL_482503, EPI_ISL_482504, EPI_ISL_482505, EPI_ISL_482506, EPI_ISL_482507, EPI_ISL_482508, EPI_ISL_482509, EPI_ISL_482510, EPI_ISL_482511, EPI_ISL_482512, EPI_ISL_482513, EPI_ISL_482514, EPI_ISL_482515, EPI_ISL_482516, EPI_ISL_482517, EPI_ISL_482518, EPI_ISL_482519, EPI_ISL_482520, EPI_ISL_482521, EPI_ISL_482522, EPI_ISL_482523, EPI_ISL_482524, EPI_ISL_482525, EPI_ISL_482526, EPI_ISL_482527, EPI_ISL_482528, EPI_ISL_482529, EPI_ISL_482530, EPI_ISL_482531, EPI_ISL_482532, EPI_ISL_482533, EPI_ISL_482534, EPI_ISL_482535, EPI_ISL_482536, EPI_ISL_482537, EPI_ISL_482538, EPI_ISL_482539, EPI_ISL_482540, EPI_ISL_482541, EPI_ISL_482542, EPI_ISL_482543, EPI_ISL_482544, EPI_ISL_482545, EPI_ISL_482546, EPI_ISL_482547, EPI_ISL_482548, EPI_ISL_482549, EPI_ISL_482550, EPI_ISL_482551, EPI_ISL_482552, EPI_ISL_482553, EPI_ISL_482554, EPI_ISL_482555, EPI_ISL_482556, EPI_ISL_482557, EPI_ISL_482558, EPI_ISL_482559, EPI_ISL_482560, EPI_ISL_482561, EPI_ISL_482562, EPI_ISL_482563, EPI_ISL_482564, EPI_ISL_482565, EPI_ISL_482566, EPI_ISL_482567, EPI_ISL_482568, EPI_ISL_482569, EPI_ISL_482570, EPI_ISL_482571, EPI_ISL_482572, EPI_ISL_482573, EPI_ISL_482574 | | | |
| see above | National Centre for Disease control (NCDC) | NCDC/CSIR-IGIB | Pramod Kumar#, Rajesh Pandey#, Pooja Sharma, Mahesh S Dhar, Vivekanand A, Bharathram Uppili, Robin Marwal, Radhakrishanan VS, Saruchi Wadhwa, Nishu Tyagi, Uma Sharma, Priyanka Singh, Hemlata Lall, Meena Datta, Varun Jaiswal, Hema Gogia, Preeti Madan, Prateek Singh, Debasis Dash, Mitali Mukerji, Sandhya Kabra, Sujeet Singh, Mohammed Faruq, Anurag Agrawal*, Partha Rakshit* |
| EPI_ISL_482575, EPI_ISL_482576, EPI_ISL_482577, EPI_ISL_482578, EPI_ISL_482579, EPI_ISL_482580, EPI_ISL_482581, EPI_ISL_482582, EPI_ISL_482583, EPI_ISL_482584, EPI_ISL_482585, EPI_ISL_482586 | | | |
| see above | Hangzhou Center for Diseases Control and Prevention | Hangzhou Center for Diseases Control and Prevention | Jun Li, Haoqiu Wang, Lingfeng Mao, Hua Yu, Xinfen Yu, Zhou Sun, Xin Qian, Shuchang Chen, Junfang Chen, Xuchu Wang |
| EPI_ISL_482587, EPI_ISL_482588, EPI_ISL_482589, EPI_ISL_482590, EPI_ISL_482591, EPI_ISL_482592, EPI_ISL_482593, EPI_ISL_482594, EPI_ISL_482595, EPI_ISL_482596, EPI_ISL_482597, EPI_ISL_482598, EPI_ISL_482599, EPI_ISL_482600, EPI_ISL_482601, EPI_ISL_482602, EPI_ISL_482603, EPI_ISL_482604, EPI_ISL_482605, EPI_ISL_482606, EPI_ISL_482607, EPI_ISL_482608, EPI_ISL_482609, EPI_ISL_482610, EPI_ISL_482611, EPI_ISL_482612, EPI_ISL_482613, EPI_ISL_482614, EPI_ISL_482615, EPI_ISL_482616, EPI_ISL_482617, EPI_ISL_482618, EPI_ISL_482619, EPI_ISL_482620, EPI_ISL_482621, EPI_ISL_482622, EPI_ISL_482623, EPI_ISL_482624, EPI_ISL_482625, EPI_ISL_482626, EPI_ISL_482627, EPI_ISL_482628, EPI_ISL_482629, EPI_ISL_482630, EPI_ISL_482631, EPI_ISL_482632, EPI_ISL_482633, EPI_ISL_482634, EPI_ISL_482635, EPI_ISL_482636, EPI_ISL_482637, EPI_ISL_482638, EPI_ISL_482639, EPI_ISL_482640, EPI_ISL_482641, EPI_ISL_482642, EPI_ISL_482643, EPI_ISL_482644, EPI_ISL_482645, EPI_ISL_482646, EPI_ISL_482647, EPI_ISL_482648, EPI_ISL_482649, EPI_ISL_482650, EPI_ISL_482651, EPI_ISL_482652, EPI_ISL_482653, EPI_ISL_482654, EPI_ISL_482655, EPI_ISL_482656, EPI_ISL_482657, EPI_ISL_482658, EPI_ISL_482659, EPI_ISL_482660, EPI_ISL_482661, EPI_ISL_482662, EPI_ISL_482663, EPI_ISL_482664, EPI_ISL_482665, EPI_ISL_482666, EPI_ISL_482667, EPI_ISL_482668, EPI_ISL_482669, EPI_ISL_482670, EPI_ISL_482671 | | | |
| see above | National Centre for Disease control (NCDC) | NCDC/CSIR-IGIB | Pramod Kumar#, Rajesh Pandey#, Pooja Sharma, Mahesh S Dhar, Vivekanand A, Bharathram Uppili, Robin Marwal, Radhakrishanan VS, Saruchi Wadhwa, Nishu Tyagi, Uma Sharma, Priyanka Singh, Hemlata Lall, Meena Datta, Varun Jaiswal, Hema Gogia, Preeti Madan, Prateek Singh, Debasis Dash, Mitali Mukerji, Sandhya Kabra, Sujeet Singh, Mohammed Faruq, Anurag Agrawal*, Partha Rakshit* |
| EPI_ISL_482672, EPI_ISL_482673, EPI_ISL_482674, EPI_ISL_482675, EPI_ISL_482676, EPI_ISL_482677, EPI_ISL_482678, EPI_ISL_482679, EPI_ISL_482680, EPI_ISL_482681, EPI_ISL_482682, EPI_ISL_482683, EPI_ISL_482684, EPI_ISL_482685, EPI_ISL_482686, EPI_ISL_482687, EPI_ISL_482688, EPI_ISL_482689, EPI_ISL_482690, EPI_ISL_482691, EPI_ISL_482692, EPI_ISL_482693, EPI_ISL_482694, EPI_ISL_482695, EPI_ISL_482696, EPI_ISL_482697, EPI_ISL_482698, EPI_ISL_482699 | | | |
| see above | Singapore General Hospital | Department of Microbiology | Nurdyana Abdul Rahman, Kun Lee Lim, Chenhao Li, Kian Sing Chan, Lynette Oon, Kern Rei Chng, Niranjan Nagarajan, Karrie Ko |
| EPI_ISL_482700, EPI_ISL_482701 | National Institute of Laboratory Medicine and Referral Center | Genomic Research Lab, BCSIR | Abu Sayeed Mohammad Mahmud, Mohammad Samir Uzzaman, Eshrar Osman, Md. Ahasan Habib, Shahina Akter, Tanjina Akhter Banu, Md. Murshed Hasan Sarkar, Barna Goswami, Iffat Jahan, Md. Saddam Hossain, Tasnim Nafisa, Md. Maruf Ahmed Molla, Mahmuda Yeasmin, Asish Kumar Ghosh, Shahjahan Siddike, A. K. M. Shamsuzzaman, Sheikh Md. Selim Al Din, Utpal Chandra Ray, Salek Ahmed Sajib, Md. Salim Khan |
| EPI_ISL_482702, EPI_ISL_482703, EPI_ISL_482704, EPI_ISL_482705, EPI_ISL_482706, EPI_ISL_482707, EPI_ISL_482708, EPI_ISL_482709 | Molecular Diagnostics Services (MDS) | KRISP, KZN Research Innovation and Sequencing Platform | Giandhari J, Pillay S, Lessells R, Chimukangara B, Mdlalose K, York D, Khan S, Tegally H, Wilkinson E, de Oliveira T |
| EPI_ISL_482710, EPI_ISL_482711, EPI_ISL_482712, EPI_ISL_482713 | NHLS-IALCH | KRISP, KZN Research Innovation and Sequencing Platform | Giandhari J, Pillay S, Lessells R, Chimukangara B, Mdlalose K, York D, Khan S, Tegally H, Wilkinson E, de Oliveira T |
| EPI_ISL_482714, EPI_ISL_482715, EPI_ISL_482716, EPI_ISL_482717, EPI_ISL_482718, EPI_ISL_482719, EPI_ISL_482720, EPI_ISL_482721, EPI_ISL_482722, EPI_ISL_482723 | Molecular Diagnostics Services (MDS) | KRISP, KZN Research Innovation and Sequencing Platform | Giandhari J, Pillay S, Lessells R, Chimukangara B, Mdlalose K, York D, Khan S, Tegally H, Wilkinson E, de Oliveira T |
| EPI_ISL_482724, EPI_ISL_482725, EPI_ISL_482726, EPI_ISL_482727, EPI_ISL_482728, EPI_ISL_482729, EPI_ISL_482730, EPI_ISL_482731 | NHLS-IALCH | KRISP, KZN Research Innovation and Sequencing Platform | Giandhari J, Pillay S, Lessells R, Chimukangara B, Mdlalose K, York D, Khan S, Tegally H, Wilkinson E, de Oliveira T |
| EPI_ISL_482732, EPI_ISL_482733, EPI_ISL_482734, EPI_ISL_482735, EPI_ISL_482736, EPI_ISL_482737, EPI_ISL_482738, EPI_ISL_482739, EPI_ISL_482740 | LNR National Reference Laboratory, Mohammed VI University of Health Sciences | Medical Biotechnology Laboratory, Rabat Medical and Pharmacy School, Mohammed The Vth University in Rabat | Meriem LAAMARTI, Souad KARTTI, Rokia LAAMARTI , M.W. CHEMAO-ELFIHRI, Loubna ALLAM, Mouna OUADGHIRI, Imane SMYEJ, Jalila RAHOUI, Houda BENRAHMA, Jalil El ATAR, Idrissa DIAWARA, Rachid EL JAOUDI, Laila SBABOU, Chakib NEJJARI, Saaid AMZAZI, Rachid MENTAG, Lahcen BELYAMANI and Azeddine IBRAHIMI |
| EPI_ISL_482742 | unknown | Molecular and Cell Biology, Globe Biotech Limited | Baray,J.C., Mahmud,A., Khan,M.R., Chowdhury,M.M.H., Roy,R., Islam,F., Nag,K. and Sultana,N. |

| | | | |
|---|---|---|---|
| EPI_ISL_482744 | unknown | Laboratory Diagnostic | Vidanovic,D., Tesovic,B., Banovic Djeri,B., Knezevic,A., Vidanovic,D., Tesovic,B., Banovic Djeri,B., Knezevic,A., Afonso,C. |
| EPI_ISL_482745 | unknown | Medical Microbiology | Snijder,E.J., Ogando,N.S., Zevenhoven,J.C., Dalebout,T.J., de Vries,J.C. and Sidorov,I. |
| EPI_ISL_482746 | unknown | Medical Microbiology | Snijder,E.J., Ogando,N.S., Zevenhoven,J.C., Dalebout,T.J., de Vries,J.J. and Sidorov,I. |
| EPI_ISL_482759, EPI_ISL_482760, EPI_ISL_482761, EPI_ISL_482762, EPI_ISL_482763, EPI_ISL_482764, EPI_ISL_482765, EPI_ISL_482766, EPI_ISL_482767, EPI_ISL_482768, EPI_ISL_482769, EPI_ISL_482770, EPI_ISL_482771, EPI_ISL_482772, EPI_ISL_482773, EPI_ISL_482774, EPI_ISL_482775 | | | |
| see above | Medical Ain Shams Research Institute (MASRI), Ain Shams University | Medical Ain Shams Research Institute (MASRI), Ain Shams University | Hesham Elghazaly, Sara Hassan Agwa, Ahmad Moustafa, Hala Hafez, Sara Elnakeep, Shaimaa Moustafa, Aya Mohamed, Reham Mamdouh, Ghada Ismael, Ashraf Omar, Osama Mansour, Mahmoud Elmeitini |
| EPI_ISL_482777 | Queen Elizabeth Hospital | Hong Kong Department of Health | Mak Gannon C.K., Cheng Peter K.C., Lam Edman T.K., Chan Rickjason C.W., Tsang Dominic N.C. |
| EPI_ISL_482778 | Tuen Mun Hospital | Hong Kong Department of Health | Mak Gannon C.K., Cheng Peter K.C., Lam Edman T.K., Chan Rickjason C.W., Tsang Dominic N.C. |
| EPI_ISL_482779, EPI_ISL_482780 | Prince of Wales Hospital | Hong Kong Department of Health | Mak Gannon C.K., Cheng Peter K.C., Lam Edman T.K., Chan Rickjason C.W., Tsang Dominic N.C. |
| EPI_ISL_482781 | Centre for Health Protection | Hong Kong Department of Health | Mak Gannon C.K., Cheng Peter K.C., Lam Edman T.K., Chan Rickjason C.W., Tsang Dominic N.C. |
| EPI_ISL_482782, EPI_ISL_482783 | Tuen Mun Hospital | Hong Kong Department of Health | Mak Gannon C.K., Cheng Peter K.C., Lam Edman T.K., Chan Rickjason C.W., Tsang Dominic N.C. |
| EPI_ISL_482784 | Prince of Wales Hospital | Hong Kong Department of Health | Mak Gannon C.K., Cheng Peter K.C., Lam Edman T.K., Chan Rickjason C.W., Tsang Dominic N.C. |
| EPI_ISL_482820 | Centre de Recerca en Sanitat Animal (IRTA-CReSA) | IrsiCaixa AIDS Research Lab | J. Segalés, M. Puig, J. Rodon, C. Avila-Nieto, J. Carrillo, G. Cantero, M.T. Terrón, S. Cruz, M. Parera ,M. Noguera-Julián, N. Izquierdo-Useros, V. Guallar, E. Vidal, A. Valencia, I. Blanco, J. Blanco, B. Clotet, J. Vergara-Alert |
| EPI_ISL_482848, EPI_ISL_482849, EPI_ISL_482850 | NHLS-IALCH | KRISP, KZN Research Innovation and Sequencing Platform | Giandhari J, Pillay S, Lessells R, Chimukangara B, Mdlalose K, York D, Khan S, Tegally H, Wilkinson E, de Oliveira T |
| EPI_ISL_482851, EPI_ISL_482852, EPI_ISL_482853, EPI_ISL_482854, EPI_ISL_482855, EPI_ISL_482856, EPI_ISL_482857, EPI_ISL_482858, EPI_ISL_482859, EPI_ISL_482860, EPI_ISL_482861, EPI_ISL_482862, EPI_ISL_482863, EPI_ISL_482864, EPI_ISL_482865, EPI_ISL_482866, EPI_ISL_482867, EPI_ISL_482868, EPI_ISL_482869, EPI_ISL_482870, EPI_ISL_482871, EPI_ISL_482872 | | | |
| see above | Molecular Diagnostics Services (MDS) | KRISP, KZN Research Innovation and Sequencing Platform | Giandhari J, Pillay S, Lessells R, Chimukangara B, Mdlalose K, York D, Khan S, Tegally H, Wilkinson E, de Oliveira T |
| EPI_ISL_482874, EPI_ISL_482875, EPI_ISL_482876, EPI_ISL_482877, EPI_ISL_482878 | Institut Pasteur Dakar | Institut Pasteur de Dakar | Ndongo Dia, Moussa Moise Diagne, Mamadou Diop, Marie Henriette Dior Ndione, Mamadou malado Jallow, Safietou Sankhe, Ousmane Faye, Amadou Alpha Sall. |
| EPI_ISL_482879, EPI_ISL_482880, EPI_ISL_482881, EPI_ISL_482882, EPI_ISL_482883, EPI_ISL_482884, EPI_ISL_482885, EPI_ISL_482886, EPI_ISL_482887, EPI_ISL_482888, EPI_ISL_482889 | | | |
| see above | CHU Purpan - Laboratoire de Virologie - Institut Fédératif de Biologie | Laboratoire de virologie - École Nationale Vétérinaire de Toulouse | Guillaume Croville, Jean-Luc Guérin, Jacques Izopet |
| EPI_ISL_482946, EPI_ISL_482947, EPI_ISL_482948, EPI_ISL_482949, EPI_ISL_482950, EPI_ISL_482951, EPI_ISL_482952, EPI_ISL_482953, EPI_ISL_482954, EPI_ISL_482955, EPI_ISL_482956, EPI_ISL_482957, EPI_ISL_482958, EPI_ISL_482959, EPI_ISL_482960, EPI_ISL_482961, EPI_ISL_482962, EPI_ISL_482963, EPI_ISL_482964, EPI_ISL_482965, EPI_ISL_482966 | | | |
| see above | Minnesota Department of Health, Public Health Laboratory | Minnesota Department of Health, Public Health Laboratory | Matt Plumb, Jacob Garfin, and Xiong Wang |
| EPI_ISL_482967, EPI_ISL_482968, EPI_ISL_482969, EPI_ISL_482970, EPI_ISL_482971, EPI_ISL_482972, EPI_ISL_482973, EPI_ISL_482974, EPI_ISL_482975, EPI_ISL_482976, EPI_ISL_482977, EPI_ISL_482978, EPI_ISL_482979, EPI_ISL_482980, EPI_ISL_482981, EPI_ISL_482982, EPI_ISL_482983, EPI_ISL_482984, EPI_ISL_482985, EPI_ISL_482986, EPI_ISL_482987 | | | |
| see above | Mayo Clinic & Mayo Clinic Laboratories | Minnesota Department of Health, Public Health Laboratory | Matt Plumb, Jacob Garfin, and Xiong Wang |
| EPI_ISL_482988, EPI_ISL_482989, EPI_ISL_482990, EPI_ISL_482991, EPI_ISL_482992, EPI_ISL_482993, EPI_ISL_482994, EPI_ISL_482995, EPI_ISL_482996, EPI_ISL_482997, EPI_ISL_482998, EPI_ISL_482999, EPI_ISL_483000, EPI_ISL_483001, EPI_ISL_483002, EPI_ISL_483003, EPI_ISL_483004, EPI_ISL_483005, EPI_ISL_483006, EPI_ISL_483007, EPI_ISL_483008, EPI_ISL_483009, EPI_ISL_483010, EPI_ISL_483011, EPI_ISL_483012, EPI_ISL_483013, EPI_ISL_483014, EPI_ISL_483015, EPI_ISL_483016, EPI_ISL_483017 | | | |
| see above | Minnesota Department of Health, Public Health Laboratory | Minnesota Department of Health, Public Health Laboratory | Matt Plumb, Jacob Garfin, and Xiong Wang |
| EPI_ISL_483018, EPI_ISL_483019, EPI_ISL_483020, EPI_ISL_483021, EPI_ISL_483022, EPI_ISL_483023, EPI_ISL_483024, EPI_ISL_483025, EPI_ISL_483026, EPI_ISL_483027, EPI_ISL_483028, EPI_ISL_483029, EPI_ISL_483030, EPI_ISL_483031, EPI_ISL_483032, EPI_ISL_483033 | | | |
| see above | Utah Public Health Laboratory | Utah Public Health Laboratory | Heidi Butz, Erin Young, Kelly Oakeson |
| EPI_ISL_483035, EPI_ISL_483036, EPI_ISL_483037, EPI_ISL_483038 | Medical Ain Shams Research Institute (MASRI), Ain Shams University | Medical Ain Shams Research Institute (MASRI), Ain Shams University | Hesham Elghazaly, Sara Hassan Agwa, Ahmad Moustafa, Hala Hafez, Sara Elnakeep, Shaimaa Moustafa, Aya Mohamed, Reham Mamdouh, Ghada Ismael, Ashraf Omar, Osama Mansour, Mahmoud Elmeitini |
| EPI_ISL_483059 | Hospital Universitari Germans Trias i Pujol | IrsiCaixa AIDS Research Lab | J. Segalés, M. Puig, J. Rodon, C. Avila-Nieto, J. Carrillo, G. Cantero, M.T. Terrón, S. Cruz, M. Parera ,M. Noguera-Julián, N. Izquierdo-Useros, V. Guallar, E. Vidal, A. Valencia, I. Blanco, J. Blanco, B. Clotet, J. Vergara-Alert |
| EPI_ISL_483060 | unknown | Microbiology, Canterbury Health Laboratories | Dilcher,M., Anderson,T. |
| EPI_ISL_483061, EPI_ISL_483062 | unknown | National Reference Center for transfusion infectious risks, INTS | Cappy,P., Candotti,D., Sauvage,V., Lucas,Q., Boizeau,L., Gomez,J., Laperche,S. |
| EPI_ISL_483063, EPI_ISL_483064 | unknown | Virology, Ecole Nationale Veterinaire de Toulouse | Bessiere,P., Cadiergues,M.-C., Croville,G., Walch,M., Dubois,M., Izopet,J., Guerin,J.-L. |
| EPI_ISL_483065 | unknown | Centro de Desenvolvimento Tecnologico em Saude, Fundacao Oswaldo Cruz | Souza,T.M., Fintelman-Rodrigues,N., De Paula,A.D., Tschoeke,D., Barroso,S.P., Gregorio,M.L., Oliveira,J.S., Saraiva,F.B., Ferreira,M.A., Sacramento,C.Q. |
| EPI_ISL_483066, EPI_ISL_483067, EPI_ISL_483068, EPI_ISL_483069, EPI_ISL_483070, EPI_ISL_483071, EPI_ISL_483072, EPI_ISL_483073, EPI_ISL_483074, EPI_ISL_483075, EPI_ISL_483076, EPI_ISL_483077, EPI_ISL_483078, EPI_ISL_483079, EPI_ISL_483080, EPI_ISL_483081, EPI_ISL_483082, EPI_ISL_483083, EPI_ISL_483084, EPI_ISL_483085, EPI_ISL_483086, EPI_ISL_483087, EPI_ISL_483088, EPI_ISL_483089, EPI_ISL_483090, EPI_ISL_483091, EPI_ISL_483092, EPI_ISL_483093, EPI_ISL_483094, EPI_ISL_483095, EPI_ISL_483096, EPI_ISL_483097, EPI_ISL_483098, EPI_ISL_483099, EPI_ISL_483100, EPI_ISL_483101, EPI_ISL_483102, EPI_ISL_483103, EPI_ISL_483104, EPI_ISL_483105, EPI_ISL_483106, EPI_ISL_483107, EPI_ISL_483108, EPI_ISL_483109, EPI_ISL_483110, EPI_ISL_483111, EPI_ISL_483112, EPI_ISL_483113, EPI_ISL_483114, EPI_ISL_483115, EPI_ISL_483116, EPI_ISL_483117, EPI_ISL_483118, EPI_ISL_483119, EPI_ISL_483120, EPI_ISL_483121, EPI_ISL_483122, EPI_ISL_483123, EPI_ISL_483124, EPI_ISL_483125, EPI_ISL_483126, EPI_ISL_483127, EPI_ISL_483128, EPI_ISL_483129, EPI_ISL_483130, EPI_ISL_483131, EPI_ISL_483132, EPI_ISL_483133, EPI_ISL_483134, EPI_ISL_483135, EPI_ISL_483136, EPI_ISL_483137, EPI_ISL_483138 | | | |
| see above | SA Pathology | SA Pathology | Lex Leong, Chuan Kok Lim, Mark Turra, Ivan Bastian, Geoff Higgins |
| EPI_ISL_483139, EPI_ISL_483140, EPI_ISL_483141, EPI_ISL_483142, EPI_ISL_483143, EPI_ISL_483144, EPI_ISL_483145, EPI_ISL_483146, EPI_ISL_483147, EPI_ISL_483148, EPI_ISL_483149, EPI_ISL_483150, EPI_ISL_483151, EPI_ISL_483152, EPI_ISL_483153, EPI_ISL_483154, EPI_ISL_483155, EPI_ISL_483156, EPI_ISL_483157 | | | |
| see above | Robert Koch Institute, ZBS1 Highly Pathogenic Viruses, Berlin, Germany | Robert Koch Institute, Bioinformatics MF1, Berlin, Germany | Janine Michel, Andrea Thuermer, Oliver Drechsel, Rene Kmiecinski, Stephan Fuchs, Max v. Kleist, Andreas Nitsche |
| EPI_ISL_483158, EPI_ISL_483159, EPI_ISL_483160, EPI_ISL_483161, EPI_ISL_483162, EPI_ISL_483163, EPI_ISL_483164 | San Diego County Public Health Laboratory | Andersen lab at Scripps Research | SEARCH Alliance San Diego with Tracy Basler, Jovan Shephard, Brett Austin |
| EPI_ISL_483165, EPI_ISL_483166, EPI_ISL_483167, EPI_ISL_483168, EPI_ISL_483169, EPI_ISL_483170, EPI_ISL_483171, EPI_ISL_483172, EPI_ISL_483173, EPI_ISL_483174, EPI_ISL_483175, EPI_ISL_483176, EPI_ISL_483177, EPI_ISL_483178, EPI_ISL_483179, EPI_ISL_483180, EPI_ISL_483181, EPI_ISL_483182, EPI_ISL_483183, EPI_ISL_483184, EPI_ISL_483185, EPI_ISL_483186, EPI_ISL_483187, EPI_ISL_483188, EPI_ISL_483189, EPI_ISL_483190, EPI_ISL_483191, EPI_ISL_483192, EPI_ISL_483193, EPI_ISL_483194, EPI_ISL_483195, EPI_ISL_483196, EPI_ISL_483197, EPI_ISL_483198, EPI_ISL_483199, EPI_ISL_483200, EPI_ISL_483201, EPI_ISL_483202, EPI_ISL_483203, EPI_ISL_483204, EPI_ISL_483205, EPI_ISL_483206, EPI_ISL_483207, EPI_ISL_483208, EPI_ISL_483209, EPI_ISL_483210, EPI_ISL_483211, EPI_ISL_483212, EPI_ISL_483213, EPI_ISL_483214, EPI_ISL_483215, EPI_ISL_483216, EPI_ISL_483217, EPI_ISL_483218, EPI_ISL_483219, EPI_ISL_483220, EPI_ISL_483221, EPI_ISL_483222, EPI_ISL_483223, EPI_ISL_483224, EPI_ISL_483225, EPI_ISL_483226, EPI_ISL_483227, EPI_ISL_483228, EPI_ISL_483229, EPI_ISL_483230, EPI_ISL_483231, EPI_ISL_483232, EPI_ISL_483233, EPI_ISL_483234, EPI_ISL_483235, EPI_ISL_483236, EPI_ISL_483237, EPI_ISL_483238, EPI_ISL_483239, EPI_ISL_483240, EPI_ISL_483241, EPI_ISL_483242, EPI_ISL_483243, EPI_ISL_483244, EPI_ISL_483245, EPI_ISL_483246, EPI_ISL_483247, EPI_ISL_483248, EPI_ISL_483249, EPI_ISL_483250, EPI_ISL_483251, EPI_ISL_483252, EPI_ISL_483253, EPI_ISL_483254, EPI_ISL_483255, EPI_ISL_483256, EPI_ISL_483257, EPI_ISL_483258, EPI_ISL_483259, EPI_ISL_483260, EPI_ISL_483261, EPI_ISL_483262, EPI_ISL_483263, EPI_ISL_483264, EPI_ISL_483265, EPI_ISL_483266, EPI_ISL_483267, EPI_ISL_483268, EPI_ISL_483269, EPI_ISL_483270, EPI_ISL_483271, EPI_ISL_483272, EPI_ISL_483273, EPI_ISL_483274, EPI_ISL_483275, EPI_ISL_483276, EPI_ISL_483277, EPI_ISL_483278, EPI_ISL_483279, EPI_ISL_483280, EPI_ISL_483281, EPI_ISL_483282, EPI_ISL_483283, EPI_ISL_483284, EPI_ISL_483285, EPI_ISL_483286, EPI_ISL_483287, EPI_ISL_483288, EPI_ISL_483289, EPI_ISL_483290, EPI_ISL_483291, EPI_ISL_483292, EPI_ISL_483293, EPI_ISL_483294, EPI_ISL_483295, EPI_ISL_483296, EPI_ISL_483297, EPI_ISL_483298, EPI_ISL_483299, EPI_ISL_483300, EPI_ISL_483301, EPI_ISL_483302, EPI_ISL_483303, EPI_ISL_483304, EPI_ISL_483305, EPI_ISL_483306, EPI_ISL_483307, EPI_ISL_483308, EPI_ISL_483309, EPI_ISL_483310, EPI_ISL_483311, EPI_ISL_483312, EPI_ISL_483313, EPI_ISL_483314, EPI_ISL_483315, EPI_ISL_483316, EPI_ISL_483317, EPI_ISL_483318, EPI_ISL_483319, EPI_ISL_483320, EPI_ISL_483321, EPI_ISL_483322, EPI_ISL_483323, EPI_ISL_483324, EPI_ISL_483325, EPI_ISL_483326, EPI_ISL_483327, EPI_ISL_483328, EPI_ISL_483329, EPI_ISL_483330, EPI_ISL_483331, EPI_ISL_483332, EPI_ISL_483333, EPI_ISL_483334, EPI_ISL_483335, EPI_ISL_483336, EPI_ISL_483337, EPI_ISL_483338, EPI_ISL_483339, EPI_ISL_483340, EPI_ISL_483341, EPI_ISL_483342, EPI_ISL_483343, EPI_ISL_483344, EPI_ISL_483345, EPI_ISL_483346, EPI_ISL_483347, EPI_ISL_483348, EPI_ISL_483349, EPI_ISL_483350, EPI_ISL_483351, EPI_ISL_483352, EPI_ISL_483353, EPI_ISL_483354, EPI_ISL_483355, EPI_ISL_483356, EPI_ISL_483357, EPI_ISL_483358, EPI_ISL_483359, EPI_ISL_483360, EPI_ISL_483361, EPI_ISL_483362, EPI_ISL_483363, EPI_ISL_483364, EPI_ISL_483365, EPI_ISL_483366, EPI_ISL_483367, EPI_ISL_483368, EPI_ISL_483369, EPI_ISL_483370, EPI_ISL_483371, EPI_ISL_483372, EPI_ISL_483373, EPI_ISL_483374, EPI_ISL_483375, EPI_ISL_483376, EPI_ISL_483377, EPI_ISL_483378, EPI_ISL_483379, EPI_ISL_483380, EPI_ISL_483381, EPI_ISL_483382, EPI_ISL_483383, EPI_ISL_483384, EPI_ISL_483385, EPI_ISL_483386, EPI_ISL_483387, EPI_ISL_483388, EPI_ISL_483389, EPI_ISL_483390, EPI_ISL_483391, EPI_ISL_483392, EPI_ISL_483393, EPI_ISL_483394, EPI_ISL_483395, EPI_ISL_483396, EPI_ISL_483397 | | | |
| see above | UC San Diego Center for Advanced Laboratory Medicine | Andersen lab at Scripps Research | SEARCH Alliance San Diego with David Pride, Ji H Shin |
| EPI_ISL_483398, EPI_ISL_483399, EPI_ISL_483400, EPI_ISL_483401, EPI_ISL_483402 | UC San Diego Center for Advanced Laboratory Medicine | Andersen lab at Scripps Research | Allison Smither, Gilberto Sabino-Santos, Patricia Snarski, Lilia Melnik, Antoinette Bell, Kaylynn Genemaras, Arnaud Drouin, Dahlene Fusco, Robert Garry with SEARCH Alliance San Diego |
| EPI_ISL_483403, EPI_ISL_483404, EPI_ISL_483405, EPI_ISL_483406, EPI_ISL_483407, EPI_ISL_483408, EPI_ISL_483409, EPI_ISL_483410, EPI_ISL_483411, EPI_ISL_483412, EPI_ISL_483413, EPI_ISL_483414, EPI_ISL_483415, EPI_ISL_483416, EPI_ISL_483417, EPI_ISL_483418, EPI_ISL_483419, EPI_ISL_483420, EPI_ISL_483421, EPI_ISL_483422, EPI_ISL_483423, EPI_ISL_483424, EPI_ISL_483425, EPI_ISL_483426, EPI_ISL_483427, EPI_ISL_483428, EPI_ISL_483429, EPI_ISL_483430, EPI_ISL_483431, EPI_ISL_483432, EPI_ISL_483433, EPI_ISL_483434, EPI_ISL_483435, EPI_ISL_483436, EPI_ISL_483437, EPI_ISL_483438, EPI_ISL_483439, EPI_ISL_483440, EPI_ISL_483441, EPI_ISL_483442, EPI_ISL_483443, EPI_ISL_483444, EPI_ISL_483445, EPI_ISL_483446, EPI_ISL_483447, EPI_ISL_483448, EPI_ISL_483449, EPI_ISL_483450, EPI_ISL_483451, EPI_ISL_483452, EPI_ISL_483453, EPI_ISL_483454, EPI_ISL_483455, EPI_ISL_483456, EPI_ISL_483457, EPI_ISL_483458, EPI_ISL_483459, EPI_ISL_483460, EPI_ISL_483461, EPI_ISL_483462, EPI_ISL_483463, EPI_ISL_483464, EPI_ISL_483465, EPI_ISL_483466, EPI_ISL_483467, EPI_ISL_483468, EPI_ISL_483469, EPI_ISL_483470, EPI_ISL_483471, EPI_ISL_483472, EPI_ISL_483473, EPI_ISL_483474, EPI_ISL_483475, EPI_ISL_483476, EPI_ISL_483477, EPI_ISL_483478, EPI_ISL_483479, EPI_ISL_483480, EPI_ISL_483481, EPI_ISL_483482, EPI_ISL_483483, EPI_ISL_483484, EPI_ISL_483485, EPI_ISL_483486, EPI_ISL_483487, EPI_ISL_483488, EPI_ISL_483489, EPI_ISL_483490, EPI_ISL_483491, EPI_ISL_483492, EPI_ISL_483493, EPI_ISL_483494, EPI_ISL_483495, EPI_ISL_483496, EPI_ISL_483497, EPI_ISL_483498, EPI_ISL_483499, EPI_ISL_483500, EPI_ISL_483501, EPI_ISL_483502, EPI_ISL_483503, EPI_ISL_483504, EPI_ISL_483505, EPI_ISL_483506, EPI_ISL_483507, EPI_ISL_483508, EPI_ISL_483509, EPI_ISL_483510, EPI_ISL_483511, EPI_ISL_483512, EPI_ISL_483513, EPI_ISL_483514, EPI_ISL_483515, EPI_ISL_483516, EPI_ISL_483517, EPI_ISL_483518, EPI_ISL_483519, EPI_ISL_483520, EPI_ISL_483521, EPI_ISL_483522, EPI_ISL_483523, EPI_ISL_483524, EPI_ISL_483525, EPI_ISL_483526, EPI_ISL_483527, EPI_ISL_483528, EPI_ISL_483529, EPI_ISL_483530 | | | |
| see above | UC San Diego Center for Advanced Laboratory Medicine | Andersen lab at Scripps Research | SEARCH Alliance San Diego with David Pride, Ji H Shin |
| EPI_ISL_483531, EPI_ISL_483532, EPI_ISL_483533, EPI_ISL_483534, EPI_ISL_483535, EPI_ISL_483536, EPI_ISL_483537, EPI_ISL_483538, EPI_ISL_483539, EPI_ISL_483540, EPI_ISL_483541 | | | |
| see above | San Diego County Public Health Laboratory | Andersen lab at Scripps Research | SEARCH Alliance San Diego with Tracy Basler, Jovan Shephard, Brett Austin |
| EPI_ISL_483542, EPI_ISL_483543, EPI_ISL_483544, EPI_ISL_483545, EPI_ISL_483546, EPI_ISL_483547, EPI_ISL_483548, EPI_ISL_483549, EPI_ISL_483550, EPI_ISL_483551, EPI_ISL_483552, EPI_ISL_483553, EPI_ISL_483554, EPI_ISL_483555, EPI_ISL_483556, EPI_ISL_483557, EPI_ISL_483558, EPI_ISL_483559, EPI_ISL_483560, EPI_ISL_483561, EPI_ISL_483562, EPI_ISL_483563, EPI_ISL_483564, EPI_ISL_483565 | | | |
| see above | Kingdom of Bahrein Ministry of Health | Erasmus Medical Center | Bas Oude Munnink, David Nieuwenhuijse, Reina Sikkema, Fatema, Ebrahim Shehad, Amjad Ghanem Mohamed, Hashmeya Al Wasti, Claudia Schapendonk, Irina Chestakova, Anne van der Linden, Theo Bestebroer, Stefan van Nieuwkoop, Mark Pronk, Pascal Lexmond, Richard Molenkamp, Marion Koopmans, on behalf of the Dutch national COVID-19 response team. |
| EPI_ISL_483566 | Clinical Microbiology Laboratory- Basurto University Hospital | Biocruces-Bizkaia | Mikel J. Urrutikoetxea-Gutierrez, Ana Belén Belén de la Hoz, Matxalen Vidal-García, Mº Carmen Nieto Toboso, Estibaliz Ugalde-Zarraga, José Luis Díaz de Tuesta del Arco |
| EPI_ISL_483570 | Clinical Microbiology Laboratory- Basurto University | Biocruces-Bizkaia | Mikel J. Urrutikoetxea-Gutierrez, Ana Belén Belén de la Hoz, Matxalen Vidal-García, Mº Carmen Nieto Toboso, Estibaliz Ugalde-Zarraga, José Luis Díaz de Tuesta del Arco |

| | | | |
|---|---|---|---|
| | Hospita | | |
| EPI_ISL_483571 | Clinical Microbiology Laboratory- Basurto University Hospital | Biocruces-Bizkaia | Mikel J. Urrutikoetxea-Gutierrez, Ana Belén Belén de la Hoz, Matxalen Vidal-García, Mº Carmen Nieto Toboso, Estibaliz Ugalde-Zarraga, José Luis Díaz de Tuesta del Arco |
| EPI_ISL_483572 | Clinical Microbiology Laboratory- Basurto University Hospital | Biocruces-Bizkaia | Mikel J. Urrutikoetxea-Gutierrez, Ana Belén Belén de la Hoz, Matxalen Vidal-García, Mº Carmen Nieto Toboso, Estibaliz Ugalde-Zarraga, José Luis Díaz de Tuesta del Arco |
| EPI_ISL_483573 | Clinical Microbiology Laboratory- Basurto University Hospital | Biocruces-Bizkaia | Mikel J. Urrutikoetxea-Gutierrez, Ana Belén Belén de la Hoz, Matxalen Vidal-García, Mº Carmen Nieto Toboso, Estibaliz Ugalde-Zarraga, José Luis Díaz de Tuesta del Arco |
| EPI_ISL_483576, EPI_ISL_483577, EPI_ISL_483578, EPI_ISL_483579, EPI_ISL_483580, EPI_ISL_483581, EPI_ISL_483582, EPI_ISL_483583, EPI_ISL_483584, EPI_ISL_483585, EPI_ISL_483586, EPI_ISL_483587, EPI_ISL_483588, EPI_ISL_483589, EPI_ISL_483590, EPI_ISL_483591, EPI_ISL_483592, EPI_ISL_483593, EPI_ISL_483594, EPI_ISL_483595, EPI_ISL_483596, EPI_ISL_483597, EPI_ISL_483598, EPI_ISL_483599, EPI_ISL_483600, EPI_ISL_483601, EPI_ISL_483602, EPI_ISL_483603, EPI_ISL_483604, EPI_ISL_483605, EPI_ISL_483606, EPI_ISL_483607, EPI_ISL_483608, EPI_ISL_483609, EPI_ISL_483610, EPI_ISL_483611, EPI_ISL_483612, EPI_ISL_483613, EPI_ISL_483614, EPI_ISL_483615, EPI_ISL_483616, EPI_ISL_483617, EPI_ISL_483618, EPI_ISL_483619, EPI_ISL_483620, EPI_ISL_483621 | | | |
| see above | National Public Health Laboratory, National Centre for Infectious Diseases | National Public Health Laboratory, National Centre for Infectious Diseases | Mak TM, Octavia S, Zhou Z, Chavatte JM, Cui L, Lin RTP |
| EPI_ISL_483622, EPI_ISL_483623 | National Institute of Laboratory Medicine and Referral Center | Genomic Research Lab, BCSIR | Tasnim Nafisa, Abu Sayeed Mohammad Mahmud, Mohammad Samir Uzzaman, Eshrar Osman, Md. Ahasan Habib, Shahina Akter, Tanjina Akhter Banu, Md. Murshed Hasan Sarkar, Barna Goswami, Iffat Jahan, Md. Saddam Hossain, Md. Maruf Ahmed Molla, Mahmuda Yeasmin, Asish Kumar Ghosh, A. K. M. Shamsuzzaman, Sheikh Md. Selim Al Din, Utpal Chandra Ray, Salek Ahmed Sajib, Md. Salim Khan |
| EPI_ISL_483624 | National Institute of Laboratory Medicine and Referral Center | Genomic Research Lab, BCSIR | Md. Maruf Ahmed Molla, Abu Sayeed Mohammad Mahmud, Mohammad Samir Uzzaman, Eshrar Osman, Md. Ahasan Habib, Shahina Akter, Tanjina Akhter Banu, Md. Murshed Hasan Sarkar, Barna Goswami, Iffat Jahan, Md. Saddam Hossain, Tasnim Nafisa, Mahmuda Yeasmin, Asish Kumar Ghosh, A. K. M. Shamsuzzaman, Sheikh Md. Selim Al Din, Utpal Chandra Ray, Salek Ahmed Sajib, Md. Salim Khan |
| EPI_ISL_483625 | unknown | Molecular Microbiology of Laboratory | Bui Thi,D.T., Nguyen,H.T., Tran,T.X., Nguyen,D.D., Pham,L.T., Vu,H.T., Le Thi,Q.M., Nguyen Le,H.K., Hoang Vu,P.M., Dang,A.D., Dong,Q.V. and Dinh,K.D. |
| EPI_ISL_483626 | National Institute of Laboratory Medicine and Referral Center | Genomic Research Lab, BCSIR | Md. Maruf Ahmed Molla, Abu Sayeed Mohammad Mahmud, Mohammad Samir Uzzaman, Eshrar Osman, Md. Ahasan Habib, Shahina Akter, Tanjina Akhter Banu, Md. Murshed Hasan Sarkar, Barna Goswami, Iffat Jahan, Md. Saddam Hossain, Tasnim Nafisa, Mahmuda Yeasmin, Asish Kumar Ghosh, A. K. M. Shamsuzzaman, Sheikh Md. Selim Al Din, Utpal Chandra Ray, Salek Ahmed Sajib, Md. Salim Khan |
| EPI_ISL_483627, EPI_ISL_483628 | National Institute of Laboratory Medicine and Referral Center | Genomic Research Lab, BCSIR | Mahmuda Yeasmin, Abu Sayeed Mohammad Mahmud, Mohammad Samir Uzzaman, Eshrar Osman, Md. Ahasan Habib, Shahina Akter, Tanjina Akhter Banu, Md. Murshed Hasan Sarkar, Barna Goswami, Iffat Jahan, Md. Saddam Hossain, Tasnim Nafisa, Md. Maruf Ahmed Molla, Asish Kumar Ghosh, A. K. M. Shamsuzzaman, Sheikh Md. Selim Al Din, Utpal Chandra Ray, Salek Ahmed Sajib, Md. Salim Khan |
| EPI_ISL_483629, EPI_ISL_483630 | National Institute of Laboratory Medicine and Referral Center | Genomic Research Lab, BCSIR | Asish Kumar Ghosh, Abu Sayeed Mohammad Mahmud, Mohammad Samir Uzzaman, Eshrar Osman, Md. Ahasan Habib, Shahina Akter, Tanjina Akhter Banu, Md. Murshed Hasan Sarkar, Barna Goswami, Iffat Jahan, Md. Saddam Hossain, Tasnim Nafisa, Md. Maruf Ahmed Molla, Mahmuda Yeasmin, A. K. M. Shamsuzzaman, Sheikh Md. Selim Al Din, Utpal Chandra Ray, Salek Ahmed Sajib, Md. Salim Khan |
| EPI_ISL_483631, EPI_ISL_483632 | National Institute of Laboratory Medicine and Referral Center | Genomic Research Lab, BCSIR | Md. Ahasan Habib, Abu Sayeed Mohammad Mahmud, Mohammad Samir Uzzaman, Eshrar Osman, Shahina Akter, Tanjina Akhter Banu, Md. Murshed Hasan Sarkar, Barna Goswami, Iffat Jahan, Md. Saddam Hossain, Tasnim Nafisa, Md. Maruf Ahmed Molla, Mahmuda Yeasmin, Asish Kumar Ghosh, A. K. M. Shamsuzzaman, Sheikh Md. Selim Al Din, Utpal Chandra Ray, Salek Ahmed Sajib, Md. Salim Khan |
| EPI_ISL_483633, EPI_ISL_483634 | National Institute of Laboratory Medicine and Referral Center | Genomic Research Lab, BCSIR | Shahina Akter, Abu Sayeed Mohammad Mahmud, Mohammad Samir Uzzaman, Eshrar Osman, Md. Ahasan Habib, Tanjina Akhter Banu, Md. Murshed Hasan Sarkar, Barna Goswami, Iffat Jahan, Md. Saddam Hossain, Tasnim Nafisa, Md. Maruf Ahmed Molla, Mahmuda Yeasmin, Asish Kumar Ghosh, A. K. M. Shamsuzzaman, Sheikh Md. Selim Al Din, Utpal Chandra Ray, Salek Ahmed Sajib, Md. Salim Khan |
| EPI_ISL_483635, EPI_ISL_483636 | National Institute of Laboratory Medicine and Referral Center | Genomic Research Lab, BCSIR | Tanjina Akhter Banu, Abu Sayeed Mohammad Mahmud, Mohammad Samir Uzzaman, Eshrar Osman, Md. Ahasan Habib, Shahina Akter, Md. Murshed Hasan Sarkar, Barna Goswami, Iffat Jahan, Md. Saddam Hossain, Tasnim Nafisa, Md. Maruf Ahmed Molla, Mahmuda Yeasmin, Asish Kumar Ghosh, A. K. M. Shamsuzzaman, Sheikh Md. Selim Al Din, Utpal Chandra Ray, Salek Ahmed Sajib, Md. Salim Khan |
| EPI_ISL_483637 | National Laboratory of Virology, Szentágothai Research Centre | National Laboratory of Virology, Szentágothai Research Centre | Endre Gábor Tóth, Balázs Somogyi, Ferenc Jakab, Gábor Kemenesi |
| EPI_ISL_483638, EPI_ISL_483639, EPI_ISL_483640 | Kingdom of Bahrein Ministry of Health | Erasmus Medical Center | Bas Oude Munnink, David Nieuwenhuijse, Reina Sikkema, Fatema, Ebrahim Shehad, Amjad Ghanem Mohamed, Hashmeya Al Wasti, Claudia Schapendonk, Irina Chestakova, Anne van der Linden, Theo Bestebroer, Stefan van Nieuwkoop, Mark Pronk, Pascal Lexmond, Richard Molenkamp, Marion Koopmans, on behalf of the Dutch national COVID-19 response team. |
| EPI_ISL_483641, EPI_ISL_483642 | National Institute of Laboratory Medicine and Referral Center | Genomic Research Lab, BCSIR | Barna Goswami, Abu Sayeed Mohammad Mahmud, Mohammad Samir Uzzaman, Eshrar Osman, Md. Ahasan Habib, Shahina Akter, Tanjina Akhter Banu, Md. Murshed Hasan Sarkar, Iffat Jahan, Md. Saddam Hossain, Tasnim Nafisa, Md. Maruf Ahmed Molla, Mahmuda Yeasmin, Asish Kumar Ghosh, A. K. M. Shamsuzzaman, Sheikh Md. Selim Al Din, Utpal Chandra Ray, Salek Ahmed Sajib, Md. Salim Khan |
| EPI_ISL_483643, EPI_ISL_483644 | National Institute of Laboratory Medicine and Referral Center | Genomic Research Lab, BCSIR | Iffat Jahan, Abu Sayeed Mohammad Mahmud, Mohammad Samir Uzzaman, Eshrar Osman, Md. Ahasan Habib, Shahina Akter, Tanjina Akhter Banu, Md. Murshed Hasan Sarkar, Barna Goswami, Md. Saddam Hossain, Tasnim Nafisa, Md. Maruf Ahmed Molla, Mahmuda Yeasmin, Asish Kumar Ghosh, A. K. M. Shamsuzzaman, Sheikh Md. Selim Al Din, Utpal Chandra Ray, Salek Ahmed Sajib, Md. Salim Khan |
| EPI_ISL_483645, EPI_ISL_483646, EPI_ISL_483647 | National Institute of Laboratory Medicine and Referral Center | Genomic Research Lab, BCSIR | Md. Saddam Hossain, Abu Sayeed Mohammad Mahmud, Mohammad Samir Uzzaman, Eshrar Osman, Md. Ahasan Habib, Shahina Akter, Tanjina Akhter Banu, Md. Murshed Hasan Sarkar, Barna Goswami, Iffat Jahan, Tasnim Nafisa, Md. Maruf Ahmed Molla, Mahmuda Yeasmin, Asish Kumar Ghosh, A. K. M. Shamsuzzaman, Sheikh Md. Selim Al Din, Utpal Chandra Ray, Salek Ahmed Sajib, Md. Salim Khan |
| EPI_ISL_483648, EPI_ISL_483649, EPI_ISL_483650, EPI_ISL_483651, EPI_ISL_483652, EPI_ISL_483653, EPI_ISL_483654, EPI_ISL_483655, EPI_ISL_483656, EPI_ISL_483657, EPI_ISL_483658, EPI_ISL_483659, EPI_ISL_483660, EPI_ISL_483661, EPI_ISL_483662, EPI_ISL_483663, EPI_ISL_483664, EPI_ISL_483665, EPI_ISL_483666, EPI_ISL_483667 | | | |
| see above | Viollier AG | Department of Biosystems Science and Engineering, ETH Zürich | Christian Beisel, Sarah Nadeau, Ivan Topolsky, Pedro Ferreira, Philipp Jablonski, Susana Posada-Céspedes, Tobias Schär, Ina Nissen, Natascha Santacroce, Elodie Burcklen, Christiane Beckmann, Maurice Redondo, Olivier Kobel, Christoph Noppen, Sophie Seidel, Noemie Santamaria de Souza, Niko Beerenwinkel, Tanja Stadler |
| EPI_ISL_483668, EPI_ISL_483669, EPI_ISL_483670, EPI_ISL_483671, EPI_ISL_483672, EPI_ISL_483673, EPI_ISL_483674, EPI_ISL_483675, EPI_ISL_483676, EPI_ISL_483677, EPI_ISL_483678, EPI_ISL_483679, EPI_ISL_483680, EPI_ISL_483681, EPI_ISL_483682, EPI_ISL_483683, EPI_ISL_483684, EPI_ISL_483685 | | | |
| see above | University Hospital Zurich | Department of Biosystems Science and Engineering, ETH Zürich | Christian Beisel, Sarah Nadeau, Ivan Topolsky, Pedro Ferreira, Philipp Jablonski, Susana Posada-Céspedes, Tobias Schär, Ina Nissen, Natascha Santacroce, Elodie Burcklen, Julia Martinez-Gomez, Phil Cheng, Mitch Levesque, Philipp Bosshard, Niko Beerenwinkel, Tanja Stadler |
| EPI_ISL_483686, EPI_ISL_483687 | National Institute of Laboratory Medicine and Referral Center | Genomic Research Lab, BCSIR | Md. Murshed Hasan Sarkar, Abu Sayeed Mohammad Mahmud, Mohammad Samir Uzzaman, Eshrar Osman, Md. Ahasan Habib, Shahina Akter, Tanjina Akhter Banu, Barna Goswami, Iffat Jahan, Md. Saddam Hossain, Tasnim Nafisa, Md. Maruf Ahmed Molla, Mahmuda Yeasmin, Asish Kumar Ghosh, A. K. M. Shamsuzzaman, Sheikh Md. Selim Al Din, Utpal Chandra Ray, Salek Ahmed Sajib, Md. Salim Khan |
| EPI_ISL_483688 | Genomic Research Lab, BCSIR | Genomic Research Lab, BCSIR | Md. Murshed Hasan Sarkar, Abu Sayeed Mohammad Mahmud, Mohammad Samir Uzzaman, Eshrar Osman, Md. Ahasan Habib, Shahina Akter, Tanjina Akhter Banu, Barna Goswami, Iffat Jahan, Md. Saddam Hossain, Tasnim Nafisa, Md. Maruf Ahmed Molla, Mahmuda Yeasmin, Asish Kumar Ghosh, A. K. M. Shamsuzzaman, Sheikh Md. Selim Al Din, Utpal Chandra Ray, Salek Ahmed Sajib, Md. Salim Khan |
| EPI_ISL_483689, EPI_ISL_483690, EPI_ISL_483691, EPI_ISL_483692 | National Institute of Laboratory Medicine and Referral Center | Genomic Research Lab, BCSIR | Md. Murshed Hasan Sarkar, Abu Sayeed Mohammad Mahmud, Mohammad Samir Uzzaman, Eshrar Osman, Md. Ahasan Habib, Shahina Akter, Tanjina Akhter Banu, Barna Goswami, Iffat Jahan, Md. Saddam Hossain, Tasnim Nafisa, Md. Maruf Ahmed Molla, Mahmuda Yeasmin, Asish Kumar Ghosh, A. K. M. Shamsuzzaman, Sheikh Md. Selim Al Din, Utpal Chandra Ray, Salek Ahmed Sajib, Md. Salim Khan |
| EPI_ISL_483693, EPI_ISL_483694, EPI_ISL_483695, EPI_ISL_483699, EPI_ISL_483700, EPI_ISL_483703 | National Institute of Laboratory Medicine and Referral Center | Genomic Research Lab, BCSIR | Abu Sayeed Mohammad Mahmud, Mohammad Samir Uzzaman, Eshrar Osman, Md. Ahasan Habib, Shahina Akter, Tanjina Akhter Banu, Md. Murshed Hasan Sarkar, Barna Goswami, Iffat Jahan, Md. Saddam Hossain, Tasnim Nafisa, Md. Maruf Ahmed Molla, Mahmuda Yeasmin, Asish Kumar Ghosh, A. K. M. Shamsuzzaman, Sheikh Md. Selim Al Din, Utpal Chandra Ray, Salek Ahmed Sajib, Md. Salim Khan |
| EPI_ISL_483704 | Israel Central Virology laboratory | Israel Central Virology laboratory | Neta Zuckerman, Efrat Dahan Bucris, Oran Erster, Ella Mendelson, Michal Mandelboim |
| EPI_ISL_483705, EPI_ISL_483707 | National Institute of Laboratory Medicine and Referral Center | Genomic Research Lab, BCSIR | Abu Sayeed Mohammad Mahmud, Mohammad Samir Uzzaman, Eshrar Osman, Md. Ahasan Habib, Shahina Akter, Tanjina Akhter Banu, Md. Murshed Hasan Sarkar, Barna Goswami, Iffat Jahan, Md. Saddam Hossain, Tasnim Nafisa, Md. Maruf Ahmed Molla, Mahmuda Yeasmin, Asish Kumar Ghosh, A. K. M. Shamsuzzaman, Sheikh Md. Selim Al Din, Utpal Chandra Ray, Salek Ahmed Sajib, Md. Salim Khan |
| EPI_ISL_483708, EPI_ISL_483709 | Israel Central Virology laboratory | Israel Central Virology laboratory | Neta Zuckerman, Efrat Dahan Bucris, Oran Erster, Ella Mendelson, Michal Mandelboim |
| EPI_ISL_483710 | National Institute of Laboratory Medicine and Referral Center | Genomic Research Lab, BCSIR | Abu Sayeed Mohammad Mahmud, Mohammad Samir Uzzaman, Eshrar Osman, Md. Ahasan Habib, Shahina Akter, Tanjina Akhter Banu, Md. Murshed Hasan Sarkar, Barna Goswami, Iffat Jahan, Md. Saddam Hossain, Tasnim Nafisa, Md. Maruf Ahmed Molla, Mahmuda Yeasmin, Asish Kumar Ghosh, A. K. M. Shamsuzzaman, Sheikh Md. Selim Al Din, Utpal Chandra Ray, Salek Ahmed Sajib, Md. Salim Khan |
| EPI_ISL_483711, EPI_ISL_483712, EPI_ISL_483713, EPI_ISL_483715, EPI_ISL_483717, EPI_ISL_483725 | Israel Central Virology laboratory | Israel Central Virology laboratory | Neta Zuckerman, Efrat Dahan Bucris, Oran Erster, Ella Mendelson, Michal Mandelboim |
| EPI_ISL_483820 | GMERS Medical College and Hospital, Gandhinagar | Gujarat Biotechnology Research Centre | Komal Patel, Labdhi Pandya, Afzal Ansari, Nikha Trivedi, Seema Bhatt, Gaurishankar Shrimali, Bhavesh Modi, Bharti Rajani, Apurvasinh Puvar, Janvi Raval, Zarna Patel, Monika Gandhi, Pinal Trivedi, Maharshi Pandya, Nidhi Patel, Nitin Savaliya, Raghawendra Kumar, Dinesh Kumar, Zuber Saiyed, R D Dixit, A M Kadri, Harsh Bakshi, Chaitanya Joshi, Madhvi Joshi |
| EPI_ISL_483821 | Government Medical College, Vadodara | Gujarat Biotechnology Research Centre | Labdhi Pandya, Afzal Ansari, Nikha Trivedi, Meenakshi Shah, Neena Doshi, Varsha Godbole, Apurvasinh Puvar, Janvi Raval, Zarna Patel, Monika Gandhi, Pinal Trivedi, Maharshi Pandya, Nidhi Patel, Nitin Savaliya, Raghawendra Kumar, Dinesh Kumar, Zuber Saiyed, Komal Patel, R D Dixit, A M Kadri, Harsh Bakshi, Chaitanya Joshi, Madhvi Joshi |
| EPI_ISL_483822 | Government Medical College, Vadodara | Gujarat Biotechnology Research Centre | Afzal Ansari, Nikha Trivedi, Meenakshi Shah, Neena Doshi, Varsha Godbole, Apurvasinh Puvar, Janvi Raval, Zarna Patel, Monika Gandhi, Pinal Trivedi, Maharshi Pandya, Nidhi Patel, Nitin Savaliya, Raghawendra Kumar, Dinesh Kumar, Zuber Saiyed, Komal Patel, Labdhi Pandya, R D Dixit, A M Kadri, Harsh Bakshi, Chaitanya Joshi, Madhvi Joshi |
| EPI_ISL_483823 | GMERS Medical College Himmatnagar | Gujarat Biotechnology Research Centre | Nikha Trivedi, Himanshu Khatri, Mayur Gandhi, Apurvasinh Puvar, Janvi Raval, Zarna Patel, Monika Gandhi, Pinal Trivedi, Maharshi Pandya, Nidhi Patel, Nitin Savaliya, Raghawendra Kumar, Dinesh Kumar, Zuber Saiyed, Komal Patel, Labdhi Pandya, Afzal Ansari, R D Dixit, A M Kadri, Harsh Bakshi, Chaitanya Joshi, Madhvi Joshi |
| EPI_ISL_483824 | GMERS Medical College Himmatnagar | Gujarat Biotechnology Research Centre | Himanshu Khatri, Mayur Gandhi, Apurvasinh Puvar, Janvi Raval, Zarna Patel, Monika Gandhi, Pinal Trivedi, Maharshi Pandya, Nidhi Patel, Nitin Savaliya, Raghawendra Kumar, Dinesh Kumar, Zuber Saiyed, Komal Patel, Labdhi Pandya, Afzal Ansari, Nikha Trivedi, R D Dixit, A M Kadri, Harsh Bakshi, Chaitanya Joshi, Madhvi Joshi |
| EPI_ISL_483825 | GMERS Medical College Himmatnagar | Gujarat Biotechnology Research Centre | Mayur Gandhi, Apurvasinh Puvar, Janvi Raval, Zarna Patel, Monika Gandhi, Pinal Trivedi, Maharshi Pandya, Nidhi Patel, Nitin Savaliya, Raghawendra Kumar, Dinesh Kumar, Zuber Saiyed, Komal Patel, Labdhi Pandya, Afzal Ansari, Nikha Trivedi, Himanshu Khatri, R D Dixit, A M Kadri, Harsh Bakshi, Chaitanya Joshi, Madhvi Joshi |
| EPI_ISL_483826 | GMERS Medical College Himmatnagar | Gujarat Biotechnology Research Centre | Apurvasinh Puvar, Janvi Raval, Zarna Patel, Monika Gandhi, Pinal Trivedi, Maharshi Pandya, Nidhi Patel, Nitin Savaliya, Raghawendra Kumar, Dinesh Kumar, Zuber Saiyed, Komal Patel, Labdhi Pandya, Afzal Ansari, Nikha Trivedi, Himanshu Khatri, Mayur Gandhi, R D Dixit, A M Kadri, Harsh Bakshi, Chaitanya Joshi, Madhvi Joshi |
| EPI_ISL_483827 | GMERS Medical College Himmatnagar | Gujarat Biotechnology Research Centre | Janvi Raval, Zarna Patel, Monika Gandhi, Pinal Trivedi, Maharshi Pandya, Nidhi Patel, Nitin Savaliya, Raghawendra Kumar, Dinesh Kumar, Zuber Saiyed, Komal Patel, Labdhi Pandya, Afzal Ansari, Nikha Trivedi, Himanshu Khatri, Mayur Gandhi, Apurvasinh Puvar, R D Dixit, A M Kadri, Harsh Bakshi, Chaitanya Joshi, Madhvi Joshi |
| EPI_ISL_483828 | GMERS Medical College Himmatnagar | Gujarat Biotechnology Research Centre | Zarna Patel, Monika Gandhi, Pinal Trivedi, Maharshi Pandya, Nidhi Patel, Nitin Savaliya, Raghawendra Kumar, Dinesh Kumar, Zuber Saiyed, Komal Patel, Labdhi Pandya, Afzal Ansari, Nikha Trivedi, Himanshu Khatri, Mayur Gandhi, Apurvasinh Puvar, Janvi Raval, R D Dixit, A M Kadri, Harsh Bakshi, Chaitanya Joshi, Madhvi Joshi |
| EPI_ISL_483829 | Pandit Deendayal Upadhyay Government Medical College, Rajkot | Gujarat Biotechnology Research Centre | Monika Gandhi, Pinal Trivedi, Maharshi Pandya, Nidhi Patel, Nitin Savaliya, Raghawendra Kumar, Dinesh Kumar, Zuber Saiyed, Komal Patel, Labdhi Pandya, Afzal Ansari, Nikha Trivedi, Gauravi Dhruv, Arti Trivedi, Apurvasinh Puvar, Janvi Raval, Zarna Patel, R D Dixit, A M Kadri, Harsh Bakshi, Chaitanya Joshi, Madhvi Joshi |
| EPI_ISL_483830 | Pandit Deendayal Upadhyay Government Medical College, Rajkot | Gujarat Biotechnology Research Centre | Pinal Trivedi, Maharshi Pandya, Nidhi Patel, Nitin Savaliya, Raghawendra Kumar, Dinesh Kumar, Zuber Saiyed, Komal Patel, Labdhi Pandya, Afzal Ansari, Nikha Trivedi, Gauravi Dhruv, Arti Trivedi, Apurvasinh Puvar, Janvi Raval, Zarna Patel, Monika Gandhi, R D Dixit, A M Kadri, Harsh Bakshi, Chaitanya Joshi, Madhvi Joshi |
| EPI_ISL_483831 | Pandit Deendayal Upadhyay Government Medical College, Rajkot | Gujarat Biotechnology Research Centre | Maharshi Pandya, Nidhi Patel, Nitin Savaliya, Raghawendra Kumar, Dinesh Kumar, Zuber Saiyed, Komal Patel, Labdhi Pandya, Afzal Ansari, Nikha Trivedi, Gauravi Dhruv, Arti Trivedi, Apurvasinh Puvar, Janvi Raval, Zarna Patel, Monika Gandhi, Pinal Trivedi, R D Dixit, A M Kadri, Harsh Bakshi, Chaitanya Joshi, Madhvi Joshi |
| EPI_ISL_483832 | Pandit Deendayal Upadhyay Government Medical College, Rajkot | Gujarat Biotechnology Research Centre | Nidhi Patel, Nitin Savaliya, Raghawendra Kumar, Dinesh Kumar, Zuber Saiyed, Komal Patel, Labdhi Pandya, Afzal Ansari, Nikha Trivedi, Gauravi Dhruv, Arti Trivedi, Apurvasinh Puvar, Janvi Raval, Zarna Patel, Monika Gandhi, Pinal Trivedi, Maharshi Pandya, R D Dixit, A M Kadri, Harsh Bakshi, Chaitanya Joshi, Madhvi Joshi |
| EPI_ISL_483833 | Pandit Deendayal Upadhyay Government Medical College, Rajkot | Gujarat Biotechnology Research Centre | Nitin Savaliya, Raghawendra Kumar, Dinesh Kumar, Zuber Saiyed, Komal Patel, Labdhi Pandya, Afzal Ansari, Nikha Trivedi, Gauravi Dhruv, Arti Trivedi, Apurvasinh Puvar, Janvi Raval, Zarna Patel, Monika Gandhi, Pinal Trivedi, Maharshi Pandya, Nidhi Patel, R D Dixit, A M Kadri, Harsh Bakshi, Chaitanya Joshi, Madhvi Joshi |

| | | | |
|---|---|---|---|
| EPI_ISL_483834 | Pandit Deendayal Upadhyay Government Medical College, Rajkot | Gujarat Biotechnology Research Centre | Raghawendra Kumar, Dinesh Kumar, Zuber Saiyed, Komal Patel, Labdhi Pandya, Afzal Ansari, Nikha Trivedi, Gauravi Dhruv, Arti Trivedi, Apurvasinh Puvar, Janvi Raval, Zarna Patel, Monika Gandhi, Pinal Trivedi, Maharshi Pandya, Nidhi Patel, Nitin Savaliya, R D Dixit, A M Kadri, Harsh Bakshi, Chaitanya Joshi, Madhvi Joshi |
| EPI_ISL_483835 | Pandit Deendayal Upadhyay Government Medical College, Rajkot | Gujarat Biotechnology Research Centre | Dinesh Kumar, Zuber Saiyed, Komal Patel, Labdhi Pandya, Afzal Ansari, Nikha Trivedi, Gauravi Dhruv, Arti Trivedi, Apurvasinh Puvar, Janvi Raval, Zarna Patel, Monika Gandhi, Pinal Trivedi, Maharshi Pandya, Nidhi Patel, Nitin Savaliya, Raghawendra Kumar, R D Dixit, A M Kadri, Harsh Bakshi, Chaitanya Joshi, Madhvi Joshi |
| EPI_ISL_483836 | Pandit Deendayal Upadhyay Government Medical College, Rajkot | Gujarat Biotechnology Research Centre | Zuber Saiyed, Komal Patel, Labdhi Pandya, Afzal Ansari, Nikha Trivedi, Gauravi Dhruv, Arti Trivedi, Apurvasinh Puvar, Janvi Raval, Zarna Patel, Monika Gandhi, Pinal Trivedi, Maharshi Pandya, Nidhi Patel, Nitin Savaliya, Raghawendra Kumar, Dinesh Kumar, R D Dixit, A M Kadri, Harsh Bakshi, Chaitanya Joshi, Madhvi Joshi |
| EPI_ISL_483837 | Pandit Deendayal Upadhyay Government Medical College, Rajkot | Gujarat Biotechnology Research Centre | Komal Patel, Labdhi Pandya, Afzal Ansari, Nikha Trivedi, Gauravi Dhruv, Arti Trivedi, Apurvasinh Puvar, Janvi Raval, Zarna Patel, Monika Gandhi, Pinal Trivedi, Maharshi Pandya, Nidhi Patel, Nitin Savaliya, Raghawendra Kumar, Dinesh Kumar, Zuber Saiyed, R D Dixit, A M Kadri, Harsh Bakshi, Chaitanya Joshi, Madhvi Joshi |
| EPI_ISL_483838 | Pandit Deendayal Upadhyay Government Medical College, Rajkot | Gujarat Biotechnology Research Centre | Labdhi Pandya, Afzal Ansari, Nikha Trivedi, Gauravi Dhruv, Arti Trivedi, Apurvasinh Puvar, Janvi Raval, Zarna Patel, Monika Gandhi, Pinal Trivedi, Maharshi Pandya, Nidhi Patel, Nitin Savaliya, Raghawendra Kumar, Dinesh Kumar, Zuber Saiyed, Komal Patel, R D Dixit, A M Kadri, Harsh Bakshi, Chaitanya Joshi, Madhvi Joshi |
| EPI_ISL_483839 | Pandit Deendayal Upadhyay Government Medical College, Rajkot | Gujarat Biotechnology Research Centre | Afzal Ansari, Nikha Trivedi, Gauravi Dhruv, Arti Trivedi, Apurvasinh Puvar, Janvi Raval, Zarna Patel, Monika Gandhi, Pinal Trivedi, Maharshi Pandya, Nidhi Patel, Nitin Savaliya, Raghawendra Kumar, Dinesh Kumar, Zuber Saiyed, Komal Patel, Labdhi Pandya, R D Dixit, A M Kadri, Harsh Bakshi, Chaitanya Joshi, Madhvi Joshi |
| EPI_ISL_483840 | Pandit Deendayal Upadhyay Government Medical College, Rajkot | Gujarat Biotechnology Research Centre | Nikha Trivedi, Gauravi Dhruv, Arti Trivedi, Apurvasinh Puvar, Janvi Raval, Zarna Patel, Monika Gandhi, Pinal Trivedi, Maharshi Pandya, Nidhi Patel, Nitin Savaliya, Raghawendra Kumar, Dinesh Kumar, Zuber Saiyed, Komal Patel, Labdhi Pandya, Afzal Ansari, R D Dixit, A M Kadri, Harsh Bakshi, Chaitanya Joshi, Madhvi Joshi |
| EPI_ISL_483841 | Pandit Deendayal Upadhyay Government Medical College, Rajkot | Gujarat Biotechnology Research Centre | Gauravi Dhruv, Arti Trivedi, Apurvasinh Puvar, Janvi Raval, Zarna Patel, Monika Gandhi, Pinal Trivedi, Maharshi Pandya, Nidhi Patel, Nitin Savaliya, Raghawendra Kumar, Dinesh Kumar, Zuber Saiyed, Komal Patel, Labdhi Pandya, Afzal Ansari, Nikha Trivedi, R D Dixit, A M Kadri, Harsh Bakshi, Chaitanya Joshi, Madhvi Joshi |
| EPI_ISL_483842 | Pandit Deendayal Upadhyay Government Medical College, Rajkot | Gujarat Biotechnology Research Centre | Arti Trivedi, Apurvasinh Puvar, Janvi Raval, Zarna Patel, Monika Gandhi, Pinal Trivedi, Maharshi Pandya, Nidhi Patel, Nitin Savaliya, Raghawendra Kumar, Dinesh Kumar, Zuber Saiyed, Komal Patel, Labdhi Pandya, Afzal Ansari, Nikha Trivedi, Gauravi Dhruv, R D Dixit, A M Kadri, Harsh Bakshi, Chaitanya Joshi, Madhvi Joshi |
| EPI_ISL_483843 | Pandit Deendayal Upadhyay Government Medical College, Rajkot | Gujarat Biotechnology Research Centre | Apurvasinh Puvar, Janvi Raval, Zarna Patel, Monika Gandhi, Pinal Trivedi, Maharshi Pandya, Nidhi Patel, Nitin Savaliya, Raghawendra Kumar, Dinesh Kumar, Zuber Saiyed, Komal Patel, Labdhi Pandya, Afzal Ansari, Nikha Trivedi, Gauravi Dhruv, Arti Trivedi, R D Dixit, A M Kadri, Harsh Bakshi, Chaitanya Joshi, Madhvi Joshi |
| EPI_ISL_483844 | Pandit Deendayal Upadhyay Government Medical College, Rajkot | Gujarat Biotechnology Research Centre | Janvi Raval, Zarna Patel, Monika Gandhi, Pinal Trivedi, Maharshi Pandya, Nidhi Patel, Nitin Savaliya, Raghawendra Kumar, Dinesh Kumar, Zuber Saiyed, Komal Patel, Labdhi Pandya, Afzal Ansari, Nikha Trivedi, Gauravi Dhruv, Arti Trivedi, Apurvasinh Puvar, R D Dixit, A M Kadri, Harsh Bakshi, Chaitanya Joshi, Madhvi Joshi |
| EPI_ISL_483845 | Pandit Deendayal Upadhyay Government Medical College, Rajkot | Gujarat Biotechnology Research Centre | Zarna Patel, Monika Gandhi, Pinal Trivedi, Maharshi Pandya, Nidhi Patel, Nitin Savaliya, Raghawendra Kumar, Dinesh Kumar, Zuber Saiyed, Komal Patel, Labdhi Pandya, Afzal Ansari, Nikha Trivedi, Gauravi Dhruv, Arti Trivedi, Apurvasinh Puvar, Janvi Raval, R D Dixit, A M Kadri, Harsh Bakshi, Chaitanya Joshi, Madhvi Joshi |
| EPI_ISL_483846 | Pandit Deendayal Upadhyay Government Medical College, Rajkot | Gujarat Biotechnology Research Centre | Monika Gandhi, Pinal Trivedi, Maharshi Pandya, Nidhi Patel, Nitin Savaliya, Raghawendra Kumar, Dinesh Kumar, Zuber Saiyed, Komal Patel, Labdhi Pandya, Afzal Ansari, Nikha Trivedi, Gauravi Dhruv, Arti Trivedi, Apurvasinh Puvar, Janvi Raval, Zarna Patel, R D Dixit, A M Kadri, Harsh Bakshi, Chaitanya Joshi, Madhvi Joshi |
| EPI_ISL_483847 | Pandit Deendayal Upadhyay Government Medical College, Rajkot | Gujarat Biotechnology Research Centre | Pinal Trivedi, Maharshi Pandya, Nidhi Patel, Nitin Savaliya, Raghawendra Kumar, Dinesh Kumar, Zuber Saiyed, Komal Patel, Labdhi Pandya, Afzal Ansari, Nikha Trivedi, Gauravi Dhruv, Arti Trivedi, Apurvasinh Puvar, Janvi Raval, Zarna Patel, Monika Gandhi, R D Dixit, A M Kadri, Harsh Bakshi, Chaitanya Joshi, Madhvi Joshi |
| EPI_ISL_483848 | Pandit Deendayal Upadhyay Government Medical College, Rajkot | Gujarat Biotechnology Research Centre | Maharshi Pandya, Nidhi Patel, Nitin Savaliya, Raghawendra Kumar, Dinesh Kumar, Zuber Saiyed, Komal Patel, Labdhi Pandya, Afzal Ansari, Nikha Trivedi, Gauravi Dhruv, Arti Trivedi, Apurvasinh Puvar, Janvi Raval, Zarna Patel, Monika Gandhi, Pinal Trivedi, R D Dixit, A M Kadri, Harsh Bakshi, Chaitanya Joshi, Madhvi Joshi |
| EPI_ISL_483849 | Pandit Deendayal Upadhyay Government Medical College, Rajkot | Gujarat Biotechnology Research Centre | Nidhi Patel, Nitin Savaliya, Raghawendra Kumar, Dinesh Kumar, Zuber Saiyed, Komal Patel, Labdhi Pandya, Afzal Ansari, Nikha Trivedi, Gauravi Dhruv, Arti Trivedi, Apurvasinh Puvar, Janvi Raval, Zarna Patel, Monika Gandhi, Pinal Trivedi, Maharshi Pandya, R D Dixit, A M Kadri, Harsh Bakshi, Chaitanya Joshi, Madhvi Joshi |
| EPI_ISL_483850 | Department of Microbiology, Government Medical College, Surat | Gujarat Biotechnology Research Centre | Nitin Savaliya, Raghawendra Kumar, Dinesh Kumar, Zuber Saiyed, Komal Patel, Labdhi Pandya, Afzal Ansari, Nikha Trivedi, Naresh Chauhan, Summaiya Mullan, Amit gamit, Apurvasinh Puvar, Janvi Raval, Zarna Patel, Monika Gandhi, Pinal Trivedi, Maharshi Pandya, Nidhi Patel, R D Dixit, A M Kadri, Harsh Bakshi, Chaitanya Joshi, Madhvi Joshi |
| EPI_ISL_483851 | Department of Microbiology, Government Medical College, Surat | Gujarat Biotechnology Research Centre | Raghawendra Kumar, Dinesh Kumar, Zuber Saiyed, Komal Patel, Labdhi Pandya, Afzal Ansari, Nikha Trivedi, Naresh Chauhan, Summaiya Mullan, Amit gamit, Apurvasinh Puvar, Janvi Raval, Zarna Patel, Monika Gandhi, Pinal Trivedi, Maharshi Pandya, Nidhi Patel, Nitin Savaliya, R D Dixit, A M Kadri, Harsh Bakshi, Chaitanya Joshi, Madhvi Joshi |
| EPI_ISL_483852 | Department of Microbiology, Government Medical College, Surat | Gujarat Biotechnology Research Centre | Dinesh Kumar, Zuber Saiyed, Komal Patel, Labdhi Pandya, Afzal Ansari, Nikha Trivedi, Naresh Chauhan, Summaiya Mullan, Amit gamit, Apurvasinh Puvar, Janvi Raval, Zarna Patel, Monika Gandhi, Pinal Trivedi, Maharshi Pandya, Nidhi Patel, Nitin Savaliya, Raghawendra Kumar, R D Dixit, A M Kadri, Harsh Bakshi, Chaitanya Joshi, Madhvi Joshi |
| EPI_ISL_483853 | Department of Microbiology, Government Medical College, Surat | Gujarat Biotechnology Research Centre | Zuber Saiyed, Komal Patel, Labdhi Pandya, Afzal Ansari, Nikha Trivedi, Naresh Chauhan, Summaiya Mullan, Amit gamit, Apurvasinh Puvar, Janvi Raval, Zarna Patel, Monika Gandhi, Pinal Trivedi, Maharshi Pandya, Nidhi Patel, Nitin Savaliya, Raghawendra Kumar, Dinesh Kumar, R D Dixit, A M Kadri, Harsh Bakshi, Chaitanya Joshi, Madhvi Joshi |
| EPI_ISL_483854 | Department of Microbiology, Government Medical College, Surat | Gujarat Biotechnology Research Centre | Komal Patel, Labdhi Pandya, Afzal Ansari, Nikha Trivedi, Naresh Chauhan, Summaiya Mullan, Amit gamit, Apurvasinh Puvar, Janvi Raval, Zarna Patel, Monika Gandhi, Pinal Trivedi, Maharshi Pandya, Nidhi Patel, Nitin Savaliya, Raghawendra Kumar, Dinesh Kumar, Zuber Saiyed, R D Dixit, A M Kadri, Harsh Bakshi, Chaitanya Joshi, Madhvi Joshi |
| EPI_ISL_483855 | Department of Microbiology, Government Medical College, Surat | Gujarat Biotechnology Research Centre | Labdhi Pandya, Afzal Ansari, Nikha Trivedi, Naresh Chauhan, Summaiya Mullan, Amit gamit, Apurvasinh Puvar, Janvi Raval, Zarna Patel, Monika Gandhi, Pinal Trivedi, Maharshi Pandya, Nidhi Patel, Nitin Savaliya, Raghawendra Kumar, Dinesh Kumar, Zuber Saiyed, Komal Patel, R D Dixit, A M Kadri, Harsh Bakshi, Chaitanya Joshi, Madhvi Joshi |
| EPI_ISL_483856 | Department of Microbiology, Government Medical College, Surat | Gujarat Biotechnology Research Centre | Afzal Ansari, Nikha Trivedi, Naresh Chauhan, Summaiya Mullan, Amit gamit, Apurvasinh Puvar, Janvi Raval, Zarna Patel, Monika Gandhi, Pinal Trivedi, Maharshi Pandya, Nidhi Patel, Nitin Savaliya, Raghawendra Kumar, Dinesh Kumar, Zuber Saiyed, Komal Patel, Labdhi Pandya, R D Dixit, A M Kadri, Harsh Bakshi, Chaitanya Joshi, Madhvi Joshi |
| EPI_ISL_483857 | Department of Microbiology, Government Medical College, Surat | Gujarat Biotechnology Research Centre | Nikha Trivedi, Naresh Chauhan, Summaiya Mullan, Amit gamit, Apurvasinh Puvar, Janvi Raval, Zarna Patel, Monika Gandhi, Pinal Trivedi, Maharshi Pandya, Nidhi Patel, Nitin Savaliya, Raghawendra Kumar, Dinesh Kumar, Zuber Saiyed, Komal Patel, Labdhi Pandya, Afzal Ansari, R D Dixit, A M Kadri, Harsh Bakshi, Chaitanya Joshi, Madhvi Joshi |
| EPI_ISL_483858 | Department of Microbiology, Government Medical College, Surat | Gujarat Biotechnology Research Centre | Naresh Chauhan, Summaiya Mullan, Amit gamit, Apurvasinh Puvar, Janvi Raval, Zarna Patel, Monika Gandhi, Pinal Trivedi, Maharshi Pandya, Nidhi Patel, Nitin Savaliya, Raghawendra Kumar, Dinesh Kumar, Zuber Saiyed, Komal Patel, Labdhi Pandya, Afzal Ansari, Nikha Trivedi, R D Dixit, A M Kadri, Harsh Bakshi, Chaitanya Joshi, Madhvi Joshi |
| EPI_ISL_483859 | Department of Microbiology, Government Medical College, Surat | Gujarat Biotechnology Research Centre | Summaiya Mullan, Amit gamit, Apurvasinh Puvar, Janvi Raval, Zarna Patel, Monika Gandhi, Pinal Trivedi, Maharshi Pandya, Nidhi Patel, Nitin Savaliya, Raghawendra Kumar, Dinesh Kumar, Zuber Saiyed, Komal Patel, Labdhi Pandya, Afzal Ansari, Nikha Trivedi, Naresh Chauhan, R D Dixit, A M Kadri, Harsh Bakshi, Chaitanya Joshi, Madhvi Joshi |
| EPI_ISL_483860 | Department of Microbiology, Government Medical College, Surat | Gujarat Biotechnology Research Centre | Amit gamit, Apurvasinh Puvar, Janvi Raval, Zarna Patel, Monika Gandhi, Pinal Trivedi, Maharshi Pandya, Nidhi Patel, Nitin Savaliya, Raghawendra Kumar, Dinesh Kumar, Zuber Saiyed, Komal Patel, Labdhi Pandya, Afzal Ansari, Nikha Trivedi, Naresh Chauhan, Summaiya Mullan, R D Dixit, A M Kadri, Harsh Bakshi, Chaitanya Joshi, Madhvi Joshi |
| EPI_ISL_483861 | Department of Microbiology, Government Medical College, Surat | Gujarat Biotechnology Research Centre | Apurvasinh Puvar, Janvi Raval, Zarna Patel, Monika Gandhi, Pinal Trivedi, Maharshi Pandya, Nidhi Patel, Nitin Savaliya, Raghawendra Kumar, Dinesh Kumar, Zuber Saiyed, Komal Patel, Labdhi Pandya, Afzal Ansari, Nikha Trivedi, Naresh Chauhan, Summaiya Mullan, Amit gamit, R D Dixit, A M Kadri, Harsh Bakshi, Chaitanya Joshi, Madhvi Joshi |
| EPI_ISL_483862 | Department of Microbiology, Government Medical College, Surat | Gujarat Biotechnology Research Centre | Janvi Raval, Zarna Patel, Monika Gandhi, Pinal Trivedi, Maharshi Pandya, Nidhi Patel, Nitin Savaliya, Raghawendra Kumar, Dinesh Kumar, Zuber Saiyed, Komal Patel, Labdhi Pandya, Afzal Ansari, Nikha Trivedi, Naresh Chauhan, Summaiya Mullan, Amit gamit, Apurvasinh Puvar, R D Dixit, A M Kadri, Harsh Bakshi, Chaitanya Joshi, Madhvi Joshi |
| EPI_ISL_483863 | Department of Microbiology, Government Medical College, Surat | Gujarat Biotechnology Research Centre | Zarna Patel, Monika Gandhi, Pinal Trivedi, Maharshi Pandya, Nidhi Patel, Nitin Savaliya, Raghawendra Kumar, Dinesh Kumar, Zuber Saiyed, Komal Patel, Labdhi Pandya, Afzal Ansari, Nikha Trivedi, Naresh Chauhan, Summaiya Mullan, Amit gamit, Apurvasinh Puvar, Janvi Raval, R D Dixit, A M Kadri, Harsh Bakshi, Chaitanya Joshi, Madhvi Joshi |
| EPI_ISL_483864 | Department of Microbiology, Government Medical College, Surat | Gujarat Biotechnology Research Centre | Monika Gandhi, Pinal Trivedi, Maharshi Pandya, Nidhi Patel, Nitin Savaliya, Raghawendra Kumar, Dinesh Kumar, Zuber Saiyed, Komal Patel, Labdhi Pandya, Afzal Ansari, Nikha Trivedi, Naresh Chauhan, Summaiya Mullan, Amit gamit, Apurvasinh Puvar, Janvi Raval, Zarna Patel, R D Dixit, A M Kadri, Harsh Bakshi, Chaitanya Joshi, Madhvi Joshi |
| EPI_ISL_483865 | Department of Microbiology, Government Medical College, Surat | Gujarat Biotechnology Research Centre | Pinal Trivedi, Maharshi Pandya, Nidhi Patel, Nitin Savaliya, Raghawendra Kumar, Dinesh Kumar, Zuber Saiyed, Komal Patel, Labdhi Pandya, Afzal Ansari, Nikha Trivedi, Naresh Chauhan, Summaiya Mullan, Amit gamit, Apurvasinh Puvar, Janvi Raval, Zarna Patel, Monika Gandhi, R D Dixit, A M Kadri, Harsh Bakshi, Chaitanya Joshi, Madhvi Joshi |
| EPI_ISL_483866 | Department of Microbiology, Government Medical College, Surat | Gujarat Biotechnology Research Centre | Maharshi Pandya, Nidhi Patel, Nitin Savaliya, Raghawendra Kumar, Dinesh Kumar, Zuber Saiyed, Komal Patel, Labdhi Pandya, Afzal Ansari, Nikha Trivedi, Naresh Chauhan, Summaiya Mullan, Amit gamit, Apurvasinh Puvar, Janvi Raval, Zarna Patel, Monika Gandhi, Pinal Trivedi, R D Dixit, A M Kadri, Harsh Bakshi, Chaitanya Joshi, Madhvi Joshi |
| EPI_ISL_483867 | Department of Microbiology, Government Medical College, Surat | Gujarat Biotechnology Research Centre | Nidhi Patel, Nitin Savaliya, Raghawendra Kumar, Dinesh Kumar, Zuber Saiyed, Komal Patel, Labdhi Pandya, Afzal Ansari, Nikha Trivedi, Naresh Chauhan, Summaiya Mullan, Amit gamit, Apurvasinh Puvar, Janvi Raval, Zarna Patel, Monika Gandhi, Pinal Trivedi, Maharshi Pandya, R D Dixit, A M Kadri, Harsh Bakshi, Chaitanya Joshi, Madhvi Joshi |
| EPI_ISL_483868 | Department of Microbiology, Government Medical College, Surat | Gujarat Biotechnology Research Centre | Nitin Savaliya, Raghawendra Kumar, Dinesh Kumar, Zuber Saiyed, Komal Patel, Labdhi Pandya, Afzal Ansari, Nikha Trivedi, Naresh Chauhan, Summaiya Mullan, Amit gamit, Apurvasinh Puvar, Janvi Raval, Zarna Patel, Monika Gandhi, Pinal Trivedi, Maharshi Pandya, Nidhi Patel, R D Dixit, A M Kadri, Harsh Bakshi, Chaitanya Joshi, Madhvi Joshi |
| EPI_ISL_483869 | Department of Microbiology, Government Medical College, Surat | Gujarat Biotechnology Research Centre | Raghawendra Kumar, Dinesh Kumar, Zuber Saiyed, Komal Patel, Labdhi Pandya, Afzal Ansari, Nikha Trivedi, Naresh Chauhan, Summaiya Mullan, Amit gamit, Apurvasinh Puvar, Janvi Raval, Zarna Patel, Monika Gandhi, Pinal Trivedi, Maharshi Pandya, Nidhi Patel, Nitin Savaliya, R D Dixit, A M Kadri, Harsh Bakshi, Chaitanya Joshi, Madhvi Joshi |
| EPI_ISL_483870 | Department of Microbiology, Government Medical College, Surat | Gujarat Biotechnology Research Centre | Dinesh Kumar, Zuber Saiyed, Komal Patel, Labdhi Pandya, Afzal Ansari, Nikha Trivedi, Naresh Chauhan, Summaiya Mullan, Amit gamit, Apurvasinh Puvar, Janvi Raval, Zarna Patel, Monika Gandhi, Pinal Trivedi, Maharshi Pandya, Nidhi Patel, Nitin Savaliya, Raghawendra Kumar, R D Dixit, A M Kadri, Harsh Bakshi, Chaitanya Joshi, Madhvi Joshi |
| EPI_ISL_483871 | Department of Microbiology, Government Medical College, Surat | Gujarat Biotechnology Research Centre | Zuber Saiyed, Komal Patel, Labdhi Pandya, Afzal Ansari, Nikha Trivedi, Naresh Chauhan, Summaiya Mullan, Amit gamit, Apurvasinh Puvar, Janvi Raval, Zarna Patel, Monika Gandhi, Pinal Trivedi, Maharshi Pandya, Nidhi Patel, Nitin Savaliya, Raghawendra Kumar, Dinesh Kumar, R D Dixit, A M Kadri, Harsh Bakshi, Chaitanya Joshi, Madhvi Joshi |
| EPI_ISL_483872 | Department of Microbiology, Government Medical College, Surat | Gujarat Biotechnology Research Centre | Komal Patel, Labdhi Pandya, Afzal Ansari, Nikha Trivedi, Naresh Chauhan, Summaiya Mullan, Amit gamit, Apurvasinh Puvar, Janvi Raval, Zarna Patel, Monika Gandhi, Pinal Trivedi, Maharshi Pandya, Nidhi Patel, Nitin Savaliya, Raghawendra Kumar, Dinesh Kumar, Zuber Saiyed, R D Dixit, A M Kadri, Harsh Bakshi, Chaitanya Joshi, Madhvi Joshi |
| EPI_ISL_483873 | Department of Microbiology, Government Medical College, Surat | Gujarat Biotechnology Research Centre | Labdhi Pandya, Afzal Ansari, Nikha Trivedi, Naresh Chauhan, Summaiya Mullan, Amit gamit, Apurvasinh Puvar, Janvi Raval, Zarna Patel, Monika Gandhi, Pinal Trivedi, Maharshi Pandya, Nidhi Patel, Nitin Savaliya, Raghawendra Kumar, Dinesh Kumar, Zuber Saiyed, Komal Patel, R D Dixit, A M Kadri, Harsh Bakshi, Chaitanya Joshi, Madhvi Joshi |
| EPI_ISL_483874 | Department of Microbiology, Government Medical College, Surat | Gujarat Biotechnology Research Centre | Afzal Ansari, Nikha Trivedi, Naresh Chauhan, Summaiya Mullan, Amit gamit, Apurvasinh Puvar, Janvi Raval, Zarna Patel, Monika Gandhi, Pinal Trivedi, Maharshi Pandya, Nidhi Patel, Nitin Savaliya, Raghawendra Kumar, Dinesh Kumar, Zuber Saiyed, Komal Patel, Labdhi Pandya, R D Dixit, A M Kadri, Harsh Bakshi, Chaitanya Joshi, Madhvi Joshi |
| EPI_ISL_483875 | Department of Microbiology, Government Medical College, Surat | Gujarat Biotechnology Research Centre | Nikha Trivedi, Naresh Chauhan, Summaiya Mullan, Amit gamit, Apurvasinh Puvar, Janvi Raval, Zarna Patel, Monika Gandhi, Pinal Trivedi, Maharshi Pandya, Nidhi Patel, Nitin Savaliya, Raghawendra Kumar, Dinesh Kumar, Zuber Saiyed, Komal Patel, Labdhi Pandya, Afzal Ansari, R D Dixit, A M Kadri, Harsh Bakshi, Chaitanya Joshi, Madhvi Joshi |
| EPI_ISL_483876 | Department of Microbiology, Government Medical College, Surat | Gujarat Biotechnology Research Centre | Naresh Chauhan, Summaiya Mullan, Amit gamit, Apurvasinh Puvar, Janvi Raval, Zarna Patel, Monika Gandhi, Pinal Trivedi, Maharshi Pandya, Nidhi Patel, Nitin Savaliya, Raghawendra Kumar, Dinesh Kumar, Zuber Saiyed, Komal Patel, Labdhi Pandya, Afzal Ansari, Nikha Trivedi, R D Dixit, A M Kadri, Harsh Bakshi, Chaitanya Joshi, Madhvi Joshi |
| EPI_ISL_483877 | Department of Microbiology, Government Medical College, Surat | Gujarat Biotechnology Research Centre | Summaiya Mullan, Amit gamit, Apurvasinh Puvar, Janvi Raval, Zarna Patel, Monika Gandhi, Pinal Trivedi, Maharshi Pandya, Nidhi Patel, Nitin Savaliya, Raghawendra Kumar, Dinesh Kumar, Zuber Saiyed, Komal Patel, Labdhi Pandya, Afzal Ansari, Nikha Trivedi, Naresh Chauhan, R D Dixit, A M Kadri, Harsh Bakshi, Chaitanya Joshi, Madhvi Joshi |
| EPI_ISL_483878 | Department of Microbiology, Government Medical College, Surat | Gujarat Biotechnology Research Centre | Amit gamit, Apurvasinh Puvar, Janvi Raval, Zarna Patel, Monika Gandhi, Pinal Trivedi, Maharshi Pandya, Nidhi Patel, Nitin Savaliya, Raghawendra Kumar, Dinesh Kumar, Zuber Saiyed, Komal Patel, Labdhi Pandya, Afzal Ansari, Nikha Trivedi, Naresh Chauhan, Summaiya Mullan, R D Dixit, A M Kadri, Harsh Bakshi, Chaitanya Joshi, Madhvi Joshi |

| | | | |
|---|---|---|---|
| EPI_ISL_483879 | Department of Microbiology, Government Medical College, Surat | Gujarat Biotechnology Research Centre | Apurvasinh Puvar, Janvi Raval, Zarna Patel, Monika Gandhi, Pinal Trivedi, Maharshi Pandya, Nidhi Patel, Nitin Savaliya, Raghawendra Kumar, Dinesh Kumar, Zuber Saiyed, Komal Patel, Labdhi Pandya, Afzal Ansari, Nikha Trivedi, Naresh Chauhan, Summaiya Mullan, Amit gamit, R D Dixit, A M Kadri, Harsh Bakshi, Chaitanya Joshi, Madhvi Joshi |
| EPI_ISL_483880, EPI_ISL_483881, EPI_ISL_483882, EPI_ISL_483883, EPI_ISL_483884, EPI_ISL_483885, EPI_ISL_483886, EPI_ISL_483887, EPI_ISL_483888, EPI_ISL_483889, EPI_ISL_483890, EPI_ISL_483891, EPI_ISL_483892, EPI_ISL_483893, EPI_ISL_483894, EPI_ISL_483895, EPI_ISL_483896, EPI_ISL_483897, EPI_ISL_483898, EPI_ISL_483899, EPI_ISL_483900, EPI_ISL_483901, EPI_ISL_483902, EPI_ISL_483903, EPI_ISL_483904, EPI_ISL_483905, EPI_ISL_483906, EPI_ISL_483907, EPI_ISL_483908, EPI_ISL_483909, EPI_ISL_483910, EPI_ISL_483911, EPI_ISL_483912 | | | |
| see above | University of Birmingham | COVID-19 Genomics UK (COG-UK) Consortium | Institute of Microbiology, University of Birmingham: Claire McMurray, Joanne Stockton, Samuel Nicholls, Radoslaw Poplawski, Will Rowe, Josh Quick, Nicholas Loman. University of Birmingham Testing Laboratory: Celina M Whalley, Andrew Bosworth, Charlotte Poxon, Kasun Wanigasooriya, Oliver Pickles, Mike Kidd, Alex Richter, Andrew D Beggs PHE Heartlands Lab: Husam Osman, Andrew Bosworth. Queen Elizabeth Hospital: Anna Casey |
| EPI_ISL_483913, EPI_ISL_483914, EPI_ISL_483915 | Department of Pathology, University of Cambridge | COVID-19 Genomics UK (COG-UK) Consortium | Luke W Meredith, M. Estée Török, Myra Hosmillo, William L. Hamilton, Martin D. Curran, Theresa Feltwell, Grant Hall, Anna Yakovleva, Fahad A Khokhar, Charlotte J. Houldcroft, Laura G Caller, Aminu S. Jahun, Sarah L. Caddy, Yasmin Chaudhry, Malte Pinckert, Ian Goodfellow |
| EPI_ISL_483916, EPI_ISL_483917, EPI_ISL_483918, EPI_ISL_483919, EPI_ISL_483920, EPI_ISL_483921, EPI_ISL_483922, EPI_ISL_483923, EPI_ISL_483924, EPI_ISL_483925, EPI_ISL_483926, EPI_ISL_483927, EPI_ISL_483928, EPI_ISL_483929, EPI_ISL_483930, EPI_ISL_483931, EPI_ISL_483932, EPI_ISL_483933, EPI_ISL_483934, EPI_ISL_483935, EPI_ISL_483936, EPI_ISL_483937, EPI_ISL_483938, EPI_ISL_483939, EPI_ISL_483940, EPI_ISL_483941, EPI_ISL_483942, EPI_ISL_483943, EPI_ISL_483944, EPI_ISL_483945, EPI_ISL_483946, EPI_ISL_483947, EPI_ISL_483948, EPI_ISL_483949, EPI_ISL_483950, EPI_ISL_483951, EPI_ISL_483952, EPI_ISL_483953, EPI_ISL_483954, EPI_ISL_483955, EPI_ISL_483956, EPI_ISL_483957, EPI_ISL_483958, EPI_ISL_483959, EPI_ISL_483960, EPI_ISL_483961, EPI_ISL_483962, EPI_ISL_483963, EPI_ISL_483964, EPI_ISL_483965, EPI_ISL_483966, EPI_ISL_483967, EPI_ISL_483968, EPI_ISL_483969, EPI_ISL_483970, EPI_ISL_483971, EPI_ISL_483972, EPI_ISL_483973, EPI_ISL_483974, EPI_ISL_483975, EPI_ISL_483976, EPI_ISL_483977, EPI_ISL_483978, EPI_ISL_483979, EPI_ISL_483980, EPI_ISL_483981, EPI_ISL_483982, EPI_ISL_483983, EPI_ISL_483984, EPI_ISL_483985, EPI_ISL_483986, EPI_ISL_483987, EPI_ISL_483988, EPI_ISL_483989, EPI_ISL_483990, EPI_ISL_483991, EPI_ISL_483992, EPI_ISL_483993, EPI_ISL_483994, EPI_ISL_483995, EPI_ISL_483996, EPI_ISL_483997, EPI_ISL_483998, EPI_ISL_483999, EPI_ISL_484000, EPI_ISL_484001, EPI_ISL_484002, EPI_ISL_484003, EPI_ISL_484004, EPI_ISL_484005, EPI_ISL_484006, EPI_ISL_484007, EPI_ISL_484008, EPI_ISL_484009, EPI_ISL_484010, EPI_ISL_484011, EPI_ISL_484012, EPI_ISL_484013, EPI_ISL_484014, EPI_ISL_484015, EPI_ISL_484016, EPI_ISL_484017, EPI_ISL_484018, EPI_ISL_484019, EPI_ISL_484020, EPI_ISL_484021, EPI_ISL_484022, EPI_ISL_484023, EPI_ISL_484024, EPI_ISL_484025, EPI_ISL_484026, EPI_ISL_484027, EPI_ISL_484028, EPI_ISL_484029, EPI_ISL_484030, EPI_ISL_484031, EPI_ISL_484032, EPI_ISL_484033, EPI_ISL_484034, EPI_ISL_484035, EPI_ISL_484036, EPI_ISL_484037, EPI_ISL_484038, EPI_ISL_484039, EPI_ISL_484040, EPI_ISL_484041, EPI_ISL_484042, EPI_ISL_484043, EPI_ISL_484044, EPI_ISL_484045, EPI_ISL_484046, EPI_ISL_484047, EPI_ISL_484048, EPI_ISL_484049, EPI_ISL_484050, EPI_ISL_484051, EPI_ISL_484052, EPI_ISL_484053, EPI_ISL_484054, EPI_ISL_484055, EPI_ISL_484056, EPI_ISL_484057, EPI_ISL_484058, EPI_ISL_484059, EPI_ISL_484060, EPI_ISL_484061, EPI_ISL_484062, EPI_ISL_484063, EPI_ISL_484064, EPI_ISL_484065, EPI_ISL_484066, EPI_ISL_484067, EPI_ISL_484068, EPI_ISL_484069, EPI_ISL_484070, EPI_ISL_484071, EPI_ISL_484072, EPI_ISL_484073, EPI_ISL_484074, EPI_ISL_484075, EPI_ISL_484076, EPI_ISL_484077, EPI_ISL_484078, EPI_ISL_484079, EPI_ISL_484080, EPI_ISL_484081, EPI_ISL_484082, EPI_ISL_484083, EPI_ISL_484084, EPI_ISL_484085, EPI_ISL_484086, EPI_ISL_484087, EPI_ISL_484088, EPI_ISL_484089, EPI_ISL_484090, EPI_ISL_484091, EPI_ISL_484092, EPI_ISL_484093, EPI_ISL_484094, EPI_ISL_484095, EPI_ISL_484096, EPI_ISL_484097, EPI_ISL_484098, EPI_ISL_484099, EPI_ISL_484100, EPI_ISL_484101, EPI_ISL_484102, EPI_ISL_484103, EPI_ISL_484104, EPI_ISL_484105, EPI_ISL_484106, EPI_ISL_484107, EPI_ISL_484108, EPI_ISL_484109, EPI_ISL_484110, EPI_ISL_484111, EPI_ISL_484112, EPI_ISL_484113, EPI_ISL_484114, EPI_ISL_484115, EPI_ISL_484116, EPI_ISL_484117, EPI_ISL_484118, EPI_ISL_484119, EPI_ISL_484120, EPI_ISL_484121, EPI_ISL_484122, EPI_ISL_484123, EPI_ISL_484124, EPI_ISL_484125, EPI_ISL_484126, EPI_ISL_484127, EPI_ISL_484128, EPI_ISL_484129, EPI_ISL_484130, EPI_ISL_484131, EPI_ISL_484132, EPI_ISL_484133, EPI_ISL_484134, EPI_ISL_484135, EPI_ISL_484136, EPI_ISL_484137, EPI_ISL_484138, EPI_ISL_484139, EPI_ISL_484140, EPI_ISL_484141, EPI_ISL_484142, EPI_ISL_484143, EPI_ISL_484144, EPI_ISL_484145, EPI_ISL_484146, EPI_ISL_484147, EPI_ISL_484148, EPI_ISL_484149, EPI_ISL_484150, EPI_ISL_484151, EPI_ISL_484152, EPI_ISL_484153, EPI_ISL_484154, EPI_ISL_484155, EPI_ISL_484156, EPI_ISL_484157, EPI_ISL_484158, EPI_ISL_484159, EPI_ISL_484160, EPI_ISL_484161, EPI_ISL_484162, EPI_ISL_484163, EPI_ISL_484164, EPI_ISL_484165, EPI_ISL_484166, EPI_ISL_484167, EPI_ISL_484168, EPI_ISL_484169, EPI_ISL_484170, EPI_ISL_484171, EPI_ISL_484172, EPI_ISL_484173, EPI_ISL_484174, EPI_ISL_484175, EPI_ISL_484176, EPI_ISL_484177, EPI_ISL_484178, EPI_ISL_484179, EPI_ISL_484180, EPI_ISL_484181, EPI_ISL_484182, EPI_ISL_484183, EPI_ISL_484184, EPI_ISL_484185, EPI_ISL_484186, EPI_ISL_484187, EPI_ISL_484188, EPI_ISL_484189, EPI_ISL_484190, EPI_ISL_484191, EPI_ISL_484192, EPI_ISL_484193, EPI_ISL_484194, EPI_ISL_484195, EPI_ISL_484196, EPI_ISL_484197, EPI_ISL_484198, EPI_ISL_484199, EPI_ISL_484200, EPI_ISL_484201, EPI_ISL_484202, EPI_ISL_484203, EPI_ISL_484204, EPI_ISL_484205, EPI_ISL_484206, EPI_ISL_484207, EPI_ISL_484208, EPI_ISL_484209, EPI_ISL_484210, EPI_ISL_484211, EPI_ISL_484212, EPI_ISL_484213, EPI_ISL_484214, EPI_ISL_484215, EPI_ISL_484216, EPI_ISL_484217 | | | |
| see above | Centre for Clinical Infection and Diagnostics Research and Genomics Innovation Unit, Guy's and St. Thomas' NHS Trust | COVID-19 Genomics UK (COG-UK) Consortium | Chloe Fisher, Luke Snell, Penny Cliff, Rahul Batra, Jonathan Edgeworth, Ali Raza Awan |
| EPI_ISL_484218, EPI_ISL_484219, EPI_ISL_484220 | University of Birmingham | COVID-19 Genomics UK (COG-UK) Consortium | Institute of Microbiology, University of Birmingham: Claire McMurray, Joanne Stockton, Samuel Nicholls, Radoslaw Poplawski, Will Rowe, Josh Quick, Nicholas Loman. University of Birmingham Testing Laboratory: Celina M Whalley, Andrew Bosworth, Charlotte Poxon, Kasun Wanigasooriya, Oliver Pickles, Mike Kidd, Alex Richter, Andrew D Beggs PHE Heartlands Lab: Husam Osman, Andrew Bosworth. Queen Elizabeth Hospital: Anna Casey |
| EPI_ISL_484221, EPI_ISL_484222, EPI_ISL_484223, EPI_ISL_484224, EPI_ISL_484225, EPI_ISL_484226, EPI_ISL_484227, EPI_ISL_484228, EPI_ISL_484229, EPI_ISL_484230, EPI_ISL_484231, EPI_ISL_484232, EPI_ISL_484233, EPI_ISL_484234, EPI_ISL_484235, EPI_ISL_484236, EPI_ISL_484237, EPI_ISL_484238, EPI_ISL_484239, EPI_ISL_484240, EPI_ISL_484241, EPI_ISL_484242, EPI_ISL_484243, EPI_ISL_484244, EPI_ISL_484245, EPI_ISL_484246, EPI_ISL_484247, EPI_ISL_484248, EPI_ISL_484249, EPI_ISL_484250, EPI_ISL_484251, EPI_ISL_484252, EPI_ISL_484253 | | | |
| see above | University Hospitals Of Leicester NHS Trust and DeepSeq Nottingham | COVID-19 Genomics UK (COG-UK) Consortium | Christopher Holmes, Paul Bird, Thomas Helmer, Karlie Fallon, Julian Tang, Jonathan Ball, Patrick McClure, Joeseph Chappell, Nadine Holmes, Matthew Carlisle, Christopher Moore, Fei Sang, Johnny Debebe, Victoria Wright, Matthew Loose |
| EPI_ISL_484254, EPI_ISL_484255, EPI_ISL_484256, EPI_ISL_484257, EPI_ISL_484258, EPI_ISL_484259, EPI_ISL_484260, EPI_ISL_484261, EPI_ISL_484262, EPI_ISL_484263 | Liverpool Clinical Laboratories | COVID-19 Genomics UK (COG-UK) Consortium | Sam Haldenby, Anita Lucaci, Steve Paterson, Julian Hiscox, Alistair Darby, M Almsaud, A Alrezaihi, Muhannad Alruwaili, Stuart D Armstrong, Jones Benjamin, Eleanor G Bentley, Anu Chawla, Jordan J Clark, Angela Cowell, Richard Eccles, Isabel García-Dorival, Matthew Gemmell, Alessandro Gerada, PKF Gilmore, Richard Gregory, Ximeng Han, Catherine Hartley, Margaret Hughes, Miren Iturriza-Gomara, James Johnson, L Luu, Jenifer Manson, Charlotte Nelson, Elaine O'Toole, Cassie Olateju, Rebekah Penrice-Randal , Lucille Rainbow, N.P Randle, Trevor Ian Robinson, Parul Sharma, Ghada T Shawli, James P Stewart, Neil Swainston, Ecaterina Vamos, Joanne Watts, Mark Whitehead |
| EPI_ISL_484264, EPI_ISL_484265, EPI_ISL_484266, EPI_ISL_484267, EPI_ISL_484268, EPI_ISL_484269, EPI_ISL_484270, EPI_ISL_484271, EPI_ISL_484272, EPI_ISL_484273, EPI_ISL_484274, EPI_ISL_484275, EPI_ISL_484276, EPI_ISL_484277, EPI_ISL_484278, EPI_ISL_484279, EPI_ISL_484280, EPI_ISL_484281, EPI_ISL_484282, EPI_ISL_484283, EPI_ISL_484284, EPI_ISL_484285, EPI_ISL_484286, EPI_ISL_484287, EPI_ISL_484288, EPI_ISL_484289, EPI_ISL_484290, EPI_ISL_484291, EPI_ISL_484292, EPI_ISL_484293, EPI_ISL_484294, EPI_ISL_484295, EPI_ISL_484296, EPI_ISL_484297, EPI_ISL_484298, EPI_ISL_484299, EPI_ISL_484300, EPI_ISL_484301, EPI_ISL_484302, EPI_ISL_484303, EPI_ISL_484304, EPI_ISL_484305, EPI_ISL_484306, EPI_ISL_484307, EPI_ISL_484308, EPI_ISL_484309, EPI_ISL_484310, EPI_ISL_484311, EPI_ISL_484312, EPI_ISL_484313, EPI_ISL_484314, EPI_ISL_484315, EPI_ISL_484316, EPI_ISL_484317, EPI_ISL_484318, EPI_ISL_484319, EPI_ISL_484320, EPI_ISL_484321, EPI_ISL_484322, EPI_ISL_484323, EPI_ISL_484324, EPI_ISL_484325, EPI_ISL_484326, EPI_ISL_484327 | | | |
| see above | Northumbria University / South Tees Hospitals NHS Foundation Trust / North Cumbria Integrated Care NHS Foundation Trust / North Tees and Hartlepool NHS Foundation Trust / Newcastle Hospitals NHS Foundation Trust | COVID-19 Genomics UK (COG-UK) Consortium | Darren L Smith,Andrew Nelson,Matthew Bashton,Greg R Young,Joshua Loh,John Allan,Mohammad A Tariq,Giles S Holt,Gary Black,Wen C Yew,Lynn Dover,Paul Baker,Steve Liggett,Sarah Essex,Jane Greenaway,Debra Padgett,Clive Graham,Garren Scott,Edward Barton,Emma Swindells,Brendan Payne,Jennifer Collins,Yusri Taha,Gary Eltringham |
| EPI_ISL_484328, EPI_ISL_484329, EPI_ISL_484330, EPI_ISL_484331, EPI_ISL_484332, EPI_ISL_484333, EPI_ISL_484334, EPI_ISL_484335, EPI_ISL_484336, EPI_ISL_484337, EPI_ISL_484338 | | | |
| see above | Quadram Institute Bioscience | COVID-19 Genomics UK (COG-UK) Consortium | Dave J. Baker, Gemma L. Kay, Alp Aydin, Thanh Le-Viet, Steven Rudder, Ana P. Tedim, Anastasia Kolyva, Maria Diaz, Leonardo de Oliveira Martins, Nabil-Fareed Alikhan, Lizzie Meadows, Rachael Stanley, Ngozi Elumogo, Muhammed Yasir, Nicholas M. Thomson, Alexander J Trotter, Rachel Gilroy, Samuel Bloomfield, Claire Stuart, Andrew Bell, Reenesh Prakash, Samir Dervisevic, Alison E. Mather, John Wain, Mark Webber, Andrew J. Page, Justin O'Grady |
| EPI_ISL_484339, EPI_ISL_484340, EPI_ISL_484341, EPI_ISL_484342, EPI_ISL_484343, EPI_ISL_484344, EPI_ISL_484345, EPI_ISL_484346, EPI_ISL_484347, EPI_ISL_484348, EPI_ISL_484349, EPI_ISL_484350, EPI_ISL_484351, EPI_ISL_484352, EPI_ISL_484353 | | | |
| see above | Queens Medical Centre, Clinical Microbiology Department / DeepSeq Nottingham | COVID-19 Genomics UK (COG-UK) Consortium | Gemma Clark, Wendy Smith, Manjinder Khakh, Vicki M Fleming, Michelle M Lister, Hannah Howson-Wells, Jonathan Ball, Patrick McClure, Joseph Chappell, Theocharis Tsoleridis, Nadine Holmes, Matthew Carlisle, Christopher Moore, Fei Sang, Johnny Debebe, Victoria Wright, Matthew Loose |
| EPI_ISL_484354, EPI_ISL_484355, EPI_ISL_484356, EPI_ISL_484357, EPI_ISL_484358, EPI_ISL_484359, EPI_ISL_484360, EPI_ISL_484361, EPI_ISL_484362, EPI_ISL_484363, EPI_ISL_484364, EPI_ISL_484365, EPI_ISL_484366, EPI_ISL_484367, EPI_ISL_484368, EPI_ISL_484369, EPI_ISL_484370, EPI_ISL_484371, EPI_ISL_484372, EPI_ISL_484373 | | | |
| see above | Lincolnshire Hospitals and DeepSeq Nottingham | COVID-19 Genomics UK (COG-UK) Consortium | Nichola Duckworth, Tim Sloan, Sarah Walsh, Jonathan Ball, Patrick McClure, Joeseph Chappell, Nadine Holmes, Matthew Carlisle, Christopher Moore, Fei Sang, Johnny Debebe, Victoria Wright, Matthew Loose |
| EPI_ISL_484374, EPI_ISL_484375 | Queens Medical Centre, Clinical Microbiology Department / DeepSeq Nottingham | COVID-19 Genomics UK (COG-UK) Consortium | Gemma Clark, Wendy Smith, Manjinder Khakh, Vicki M Fleming, Michelle M Lister, Hannah Howson-Wells, Jonathan Ball, Patrick McClure, Joseph Chappell, Theocharis Tsoleridis, Nadine Holmes, Matthew Carlisle, Christopher Moore, Fei Sang, Johnny Debebe, Victoria Wright, Matthew Loose |
| EPI_ISL_484376, EPI_ISL_484377, EPI_ISL_484378, EPI_ISL_484379, EPI_ISL_484380, EPI_ISL_484381, EPI_ISL_484382, EPI_ISL_484383, EPI_ISL_484384, EPI_ISL_484385, EPI_ISL_484386, EPI_ISL_484387, EPI_ISL_484388, EPI_ISL_484389 | | | |
| see above | University Hospitals Of Leicester NHS Trust and DeepSeq Nottingham | COVID-19 Genomics UK (COG-UK) Consortium | Christopher Holmes, Paul Bird, Thomas Helmer, Karlie Fallon, Julian Tang, Jonathan Ball, Patrick McClure, Joeseph Chappell, Nadine Holmes, Matthew Carlisle, Christopher Moore, Fei Sang, Johnny Debebe, Victoria Wright, Matthew Loose |
| EPI_ISL_484390, EPI_ISL_484391 | Queens Medical Centre, Clinical Microbiology Department / DeepSeq Nottingham | COVID-19 Genomics UK (COG-UK) Consortium | Gemma Clark, Wendy Smith, Manjinder Khakh, Vicki M Fleming, Michelle M Lister, Hannah Howson-Wells, Jonathan Ball, Patrick McClure, Joseph Chappell, Theocharis Tsoleridis, Nadine Holmes, Matthew Carlisle, Christopher Moore, Fei Sang, Johnny Debebe, Victoria Wright, Matthew Loose |
| EPI_ISL_484392, EPI_ISL_484393, EPI_ISL_484394, EPI_ISL_484395, EPI_ISL_484396, EPI_ISL_484397, EPI_ISL_484398, EPI_ISL_484399, EPI_ISL_484400, EPI_ISL_484401, EPI_ISL_484402, EPI_ISL_484403, EPI_ISL_484404, EPI_ISL_484405, EPI_ISL_484406 | | | |
| see above | Lincolnshire Hospitals and DeepSeq Nottingham | COVID-19 Genomics UK (COG-UK) Consortium | Nichola Duckworth, Tim Sloan, Sarah Walsh, Jonathan Ball, Patrick McClure, Joeseph Chappell, Nadine Holmes, Matthew Carlisle, Christopher Moore, Fei Sang, Johnny Debebe, Victoria Wright, Matthew Loose |
| EPI_ISL_484407, EPI_ISL_484408, EPI_ISL_484409, EPI_ISL_484410, EPI_ISL_484411, EPI_ISL_484412, EPI_ISL_484413, EPI_ISL_484414, EPI_ISL_484415, EPI_ISL_484416, EPI_ISL_484417, EPI_ISL_484418, EPI_ISL_484419, EPI_ISL_484420, EPI_ISL_484421, EPI_ISL_484422, EPI_ISL_484423, EPI_ISL_484424, EPI_ISL_484425, EPI_ISL_484426, EPI_ISL_484427, EPI_ISL_484428, EPI_ISL_484429, EPI_ISL_484430, EPI_ISL_484431, EPI_ISL_484432 | | | |
| see above | Centre for Enzyme Innovation, University of Portsmouth / Translational Research Laboratory, Portsmouth Hospitals NHS Trust | COVID-19 Genomics UK (COG-UK) Consortium | Angela Beckett,Yann Bourgeois,Garry Scarlett,Sharon Glaysher,Scott Elliott,Kelly Bicknell,Robert Impey,Allyson Lloyd,Sarah Wyllie,Ethan Butcher,Anoop Chauhan,Samuel Robson |
| EPI_ISL_484433, EPI_ISL_484434, EPI_ISL_484435, EPI_ISL_484436, EPI_ISL_484437, EPI_ISL_484438, EPI_ISL_484439, EPI_ISL_484440, EPI_ISL_484441, EPI_ISL_484442, EPI_ISL_484443, EPI_ISL_484444, EPI_ISL_484445, EPI_ISL_484446, EPI_ISL_484447, EPI_ISL_484448, EPI_ISL_484449, EPI_ISL_484450, EPI_ISL_484451, EPI_ISL_484452, EPI_ISL_484453, EPI_ISL_484454, EPI_ISL_484455, EPI_ISL_484456, EPI_ISL_484457, EPI_ISL_484458, EPI_ISL_484459, EPI_ISL_484460, EPI_ISL_484461, EPI_ISL_484462, EPI_ISL_484463, EPI_ISL_484464, EPI_ISL_484465, EPI_ISL_484466, EPI_ISL_484467, EPI_ISL_484468, EPI_ISL_484469, EPI_ISL_484470, EPI_ISL_484471, EPI_ISL_484472, EPI_ISL_484473, EPI_ISL_484474, EPI_ISL_484475, EPI_ISL_484476, EPI_ISL_484477, EPI_ISL_484478, EPI_ISL_484479, EPI_ISL_484480, EPI_ISL_484481, EPI_ISL_484482, EPI_ISL_484483, EPI_ISL_484484, EPI_ISL_484485, EPI_ISL_484486, EPI_ISL_484487, EPI_ISL_484488, EPI_ISL_484489, EPI_ISL_484490, EPI_ISL_484491, EPI_ISL_484492, EPI_ISL_484493, EPI_ISL_484494, EPI_ISL_484495, EPI_ISL_484496, EPI_ISL_484497, EPI_ISL_484498, EPI_ISL_484499, EPI_ISL_484500, EPI_ISL_484501, EPI_ISL_484502, EPI_ISL_484503, EPI_ISL_484504, EPI_ISL_484505, EPI_ISL_484506, EPI_ISL_484507, EPI_ISL_484508, EPI_ISL_484509, EPI_ISL_484510, EPI_ISL_484511, EPI_ISL_484512, EPI_ISL_484513, EPI_ISL_484514, EPI_ISL_484515, EPI_ISL_484516, EPI_ISL_484517, EPI_ISL_484518 | | | |
| see above | Virology Department, Sheffield Teaching Hospitals NHS Foundation Trust/Department of Infection, Immunity and Cardiovascular Disease, The Medical School, University of Sheffield | COVID-19 Genomics UK (COG-UK) Consortium | Thushan de Silva, Matthew Parker, Nikki Smith, Adri Angyal, Rebecca Brown, Luke Green, Rachel Tucker, Paul Parsons, Danielle Groves, Katie Johnson, Laura Carrilero, Alex Keeley, Dave Partridge, Matthew Wyles, Benjamin Lindsey, Mehmet Yavuz, Mohammad Raza, Cariad Evans |
| EPI_ISL_484519, EPI_ISL_484520, EPI_ISL_484521, EPI_ISL_484522, EPI_ISL_484523, EPI_ISL_484524, EPI_ISL_484525, EPI_ISL_484526, EPI_ISL_484527, EPI_ISL_484528, EPI_ISL_484529, EPI_ISL_484530, EPI_ISL_484531, EPI_ISL_484532, EPI_ISL_484533, EPI_ISL_484534, EPI_ISL_484535, EPI_ISL_484536, EPI_ISL_484537, EPI_ISL_484538, EPI_ISL_484539, EPI_ISL_484540, EPI_ISL_484541, EPI_ISL_484542, EPI_ISL_484543, EPI_ISL_484544, EPI_ISL_484545, EPI_ISL_484546, EPI_ISL_484547, EPI_ISL_484548, EPI_ISL_484549, EPI_ISL_484550, EPI_ISL_484551, EPI_ISL_484552, EPI_ISL_484553, EPI_ISL_484554, EPI_ISL_484555, EPI_ISL_484556, EPI_ISL_484557, EPI_ISL_484558, EPI_ISL_484559, EPI_ISL_484560, EPI_ISL_484561, EPI_ISL_484562, EPI_ISL_484563, EPI_ISL_484564, EPI_ISL_484565, EPI_ISL_484566, EPI_ISL_484567, EPI_ISL_484568, EPI_ISL_484569, EPI_ISL_484570, EPI_ISL_484571, EPI_ISL_484572, EPI_ISL_484573, EPI_ISL_484574, EPI_ISL_484575, EPI_ISL_484576, EPI_ISL_484577, EPI_ISL_484578, EPI_ISL_484579, EPI_ISL_484580, EPI_ISL_484581, EPI_ISL_484582, EPI_ISL_484583, EPI_ISL_484584, EPI_ISL_484585, EPI_ISL_484586, EPI_ISL_484587, EPI_ISL_484588, EPI_ISL_484589, EPI_ISL_484590, EPI_ISL_484591, EPI_ISL_484592, EPI_ISL_484593, EPI_ISL_484594, EPI_ISL_484595, EPI_ISL_484596, EPI_ISL_484597, EPI_ISL_484598, EPI_ISL_484599, EPI_ISL_484600, EPI_ISL_484601, EPI_ISL_484602, EPI_ISL_484603, EPI_ISL_484604, EPI_ISL_484605, EPI_ISL_484606, EPI_ISL_484607, EPI_ISL_484608, EPI_ISL_484609, EPI_ISL_484610, EPI_ISL_484611, EPI_ISL_484612, EPI_ISL_484613, EPI_ISL_484614, EPI_ISL_484615, EPI_ISL_484616, EPI_ISL_484617, EPI_ISL_484618, EPI_ISL_484619, EPI_ISL_484620, EPI_ISL_484621, EPI_ISL_484622, EPI_ISL_484623, EPI_ISL_484624, EPI_ISL_484625, EPI_ISL_484626, EPI_ISL_484627, EPI_ISL_484628, EPI_ISL_484629, EPI_ISL_484630, EPI_ISL_484631, EPI_ISL_484632, EPI_ISL_484633, EPI_ISL_484634, EPI_ISL_484635, EPI_ISL_484636, EPI_ISL_484637, EPI_ISL_484638, EPI_ISL_484639, EPI_ISL_484640, EPI_ISL_484641, EPI_ISL_484642, EPI_ISL_484643, EPI_ISL_484644, EPI_ISL_484645, EPI_ISL_484646, EPI_ISL_484647, EPI_ISL_484648, EPI_ISL_484649, EPI_ISL_484650, EPI_ISL_484651, EPI_ISL_484652, EPI_ISL_484653, EPI_ISL_484654, EPI_ISL_484655, EPI_ISL_484656, EPI_ISL_484657, EPI_ISL_484658, EPI_ISL_484659, EPI_ISL_484660, EPI_ISL_484661, EPI_ISL_484662, EPI_ISL_484663, EPI_ISL_484664, EPI_ISL_484665, EPI_ISL_484666, EPI_ISL_484667, EPI_ISL_484668, EPI_ISL_484669, EPI_ISL_484670, EPI_ISL_484671, EPI_ISL_484672, EPI_ISL_484673, EPI_ISL_484674, EPI_ISL_484675, EPI_ISL_484676, EPI_ISL_484677, EPI_ISL_484678, EPI_ISL_484679 | | | |
| see above | West of Scotland Specialist Virology Centre, NHSGGC / MRC-University of Glasgow Centre for Virus Research | COVID-19 Genomics UK (COG-UK) Consortium | Ana da Silva Filipe, Natasha Johnson, Kathy Smollett, Daniel Mair, Stephen Carmichael, Lily Tong, Jenna Nichols, Elihu Aranday-Cortes, Kirstyn Brunker, Yasmin Parr, Alice Broos, Kyriaki Nomikou; Sarah McDonald, Marc Niebel, Patawee Asamaphan; Richard Orton, Joseph Hughes, Sreenu Vattipally, David L Robertson; Alasdair MacLean, Rory Gunson; Kathy Li, Natasha Jesudason, Rajiv Shah, James Shepherd, Antonia Ho, Emma Thomson |
| EPI_ISL_484680, EPI_ISL_484681, EPI_ISL_484682, EPI_ISL_484683, EPI_ISL_484684 | Virology Department, Royal Infirmary of Edinburgh, NHS Lothian / School of Biological Sciences, University of Edinburgh / Institute of Genetics and Molecular Medicine, University of Edinburgh | COVID-19 Genomics UK (COG-UK) Consortium | McHugh M, Dewar R, Rooke S, Gallagher M, Balcaza C, O'Toole Á, Scher E, Hill V, McCrone JT, Colquhoun R, Yu X, Jackson B, Rambaut A, Williams TC, Templeton K |
| EPI_ISL_484685, EPI_ISL_484686, EPI_ISL_484687, EPI_ISL_484688, EPI_ISL_484689, EPI_ISL_484690, EPI_ISL_484691 | Originating lab: Wales Specialist Virology Centre Sequencing lab: Pathogen Genomics Unit | COVID-19 Genomics UK (COG-UK) Consortium | Catherine Moore, Johnathan Evans, Laura Gifford, Malorie Perry, Simon Cottrell, Angela Marchbank, Alec Birchley, Alexander Adams, Amy Gaskin, Bree Gatica-Wilcox, Jason Coombes, Joel Southgate, Lauren Gilbert, Lee Graham, Nicole Pacchiarini, Sara Kumziene-Summerhayes, Sarah Taylor, Sophie Jones, Sara Rey, Matthew Bull, Joanne Watkins, Sally Corden, Tom Connor |
| EPI_ISL_484692 | Minnesota Department of Health, Public Health Laboratory | Minnesota Department of Health, Public Health Laboratory | Matt Plumb, Jacob Garfin, and Xiong Wang |
| EPI_ISL_484693, EPI_ISL_484694, EPI_ISL_484695, EPI_ISL_484696, EPI_ISL_484697, EPI_ISL_484698, EPI_ISL_484699, EPI_ISL_484700, EPI_ISL_484701, EPI_ISL_484702, EPI_ISL_484703, EPI_ISL_484704, EPI_ISL_484705, EPI_ISL_484706, EPI_ISL_484707, EPI_ISL_484708 | | | |
| see above | Department of Clinical Microbiology | GIGA Medical Genomics | Keith Durkin, Maria Artesi, Sébastien Bontems, Raphaël Boreux, Cécile Meex, Axelle Chaslain, Céline Fombellida-Lopez, Pierrette Melin, Marie-Pierre Hayette, Vincent Bours. |
| EPI_ISL_484709, EPI_ISL_484710, EPI_ISL_484711, EPI_ISL_484712, EPI_ISL_484713, EPI_ISL_484714, EPI_ISL_484715, EPI_ISL_484716, EPI_ISL_484717, EPI_ISL_484718, EPI_ISL_484719, EPI_ISL_484720, EPI_ISL_484721, EPI_ISL_484722, EPI_ISL_484723, EPI_ISL_484724, EPI_ISL_484725, EPI_ISL_484726, EPI_ISL_484727, EPI_ISL_484728, EPI_ISL_484729, EPI_ISL_484730, EPI_ISL_484731, EPI_ISL_484732, | | | |

| | | | |
|---|---|---|---|
| EPI_ISL_484733, EPI_ISL_484734, EPI_ISL_484735, EPI_ISL_484736, EPI_ISL_484737, EPI_ISL_484738, EPI_ISL_484739, EPI_ISL_484740, EPI_ISL_484741, EPI_ISL_484742, EPI_ISL_484743, EPI_ISL_484744, EPI_ISL_484745, EPI_ISL_484746, EPI_ISL_484747, EPI_ISL_484748, EPI_ISL_484749, EPI_ISL_484750, EPI_ISL_484751, EPI_ISL_484752, EPI_ISL_484753, EPI_ISL_484754, EPI_ISL_484755, EPI_ISL_484756, EPI_ISL_484757, EPI_ISL_484758, EPI_ISL_484759, EPI_ISL_484760, EPI_ISL_484761, EPI_ISL_484762, EPI_ISL_484763, EPI_ISL_484764, EPI_ISL_484765, EPI_ISL_484766, EPI_ISL_484767, EPI_ISL_484768, EPI_ISL_484769, EPI_ISL_484770, EPI_ISL_484771, EPI_ISL_484772, EPI_ISL_484773, EPI_ISL_484774, EPI_ISL_484775, EPI_ISL_484776, EPI_ISL_484777, EPI_ISL_484778, EPI_ISL_484779, EPI_ISL_484780, EPI_ISL_484781, EPI_ISL_484782, EPI_ISL_484783, EPI_ISL_484784, EPI_ISL_484785, EPI_ISL_484786, EPI_ISL_484787, EPI_ISL_484788, EPI_ISL_484789, EPI_ISL_484790, EPI_ISL_484791, EPI_ISL_484792, EPI_ISL_484793, EPI_ISL_484794, EPI_ISL_484795, EPI_ISL_484796, EPI_ISL_484797, EPI_ISL_484798, EPI_ISL_484799, EPI_ISL_484800, EPI_ISL_484801, EPI_ISL_484802, EPI_ISL_484803, EPI_ISL_484804, EPI_ISL_484805, EPI_ISL_484806 | | | |
| see above | University of Michigan Clinical Microbiology Laboratory | Lauring Lab, University of Michigan, Department of Microbiology and Immunology | Valesano et al. |
| EPI_ISL_484807, EPI_ISL_484808, EPI_ISL_484809, EPI_ISL_484810, EPI_ISL_484811, EPI_ISL_484812, EPI_ISL_484813, EPI_ISL_484814, EPI_ISL_484815, EPI_ISL_484816, EPI_ISL_484817, EPI_ISL_484818, EPI_ISL_484819, EPI_ISL_484820, EPI_ISL_484821, EPI_ISL_484822, EPI_ISL_484823, EPI_ISL_484824, EPI_ISL_484825, EPI_ISL_484826, EPI_ISL_484827, EPI_ISL_484828, EPI_ISL_484829, EPI_ISL_484830, EPI_ISL_484832, EPI_ISL_484833, EPI_ISL_484834, EPI_ISL_484835, EPI_ISL_484836, EPI_ISL_484837, EPI_ISL_484838, EPI_ISL_484839, EPI_ISL_484840, EPI_ISL_484841, EPI_ISL_484842, EPI_ISL_484843, EPI_ISL_484844, EPI_ISL_484845, EPI_ISL_484846, EPI_ISL_484847, EPI_ISL_484849, EPI_ISL_484850, EPI_ISL_484851, EPI_ISL_484852, EPI_ISL_484853, EPI_ISL_484854, EPI_ISL_484855, EPI_ISL_484856, EPI_ISL_484857, EPI_ISL_484858, EPI_ISL_484859, EPI_ISL_484860, EPI_ISL_484861, EPI_ISL_484862, EPI_ISL_484863, EPI_ISL_484864, EPI_ISL_484865, EPI_ISL_484866, EPI_ISL_484867, EPI_ISL_484868, EPI_ISL_484869, EPI_ISL_484870, EPI_ISL_484871, EPI_ISL_484872, EPI_ISL_484873, EPI_ISL_484874, EPI_ISL_484875, EPI_ISL_484876, EPI_ISL_484877, EPI_ISL_484878, EPI_ISL_484879, EPI_ISL_484880, EPI_ISL_484881, EPI_ISL_484882, EPI_ISL_484883, EPI_ISL_484884, EPI_ISL_484885, EPI_ISL_484886, EPI_ISL_484887, EPI_ISL_484888, EPI_ISL_484889, EPI_ISL_484890, EPI_ISL_484891, EPI_ISL_484892, EPI_ISL_484893, EPI_ISL_484894, EPI_ISL_484895, EPI_ISL_484896, EPI_ISL_484897, EPI_ISL_484898, EPI_ISL_484899, EPI_ISL_484900, EPI_ISL_484901, EPI_ISL_484902, EPI_ISL_484903, EPI_ISL_484904, EPI_ISL_484905, EPI_ISL_484906, EPI_ISL_484907, EPI_ISL_484908, EPI_ISL_484909, EPI_ISL_484910, EPI_ISL_484911, EPI_ISL_484912, EPI_ISL_484913, EPI_ISL_484914, EPI_ISL_484915, EPI_ISL_484916, EPI_ISL_484917, EPI_ISL_484918, EPI_ISL_484919, EPI_ISL_484920, EPI_ISL_484921, EPI_ISL_484922, EPI_ISL_484923, EPI_ISL_484924, EPI_ISL_484925, EPI_ISL_484926, EPI_ISL_484927, EPI_ISL_484928, EPI_ISL_484929, EPI_ISL_484930, EPI_ISL_484931, EPI_ISL_484932, EPI_ISL_484933, EPI_ISL_484934, EPI_ISL_484935, EPI_ISL_484936, EPI_ISL_484937, EPI_ISL_484938, EPI_ISL_484939, EPI_ISL_484940, EPI_ISL_484941, EPI_ISL_484942, EPI_ISL_484943, EPI_ISL_484944, EPI_ISL_484945, EPI_ISL_484946, EPI_ISL_484947, EPI_ISL_484948, EPI_ISL_484949, EPI_ISL_484950, EPI_ISL_484951, EPI_ISL_484952, EPI_ISL_484953, EPI_ISL_484954, EPI_ISL_484955, EPI_ISL_484956, EPI_ISL_484957, EPI_ISL_484958, EPI_ISL_484959, EPI_ISL_484960, EPI_ISL_484961, EPI_ISL_484962, EPI_ISL_484963, EPI_ISL_484964, EPI_ISL_484965, EPI_ISL_484966, EPI_ISL_484967, EPI_ISL_484969, EPI_ISL_484970, EPI_ISL_484971, EPI_ISL_484972, EPI_ISL_484973, EPI_ISL_484974, EPI_ISL_484975, EPI_ISL_484976, EPI_ISL_484978, EPI_ISL_484979, EPI_ISL_484981, EPI_ISL_484982, EPI_ISL_484983, EPI_ISL_484984, EPI_ISL_484985, EPI_ISL_484986, EPI_ISL_484987, EPI_ISL_484988, EPI_ISL_484989, EPI_ISL_484990, EPI_ISL_484991, EPI_ISL_484992, EPI_ISL_484993, EPI_ISL_484994, EPI_ISL_484995, EPI_ISL_484996, EPI_ISL_484997, EPI_ISL_484998, EPI_ISL_484999, EPI_ISL_485000 | | | |
| see above | University of Wisconsin-Madison AIDS Vaccine Research Laboratories | University of Wisconsin-Madison AIDS Vaccine Research Laboratories | Gage Moreno, Katarina Braun, et al. AIDS Vaccine Research Laboratories |
| EPI_ISL_485001, EPI_ISL_485002 | University of Ulsan College of Medicine and Asan Medical Center | University of Ulsan College of Medicine and Asan Medical Center | Kuenyoul Park, Jaewoong Lee, Kihyun Lee, Jiwon Jung, Sung-Han Kim, Jina Lee, Mauricio Chalita, Seok-Hwan Yoon, Jongsik Chun, Kyu-Hwa Hur, Heungsup Sung, Mi-Na Kim, and Hae Kyung Lee |
| EPI_ISL_485003, EPI_ISL_485004, EPI_ISL_485005, EPI_ISL_485006, EPI_ISL_485007, EPI_ISL_485008, EPI_ISL_485009, EPI_ISL_485010, EPI_ISL_485011, EPI_ISL_485012, EPI_ISL_485013, EPI_ISL_485014, EPI_ISL_485015, EPI_ISL_485016, EPI_ISL_485017, EPI_ISL_485018, EPI_ISL_485019, EPI_ISL_485020, EPI_ISL_485021, EPI_ISL_485022, EPI_ISL_485023, EPI_ISL_485024, EPI_ISL_485025, EPI_ISL_485026, EPI_ISL_485027, EPI_ISL_485028, EPI_ISL_485029, EPI_ISL_485030, EPI_ISL_485031, EPI_ISL_485032, EPI_ISL_485033, EPI_ISL_485034, EPI_ISL_485035, EPI_ISL_485036, EPI_ISL_485037, EPI_ISL_485038, EPI_ISL_485039, EPI_ISL_485040, EPI_ISL_485041, EPI_ISL_485042, EPI_ISL_485043, EPI_ISL_485044, EPI_ISL_485045, EPI_ISL_485046, EPI_ISL_485047, EPI_ISL_485048, EPI_ISL_485049, EPI_ISL_485050, EPI_ISL_485051, EPI_ISL_485052, EPI_ISL_485053, EPI_ISL_485054, EPI_ISL_485055, EPI_ISL_485056, EPI_ISL_485057, EPI_ISL_485058, EPI_ISL_485059, EPI_ISL_485060, EPI_ISL_485061, EPI_ISL_485062, EPI_ISL_485063, EPI_ISL_485064, EPI_ISL_485065, EPI_ISL_485066, EPI_ISL_485067, EPI_ISL_485068, EPI_ISL_485069, EPI_ISL_485070, EPI_ISL_485071, EPI_ISL_485072, EPI_ISL_485073, EPI_ISL_485074, EPI_ISL_485075, EPI_ISL_485076, EPI_ISL_485077, EPI_ISL_485078, EPI_ISL_485079, EPI_ISL_485080, EPI_ISL_485081, EPI_ISL_485082, EPI_ISL_485083, EPI_ISL_485084, EPI_ISL_485085, EPI_ISL_485086, EPI_ISL_485087, EPI_ISL_485088, EPI_ISL_485089, EPI_ISL_485090, EPI_ISL_485091, EPI_ISL_485092, EPI_ISL_485093, EPI_ISL_485094, EPI_ISL_485095, EPI_ISL_485096, EPI_ISL_485097, EPI_ISL_485098, EPI_ISL_485099, EPI_ISL_485100, EPI_ISL_485101, EPI_ISL_485102, EPI_ISL_485103, EPI_ISL_485104, EPI_ISL_485105, EPI_ISL_485106, EPI_ISL_485107, EPI_ISL_485108, EPI_ISL_485109, EPI_ISL_485110, EPI_ISL_485111, EPI_ISL_485112, EPI_ISL_485113, EPI_ISL_485114, EPI_ISL_485115, EPI_ISL_485116, EPI_ISL_485117, EPI_ISL_485118, EPI_ISL_485119, EPI_ISL_485120, EPI_ISL_485121, EPI_ISL_485122, EPI_ISL_485123, EPI_ISL_485124, EPI_ISL_485125, EPI_ISL_485126, EPI_ISL_485127, EPI_ISL_485128, EPI_ISL_485129, EPI_ISL_485130, EPI_ISL_485131, EPI_ISL_485132, EPI_ISL_485133, EPI_ISL_485134, EPI_ISL_485135, EPI_ISL_485136, EPI_ISL_485137, EPI_ISL_485138, EPI_ISL_485139, EPI_ISL_485140, EPI_ISL_485141, EPI_ISL_485142, EPI_ISL_485143, EPI_ISL_485144, EPI_ISL_485145, EPI_ISL_485146, EPI_ISL_485147, EPI_ISL_485148, EPI_ISL_485149, EPI_ISL_485150, EPI_ISL_485151, EPI_ISL_485152, EPI_ISL_485153, EPI_ISL_485154, EPI_ISL_485155, EPI_ISL_485156, EPI_ISL_485157, EPI_ISL_485158, EPI_ISL_485159, EPI_ISL_485160, EPI_ISL_485161, EPI_ISL_485162, EPI_ISL_485163, EPI_ISL_485164, EPI_ISL_485165, EPI_ISL_485166, EPI_ISL_485167, EPI_ISL_485168, EPI_ISL_485169, EPI_ISL_485170, EPI_ISL_485171, EPI_ISL_485172, EPI_ISL_485173, EPI_ISL_485174, EPI_ISL_485175, EPI_ISL_485176, EPI_ISL_485177, EPI_ISL_485178, EPI_ISL_485179, EPI_ISL_485180, EPI_ISL_485181, EPI_ISL_485182, EPI_ISL_485183, EPI_ISL_485184, EPI_ISL_485185, EPI_ISL_485186, EPI_ISL_485187, EPI_ISL_485188, EPI_ISL_485189, EPI_ISL_485190, EPI_ISL_485191, EPI_ISL_485192, EPI_ISL_485193, EPI_ISL_485194, EPI_ISL_485195, EPI_ISL_485196, EPI_ISL_485197, EPI_ISL_485198, EPI_ISL_485199, EPI_ISL_485200, EPI_ISL_485201, EPI_ISL_485202, EPI_ISL_485203, EPI_ISL_485204, EPI_ISL_485205, EPI_ISL_485206, EPI_ISL_485207, EPI_ISL_485208, EPI_ISL_485209, EPI_ISL_485210, EPI_ISL_485211, EPI_ISL_485212, EPI_ISL_485213, EPI_ISL_485214, EPI_ISL_485215, EPI_ISL_485216, EPI_ISL_485217, EPI_ISL_485218, EPI_ISL_485219, EPI_ISL_485220, EPI_ISL_485221, EPI_ISL_485222, EPI_ISL_485223, EPI_ISL_485224, EPI_ISL_485225, EPI_ISL_485226, EPI_ISL_485227, EPI_ISL_485228, EPI_ISL_485229, EPI_ISL_485230, EPI_ISL_485231, EPI_ISL_485232, EPI_ISL_485233, EPI_ISL_485234, EPI_ISL_485235, EPI_ISL_485236, EPI_ISL_485237, EPI_ISL_485238, EPI_ISL_485239, EPI_ISL_485240, EPI_ISL_485241, EPI_ISL_485242, EPI_ISL_485243, EPI_ISL_485244, EPI_ISL_485245, EPI_ISL_485246, EPI_ISL_485247, EPI_ISL_485248, EPI_ISL_485249, EPI_ISL_485250, EPI_ISL_485251, EPI_ISL_485252, EPI_ISL_485253, EPI_ISL_485254, EPI_ISL_485255, EPI_ISL_485256, EPI_ISL_485257, EPI_ISL_485258, EPI_ISL_485259, EPI_ISL_485260, EPI_ISL_485261, EPI_ISL_485262, EPI_ISL_485263, EPI_ISL_485264, EPI_ISL_485265, EPI_ISL_485266, EPI_ISL_485267, EPI_ISL_485268, EPI_ISL_485269, EPI_ISL_485270, EPI_ISL_485271, EPI_ISL_485272, EPI_ISL_485273, EPI_ISL_485274, EPI_ISL_485275, EPI_ISL_485276, EPI_ISL_485277, EPI_ISL_485278, EPI_ISL_485279, EPI_ISL_485280, EPI_ISL_485281, EPI_ISL_485282, EPI_ISL_485283, EPI_ISL_485284, EPI_ISL_485285, EPI_ISL_485286, EPI_ISL_485287, EPI_ISL_485288, EPI_ISL_485289, EPI_ISL_485290, EPI_ISL_485291, EPI_ISL_485292, EPI_ISL_485293, EPI_ISL_485294, EPI_ISL_485295, EPI_ISL_485296, EPI_ISL_485297, EPI_ISL_485298, EPI_ISL_485299, EPI_ISL_485300, EPI_ISL_485301, EPI_ISL_485302, EPI_ISL_485303, EPI_ISL_485304, EPI_ISL_485305, EPI_ISL_485306, EPI_ISL_485307, EPI_ISL_485308, EPI_ISL_485309, EPI_ISL_485310, EPI_ISL_485311, EPI_ISL_485312, EPI_ISL_485313, EPI_ISL_485314, EPI_ISL_485315, EPI_ISL_485316, EPI_ISL_485317, EPI_ISL_485318, EPI_ISL_485319, EPI_ISL_485320, EPI_ISL_485321, EPI_ISL_485322, EPI_ISL_485323, EPI_ISL_485324, EPI_ISL_485325, EPI_ISL_485326, EPI_ISL_485327, EPI_ISL_485328, EPI_ISL_485329, EPI_ISL_485330, EPI_ISL_485331, EPI_ISL_485332, EPI_ISL_485333, EPI_ISL_485334, EPI_ISL_485335, EPI_ISL_485336, EPI_ISL_485337, EPI_ISL_485338, EPI_ISL_485339, EPI_ISL_485340, EPI_ISL_485341, EPI_ISL_485342, EPI_ISL_485343, EPI_ISL_485344, EPI_ISL_485345, EPI_ISL_485346, EPI_ISL_485347, EPI_ISL_485348, EPI_ISL_485349, EPI_ISL_485350, EPI_ISL_485351, EPI_ISL_485352, EPI_ISL_485353, EPI_ISL_485354, EPI_ISL_485355, EPI_ISL_485356, EPI_ISL_485357, EPI_ISL_485358, EPI_ISL_485359, EPI_ISL_485360, EPI_ISL_485361, EPI_ISL_485362, EPI_ISL_485363, EPI_ISL_485364, EPI_ISL_485365, EPI_ISL_485366, EPI_ISL_485367, EPI_ISL_485368, EPI_ISL_485369, EPI_ISL_485370, EPI_ISL_485371, EPI_ISL_485372, EPI_ISL_485373, EPI_ISL_485374, EPI_ISL_485375, EPI_ISL_485376, EPI_ISL_485377, EPI_ISL_485378, EPI_ISL_485379, EPI_ISL_485380, EPI_ISL_485381, EPI_ISL_485382, EPI_ISL_485383, EPI_ISL_485384, EPI_ISL_485385, EPI_ISL_485386, EPI_ISL_485387 | | | |
| see above | River Road Testing Lab | Ginkgo Bioworks Clinical Laboratory | Rebecca C. Christofferson, Stephania A. Cormier, Luan V. Dinh, E. Handly Mayton, Hollis R. O'Neil, Thaya Stoufflet, Malaika Mckenzie-Bennett, James McGann, Jim Griffin, Keith Robison, Alex Plocik, Becky Schilling, Rebecca Littlefield, Michelle Spencer, Birgitte Simen |
| EPI_ISL_485388, EPI_ISL_485389, EPI_ISL_485390, EPI_ISL_485391, EPI_ISL_485392, EPI_ISL_485393, EPI_ISL_485394, EPI_ISL_485395, EPI_ISL_485396 | University of Ulsan College of Medicine and Asan Medical Center | University of Ulsan College of Medicine and Asan Medical Center | Kuenyoul Park, Jaewoong Lee, Kihyun Lee, Jiwon Jung, Sung-Han Kim, Jina Lee, Mauricio Chalita, Seok-Hwan Yoon, Jongsik Chun, Kyu-Hwa Hur, Heungsup Sung, Mi-Na Kim, and Hae Kyung Lee |
| EPI_ISL_485398 | unknown | Department of Internal Medicine | Kim,D.-M. |
| EPI_ISL_485399 | Institute of Human Genetics, Polish Academy of Sciences | Institute of Human Genetics, Polish Academy of Sciences | Szymon Hryhorowicz, Adam Ustaszewski, Marta Kaczmarek-Ryś, Emilia Lis, Ewa Ziętkiewicz, Michał Witt, Andrzej Pławski |
| EPI_ISL_485400 | Institute of Human Genetics, Polish Academy of Sciences | Institute of Human Genetics, Polish Academy of Sciences, | Szymon Hryhorowicz, Adam Ustaszewski, Marta Kaczmarek-Ryś, Emilia Lis, Ewa Ziętkiewicz, Michał Witt, Andrzej Pławski |
| EPI_ISL_485401 | unknown | Communicable Disease Laboratory | Zaed,A., Al-Wasti,H., Al-Taif,Z. and Shehab,F. |
| EPI_ISL_485575, EPI_ISL_485576, EPI_ISL_485577, EPI_ISL_485578, EPI_ISL_485579, EPI_ISL_485580, EPI_ISL_485581, EPI_ISL_485582, EPI_ISL_485583, EPI_ISL_485584, EPI_ISL_485585, EPI_ISL_485586, EPI_ISL_485587, EPI_ISL_485588, EPI_ISL_485589, EPI_ISL_485590, EPI_ISL_485591, EPI_ISL_485592, EPI_ISL_485593, EPI_ISL_485594, EPI_ISL_485595, EPI_ISL_485596, EPI_ISL_485597, EPI_ISL_485598, EPI_ISL_485599, EPI_ISL_485600, EPI_ISL_485601, EPI_ISL_485602 | | | |
| see above | Instituto de diagnóstico y Referencia Epidemiologicos (INDRE) | Instituto de diagnóstico y Referencia Epidemiologicos (INDRE) | Barrera-Badillo,G., Ramirez-Gonzalez,E. |
| EPI_ISL_485603 | Division of Infectious Disease | Steininger Lab | Jakob Thannesberger, Ingeborg Klymiuk, Nicolas Rascovan, Lorenz Schubert, Oliver Robak, Christoph Steininger |
| EPI_ISL_485604, EPI_ISL_485605, EPI_ISL_485606, EPI_ISL_485607, EPI_ISL_485608, EPI_ISL_485609, EPI_ISL_485610, EPI_ISL_485611 | Respiratory Virus Unit, Microbiology Services Colindale, Public Health England | Respiratory Virus Unit, Microbiology Services Colindale, Public Health England | PHE Covid Sequencing Team |
| EPI_ISL_485635, EPI_ISL_485708, EPI_ISL_485710, EPI_ISL_485711 | Institut Pasteur Dakar | Institut Pasteur de Dakar | Ndongo Dia, Moussa Moise Diagne, Mamadou diop, Marie Henriette Dior Ndione, Mamadou Malado Jallow, Safietou Sanke, Ousmane Faye, Amadou Alpha Sall. |
| EPI_ISL_485712 | Institut Pasteur | Institut Pasteur de Dakar | Ndongo Dia, Moussa Moise Diagne, Mamadou diop, Marie Henriette Dior Ndione, Mamadou Malado Jallow, Safietou Sanke, Ousmane Faye, Amadou Alpha Sall. |
| EPI_ISL_485713, EPI_ISL_485715, EPI_ISL_485716, EPI_ISL_485717 | Institut Pasteur Dakar | Institut Pasteur de Dakar | Ndongo Dia, Moussa Moise Diagne, Mamadou diop, Marie Henriette Dior Ndione, Mamadou Malado Jallow, Safietou Sanke, Ousmane Faye, Amadou Alpha Sall. |
| EPI_ISL_485809 | Institut für Virologie und Epidemiologie der Viruskrankheiten, Universitätsklinikum Tübingen | NGS Competence Center Tübingen, Institut für Medizinische Mikrobiologie und Hygiene, Universitätsklinikum Tübingen | Angelov at al. |
| EPI_ISL_485810, EPI_ISL_485811, EPI_ISL_485812, EPI_ISL_485813 | Institut für Virologie und Epidemiologie der Viruskrankheiten, Universitätsklinikum Tübingen | NGS Competence Center Tübingen, Institut für Medizinische Mikrobiologie und Hygiene, Universitätsklinikum Tübingen | Angelov et al. |
| EPI_ISL_485814, EPI_ISL_485815, EPI_ISL_485816, EPI_ISL_485817, EPI_ISL_485818, EPI_ISL_485819, EPI_ISL_485820, EPI_ISL_485821, EPI_ISL_485822, EPI_ISL_485823, EPI_ISL_485824, EPI_ISL_485825, EPI_ISL_485826, EPI_ISL_485827, EPI_ISL_485828, EPI_ISL_485829, EPI_ISL_485830, EPI_ISL_485831, EPI_ISL_485832, EPI_ISL_485833, EPI_ISL_485834, EPI_ISL_485835, EPI_ISL_485836, EPI_ISL_485837, EPI_ISL_485838, EPI_ISL_485839, EPI_ISL_485840, EPI_ISL_485841, EPI_ISL_485842, EPI_ISL_485843, EPI_ISL_485844, EPI_ISL_485845, EPI_ISL_485846, EPI_ISL_485847, EPI_ISL_485848, EPI_ISL_485849, EPI_ISL_485850, EPI_ISL_485851, EPI_ISL_485852, EPI_ISL_485853, EPI_ISL_485854, EPI_ISL_485855, EPI_ISL_485856, EPI_ISL_485857, EPI_ISL_485858, EPI_ISL_485859, EPI_ISL_485860, EPI_ISL_485861, EPI_ISL_485862, EPI_ISL_485863, EPI_ISL_485864, EPI_ISL_485865, EPI_ISL_485866, EPI_ISL_485867, EPI_ISL_485868, EPI_ISL_485869, EPI_ISL_485870, EPI_ISL_485871 | | | |
| see above | Virginia DCLS | Virginia DCLS | Virginia DCLS |
| EPI_ISL_485872 | New Mexico Department of Health Scientific Laboratory Division | Center for Global Health, University of New Mexico Health Sciences Center | Daryl Domman, Kurt Schwalm, Twila Kunde, Joseph Hicks, Michael Edwards, Darrell Dinwiddie |
| EPI_ISL_485873, EPI_ISL_485874, EPI_ISL_485875, EPI_ISL_485876, EPI_ISL_485877, EPI_ISL_485878, EPI_ISL_485879, EPI_ISL_485880, EPI_ISL_485881, EPI_ISL_485882, EPI_ISL_485883, EPI_ISL_485884, EPI_ISL_485885, EPI_ISL_485886, EPI_ISL_485887, EPI_ISL_485888, EPI_ISL_485889, EPI_ISL_485890, EPI_ISL_485891, EPI_ISL_485892, EPI_ISL_485893, EPI_ISL_485894, EPI_ISL_485895, EPI_ISL_485896, EPI_ISL_485897, EPI_ISL_485898, EPI_ISL_485899, EPI_ISL_485900, EPI_ISL_485901, EPI_ISL_485902, EPI_ISL_485903, EPI_ISL_485904, EPI_ISL_485905, EPI_ISL_485906 | | | |
| see above | River Road Testing Lab | Ginkgo Bioworks Clinical Laboratory | Rebecca C. Christofferson, Stephania A. Cormier, Luan V. Dinh, E. Handly Mayton, Hollis R. O'Neil, Thaya Stoufflet, Malaika Mckenzie-Bennett, James McGann, Jim Griffin, Keith Robison, Alex Plocik, Becky Schilling, Rebecca Littlefield, Michelle Spencer, Birgitte Simen |
| EPI_ISL_485917, EPI_ISL_485918, EPI_ISL_485919, EPI_ISL_485920, EPI_ISL_485921, EPI_ISL_485922, EPI_ISL_485923, EPI_ISL_485924, EPI_ISL_485925, EPI_ISL_485926, EPI_ISL_485927, EPI_ISL_485928, EPI_ISL_485929, EPI_ISL_485930, EPI_ISL_485931, EPI_ISL_485932, EPI_ISL_485933, EPI_ISL_485934, EPI_ISL_485935, EPI_ISL_485936, EPI_ISL_485937, EPI_ISL_485938, EPI_ISL_485939, EPI_ISL_485940, EPI_ISL_485941, EPI_ISL_485942, EPI_ISL_485944, EPI_ISL_485945, EPI_ISL_485946, EPI_ISL_485947, EPI_ISL_485948, EPI_ISL_485949, EPI_ISL_485950, EPI_ISL_485951, EPI_ISL_485952, EPI_ISL_485953, EPI_ISL_485954, EPI_ISL_485955, EPI_ISL_485956, EPI_ISL_485957, EPI_ISL_485958, EPI_ISL_485959, EPI_ISL_485960, EPI_ISL_485961, EPI_ISL_485963, EPI_ISL_485964, EPI_ISL_485967, EPI_ISL_485968, EPI_ISL_485969, EPI_ISL_485971, EPI_ISL_485972, EPI_ISL_485973, EPI_ISL_485974, EPI_ISL_485975, EPI_ISL_485976, EPI_ISL_485977, EPI_ISL_485978, EPI_ISL_485979, EPI_ISL_485980, EPI_ISL_485981, EPI_ISL_485982, EPI_ISL_485983, EPI_ISL_485984, EPI_ISL_485985, EPI_ISL_485986, EPI_ISL_485987, EPI_ISL_485988, EPI_ISL_485989, EPI_ISL_485990, EPI_ISL_485991, EPI_ISL_485992, EPI_ISL_485993, EPI_ISL_485994, EPI_ISL_485995, EPI_ISL_485996, EPI_ISL_485997, EPI_ISL_485998, EPI_ISL_485999, EPI_ISL_486000, EPI_ISL_486001, EPI_ISL_486002, EPI_ISL_486003, EPI_ISL_486004, EPI_ISL_486005, EPI_ISL_486006, EPI_ISL_486007, EPI_ISL_486008, EPI_ISL_486009, EPI_ISL_486010, EPI_ISL_486011, EPI_ISL_486012, EPI_ISL_486013, EPI_ISL_486014, EPI_ISL_486015, EPI_ISL_486016, EPI_ISL_486017, EPI_ISL_486018, EPI_ISL_486019, EPI_ISL_486020, EPI_ISL_486021, EPI_ISL_486022, EPI_ISL_486023, EPI_ISL_486024, EPI_ISL_486025, EPI_ISL_486026, EPI_ISL_486027, EPI_ISL_486028, EPI_ISL_486029, EPI_ISL_486030, EPI_ISL_486032, EPI_ISL_486033, EPI_ISL_486034, EPI_ISL_486035, EPI_ISL_486036, EPI_ISL_486037, EPI_ISL_486038, EPI_ISL_486039, EPI_ISL_486040, EPI_ISL_486041, EPI_ISL_486042, EPI_ISL_486043, EPI_ISL_486044, EPI_ISL_486045, EPI_ISL_486046, EPI_ISL_486047, EPI_ISL_486048, EPI_ISL_486049, EPI_ISL_486050, EPI_ISL_486051, EPI_ISL_486052, EPI_ISL_486053, EPI_ISL_486054, EPI_ISL_486055, EPI_ISL_486056, EPI_ISL_486057, EPI_ISL_486058, EPI_ISL_486059, EPI_ISL_486060, EPI_ISL_486061, EPI_ISL_486062, EPI_ISL_486063, EPI_ISL_486064, EPI_ISL_486065, EPI_ISL_486066, EPI_ISL_486067, EPI_ISL_486068, EPI_ISL_486069, EPI_ISL_486070, EPI_ISL_486071, EPI_ISL_486072, EPI_ISL_486073, EPI_ISL_486074, EPI_ISL_486075, EPI_ISL_486076, EPI_ISL_486077, EPI_ISL_486078, EPI_ISL_486079, EPI_ISL_486080, EPI_ISL_486081, EPI_ISL_486082, EPI_ISL_486083, EPI_ISL_486084, EPI_ISL_486085, EPI_ISL_486086, EPI_ISL_486087, EPI_ISL_486088, EPI_ISL_486089, EPI_ISL_486090, EPI_ISL_486093, EPI_ISL_486094, EPI_ISL_486095, EPI_ISL_486097, EPI_ISL_486098, EPI_ISL_486099, EPI_ISL_486102, EPI_ISL_486103, EPI_ISL_486105, EPI_ISL_486106, EPI_ISL_486108, EPI_ISL_486109, EPI_ISL_486110, EPI_ISL_486112, EPI_ISL_486113, EPI_ISL_486114 | | | |
| see above | UW Virology Lab | UW Virology Lab | Pavitra Roychoudhury, Hong Xie, Lasata Shrestha, Amin Addetia, Truong Nguyen, Victoria M Rachleff, Meei-Li Huang, Keith R Jerome, Alexander Greninger |
| EPI_ISL_486115, EPI_ISL_486116, EPI_ISL_486117, EPI_ISL_486118, EPI_ISL_486119 | County of Santa Clara Public Health Department | Chan-Zuckerberg Biohub | CZB Cliahub Consortium |
| EPI_ISL_486120, EPI_ISL_486121, EPI_ISL_486122, EPI_ISL_486123, EPI_ISL_486124, EPI_ISL_486125, EPI_ISL_486126, EPI_ISL_486127, EPI_ISL_486128, EPI_ISL_486129, EPI_ISL_486130, EPI_ISL_486131, EPI_ISL_486132, EPI_ISL_486133, EPI_ISL_486134, EPI_ISL_486135, EPI_ISL_486136, EPI_ISL_486137, EPI_ISL_486138, EPI_ISL_486139, EPI_ISL_486140, EPI_ISL_486141, EPI_ISL_486142, EPI_ISL_486143, EPI_ISL_486144, EPI_ISL_486145, EPI_ISL_486146, EPI_ISL_486147, EPI_ISL_486148, EPI_ISL_486149, EPI_ISL_486150, EPI_ISL_486151, EPI_ISL_486152, EPI_ISL_486153, EPI_ISL_486154, EPI_ISL_486155, EPI_ISL_486156, EPI_ISL_486157, EPI_ISL_486158, EPI_ISL_486159, EPI_ISL_486160, EPI_ISL_486161, EPI_ISL_486162, EPI_ISL_486163, EPI_ISL_486164, EPI_ISL_486165, EPI_ISL_486166, EPI_ISL_486167, EPI_ISL_486168, EPI_ISL_486169, EPI_ISL_486170, EPI_ISL_486171, EPI_ISL_486172, EPI_ISL_486173, EPI_ISL_486174, EPI_ISL_486175, EPI_ISL_486176, EPI_ISL_486177, EPI_ISL_486178, EPI_ISL_486179, EPI_ISL_486180, EPI_ISL_486181, EPI_ISL_486182, EPI_ISL_486183, EPI_ISL_486184, EPI_ISL_486185, EPI_ISL_486186, EPI_ISL_486187, EPI_ISL_486188, EPI_ISL_486189, EPI_ISL_486190, EPI_ISL_486191, EPI_ISL_486192, EPI_ISL_486193, EPI_ISL_486194, EPI_ISL_486195, EPI_ISL_486196, EPI_ISL_486197, EPI_ISL_486198, EPI_ISL_486199, EPI_ISL_486200, EPI_ISL_486201, EPI_ISL_486202, EPI_ISL_486203, EPI_ISL_486204, EPI_ISL_486205, EPI_ISL_486206, EPI_ISL_486207, EPI_ISL_486208, EPI_ISL_486209, EPI_ISL_486210, EPI_ISL_486211, EPI_ISL_486212, EPI_ISL_486213, EPI_ISL_486214, EPI_ISL_486215, EPI_ISL_486216, EPI_ISL_486217, EPI_ISL_486218, EPI_ISL_486219, EPI_ISL_486220, EPI_ISL_486221, EPI_ISL_486222, EPI_ISL_486223, EPI_ISL_486224, EPI_ISL_486225, EPI_ISL_486226, EPI_ISL_486227, EPI_ISL_486228, EPI_ISL_486229, EPI_ISL_486230, EPI_ISL_486231, EPI_ISL_486232, EPI_ISL_486233, EPI_ISL_486234, EPI_ISL_486235, EPI_ISL_486236, EPI_ISL_486237, EPI_ISL_486238, EPI_ISL_486239, EPI_ISL_486240, EPI_ISL_486241, EPI_ISL_486242, EPI_ISL_486243, EPI_ISL_486244, EPI_ISL_486245, EPI_ISL_486246, EPI_ISL_486247, EPI_ISL_486248, EPI_ISL_486249, EPI_ISL_486250, EPI_ISL_486251, EPI_ISL_486252, EPI_ISL_486253, EPI_ISL_486254, EPI_ISL_486255, EPI_ISL_486256, EPI_ISL_486257, EPI_ISL_486258, EPI_ISL_486259, EPI_ISL_486260, EPI_ISL_486261, EPI_ISL_486262, EPI_ISL_486263, | | | |

| | | | |
|---|---|---|---|
| EPI_ISL_486264, EPI_ISL_486265, EPI_ISL_486266, EPI_ISL_486267, EPI_ISL_486268, EPI_ISL_486269, EPI_ISL_486270, EPI_ISL_486271, EPI_ISL_486272, EPI_ISL_486273, EPI_ISL_486274, EPI_ISL_486275, EPI_ISL_486276, EPI_ISL_486277, EPI_ISL_486278, EPI_ISL_486279 | | | |
| see above | Orange County Public Health Laboratory | Chan-Zuckerberg Biohub | CZB Cliahub Consortium |
| EPI_ISL_486280, EPI_ISL_486281, EPI_ISL_486282, EPI_ISL_486283, EPI_ISL_486284, EPI_ISL_486285, EPI_ISL_486286 | Humboldt County Public Health Laboratory | Chan-Zuckerberg Biohub | CZB Cliahub Consortium |
| EPI_ISL_486287, EPI_ISL_486288, EPI_ISL_486289, EPI_ISL_486290, EPI_ISL_486291, EPI_ISL_486292, EPI_ISL_486293, EPI_ISL_486294, EPI_ISL_486295, EPI_ISL_486296, EPI_ISL_486297, EPI_ISL_486298, EPI_ISL_486299, EPI_ISL_486300, EPI_ISL_486301, EPI_ISL_486302, EPI_ISL_486303, EPI_ISL_486304, EPI_ISL_486305, EPI_ISL_486306, EPI_ISL_486307, EPI_ISL_486308, EPI_ISL_486309, EPI_ISL_486310, EPI_ISL_486311, EPI_ISL_486312, EPI_ISL_486313, EPI_ISL_486314, EPI_ISL_486315, EPI_ISL_486316, EPI_ISL_486317, EPI_ISL_486318, EPI_ISL_486319, EPI_ISL_486320, EPI_ISL_486321, EPI_ISL_486322, EPI_ISL_486323, EPI_ISL_486324, EPI_ISL_486325, EPI_ISL_486326, EPI_ISL_486327, EPI_ISL_486328, EPI_ISL_486329, EPI_ISL_486330, EPI_ISL_486331, EPI_ISL_486332, EPI_ISL_486333, EPI_ISL_486334, EPI_ISL_486335, EPI_ISL_486336, EPI_ISL_486337, EPI_ISL_486338 | | | |
| see above | San Joaquin County Public Health Lab | Chan-Zuckerberg Biohub | CZB Cliahub Consortium |
| EPI_ISL_486339, EPI_ISL_486340, EPI_ISL_486341, EPI_ISL_486342, EPI_ISL_486343, EPI_ISL_486344, EPI_ISL_486345, EPI_ISL_486346, EPI_ISL_486347, EPI_ISL_486348, EPI_ISL_486349, EPI_ISL_486350, EPI_ISL_486351, EPI_ISL_486352, EPI_ISL_486353, EPI_ISL_486354, EPI_ISL_486355, EPI_ISL_486356, EPI_ISL_486357, EPI_ISL_486358, EPI_ISL_486359, EPI_ISL_486360, EPI_ISL_486361, EPI_ISL_486362, EPI_ISL_486363, EPI_ISL_486364, EPI_ISL_486365 | | | |
| see above | UCSF Clinical Microbiology Laboratory | Chan-Zuckerberg Biohub | CZB Cliahub Consortium |
| EPI_ISL_486382 | District Surveillence Unit | Department of Neurovirology, National Institute of Mental Health and Neuroscience (NIMHANS) | Chitra Pattabiraman, Vijayalakshmi Reddy, Harsha PK, Risha Rasheed, Shafeeq S Hameed, Manjunatha Venkataswamy, Anita Desai, Ravi Vasanthapuram |
| EPI_ISL_486383 | CV Raman Hospital | Department of Neurovirology, National Institute of Mental Health and Neuroscience (NIMHANS) | Chitra Pattabiraman, Vijayalakshmi Reddy, Harsha PK, Risha Rasheed, Shafeeq S Hameed, Manjunatha Venkataswamy, Anita Desai, Ravi Vasanthapuram |
| EPI_ISL_486384 | DH | Department of Neurovirology, National Institute of Mental Health and Neuroscience (NIMHANS) | Chitra Pattabiraman, Vijayalakshmi Reddy, Harsha PK, Risha Rasheed, Shafeeq S Hameed, Manjunatha Venkataswamy, Anita Desai, Ravi Vasanthapuram |
| EPI_ISL_486385, EPI_ISL_486386 | Victoria Hospital | Department of Neurovirology, National Institute of Mental Health and Neuroscience (NIMHANS) | Chitra Pattabiraman, Vijayalakshmi Reddy, Harsha PK, Risha Rasheed, Shafeeq S Hameed, Manjunatha Venkataswamy, Anita Desai, Ravi Vasanthapuram |
| EPI_ISL_486387, EPI_ISL_486388, EPI_ISL_486389 | DH | Department of Neurovirology, National Institute of Mental Health and Neuroscience (NIMHANS) | Chitra Pattabiraman, Vijayalakshmi Reddy, Harsha PK, Risha Rasheed, Shafeeq S Hameed, Manjunatha Venkataswamy, Anita Desai, Ravi Vasanthapuram |
| EPI_ISL_486390, EPI_ISL_486391 | Centrālā laboratorija | Latvian Biomedical Research and Study Centre | Ivars Silamiķelis, Kaspars Megnis, Monta Ustinova, Ņikita Zrelovs, Vita Rovīte, Stella Lapiņa, Jana Osīte, Marta Priedīte, Uga Dumpis, Jānis Kloviņš |
| EPI_ISL_486392 | Victoria Hospital | Department of Neurovirology, National Institute of Mental Health and Neuroscience (NIMHANS) | Chitra Pattabiraman, Vijayalakshmi Reddy, Harsha PK, Risha Rasheed, Shafeeq S Hameed, Manjunatha Venkataswamy, Anita Desai, Ravi Vasanthapuram |
| EPI_ISL_486393 | SJMCH | Department of Neurovirology, National Institute of Mental Health and Neuroscience (NIMHANS) | Chitra Pattabiraman, Vijayalakshmi Reddy, Harsha PK, Risha Rasheed, Shafeeq S Hameed, Manjunatha Venkataswamy, Anita Desai, Ravi Vasanthapuram |
| EPI_ISL_486394 | MIMS | Department of Neurovirology, National Institute of Mental Health and Neuroscience (NIMHANS) | Chitra Pattabiraman, Vijayalakshmi Reddy, Harsha PK, Risha Rasheed, Shafeeq S Hameed, Manjunatha Venkataswamy, Anita Desai, Ravi Vasanthapuram |
| EPI_ISL_486395 | BIMS | Department of Neurovirology, National Institute of Mental Health and Neuroscience (NIMHANS) | Chitra Pattabiraman, Vijayalakshmi Reddy, Harsha PK, Risha Rasheed, Shafeeq S Hameed, Manjunatha Venkataswamy, Anita Desai, Ravi Vasanthapuram |
| EPI_ISL_486396 | Jayanagar General Hospital to Victoria Hospital | Department of Neurovirology, National Institute of Mental Health and Neuroscience (NIMHANS) | Chitra Pattabiraman, Vijayalakshmi Reddy, Harsha PK, Risha Rasheed, Shafeeq S Hameed, Manjunatha Venkataswamy, Anita Desai, Ravi Vasanthapuram |
| EPI_ISL_486397 | KC General Hospital | Department of Neurovirology, National Institute of Mental Health and Neuroscience (NIMHANS) | Chitra Pattabiraman, Vijayalakshmi Reddy, Harsha PK, Risha Rasheed, Shafeeq S Hameed, Manjunatha Venkataswamy, Anita Desai, Ravi Vasanthapuram |
| EPI_ISL_486398, EPI_ISL_486399 | MIMS | Department of Neurovirology, National Institute of Mental Health and Neuroscience (NIMHANS) | Chitra Pattabiraman, Vijayalakshmi Reddy, Harsha PK, Risha Rasheed, Shafeeq S Hameed, Manjunatha Venkataswamy, Anita Desai, Ravi Vasanthapuram |
| EPI_ISL_486400 | Victoria Hospital | Department of Neurovirology, National Institute of Mental Health and Neuroscience (NIMHANS) | Chitra Pattabiraman, Vijayalakshmi Reddy, Harsha PK, Risha Rasheed, Shafeeq S Hameed, Manjunatha Venkataswamy, Anita Desai, Ravi Vasanthapuram |
| EPI_ISL_486401, EPI_ISL_486402, EPI_ISL_486403 | DH | Department of Neurovirology, National Institute of Mental Health and Neuroscience (NIMHANS) | Chitra Pattabiraman, Vijayalakshmi Reddy, Harsha PK, Risha Rasheed, Shafeeq S Hameed, Manjunatha Venkataswamy, Anita Desai, Ravi Vasanthapuram |
| EPI_ISL_486404 | Victoria Hospital | Department of Neurovirology, National Institute of Mental Health and Neuroscience (NIMHANS) | Chitra Pattabiraman, Vijayalakshmi Reddy, Harsha PK, Risha Rasheed, Shafeeq S Hameed, Manjunatha Venkataswamy, Anita Desai, Ravi Vasanthapuram |
| EPI_ISL_486405, EPI_ISL_486406, EPI_ISL_486407, EPI_ISL_486408, EPI_ISL_486409 | DH | Department of Neurovirology, National Institute of Mental Health and Neuroscience (NIMHANS) | Chitra Pattabiraman, Vijayalakshmi Reddy, Harsha PK, Risha Rasheed, Shafeeq S Hameed, Manjunatha Venkataswamy, Anita Desai, Ravi Vasanthapuram |
| EPI_ISL_486410, EPI_ISL_486411 | Centrālā laboratorija | Latvian Biomedical Research and Study Centre | Ivars Silamiķelis, Kaspars Megnis, Monta Ustinova, Ņikita Zrelovs, Vita Rovīte, Stella Lapiņa, Jana Osīte, Marta Priedīte, Uga Dumpis, Jānis Kloviņš |
| EPI_ISL_486412 | Centrālā laboratorija | Latvian Biomedical Research and Study Centre | Ivars Silamiķelis, Kaspars Megnis, Monta Ustinova, Ņikita Zrelovs, Vita Rovīte, Stella Lapiņa, Jana Osīte, Marta Priedīte, Uga Dumpis, Jānis Kloviņš |
| EPI_ISL_486413, EPI_ISL_486414 | Centrālā laboratorija | Latvian Biomedical Research and Study Centre | Ivars Silamiķelis, Kaspars Megnis, Monta Ustinova, Ņikita Zrelovs, Vita Rovīte, Stella Lapiņa, Jana Osīte, Marta Priedīte, Uga Dumpis, Jānis Kloviņš |
| EPI_ISL_486415 | Centrālā laboratorija | Latvian Biomedical Research and Study Centre | Ivars Silamiķelis, Kaspars Megnis, Monta Ustinova, Ņikita Zrelovs, Vita Rovīte, Stella Lapiņa, Jana Osīte, Marta Priedīte, Uga Dumpis, Jānis Kloviņš |
| EPI_ISL_486416 | Centrālā laboratorija | Latvian Biomedical Research and Study Centre | Ivars Silamiķelis, Kaspars Megnis, Monta Ustinova, Ņikita Zrelovs, Vita Rovīte, Stella Lapiņa, Jana Osīte, Marta Priedīte, Uga Dumpis, Jānis Kloviņš |
| EPI_ISL_486417, EPI_ISL_486418 | Centrālā laboratorija | Latvian Biomedical Research and Study Centre | Ivars Silamiķelis, Kaspars Megnis, Monta Ustinova, Ņikita Zrelovs, Vita Rovīte, Stella Lapiņa, Jana Osīte, Marta Priedīte, Uga Dumpis, Jānis Kloviņš |
| EPI_ISL_486419 | Centrālā laboratorija | Latvian Biomedical Research and Study Centre | Ivars Silamiķelis, Kaspars Megnis, Monta Ustinova, Ņikita Zrelovs, Vita Rovīte, Stella Lapiņa, Jana Osīte, Marta Priedīte, Uga Dumpis, Jānis Kloviņš |
| EPI_ISL_486420, EPI_ISL_486421 | Centrālā laboratorija | Latvian Biomedical Research and Study Centre | Ivars Silamiķelis, Kaspars Megnis, Monta Ustinova, Ņikita Zrelovs, Vita Rovīte, Stella Lapiņa, Jana Osīte, Marta Priedīte, Uga Dumpis, Jānis Kloviņš |
| EPI_ISL_486422, EPI_ISL_486423, EPI_ISL_486424, EPI_ISL_486425, EPI_ISL_486426 | Latvijas Infektoloģijas centrs | Latvian Biomedical Research and Study Centre | Ivars Silamiķelis, Kaspars Megnis, Monta Ustinova, Ņikita Zrelovs, Vita Rovīte, Jeļena Storoženko, Tatjana Kolupajeva, Oksana Savicka, Uga Dumpis, Jānis Kloviņš |
| EPI_ISL_486427 | unknown | Clinical Laboratory, Hospital Israelita Albert Einstein | Amgarteb,D., Malta,F., Guedes,R.L., Santana,R.A., de Menezes,F.G., Mangueira,C.L. and Pinho,J.R. |
| EPI_ISL_486428 | Latvijas Infektoloģijas centrs | Latvian Biomedical Research and Study Centre | Ivars Silamiķelis, Kaspars Megnis, Monta Ustinova, Ņikita Zrelovs, Vita Rovīte, Jeļena Storoženko, Tatjana Kolupajeva, Oksana Savicka, Uga Dumpis, Jānis Kloviņš |
| EPI_ISL_486429 | unknown | Clinical Laboratory, Hospital Israelita Albert Einstein | Malta,F., Amgarten,D., Guedes,R.L., Santana,R.A., de Menezes,F.G., Mangueira,C.L. and Pinho,J.R. |
| EPI_ISL_486430, EPI_ISL_486431, EPI_ISL_486432, EPI_ISL_486433, EPI_ISL_486434, EPI_ISL_486435 | Latvijas Infektoloģijas centrs | Latvian Biomedical Research and Study Centre | Ivars Silamiķelis, Kaspars Megnis, Monta Ustinova, Ņikita Zrelovs, Vita Rovīte, Jeļena Storoženko, Tatjana Kolupajeva, Oksana Savicka, Uga Dumpis, Jānis Kloviņš |
| EPI_ISL_486436 | Latvijas Infektoloģijas centrs | Latvian Biomedical Research and Study Centre | Ivars Silamiķelis, Kaspars Megnis, Monta Ustinova, Ņikita Zrelovs, Vita Rovīte, Jeļena Storoženko, Tatjana Kolupajeva, Oksana Savicka, Uga Dumpis, Jānis Kloviņš |

| | | | |
|---|---|---|---|
| EPI_ISL_486437 | Centrālā laboratorija | Latvian Biomedical Research and Study Centre | Ivars Silamiķelis, Kaspars Megnis, Monta Ustinova, Ņikita Zrelovs, Vita Rovīte, Stella Lapiņa, Jana Osīte, Marta Priedīte, Uga Dumpis, Jānis Kloviņš |
| EPI_ISL_486438 | E. Gulbja laboratorija | Latvian Biomedical Research and Study Centre | Ivars Silamiķelis, Kaspars Megnis, Monta Ustinova, Ņikita Zrelovs, Vita Rovīte, Mikus Gavars, Dmitrijs Perminovs, Uga Dumpis, Jānis Kloviņš |
| EPI_ISL_486442, EPI_ISL_486443, EPI_ISL_486444, EPI_ISL_486445, EPI_ISL_486446, EPI_ISL_486447, EPI_ISL_486448, EPI_ISL_486449, EPI_ISL_486450, EPI_ISL_486451, EPI_ISL_486452, EPI_ISL_486453, EPI_ISL_486454, EPI_ISL_486455, EPI_ISL_486456, EPI_ISL_486457, EPI_ISL_486458, EPI_ISL_486459, EPI_ISL_486460, EPI_ISL_486461, EPI_ISL_486462, EPI_ISL_486463, EPI_ISL_486464, EPI_ISL_486465, EPI_ISL_486466, EPI_ISL_486467, EPI_ISL_486468, EPI_ISL_486469, EPI_ISL_486470, EPI_ISL_486471, EPI_ISL_486472, EPI_ISL_486473, EPI_ISL_486474, EPI_ISL_486475, EPI_ISL_486476, EPI_ISL_486477, EPI_ISL_486478, EPI_ISL_486479, EPI_ISL_486480, EPI_ISL_486481, EPI_ISL_486482, EPI_ISL_486483, EPI_ISL_486484, EPI_ISL_486485, EPI_ISL_486486, EPI_ISL_486487, EPI_ISL_486488, EPI_ISL_486489, EPI_ISL_486490, EPI_ISL_486491, EPI_ISL_486492, EPI_ISL_486493, EPI_ISL_486494, EPI_ISL_486495, EPI_ISL_486496, EPI_ISL_486497, EPI_ISL_486498, EPI_ISL_486499, EPI_ISL_486500, EPI_ISL_486501, EPI_ISL_486502, EPI_ISL_486503, EPI_ISL_486504, EPI_ISL_486505, EPI_ISL_486506, EPI_ISL_486507, EPI_ISL_486508, EPI_ISL_486509, EPI_ISL_486510, EPI_ISL_486511, EPI_ISL_486512, EPI_ISL_486513, EPI_ISL_486514, EPI_ISL_486515, EPI_ISL_486516, EPI_ISL_486517, EPI_ISL_486518, EPI_ISL_486519, EPI_ISL_486520, EPI_ISL_486521, EPI_ISL_486522, EPI_ISL_486523, EPI_ISL_486524, EPI_ISL_486525, EPI_ISL_486526, EPI_ISL_486527, EPI_ISL_486528, EPI_ISL_486529, EPI_ISL_486530, EPI_ISL_486531, EPI_ISL_486532, EPI_ISL_486533, EPI_ISL_486534, EPI_ISL_486535, EPI_ISL_486536, EPI_ISL_486537, EPI_ISL_486538 | | | |
| see above | Viollier AG | Department of Biosystems Science and Engineering, ETH Zürich | Christian Beisel, Sarah Nadeau, Ivan Topolsky, Pedro Ferreira, Philipp Jablonski, Susana Posada-Céspedes, Tobias Schär, Ina Nissen, Natascha Santacroce, Elodie Burcklen, Christiane Beckmann, Maurice Redondo, Olivier Kobel, Christoph Noppen, Sophie Seidel, Noemie Santamaria de Souza, Niko Beerenwinkel, Tanja Stadler |
| EPI_ISL_486646, EPI_ISL_486647 | Microbiology, Virology and Biemergency Laboratory-ASST FBF Sacco | Microbiology, Virology and Biemergency Laboratory-ASST FBF Sacco | Mancon A, Comandatore F, Romeri F, Micheli V, Rimoldi SG |
| EPI_ISL_486648 | Microbiology, Virology and Biemergency Laboratory-ASST FBF Sacco | Microbiology, Virology and Biemergency Laboratory-ASST FBF Sacco | Micheli V, Comandatore F, Romeri F, Mancon A, Rimoldi SG |
| EPI_ISL_486649 | Microbiology, Virology and Biemergency Laboratory-ASST FBF Sacco | Microbiology, Virology and Biemergency Laboratory-ASST FBF Sacco | Rimoldi SG, Comandatore F, Romeri F, Mancon A, Micheli V |
| EPI_ISL_486650 | Microbiology, Virology and Biemergency Laboratory-ASST FBF Sacco | Microbiology, Virology and Biemergency Laboratory-ASST FBF Sacco | Romeri F, Comandatore F, Mancon A, Micheli V, Rimoldi SG |
| EPI_ISL_486651 | Microbiology, Virology and Biemergency Laboratory-ASST FBF Sacco | Microbiology, Virology and Biemergency Laboratory-ASST FBF Sacco | Mancon A, Comandatore F, Romeri F, Micheli V, Rimoldi SG |
| EPI_ISL_486652 | Microbiology, Virology and Biemergency Laboratory-ASST FBF Sacco | Microbiology, Virology and Biemergency Laboratory-ASST FBF Sacco | Micheli V, Comandatore F, Romeri F, Mancon A, Rimoldi SG |
| EPI_ISL_486653 | Microbiology, Virology and Biemergency Laboratory-ASST FBF Sacco | Microbiology, Virology and Biemergency Laboratory-ASST FBF Sacco | Rimoldi SG, Comandatore F, Romeri F, Mancon A, Micheli V |
| EPI_ISL_486654 | Microbiology, Virology and Biemergency Laboratory-ASST FBF Sacco | Microbiology, Virology and Biemergency Laboratory-ASST FBF Sacco | Romeri F, Comandatore F, Mancon A, Micheli V, Rimoldi SG |
| EPI_ISL_486655 | Microbiology, Virology and Biemergency Laboratory-ASST FBF Sacco | Microbiology, Virology and Biemergency Laboratory-ASST FBF Sacco | Mancon A, Comandatore F, Romeri F, Micheli V, Rimoldi SG |
| EPI_ISL_486656 | Microbiology, Virology and Biemergency Laboratory-ASST FBF Sacco | Microbiology, Virology and Biemergency Laboratory-ASST FBF Sacco | Micheli V, Comandatore F, Romeri F, Mancon A, Rimoldi SG |
| EPI_ISL_486657 | Microbiology, Virology and Biemergency Laboratory-ASST FBF Sacco | Microbiology, Virology and Biemergency Laboratory-ASST FBF Sacco | Rimoldi SG, Comandatore F, Romeri F, Mancon A, Micheli V |
| EPI_ISL_486658 | Microbiology, Virology and Biemergency Laboratory-ASST FBF Sacco | Microbiology, Virology and Biemergency Laboratory-ASST FBF Sacco | Romeri F, Comandatore F, Mancon A, Micheli V, Rimoldi SG |
| EPI_ISL_486659 | Microbiology, Virology and Biemergency Laboratory-ASST FBF Sacco | Microbiology, Virology and Biemergency Laboratory-ASST FBF Sacco | Micheli V, Comandatore F, Romeri F, Mancon A, Rimoldi SG |
| EPI_ISL_486660 | Microbiology, Virology and Biemergency Laboratory-ASST FBF Sacco | Microbiology, Virology and Biemergency Laboratory-ASST FBF Sacco | Rimoldi SG, Comandatore F, Romeri F, Mancon A, Micheli V |
| EPI_ISL_486661 | Microbiology, Virology and Biemergency Laboratory-ASST FBF Sacco | Microbiology, Virology and Biemergency Laboratory-ASST FBF Sacco | Romeri F, Comandatore F, Mancon A, Micheli V, Rimoldi SG |
| EPI_ISL_486662 | Microbiology, Virology and Biemergency Laboratory-ASST FBF Sacco | Microbiology, Virology and Biemergency Laboratory-ASST FBF Sacco | Mancon A, Comandatore F, Romeri F, Micheli V, Rimoldi SG |
| EPI_ISL_486663 | Microbiology, Virology and Biemergency Laboratory-ASST FBF Sacco | Microbiology, Virology and Biemergency Laboratory-ASST FBF Sacco | Micheli V, Comandatore F, Romeri F, Mancon A, Rimoldi SG |
| EPI_ISL_486664 | Microbiology, Virology and Biemergency Laboratory-ASST FBF Sacco | Microbiology, Virology and Biemergency Laboratory-ASST FBF Sacco | Rimoldi SG, Comandatore F, Romeri F, Mancon A, Micheli V |
| EPI_ISL_486665 | Microbiology, Virology and Biemergency Laboratory-ASST FBF Sacco | Microbiology, Virology and Biemergency Laboratory-ASST FBF Sacco | Micheli V, Rimoldi SG, Comandatore F, Mancon A, Romeri F |
| EPI_ISL_486666, EPI_ISL_486667, EPI_ISL_486668, EPI_ISL_486669, EPI_ISL_486670, EPI_ISL_486671, EPI_ISL_486672 | Institute for Stem Cell Science and Regenerative Medicine | National Centre for Biological Sciences | Farhan Ali, Vanessa Molin Paynter, Srikar Krishna, Mohak Sharda, Shah-e-Jahan Gulzar, Awadhesh Pandit, Varadha Sundarmurthy, Uma Ramakrishnan, Dasaradhi Palakodeti, Aswin Seshasayee |
| EPI_ISL_486673, EPI_ISL_486674 | Institute for Stem Cell Science and Regenerative Medicine | National Centre for Biological Sciences | Farhan Ali, Vanessa Molin Paynter, Srikar Krishna, Mohak Sharda, Shah-e-Jahan Gulzar, Awadhesh Pandit, Varadha Sundarmurthy, Uma Ramakrishnan, Dasaradhi Palakodeti, Aswin Seshasayee |
| EPI_ISL_486815, EPI_ISL_486816, EPI_ISL_486817, EPI_ISL_486818, EPI_ISL_486819, EPI_ISL_486820, EPI_ISL_486821, EPI_ISL_486822, EPI_ISL_486823, EPI_ISL_486824, EPI_ISL_486825, EPI_ISL_486826, EPI_ISL_486827, EPI_ISL_486828, EPI_ISL_486829 | | | |
| see above | Molecular diagnostic laboratory of Federal Budget Institution of Science "Central Research Institute of Epidemiology" of The Federal Service on Customers' Rights Protection and Human Well-being Surveillance | Group of Genomics and Postgenomic Technologies of Central Research Institute of Epidemilology | Speranskaya AS, Kaptelova VV, Valdokhina AV, Bulanenko VP, Samoilov AE, Korneenko EV, Tivanova EV, Shipulina OY, Akimkin VG |
| EPI_ISL_486830, EPI_ISL_486831 | Providence St. Joseph Health Molecular Genomics Laboratory | Providence St. Joseph Health Molecular Genomics Laboratory | Alexa K Dowdell, Brian D Piening, Fred L Robinson, Carlo B Bifulco, Mary Campbell |
| EPI_ISL_486834 | Suceava County Energency Hospital "Sf. Ioan cel Nou" | SMU Metagenomics lab | Lobiuc Andrei, Antoniadis Panagiotis |
| EPI_ISL_486835, EPI_ISL_486836, EPI_ISL_486837, EPI_ISL_486838, EPI_ISL_486839, EPI_ISL_486840, EPI_ISL_486841 | Institute for Stem Cell Science and Regenerative Medicine | National Centre for Biological Sciences | Farhan Ali, Vanessa Molin Paynter, Srikar Krishna, Mohak Sharda, Shah-e-Jahan Gulzar, Awadhesh Pandit, Varadha Sundarmurthy, Uma Ramakrishnan, Dasaradhi Palakodeti, Aswin Seshasayee |
| EPI_ISL_486842, EPI_ISL_486843, EPI_ISL_486844 | Institute of Microbiology, Universidad San Francisco de Quito | Institute of Microbiology, Universidad San Francisco de Quito | Belén Prado-Vivar, Sully Márquez, Juan José Guadalupe, Monica Becerra-Wong, Carla Torres, Bernardo Gutiérrez, Fausto Maldonado, Geovanny Carzola, Verónica Barragán, Patricio Rojas-Silva, Gabriel Trueba, Michelle Grunauer, Paúl Cárdenas |
| EPI_ISL_486845, EPI_ISL_486846, EPI_ISL_486847, EPI_ISL_486848, EPI_ISL_486849, EPI_ISL_486850, EPI_ISL_486851 | Institute of Microbiology, Universidad San Francisco de Quito | Institute of Microbiology, Universidad San Francisco de Quito | Belén Prado-Vivar, Sully Márquez, Juan José Guadalupe, Monica Becerra-Wong, Carla Torres, Bernardo Gutiérrez, Jonathan Araujo, Verónica Barragán, Patricio Rojas-Silva, Gabriel Trueba, Michelle Grunauer, Paúl Cárdenas |
| EPI_ISL_486852 | CDRI/SGPGI | CSIR-CDRI/SGPGI | Saumya Sarkar, Dharam Veer Singh, Rahul Vishvkarma, Ujjala Ghoshal, Uday Ghoshal, Ravishankar Ramachandran, Tapas Kumar Kundu, Rajender Singh |
| EPI_ISL_486853 | CSIR-CDRI/SGPGI | CSIR-CDRI/SGPGI | Saumya Sarkar, Dharam Veer Singh, Rahul Vishvkarma, Ujjala Ghoshal, Uday Ghoshal, Ravishankar Ramachandran, Tapas Kumar Kundu, Rajender Singh |
| EPI_ISL_486855 | Emergency county Hospital Suceava | "Stefan cel Mare" University Metagenomics Lab | Lobiuc Andrei et al. |
| EPI_ISL_486857, EPI_ISL_486858, EPI_ISL_486859, EPI_ISL_486860, EPI_ISL_486861, EPI_ISL_486862, EPI_ISL_486863, EPI_ISL_486864, EPI_ISL_486865, EPI_ISL_486866, EPI_ISL_486867, EPI_ISL_486868, EPI_ISL_486869, EPI_ISL_486870, EPI_ISL_486871, EPI_ISL_486872, EPI_ISL_486873 | | | |
| see above | Institut Pasteur Dakar | Institut Pasteur de Dakar | Ndongo Dia, Moussa Moise Diagne, Mamadou Diop, Marie Henriette Dior Ndione, Mamadou Malado Jallow, Safietou Sanke, Ousmane Faye, Amadou Alpha Sall. |
| EPI_ISL_486876 | Clinical Microbiology Laboratory- Basurto University Hospital | Biocruces-Bizkaia | Mikel J. Urrutikoetxea-Gutierrez, Ana Belén Belén de la Hoz, Matxalen Vidal-García, Mº Carmen Nieto Toboso, Estibaliz Ugalde-Zarraga, José Luis Díaz de Tuesta del Arco |
