## Supplementary material for "Ultrafast Sample Placement on Existing Trees (UShER) Empowers Real-Time Phylogenetics for the SARS-CoV-2 Pandemic": File_S3

We gratefully acknowledge the following Authors from the Originating laboratories responsible for obtaining the specimens, as well as the Submitting laboratories where the genome data were generated and shared via GISAID, on which this research is based.

All Submitters of data may be contacted directly via www.gisaid.org

| Accession ID | Originating Laboratory | Submitting Laboratory | Authors |
|---|---|---|---|
| EPI_ISL_470719, EPI_ISL_470720, EPI_ISL_470721, EPI_ISL_470722, EPI_ISL_470723, EPI_ISL_470724, EPI_ISL_470725, EPI_ISL_470726, EPI_ISL_470727, EPI_ISL_470728, EPI_ISL_470729, EPI_ISL_470730, EPI_ISL_470731, EPI_ISL_470732, EPI_ISL_470733, EPI_ISL_470734, EPI_ISL_470735, EPI_ISL_470736, EPI_ISL_470737, EPI_ISL_470738, EPI_ISL_470739, EPI_ISL_470740, EPI_ISL_470741, EPI_ISL_470742, EPI_ISL_470743, EPI_ISL_470744, EPI_ISL_470745, EPI_ISL_470746 | | | |
| see above | Utah Public Health Laboratory | Utah Public Health Laboratory | Heidi Butz, Erin Young, Kelly Oakeson |
| EPI_ISL_470747, EPI_ISL_470748, EPI_ISL_470749, EPI_ISL_470750, EPI_ISL_470751, EPI_ISL_470752, EPI_ISL_470753, EPI_ISL_470754, EPI_ISL_470755, EPI_ISL_470756, EPI_ISL_470757, EPI_ISL_470758, EPI_ISL_470759, EPI_ISL_470760, EPI_ISL_470761, EPI_ISL_470762, EPI_ISL_470763, EPI_ISL_470764, EPI_ISL_470765, EPI_ISL_470766, EPI_ISL_470767, EPI_ISL_470768, EPI_ISL_470769, EPI_ISL_470770, EPI_ISL_470771, EPI_ISL_470772, EPI_ISL_470773, EPI_ISL_470774, EPI_ISL_470775, EPI_ISL_470776, EPI_ISL_470777, EPI_ISL_470778, EPI_ISL_470779, EPI_ISL_470780, EPI_ISL_470781, EPI_ISL_470782, EPI_ISL_470783, EPI_ISL_470784, EPI_ISL_470785, EPI_ISL_470786, EPI_ISL_470787, EPI_ISL_470788, EPI_ISL_470789 | | | |
| see above | Minnesota Department of Health, Public Health Laboratory | Minnesota Department of Health, Public Health Laboratory | Matt Plumb, Jacob Garfin, and Xiong Wang |
| EPI_ISL_470790, EPI_ISL_470791, EPI_ISL_470792, EPI_ISL_470793, EPI_ISL_470794, EPI_ISL_470795, EPI_ISL_470796, EPI_ISL_470797, EPI_ISL_470798, EPI_ISL_470799, EPI_ISL_470800 | | | |
| see above | M Health Fairview | Minnesota Department of Health, Public Health Laboratory | Matt Plumb, Jacob Garfin, and Xiong Wang |
| EPI_ISL_470802 | State Key Laboratory of Agriculture Microbiology | State Key Laboratory of Agriculture Microbiology, Huazhong Agric | Zhong Zou |
| EPI_ISL_470830, EPI_ISL_470831, EPI_ISL_470832, EPI_ISL_470833, EPI_ISL_470834, EPI_ISL_470835, EPI_ISL_470836, EPI_ISL_470837, EPI_ISL_470838, EPI_ISL_470839, EPI_ISL_470840, EPI_ISL_470841, EPI_ISL_470842, EPI_ISL_470843, EPI_ISL_470844, EPI_ISL_470845, EPI_ISL_470846, EPI_ISL_470847, EPI_ISL_470848, EPI_ISL_470849, EPI_ISL_470850, EPI_ISL_470851, EPI_ISL_470852, EPI_ISL_470853, EPI_ISL_470854, EPI_ISL_470855, EPI_ISL_470856, EPI_ISL_470857, EPI_ISL_470858, EPI_ISL_470859, EPI_ISL_470860, EPI_ISL_470861, EPI_ISL_470862, EPI_ISL_470863, EPI_ISL_470864, EPI_ISL_470865, EPI_ISL_470866, EPI_ISL_470867, EPI_ISL_470868, EPI_ISL_470869, EPI_ISL_470870, EPI_ISL_470871, EPI_ISL_470872, EPI_ISL_470873, EPI_ISL_470874, EPI_ISL_470875 | | | |
| see above | PathWest Laboratory Medicine WA | PathWest Laboratory Medicine WA | Chisha Sikazwe, Jurissa Lang, Avram Levy, David Smith and David Speers |
| EPI_ISL_470876 | Department for Virology, Molecular Biology and Genome Research, R. G. Lugar Center for Public Health Research, National Center for Disease Control and Public Health (NCDC) of Georgia. | Department for Virology, Molecular Biology and Genome Research, R. G. Lugar Center for Public Health Research, National Center for Disease Control and Public Health (NCDC) of Georgia. | Giorgi Tomashvili, Meri Pantsulaia, Gvantsa Brachveli, Gvantsa Chanturia, Ann Machablishvili, Nato Kotaria, Marine Murtskhvaladze, Lela Sabadze, Mari Gavashelidze, Ana Papkiauri, Tata Imnadze, Tamar Jashiashvili, Tea Tevdoradze, Ketevan Sidamonidze, Ekaterine Khmaladze, Ekaterine Zhghenti, Roena Sukhiashvili, Mariam Zakalashvili, Lela Urushadze, Magda Dgebuadze, Davit Tsaguria, Ekaterine Zangaladze, Nino Berishvili, Adam Kotorashvili, Maia Alkhazashvili, Irma Burjanadze, Anna Kasradze, Khatuna Zakhashvili, Paata Imnadze, Amiran Gamkrelidze. |
| EPI_ISL_470877 | Department for Virology, Molecular Biology and Genome Research, R. G. Lugar Center for Public Health Research, National Center for Disease Control and Public Health (NCDC) of Georgia. | Department for Virology, Molecular Biology and Genome Research, R. G. Lugar Center for Public Health Research, National Center for Disease Control and Public Health (NCDC) of Georgia. | Gvantsa Brachveli, Meri Pantsulaia, Giorgi Tomashvili, Gvantsa Chanturia, Ann Machablishvili, Nato Kotaria, Marine Murtskhvaladze, Lela Sabadze, Mari Gavashelidze, Ana Papkiauri, Tata Imnadze, Tamar Jashiashvili, Tea Tevdoradze, Ketevan Sidamonidze, Ekaterine Khmaladze, Ekaterine Zhghenti, Roena Sukhiashvili, Mariam Zakalashvili, Lela Urushadze, Magda Dgebuadze, Davit Tsaguria, Ekaterine Zangaladze, Nino Berishvili, Adam Kotorashvili, Maia Alkhazashvili, Irma Burjanadze, Anna Kasradze, Khatuna Zakhashvili, Paata Imnadze, Amiran Gamkrelidze. |
| EPI_ISL_470878, EPI_ISL_470879, EPI_ISL_470880 | National Institute for Communicable Diseases of the National Health Laboratory Service | National Institute for Communicable Diseases of the National Health Laboratory Service | Allam M, Ismail A, Khumalo Z, Kwenda S, van Heusden P, Mtshali P, Mnyameni F, Mohale T, Subramoney K, Bhiman JN |
| EPI_ISL_470882 | Foerde Hospital, Department of Microbiology | Norwegian Institute of Public Health, Department of Virology | Kathrine Stene-Johansen, Kamilla Heddeland Instefjord, Hilde Elshaug, Rasmus Riis Kopperud, Karoline Bragstad, Olav Hungnes |
| EPI_ISL_470896 | Russian State Collection of Viruses | Pathogenic Microorganisms Variability Laboratory | Alexey Shchetinin, Maria Nikiforova, Elena Shidlovskaya, Nadezhda Kuznetsova, Inna Dolzhikova, Daria Grousova, Andrey Botikov, Denis Logunov, Alexander Gintsburg, Vladimir Gushchin |
| EPI_ISL_470897, EPI_ISL_470898, EPI_ISL_470899 | Pathogenic Microorganisms Variability Laboratory | Pathogenic Microorganisms Variability Laboratory | Alexey Shchetinin, Maria Nikiforova, Elena Shidlovskaya, Nadezhda Kuznetsova, Andrey Botikov, Alexander Gintsburg, Vladimir Gushchin |
| EPI_ISL_470900, EPI_ISL_470901, EPI_ISL_470902 | Influenza etiology and epidemiology laboratory | Pathogenic Microorganisms Variability Laboratory | Alexey Shchetinin, Maria Nikiforova, Elena Shidlovskaya, Nadezhda Kuznetsova, Vladimir Gushchin, Inna Dolzhikova, Daria Grousova, Andrey Botikov, Denis Logunov, Kirill Krasnoslobotsev, Svetlana Trushakova, Elena Burtseva, Ludmila Kolobukhina, Svetlana Smetanina, Alexander Gintsburg |
| EPI_ISL_470903, EPI_ISL_470904 | Influenza etiology and epidemiology laboratory | Pathogenic Microorganisms Variability Laboratory | Alexey Shchetinin, Maria Nikiforova, Elena Shidlovskaya, Nadezhda Kuznetsova, Vladimir Gushchin, Inna Dolzhikova, Daria Grousova, Andrey Botikov, Denis Logunov, Anna Ignatjeva, Evgeniya Mukasheva, Elena Burtseva, Ludmila Kolobukhina, Svetlana Smetanina, Alexander Gintsburg |
| EPI_ISL_471144, EPI_ISL_471145, EPI_ISL_471146, EPI_ISL_471147, EPI_ISL_471148, EPI_ISL_471149, EPI_ISL_471150, EPI_ISL_471151, EPI_ISL_471152, EPI_ISL_471153, EPI_ISL_471154, EPI_ISL_471155, EPI_ISL_471156 | | | |
| see above | Gundersen Molecular Diagnostics Laboratory | Kabara Cancer Research Institute | Craig S. Richmond, Paraic A. Kenny |
| EPI_ISL_471157 | Gundersen Clinical Microbiology Laboratory | Kabara Cancer Research Institute | Craig S. Richmond, Paraic A. Kenny |
| EPI_ISL_471158, EPI_ISL_471159, EPI_ISL_471160, EPI_ISL_471161, EPI_ISL_471162, EPI_ISL_471163, EPI_ISL_471164, EPI_ISL_471165, EPI_ISL_471166, EPI_ISL_471167, EPI_ISL_471168, EPI_ISL_471169, EPI_ISL_471170, EPI_ISL_471171 | | | |
| see above | MRCG at LSHTM Genomics lab | MRCG at LSHTM Genomics lab | Sesay et al |
| EPI_ISL_471172 | Unilabs Laboratory Medicine | Norwegian Institute of Public Health, Department of Virology | Kathrine Stene-Johansen, Kamilla Heddeland Instefjord, Hilde Elshaug, Rasmus Riis Kopperud, Karoline Bragstad, Olav Hungnes |
| EPI_ISL_471173 | Hospital of Southern Norway - Kristiansand, Department of Medical Microbiology | Norwegian Institute of Public Health, Department of Virology | Kathrine Stene-Johansen, Kamilla Heddeland Instefjord, Hilde Elshaug, Rasmus Riis Kopperud, Karoline Bragstad, Olav Hungnes |
| EPI_ISL_471174 | Ostfold Hospital Trust - Kalnes, Centre for Laboratory Medicine, Section for gene technology and infection serology | Norwegian Institute of Public Health, Department of Virology | Kathrine Stene-Johansen, Kamilla Heddeland Instefjord, Hilde Elshaug, Rasmus Riis Kopperud, Karoline Bragstad, Olav Hungnes |
| EPI_ISL_471175 | Oslo University Hospital, Department of Medical Microbiology | Norwegian Institute of Public Health, Department of Virology | Kathrine Stene-Johansen, Kamilla Heddeland Instefjord, Hilde Elshaug, Rasmus Riis Kopperud, Karoline Bragstad, Olav Hungnes |
| EPI_ISL_471176 | Hospital of Southern Norway - Kristiansand, Department of Medical Microbiology | Norwegian Institute of Public Health, Department of Virology | Kathrine Stene-Johansen, Kamilla Heddeland Instefjord, Hilde Elshaug, Rasmus Riis Kopperud, Karoline Bragstad, Olav Hungnes |
| EPI_ISL_471177 | Oslo University Hospital, Department of Medical Microbiology | Norwegian Institute of Public Health, Department of Virology | Kathrine Stene-Johansen, Kamilla Heddeland Instefjord, Hilde Elshaug, Rasmus Riis Kopperud, Karoline Bragstad, Olav Hungnes |
| EPI_ISL_471178, EPI_ISL_471179, EPI_ISL_471180, EPI_ISL_471181, EPI_ISL_471182, EPI_ISL_471183, EPI_ISL_471184, EPI_ISL_471185, EPI_ISL_471186, EPI_ISL_471187, EPI_ISL_471188, EPI_ISL_471189, EPI_ISL_471190, EPI_ISL_471191, EPI_ISL_471192, EPI_ISL_471193, EPI_ISL_471194, EPI_ISL_471195, EPI_ISL_471196, EPI_ISL_471197, EPI_ISL_471198, EPI_ISL_471199, EPI_ISL_471200, EPI_ISL_471201, EPI_ISL_471202, EPI_ISL_471203, EPI_ISL_471204, EPI_ISL_471205, EPI_ISL_471206, EPI_ISL_471207, EPI_ISL_471208, EPI_ISL_471209, EPI_ISL_471210, EPI_ISL_471211, EPI_ISL_471212, EPI_ISL_471213, EPI_ISL_471214, EPI_ISL_471215, EPI_ISL_471216, EPI_ISL_471217, EPI_ISL_471218, EPI_ISL_471219, EPI_ISL_471220, EPI_ISL_471221, EPI_ISL_471222, EPI_ISL_471223, EPI_ISL_471224, EPI_ISL_471225, EPI_ISL_471226, EPI_ISL_471227, EPI_ISL_471228, EPI_ISL_471229, EPI_ISL_471230, EPI_ISL_471231, EPI_ISL_471232, EPI_ISL_471233, EPI_ISL_471234, EPI_ISL_471235, EPI_ISL_471236, EPI_ISL_471237, EPI_ISL_471238, EPI_ISL_471239, EPI_ISL_471240, EPI_ISL_471241, EPI_ISL_471242, EPI_ISL_471243, EPI_ISL_471244, EPI_ISL_471245, EPI_ISL_471246, EPI_ISL_471247, EPI_ISL_471248, EPI_ISL_471249, EPI_ISL_471250, EPI_ISL_471251, EPI_ISL_471252, EPI_ISL_471253, EPI_ISL_471254, EPI_ISL_471255, EPI_ISL_471256, EPI_ISL_471257, EPI_ISL_471258, EPI_ISL_471259, EPI_ISL_471260, EPI_ISL_471261, EPI_ISL_471262, EPI_ISL_471263, EPI_ISL_471264, EPI_ISL_471265, EPI_ISL_471266 | | | |

| | | | |
|---|---|---|---|
| see above | Wisconsin State Laboratory of Hygiene Communicable Disease Division | Wisconsin State Laboratory of Hygiene Communicable Disease Division | Kelsey R. Florek, Abigail C. Shockey |
| EPI_ISL_471267 | Hospital IESS Babahoyo | Institute of Microbiology, Universidad San Francisco de Quito | Sully Márquez, Belén Prado-Vivar, Juan José Guadalupe, Bernardo Gutiérrez, Francisco Cordova, Ninfa Henriquez, Killen Briones-Zamora, Killen Briones-Claudette, Verónica Barragán, Patricio Rojas-Silva, Gabriel Trueba, Michelle Grunauer, Paúl Cárdenas |
| EPI_ISL_471268 | Hospital IESS Babahoyo | Institute of Microbiology, Universidad San Francisco de Quito | Belén Prado-Vivar, Sully Márquez, Juan José Guadalupe, Bernardo Gutiérrez, Francisco Cordova, Ninfa Henriquez, Killen Briones-Zamora, Killen Briones-Claudette, Verónica Barragán, Patricio Rojas-Silva, Gabriel Trueba, Michelle Grunauer, Paúl Cárdenas |
| EPI_ISL_471269 | Hospital Oncológico Solca Núcleo de Quito | Institute of Microbiology, Universidad San Francisco de Quito | Sully Márquez, Belén Prado-Vivar, Juan José Guadalupe, Bernardo Gutiérrez, Marcos Di Stefano, Grace Salazar, Verónica Barragán, Patricio Rojas-Silva, Gabriel Trueba, Michelle Grunauer, Paúl Cárdenas |
| EPI_ISL_471270 | Hospital Oncológico Solca Núcleo de Quito | Institute of Microbiology, Universidad San Francisco de Quito | Sully Márquez, Belén Prado-Vivar, Juan José Guadalupe, Bernardo Gutiérrez, Marcos Di Stefano, Grace Salazar, Verónica Barragán, Patricio Rojas-Silva, Gabriel Trueba, Michelle Grunauer, Paúl Cárdenas |
| EPI_ISL_471271 | Hospital Oncológico Solca Núcleo de Quito | Institute of Microbiology, Universidad San Francisco de Quito | Sully Márquez, Belén Prado-Vivar, Juan José Guadalupe, Bernardo Gutiérrez, Marcos Di Stefano, Grace Salazar, Verónica Barragán, Patricio Rojas-Silva, Gabriel Trueba, Michelle Grunauer, Paúl Cárdenas |
| EPI_ISL_471396, EPI_ISL_471397, EPI_ISL_471398, EPI_ISL_471399, EPI_ISL_471400, EPI_ISL_471401, EPI_ISL_471402, EPI_ISL_471403, EPI_ISL_471404, EPI_ISL_471405, EPI_ISL_471406, EPI_ISL_471407, EPI_ISL_471408, EPI_ISL_471409, EPI_ISL_471410, EPI_ISL_471411, EPI_ISL_471412, EPI_ISL_471413, EPI_ISL_471414, EPI_ISL_471415 | | | |
| see above | Viral Respiratory Lab, National Institute for Biomedical Research (INRB) | Pathogen Sequencing Lab, National Institute for Biomedical Research (INRB) | Placide Mbala-Kingebeni, Edith Nkwembe, Eddy Kinganda-Lusamaki, Amuri Aziza, Francisca Muyembe Mawete, Catherine Pratt, Matthias Pauthner, Josh Quick, Allison Black, James Hadfield, Trevor Bedford, Ian Goodfellow, Andrew Rambaut, Nick Loman, Kristian Andersen, Michael Wiley, Steve Ahuka-Mundeke, Jean-Jacques Muyembe Tamfum |
| EPI_ISL_471416, EPI_ISL_471417, EPI_ISL_471418, EPI_ISL_471419, EPI_ISL_471420, EPI_ISL_471421, EPI_ISL_471422, EPI_ISL_471423, EPI_ISL_471424 | Laboratory for Respiratory Viruses, National Influenza Centre, Cantacuzino National Military-Medical Institute for Research and Development | Cantacuzino Institute | Luiza Ustea, Nicoleta Paraschiv, Tim Durfee, Mihaela Lazar |
| EPI_ISL_471425 | Division of Viral Diseases, Center for Laboratory Control of Infectious Diseases, Korea Centers for Diseases Control and Prevention | Division of Viral Diseases, Center for Laboratory Control of Infectious Diseases, Korea Centers for Diseases Control and Prevention | Jeong-Min Kim, Yoon-Seok Chung, Namjoo Lee, Mi-Seon Kim, Sang Hee Woo, Hye-Jun Jo, Sehee Park, Heui Man Kim, Jun-Sub Kim, Junhyeong Jang, Dong Hyun Song, Daesang Lee, Seong Tae Jeong, Myung Guk Han |
| EPI_ISL_471426 | Division of Viral Diseases, Center for Laboratory Control of Infectious Diseases, Korea Centers for Diseases Control and Prevention | Division of Viral Diseases, Center for Laboratory Control of Infectious Diseases, Korea Centers for Diseases Control and Prevention | Jeong-Min Kim, Yoon-Seok Chung, Namjoo Lee, Mi-Seon Kim, Sang Hee Woo, Hye-Jun Jo, Sehee Park, Heui Man Kim, Jun-Sub Kim, Junhyeong Jang, Dong Hyun Song, Daesang Lee, Seong Tae Jeong, Myung Guk Han |
| EPI_ISL_471427, EPI_ISL_471428, EPI_ISL_471429, EPI_ISL_471430, EPI_ISL_471431, EPI_ISL_471432, EPI_ISL_471433, EPI_ISL_471434, EPI_ISL_471435, EPI_ISL_471436, EPI_ISL_471437 | | | |
| see above | Department of Clinical Microbiology | GIGA Medical Genomics | Keith Durkin, Maria Artesi, Sébastien Bontems, Raphaël Boreux, Cécile Meex, Axelle Chaslain, Céline Fombellida-Lopez, Pierrette Melin, Marie-Pierre Hayette, Vincent Bours. |
| EPI_ISL_471438, EPI_ISL_471439, EPI_ISL_471440, EPI_ISL_471441, EPI_ISL_471442, EPI_ISL_471443, EPI_ISL_471444 | Division of Viral Diseases, Center for Laboratory Control of Infectious Diseases, Korea Centers for Diseases Control and Prevention | Division of Viral Diseases, Center for Laboratory Control of Infectious Diseases, Korea Centers for Diseases Control and Prevention | Jeong-Min Kim, Yoon-Seok Chung, Namjoo Lee, Sang Hee Woo, Hye-Jun Jo, Heui Man Kim, Jun-Sub Kim, Dong Hyun Song, Daesang Lee, Seong Tae Jeong, Myung Guk Han |
| EPI_ISL_471445 | Division of Viral Diseases, Center for Laboratory Control of Infectious Diseases, Korea Centers for Diseases Control and Prevention | Division of Viral Diseases, Center for Laboratory Control of Infectious Diseases, Korea Centers for Diseases Control and Prevention | Jeong-Min Kim, Yoon-Seok Chung, Namjoo Lee, Sang Hee Woo, Hye-Jun Jo, Heui Man Kim, Jun-Sub Kim, Myung Guk Han |
| EPI_ISL_471446, EPI_ISL_471447, EPI_ISL_471448, EPI_ISL_471449, EPI_ISL_471450, EPI_ISL_471451, EPI_ISL_471452 | Division of Viral Diseases, Center for Laboratory Control of Infectious Diseases, Korea Centers for Diseases Control and Prevention | Division of Viral Diseases, Center for Laboratory Control of Infectious Diseases, Korea Centers for Diseases Control and Prevention | Jeong-Min Kim, Yoon-Seok Chung, Namjoo Lee, Sang Hee Woo, Hye-Jun Jo, Heui Man Kim, Jun-Sub Kim, Dong Hyun Song, Daesang Lee, Seong Tae Jeong, Myung Guk Han |
| EPI_ISL_471453 | Division of Viral Diseases, Center for Laboratory Control of Infectious Diseases, Korea Centers for Diseases Control and Prevention | Division of Viral Diseases, Center for Laboratory Control of Infectious Diseases, Korea Centers for Diseases Control and Prevention | Jeong-Min Kim, Yoon-Seok Chung, Namjoo Lee, Sang Hee Woo, Hye-Jun Jo, Heui Man Kim, Jun-Sub Kim, Myung Guk Han |
| EPI_ISL_471454, EPI_ISL_471455 | Division of Viral Diseases, Center for Laboratory Control of Infectious Diseases, Korea Centers for Diseases Control and Prevention | Division of Viral Diseases, Center for Laboratory Control of Infectious Diseases, Korea Centers for Diseases Control and Prevention | Jeong-Min Kim, Yoon-Seok Chung, Namjoo Lee, Sang Hee Woo, Hye-Jun Jo, Heui Man Kim, Jun-Sub Kim, Dong Hyun Song, Daesang Lee, Seong Tae Jeong, Myung Guk Han |
| EPI_ISL_471456, EPI_ISL_471457, EPI_ISL_471458, EPI_ISL_471459, EPI_ISL_471460 | Centre de Virologie des Maladies Tropicales | Functional Genomic Platform/Service Analyses Biologique/UATRS/ Centre National Pour la Recherche Scientifique Et Technique (CNRST) | Hicham ANNAZ, Elmostafa EL FAHIME, Marouane MELLOUL, Yassine AKHOUAD, Mly Abdelaziz ELALAOUI, Ahmed REGGAD, Sanaa ALAOUI-Amine , Rachid ABI, Rida TAGAJDID, Zhor KASMY, Safaa ELKORCHI, Nadia TOUIL, Farida HILALI, Abdelkader LAATIRIS , Abdelillah LARAQUI, Tahra BAJJOU , Yassine SEKHSOKH , Idriss-Amine LAHLOU, Mostafa ELOUENNASS, Khalid ENNIBI |
| EPI_ISL_471461 | South China Agricultural University | South China Agricultural University | Yongyi Shen, Lihua Xiao, Wu Chen |
| EPI_ISL_471462, EPI_ISL_471463 | South China Agricultural University | South China Agricultural University | Yongyi Shen, Lihua Xiao, Wu Chen |
| EPI_ISL_471464 | South China Agricultural University | South China Agricultural University | Yongyi Shen, Lihua Xiao, Wu Chen |
| EPI_ISL_471465, EPI_ISL_471466 | South China Agricultural University | South China Agricultural University | Yongyi Shen, Lihua Xiao, Wu Chen |
| EPI_ISL_471467, EPI_ISL_471468, EPI_ISL_471469, EPI_ISL_471470 | South China Agricultural University | South China Agricultural University | Yongyi Shen, Wu Chen |
| EPI_ISL_471472 | Hospital Universitari Germans Trias i Pujol(HUGTiP)/Fundació Lluita contra la SIDA (FLSida)/IRTA-CReSA | IrsiCaixa AIDS Research Lab | Marc Noguera-Julian, Pilar Armengol, Jordi Rodón, Julia Vergara, Lidia Ruiz, Nuria Izquierdo, Jorge Carrillo, Roger Paredes, Albert Bensaid, Julia Blanco, Joaquim Segalés, Bonaventura Clotet |
| EPI_ISL_471510, EPI_ISL_471511, EPI_ISL_471512, EPI_ISL_471513, EPI_ISL_471514, EPI_ISL_471515, EPI_ISL_471516, EPI_ISL_471517, EPI_ISL_471518, EPI_ISL_471519, EPI_ISL_471520, EPI_ISL_471521, EPI_ISL_471522, EPI_ISL_471523, EPI_ISL_471524, EPI_ISL_471525, EPI_ISL_471526, EPI_ISL_471527 | | | |
| see above | Respiratory Virus Unit, Microbiology Services Colindale, Public Health England | Respiratory Virus Unit, Microbiology Services Colindale, Public Health England | PHE Covid Sequencing Team |
| EPI_ISL_471528 | The National Institute of Public Health | State Veterinary Institute Prague and The National Institute of Public Health | Nagy,A;Jirincova,H;Novakova,L;Trnka,D;Vecerova,J |
| EPI_ISL_471529 | Department for Virology, Molecular Biology and Genome Research, R. G. Lugar Center for Public Health Research, National Center for Disease Control and Public Health (NCDC) of Georgia. | Department for Virology, Molecular Biology and Genome Research, R. G. Lugar Center for Public Health Research, National Center for Disease Control and Public Health (NCDC) of Georgia. | Meri Pantsulaia, Gvantsa Brachveli, Giorgi Tomashvili, Gvantsa Chanturia, Ann Machablishvili, Nato Kotaria, Marine Murtskhvaladze, Lela Sabadze, Mari Gavashelidze, Ana Papkiauri, Gvantsa Brachveli, Tata Imnadze, Tamar Jashiashvili, Tea Tevdoradze, Ketevan Sidamonidze, Ekaterine Khmaladze, Ekaterine Zhghenti, Roena Sukhiashvili, Mariam Zakalashvili, Lela Urushadze, Magda Dgebuadze, Davit Tsaguria, Ekaterine Zangaladze, Nino Berishvili, Adam Kotorashvili, Maia Alkhazashvili, Irma Burjanadze, Anna Kasradze, Khatuna Zakhashvili, Paata Imnadze, Amiran Gamkrelidze. |
| EPI_ISL_471530 | The National Institute of Public Health | State Veterinary Institute Prague and The National Institute of Public Health | Nagy,A;Jirincova,H;Novakova,L;Trnka,D;Vecerova,J |
| EPI_ISL_471539 | Hospital Universitario da USP Sao Paulo | Instituto Adolfo Lutz, Interdiciplinary Procedures Center, Strategic Laboratory | Claudio Tavares Sacchi, Claudia Regina Gonçalves, Erica Valessa Ramos Gomes |
| EPI_ISL_471540 | The National Institute of Public Health | State Veterinary Institute Prague and The National Institute of Public Health | Nagy,A;Jirincova,H;Novakova,L;Trnka,D;Vecerova,J |

| | | | |
|---|---|---|---|
| EPI_ISL_471541 | Hospital Geral Santa Marcelina | Instituto Adolfo Lutz, Interdiciplinary Procedures Center, Strategic Laboratory | Claudio Tavares Sacchi, Claudia Regina Gonçalves, Erica Valessa Ramos Gomes |
| EPI_ISL_471542 | Secretaria de Saude de Mogi das Cruzes | Instituto Adolfo Lutz, Interdiciplinary Procedures Center, Strategic Laboratory | Claudio Tavares Sacchi, Claudia Regina Gonçalves, Erica Valessa Ramos Gomes |
| EPI_ISL_471543 | Centro de Saude I Tacito Leite de Carvalho e Silva | Instituto Adolfo Lutz, Interdiciplinary Procedures Center, Strategic Laboratory | Claudio Tavares Sacchi, Claudia Regina Gonçalves, Erica Valessa Ramos Gomes |
| EPI_ISL_471544 | The National Institute of Public Health | State Veterinary Institute Prague and The National Institute of Public Health | Nagy,A;Jirincova,H;Novakova,L;Trnka,D;Vecerova,J |
| EPI_ISL_471545 | Hospital Sao Paulo de Ensino da Unifesp | Instituto Adolfo Lutz, Interdiciplinary Procedures Center, Strategic Laboratory | Claudio Tavares Sacchi, Claudia Regina Gonçalves, Erica Valessa Ramos Gomes |
| EPI_ISL_471546 | AMA DR Jose Soares Hungria | Instituto Adolfo Lutz, Interdiciplinary Procedures Center, Strategic Laboratory | Claudio Tavares Sacchi, Claudia Regina Gonçalves, Erica Valessa Ramos Gomes |
| EPI_ISL_471547 | The National Institute of Public Health | State Veterinary Institute Prague and The National Institute of Public Health | Nagy,A;Jirincova,H;Novakova,L;Trnka,D;Vecerova,J |
| EPI_ISL_471548 | Hospital do Servidor Público Estadual Francisco Morato de Oliveira | Instituto Adolfo Lutz, Interdiciplinary Procedures Center, Strategic Laboratory | Claudio Tavares Sacchi, Claudia Regina Gonçalves, Erica Valessa Ramos Gomes |
| EPI_ISL_471549 | Hospital Municipal Carmen Prudente | Instituto Adolfo Lutz, Interdiciplinary Procedures Center, Strategic Laboratory | Claudio Tavares Sacchi, Claudia Regina Gonçalves, Erica Valessa Ramos Gomes |
| EPI_ISL_471550 | The National Institute of Public Health | State Veterinary Institute Prague and The National Institute of Public Health | Nagy,A;Jirincova,H;Novakova,L;Trnka,D;Vecerova,J |
| EPI_ISL_471551 | Hospital Sao Paulo de Ensino da Unifesp | Instituto Adolfo Lutz, Interdiciplinary Procedures Center, Strategic Laboratory | Claudio Tavares Sacchi, Claudia Regina Gonçalves, Erica Valessa Ramos Gomes |
| EPI_ISL_471552 | Hospital Sancta Maggiore | Instituto Adolfo Lutz, Interdiciplinary Procedures Center, Strategic Laboratory | Claudio Tavares Sacchi, Claudia Regina Gonçalves, Erica Valessa Ramos Gomes |
| EPI_ISL_471553 | The National Institute of Public Health | State Veterinary Institute Prague and The National Institute of Public Health | Nagy,A;Jirincova,H;Novakova,L;Trnka,D;Vecerova,J |
| EPI_ISL_471554 | Hospital Bosque da Saúde | Instituto Adolfo Lutz, Interdiciplinary Procedures Center, Strategic Laboratory | Claudio Tavares Sacchi, Claudia Regina Gonçalves, Erica Valessa Ramos Gomes |
| EPI_ISL_471555 | The National Institute of Public Health | State Veterinary Institute Prague and The National Institute of Public Health | Nagy,A;Jirincova,H;Novakova,L;Trnka,D;Vecerova,J |
| EPI_ISL_471556 | Pronto Socorro Jose Ibrahin | Instituto Adolfo Lutz, Interdiciplinary Procedures Center, Strategic Laboratory | Claudio Tavares Sacchi, Claudia Regina Gonçalves, Erica Valessa Ramos Gomes |
| EPI_ISL_471562, EPI_ISL_471581, EPI_ISL_471582 | Hosp. Municipal Prof. Dr. Alípio Corrêa Netto | Instituto Adolfo Lutz, Interdiciplinary Procedures Center, Strategic Laboratory | Claudio Tavares Sacchi, Claudia Regina Gonçalves, Erica Valessa Ramos Gomes |
| EPI_ISL_471583, EPI_ISL_471584 | King Institute of Preventive Medicine & Research | CSIR-Centre for Cellular and Molecular Biology | K.Kaveri,S.Sivasubramanian,S.Vennila,P.Padmapriya,R.Kiruba,S.Magesh,G. Dhinakar Raj, G. Ravikumar, P. Azhahianambi,K Thangaraj,Payel Mukherjee, Sofia Banu, Priya Singh, Dhiviya Vedagiri, Divya Gupta, Vishal Sah, Santosh Kumar Kuncha, Krishnan Harinivas Harshan, Archana Bharadwaj Siva, Karthik Bharadwaj Tallapaka, Shagufta Khan, Lamuk Zaveri, Namami Gaur, Sakshi Shambhavi, Tulasi Nagabandi, Purushotham Vodnala, Rakesh K Mishra, Divya Tej Sowpati |
| EPI_ISL_471585 | CSIR-Centre for Cellular and Molecular Biology | CSIR-Centre for Cellular and Molecular Biology | Dhiviya Vedagiri, Divya Gupta, Vishal Sah, Payel Mukherjee, Sofia Banu, Priya Singh, Santosh Kumar Kuncha, Archana Bharadwaj Siva, Karthik Bharadwaj Tallapaka, Shagufta Khan, Lamuk Zaveri, Namami Gaur, Sakshi Shambhavi, Tulasi Nagabandi, Purushotham Vodnala, Rakesh K Mishra, Divya Tej Sowpati, Krishnan Harinivas Harshan |
| EPI_ISL_471586 | CSIR-Centre for Cellular and Molecular Biology | CSIR-Centre for Cellular and Molecular Biology | Lamuk Zaveri, Shagufta Khan, Namami Gaur, Sakshi Shambhavi, Tulasi Nagabandi, Purushotham Vodnala, Payel Mukherjee, Sofia Banu, Priya Singh, Dhiviya Vedagiri, Divya Gupta, Vishal Sah, Santosh Kumar Kuncha, Krishnan Harinivas Harshan, Archana Bharadwaj Siva, Karthik Bharadwaj Tallapaka,Zeba Rizvi, Zuberwasim Sayyad, Kakade Aishwarya Arun, Amrutha H C, Ananga Ghosh, Rakesh K Mishra, Divya Tej Sowpati |
| EPI_ISL_471587 | CSIR-Centre for Cellular and Molecular Biology | CSIR-Centre for Cellular and Molecular Biology | Dhiviya Vedagiri, Divya Gupta, Vishal Sah, Payel Mukherjee, Sofia Banu, Priya Singh, Santosh Kumar Kuncha, Archana Bharadwaj Siva, Karthik Bharadwaj Tallapaka, Shagufta Khan, Lamuk Zaveri, Namami Gaur, Sakshi Shambhavi, Tulasi Nagabandi, Purushotham Vodnala, Rakesh K Mishra, Divya Tej Sowpati, Krishnan Harinivas Harshan |
| EPI_ISL_471588 | CSIR-Centre for Cellular and Molecular Biology | CSIR-Centre for Cellular and Molecular Biology | Lamuk Zaveri, Shagufta Khan, Namami Gaur, Sakshi Shambhavi, Tulasi Nagabandi, Purushotham Vodnala, Payel Mukherjee, Sofia Banu, Priya Singh, Dhiviya Vedagiri, Divya Gupta, Vishal Sah, Santosh Kumar Kuncha, Krishnan Harinivas Harshan, Archana Bharadwaj Siva, Karthik Bharadwaj Tallapaka, Renu Sudhakar, Somesh Gorde, Gangumala Srinivas Reddy, Sujoy Deb, Swati Bayyana, Rakesh K Mishra, Divya Tej Sowpati |
| EPI_ISL_471589 | CSIR-Centre for Cellular and Molecular Biology | CSIR-Centre for Cellular and Molecular Biology | Lamuk Zaveri, Shagufta Khan, Namami Gaur, Sakshi Shambhavi, Tulasi Nagabandi, Purushotham Vodnala, Payel Mukherjee, Sofia Banu, Priya Singh, Dhiviya Vedagiri, Divya Gupta, Vishal Sah, Santosh Kumar Kuncha, Krishnan Harinivas Harshan, Archana Bharadwaj Siva, Karthik Bharadwaj Tallapaka,Umesh Kumar, Unis Ahmad Bhat, Ajay Sarawagi, Priyanka Pant, Rajkanwar Nathawat, Rakesh K Mishra, Divya Tej Sowpati |
| EPI_ISL_471590 | CSIR-Centre for Cellular and Molecular Biology | CSIR-Centre for Cellular and Molecular Biology | Lamuk Zaveri, Shagufta Khan, Namami Gaur, Sakshi Shambhavi, Tulasi Nagabandi, Purushotham Vodnala, Payel Mukherjee, Sofia Banu, Priya Singh, Dhiviya Vedagiri, Divya Gupta, Vishal Sah, Santosh Kumar Kuncha, Krishnan Harinivas Harshan, Archana Bharadwaj Siva, Karthik Bharadwaj Tallapaka,Zeba Rizvi, Zuberwasim Sayyad, Kakade Aishwarya Arun, Amrutha H C, Ananga Ghosh, Rakesh K Mishra, Divya Tej Sowpati |
| EPI_ISL_471591 | CSIR-Centre for Cellular and Molecular Biology | CSIR-Centre for Cellular and Molecular Biology | Namami Gaur, Sakshi Shambhavi, Lamuk Zaveri, Shagufta Khan, Tulasi Nagabandi, Purushotham Vodnala, Payel Mukherjee, Sofia Banu, Priya Singh, Dhiviya Vedagiri, Divya Gupta, Vishal Sah, Santosh Kumar Kuncha, Krishnan Harinivas Harshan, Archana Bharadwaj Siva, Karthik Bharadwaj Tallapaka, Zeba Rizvi, Zuberwasim Sayyad, Kakade Aishwarya Arun, Amrutha H C, Ananga Ghosh, Rakesh K Mishra, Divya Tej Sowpati |
| EPI_ISL_471592 | CSIR-Centre for Cellular and Molecular Biology | CSIR-Centre for Cellular and Molecular Biology | Namami Gaur, Sakshi Shambhavi, Lamuk Zaveri, Shagufta Khan, Tulasi Nagabandi, Purushotham Vodnala, Payel Mukherjee, Sofia Banu, Priya Singh, Dhiviya Vedagiri, Divya Gupta, Vishal Sah, Santosh Kumar Kuncha, Krishnan Harinivas Harshan, Archana Bharadwaj Siva, Karthik Bharadwaj Tallapaka, Nikhil Hajirnis, Pratheusa Maccha, M Soujanya Reddy,G. Aditya Kumar, Koushick Sivakumar, Rakesh K Mishra, Divya Tej Sowpati |
| EPI_ISL_471593 | CSIR-Centre for Cellular and Molecular Biology | CSIR-Centre for Cellular and Molecular Biology | Namami Gaur, Sakshi Shambhavi, Lamuk Zaveri, Shagufta Khan, Tulasi Nagabandi, Purushotham Vodnala, Payel Mukherjee, Sofia Banu, Priya Singh, Dhiviya Vedagiri, Divya Gupta, Vishal Sah, Santosh Kumar Kuncha, Krishnan Harinivas Harshan, Archana Bharadwaj Siva, Karthik Bharadwaj Tallapaka, Zeba Rizvi, Zuberwasim Sayyad, Kakade Aishwarya Arun, Amrutha H C, Ananga Ghosh, Rakesh K Mishra, Divya Tej Sowpati |
| EPI_ISL_471594 | CSIR-Centre for Cellular and Molecular Biology | CSIR-Centre for Cellular and Molecular Biology | Namami Gaur, Sakshi Shambhavi, Lamuk Zaveri, Shagufta Khan, Tulasi Nagabandi, Purushotham Vodnala, Payel Mukherjee, Sofia Banu, Priya Singh, Dhiviya Vedagiri, Divya Gupta, Vishal Sah, Santosh Kumar Kuncha, Krishnan Harinivas Harshan, Archana Bharadwaj Siva, Karthik Bharadwaj Tallapaka,Kezia J Ann, Radhika Khandelwal, Roshan Maku Venkata, Shemin Mansuri, Sonu Uday, Rakesh K Mishra, Divya Tej Sowpati |
| EPI_ISL_471595 | CSIR-Centre for Cellular and Molecular Biology | CSIR-Centre for Cellular and Molecular Biology | Payel Mukherjee, Sofia Banu, Priya Singh, Dhiviya Vedagiri, Divya Gupta, Vishal Sah, Santosh Kumar Kuncha, Krishnan Harinivas Harshan, Archana Bharadwaj Siva, Karthik Bharadwaj Tallapaka, Shagufta Khan, Lamuk Zaveri, Namami Gaur, Sakshi Shambhavi, Tulasi Nagabandi, Purushotham Vodnala, G. Aditya Kumar, Koushick Sivakumar, Pooja Ramesh Gupta, Rajan Kumar Jha, Shraddha Vijay Lahoti, Rakesh K Mishra, Divya Tej Sowpati |
| EPI_ISL_471596 | CSIR-Centre for Cellular and Molecular Biology | CSIR-Centre for Cellular and Molecular Biology | Payel Mukherjee, Sofia Banu, Priya Singh, Dhiviya Vedagiri, Divya Gupta, Vishal Sah, Santosh Kumar Kuncha, Krishnan Harinivas Harshan, Archana Bharadwaj Siva, Karthik Bharadwaj Tallapaka, Shagufta Khan, Lamuk Zaveri, Namami Gaur, Sakshi Shambhavi, Tulasi Nagabandi, Purushotham Vodnala, |

| | | | |
|---|---|---|---|
| | | | Gokulan C G, Gunjan Purohit, Hanuman Tulashiram Kale, Pankaj Kumar, Prachand Issarapu, Rakesh K Mishra, Divya Tej Sowpati |
| EPI_ISL_471597 | CSIR-Centre for Cellular and Molecular Biology | CSIR-Centre for Cellular and Molecular Biology | Payel Mukherjee, Sofia Banu, Priya Singh, Dhiviya Vedagiri, Divya Gupta, Vishal Sah, Santosh Kumar Kuncha, Krishnan Harinivas Harshan, Archana Bharadwaj Siva, Karthik Bharadwaj Tallapaka, Shagufta Khan, Lamuk Zaveri, Namami Gaur, Sakshi Shambhavi, Tulasi Nagabandi, Purushotham Vodnala, Rakesh K Mishra, Sonu Uday, Sudipta Mondal, Annapoorna P Karthyayani, Debabrata Jana, Debrya Saha, Divya Tej Sowpati |
| EPI_ISL_471598 | CSIR-Centre for Cellular and Molecular Biology | CSIR-Centre for Cellular and Molecular Biology | Payel Mukherjee, Sofia Banu, Priya Singh, Dhiviya Vedagiri, Divya Gupta, Vishal Sah, Santosh Kumar Kuncha, Krishnan Harinivas Harshan, Archana Bharadwaj Siva, Karthik Bharadwaj Tallapaka, Shagufta Khan, Lamuk Zaveri, Namami Gaur, Sakshi Shambhavi, Tulasi Nagabandi, Purushotham Vodnala,Deepak Kumar, Devi Prasad Vijayashankar, Disha Nanda, Divya Das, Jotin Gogoi, Manish Bhattacharjee, Rakesh K Mishra, Divya Tej Sowpati |
| EPI_ISL_471599 | CSIR-Centre for Cellular and Molecular Biology | CSIR-Centre for Cellular and Molecular Biology | Sakshi Shambhavi, Lamuk Zaveri, Shagufta Khan, Namami Gaur, Tulasi Nagabandi, Purushotham Vodnala, Payel Mukherjee, Sofia Banu, Priya Singh, Dhiviya Vedagiri, Divya Gupta, Vishal Sah, Santosh Kumar Kuncha, Krishnan Harinivas Harshan, Archana Bharadwaj Siva, Karthik Bharadwaj Tallapaka, Deepak Kumar, Devi Prasad Vijayashankar, Disha Nanda, Divya Das, Jotin Gogoi, Manish Bhattacharjee, Rakesh K Mishra, Divya Tej Sowpati |
| EPI_ISL_471600 | CSIR-Centre for Cellular and Molecular Biology | CSIR-Centre for Cellular and Molecular Biology | Sakshi Shambhavi, Lamuk Zaveri, Shagufta Khan, Namami Gaur, Tulasi Nagabandi, Purushotham Vodnala, Payel Mukherjee, Sofia Banu, Priya Singh, Dhiviya Vedagiri, Divya Gupta, Vishal Sah, Santosh Kumar Kuncha, Krishnan Harinivas Harshan, Archana Bharadwaj Siva, Karthik Bharadwaj Tallapaka, G. Aditya Kumar, Koushick Sivakumar, Pooja Ramesh Gupta, Rajan Kumar Jha, Shraddha Vijay Lahoti, Rakesh K Mishra, Divya Tej Sowpati |
| EPI_ISL_471601 | CSIR-Centre for Cellular and Molecular Biology | CSIR-Centre for Cellular and Molecular Biology | Sakshi Shambhavi, Lamuk Zaveri, Shagufta Khan, Namami Gaur, Tulasi Nagabandi, Purushotham Vodnala, Payel Mukherjee, Sofia Banu, Priya Singh, Dhiviya Vedagiri, Divya Gupta, Vishal Sah, Santosh Kumar Kuncha, Krishnan Harinivas Harshan, Archana Bharadwaj Siva, Karthik Bharadwaj Tallapaka,Nikhil Hajirnis, Pratheusa Maccha, M Soujanya Reddy,G. Aditya Kumar, Koushick Sivakumar, Rakesh K Mishra, Divya Tej Sowpati |
| EPI_ISL_471602 | CSIR-Centre for Cellular and Molecular Biology | CSIR-Centre for Cellular and Molecular Biology | Sakshi Shambhavi, Lamuk Zaveri, Shagufta Khan, Namami Gaur, Tulasi Nagabandi, Purushotham Vodnala, Payel Mukherjee, Sofia Banu, Priya Singh, Dhiviya Vedagiri, Divya Gupta, Vishal Sah, Santosh Kumar Kuncha, Krishnan Harinivas Harshan, Archana Bharadwaj Siva, Karthik Bharadwaj Tallapaka,Nikhil Hajirnis, Pratheusa Maccha, M Soujanya Reddy,G. Aditya Kumar, Koushick Sivakumar,Disha Nanda, Divya Das, Jotin Gogoi, Manish Bhattacharjee, Ravi Prasad Mukku, Rakesh K Mishra, Divya Tej Sowpati |
| EPI_ISL_471603 | CSIR-Centre for Cellular and Molecular Biology | CSIR-Centre for Cellular and Molecular Biology | Shagufta Khan, Lamuk Zaveri, Namami Gaur, Sakshi Shambhavi, Tulasi Nagabandi, Purushotham Vodnala, Payel Mukherjee, Sofia Banu, Priya Singh, Dhiviya Vedagiri, Divya Gupta, Vishal Sah, Santosh Kumar Kuncha, Krishnan Harinivas Harshan, Archana Bharadwaj Siva, Karthik Bharadwaj Tallapaka, Disha Nanda, Divya Das, Jotin Gogoi, Manish Bhattacharjee, Ravi Prasad Mukku, Rakesh K Mishra, Divya Tej Sowpati |
| EPI_ISL_471604 | CSIR-Centre for Cellular and Molecular Biology | CSIR-Centre for Cellular and Molecular Biology | Shagufta Khan, Lamuk Zaveri, Namami Gaur, Sakshi Shambhavi, Tulasi Nagabandi, Purushotham Vodnala, Payel Mukherjee, Sofia Banu, Priya Singh, Dhiviya Vedagiri, Divya Gupta, Vishal Sah, Santosh Kumar Kuncha, Krishnan Harinivas Harshan, Archana Bharadwaj Siva, Karthik Bharadwaj Tallapaka, Renu Sudhakar, Somesh Gorde, Gangumala Srinivas Reddy, Sujoy Deb, Swati Bayyana, Rakesh K Mishra, Divya Tej Sowpati |
| EPI_ISL_471605 | CSIR-Centre for Cellular and Molecular Biology | CSIR-Centre for Cellular and Molecular Biology | Shagufta Khan, Lamuk Zaveri, Namami Gaur, Sakshi Shambhavi, Tulasi Nagabandi, Purushotham Vodnala, Payel Mukherjee, Sofia Banu, Priya Singh, Dhiviya Vedagiri, Divya Gupta, Vishal Sah, Santosh Kumar Kuncha, Krishnan Harinivas Harshan, Archana Bharadwaj Siva, Karthik Bharadwaj Tallapaka,Preethi Jampala, Sharada Ravi Iyer, Sulagana Mukherjee, Swetha Sundar, Peddapuvala Sai Uday Kiran Rakesh K Mishra, Divya Tej Sowpati |
| EPI_ISL_471606 | CSIR-Centre for Cellular and Molecular Biology | CSIR-Centre for Cellular and Molecular Biology | Shagufta Khan, Lamuk Zaveri, Namami Gaur, Sakshi Shambhavi, Tulasi Nagabandi, Purushotham Vodnala, Payel Mukherjee, Sofia Banu, Priya Singh, Dhiviya Vedagiri, Divya Gupta, Vishal Sah, Santosh Kumar Kuncha, Krishnan Harinivas Harshan, Archana Bharadwaj Siva, Karthik Bharadwaj Tallapaka,Umesh Kumar, Unis Ahmad Bhat, Ajay Sarawagi, Priyanka Pant, Rajkanwar Nathawat, Rakesh K Mishra, Divya Tej Sowpati |
| EPI_ISL_471607 | CSIR-Centre for Cellular and Molecular Biology | CSIR-Centre for Cellular and Molecular Biology | Sofia Banu, Payel Mukherjee, Priya Singh, Dhiviya Vedagiri, Divya Gupta, Vishal Sah, Santosh Kumar Kuncha, Krishnan Harinivas Harshan, Archana Bharadwaj Siva, Karthik Bharadwaj Tallapaka, Shagufta Khan, Lamuk Zaveri, Namami Gaur, Sakshi Shambhavi, Tulasi Nagabandi, Purushotham Vodnala, Deepak Kumar, Devi Prasad Vijayashankar, Disha Nanda, Divya Das, Jotin Gogoi, Manish Bhattacharjee, Rakesh K Mishra, Divya Tej Sowpati |
| EPI_ISL_471608 | CSIR-Centre for Cellular and Molecular Biology | CSIR-Centre for Cellular and Molecular Biology | Sofia Banu, Payel Mukherjee, Priya Singh, Dhiviya Vedagiri, Divya Gupta, Vishal Sah, Santosh Kumar Kuncha, Krishnan Harinivas Harshan, Archana Bharadwaj Siva, Karthik Bharadwaj Tallapaka, Shagufta Khan, Lamuk Zaveri, Namami Gaur, Sakshi Shambhavi, Tulasi Nagabandi, Purushotham Vodnala, Disha Nanda, Divya Das, Jotin Gogoi, Manish Bhattacharjee, Ravi Prasad Mukku, Rakesh K Mishra, Divya Tej Sowpati |
| EPI_ISL_471609 | CSIR-Centre for Cellular and Molecular Biology | CSIR-Centre for Cellular and Molecular Biology | Sofia Banu, Payel Mukherjee, Priya Singh, Dhiviya Vedagiri, Divya Gupta, Vishal Sah, Santosh Kumar Kuncha, Krishnan Harinivas Harshan, Archana Bharadwaj Siva, Karthik Bharadwaj Tallapaka, Shagufta Khan, Lamuk Zaveri, Namami Gaur, Sakshi Shambhavi, Tulasi Nagabandi, Purushotham Vodnala, Gokulan C G, Gunjan Purohit, Hanuman Tulashiram Kale, Pankaj Kumar, Prachand Issarapu, Rakesh K Mishra, Divya Tej Sowpati |
| EPI_ISL_471610 | CSIR-Centre for Cellular and Molecular Biology | CSIR-Centre for Cellular and Molecular Biology | Sofia Banu, Payel Mukherjee, Priya Singh, Dhiviya Vedagiri, Divya Gupta, Vishal Sah, Santosh Kumar Kuncha, Krishnan Harinivas Harshan, Archana Bharadwaj Siva, Karthik Bharadwaj Tallapaka, Shagufta Khan, Lamuk Zaveri, Namami Gaur, Sakshi Shambhavi, Tulasi Nagabandi, Purushotham Vodnala,Preethi Jampala, Sharada Ravi Iyer, Sulagana Mukherjee, Swetha Sundar, Peddapuvala Sai Uday Kiran, Rakesh K Mishra, Divya Tej Sowpati |
| EPI_ISL_471611 | CSIR-Centre for Cellular and Molecular Biology | CSIR-Centre for Cellular and Molecular Biology | Tulasi Nagabandi, Namami Gaur, Sakshi Shambhavi, Lamuk Zaveri, Shagufta Khan, Purushotham Vodnala, Payel Mukherjee, Sofia Banu, Priya Singh, Dhiviya Vedagiri, Divya Gupta, Vishal Sah, Santosh Kumar Kuncha, Krishnan Harinivas Harshan, Archana Bharadwaj Siva, Karthik Bharadwaj Tallapaka,G. Aditya Kumar, Koushick Sivakumar, Pooja Ramesh Gupta, Rajan Kumar Jha, Shraddha Vijay Lahoti, Rakesh K Mishra, Divya Tej Sowpati |
| EPI_ISL_471612 | CSIR-Centre for Cellular and Molecular Biology | CSIR-Centre for Cellular and Molecular Biology | Tulasi Nagabandi, Namami Gaur, Sakshi Shambhavi, Lamuk Zaveri, Shagufta Khan, Purushotham Vodnala, Payel Mukherjee, Sofia Banu, Priya Singh, Dhiviya Vedagiri, Divya Gupta, Vishal Sah, Santosh Kumar Kuncha, Krishnan Harinivas Harshan, Archana Bharadwaj Siva, Karthik Bharadwaj Tallapaka,Kezia J Ann, Radhika Khandelwal, Roshan Maku Venkata, Shemin Mansuri, Sonu Uday, Rakesh K Mishra, Divya Tej Sowpati |
| EPI_ISL_471613 | CSIR-Centre for Cellular and Molecular Biology | CSIR-Centre for Cellular and Molecular Biology | Tulasi Nagabandi, Namami Gaur, Sakshi Shambhavi, Lamuk Zaveri, Shagufta Khan, Purushotham Vodnala, Payel Mukherjee, Sofia Banu, Priya Singh, Dhiviya Vedagiri, Divya Gupta, Vishal Sah, Santosh Kumar Kuncha, Krishnan Harinivas Harshan, Archana Bharadwaj Siva, Karthik Bharadwaj Tallapaka,G. Aditya Kumar, Koushick Sivakumar, Pooja Ramesh Gupta, Rajan Kumar Jha, Shraddha Vijay Lahoti, Rakesh K Mishra, Divya Tej Sowpati |
| EPI_ISL_471614 | CSIR-Centre for Cellular and Molecular Biology | CSIR-Centre for Cellular and Molecular Biology | Tulasi Nagabandi, Namami Gaur, Sakshi Shambhavi, Lamuk Zaveri, Shagufta Khan, Purushotham Vodnala, Payel Mukherjee, Sofia Banu, Priya Singh, Dhiviya Vedagiri, Divya Gupta, Vishal Sah, Santosh Kumar Kuncha, Krishnan Harinivas Harshan, Archana Bharadwaj Siva, Karthik Bharadwaj Tallapaka,Kezia J Ann, Radhika Khandelwal, Roshan Maku Venkata, Shemin Mansuri, Sonu Uday, Rakesh K Mishra, Divya Tej Sowpati |
| EPI_ISL_471615 | CSIR-Centre for Cellular and Molecular Biology | CSIR-Centre for Cellular and Molecular Biology | Lamuk Zaveri, Shagufta Khan, Namami Gaur, Sakshi Shambhavi, Tulasi Nagabandi, Purushotham Vodnala, Payel Mukherjee, Sofia Banu, Priya Singh, Dhiviya Vedagiri, Divya Gupta, Vishal Sah, Santosh Kumar Kuncha, Krishnan Harinivas Harshan, Archana Bharadwaj Siva, Karthik Bharadwaj Tallapaka,Zeba Rizvi, Zuberwasim Sayyad, Kakade Aishwarya Arun, Amrutha H C, Ananga Ghosh, Rakesh K Mishra, Divya Tej Sowpati |
| EPI_ISL_471616 | CSIR-Centre for Cellular and Molecular Biology | CSIR-Centre for Cellular and Molecular Biology | Lamuk Zaveri, Shagufta Khan, Namami Gaur, Sakshi Shambhavi, Tulasi Nagabandi, Purushotham Vodnala, Payel Mukherjee, Sofia Banu, Priya Singh, Dhiviya Vedagiri, Divya Gupta, Vishal Sah, Santosh Kumar Kuncha, Krishnan Harinivas Harshan, Archana Bharadwaj Siva, Karthik Bharadwaj Tallapaka, Renu Sudhakar, Somesh Gorde, Gangumala Srinivas Reddy, Sujoy Deb, Swati Bayyana, Rakesh K Mishra, Divya Tej Sowpati |
| EPI_ISL_471617 | CSIR-Centre for Cellular and Molecular Biology | CSIR-Centre for Cellular and Molecular Biology | Lamuk Zaveri, Shagufta Khan, Namami Gaur, Sakshi Shambhavi, Tulasi Nagabandi, Purushotham Vodnala, Payel Mukherjee, Sofia Banu, Priya Singh, Dhiviya Vedagiri, Divya Gupta, Vishal Sah, Santosh Kumar Kuncha, Krishnan Harinivas Harshan, Archana Bharadwaj Siva, Karthik Bharadwaj Tallapaka,Umesh Kumar, Unis Ahmad Bhat, Ajay Sarawagi, Priyanka Pant, Rajkanwar Nathawat, Rakesh K Mishra, Divya Tej Sowpati |
| EPI_ISL_471618 | CSIR-Centre for Cellular and Molecular Biology | CSIR-Centre for Cellular and Molecular Biology | Lamuk Zaveri, Shagufta Khan, Namami Gaur, Sakshi Shambhavi, Tulasi Nagabandi, Purushotham Vodnala, Payel Mukherjee, Sofia Banu, Priya Singh, Dhiviya Vedagiri, Divya Gupta, Vishal Sah, Santosh Kumar Kuncha, Krishnan Harinivas Harshan, Archana Bharadwaj Siva, Karthik Bharadwaj Tallapaka,Zeba Rizvi, Zuberwasim Sayyad, Kakade Aishwarya Arun, Amrutha H C, Ananga Ghosh, Rakesh K Mishra, Divya Tej Sowpati |
| EPI_ISL_471619 | CSIR-Centre for Cellular and Molecular Biology | CSIR-Centre for Cellular and Molecular Biology | Namami Gaur, Sakshi Shambhavi, Lamuk Zaveri, Shagufta Khan, Tulasi Nagabandi, Purushotham Vodnala, Payel Mukherjee, Sofia Banu, Priya Singh, Dhiviya Vedagiri, Divya Gupta, Vishal Sah, Santosh Kumar Kuncha, Krishnan Harinivas Harshan, Archana Bharadwaj Siva, Karthik Bharadwaj Tallapaka, Zeba Rizvi, Zuberwasim Sayyad, Kakade Aishwarya Arun, Amrutha H C, Ananga Ghosh, Rakesh K Mishra, Divya Tej Sowpati |
| EPI_ISL_471620 | CSIR-Centre for Cellular and Molecular Biology | CSIR-Centre for Cellular and Molecular Biology | Namami Gaur, Sakshi Shambhavi, Lamuk Zaveri, Shagufta Khan, Tulasi Nagabandi, Purushotham Vodnala, Payel Mukherjee, Sofia Banu, Priya Singh, Dhiviya Vedagiri, Divya Gupta, Vishal Sah, Santosh Kumar Kuncha, Krishnan Harinivas Harshan, Archana Bharadwaj Siva, Karthik Bharadwaj Tallapaka, Nikhil Hajirnis, Pratheusa Maccha, M Soujanya Reddy,G. Aditya Kumar, Koushick Sivakumar, Rakesh K Mishra, Divya Tej Sowpati |
| EPI_ISL_471621 | CSIR-Centre for Cellular and Molecular Biology | CSIR-Centre for Cellular and Molecular Biology | Namami Gaur, Sakshi Shambhavi, Lamuk Zaveri, Shagufta Khan, Tulasi Nagabandi, Purushotham Vodnala, Payel Mukherjee, Sofia Banu, Priya Singh, |

| | | | |
|---|---|---|---|
| | | | Dhiviya Vedagiri, Divya Gupta, Vishal Sah, Santosh Kumar Kuncha, Krishnan Harinivas Harshan, Archana Bharadwaj Siva, Karthik Bharadwaj Tallapaka, Zeba Rizvi, Zuberwasim Sayyad, Kakade Aishwarya Arun, Amrutha H C, Ananga Ghosh, Rakesh K Mishra, Divya Tej Sowpati |
| EPI_ISL_471622 | CSIR-Centre for Cellular and Molecular Biology | CSIR-Centre for Cellular and Molecular Biology | Namami Gaur, Sakshi Shambhavi, Lamuk Zaveri, Shagufta Khan, Tulasi Nagabandi, Purushotham Vodnala, Payel Mukherjee, Sofia Banu, Priya Singh, Dhiviya Vedagiri, Divya Gupta, Vishal Sah, Santosh Kumar Kuncha, Krishnan Harinivas Harshan, Archana Bharadwaj Siva, Karthik Bharadwaj Tallapaka,Kezia J Ann, Radhika Khandelwal, Roshan Maku Venkata, Shemin Mansuri, Sonu Uday, Rakesh K Mishra, Divya Tej Sowpati |
| EPI_ISL_471623 | CSIR-Centre for Cellular and Molecular Biology | CSIR-Centre for Cellular and Molecular Biology | Payel Mukherjee, Sofia Banu, Priya Singh, Dhiviya Vedagiri, Divya Gupta, Vishal Sah, Santosh Kumar Kuncha, Krishnan Harinivas Harshan, Archana Bharadwaj Siva, Karthik Bharadwaj Tallapaka, Shagufta Khan, Lamuk Zaveri, Namami Gaur, Sakshi Shambhavi, Tulasi Nagabandi, Purushotham Vodnala, G. Aditya Kumar, Koushick Sivakumar, Pooja Ramesh Gupta, Rajan Kumar Jha, Shraddha Vijay Lahoti, Rakesh K Mishra, Divya Tej Sowpati |
| EPI_ISL_471624 | CSIR-Centre for Cellular and Molecular Biology | CSIR-Centre for Cellular and Molecular Biology | Payel Mukherjee, Sofia Banu, Priya Singh, Dhiviya Vedagiri, Divya Gupta, Vishal Sah, Santosh Kumar Kuncha, Krishnan Harinivas Harshan, Archana Bharadwaj Siva, Karthik Bharadwaj Tallapaka, Shagufta Khan, Lamuk Zaveri, Namami Gaur, Sakshi Shambhavi, Tulasi Nagabandi, Purushotham Vodnala, Gokulan C G, Gunjan Purohit, Hanuman Tulashiram Kale, Pankaj Kumar, Prachand Issarapu, Rakesh K Mishra, Divya Tej Sowpati |
| EPI_ISL_471625 | CSIR-Centre for Cellular and Molecular Biology | CSIR-Centre for Cellular and Molecular Biology | Payel Mukherjee, Sofia Banu, Priya Singh, Dhiviya Vedagiri, Divya Gupta, Vishal Sah, Santosh Kumar Kuncha, Krishnan Harinivas Harshan, Archana Bharadwaj Siva, Karthik Bharadwaj Tallapaka, Shagufta Khan, Lamuk Zaveri, Namami Gaur, Sakshi Shambhavi, Tulasi Nagabandi, Purushotham Vodnala, Rakesh K Mishra, Sonu Uday, Sudipta Mondal, Annapoorna P Karthyayani, Debabrata Jana, Debrya Saha, Divya Tej Sowpati |
| EPI_ISL_471626 | CSIR-Centre for Cellular and Molecular Biology | CSIR-Centre for Cellular and Molecular Biology | Payel Mukherjee, Sofia Banu, Priya Singh, Dhiviya Vedagiri, Divya Gupta, Vishal Sah, Santosh Kumar Kuncha, Krishnan Harinivas Harshan, Archana Bharadwaj Siva, Karthik Bharadwaj Tallapaka, Shagufta Khan, Lamuk Zaveri, Namami Gaur, Sakshi Shambhavi, Tulasi Nagabandi, Purushotham Vodnala,Deepak Kumar, Devi Prasad Vijayashankar, Disha Nanda, Divya Das, Jotin Gogoi, Manish Bhattacharjee, Rakesh K Mishra, Divya Tej Sowpati |
| EPI_ISL_471627 | CSIR-Centre for Cellular and Molecular Biology | CSIR-Centre for Cellular and Molecular Biology | Sakshi Shambhavi, Lamuk Zaveri, Shagufta Khan, Namami Gaur, Tulasi Nagabandi, Purushotham Vodnala, Payel Mukherjee, Sofia Banu, Priya Singh, Dhiviya Vedagiri, Divya Gupta, Vishal Sah, Santosh Kumar Kuncha, Krishnan Harinivas Harshan, Archana Bharadwaj Siva, Karthik Bharadwaj Tallapaka, Deepak Kumar, Devi Prasad Vijayashankar, Disha Nanda, Divya Das, Jotin Gogoi, Manish Bhattacharjee, Rakesh K Mishra, Divya Tej Sowpati |
| EPI_ISL_471628 | CSIR-Centre for Cellular and Molecular Biology | CSIR-Centre for Cellular and Molecular Biology | Sakshi Shambhavi, Lamuk Zaveri, Shagufta Khan, Namami Gaur, Tulasi Nagabandi, Purushotham Vodnala, Payel Mukherjee, Sofia Banu, Priya Singh, Dhiviya Vedagiri, Divya Gupta, Vishal Sah, Santosh Kumar Kuncha, Krishnan Harinivas Harshan, Archana Bharadwaj Siva, Karthik Bharadwaj Tallapaka, G. Aditya Kumar, Koushick Sivakumar, Pooja Ramesh Gupta, Rajan Kumar Jha, Shraddha Vijay Lahoti, Rakesh K Mishra, Divya Tej Sowpati |
| EPI_ISL_471629 | CSIR-Centre for Cellular and Molecular Biology | CSIR-Centre for Cellular and Molecular Biology | Sakshi Shambhavi, Lamuk Zaveri, Shagufta Khan, Namami Gaur, Tulasi Nagabandi, Purushotham Vodnala, Payel Mukherjee, Sofia Banu, Priya Singh, Dhiviya Vedagiri, Divya Gupta, Vishal Sah, Santosh Kumar Kuncha, Krishnan Harinivas Harshan, Archana Bharadwaj Siva, Karthik Bharadwaj Tallapaka,Nikhil Hajirnis, Pratheusa Maccha, M Soujanya Reddy,G. Aditya Kumar, Koushick Sivakumar, Rakesh K Mishra, Divya Tej Sowpati |
| EPI_ISL_471630 | CSIR-Centre for Cellular and Molecular Biology | CSIR-Centre for Cellular and Molecular Biology | Sakshi Shambhavi, Lamuk Zaveri, Shagufta Khan, Namami Gaur, Tulasi Nagabandi, Purushotham Vodnala, Payel Mukherjee, Sofia Banu, Priya Singh, Dhiviya Vedagiri, Divya Gupta, Vishal Sah, Santosh Kumar Kuncha, Krishnan Harinivas Harshan, Archana Bharadwaj Siva, Karthik Bharadwaj Tallapaka,Nikhil Hajirnis, Pratheusa Maccha, M Soujanya Reddy,G. Aditya Kumar, Koushick Sivakumar,Disha Nanda, Divya Das, Jotin Gogoi, Manish Bhattacharjee, Ravi Prasad Mukku, Rakesh K Mishra, Divya Tej Sowpati |
| EPI_ISL_471631 | CSIR-Centre for Cellular and Molecular Biology | CSIR-Centre for Cellular and Molecular Biology | Shagufta Khan, Lamuk Zaveri, Namami Gaur, Sakshi Shambhavi, Tulasi Nagabandi, Purushotham Vodnala, Payel Mukherjee, Sofia Banu, Priya Singh, Dhiviya Vedagiri, Divya Gupta, Vishal Sah, Santosh Kumar Kuncha, Krishnan Harinivas Harshan, Archana Bharadwaj Siva, Karthik Bharadwaj Tallapaka, Disha Nanda, Divya Das, Jotin Gogoi, Manish Bhattacharjee, Ravi Prasad Mukku, Rakesh K Mishra, Divya Tej Sowpati |
| EPI_ISL_471632 | CSIR-Centre for Cellular and Molecular Biology | CSIR-Centre for Cellular and Molecular Biology | Shagufta Khan, Lamuk Zaveri, Namami Gaur, Sakshi Shambhavi, Tulasi Nagabandi, Purushotham Vodnala, Payel Mukherjee, Sofia Banu, Priya Singh, Dhiviya Vedagiri, Divya Gupta, Vishal Sah, Santosh Kumar Kuncha, Krishnan Harinivas Harshan, Archana Bharadwaj Siva, Karthik Bharadwaj Tallapaka, Renu Sudhakar, Somesh Gorde, Gangumala Srinivas Reddy, Sujoy Deb, Swati Bayyana, Rakesh K Mishra, Divya Tej Sowpati |
| EPI_ISL_471633 | CSIR-Centre for Cellular and Molecular Biology | CSIR-Centre for Cellular and Molecular Biology | Shagufta Khan, Lamuk Zaveri, Namami Gaur, Sakshi Shambhavi, Tulasi Nagabandi, Purushotham Vodnala, Payel Mukherjee, Sofia Banu, Priya Singh, Dhiviya Vedagiri, Divya Gupta, Vishal Sah, Santosh Kumar Kuncha, Krishnan Harinivas Harshan, Archana Bharadwaj Siva, Karthik Bharadwaj Tallapaka,Preethi Jampala, Sharada Ravi Iyer, Sulagana Mukherjee, Swetha Sundar, Peddapuvala Sai Uday Kiran Rakesh K Mishra, Divya Tej Sowpati |
| EPI_ISL_471634 | CSIR-Centre for Cellular and Molecular Biology | CSIR-Centre for Cellular and Molecular Biology | Shagufta Khan, Lamuk Zaveri, Namami Gaur, Sakshi Shambhavi, Tulasi Nagabandi, Purushotham Vodnala, Payel Mukherjee, Sofia Banu, Priya Singh, Dhiviya Vedagiri, Divya Gupta, Vishal Sah, Santosh Kumar Kuncha, Krishnan Harinivas Harshan, Archana Bharadwaj Siva, Karthik Bharadwaj Tallapaka,Umesh Kumar, Unis Ahmad Bhat, Ajay Sarawagi, Priyanka Pant, Rajkanwar Nathawat, Rakesh K Mishra, Divya Tej Sowpati |
| EPI_ISL_471635 | CSIR-Centre for Cellular and Molecular Biology | CSIR-Centre for Cellular and Molecular Biology | Sofia Banu, Payel Mukherjee, Priya Singh, Dhiviya Vedagiri, Divya Gupta, Vishal Sah, Santosh Kumar Kuncha, Krishnan Harinivas Harshan, Archana Bharadwaj Siva, Karthik Bharadwaj Tallapaka, Shagufta Khan, Lamuk Zaveri, Namami Gaur, Sakshi Shambhavi, Tulasi Nagabandi, Purushotham Vodnala, Deepak Kumar, Devi Prasad Vijayashankar, Disha Nanda, Divya Das, Jotin Gogoi, Manish Bhattacharjee, Rakesh K Mishra, Divya Tej Sowpati |
| EPI_ISL_471636 | CSIR-Centre for Cellular and Molecular Biology | CSIR-Centre for Cellular and Molecular Biology | Sofia Banu, Payel Mukherjee, Priya Singh, Dhiviya Vedagiri, Divya Gupta, Vishal Sah, Santosh Kumar Kuncha, Krishnan Harinivas Harshan, Archana Bharadwaj Siva, Karthik Bharadwaj Tallapaka, Shagufta Khan, Lamuk Zaveri, Namami Gaur, Sakshi Shambhavi, Tulasi Nagabandi, Purushotham Vodnala, Disha Nanda, Divya Das, Jotin Gogoi, Manish Bhattacharjee, Ravi Prasad Mukku, Rakesh K Mishra, Divya Tej Sowpati |
| EPI_ISL_471637 | CSIR-Centre for Cellular and Molecular Biology | CSIR-Centre for Cellular and Molecular Biology | Sofia Banu, Payel Mukherjee, Priya Singh, Dhiviya Vedagiri, Divya Gupta, Vishal Sah, Santosh Kumar Kuncha, Krishnan Harinivas Harshan, Archana Bharadwaj Siva, Karthik Bharadwaj Tallapaka, Shagufta Khan, Lamuk Zaveri, Namami Gaur, Sakshi Shambhavi, Tulasi Nagabandi, Purushotham Vodnala, Gokulan C G, Gunjan Purohit, Hanuman Tulashiram Kale, Pankaj Kumar, Prachand Issarapu, Rakesh K Mishra, Divya Tej Sowpati |
| EPI_ISL_471638 | CSIR-Centre for Cellular and Molecular Biology | CSIR-Centre for Cellular and Molecular Biology | Sofia Banu, Payel Mukherjee, Priya Singh, Dhiviya Vedagiri, Divya Gupta, Vishal Sah, Santosh Kumar Kuncha, Krishnan Harinivas Harshan, Archana Bharadwaj Siva, Karthik Bharadwaj Tallapaka, Shagufta Khan, Lamuk Zaveri, Namami Gaur, Sakshi Shambhavi, Tulasi Nagabandi, Purushotham Vodnala,Preethi Jampala, Sharada Ravi Iyer, Sulagana Mukherjee, Swetha Sundar, Peddapuvala Sai Uday Kiran, Rakesh K Mishra, Divya Tej Sowpati |
| EPI_ISL_471639 | CSIR-Centre for Cellular and Molecular Biology | CSIR-Centre for Cellular and Molecular Biology | Tulasi Nagabandi, Namami Gaur, Sakshi Shambhavi, Lamuk Zaveri, Shagufta Khan, Purushotham Vodnala, Payel Mukherjee, Sofia Banu, Priya Singh, Dhiviya Vedagiri, Divya Gupta, Vishal Sah, Santosh Kumar Kuncha, Krishnan Harinivas Harshan, Archana Bharadwaj Siva, Karthik Bharadwaj Tallapaka,G. Aditya Kumar, Koushick Sivakumar, Pooja Ramesh Gupta, Rajan Kumar Jha, Shraddha Vijay Lahoti, Rakesh K Mishra, Divya Tej Sowpati |
| EPI_ISL_471640 | CSIR-Centre for Cellular and Molecular Biology | CSIR-Centre for Cellular and Molecular Biology | Tulasi Nagabandi, Namami Gaur, Sakshi Shambhavi, Lamuk Zaveri, Shagufta Khan, Purushotham Vodnala, Payel Mukherjee, Sofia Banu, Priya Singh, Dhiviya Vedagiri, Divya Gupta, Vishal Sah, Santosh Kumar Kuncha, Krishnan Harinivas Harshan, Archana Bharadwaj Siva, Karthik Bharadwaj Tallapaka,Kezia J Ann, Radhika Khandelwal, Roshan Maku Venkata, Shemin Mansuri, Sonu Uday, Rakesh K Mishra, Divya Tej Sowpati |
| EPI_ISL_471641, EPI_ISL_471642 | CSIR-Centre for Cellular and Molecular Biology | CSIR-Centre for Cellular and Molecular Biology | Dhiviya Vedagiri, Divya Gupta, Vishal Sah, Payel Mukherjee, Sofia Banu, Priya Singh, Santosh Kumar Kuncha, Archana Bharadwaj Siva, Karthik Bharadwaj Tallapaka, Shagufta Khan, Lamuk Zaveri, Namami Gaur, Sakshi Shambhavi, Tulasi Nagabandi, Purushotham Vodnala, Rakesh K Mishra, Divya Tej Sowpati, Krishnan Harinivas Harshan |
| EPI_ISL_471643 | CSIR-Centre for Cellular and Molecular Biology | CSIR-Centre for Cellular and Molecular Biology | Tulasi Nagabandi, Namami Gaur, Sakshi Shambhavi, Lamuk Zaveri, Shagufta Khan, Purushotham Vodnala, Payel Mukherjee, Sofia Banu, Priya Singh, Dhiviya Vedagiri, Divya Gupta, Vishal Sah, Santosh Kumar Kuncha, Krishnan Harinivas Harshan, Archana Bharadwaj Siva, Karthik Bharadwaj Tallapaka,G. Aditya Kumar, Koushick Sivakumar, Pooja Ramesh Gupta, Rajan Kumar Jha, Shraddha Vijay Lahoti, Rakesh K Mishra, Divya Tej Sowpati |
| EPI_ISL_471644 | CSIR-Centre for Cellular and Molecular Biology | CSIR-Centre for Cellular and Molecular Biology | Tulasi Nagabandi, Namami Gaur, Sakshi Shambhavi, Lamuk Zaveri, Shagufta Khan, Purushotham Vodnala, Payel Mukherjee, Sofia Banu, Priya Singh, Dhiviya Vedagiri, Divya Gupta, Vishal Sah, Santosh Kumar Kuncha, Krishnan Harinivas Harshan, Archana Bharadwaj Siva, Karthik Bharadwaj Tallapaka,Kezia J Ann, Radhika Khandelwal, Roshan Maku Venkata, Shemin Mansuri, Sonu Uday, Rakesh K Mishra, Divya Tej Sowpati |
| EPI_ISL_471645, EPI_ISL_471646 | CSIR-Centre for Cellular and Molecular Biology | CSIR-Centre for Cellular and Molecular Biology | Dhiviya Vedagiri, Divya Gupta, Vishal Sah, Payel Mukherjee, Sofia Banu, Priya Singh, Santosh Kumar Kuncha, Archana Bharadwaj Siva, Karthik Bharadwaj Tallapaka, Shagufta Khan, Lamuk Zaveri, Namami Gaur, Sakshi Shambhavi, Tulasi Nagabandi, Purushotham Vodnala, Rakesh K Mishra, Divya Tej Sowpati, Krishnan Harinivas Harshan |
| EPI_ISL_471647 | Hospital Municipal de Barueri Dr. Francisco Moran | Instituto Adolfo Lutz, Interdiciplinary Procedures Center, Strategic Laboratory | Claudio Tavares Sacchi, Claudia Regina Gonçalves, Erica Valessa Ramos Gomes |
| EPI_ISL_471648 | UBS e Pronto Socorro Jd. Jacira | Instituto Adolfo Lutz, Interdiciplinary Procedures Center, | Claudio Tavares Sacchi, Claudia Regina Gonçalves, Erica Valessa Ramos Gomes |

| | | | |
|---|---|---|---|
| | | Strategic Laboratory | |
| EPI_ISL_471676, EPI_ISL_471677, EPI_ISL_471678, EPI_ISL_471679, EPI_ISL_471680, EPI_ISL_471681, EPI_ISL_471682, EPI_ISL_471683, EPI_ISL_471684, EPI_ISL_471685, EPI_ISL_471686, EPI_ISL_471687, EPI_ISL_471688, EPI_ISL_471689, EPI_ISL_471690, EPI_ISL_471691, EPI_ISL_471692, EPI_ISL_471693, EPI_ISL_471694, EPI_ISL_471695, EPI_ISL_471696, EPI_ISL_471697, EPI_ISL_471698, EPI_ISL_471699, EPI_ISL_471700, EPI_ISL_471701, EPI_ISL_471702, EPI_ISL_471703, EPI_ISL_471704, EPI_ISL_471705, EPI_ISL_471706, EPI_ISL_471707, EPI_ISL_471708, EPI_ISL_471709, EPI_ISL_471710, EPI_ISL_471711, EPI_ISL_471712, EPI_ISL_471713, EPI_ISL_471714, EPI_ISL_471715, EPI_ISL_471716, EPI_ISL_471717, EPI_ISL_471718, EPI_ISL_471719, EPI_ISL_471720, EPI_ISL_471721, EPI_ISL_471722, EPI_ISL_471723, EPI_ISL_471724, EPI_ISL_471725, EPI_ISL_471726, EPI_ISL_471727, EPI_ISL_471728, EPI_ISL_471729, EPI_ISL_471730, EPI_ISL_471731, EPI_ISL_471732, EPI_ISL_471733, EPI_ISL_471734, EPI_ISL_471735, EPI_ISL_471736, EPI_ISL_471737, EPI_ISL_471738, EPI_ISL_471739, EPI_ISL_471740, EPI_ISL_471741, EPI_ISL_471742, EPI_ISL_471743, EPI_ISL_471744, EPI_ISL_471745, EPI_ISL_471746, EPI_ISL_471747, EPI_ISL_471748, EPI_ISL_471749, EPI_ISL_471750, EPI_ISL_471751, EPI_ISL_471752, EPI_ISL_471753, EPI_ISL_471754, EPI_ISL_471755, EPI_ISL_471756, EPI_ISL_471757, EPI_ISL_471758, EPI_ISL_471759, EPI_ISL_471760, EPI_ISL_471761, EPI_ISL_471762, EPI_ISL_471763, EPI_ISL_471764, EPI_ISL_471765, EPI_ISL_471766, EPI_ISL_471767, EPI_ISL_471768, EPI_ISL_471769, EPI_ISL_471770, EPI_ISL_471771, EPI_ISL_471772, EPI_ISL_471773, EPI_ISL_471774, EPI_ISL_471775, EPI_ISL_471776, EPI_ISL_471777, EPI_ISL_471778, EPI_ISL_471779, EPI_ISL_471780, EPI_ISL_471781, EPI_ISL_471782, EPI_ISL_471783, EPI_ISL_471784, EPI_ISL_471785, EPI_ISL_471786, EPI_ISL_471787, EPI_ISL_471788, EPI_ISL_471789, EPI_ISL_471790, EPI_ISL_471791, EPI_ISL_471792, EPI_ISL_471793, EPI_ISL_471794, EPI_ISL_471795, EPI_ISL_471796, EPI_ISL_471797, EPI_ISL_471798, EPI_ISL_471799, EPI_ISL_471800, EPI_ISL_471801, EPI_ISL_471802, EPI_ISL_471803, EPI_ISL_471804, EPI_ISL_471805, EPI_ISL_471806, EPI_ISL_471807, EPI_ISL_471808, EPI_ISL_471809, EPI_ISL_471810, EPI_ISL_471811, EPI_ISL_471812, EPI_ISL_471813, EPI_ISL_471814, EPI_ISL_471815, EPI_ISL_471816, EPI_ISL_471817, EPI_ISL_471818, EPI_ISL_471819, EPI_ISL_471820, EPI_ISL_471821, EPI_ISL_471822, EPI_ISL_471823, EPI_ISL_471824, EPI_ISL_471825, EPI_ISL_471826, EPI_ISL_471827, EPI_ISL_471828, EPI_ISL_471829, EPI_ISL_471830, EPI_ISL_471831, EPI_ISL_471832, EPI_ISL_471833, EPI_ISL_471834, EPI_ISL_471835, EPI_ISL_471836, EPI_ISL_471837, EPI_ISL_471838, EPI_ISL_471839, EPI_ISL_471840, EPI_ISL_471841, EPI_ISL_471842, EPI_ISL_471843, EPI_ISL_471844, EPI_ISL_471845, EPI_ISL_471846, EPI_ISL_471847, EPI_ISL_471848, EPI_ISL_471849, EPI_ISL_471850, EPI_ISL_471851, EPI_ISL_471852, EPI_ISL_471853, EPI_ISL_471854, EPI_ISL_471855, EPI_ISL_471856, EPI_ISL_471857, EPI_ISL_471858, EPI_ISL_471859, EPI_ISL_471860, EPI_ISL_471861, EPI_ISL_471862, EPI_ISL_471863, EPI_ISL_471864, EPI_ISL_471865, EPI_ISL_471866, EPI_ISL_471867, EPI_ISL_471868, EPI_ISL_471869, EPI_ISL_471870, EPI_ISL_471871, EPI_ISL_471872, EPI_ISL_471873, EPI_ISL_471874, EPI_ISL_471875, EPI_ISL_471876, EPI_ISL_471877, EPI_ISL_471878, EPI_ISL_471879, EPI_ISL_471880, EPI_ISL_471881, EPI_ISL_471882, EPI_ISL_471883, EPI_ISL_471884, EPI_ISL_471885, EPI_ISL_471886, EPI_ISL_471887, EPI_ISL_471888, EPI_ISL_471889, EPI_ISL_471890, EPI_ISL_471891, EPI_ISL_471892, EPI_ISL_471893, EPI_ISL_471894, EPI_ISL_471895, EPI_ISL_471896, EPI_ISL_471897, EPI_ISL_471898, EPI_ISL_471899, EPI_ISL_471900, EPI_ISL_471901, EPI_ISL_471902, EPI_ISL_471903, EPI_ISL_471904, EPI_ISL_471905, EPI_ISL_471906, EPI_ISL_471907, EPI_ISL_471908, EPI_ISL_471909, EPI_ISL_471910 | | | |
| see above | Michigan Department of Health and Human Services, Bureau of Laboratories | Michigan Department of Health and Human Services, Bureau of Laboratories | Blankenship HM, Riner D, Soehnlen MK |
| EPI_ISL_471911, EPI_ISL_471912, EPI_ISL_471913, EPI_ISL_471914, EPI_ISL_471915, EPI_ISL_471916, EPI_ISL_471917, EPI_ISL_471918, EPI_ISL_471919, EPI_ISL_471920, EPI_ISL_471921, EPI_ISL_471922, EPI_ISL_471923, EPI_ISL_471924, EPI_ISL_471925, EPI_ISL_471926, EPI_ISL_471927, EPI_ISL_471928, EPI_ISL_471929, EPI_ISL_471930, EPI_ISL_471931, EPI_ISL_471932, EPI_ISL_471933, EPI_ISL_471934, EPI_ISL_471935, EPI_ISL_471936, EPI_ISL_471937, EPI_ISL_471938, EPI_ISL_471939, EPI_ISL_471940, EPI_ISL_471941, EPI_ISL_471942, EPI_ISL_471943, EPI_ISL_471944, EPI_ISL_471945, EPI_ISL_471946, EPI_ISL_471947, EPI_ISL_471948, EPI_ISL_471949, EPI_ISL_471950, EPI_ISL_471951, EPI_ISL_471952, EPI_ISL_471953, EPI_ISL_471954, EPI_ISL_471955, EPI_ISL_471956, EPI_ISL_471957, EPI_ISL_471958, EPI_ISL_471959, EPI_ISL_471960, EPI_ISL_471961, EPI_ISL_471962, EPI_ISL_471963, EPI_ISL_471964, EPI_ISL_471965, EPI_ISL_471966, EPI_ISL_471967, EPI_ISL_471968, EPI_ISL_471969, EPI_ISL_471970, EPI_ISL_471971, EPI_ISL_471972, EPI_ISL_471973, EPI_ISL_471974, EPI_ISL_471975, EPI_ISL_471976, EPI_ISL_471977, EPI_ISL_471978, EPI_ISL_471979, EPI_ISL_471980, EPI_ISL_471981, EPI_ISL_471982, EPI_ISL_471983, EPI_ISL_471984, EPI_ISL_471985, EPI_ISL_471986, EPI_ISL_471987, EPI_ISL_471988, EPI_ISL_471989, EPI_ISL_471990, EPI_ISL_471991, EPI_ISL_471992 | | | |
| see above | University of Exeter | COVID-19 Genomics UK (COG-UK) Consortium | Ben Temperton,Aaron Jeffries,Michelle Michelsen,Joanna Warwick-Dugdale,Audrey Farbos,Robyn Manley,Stephen Michell,Jane Masoli |
| EPI_ISL_471993, EPI_ISL_471994, EPI_ISL_471995, EPI_ISL_471996, EPI_ISL_471997, EPI_ISL_471998, EPI_ISL_471999, EPI_ISL_472000, EPI_ISL_472001, EPI_ISL_472002, EPI_ISL_472003, EPI_ISL_472004, EPI_ISL_472005, EPI_ISL_472006, EPI_ISL_472007, EPI_ISL_472008, EPI_ISL_472009, EPI_ISL_472010, EPI_ISL_472011, EPI_ISL_472012, EPI_ISL_472013, EPI_ISL_472014, EPI_ISL_472015, EPI_ISL_472016, EPI_ISL_472017, EPI_ISL_472018, EPI_ISL_472019, EPI_ISL_472020, EPI_ISL_472021, EPI_ISL_472022, EPI_ISL_472023, EPI_ISL_472024, EPI_ISL_472025, EPI_ISL_472026, EPI_ISL_472027, EPI_ISL_472028, EPI_ISL_472029, EPI_ISL_472030, EPI_ISL_472031, EPI_ISL_472032, EPI_ISL_472033, EPI_ISL_472034, EPI_ISL_472035, EPI_ISL_472036, EPI_ISL_472037, EPI_ISL_472038, EPI_ISL_472039, EPI_ISL_472040, EPI_ISL_472041, EPI_ISL_472042, EPI_ISL_472043, EPI_ISL_472044, EPI_ISL_472045, EPI_ISL_472046, EPI_ISL_472047, EPI_ISL_472048, EPI_ISL_472049, EPI_ISL_472050, EPI_ISL_472051, EPI_ISL_472052, EPI_ISL_472053, EPI_ISL_472054, EPI_ISL_472055, EPI_ISL_472056, EPI_ISL_472057, EPI_ISL_472058, EPI_ISL_472059, EPI_ISL_472060, EPI_ISL_472061, EPI_ISL_472062, EPI_ISL_472063, EPI_ISL_472064, EPI_ISL_472065, EPI_ISL_472066, EPI_ISL_472067, EPI_ISL_472068, EPI_ISL_472069, EPI_ISL_472070, EPI_ISL_472071, EPI_ISL_472072, EPI_ISL_472073, EPI_ISL_472074, EPI_ISL_472075, EPI_ISL_472076, EPI_ISL_472077, EPI_ISL_472078, EPI_ISL_472079, EPI_ISL_472080, EPI_ISL_472081, EPI_ISL_472082, EPI_ISL_472083, EPI_ISL_472084, EPI_ISL_472085, EPI_ISL_472086, EPI_ISL_472087, EPI_ISL_472088, EPI_ISL_472089, EPI_ISL_472090, EPI_ISL_472091, EPI_ISL_472092, EPI_ISL_472093, EPI_ISL_472094, EPI_ISL_472095, EPI_ISL_472096, EPI_ISL_472097, EPI_ISL_472098, EPI_ISL_472099, EPI_ISL_472100, EPI_ISL_472101, EPI_ISL_472102, EPI_ISL_472103, EPI_ISL_472104, EPI_ISL_472105, EPI_ISL_472106, EPI_ISL_472107, EPI_ISL_472108, EPI_ISL_472109, EPI_ISL_472110, EPI_ISL_472111, EPI_ISL_472112, EPI_ISL_472113, EPI_ISL_472114, EPI_ISL_472115, EPI_ISL_472116, EPI_ISL_472117, EPI_ISL_472118, EPI_ISL_472119, EPI_ISL_472120, EPI_ISL_472121, EPI_ISL_472122, EPI_ISL_472123, EPI_ISL_472124, EPI_ISL_472125, EPI_ISL_472126, EPI_ISL_472127, EPI_ISL_472128, EPI_ISL_472129, EPI_ISL_472130, EPI_ISL_472131, EPI_ISL_472132, EPI_ISL_472133, EPI_ISL_472134, EPI_ISL_472135 | | | |
| see above | Liverpool Clinical Laboratories | COVID-19 Genomics UK (COG-UK) Consortium | Sam Haldenby, Anita Lucaci, Steve Paterson, Julian Hiscox, Alistair Darby, M Almsaud, A Alrezaihi, Muhannad Alruwaili, Stuart D Armstrong, Jones Benjamin, Eleanor G Bentley, Anu Chawla, Jordan J Clark, Angela Cowell, Richard Eccles, Isabel García-Dorival, Matthew Gemmell, Alessandro Gerada, PKF Gilmore, Richard Gregory, Ximeng Han, Catherine Hartley, Margaret Hughes, Miren Iturriza-Gomara, James Johnson, L Luu, Jenifer Manson, Charlotte Nelson, Elaine O'Toole, Cassie Olateju, Rebekah Penrice-Randal , Lucille Rainbow, N.P Randle, Trevor Ian Robinson, Parul Sharma, Ghada T Shawli, James P Stewart, Neil Swainston, Ecaterina Vamos, Joanne Watts, Mark Whitehead |
| EPI_ISL_472136, EPI_ISL_472137, EPI_ISL_472138, EPI_ISL_472139, EPI_ISL_472140, EPI_ISL_472141, EPI_ISL_472142, EPI_ISL_472143, EPI_ISL_472144, EPI_ISL_472145, EPI_ISL_472146, EPI_ISL_472147, EPI_ISL_472148, EPI_ISL_472149, EPI_ISL_472150, EPI_ISL_472151, EPI_ISL_472152, EPI_ISL_472153, EPI_ISL_472154 | | | |
| see above | Regional Virus Laboratory, Belfast Health and Social Care Trust | COVID-19 Genomics UK (COG-UK) Consortium | Conall McCaughey, James McKenna, Tanya Curran, Susan Feeney, Alison Watt, Ciara Cox, Mairead Connor, Zoltan Molnar, David Simpson, Derek Fairley |
| EPI_ISL_472155, EPI_ISL_472156, EPI_ISL_472157, EPI_ISL_472158, EPI_ISL_472159, EPI_ISL_472160, EPI_ISL_472161, EPI_ISL_472162, EPI_ISL_472163, EPI_ISL_472164, EPI_ISL_472165, EPI_ISL_472166, EPI_ISL_472167, EPI_ISL_472168, EPI_ISL_472169, EPI_ISL_472170, EPI_ISL_472171, EPI_ISL_472172, EPI_ISL_472173, EPI_ISL_472174, EPI_ISL_472175, EPI_ISL_472176, EPI_ISL_472177, EPI_ISL_472178, EPI_ISL_472179, EPI_ISL_472180, EPI_ISL_472181, EPI_ISL_472182, EPI_ISL_472183, EPI_ISL_472184, EPI_ISL_472185, EPI_ISL_472186, EPI_ISL_472187, EPI_ISL_472188, EPI_ISL_472189, EPI_ISL_472190, EPI_ISL_472191, EPI_ISL_472192, EPI_ISL_472193, EPI_ISL_472194, EPI_ISL_472195, EPI_ISL_472196, EPI_ISL_472197, EPI_ISL_472198, EPI_ISL_472199, EPI_ISL_472200, EPI_ISL_472201, EPI_ISL_472202, EPI_ISL_472203, EPI_ISL_472204, EPI_ISL_472205, EPI_ISL_472206, EPI_ISL_472207, EPI_ISL_472208, EPI_ISL_472209, EPI_ISL_472210, EPI_ISL_472211, EPI_ISL_472212, EPI_ISL_472213, EPI_ISL_472214, EPI_ISL_472215, EPI_ISL_472216, EPI_ISL_472217, EPI_ISL_472218, EPI_ISL_472219, EPI_ISL_472220, EPI_ISL_472221, EPI_ISL_472222, EPI_ISL_472223, EPI_ISL_472224, EPI_ISL_472225, EPI_ISL_472226, EPI_ISL_472227, EPI_ISL_472228, EPI_ISL_472229, EPI_ISL_472230, EPI_ISL_472231, EPI_ISL_472232, EPI_ISL_472233, EPI_ISL_472234, EPI_ISL_472235, EPI_ISL_472236, EPI_ISL_472237, EPI_ISL_472238, EPI_ISL_472239, EPI_ISL_472240, EPI_ISL_472241, EPI_ISL_472242, EPI_ISL_472243, EPI_ISL_472244, EPI_ISL_472245, EPI_ISL_472246, EPI_ISL_472247, EPI_ISL_472248, EPI_ISL_472249, EPI_ISL_472250, EPI_ISL_472251, EPI_ISL_472252, EPI_ISL_472253, EPI_ISL_472254, EPI_ISL_472255, EPI_ISL_472256, EPI_ISL_472257, EPI_ISL_472258, EPI_ISL_472259, EPI_ISL_472260, EPI_ISL_472261, EPI_ISL_472262, EPI_ISL_472263, EPI_ISL_472264, EPI_ISL_472265, EPI_ISL_472266, EPI_ISL_472267, EPI_ISL_472268, EPI_ISL_472269, EPI_ISL_472270, EPI_ISL_472271, EPI_ISL_472272, EPI_ISL_472273, EPI_ISL_472274, EPI_ISL_472275, EPI_ISL_472276, EPI_ISL_472277, EPI_ISL_472278, EPI_ISL_472279, EPI_ISL_472280, EPI_ISL_472281, EPI_ISL_472282, EPI_ISL_472283, EPI_ISL_472284, EPI_ISL_472285, EPI_ISL_472286, EPI_ISL_472287, EPI_ISL_472288, EPI_ISL_472289, EPI_ISL_472290 | | | |
| see above | Northumbria University / South Tees Hospitals NHS Foundation Trust / North Cumbria Integrated Care NHS Foundation Trust / North Tees and Hartlepool NHS Foundation Trust / Newcastle Hospitals NHS Foundation Trust | COVID-19 Genomics UK (COG-UK) Consortium | Darren L Smith,Andrew Nelson,Matthew Bashton,Greg R Young,Joshua Loh,John Allan,Mohammad A Tariq,Giles S Holt,Gary Black,Wen C Yew,Lynn Dover,Paul Baker,Steve Liggett,Sarah Essex,Jane Greenaway,Debra Padgett,Clive Graham,Garren Scott,Edward Barton,Emma Swindells,Brendan Payne,Jennifer Collins,Yusri Taha,Gary Eltringham |
| EPI_ISL_472291, EPI_ISL_472292, EPI_ISL_472293, EPI_ISL_472294, EPI_ISL_472295, EPI_ISL_472296, EPI_ISL_472297, EPI_ISL_472298, EPI_ISL_472299, EPI_ISL_472300, EPI_ISL_472301, EPI_ISL_472302, EPI_ISL_472303, EPI_ISL_472304, EPI_ISL_472305, EPI_ISL_472306, EPI_ISL_472307, EPI_ISL_472308, EPI_ISL_472309, EPI_ISL_472310, EPI_ISL_472311, EPI_ISL_472312, EPI_ISL_472313, EPI_ISL_472314, EPI_ISL_472315, EPI_ISL_472316, EPI_ISL_472317, EPI_ISL_472318, EPI_ISL_472319, EPI_ISL_472320, EPI_ISL_472321, EPI_ISL_472322, EPI_ISL_472323, EPI_ISL_472324, EPI_ISL_472325, EPI_ISL_472326, EPI_ISL_472327, EPI_ISL_472328, EPI_ISL_472329, EPI_ISL_472330, EPI_ISL_472331, EPI_ISL_472332, EPI_ISL_472333, EPI_ISL_472334, EPI_ISL_472335, EPI_ISL_472336, EPI_ISL_472337, EPI_ISL_472338, EPI_ISL_472339, EPI_ISL_472340, EPI_ISL_472341, EPI_ISL_472342, EPI_ISL_472343, EPI_ISL_472344, EPI_ISL_472345, EPI_ISL_472346, EPI_ISL_472347, EPI_ISL_472348, EPI_ISL_472349, EPI_ISL_472350, EPI_ISL_472351, EPI_ISL_472352, EPI_ISL_472353, EPI_ISL_472354, EPI_ISL_472355, EPI_ISL_472356, EPI_ISL_472357, EPI_ISL_472358, EPI_ISL_472359, EPI_ISL_472360, EPI_ISL_472361, EPI_ISL_472362, EPI_ISL_472363, EPI_ISL_472364, EPI_ISL_472365, EPI_ISL_472366, EPI_ISL_472367, EPI_ISL_472368, EPI_ISL_472369, EPI_ISL_472370, EPI_ISL_472371, EPI_ISL_472372, EPI_ISL_472373, EPI_ISL_472374, EPI_ISL_472375, EPI_ISL_472376, EPI_ISL_472377, EPI_ISL_472378, EPI_ISL_472379, EPI_ISL_472380, EPI_ISL_472381, EPI_ISL_472382, EPI_ISL_472383 | | | |
| see above | Quadram Institute Bioscience | COVID-19 Genomics UK (COG-UK) Consortium | Dave J. Baker, Gemma L. Kay, Alp Aydin, Thanh Le-Viet, Steven Rudder, Ana P. Tedim, Anastasia Kolyva, Maria Diaz, Leonardo de Oliveira Martins, Nabil-Fareed Alikhan, Lizzie Meadows, Rachael Stanley, Ngozi Elumogo, Muhammed Yasir, Nicholas M. Thomson, Alexander J Trotter, Rachel Gilroy, Samuel Bloomfield, Claire Stuart, Andrew Bell, Reenesh Prakash, Samir Dervisevic, Alison E. Mather, John Wain, Mark Webber, Andrew J. Page, Justin O'Grady |
| EPI_ISL_472384, EPI_ISL_472385, EPI_ISL_472386, EPI_ISL_472387, EPI_ISL_472388, EPI_ISL_472389, EPI_ISL_472390, EPI_ISL_472391, EPI_ISL_472392, EPI_ISL_472393, EPI_ISL_472394, EPI_ISL_472395, EPI_ISL_472396, EPI_ISL_472397, EPI_ISL_472398, EPI_ISL_472399, EPI_ISL_472400, EPI_ISL_472401, EPI_ISL_472402, EPI_ISL_472403, EPI_ISL_472404, EPI_ISL_472405, EPI_ISL_472406, EPI_ISL_472407, EPI_ISL_472408, EPI_ISL_472409, EPI_ISL_472410, EPI_ISL_472411, EPI_ISL_472412, EPI_ISL_472413, EPI_ISL_472414, EPI_ISL_472415, EPI_ISL_472416, EPI_ISL_472417, EPI_ISL_472418, EPI_ISL_472419, EPI_ISL_472420, EPI_ISL_472421, EPI_ISL_472422, EPI_ISL_472423, EPI_ISL_472424, EPI_ISL_472425, EPI_ISL_472426, EPI_ISL_472427, EPI_ISL_472428, EPI_ISL_472429 | | | |
| see above | Queens Medical Centre, Clinical Microbiology Department / DeepSeq Nottingham | COVID-19 Genomics UK (COG-UK) Consortium | Gemma Clark, Wendy Smith, Manjinder Khakh, Vicki M Fleming, Michelle M Lister, Hannah Howson-Wells, Jonathan Ball, Patrick McClure, Joseph Chappell, Theocharis Tsoleridis, Nadine Holmes, Matthew Carlisle, Christopher Moore, Fei Sang, Johnny Debebe, Victoria Wright, Matthew Loose |
| EPI_ISL_472430, EPI_ISL_472431 | Queens Medical Centre, Clinical Microbiology Department / DeepSeq Nottingham | COVID-19 Genomics UK (COG-UK) Consortium | Nichola Duckworth, Tim Sloan, Sarah Walsh, Jonathan Ball, Patrick McClure, Joeseph Chappell, Nadine Holmes, Matthew Carlisle, Christopher Moore, Fei Sang, Johnny Debebe, Victoria Wright, Matthew Loose |
| EPI_ISL_472432, EPI_ISL_472433, EPI_ISL_472434, EPI_ISL_472435, EPI_ISL_472436, EPI_ISL_472437, EPI_ISL_472438, EPI_ISL_472439, EPI_ISL_472440, EPI_ISL_472441, EPI_ISL_472442, EPI_ISL_472443, EPI_ISL_472444, EPI_ISL_472445, EPI_ISL_472446, EPI_ISL_472447, EPI_ISL_472448, EPI_ISL_472449, EPI_ISL_472450, EPI_ISL_472451, EPI_ISL_472452, EPI_ISL_472453, EPI_ISL_472454, EPI_ISL_472455, EPI_ISL_472456, EPI_ISL_472457, EPI_ISL_472458, EPI_ISL_472459, EPI_ISL_472460, EPI_ISL_472461, EPI_ISL_472462, EPI_ISL_472463, EPI_ISL_472464, EPI_ISL_472465, EPI_ISL_472466, EPI_ISL_472467, EPI_ISL_472468, EPI_ISL_472469, EPI_ISL_472470, EPI_ISL_472471, EPI_ISL_472472, EPI_ISL_472473, EPI_ISL_472474, EPI_ISL_472475, EPI_ISL_472476, EPI_ISL_472477, EPI_ISL_472478, EPI_ISL_472479, EPI_ISL_472480, EPI_ISL_472481, EPI_ISL_472482, EPI_ISL_472483, EPI_ISL_472484, EPI_ISL_472485, EPI_ISL_472486, EPI_ISL_472487, EPI_ISL_472488, EPI_ISL_472489, EPI_ISL_472490, EPI_ISL_472491, EPI_ISL_472492, EPI_ISL_472493, EPI_ISL_472494, EPI_ISL_472495, EPI_ISL_472496, EPI_ISL_472497, EPI_ISL_472498, EPI_ISL_472499, EPI_ISL_472500, EPI_ISL_472501, EPI_ISL_472502, EPI_ISL_472503, EPI_ISL_472504, EPI_ISL_472505, EPI_ISL_472506, EPI_ISL_472507, EPI_ISL_472508, EPI_ISL_472509, EPI_ISL_472510, EPI_ISL_472511, EPI_ISL_472512, EPI_ISL_472513, EPI_ISL_472514, EPI_ISL_472515, EPI_ISL_472516, EPI_ISL_472517, EPI_ISL_472518, EPI_ISL_472519, EPI_ISL_472520, EPI_ISL_472521, EPI_ISL_472522, EPI_ISL_472523, EPI_ISL_472524, EPI_ISL_472525, EPI_ISL_472526, EPI_ISL_472527, EPI_ISL_472528, EPI_ISL_472529, EPI_ISL_472530, EPI_ISL_472531, EPI_ISL_472532, EPI_ISL_472533, EPI_ISL_472534, EPI_ISL_472535, EPI_ISL_472536, EPI_ISL_472537, EPI_ISL_472538, EPI_ISL_472539, EPI_ISL_472540, EPI_ISL_472541, EPI_ISL_472542, EPI_ISL_472543, EPI_ISL_472544, EPI_ISL_472545, EPI_ISL_472546, EPI_ISL_472547, EPI_ISL_472548, EPI_ISL_472549, EPI_ISL_472550, EPI_ISL_472551, EPI_ISL_472552, EPI_ISL_472553, EPI_ISL_472554, EPI_ISL_472555, EPI_ISL_472556, EPI_ISL_472557, EPI_ISL_472558, EPI_ISL_472559, EPI_ISL_472560, EPI_ISL_472561, EPI_ISL_472562, EPI_ISL_472563, EPI_ISL_472564, EPI_ISL_472565, EPI_ISL_472566, EPI_ISL_472567, EPI_ISL_472568, EPI_ISL_472569, EPI_ISL_472570, EPI_ISL_472571, EPI_ISL_472572, EPI_ISL_472573, EPI_ISL_472574, EPI_ISL_472575, EPI_ISL_472576, EPI_ISL_472577, EPI_ISL_472578, EPI_ISL_472579, EPI_ISL_472580, EPI_ISL_472581, EPI_ISL_472582, EPI_ISL_472583, EPI_ISL_472584, EPI_ISL_472585, EPI_ISL_472586, EPI_ISL_472587, EPI_ISL_472588, EPI_ISL_472589, EPI_ISL_472590, EPI_ISL_472591, EPI_ISL_472592, EPI_ISL_472593, | | | |

EPI_ISL_472594, EPI_ISL_472595, EPI_ISL_472596, EPI_ISL_472597, EPI_ISL_472598, EPI_ISL_472599, EPI_ISL_472600, EPI_ISL_472601, EPI_ISL_472602, EPI_ISL_472603, EPI_ISL_472604, EPI_ISL_472605, EPI_ISL_472606, EPI_ISL_472607, EPI_ISL_472608, EPI_ISL_472609, EPI_ISL_472610, EPI_ISL_472611, EPI_ISL_472612, EPI_ISL_472613, EPI_ISL_472614, EPI_ISL_472615, EPI_ISL_472616, EPI_ISL_472617, EPI_ISL_472618, EPI_ISL_472619, EPI_ISL_472620, EPI_ISL_472621, EPI_ISL_472622, EPI_ISL_472623, EPI_ISL_472624, EPI_ISL_472625, EPI_ISL_472626, EPI_ISL_472627, EPI_ISL_472628, EPI_ISL_472629, EPI_ISL_472630, EPI_ISL_472631, EPI_ISL_472632, EPI_ISL_472633, EPI_ISL_472634, EPI_ISL_472635, EPI_ISL_472636, EPI_ISL_472637, EPI_ISL_472638, EPI_ISL_472639, EPI_ISL_472640, EPI_ISL_472641, EPI_ISL_472642, EPI_ISL_472643, EPI_ISL_472644, EPI_ISL_472645, EPI_ISL_472646, EPI_ISL_472647, EPI_ISL_472648, EPI_ISL_472649, EPI_ISL_472650, EPI_ISL_472651, EPI_ISL_472652, EPI_ISL_472653, EPI_ISL_472654, EPI_ISL_472655, EPI_ISL_472656, EPI_ISL_472657, EPI_ISL_472658, EPI_ISL_472659, EPI_ISL_472660, EPI_ISL_472661, EPI_ISL_472662, EPI_ISL_472663, EPI_ISL_472664, EPI_ISL_472665, EPI_ISL_472666, EPI_ISL_472667, EPI_ISL_472668, EPI_ISL_472669, EPI_ISL_472670, EPI_ISL_472671, EPI_ISL_472672, EPI_ISL_472673, EPI_ISL_472674, EPI_ISL_472675, EPI_ISL_472676, EPI_ISL_472677, EPI_ISL_472678, EPI_ISL_472679, EPI_ISL_472680, EPI_ISL_472681, EPI_ISL_472682, EPI_ISL_472683, EPI_ISL_472684, EPI_ISL_472685, EPI_ISL_472686, EPI_ISL_472687, EPI_ISL_472688, EPI_ISL_472689, EPI_ISL_472690, EPI_ISL_472691, EPI_ISL_472692, EPI_ISL_472693, EPI_ISL_472694, EPI_ISL_472695, EPI_ISL_472696, EPI_ISL_472697, EPI_ISL_472698, EPI_ISL_472699, EPI_ISL_472700, EPI_ISL_472701, EPI_ISL_472702, EPI_ISL_472703, EPI_ISL_472704, EPI_ISL_472705, EPI_ISL_472706, EPI_ISL_472707, EPI_ISL_472708, EPI_ISL_472709, EPI_ISL_472710, EPI_ISL_472711, EPI_ISL_472712, EPI_ISL_472713, EPI_ISL_472714, EPI_ISL_472715, EPI_ISL_472716, EPI_ISL_472717, EPI_ISL_472718, EPI_ISL_472719, EPI_ISL_472720, EPI_ISL_472721, EPI_ISL_472722, EPI_ISL_472723, EPI_ISL_472724, EPI_ISL_472725, EPI_ISL_472726, EPI_ISL_472727, EPI_ISL_472728, EPI_ISL_472729, EPI_ISL_472730, EPI_ISL_472731, EPI_ISL_472732, EPI_ISL_472733, EPI_ISL_472734, EPI_ISL_472735, EPI_ISL_472736, EPI_ISL_472737, EPI_ISL_472738, EPI_ISL_472739, EPI_ISL_472740, EPI_ISL_472741, EPI_ISL_472742, EPI_ISL_472743, EPI_ISL_472744, EPI_ISL_472745, EPI_ISL_472746, EPI_ISL_472747, EPI_ISL_472748, EPI_ISL_472749, EPI_ISL_472750, EPI_ISL_472751, EPI_ISL_472752, EPI_ISL_472753, EPI_ISL_472754, EPI_ISL_472755, EPI_ISL_472756, EPI_ISL_472757, EPI_ISL_472758, EPI_ISL_472759, EPI_ISL_472760, EPI_ISL_472761, EPI_ISL_472762, EPI_ISL_472763, EPI_ISL_472764, EPI_ISL_472765, EPI_ISL_472766, EPI_ISL_472767, EPI_ISL_472768, EPI_ISL_472769, EPI_ISL_472770, EPI_ISL_472771, EPI_ISL_472772, EPI_ISL_472773, EPI_ISL_472774, EPI_ISL_472775, EPI_ISL_472776, EPI_ISL_472777, EPI_ISL_472778, EPI_ISL_472779, EPI_ISL_472780, EPI_ISL_472781, EPI_ISL_472782, EPI_ISL_472783, EPI_ISL_472784, EPI_ISL_472785, EPI_ISL_472786, EPI_ISL_472787, EPI_ISL_472788, EPI_ISL_472789, EPI_ISL_472790, EPI_ISL_472791, EPI_ISL_472792, EPI_ISL_472793, EPI_ISL_472794, EPI_ISL_472795, EPI_ISL_472796, EPI_ISL_472797, EPI_ISL_472798, EPI_ISL_472799, EPI_ISL_472800, EPI_ISL_472801, EPI_ISL_472802, EPI_ISL_472803, EPI_ISL_472804, EPI_ISL_472805, EPI_ISL_472806, EPI_ISL_472807, EPI_ISL_472808, EPI_ISL_472809, EPI_ISL_472810, EPI_ISL_472811, EPI_ISL_472812, EPI_ISL_472813, EPI_ISL_472814, EPI_ISL_472815, EPI_ISL_472816, EPI_ISL_472817, EPI_ISL_472818, EPI_ISL_472819, EPI_ISL_472820, EPI_ISL_472821, EPI_ISL_472822, EPI_ISL_472823, EPI_ISL_472824, EPI_ISL_472825, EPI_ISL_472826, EPI_ISL_472827, EPI_ISL_472828, EPI_ISL_472829, EPI_ISL_472830, EPI_ISL_472831, EPI_ISL_472832, EPI_ISL_472833, EPI_ISL_472834, EPI_ISL_472835, EPI_ISL_472836, EPI_ISL_472837, EPI_ISL_472838, EPI_ISL_472839, EPI_ISL_472840, EPI_ISL_472841, EPI_ISL_472842, EPI_ISL_472843, EPI_ISL_472844, EPI_ISL_472845, EPI_ISL_472846, EPI_ISL_472847, EPI_ISL_472848, EPI_ISL_472849, EPI_ISL_472850, EPI_ISL_472851, EPI_ISL_472852, EPI_ISL_472853, EPI_ISL_472854, EPI_ISL_472855, EPI_ISL_472856, EPI_ISL_472857, EPI_ISL_472858, EPI_ISL_472859, EPI_ISL_472860, EPI_ISL_472861, EPI_ISL_472862, EPI_ISL_472863, EPI_ISL_472864, EPI_ISL_472865, EPI_ISL_472866, EPI_ISL_472867, EPI_ISL_472868, EPI_ISL_472869, EPI_ISL_472870, EPI_ISL_472871, EPI_ISL_472872, EPI_ISL_472873, EPI_ISL_472874, EPI_ISL_472875, EPI_ISL_472876, EPI_ISL_472877, EPI_ISL_472878, EPI_ISL_472879, EPI_ISL_472880, EPI_ISL_472881, EPI_ISL_472882, EPI_ISL_472883, EPI_ISL_472884, EPI_ISL_472885, EPI_ISL_472886, EPI_ISL_472887, EPI_ISL_472888, EPI_ISL_472889, EPI_ISL_472890, EPI_ISL_472891, EPI_ISL_472892, EPI_ISL_472893, EPI_ISL_472894, EPI_ISL_472895, EPI_ISL_472896, EPI_ISL_472897, EPI_ISL_472898, EPI_ISL_472899, EPI_ISL_472900, EPI_ISL_472901, EPI_ISL_472902, EPI_ISL_472903, EPI_ISL_472904, EPI_ISL_472905, EPI_ISL_472906, EPI_ISL_472907, EPI_ISL_472908, EPI_ISL_472909, EPI_ISL_472910, EPI_ISL_472911, EPI_ISL_472912, EPI_ISL_472913, EPI_ISL_472914, EPI_ISL_472915, EPI_ISL_472916, EPI_ISL_472917, EPI_ISL_472918, EPI_ISL_472919, EPI_ISL_472920, EPI_ISL_472921, EPI_ISL_472922, EPI_ISL_472923, EPI_ISL_472924, EPI_ISL_472925, EPI_ISL_472926, EPI_ISL_472927, EPI_ISL_472928, EPI_ISL_472929, EPI_ISL_472930, EPI_ISL_472931, EPI_ISL_472932, EPI_ISL_472933, EPI_ISL_472934, EPI_ISL_472935, EPI_ISL_472936, EPI_ISL_472937, EPI_ISL_472938, EPI_ISL_472939, EPI_ISL_472940, EPI_ISL_472941, EPI_ISL_472942, EPI_ISL_472943, EPI_ISL_472944, EPI_ISL_472945, EPI_ISL_472946, EPI_ISL_472947, EPI_ISL_472948, EPI_ISL_472949, EPI_ISL_472950, EPI_ISL_472951, EPI_ISL_472952, EPI_ISL_472953, EPI_ISL_472954, EPI_ISL_472955, EPI_ISL_472956, EPI_ISL_472957, EPI_ISL_472958, EPI_ISL_472959, EPI_ISL_472960, EPI_ISL_472961, EPI_ISL_472962, EPI_ISL_472963, EPI_ISL_472964, EPI_ISL_472965, EPI_ISL_472966, EPI_ISL_472967, EPI_ISL_472968, EPI_ISL_472969, EPI_ISL_472970, EPI_ISL_472971, EPI_ISL_472972, EPI_ISL_472973, EPI_ISL_472974, EPI_ISL_472975, EPI_ISL_472976, EPI_ISL_472977, EPI_ISL_472978, EPI_ISL_472979, EPI_ISL_472980, EPI_ISL_472981, EPI_ISL_472982, EPI_ISL_472983, EPI_ISL_472984, EPI_ISL_472985, EPI_ISL_472986, EPI_ISL_472987, EPI_ISL_472988, EPI_ISL_472989, EPI_ISL_472990, EPI_ISL_472991, EPI_ISL_472992, EPI_ISL_472993, EPI_ISL_472994, EPI_ISL_472995, EPI_ISL_472996, EPI_ISL_472997, EPI_ISL_472998, EPI_ISL_472999, EPI_ISL_473000, EPI_ISL_473001, EPI_ISL_473002, EPI_ISL_473003, EPI_ISL_473004, EPI_ISL_473005, EPI_ISL_473006, EPI_ISL_473007, EPI_ISL_473008, EPI_ISL_473009, EPI_ISL_473010, EPI_ISL_473011, EPI_ISL_473012, EPI_ISL_473013, EPI_ISL_473014, EPI_ISL_473015, EPI_ISL_473016, EPI_ISL_473017, EPI_ISL_473018, EPI_ISL_473019, EPI_ISL_473020, EPI_ISL_473021, EPI_ISL_473022, EPI_ISL_473023, EPI_ISL_473024, EPI_ISL_473025, EPI_ISL_473026, EPI_ISL_473027, EPI_ISL_473028, EPI_ISL_473029, EPI_ISL_473030, EPI_ISL_473031, EPI_ISL_473032, EPI_ISL_473033, EPI_ISL_473034, EPI_ISL_473035, EPI_ISL_473036, EPI_ISL_473037, EPI_ISL_473038, EPI_ISL_473039, EPI_ISL_473040, EPI_ISL_473041, EPI_ISL_473042, EPI_ISL_473043, EPI_ISL_473044, EPI_ISL_473045, EPI_ISL_473046, EPI_ISL_473047, EPI_ISL_473048, EPI_ISL_473049, EPI_ISL_473050, EPI_ISL_473051, EPI_ISL_473052, EPI_ISL_473053, EPI_ISL_473054, EPI_ISL_473055, EPI_ISL_473056, EPI_ISL_473057, EPI_ISL_473058, EPI_ISL_473059, EPI_ISL_473060, EPI_ISL_473061, EPI_ISL_473062, EPI_ISL_473063, EPI_ISL_473064, EPI_ISL_473065, EPI_ISL_473066, EPI_ISL_473067, EPI_ISL_473068, EPI_ISL_473069, EPI_ISL_473070, EPI_ISL_473071, EPI_ISL_473072, EPI_ISL_473073, EPI_ISL_473074, EPI_ISL_473075, EPI_ISL_473076, EPI_ISL_473077, EPI_ISL_473078, EPI_ISL_473079, EPI_ISL_473080, EPI_ISL_473081, EPI_ISL_473082, EPI_ISL_473083, EPI_ISL_473084, EPI_ISL_473085, EPI_ISL_473086, EPI_ISL_473087, EPI_ISL_473088, EPI_ISL_473089, EPI_ISL_473090, EPI_ISL_473091, EPI_ISL_473092, EPI_ISL_473093, EPI_ISL_473094, EPI_ISL_473095, EPI_ISL_473096, EPI_ISL_473097, EPI_ISL_473098, EPI_ISL_473099, EPI_ISL_473100, EPI_ISL_473101, EPI_ISL_473102, EPI_ISL_473103, EPI_ISL_473104, EPI_ISL_473105, EPI_ISL_473106, EPI_ISL_473107, EPI_ISL_473108, EPI_ISL_473109, EPI_ISL_473110, EPI_ISL_473111, EPI_ISL_473112, EPI_ISL_473113, EPI_ISL_473114, EPI_ISL_473115, EPI_ISL_473116, EPI_ISL_473117, EPI_ISL_473118, EPI_ISL_473119, EPI_ISL_473120, EPI_ISL_473121, EPI_ISL_473122, EPI_ISL_473123, EPI_ISL_473124, EPI_ISL_473125, EPI_ISL_473126, EPI_ISL_473127, EPI_ISL_473128, EPI_ISL_473129, EPI_ISL_473130, EPI_ISL_473131, EPI_ISL_473132, EPI_ISL_473133, EPI_ISL_473134, EPI_ISL_473135, EPI_ISL_473136, EPI_ISL_473137, EPI_ISL_473138, EPI_ISL_473139, EPI_ISL_473140, EPI_ISL_473141, EPI_ISL_473142, EPI_ISL_473143, EPI_ISL_473144, EPI_ISL_473145, EPI_ISL_473146, EPI_ISL_473147, EPI_ISL_473148, EPI_ISL_473149, EPI_ISL_473150, EPI_ISL_473151, EPI_ISL_473152, EPI_ISL_473153, EPI_ISL_473154, EPI_ISL_473155, EPI_ISL_473156, EPI_ISL_473157, EPI_ISL_473158, EPI_ISL_473159, EPI_ISL_473160, EPI_ISL_473161, EPI_ISL_473162, EPI_ISL_473163, EPI_ISL_473164, EPI_ISL_473165, EPI_ISL_473166, EPI_ISL_473167, EPI_ISL_473168, EPI_ISL_473169, EPI_ISL_473170, EPI_ISL_473171, EPI_ISL_473172, EPI_ISL_473173, EPI_ISL_473174, EPI_ISL_473175, EPI_ISL_473176, EPI_ISL_473177, EPI_ISL_473178, EPI_ISL_473179, EPI_ISL_473180, EPI_ISL_473181, EPI_ISL_473182, EPI_ISL_473183, EPI_ISL_473184, EPI_ISL_473185, EPI_ISL_473186, EPI_ISL_473187, EPI_ISL_473188, EPI_ISL_473189, EPI_ISL_473190, EPI_ISL_473191, EPI_ISL_473192, EPI_ISL_473193, EPI_ISL_473194, EPI_ISL_473195, EPI_ISL_473196, EPI_ISL_473197, EPI_ISL_473198, EPI_ISL_473199, EPI_ISL_473200, EPI_ISL_473201, EPI_ISL_473202, EPI_ISL_473203, EPI_ISL_473204, EPI_ISL_473205, EPI_ISL_473206, EPI_ISL_473207, EPI_ISL_473208, EPI_ISL_473209, EPI_ISL_473210, EPI_ISL_473211, EPI_ISL_473212, EPI_ISL_473213, EPI_ISL_473214, EPI_ISL_473215, EPI_ISL_473216, EPI_ISL_473217, EPI_ISL_473218, EPI_ISL_473219, EPI_ISL_473220, EPI_ISL_473221, EPI_ISL_473222, EPI_ISL_473223, EPI_ISL_473224, EPI_ISL_473225, EPI_ISL_473226, EPI_ISL_473227, EPI_ISL_473228, EPI_ISL_473229, EPI_ISL_473230, EPI_ISL_473231, EPI_ISL_473232, EPI_ISL_473233, EPI_ISL_473234, EPI_ISL_473235, EPI_ISL_473236, EPI_ISL_473237, EPI_ISL_473238, EPI_ISL_473239, EPI_ISL_473240, EPI_ISL_473241, EPI_ISL_473242, EPI_ISL_473243, EPI_ISL_473244, EPI_ISL_473245, EPI_ISL_473246, EPI_ISL_473247, EPI_ISL_473248, EPI_ISL_473249, EPI_ISL_473250, EPI_ISL_473251, EPI_ISL_473252, EPI_ISL_473253, EPI_ISL_473254, EPI_ISL_473255, EPI_ISL_473256, EPI_ISL_473257, EPI_ISL_473258, EPI_ISL_473259, EPI_ISL_473260, EPI_ISL_473261, EPI_ISL_473262, EPI_ISL_473263, EPI_ISL_473264, EPI_ISL_473265, EPI_ISL_473266, EPI_ISL_473267, EPI_ISL_473268, EPI_ISL_473269, EPI_ISL_473270, EPI_ISL_473271, EPI_ISL_473272, EPI_ISL_473273, EPI_ISL_473274, EPI_ISL_473275, EPI_ISL_473276, EPI_ISL_473277, EPI_ISL_473278, EPI_ISL_473279, EPI_ISL_473280, EPI_ISL_473281, EPI_ISL_473282, EPI_ISL_473283, EPI_ISL_473284, EPI_ISL_473285, EPI_ISL_473286, EPI_ISL_473287, EPI_ISL_473288, EPI_ISL_473289, EPI_ISL_473290, EPI_ISL_473291, EPI_ISL_473292, EPI_ISL_473293, EPI_ISL_473294, EPI_ISL_473295, EPI_ISL_473296, EPI_ISL_473297, EPI_ISL_473298, EPI_ISL_473299, EPI_ISL_473300, EPI_ISL_473301, EPI_ISL_473302, EPI_ISL_473303, EPI_ISL_473304, EPI_ISL_473305

| | | | |
|---|---|---|---|
| see above | Wales Specialist Virology Centre Sequencing lab: Pathogen Genomics Unit | COVID-19 Genomics UK (COG-UK) Consortium | Catherine Moore, Johnathan Evans, Laura Gifford, Malorie Perry, Simon Cottrell, Angela Marchbank, Alec Birchley, Alexander Adams, Amy Gaskin, Bree Gatica-Wilcox, Jason Coombes, Joel Southgate, Lauren Gilbert, Lee Graham, Nicole Pacchiarini, Sara Kumziene-Summerhayes, Sarah Taylor, Sophie Jones, Sara Rey, Matthew Bull, Joanne Watkins, Sally Corden, Tom Connor |

EPI_ISL_473306, EPI_ISL_473307, EPI_ISL_473308, EPI_ISL_473309, EPI_ISL_473310, EPI_ISL_473311, EPI_ISL_473312, EPI_ISL_473313, EPI_ISL_473314, EPI_ISL_473315, EPI_ISL_473316, EPI_ISL_473317, EPI_ISL_473318, EPI_ISL_473319, EPI_ISL_473320, EPI_ISL_473321, EPI_ISL_473322, EPI_ISL_473323, EPI_ISL_473324, EPI_ISL_473325, EPI_ISL_473326, EPI_ISL_473327, EPI_ISL_473328, EPI_ISL_473329, EPI_ISL_473330, EPI_ISL_473331, EPI_ISL_473332, EPI_ISL_473333, EPI_ISL_473334, EPI_ISL_473335, EPI_ISL_473336, EPI_ISL_473337, EPI_ISL_473338, EPI_ISL_473339, EPI_ISL_473340, EPI_ISL_473341, EPI_ISL_473342, EPI_ISL_473343, EPI_ISL_473344, EPI_ISL_473345, EPI_ISL_473346, EPI_ISL_473347, EPI_ISL_473348, EPI_ISL_473349, EPI_ISL_473350, EPI_ISL_473351, EPI_ISL_473352, EPI_ISL_473353, EPI_ISL_473354, EPI_ISL_473355, EPI_ISL_473356, EPI_ISL_473357, EPI_ISL_473358, EPI_ISL_473359, EPI_ISL_473360, EPI_ISL_473361, EPI_ISL_473362, EPI_ISL_473363, EPI_ISL_473364, EPI_ISL_473365, EPI_ISL_473366, EPI_ISL_473367, EPI_ISL_473368, EPI_ISL_473369, EPI_ISL_473370, EPI_ISL_473371, EPI_ISL_473372, EPI_ISL_473373, EPI_ISL_473374, EPI_ISL_473375, EPI_ISL_473376, EPI_ISL_473377, EPI_ISL_473378, EPI_ISL_473379, EPI_ISL_473380, EPI_ISL_473381, EPI_ISL_473382, EPI_ISL_473383, EPI_ISL_473384, EPI_ISL_473385, EPI_ISL_473386, EPI_ISL_473387, EPI_ISL_473388, EPI_ISL_473389, EPI_ISL_473390, EPI_ISL_473391, EPI_ISL_473392, EPI_ISL_473393, EPI_ISL_473394, EPI_ISL_473395, EPI_ISL_473396, EPI_ISL_473397, EPI_ISL_473398, EPI_ISL_473399, EPI_ISL_473400, EPI_ISL_473401, EPI_ISL_473402, EPI_ISL_473403, EPI_ISL_473404, EPI_ISL_473405, EPI_ISL_473406, EPI_ISL_473407, EPI_ISL_473408, EPI_ISL_473409, EPI_ISL_473410, EPI_ISL_473411, EPI_ISL_473412, EPI_ISL_473413, EPI_ISL_473414, EPI_ISL_473415, EPI_ISL_473416, EPI_ISL_473417, EPI_ISL_473418, EPI_ISL_473419, EPI_ISL_473420, EPI_ISL_473421, EPI_ISL_473422, EPI_ISL_473423, EPI_ISL_473424, EPI_ISL_473425, EPI_ISL_473426, EPI_ISL_473427, EPI_ISL_473428, EPI_ISL_473429, EPI_ISL_473430, EPI_ISL_473431, EPI_ISL_473432, EPI_ISL_473433, EPI_ISL_473434, EPI_ISL_473435, EPI_ISL_473436, EPI_ISL_473437, EPI_ISL_473438, EPI_ISL_473439, EPI_ISL_473440, EPI_ISL_473441, EPI_ISL_473442, EPI_ISL_473443, EPI_ISL_473444, EPI_ISL_473445, EPI_ISL_473446, EPI_ISL_473447, EPI_ISL_473448, EPI_ISL_473449, EPI_ISL_473450

| | | | |
|---|---|---|---|
| see above | University of Birmingham | COVID-19 Genomics UK (COG-UK) Consortium | Institute of Microbiology, University of Birmingham: Claire McMurray, Joanne Stockton, Samuel Nicholls, Radoslaw Poplawski, Will Rowe, Josh Quick, Nicholas Loman, University of Birmingham Testing Laboratory: Celina M Whalley, Andrew Bosworth, Charlotte Poxon, Kasun Wanigasooriya, Oliver Pickles, Mike Kidd, Alex Richter, Andrew D Beggs PHE Heartlands Lab: Husam Osman, Andrew Bosworth. Queen Elizabeth Hospital: Anna Casey |

EPI_ISL_473451, EPI_ISL_473452, EPI_ISL_473453, EPI_ISL_473454, EPI_ISL_473455, EPI_ISL_473456, EPI_ISL_473457, EPI_ISL_473458, EPI_ISL_473459, EPI_ISL_473460, EPI_ISL_473461, EPI_ISL_473462, EPI_ISL_473463, EPI_ISL_473464, EPI_ISL_473465, EPI_ISL_473466, EPI_ISL_473467, EPI_ISL_473468, EPI_ISL_473469, EPI_ISL_473470, EPI_ISL_473471, EPI_ISL_473472, EPI_ISL_473473, EPI_ISL_473474, EPI_ISL_473475, EPI_ISL_473476, EPI_ISL_473477, EPI_ISL_473478, EPI_ISL_473479, EPI_ISL_473480, EPI_ISL_473481, EPI_ISL_473482, EPI_ISL_473483, EPI_ISL_473484, EPI_ISL_473485, EPI_ISL_473486, EPI_ISL_473487, EPI_ISL_473488, EPI_ISL_473489, EPI_ISL_473490, EPI_ISL_473491, EPI_ISL_473492, EPI_ISL_473493, EPI_ISL_473494, EPI_ISL_473495, EPI_ISL_473496, EPI_ISL_473497, EPI_ISL_473498, EPI_ISL_473499, EPI_ISL_473500, EPI_ISL_473501, EPI_ISL_473502, EPI_ISL_473503, EPI_ISL_473504, EPI_ISL_473505, EPI_ISL_473506, EPI_ISL_473507

| | | | |
|---|---|---|---|
| see above | Department of Pathology, University of Cambridge | COVID-19 Genomics UK (COG-UK) Consortium | Luke W Meredith, M. Estée Török, Myra Hosmillo, William L. Hamilton, Martin D. Curran, Theresa Feltwell, Grant Hall, Anna Yakovleva, Fahad A Khokhar, Charlotte J. Houldcroft, Laura G Caller, Aminu S. Jahun, Sarah L. Caddy, Yasmin Chaudhry, Malte Pinckert, Ian Goodfellow |

EPI_ISL_473508, EPI_ISL_473509, EPI_ISL_473510, EPI_ISL_473511, EPI_ISL_473512, EPI_ISL_473513, EPI_ISL_473514, EPI_ISL_473515, EPI_ISL_473516, EPI_ISL_473517, EPI_ISL_473518, EPI_ISL_473519, EPI_ISL_473520, EPI_ISL_473521, EPI_ISL_473522, EPI_ISL_473523, EPI_ISL_473524, EPI_ISL_473525, EPI_ISL_473526, EPI_ISL_473527, EPI_ISL_473528, EPI_ISL_473529, EPI_ISL_473530, EPI_ISL_473531, EPI_ISL_473532, EPI_ISL_473533, EPI_ISL_473534, EPI_ISL_473535, EPI_ISL_473536, EPI_ISL_473537, EPI_ISL_473538, EPI_ISL_473539, EPI_ISL_473540, EPI_ISL_473541, EPI_ISL_473542, EPI_ISL_473543, EPI_ISL_473544, EPI_ISL_473545, EPI_ISL_473546, EPI_ISL_473547, EPI_ISL_473548, EPI_ISL_473549, EPI_ISL_473550, EPI_ISL_473551, EPI_ISL_473552, EPI_ISL_473553, EPI_ISL_473554, EPI_ISL_473555, EPI_ISL_473556, EPI_ISL_473557, EPI_ISL_473558, EPI_ISL_473559, EPI_ISL_473560, EPI_ISL_473561, EPI_ISL_473562, EPI_ISL_473563, EPI_ISL_473564, EPI_ISL_473565, EPI_ISL_473566, EPI_ISL_473567, EPI_ISL_473568, EPI_ISL_473569, EPI_ISL_473570, EPI_ISL_473571, EPI_ISL_473572, EPI_ISL_473573, EPI_ISL_473574, EPI_ISL_473575, EPI_ISL_473576, EPI_ISL_473577, EPI_ISL_473578, EPI_ISL_473579, EPI_ISL_473580, EPI_ISL_473581, EPI_ISL_473582, EPI_ISL_473583, EPI_ISL_473584, EPI_ISL_473585, EPI_ISL_473586, EPI_ISL_473587, EPI_ISL_473588, EPI_ISL_473589, EPI_ISL_473590, EPI_ISL_473591, EPI_ISL_473592, EPI_ISL_473593, EPI_ISL_473594, EPI_ISL_473595, EPI_ISL_473596, EPI_ISL_473597, EPI_ISL_473598, EPI_ISL_473599, EPI_ISL_473600, EPI_ISL_473601, EPI_ISL_473602, EPI_ISL_473603, EPI_ISL_473604, EPI_ISL_473605, EPI_ISL_473606, EPI_ISL_473607, EPI_ISL_473608, EPI_ISL_473609, EPI_ISL_473610, EPI_ISL_473611, EPI_ISL_473612, EPI_ISL_473613, EPI_ISL_473614, EPI_ISL_473615, EPI_ISL_473616, EPI_ISL_473617, EPI_ISL_473618, EPI_ISL_473619, EPI_ISL_473620, EPI_ISL_473621, EPI_ISL_473622, EPI_ISL_473623, EPI_ISL_473624, EPI_ISL_473625, EPI_ISL_473626, EPI_ISL_473627, EPI_ISL_473628, EPI_ISL_473629, EPI_ISL_473630, EPI_ISL_473631, EPI_ISL_473632, EPI_ISL_473633, EPI_ISL_473634, EPI_ISL_473635, EPI_ISL_473636, EPI_ISL_473637, EPI_ISL_473638, EPI_ISL_473639, EPI_ISL_473640, EPI_ISL_473641, EPI_ISL_473642, EPI_ISL_473643, EPI_ISL_473644, EPI_ISL_473645, EPI_ISL_473646, EPI_ISL_473647, EPI_ISL_473648, EPI_ISL_473649, EPI_ISL_473650, EPI_ISL_473651, EPI_ISL_473652, EPI_ISL_473653, EPI_ISL_473654, EPI_ISL_473655, EPI_ISL_473656, EPI_ISL_473657, EPI_ISL_473658, EPI_ISL_473659, EPI_ISL_473660, EPI_ISL_473661, EPI_ISL_473662, EPI_ISL_473663, EPI_ISL_473664, EPI_ISL_473665, EPI_ISL_473666, EPI_ISL_473667, EPI_ISL_473668, EPI_ISL_473669, EPI_ISL_473670, EPI_ISL_473671, EPI_ISL_473672, EPI_ISL_473673, EPI_ISL_473674, EPI_ISL_473675, EPI_ISL_473676, EPI_ISL_473677, EPI_ISL_473678, EPI_ISL_473679, EPI_ISL_473680, EPI_ISL_473681, EPI_ISL_473682, EPI_ISL_473683, EPI_ISL_473684, EPI_ISL_473685, EPI_ISL_473686, EPI_ISL_473687, EPI_ISL_473688, EPI_ISL_473689, EPI_ISL_473690, EPI_ISL_473691, EPI_ISL_473692, EPI_ISL_473693, EPI_ISL_473694, EPI_ISL_473695, EPI_ISL_473696, EPI_ISL_473697, EPI_ISL_473698, EPI_ISL_473699, EPI_ISL_473700, EPI_ISL_473701, EPI_ISL_473702, EPI_ISL_473703, EPI_ISL_473704, EPI_ISL_473705, EPI_ISL_473706, EPI_ISL_473707, EPI_ISL_473708, EPI_ISL_473709, EPI_ISL_473710, EPI_ISL_473711, EPI_ISL_473712, EPI_ISL_473713, EPI_ISL_473714, EPI_ISL_473715, EPI_ISL_473716, EPI_ISL_473717, EPI_ISL_473718, EPI_ISL_473719, EPI_ISL_473720, EPI_ISL_473721, EPI_ISL_473722, EPI_ISL_473723, EPI_ISL_473724, EPI_ISL_473725, EPI_ISL_473726, EPI_ISL_473727, EPI_ISL_473728, EPI_ISL_473729, EPI_ISL_473730, EPI_ISL_473731, EPI_ISL_473732, EPI_ISL_473733, EPI_ISL_473734, EPI_ISL_473735, EPI_ISL_473736, EPI_ISL_473737, EPI_ISL_473738, EPI_ISL_473739, EPI_ISL_473740, EPI_ISL_473741, EPI_ISL_473742, EPI_ISL_473743, EPI_ISL_473744, EPI_ISL_473745, EPI_ISL_473746, EPI_ISL_473747, EPI_ISL_473748, EPI_ISL_473749, EPI_ISL_473750, EPI_ISL_473751, EPI_ISL_473752, EPI_ISL_473753, EPI_ISL_473754, EPI_ISL_473755, EPI_ISL_473756, EPI_ISL_473757, EPI_ISL_473758, EPI_ISL_473759, EPI_ISL_473760, EPI_ISL_473761, EPI_ISL_473762, EPI_ISL_473763, EPI_ISL_473764, EPI_ISL_473765, EPI_ISL_473766, EPI_ISL_473767, EPI_ISL_473768, EPI_ISL_473769, EPI_ISL_473770, EPI_ISL_473771, EPI_ISL_473772, EPI_ISL_473773, EPI_ISL_473774, EPI_ISL_473775, EPI_ISL_473776, EPI_ISL_473777, EPI_ISL_473778, EPI_ISL_473779, EPI_ISL_473780, EPI_ISL_473781, EPI_ISL_473782

| | | | |
|---|---|---|---|
| see above | West of Scotland Specialist Virology Centre, NHSGGC / MRC-University of Glasgow Centre for Virus Research | COVID-19 Genomics UK (COG-UK) Consortium | Ana da Silva Filipe, Natasha Johnson, Kathy Smollett, Daniel Mair, Stephen Carmichael, Lily Tong, Jenna Nichols, Elihu Aranday-Cortes, Kirstyn Brunker, Yasmin Parr, Alice Broos, Kyriaki Nomikou; Sarah McDonald, Marc Niebel, Patawee Asamaphan; Richard Orton, Joseph Hughes, Sreenu Vattipally, David L Robertson; Alasdair MacLean, Rory Gunson; Kathy Li, Natasha Jesudason, Rajiv Shah, James Shepherd, Antonia Ho, Emma Thomson |

EPI_ISL_473783, EPI_ISL_473784, EPI_ISL_473785, EPI_ISL_473786, EPI_ISL_473787, EPI_ISL_473788, EPI_ISL_473789, EPI_ISL_473790, EPI_ISL_473791, EPI_ISL_473792, EPI_ISL_473793, EPI_ISL_473794, EPI_ISL_473795, EPI_ISL_473796, EPI_ISL_473797, EPI_ISL_473798, EPI_ISL_473799, EPI_ISL_473800, EPI_ISL_473801, EPI_ISL_473802, EPI_ISL_473803, EPI_ISL_473804, EPI_ISL_473805, EPI_ISL_473806, EPI_ISL_473807, EPI_ISL_473808, EPI_ISL_473809, EPI_ISL_473810, EPI_ISL_473811, EPI_ISL_473812, EPI_ISL_473813, EPI_ISL_473814, EPI_ISL_473815, EPI_ISL_473816, EPI_ISL_473817, EPI_ISL_473818, EPI_ISL_473819, EPI_ISL_473820, EPI_ISL_473821, EPI_ISL_473822, EPI_ISL_473823, EPI_ISL_473824, EPI_ISL_473825, EPI_ISL_473826, EPI_ISL_473827, EPI_ISL_473828, EPI_ISL_473829, EPI_ISL_473830, EPI_ISL_473831, EPI_ISL_473832, EPI_ISL_473833, EPI_ISL_473834, EPI_ISL_473835, EPI_ISL_473836,

EPI_ISL_473837, EPI_ISL_473838, EPI_ISL_473839, EPI_ISL_473840, EPI_ISL_473841, EPI_ISL_473842, EPI_ISL_473843, EPI_ISL_473844, EPI_ISL_473845, EPI_ISL_473846, EPI_ISL_473847, EPI_ISL_473848, EPI_ISL_473849, EPI_ISL_473850, EPI_ISL_473851, EPI_ISL_473852, EPI_ISL_473853, EPI_ISL_473854, EPI_ISL_473855, EPI_ISL_473856, EPI_ISL_473857, EPI_ISL_473858, EPI_ISL_473859, EPI_ISL_473860, EPI_ISL_473861, EPI_ISL_473862, EPI_ISL_473863, EPI_ISL_473864, EPI_ISL_473865, EPI_ISL_473866, EPI_ISL_473867, EPI_ISL_473868, EPI_ISL_473869, EPI_ISL_473870, EPI_ISL_473871, EPI_ISL_473872, EPI_ISL_473873, EPI_ISL_473874, EPI_ISL_473875, EPI_ISL_473876, EPI_ISL_473877, EPI_ISL_473878, EPI_ISL_473879, EPI_ISL_473880, EPI_ISL_473881, EPI_ISL_473882, EPI_ISL_473883, EPI_ISL_473884, EPI_ISL_473885, EPI_ISL_473886, EPI_ISL_473887, EPI_ISL_473888, EPI_ISL_473889, EPI_ISL_473890, EPI_ISL_473891, EPI_ISL_473892, EPI_ISL_473893, EPI_ISL_473894, EPI_ISL_473895, EPI_ISL_473896, EPI_ISL_473897, EPI_ISL_473898, EPI_ISL_473899, EPI_ISL_473900, EPI_ISL_473901, EPI_ISL_473902, EPI_ISL_473903, EPI_ISL_473904, EPI_ISL_473905, EPI_ISL_473906, EPI_ISL_473907, EPI_ISL_473908, EPI_ISL_473909, EPI_ISL_473910, EPI_ISL_473911, EPI_ISL_473912, EPI_ISL_473913, EPI_ISL_473914, EPI_ISL_473915, EPI_ISL_473916, EPI_ISL_473917, EPI_ISL_473918, EPI_ISL_473919, EPI_ISL_473920, EPI_ISL_473921, EPI_ISL_473922, EPI_ISL_473923, EPI_ISL_473924, EPI_ISL_473925, EPI_ISL_473926, EPI_ISL_473927, EPI_ISL_473928, EPI_ISL_473929, EPI_ISL_473930, EPI_ISL_473931, EPI_ISL_473932, EPI_ISL_473933, EPI_ISL_473934, EPI_ISL_473935, EPI_ISL_473936, EPI_ISL_473937, EPI_ISL_473938, EPI_ISL_473939, EPI_ISL_473940, EPI_ISL_473941, EPI_ISL_473942, EPI_ISL_473943, EPI_ISL_473944, EPI_ISL_473945, EPI_ISL_473946, EPI_ISL_473947, EPI_ISL_473948, EPI_ISL_473949, EPI_ISL_473950, EPI_ISL_473951, EPI_ISL_473952, EPI_ISL_473953, EPI_ISL_473954, EPI_ISL_473955

| | | | |
|---|---|---|---|
| see above | Virology Department, Royal Infirmary of Edinburgh, NHS Lothian / School of Biological Sciences, University of Edinburgh / Institute of Genetics and Molecular Medicine, University of Edinburgh | COVID-19 Genomics UK (COG-UK) Consortium | McHugh M, Dewar R, Rooke S, Gallagher M, Balcaza C, O'Toole Á, Scher E, Hill V, McCrone JT, Colquhoun R, Yu X, Jackson B, Rambaut A, Williams TC, Templeton K |

EPI_ISL_473956, EPI_ISL_473957, EPI_ISL_473958, EPI_ISL_473959, EPI_ISL_473960, EPI_ISL_473961, EPI_ISL_473962, EPI_ISL_473963, EPI_ISL_473964, EPI_ISL_473965, EPI_ISL_473966, EPI_ISL_473967, EPI_ISL_473968, EPI_ISL_473969, EPI_ISL_473970, EPI_ISL_473971, EPI_ISL_473972, EPI_ISL_473973, EPI_ISL_473974, EPI_ISL_473975, EPI_ISL_473976, EPI_ISL_473977, EPI_ISL_473978, EPI_ISL_473979, EPI_ISL_473980, EPI_ISL_473981, EPI_ISL_473982, EPI_ISL_473983, EPI_ISL_473984, EPI_ISL_473985, EPI_ISL_473986, EPI_ISL_473987, EPI_ISL_473988, EPI_ISL_473989, EPI_ISL_473990, EPI_ISL_473991, EPI_ISL_473992, EPI_ISL_473993, EPI_ISL_473994, EPI_ISL_473995, EPI_ISL_473996, EPI_ISL_473997, EPI_ISL_473998, EPI_ISL_473999, EPI_ISL_474000, EPI_ISL_474001, EPI_ISL_474002, EPI_ISL_474003, EPI_ISL_474004, EPI_ISL_474005, EPI_ISL_474006, EPI_ISL_474007, EPI_ISL_474008, EPI_ISL_474009, EPI_ISL_474010, EPI_ISL_474011, EPI_ISL_474012, EPI_ISL_474013, EPI_ISL_474014, EPI_ISL_474015, EPI_ISL_474016, EPI_ISL_474017, EPI_ISL_474018, EPI_ISL_474019, EPI_ISL_474020, EPI_ISL_474021, EPI_ISL_474022, EPI_ISL_474023, EPI_ISL_474024, EPI_ISL_474025, EPI_ISL_474026, EPI_ISL_474027, EPI_ISL_474028, EPI_ISL_474029, EPI_ISL_474030, EPI_ISL_474031, EPI_ISL_474032, EPI_ISL_474033, EPI_ISL_474034, EPI_ISL_474035, EPI_ISL_474036, EPI_ISL_474037, EPI_ISL_474038, EPI_ISL_474039, EPI_ISL_474040, EPI_ISL_474041, EPI_ISL_474042, EPI_ISL_474043, EPI_ISL_474044, EPI_ISL_474045, EPI_ISL_474046, EPI_ISL_474047, EPI_ISL_474048, EPI_ISL_474049, EPI_ISL_474050, EPI_ISL_474051, EPI_ISL_474052, EPI_ISL_474053, EPI_ISL_474054, EPI_ISL_474055, EPI_ISL_474056, EPI_ISL_474057, EPI_ISL_474058, EPI_ISL_474059, EPI_ISL_474060, EPI_ISL_474061, EPI_ISL_474062, EPI_ISL_474063, EPI_ISL_474064, EPI_ISL_474065, EPI_ISL_474066, EPI_ISL_474067, EPI_ISL_474068, EPI_ISL_474069, EPI_ISL_474070, EPI_ISL_474071, EPI_ISL_474072, EPI_ISL_474073, EPI_ISL_474074, EPI_ISL_474075, EPI_ISL_474076, EPI_ISL_474077, EPI_ISL_474078, EPI_ISL_474079, EPI_ISL_474080, EPI_ISL_474081, EPI_ISL_474082, EPI_ISL_474083, EPI_ISL_474084, EPI_ISL_474085, EPI_ISL_474086, EPI_ISL_474087, EPI_ISL_474088, EPI_ISL_474089, EPI_ISL_474090, EPI_ISL_474091, EPI_ISL_474092, EPI_ISL_474093, EPI_ISL_474094, EPI_ISL_474095, EPI_ISL_474096, EPI_ISL_474097, EPI_ISL_474098, EPI_ISL_474099, EPI_ISL_474100, EPI_ISL_474101, EPI_ISL_474102, EPI_ISL_474103, EPI_ISL_474104, EPI_ISL_474105, EPI_ISL_474106, EPI_ISL_474107, EPI_ISL_474108, EPI_ISL_474109, EPI_ISL_474110, EPI_ISL_474111, EPI_ISL_474112, EPI_ISL_474113, EPI_ISL_474114, EPI_ISL_474115, EPI_ISL_474116, EPI_ISL_474117, EPI_ISL_474118, EPI_ISL_474119, EPI_ISL_474120, EPI_ISL_474121, EPI_ISL_474122, EPI_ISL_474123, EPI_ISL_474124, EPI_ISL_474125, EPI_ISL_474126, EPI_ISL_474127, EPI_ISL_474128, EPI_ISL_474129, EPI_ISL_474130, EPI_ISL_474131, EPI_ISL_474132, EPI_ISL_474133, EPI_ISL_474134, EPI_ISL_474135, EPI_ISL_474136, EPI_ISL_474137, EPI_ISL_474138, EPI_ISL_474139, EPI_ISL_474140, EPI_ISL_474141, EPI_ISL_474142, EPI_ISL_474143, EPI_ISL_474144, EPI_ISL_474145, EPI_ISL_474146, EPI_ISL_474147, EPI_ISL_474148, EPI_ISL_474149, EPI_ISL_474150, EPI_ISL_474151, EPI_ISL_474152, EPI_ISL_474153, EPI_ISL_474154, EPI_ISL_474155, EPI_ISL_474156, EPI_ISL_474157, EPI_ISL_474158, EPI_ISL_474159, EPI_ISL_474160, EPI_ISL_474161, EPI_ISL_474162, EPI_ISL_474163, EPI_ISL_474164, EPI_ISL_474165, EPI_ISL_474166, EPI_ISL_474167, EPI_ISL_474168, EPI_ISL_474169, EPI_ISL_474170, EPI_ISL_474171, EPI_ISL_474172, EPI_ISL_474173, EPI_ISL_474174, EPI_ISL_474175, EPI_ISL_474176, EPI_ISL_474177, EPI_ISL_474178, EPI_ISL_474179, EPI_ISL_474180, EPI_ISL_474181, EPI_ISL_474182, EPI_ISL_474183, EPI_ISL_474184, EPI_ISL_474185, EPI_ISL_474186, EPI_ISL_474187, EPI_ISL_474188, EPI_ISL_474189, EPI_ISL_474190, EPI_ISL_474191, EPI_ISL_474192, EPI_ISL_474193, EPI_ISL_474194, EPI_ISL_474195, EPI_ISL_474196, EPI_ISL_474197, EPI_ISL_474198, EPI_ISL_474199, EPI_ISL_474200, EPI_ISL_474201, EPI_ISL_474202, EPI_ISL_474203, EPI_ISL_474204, EPI_ISL_474205, EPI_ISL_474206, EPI_ISL_474207, EPI_ISL_474208, EPI_ISL_474209, EPI_ISL_474210, EPI_ISL_474211, EPI_ISL_474212, EPI_ISL_474213, EPI_ISL_474214, EPI_ISL_474215, EPI_ISL_474216, EPI_ISL_474217, EPI_ISL_474218, EPI_ISL_474219, EPI_ISL_474220, EPI_ISL_474221, EPI_ISL_474222, EPI_ISL_474223, EPI_ISL_474224, EPI_ISL_474225, EPI_ISL_474226, EPI_ISL_474227, EPI_ISL_474228, EPI_ISL_474229, EPI_ISL_474230, EPI_ISL_474231, EPI_ISL_474232, EPI_ISL_474233, EPI_ISL_474234, EPI_ISL_474235, EPI_ISL_474236, EPI_ISL_474237, EPI_ISL_474238, EPI_ISL_474239, EPI_ISL_474240, EPI_ISL_474241, EPI_ISL_474242, EPI_ISL_474243, EPI_ISL_474244, EPI_ISL_474245, EPI_ISL_474246, EPI_ISL_474247, EPI_ISL_474248, EPI_ISL_474249, EPI_ISL_474250, EPI_ISL_474251, EPI_ISL_474252, EPI_ISL_474253, EPI_ISL_474254, EPI_ISL_474255, EPI_ISL_474256, EPI_ISL_474257, EPI_ISL_474258, EPI_ISL_474259, EPI_ISL_474260, EPI_ISL_474261, EPI_ISL_474262, EPI_ISL_474263, EPI_ISL_474264, EPI_ISL_474265, EPI_ISL_474266, EPI_ISL_474267, EPI_ISL_474268, EPI_ISL_474269, EPI_ISL_474270, EPI_ISL_474271, EPI_ISL_474272, EPI_ISL_474273, EPI_ISL_474274, EPI_ISL_474275, EPI_ISL_474276, EPI_ISL_474277, EPI_ISL_474278, EPI_ISL_474279, EPI_ISL_474280, EPI_ISL_474281, EPI_ISL_474282, EPI_ISL_474283, EPI_ISL_474284, EPI_ISL_474285, EPI_ISL_474286, EPI_ISL_474287, EPI_ISL_474288, EPI_ISL_474289, EPI_ISL_474290, EPI_ISL_474291, EPI_ISL_474292, EPI_ISL_474293, EPI_ISL_474294, EPI_ISL_474295, EPI_ISL_474296, EPI_ISL_474297, EPI_ISL_474298, EPI_ISL_474299, EPI_ISL_474300, EPI_ISL_474301, EPI_ISL_474302, EPI_ISL_474303, EPI_ISL_474304, EPI_ISL_474305, EPI_ISL_474306, EPI_ISL_474307, EPI_ISL_474308, EPI_ISL_474309, EPI_ISL_474310, EPI_ISL_474311, EPI_ISL_474312, EPI_ISL_474313, EPI_ISL_474314, EPI_ISL_474315, EPI_ISL_474316, EPI_ISL_474317, EPI_ISL_474318, EPI_ISL_474319, EPI_ISL_474320, EPI_ISL_474321, EPI_ISL_474322, EPI_ISL_474323, EPI_ISL_474324, EPI_ISL_474325, EPI_ISL_474326, EPI_ISL_474327, EPI_ISL_474328, EPI_ISL_474329, EPI_ISL_474330, EPI_ISL_474331, EPI_ISL_474332, EPI_ISL_474333, EPI_ISL_474334, EPI_ISL_474335, EPI_ISL_474336, EPI_ISL_474337, EPI_ISL_474338, EPI_ISL_474339, EPI_ISL_474340, EPI_ISL_474341, EPI_ISL_474342, EPI_ISL_474343, EPI_ISL_474344, EPI_ISL_474345, EPI_ISL_474346, EPI_ISL_474347, EPI_ISL_474348, EPI_ISL_474349, EPI_ISL_474350, EPI_ISL_474351, EPI_ISL_474352, EPI_ISL_474353, EPI_ISL_474354, EPI_ISL_474355, EPI_ISL_474356, EPI_ISL_474357, EPI_ISL_474358, EPI_ISL_474359, EPI_ISL_474360, EPI_ISL_474361, EPI_ISL_474362, EPI_ISL_474363, EPI_ISL_474364, EPI_ISL_474365, EPI_ISL_474366, EPI_ISL_474367, EPI_ISL_474368, EPI_ISL_474369, EPI_ISL_474370, EPI_ISL_474371, EPI_ISL_474372, EPI_ISL_474373, EPI_ISL_474374, EPI_ISL_474375, EPI_ISL_474376, EPI_ISL_474377, EPI_ISL_474378, EPI_ISL_474379, EPI_ISL_474380, EPI_ISL_474381, EPI_ISL_474382, EPI_ISL_474383, EPI_ISL_474384, EPI_ISL_474385, EPI_ISL_474386, EPI_ISL_474387, EPI_ISL_474388, EPI_ISL_474389, EPI_ISL_474390, EPI_ISL_474391, EPI_ISL_474392, EPI_ISL_474393, EPI_ISL_474394, EPI_ISL_474395, EPI_ISL_474396, EPI_ISL_474397, EPI_ISL_474398, EPI_ISL_474399, EPI_ISL_474400, EPI_ISL_474401, EPI_ISL_474402, EPI_ISL_474403, EPI_ISL_474404, EPI_ISL_474405, EPI_ISL_474406, EPI_ISL_474407, EPI_ISL_474408, EPI_ISL_474409, EPI_ISL_474410, EPI_ISL_474411, EPI_ISL_474412, EPI_ISL_474413, EPI_ISL_474414, EPI_ISL_474415, EPI_ISL_474416, EPI_ISL_474417, EPI_ISL_474418, EPI_ISL_474419, EPI_ISL_474420, EPI_ISL_474421, EPI_ISL_474422, EPI_ISL_474423, EPI_ISL_474424, EPI_ISL_474425, EPI_ISL_474426, EPI_ISL_474427, EPI_ISL_474428, EPI_ISL_474429, EPI_ISL_474430, EPI_ISL_474431, EPI_ISL_474432, EPI_ISL_474433, EPI_ISL_474434, EPI_ISL_474435, EPI_ISL_474436, EPI_ISL_474437, EPI_ISL_474438, EPI_ISL_474439, EPI_ISL_474440, EPI_ISL_474441, EPI_ISL_474442, EPI_ISL_474443, EPI_ISL_474444, EPI_ISL_474445, EPI_ISL_474446, EPI_ISL_474447, EPI_ISL_474448, EPI_ISL_474449, EPI_ISL_474450, EPI_ISL_474451, EPI_ISL_474452, EPI_ISL_474453, EPI_ISL_474454, EPI_ISL_474455, EPI_ISL_474456, EPI_ISL_474457, EPI_ISL_474458, EPI_ISL_474459, EPI_ISL_474460, EPI_ISL_474461, EPI_ISL_474462, EPI_ISL_474463, EPI_ISL_474464, EPI_ISL_474465, EPI_ISL_474466, EPI_ISL_474467, EPI_ISL_474468, EPI_ISL_474469, EPI_ISL_474470, EPI_ISL_474471, EPI_ISL_474472, EPI_ISL_474473, EPI_ISL_474474, EPI_ISL_474475, EPI_ISL_474476, EPI_ISL_474477, EPI_ISL_474478, EPI_ISL_474479, EPI_ISL_474480, EPI_ISL_474481, EPI_ISL_474482, EPI_ISL_474483, EPI_ISL_474484, EPI_ISL_474485, EPI_ISL_474486, EPI_ISL_474487, EPI_ISL_474488, EPI_ISL_474489, EPI_ISL_474490, EPI_ISL_474491, EPI_ISL_474492, EPI_ISL_474493, EPI_ISL_474494, EPI_ISL_474495, EPI_ISL_474496, EPI_ISL_474497, EPI_ISL_474498, EPI_ISL_474499, EPI_ISL_474500, EPI_ISL_474501, EPI_ISL_474502, EPI_ISL_474503, EPI_ISL_474504, EPI_ISL_474505, EPI_ISL_474506, EPI_ISL_474507, EPI_ISL_474508, EPI_ISL_474509, EPI_ISL_474510, EPI_ISL_474511, EPI_ISL_474512, EPI_ISL_474513, EPI_ISL_474514, EPI_ISL_474515, EPI_ISL_474516, EPI_ISL_474517, EPI_ISL_474518, EPI_ISL_474519, EPI_ISL_474520, EPI_ISL_474521, EPI_ISL_474522, EPI_ISL_474523, EPI_ISL_474524, EPI_ISL_474525, EPI_ISL_474526, EPI_ISL_474527, EPI_ISL_474528, EPI_ISL_474529, EPI_ISL_474530, EPI_ISL_474531, EPI_ISL_474532, EPI_ISL_474533, EPI_ISL_474534, EPI_ISL_474535, EPI_ISL_474536, EPI_ISL_474537, EPI_ISL_474538, EPI_ISL_474539, EPI_ISL_474540, EPI_ISL_474541, EPI_ISL_474542, EPI_ISL_474543, EPI_ISL_474544, EPI_ISL_474545, EPI_ISL_474546, EPI_ISL_474547, EPI_ISL_474548, EPI_ISL_474549, EPI_ISL_474550, EPI_ISL_474551, EPI_ISL_474552, EPI_ISL_474553, EPI_ISL_474554, EPI_ISL_474555, EPI_ISL_474556, EPI_ISL_474557, EPI_ISL_474558, EPI_ISL_474559, EPI_ISL_474560, EPI_ISL_474561, EPI_ISL_474562, EPI_ISL_474563, EPI_ISL_474564, EPI_ISL_474565, EPI_ISL_474566, EPI_ISL_474567, EPI_ISL_474568, EPI_ISL_474569, EPI_ISL_474570, EPI_ISL_474571, EPI_ISL_474572, EPI_ISL_474573, EPI_ISL_474574, EPI_ISL_474575, EPI_ISL_474576, EPI_ISL_474577, EPI_ISL_474578, EPI_ISL_474579, EPI_ISL_474580, EPI_ISL_474581, EPI_ISL_474582, EPI_ISL_474583, EPI_ISL_474584, EPI_ISL_474585, EPI_ISL_474586, EPI_ISL_474587, EPI_ISL_474588, EPI_ISL_474589, EPI_ISL_474590, EPI_ISL_474591, EPI_ISL_474592, EPI_ISL_474593, EPI_ISL_474594, EPI_ISL_474595, EPI_ISL_474596, EPI_ISL_474597, EPI_ISL_474598, EPI_ISL_474599, EPI_ISL_474600, EPI_ISL_474601, EPI_ISL_474602, EPI_ISL_474603, EPI_ISL_474604, EPI_ISL_474605, EPI_ISL_474606, EPI_ISL_474607, EPI_ISL_474608, EPI_ISL_474609, EPI_ISL_474610, EPI_ISL_474611, EPI_ISL_474612, EPI_ISL_474613, EPI_ISL_474614, EPI_ISL_474615, EPI_ISL_474616, EPI_ISL_474617, EPI_ISL_474618, EPI_ISL_474619, EPI_ISL_474620, EPI_ISL_474621, EPI_ISL_474622, EPI_ISL_474623, EPI_ISL_474624, EPI_ISL_474625, EPI_ISL_474626, EPI_ISL_474627, EPI_ISL_474628, EPI_ISL_474629, EPI_ISL_474630, EPI_ISL_474631, EPI_ISL_474632, EPI_ISL_474633, EPI_ISL_474634, EPI_ISL_474635, EPI_ISL_474636, EPI_ISL_474637, EPI_ISL_474638, EPI_ISL_474639, EPI_ISL_474640, EPI_ISL_474641, EPI_ISL_474642, EPI_ISL_474643, EPI_ISL_474644, EPI_ISL_474645, EPI_ISL_474646, EPI_ISL_474647, EPI_ISL_474648, EPI_ISL_474649, EPI_ISL_474650, EPI_ISL_474651, EPI_ISL_474652, EPI_ISL_474653, EPI_ISL_474654, EPI_ISL_474655, EPI_ISL_474656, EPI_ISL_474657, EPI_ISL_474658, EPI_ISL_474659, EPI_ISL_474660, EPI_ISL_474661, EPI_ISL_474662, EPI_ISL_474663, EPI_ISL_474664, EPI_ISL_474665, EPI_ISL_474666, EPI_ISL_474667, EPI_ISL_474668, EPI_ISL_474669, EPI_ISL_474670, EPI_ISL_474671, EPI_ISL_474672, EPI_ISL_474673, EPI_ISL_474674, EPI_ISL_474675, EPI_ISL_474676, EPI_ISL_474677, EPI_ISL_474678, EPI_ISL_474679, EPI_ISL_474680, EPI_ISL_474681, EPI_ISL_474682, EPI_ISL_474683, EPI_ISL_474684, EPI_ISL_474685, EPI_ISL_474686, EPI_ISL_474687, EPI_ISL_474688, EPI_ISL_474689, EPI_ISL_474690, EPI_ISL_474691, EPI_ISL_474692, EPI_ISL_474693, EPI_ISL_474694, EPI_ISL_474695, EPI_ISL_474696, EPI_ISL_474697, EPI_ISL_474698, EPI_ISL_474699, EPI_ISL_474700, EPI_ISL_474701, EPI_ISL_474702, EPI_ISL_474703, EPI_ISL_474704, EPI_ISL_474705, EPI_ISL_474706, EPI_ISL_474707, EPI_ISL_474708, EPI_ISL_474709, EPI_ISL_474710, EPI_ISL_474711, EPI_ISL_474712, EPI_ISL_474713, EPI_ISL_474714, EPI_ISL_474715, EPI_ISL_474716, EPI_ISL_474717, EPI_ISL_474718, EPI_ISL_474719, EPI_ISL_474720, EPI_ISL_474721, EPI_ISL_474722, EPI_ISL_474723, EPI_ISL_474724, EPI_ISL_474725, EPI_ISL_474726, EPI_ISL_474727, EPI_ISL_474728, EPI_ISL_474729, EPI_ISL_474730, EPI_ISL_474731, EPI_ISL_474732, EPI_ISL_474733, EPI_ISL_474734, EPI_ISL_474735, EPI_ISL_474736, EPI_ISL_474737, EPI_ISL_474738, EPI_ISL_474739, EPI_ISL_474740, EPI_ISL_474741, EPI_ISL_474742, EPI_ISL_474743, EPI_ISL_474744, EPI_ISL_474745, EPI_ISL_474746, EPI_ISL_474747, EPI_ISL_474748, EPI_ISL_474749, EPI_ISL_474750, EPI_ISL_474751, EPI_ISL_474752, EPI_ISL_474753, EPI_ISL_474754, EPI_ISL_474755, EPI_ISL_474756, EPI_ISL_474757, EPI_ISL_474758, EPI_ISL_474759, EPI_ISL_474760, EPI_ISL_474761, EPI_ISL_474762, EPI_ISL_474763, EPI_ISL_474764, EPI_ISL_474765, EPI_ISL_474766, EPI_ISL_474767, EPI_ISL_474768, EPI_ISL_474769, EPI_ISL_474770, EPI_ISL_474771, EPI_ISL_474772, EPI_ISL_474773, EPI_ISL_474774, EPI_ISL_474775, EPI_ISL_474776, EPI_ISL_474777, EPI_ISL_474778, EPI_ISL_474779, EPI_ISL_474780, EPI_ISL_474781, EPI_ISL_474782, EPI_ISL_474783, EPI_ISL_474784, EPI_ISL_474785, EPI_ISL_474786, EPI_ISL_474787, EPI_ISL_474788, EPI_ISL_474789, EPI_ISL_474790, EPI_ISL_474791, EPI_ISL_474792, EPI_ISL_474793, EPI_ISL_474794, EPI_ISL_474795, EPI_ISL_474796

| | | | |
|---|---|---|---|
| see above | Originating lab: Wales Specialist Virology Centre<br>Sequencing lab: Pathogen Genomics Unit | COVID-19 Genomics UK (COG-UK) Consortium | Catherine Moore, Johnathan Evans, Laura Gifford, Malorie Perry, Simon Cottrell, Angela Marchbank, Alec Birchley, Alexander Adams, Amy Gaskin, Bree Gatica-Wilcox, Jason Coombes, Joel Southgate, Lauren Gilbert, Lee Graham, Nicole Pacchiarini, Sara Kumziene-Summerhayes, Sarah Taylor, Sophie Jones, Sara Rey, Matthew Bull, Joanne Watkins, Sally Corden, Tom Connor |
| EPI_ISL_474797 | Hospital Universitario Virgen de las Nieves de Granada-SAS | SeqCOVID-SPAIN consortium/IBV(CSIC) | Mercedes Pérez Ruiz, Sara Sanbonmatsu Gámez, Irene Pedrosa Corral, José M. Navarro-Marí and SeqCOVID-SPAIN consortium |

EPI_ISL_474798, EPI_ISL_474799, EPI_ISL_474800, EPI_ISL_474801, EPI_ISL_474802, EPI_ISL_474803, EPI_ISL_474804, EPI_ISL_474805, EPI_ISL_474806, EPI_ISL_474807, EPI_ISL_474808, EPI_ISL_474809, EPI_ISL_474810, EPI_ISL_474811, EPI_ISL_474812, EPI_ISL_474813, EPI_ISL_474814, EPI_ISL_474815, EPI_ISL_474816, EPI_ISL_474817, EPI_ISL_474818, EPI_ISL_474819, EPI_ISL_474820, EPI_ISL_474821, EPI_ISL_474822, EPI_ISL_474823, EPI_ISL_474824, EPI_ISL_474825, EPI_ISL_474826, EPI_ISL_474827, EPI_ISL_474828, EPI_ISL_474829, EPI_ISL_474830, EPI_ISL_474831

| | | | |
|---|---|---|---|
| see above | Complejo Hospitalario Universitario de Albacete | SeqCOVID-SPAIN consortium/IBV(CSIC) | Encarnacion Simarro Córdoba, Julia Lozano Serra, Lorena Robles Fonseca , Monica Parra Grandes, Caridad Sainz de Baranda Camino and SeqCOVID-SPAIN consortium |
| EPI_ISL_474832, EPI_ISL_474833, EPI_ISL_474834, EPI_ISL_474835, EPI_ISL_474836 | Hospital Universitario Virgen de las Nieves de Granada-SAS | SeqCOVID-SPAIN consortium/IBV(CSIC) | Mercedes Pérez Ruiz, Sara Sanbonmatsu Gámez, Irene Pedrosa Corral, José M. Navarro-Marí and SeqCOVID-SPAIN consortium |
| EPI_ISL_474837, EPI_ISL_474838, EPI_ISL_474839, EPI_ISL_474840 | Complejo Hospitalario Universitario de Albacete | SeqCOVID-SPAIN consortium/IBV(CSIC) | Encarnacion Simarro Córdoba, Julia Lozano Serra, Lorena Robles Fonseca , Monica Parra Grandes, Caridad Sainz de Baranda Camino and SeqCOVID-SPAIN consortium |
| EPI_ISL_474841, EPI_ISL_474842, EPI_ISL_474843, EPI_ISL_474844, EPI_ISL_474845, EPI_ISL_474846 | Hospital Universitario Virgen de las Nieves de Granada-SAS | SeqCOVID-SPAIN consortium/IBV(CSIC) | Mercedes Pérez Ruiz, Sara Sanbonmatsu Gámez, Irene Pedrosa Corral, José M. Navarro-Marí and SeqCOVID-SPAIN consortium |
| EPI_ISL_474847, EPI_ISL_474848, EPI_ISL_474849 | Complejo Hospitalario Universitario de Albacete | SeqCOVID-SPAIN consortium/IBV(CSIC) | Encarnacion Simarro Córdoba, Julia Lozano Serra, Lorena Robles Fonseca , Monica Parra Grandes, Caridad Sainz de Baranda Camino and SeqCOVID-SPAIN consortium |
| EPI_ISL_474850, EPI_ISL_474851, EPI_ISL_474852 | Hospital Universitario Virgen de las Nieves de Granada-SAS | SeqCOVID-SPAIN consortium/IBV(CSIC) | Mercedes Pérez Ruiz, Sara Sanbonmatsu Gámez, Irene Pedrosa Corral, José M. Navarro-Marí and SeqCOVID-SPAIN consortium |
| EPI_ISL_474853 | Complejo Hospitalario Universitario de Albacete | SeqCOVID-SPAIN consortium/IBV(CSIC) | Encarnacion Simarro Córdoba, Julia Lozano Serra, Lorena Robles Fonseca , Monica Parra Grandes, Caridad Sainz de Baranda Camino and SeqCOVID-SPAIN consortium |

EPI_ISL_474854, EPI_ISL_474855, EPI_ISL_474856, EPI_ISL_474857, EPI_ISL_474858, EPI_ISL_474859, EPI_ISL_474860, EPI_ISL_474861, EPI_ISL_474862, EPI_ISL_474863, EPI_ISL_474864, EPI_ISL_474865, EPI_ISL_474866, EPI_ISL_474867, EPI_ISL_474868, EPI_ISL_474869, EPI_ISL_474870, EPI_ISL_474871, EPI_ISL_474872, EPI_ISL_474873, EPI_ISL_474874, EPI_ISL_474875, EPI_ISL_474876, EPI_ISL_474877, EPI_ISL_474878, EPI_ISL_474879, EPI_ISL_474880, EPI_ISL_474881, EPI_ISL_474882, EPI_ISL_474883, EPI_ISL_474884, EPI_ISL_474885, EPI_ISL_474886, EPI_ISL_474887, EPI_ISL_474888, EPI_ISL_474889,

| | | | |
|---|---|---|---|
| EPI_ISL_474890, EPI_ISL_474891, EPI_ISL_474892, EPI_ISL_474893, EPI_ISL_474894, EPI_ISL_474895, EPI_ISL_474896, EPI_ISL_474897, EPI_ISL_474898, EPI_ISL_474899, EPI_ISL_474900 | | | |
| see above | Hospital Universitario Virgen de las Nieves de Granada-SAS | SeqCOVID-SPAIN consortium/IBV(CSIC) | Mercedes Pérez Ruiz, Sara Sanbonmatsu Gámez, Irene Pedrosa Corral, José M. Navarro-Marí and SeqCOVID-SPAIN consortium |
| EPI_ISL_474901, EPI_ISL_474902, EPI_ISL_474903, EPI_ISL_474904, EPI_ISL_474905, EPI_ISL_474906 | Complejo Hospitalario Universitario de Albacete | SeqCOVID-SPAIN consortium/IBV(CSIC) | Encarnacion Simarro Córdoba, Julia Lozano Serra, Lorena Robles Fonseca , Monica Parra Grandes, Caridad Sainz de Baranda Camino and SeqCOVID-SPAIN consortium |
| EPI_ISL_474907, EPI_ISL_474908, EPI_ISL_474909 | Hospital Universitario Virgen de las Nieves de Granada-SAS | SeqCOVID-SPAIN consortium/IBV(CSIC) | Mercedes Pérez Ruiz, Sara Sanbonmatsu Gámez, Irene Pedrosa Corral, José M. Navarro-Marí and SeqCOVID-SPAIN consortium |
| EPI_ISL_474910, EPI_ISL_474911, EPI_ISL_474912, EPI_ISL_474913, EPI_ISL_474914, EPI_ISL_474915, EPI_ISL_474916, EPI_ISL_474917, EPI_ISL_474918 | Hospital Universitario de Gran Canaria Dr. Negrín | SeqCOVID-SPAIN consortium/IBV(CSIC) | M. Carmen Pérez González, Francisco J. Chamizo López, Ana Bordes Benítez and SeqCOVID-SPAIN consortium |
| EPI_ISL_474919 | Complejo Hospitalario Universitario de Albacete | SeqCOVID-SPAIN consortium/IBV(CSIC) | Encarnacion Simarro Córdoba, Julia Lozano Serra, Lorena Robles Fonseca , Monica Parra Grandes, Caridad Sainz de Baranda Camino and SeqCOVID-SPAIN consortium |
| EPI_ISL_474920 | Hospital Universitario Virgen de las Nieves de Granada-SAS | SeqCOVID-SPAIN consortium/IBV(CSIC) | Mercedes Pérez Ruiz, Sara Sanbonmatsu Gámez, Irene Pedrosa Corral, José M. Navarro-Marí and SeqCOVID-SPAIN consortium |
| EPI_ISL_474921 | Complejo Hospitalario Universitario de Albacete | SeqCOVID-SPAIN consortium/IBV(CSIC) | Encarnacion Simarro Córdoba, Julia Lozano Serra, Lorena Robles Fonseca , Monica Parra Grandes, Caridad Sainz de Baranda Camino and SeqCOVID-SPAIN consortium |
| EPI_ISL_474922, EPI_ISL_474923, EPI_ISL_474924, EPI_ISL_474925, EPI_ISL_474926, EPI_ISL_474927, EPI_ISL_474928, EPI_ISL_474929, EPI_ISL_474930, EPI_ISL_474931, EPI_ISL_474932 | | | |
| see above | Hospital Universitario Virgen de las Nieves de Granada-SAS | SeqCOVID-SPAIN consortium/IBV(CSIC) | Mercedes Pérez Ruiz, Sara Sanbonmatsu Gámez, Irene Pedrosa Corral, José M. Navarro-Marí and SeqCOVID-SPAIN consortium |
| EPI_ISL_474933 | Complejo Hospitalario Universitario de Albacete | SeqCOVID-SPAIN consortium/IBV(CSIC) | Encarnacion Simarro Córdoba, Julia Lozano Serra, Lorena Robles Fonseca , Monica Parra Grandes, Caridad Sainz de Baranda Camino and SeqCOVID-SPAIN consortium |
| EPI_ISL_474934, EPI_ISL_474935, EPI_ISL_474936, EPI_ISL_474937, EPI_ISL_474938, EPI_ISL_474939 | Hospital Universitario Virgen de las Nieves de Granada-SAS | SeqCOVID-SPAIN consortium/IBV(CSIC) | Mercedes Pérez Ruiz, Sara Sanbonmatsu Gámez, Irene Pedrosa Corral, José M. Navarro-Marí and SeqCOVID-SPAIN consortium |
| EPI_ISL_474940, EPI_ISL_474941 | Complejo Hospitalario Universitario de Albacete | SeqCOVID-SPAIN consortium/IBV(CSIC) | Encarnacion Simarro Córdoba, Julia Lozano Serra, Lorena Robles Fonseca , Monica Parra Grandes, Caridad Sainz de Baranda Camino and SeqCOVID-SPAIN consortium |
| EPI_ISL_474942, EPI_ISL_474943, EPI_ISL_474944 | Hospital Universitario Virgen de las Nieves de Granada-SAS | SeqCOVID-SPAIN consortium/IBV(CSIC) | Mercedes Pérez Ruiz, Sara Sanbonmatsu Gámez, Irene Pedrosa Corral, José M. Navarro-Marí and SeqCOVID-SPAIN consortium |
| EPI_ISL_474945, EPI_ISL_474946, EPI_ISL_474947 | Complejo Hospitalario Universitario de Albacete | SeqCOVID-SPAIN consortium/IBV(CSIC) | Encarnacion Simarro Córdoba, Julia Lozano Serra, Lorena Robles Fonseca , Monica Parra Grandes, Caridad Sainz de Baranda Camino and SeqCOVID-SPAIN consortium |
| EPI_ISL_474948, EPI_ISL_474949, EPI_ISL_474950 | Hospital Universitario Virgen de las Nieves de Granada-SAS | SeqCOVID-SPAIN consortium/IBV(CSIC) | Mercedes Pérez Ruiz, Sara Sanbonmatsu Gámez, Irene Pedrosa Corral, José M. Navarro-Marí and SeqCOVID-SPAIN consortium |
| EPI_ISL_474951, EPI_ISL_474952, EPI_ISL_474953, EPI_ISL_474954, EPI_ISL_474955, EPI_ISL_474956 | Complejo Hospitalario Universitario de Albacete | SeqCOVID-SPAIN consortium/IBV(CSIC) | Encarnacion Simarro Córdoba, Julia Lozano Serra, Lorena Robles Fonseca , Monica Parra Grandes, Caridad Sainz de Baranda Camino and SeqCOVID-SPAIN consortium |
| EPI_ISL_474957 | Hospital Universitario Virgen de las Nieves de Granada-SAS | SeqCOVID-SPAIN consortium/IBV(CSIC) | Mercedes Pérez Ruiz, Sara Sanbonmatsu Gámez, Irene Pedrosa Corral, José M. Navarro-Marí and SeqCOVID-SPAIN consortium |
| EPI_ISL_474958 | Israeli Central Virology laboratory | Israel Central Virology laboratory | Neta Zuckerman, Efrat Dahan Bucris, Oran Erster, Ella Mendelson, Michal Mandelboim |
| EPI_ISL_474959, EPI_ISL_474960, EPI_ISL_474961, EPI_ISL_474962, EPI_ISL_474963, EPI_ISL_474964, EPI_ISL_474965, EPI_ISL_474966, EPI_ISL_474967, EPI_ISL_474968, EPI_ISL_474969, EPI_ISL_474970, EPI_ISL_474971, EPI_ISL_474972, EPI_ISL_474973, EPI_ISL_474974, EPI_ISL_474975, EPI_ISL_474976, EPI_ISL_474977, EPI_ISL_474978, EPI_ISL_474979, EPI_ISL_474980, EPI_ISL_474981, EPI_ISL_474982, EPI_ISL_474983, EPI_ISL_474984, EPI_ISL_474985, EPI_ISL_474986, EPI_ISL_474987, EPI_ISL_474988, EPI_ISL_474989, EPI_ISL_474990, EPI_ISL_474991, EPI_ISL_474992, EPI_ISL_474993, EPI_ISL_474994, EPI_ISL_474995, EPI_ISL_474996, EPI_ISL_474997, EPI_ISL_474998, EPI_ISL_474999, EPI_ISL_475000, EPI_ISL_475001, EPI_ISL_475002, EPI_ISL_475003, EPI_ISL_475004, EPI_ISL_475005, EPI_ISL_475006, EPI_ISL_475007, EPI_ISL_475008, EPI_ISL_475009, EPI_ISL_475010, EPI_ISL_475011, EPI_ISL_475012, EPI_ISL_475013, EPI_ISL_475014, EPI_ISL_475015, EPI_ISL_475016, EPI_ISL_475017, EPI_ISL_475018, EPI_ISL_475019, EPI_ISL_475020, EPI_ISL_475021, EPI_ISL_475022, EPI_ISL_475023, EPI_ISL_475024, EPI_ISL_475025 | | | |
| see above | Israel Central Virology laboratory | Israel Central Virology laboratory | Neta Zuckerman, Efrat Dahan Bucris, Oran Erster, Ella Mendelson, Michal Mandelboim |
| EPI_ISL_475026 | Banas Medical College and Research Institute | Gujarat Biotechnology Research Centre | Sunil R Joshi, Viren s Doshi, Pritesh Sabara, Apurvasinh Puvar, Janvi Raval, Zarna Patel, Monika Gandhi, Pinal Trivedi, Maharshi Pandya, Nidhi Patel, Nitin Savaliya, Raghawendra Kumar, Dinesh Kumar, Zuber Saiyed, Komal Patel, Labdhi Pandya, Snehal Bagatharia, Radhika Khara, Neha Rajpara, R D Dixit, A M Kadri, Harsh Bakshi, Chaitanya Joshi, Madhvi Joshi |
| EPI_ISL_475027 | Banas Medical College and Research Institute | Gujarat Biotechnology Research Centre | Viren s Doshi, Pritesh Sabara, Apurvasinh Puvar, Janvi Raval, Zarna Patel, Monika Gandhi, Pinal Trivedi, Maharshi Pandya, Nidhi Patel, Nitin Savaliya, Raghawendra Kumar, Dinesh Kumar, Zuber Saiyed, Komal Patel, Labdhi Pandya, Snehal Bagatharia, Radhika Khara, Sunil R Joshi, Afzal Ansari, R D Dixit, A M Kadri, Harsh Bakshi, Chaitanya Joshi, Madhvi Joshi |
| EPI_ISL_475028 | Banas Medical College and Research Institute | Gujarat Biotechnology Research Centre | Pritesh Sabara, Apurvasinh Puvar, Janvi Raval, Zarna Patel, Monika Gandhi, Pinal Trivedi, Maharshi Pandya, Nidhi Patel, Nitin Savaliya, Raghawendra Kumar, Dinesh Kumar, Zuber Saiyed, Komal Patel, Labdhi Pandya, Snehal Bagatharia, Radhika Khara, Sunil R Joshi, Viren s Doshi, Fenil Patel, R D Dixit, A M Kadri, Harsh Bakshi, Chaitanya Joshi, Madhvi Joshi |
| EPI_ISL_475029 | Banas Medical College and Research Institute | Gujarat Biotechnology Research Centre | Apurvasinh Puvar, Janvi Raval, Zarna Patel, Monika Gandhi, Pinal Trivedi, Maharshi Pandya, Nidhi Patel, Nitin Savaliya, Raghawendra Kumar, Dinesh Kumar, Zuber Saiyed, Komal Patel, Labdhi Pandya, Snehal Bagatharia, Radhika Khara, Sunil R Joshi, Viren s Doshi, Pritesh Sabara, Neelam Nathani, R D Dixit, A M Kadri, Harsh Bakshi, Chaitanya Joshi, Madhvi Joshi |
| EPI_ISL_475030 | Department of MicroBiology, Government Medical College, Surat | Gujarat Biotechnology Research Centre | Janvi Raval, Zarna Patel, Monika Gandhi, Pinal Trivedi, Maharshi Pandya, Nidhi Patel, Nitin Savaliya, Raghawendra Kumar, Dinesh Kumar, Zuber Saiyed, Komal Patel, Labdhi Pandya, Snehal Bagatharia, Naresh Chauhan, Summaiya Mullan, Amit gamit, Pritesh Sabara, Apurvasinh Puvar, Armi Chaudhari, R D Dixit, A M Kadri, Harsh Bakshi, Chaitanya Joshi, Madhvi Joshi |
| EPI_ISL_475031 | Department of MicroBiology, Government Medical College, Surat | Gujarat Biotechnology Research Centre | Zarna Patel, Monika Gandhi, Pinal Trivedi, Maharshi Pandya, Nidhi Patel, Nitin Savaliya, Raghawendra Kumar, Dinesh Kumar, Zuber Saiyed, Komal Patel, Labdhi Pandya, Snehal Bagatharia, Naresh Chauhan, Summaiya Mullan, Amit gamit, Pritesh Sabara, Apurvasinh Puvar, Janvi Raval, Bhavya Jindal, R D Dixit, A M Kadri, Harsh Bakshi, Chaitanya Joshi, Madhvi Joshi |
| EPI_ISL_475032 | Department of MicroBiology, Government Medical College, Surat | Gujarat Biotechnology Research Centre | Monika Gandhi, Pinal Trivedi, Maharshi Pandya, Nidhi Patel, Nitin Savaliya, Raghawendra Kumar, Dinesh Kumar, Zuber Saiyed, Komal Patel, Labdhi Pandya, Snehal Bagatharia, Naresh Chauhan, Summaiya Mullan, Amit gamit, Pritesh Sabara, Apurvasinh Puvar, Janvi Raval, Zarna Patel, Priyanka P Vatsa, R D Dixit, A M Kadri, Harsh Bakshi, Chaitanya Joshi, Madhvi Joshi |
| EPI_ISL_475033 | Department of MicroBiology, Government Medical College, Surat | Gujarat Biotechnology Research Centre | Pinal Trivedi, Maharshi Pandya, Nidhi Patel, Nitin Savaliya, Raghawendra Kumar, Dinesh Kumar, Zuber Saiyed, Komal Patel, Labdhi Pandya, Snehal Bagatharia, Naresh Chauhan, Summaiya Mullan, Amit gamit, Pritesh Sabara, Apurvasinh Puvar, Janvi Raval, Zarna Patel, Monika Gandhi, Pooja P Doshi, R D Dixit, A M Kadri, Harsh Bakshi, Chaitanya Joshi, Madhvi Joshi |
| EPI_ISL_475034 | Department of MicroBiology, Government Medical College, Surat | Gujarat Biotechnology Research Centre | Maharshi Pandya, Nidhi Patel, Nitin Savaliya, Raghawendra Kumar, Dinesh Kumar, Zuber Saiyed, Komal Patel, Labdhi Pandya, Snehal Bagatharia, Naresh Chauhan, Summaiya Mullan, Amit gamit, Pritesh Sabara, Apurvasinh Puvar, Janvi Raval, Zarna Patel, Monika Gandhi, Pinal Trivedi, Akanksha Verma, R D Dixit, A M Kadri, Harsh Bakshi, Chaitanya Joshi, Madhvi Joshi |
| EPI_ISL_475035 | Department of MicroBiology, Government Medical College, Surat | Gujarat Biotechnology Research Centre | Nidhi Patel, Nitin Savaliya, Raghawendra Kumar, Dinesh Kumar, Zuber Saiyed, Komal Patel, Labdhi Pandya, Snehal Bagatharia, Naresh Chauhan, Summaiya Mullan, Amit gamit, Pritesh Sabara, Apurvasinh Puvar, Janvi Raval, Zarna Patel, Monika Gandhi, Pinal Trivedi, Maharshi Pandya, Priti Pandita, R D Dixit, A M Kadri, Harsh Bakshi, Chaitanya Joshi, Madhvi Joshi |
| EPI_ISL_475036 | Department of MicroBiology, Government Medical | Gujarat Biotechnology Research Centre | Nitin Savaliya, Raghawendra Kumar, Dinesh Kumar, Zuber Saiyed, Komal Patel, Labdhi Pandya, Snehal Bagatharia, Naresh Chauhan, Summaiya Mullan, |

| | | | |
|---|---|---|---|
| | College, Surat | | Amit gamit, Pritesh Sabara, Apurvasinh Puvar, Janvi Raval, Zarna Patel, Monika Gandhi, Pinal Trivedi, Maharshi Pandya, Nidhi Patel, Pragya Sharma, R D Dixit, A M Kadri, Harsh Bakshi, Chaitanya Joshi, Madhvi Joshi |
| EPI_ISL_475037 | Department of MicroBiology, Government Medical College, Surat | Gujarat Biotechnology Research Centre | Raghawendra Kumar, Dinesh Kumar, Zuber Saiyed, Komal Patel, Labdhi Pandya, Snehal Bagatharia, Naresh Chauhan, Summaiya Mullan, Amit gamit, Pritesh Sabara, Apurvasinh Puvar, Janvi Raval, Zarna Patel, Monika Gandhi, Pinal Trivedi, Maharshi Pandya, Nidhi Patel, Nitin Savaliya, Neha Rajpara, R D Dixit, A M Kadri, Harsh Bakshi, Chaitanya Joshi, Madhvi Joshi |
| EPI_ISL_475038 | Department of MicroBiology, Government Medical College, Surat | Gujarat Biotechnology Research Centre | Dinesh Kumar, Zuber Saiyed, Komal Patel, Labdhi Pandya, Snehal Bagatharia, Naresh Chauhan, Summaiya Mullan, Amit gamit, Pritesh Sabara, Apurvasinh Puvar, Janvi Raval, Zarna Patel, Monika Gandhi, Pinal Trivedi, Maharshi Pandya, Nidhi Patel, Nitin Savaliya, Raghawendra Kumar, Afzal Ansari, R D Dixit, A M Kadri, Harsh Bakshi, Chaitanya Joshi, Madhvi Joshi |
| EPI_ISL_475039 | Department of MicroBiology, Government Medical College, Surat | Gujarat Biotechnology Research Centre | Zuber Saiyed, Komal Patel, Labdhi Pandya, Snehal Bagatharia, Naresh Chauhan, Summaiya Mullan, Amit gamit, Pritesh Sabara, Apurvasinh Puvar, Janvi Raval, Zarna Patel, Monika Gandhi, Pinal Trivedi, Maharshi Pandya, Nidhi Patel, Nitin Savaliya, Raghawendra Kumar, Dinesh Kumar, Fenil Patel, R D Dixit, A M Kadri, Harsh Bakshi, Chaitanya Joshi, Madhvi Joshi |
| EPI_ISL_475040 | Department of MicroBiology, Government Medical College, Surat | Gujarat Biotechnology Research Centre | Komal Patel, Labdhi Pandya, Snehal Bagatharia, Naresh Chauhan, Summaiya Mullan, Amit gamit, Pritesh Sabara, Apurvasinh Puvar, Janvi Raval, Zarna Patel, Monika Gandhi, Pinal Trivedi, Maharshi Pandya, Nidhi Patel, Nitin Savaliya, Raghawendra Kumar, Dinesh Kumar, Zuber Saiyed, Neelam Nathani, R D Dixit, A M Kadri, Harsh Bakshi, Chaitanya Joshi, Madhvi Joshi |
| EPI_ISL_475041 | Department of MicroBiology, Government Medical College, Surat | Gujarat Biotechnology Research Centre | Labdhi Pandya, Snehal Bagatharia, Naresh Chauhan, Summaiya Mullan, Amit gamit, Pritesh Sabara, Apurvasinh Puvar, Janvi Raval, Zarna Patel, Monika Gandhi, Pinal Trivedi, Maharshi Pandya, Nidhi Patel, Nitin Savaliya, Raghawendra Kumar, Dinesh Kumar, Zuber Saiyed, Komal Patel, Armi Chaudhari, R D Dixit, A M Kadri, Harsh Bakshi, Chaitanya Joshi, Madhvi Joshi |
| EPI_ISL_475042 | Department of MicroBiology, Government Medical College, Surat | Gujarat Biotechnology Research Centre | Snehal Bagatharia, Naresh Chauhan, Summaiya Mullan, Amit gamit, Pritesh Sabara, Apurvasinh Puvar, Janvi Raval, Zarna Patel, Monika Gandhi, Pinal Trivedi, Maharshi Pandya, Nidhi Patel, Nitin Savaliya, Raghawendra Kumar, Dinesh Kumar, Zuber Saiyed, Komal Patel, Labdhi Pandya, Bhavya Jindal, R D Dixit, A M Kadri, Harsh Bakshi, Chaitanya Joshi, Madhvi Joshi |
| EPI_ISL_475043 | Department of MicroBiology, Government Medical College, Surat | Gujarat Biotechnology Research Centre | Naresh Chauhan, Summaiya Mullan, Amit gamit, Pritesh Sabara, Apurvasinh Puvar, Janvi Raval, Zarna Patel, Monika Gandhi, Pinal Trivedi, Maharshi Pandya, Nidhi Patel, Nitin Savaliya, Raghawendra Kumar, Dinesh Kumar, Zuber Saiyed, Komal Patel, Labdhi Pandya, Snehal Bagatharia, Priyanka P Vatsa, R D Dixit, A M Kadri, Harsh Bakshi, Chaitanya Joshi, Madhvi Joshi |
| EPI_ISL_475044 | Department of MicroBiology, Government Medical College, Surat | Gujarat Biotechnology Research Centre | Summaiya Mullan, Amit gamit, Pritesh Sabara, Apurvasinh Puvar, Janvi Raval, Zarna Patel, Monika Gandhi, Pinal Trivedi, Maharshi Pandya, Nidhi Patel, Nitin Savaliya, Raghawendra Kumar, Dinesh Kumar, Zuber Saiyed, Komal Patel, Labdhi Pandya, Snehal Bagatharia, Naresh Chauhan, Pooja P Doshi, R D Dixit, A M Kadri, Harsh Bakshi, Chaitanya Joshi, Madhvi Joshi |
| EPI_ISL_475045 | Department of MicroBiology, Government Medical College, Surat | Gujarat Biotechnology Research Centre | Amit gamit, Pritesh Sabara, Apurvasinh Puvar, Janvi Raval, Zarna Patel, Monika Gandhi, Pinal Trivedi, Maharshi Pandya, Nidhi Patel, Nitin Savaliya, Raghawendra Kumar, Dinesh Kumar, Zuber Saiyed, Komal Patel, Labdhi Pandya, Snehal Bagatharia, Naresh Chauhan, Summaiya Mullan, Akanksha Verma, R D Dixit, A M Kadri, Harsh Bakshi, Chaitanya Joshi, Madhvi Joshi |
| EPI_ISL_475046 | Department of MicroBiology, Government Medical College, Surat | Gujarat Biotechnology Research Centre | Pritesh Sabara, Apurvasinh Puvar, Janvi Raval, Zarna Patel, Monika Gandhi, Pinal Trivedi, Maharshi Pandya, Nidhi Patel, Nitin Savaliya, Raghawendra Kumar, Dinesh Kumar, Zuber Saiyed, Komal Patel, Labdhi Pandya, Snehal Bagatharia, Naresh Chauhan, Summaiya Mullan, Amit gamit, Priti Pandita, R D Dixit, A M Kadri, Harsh Bakshi, Chaitanya Joshi, Madhvi Joshi |
| EPI_ISL_475047 | GMERS Medical College & Hospital | Gujarat Biotechnology Research Centre | Apurvasinh Puvar, Janvi Raval, Zarna Patel, Monika Gandhi, Pinal Trivedi, Maharshi Pandya, Nidhi Patel, Nitin Savaliya, Raghawendra Kumar, Dinesh Kumar, Zuber Saiyed, Komal Patel, Labdhi Pandya, Snehal Bagatharia, Meenakshi Shah, Neena Doshi, Varsha Godbole, Pritesh Sabara, Pragya Sharma, R D Dixit, A M Kadri, Harsh Bakshi, Chaitanya Joshi, Madhvi Joshi |
| EPI_ISL_475048 | GMERS Medical College & Hospital | Gujarat Biotechnology Research Centre | Janvi Raval, Zarna Patel, Monika Gandhi, Pinal Trivedi, Maharshi Pandya, Nidhi Patel, Nitin Savaliya, Raghawendra Kumar, Dinesh Kumar, Zuber Saiyed, Komal Patel, Labdhi Pandya, Snehal Bagatharia, Meenakshi Shah, Neena Doshi, Varsha Godbole, Pritesh Sabara, Apurvasinh Puvar, Neha Rajpara, R D Dixit, A M Kadri, Harsh Bakshi, Chaitanya Joshi, Madhvi Joshi |
| EPI_ISL_475049 | GMERS Medical College & Hospital | Gujarat Biotechnology Research Centre | Zarna Patel, Monika Gandhi, Pinal Trivedi, Maharshi Pandya, Nidhi Patel, Nitin Savaliya, Raghawendra Kumar, Dinesh Kumar, Zuber Saiyed, Komal Patel, Labdhi Pandya, Snehal Bagatharia, Meenakshi Shah, Neena Doshi, Varsha Godbole, Pritesh Sabara, Apurvasinh Puvar, Janvi Raval, Afzal Ansari, R D Dixit, A M Kadri, Harsh Bakshi, Chaitanya Joshi, Madhvi Joshi |
| EPI_ISL_475050 | GMERS Medical College & Hospital | Gujarat Biotechnology Research Centre | Monika Gandhi, Pinal Trivedi, Maharshi Pandya, Nidhi Patel, Nitin Savaliya, Raghawendra Kumar, Dinesh Kumar, Zuber Saiyed, Komal Patel, Labdhi Pandya, Snehal Bagatharia, Meenakshi Shah, Neena Doshi, Varsha Godbole, Pritesh Sabara, Apurvasinh Puvar, Janvi Raval, Zarna Patel, Fenil Patel, R D Dixit, A M Kadri, Harsh Bakshi, Chaitanya Joshi, Madhvi Joshi |
| EPI_ISL_475051 | GMERS Medical College & Hospital | Gujarat Biotechnology Research Centre | Pinal Trivedi, Maharshi Pandya, Nidhi Patel, Nitin Savaliya, Raghawendra Kumar, Dinesh Kumar, Zuber Saiyed, Komal Patel, Labdhi Pandya, Snehal Bagatharia, Meenakshi Shah, Neena Doshi, Varsha Godbole, Pritesh Sabara, Apurvasinh Puvar, Janvi Raval, Zarna Patel, Monika Gandhi, Neelam Nathani, R D Dixit, A M Kadri, Harsh Bakshi, Chaitanya Joshi, Madhvi Joshi |
| EPI_ISL_475052 | GMERS Medical College & Hospital | Gujarat Biotechnology Research Centre | Maharshi Pandya, Nidhi Patel, Nitin Savaliya, Raghawendra Kumar, Dinesh Kumar, Zuber Saiyed, Komal Patel, Labdhi Pandya, Snehal Bagatharia, Meenakshi Shah, Neena Doshi, Varsha Godbole, Pritesh Sabara, Apurvasinh Puvar, Janvi Raval, Zarna Patel, Monika Gandhi, Pinal Trivedi, Armi Chaudhari, R D Dixit, A M Kadri, Harsh Bakshi, Chaitanya Joshi, Madhvi Joshi |
| EPI_ISL_475053 | GMERS Medical College & Hospital | Gujarat Biotechnology Research Centre | Nidhi Patel, Nitin Savaliya, Raghawendra Kumar, Dinesh Kumar, Zuber Saiyed, Komal Patel, Labdhi Pandya, Snehal Bagatharia, Meenakshi Shah, Neena Doshi, Varsha Godbole, Pritesh Sabara, Apurvasinh Puvar, Janvi Raval, Zarna Patel, Monika Gandhi, Pinal Trivedi, Maharshi Pandya, Bhavya Jindal, R D Dixit, A M Kadri, Harsh Bakshi, Chaitanya Joshi, Madhvi Joshi |
| EPI_ISL_475054 | GMERS Medical College & Hospital | Gujarat Biotechnology Research Centre | Nitin Savaliya, Raghawendra Kumar, Dinesh Kumar, Zuber Saiyed, Komal Patel, Labdhi Pandya, Snehal Bagatharia, Meenakshi Shah, Neena Doshi, Varsha Godbole, Pritesh Sabara, Apurvasinh Puvar, Janvi Raval, Zarna Patel, Monika Gandhi, Pinal Trivedi, Maharshi Pandya, Nidhi Patel, Priyanka P Vatsa, R D Dixit, A M Kadri, Harsh Bakshi, Chaitanya Joshi, Madhvi Joshi |
| EPI_ISL_475055 | GMERS Medical College & Hospital | Gujarat Biotechnology Research Centre | Raghawendra Kumar, Dinesh Kumar, Zuber Saiyed, Komal Patel, Labdhi Pandya, Snehal Bagatharia, Meenakshi Shah, Neena Doshi, Varsha Godbole, Pritesh Sabara, Apurvasinh Puvar, Janvi Raval, Zarna Patel, Monika Gandhi, Pinal Trivedi, Maharshi Pandya, Nidhi Patel, Nitin Savaliya, Pooja P Doshi, R D Dixit, A M Kadri, Harsh Bakshi, Chaitanya Joshi, Madhvi Joshi |
| EPI_ISL_475056 | Dr. N. D. Desai Medical College & Hospital | Gujarat Biotechnology Research Centre | Dinesh Kumar, Zuber Saiyed, Komal Patel, Labdhi Pandya, Snehal Bagatharia, J G Buch, Jigar Gusani, Supreet Prabhu, Pritesh Sabara, Apurvasinh Puvar, Janvi Raval, Zarna Patel, Monika Gandhi, Pinal Trivedi, Maharshi Pandya, Nidhi Patel, Nitin Savaliya, Raghawendra Kumar, Akanksha Verma, R D Dixit, A M Kadri, Harsh Bakshi, Chaitanya Joshi, Madhvi Joshi |
| EPI_ISL_475057 | Dr. N. D. Desai Medical College & Hospital | Gujarat Biotechnology Research Centre | Zuber Saiyed, Komal Patel, Labdhi Pandya, Supreet Prabhu, Snehal Bagatharia, Jigar Gusani, J G Buch, Pritesh Sabara, Apurvasinh Puvar, Janvi Raval, Zarna Patel, Monika Gandhi, Pinal Trivedi, Maharshi Pandya, Nidhi Patel, Nitin Savaliya, Raghawendra Kumar, Dinesh Kumar, Priti Pandita, R D Dixit, A M Kadri, Harsh Bakshi, Chaitanya Joshi, Madhvi Joshi |
| EPI_ISL_475058 | GAIMS & G K General Hospital | Gujarat Biotechnology Research Centre | Babulal Babhoria, Hitesh Assudani, Komal Patel, Labdhi Pandya, Snehal Bagatharia, Pritesh Sabara, Apurvasinh Puvar, Janvi Raval, Zarna Patel, Monika Gandhi, Pinal Trivedi, Maharshi Pandya, Nidhi Patel, Nitin Savaliya, Raghawendra Kumar, Dinesh Kumar, Zuber Saiyed, Pragya Sharma, R D Dixit, A M Kadri, Harsh Bakshi, Chaitanya Joshi, Madhvi Joshi |
| EPI_ISL_475059 | GAIMS & G K General Hospital | Gujarat Biotechnology Research Centre | Hitesh Assudani, Babulal Babhoria, Labdhi Pandya, Snehal Bagatharia, Pritesh Sabara, Apurvasinh Puvar, Janvi Raval, Zarna Patel, Monika Gandhi, Pinal Trivedi, Maharshi Pandya, Nidhi Patel, Nitin Savaliya, Raghawendra Kumar, Dinesh Kumar, Zuber Saiyed, Komal Patel, Neha Rajpara, R D Dixit, A M Kadri, Harsh Bakshi, Chaitanya Joshi, Madhvi Joshi |
| EPI_ISL_475060, EPI_ISL_475061, EPI_ISL_475062, EPI_ISL_475063, EPI_ISL_475064, EPI_ISL_475065, EPI_ISL_475066, EPI_ISL_475067, EPI_ISL_475068, EPI_ISL_475069, EPI_ISL_475070 | | | |
| see above | Lab voor klinische biologie | Onderzoeksgroep Virologie | Laurens Lambrechts, Nick Vereecke, Marthe Pauwels, Bruno Verhasselt, Linos Vandekerckhove, Hans Nauwynck, Sebastiaan Theuns |
| EPI_ISL_475071, EPI_ISL_475072, EPI_ISL_475073, EPI_ISL_475074, EPI_ISL_475075, EPI_ISL_475076, EPI_ISL_475077, EPI_ISL_475078, EPI_ISL_475079, EPI_ISL_475080, EPI_ISL_475081, EPI_ISL_475082 | | | |

| | | | |
|---|---|---|---|
| see above | Lab voor klinische biologie | Onderzoeksgroep Virologie | Nick Vereecke, Laurens Lambrechts, Marthe Pauwels, Bruno Verhasselt, Linos Vandekerckhove, Hans Nauwynck, Sebastiaan Theuns |
| EPI_ISL_475083, EPI_ISL_475084 | National Institute of Laboratory Medicine and Referral Center | Genomic Research Lab, BCSIR | Md. Murshed Hasan Sarkar, Abu Sayeed Mohammad Mahmud, Mohammad Samir Uzzaman, Eshrar Osman, Md. Ahasan Habib, Shahina Akter, Tanjina Akhter Banu, Barna Goswami, Iffat Jahan, Md. Saddam Hossain, Tasnim Nafisa, Md. Maruf Ahmed Molla, Mahmuda Yeasmin, Asish Kumar Ghosh, Bayzid Bin Monir, A. K. M. Shamsuzzaman, Sheikh Md. Selim Al Din, Utpal Chandra Ray, Salek Ahmed Sajib, Md. Salim Khan |
| EPI_ISL_475085 | Skovde/Unilabs | The Public Health Agency of Sweden | Oskar Karlsson Lindsjo, Maria Lind Karlberg, Mattias Haukland, Reza Advani, Olov Svartstrom, Anna-Malin Linde, Sandra Broddesson, Petra Edquist, Shamam Muradrasoli, Anna Risberg, Karin Tegmark-Wisell |
| EPI_ISL_475086, EPI_ISL_475087, EPI_ISL_475088, EPI_ISL_475089, EPI_ISL_475090, EPI_ISL_475091 | Karolinska Universitetslaboratoriet | The Public Health Agency of Sweden | Oskar Karlsson Lindsjo, Maria Lind Karlberg, Mattias Haukland, Reza Advani, Olov Svartstrom, Anna-Malin Linde, Sandra Broddesson, Petra Edquist, Shamam Muradrasoli, Anna Risberg, Karin Tegmark-Wisell |
| EPI_ISL_475092 | Skovde/Unilabs | The Public Health Agency of Sweden | Oskar Karlsson Lindsjo, Maria Lind Karlberg, Mattias Haukland, Reza Advani, Olov Svartstrom, Anna-Malin Linde, Sandra Broddesson, Petra Edquist, Shamam Muradrasoli, Anna Risberg, Karin Tegmark-Wisell |
| EPI_ISL_475093, EPI_ISL_475094 | Halmstad klinisk mikrobiologi | The Public Health Agency of Sweden | Oskar Karlsson Lindsjo, Maria Lind Karlberg, Mattias Haukland, Reza Advani, Olov Svartstrom, Anna-Malin Linde, Sandra Broddesson, Petra Edquist, Shamam Muradrasoli, Anna Risberg, Karin Tegmark-Wisell |
| EPI_ISL_475095 | Karolinska Universitetslaboratoriet | The Public Health Agency of Sweden | Oskar Karlsson Lindsjo, Maria Lind Karlberg, Mattias Haukland, Reza Advani, Olov Svartstrom, Anna-Malin Linde, Sandra Broddesson, Petra Edquist, Shamam Muradrasoli, Anna Risberg, Karin Tegmark-Wisell |
| EPI_ISL_475096, EPI_ISL_475097, EPI_ISL_475098 | Halmstad klinisk mikrobiologi | The Public Health Agency of Sweden | Oskar Karlsson Lindsjo, Maria Lind Karlberg, Mattias Haukland, Reza Advani, Olov Svartstrom, Anna-Malin Linde, Sandra Broddesson, Petra Edquist, Shamam Muradrasoli, Anna Risberg, Karin Tegmark-Wisell |
| EPI_ISL_475099 | Skovde/Unilabs | The Public Health Agency of Sweden | Oskar Karlsson Lindsjo, Maria Lind Karlberg, Mattias Haukland, Reza Advani, Olov Svartstrom, Anna-Malin Linde, Sandra Broddesson, Petra Edquist, Shamam Muradrasoli, Anna Risberg, Karin Tegmark-Wisell |
| EPI_ISL_475100 | Halmstad klinisk mikrobiologi | The Public Health Agency of Sweden | Oskar Karlsson Lindsjo, Maria Lind Karlberg, Mattias Haukland, Reza Advani, Olov Svartstrom, Anna-Malin Linde, Sandra Broddesson, Petra Edquist, Shamam Muradrasoli, Anna Risberg, Karin Tegmark-Wisell |
| EPI_ISL_475101, EPI_ISL_475102, EPI_ISL_475103, EPI_ISL_475104 | Klinisk Mikrobiologi | The Public Health Agency of Sweden | Oskar Karlsson Lindsjo, Maria Lind Karlberg, Mattias Haukland, Reza Advani, Olov Svartstrom, Anna-Malin Linde, Sandra Broddesson, Petra Edquist, Shamam Muradrasoli, Anna Risberg, Karin Tegmark-Wisell |
| EPI_ISL_475105, EPI_ISL_475106, EPI_ISL_475107, EPI_ISL_475108, EPI_ISL_475109 | Skovde/Unilabs | The Public Health Agency of Sweden | Oskar Karlsson Lindsjo, Maria Lind Karlberg, Mattias Haukland, Reza Advani, Olov Svartstrom, Anna-Malin Linde, Sandra Broddesson, Petra Edquist, Shamam Muradrasoli, Anna Risberg, Karin Tegmark-Wisell |
| EPI_ISL_475110 | Gavle klinisk mikrobiologi | The Public Health Agency of Sweden | Oskar Karlsson Lindsjo, Maria Lind Karlberg, Mattias Haukland, Reza Advani, Olov Svartstrom, Anna-Malin Linde, Sandra Broddesson, Petra Edquist, Shamam Muradrasoli, Anna Risberg, Karin Tegmark-Wisell |
| EPI_ISL_475111, EPI_ISL_475112, EPI_ISL_475113 | Skovde/Unilabs | The Public Health Agency of Sweden | Oskar Karlsson Lindsjo, Maria Lind Karlberg, Mattias Haukland, Reza Advani, Olov Svartstrom, Anna-Malin Linde, Sandra Broddesson, Petra Edquist, Shamam Muradrasoli, Anna Risberg, Karin Tegmark-Wisell |
| EPI_ISL_475114 | Halmstad klinisk mikrobiologi | The Public Health Agency of Sweden | Oskar Karlsson Lindsjo, Maria Lind Karlberg, Mattias Haukland, Reza Advani, Olov Svartstrom, Anna-Malin Linde, Sandra Broddesson, Petra Edquist, Shamam Muradrasoli, Anna Risberg, Karin Tegmark-Wisell |
| EPI_ISL_475115 | Gavle klinisk mikrobiologi | The Public Health Agency of Sweden | Oskar Karlsson Lindsjo, Maria Lind Karlberg, Mattias Haukland, Reza Advani, Olov Svartstrom, Anna-Malin Linde, Sandra Broddesson, Petra Edquist, Shamam Muradrasoli, Anna Risberg, Karin Tegmark-Wisell |
| EPI_ISL_475116, EPI_ISL_475117 | Halmstad klinisk mikrobiologi | The Public Health Agency of Sweden | Oskar Karlsson Lindsjo, Maria Lind Karlberg, Mattias Haukland, Reza Advani, Olov Svartstrom, Anna-Malin Linde, Sandra Broddesson, Petra Edquist, Shamam Muradrasoli, Anna Risberg, Karin Tegmark-Wisell |
| EPI_ISL_475118, EPI_ISL_475119 | Uppsala klinisk mikrobiologi | The Public Health Agency of Sweden | Oskar Karlsson Lindsjo, Maria Lind Karlberg, Mattias Haukland, Reza Advani, Olov Svartstrom, Anna-Malin Linde, Sandra Broddesson, Petra Edquist, Shamam Muradrasoli, Anna Risberg, Karin Tegmark-Wisell |
| EPI_ISL_475120, EPI_ISL_475121 | Halmstad klinisk mikrobiologi | The Public Health Agency of Sweden | Oskar Karlsson Lindsjo, Maria Lind Karlberg, Mattias Haukland, Reza Advani, Olov Svartstrom, Anna-Malin Linde, Sandra Broddesson, Petra Edquist, Shamam Muradrasoli, Anna Risberg, Karin Tegmark-Wisell |
| EPI_ISL_475122 | Umea klinisk mikrobiologi | The Public Health Agency of Sweden | Oskar Karlsson Lindsjo, Maria Lind Karlberg, Mattias Haukland, Reza Advani, Olov Svartstrom, Anna-Malin Linde, Sandra Broddesson, Petra Edquist, Shamam Muradrasoli, Anna Risberg, Karin Tegmark-Wisell |
| EPI_ISL_475123 | Halmstad klinisk mikrobiologi | The Public Health Agency of Sweden | Oskar Karlsson Lindsjo, Maria Lind Karlberg, Mattias Haukland, Reza Advani, Olov Svartstrom, Anna-Malin Linde, Sandra Broddesson, Petra Edquist, Shamam Muradrasoli, Anna Risberg, Karin Tegmark-Wisell |
| EPI_ISL_475124 | Ostersund klinisk mikrobiologi | The Public Health Agency of Sweden | Oskar Karlsson Lindsjo, Maria Lind Karlberg, Mattias Haukland, Reza Advani, Olov Svartstrom, Anna-Malin Linde, Sandra Broddesson, Petra Edquist, Shamam Muradrasoli, Anna Risberg, Karin Tegmark-Wisell |
| EPI_ISL_475125 | Halmstad klinisk mikrobiologi | The Public Health Agency of Sweden | Oskar Karlsson Lindsjo, Maria Lind Karlberg, Mattias Haukland, Reza Advani, Olov Svartstrom, Anna-Malin Linde, Sandra Broddesson, Petra Edquist, Shamam Muradrasoli, Anna Risberg, Karin Tegmark-Wisell |
| EPI_ISL_475126, EPI_ISL_475127 | Umea klinisk mikrobiologi | The Public Health Agency of Sweden | Oskar Karlsson Lindsjo, Maria Lind Karlberg, Mattias Haukland, Reza Advani, Olov Svartstrom, Anna-Malin Linde, Sandra Broddesson, Petra Edquist, Shamam Muradrasoli, Anna Risberg, Karin Tegmark-Wisell |
| EPI_ISL_475128, EPI_ISL_475129, EPI_ISL_475130, EPI_ISL_475131, EPI_ISL_475132, EPI_ISL_475133, EPI_ISL_475134, EPI_ISL_475135 | Orebro klinisk mikrobiologi | The Public Health Agency of Sweden | Oskar Karlsson Lindsjo, Maria Lind Karlberg, Mattias Haukland, Reza Advani, Olov Svartstrom, Anna-Malin Linde, Sandra Broddesson, Petra Edquist, Shamam Muradrasoli, Anna Risberg, Karin Tegmark-Wisell |
| EPI_ISL_475136 | Karolinska Universitetslaboratoriet | The Public Health Agency of Sweden | Oskar Karlsson Lindsjo, Maria Lind Karlberg, Mattias Haukland, Reza Advani, Olov Svartstrom, Anna-Malin Linde, Sandra Broddesson, Petra Edquist, Shamam Muradrasoli, Anna Risberg, Karin Tegmark-Wisell |
| EPI_ISL_475137, EPI_ISL_475138 | Kalmar klinisk mikrobiologi | The Public Health Agency of Sweden | Oskar Karlsson Lindsjo, Maria Lind Karlberg, Mattias Haukland, Reza Advani, Olov Svartstrom, Anna-Malin Linde, Sandra Broddesson, Petra Edquist, Shamam Muradrasoli, Anna Risberg, Karin Tegmark-Wisell |
| EPI_ISL_475139 | Orebro klinisk mikrobiologi | The Public Health Agency of Sweden | Oskar Karlsson Lindsjo, Maria Lind Karlberg, Mattias Haukland, Reza Advani, Olov Svartstrom, Anna-Malin Linde, Sandra Broddesson, Petra Edquist, Shamam Muradrasoli, Anna Risberg, Karin Tegmark-Wisell |
| EPI_ISL_475140 | Skovde/Unilabs | The Public Health Agency of Sweden | Oskar Karlsson Lindsjo, Maria Lind Karlberg, Mattias Haukland, Reza Advani, Olov Svartstrom, Anna-Malin Linde, Sandra Broddesson, Petra Edquist, Shamam Muradrasoli, Anna Risberg, Karin Tegmark-Wisell |
| EPI_ISL_475141, EPI_ISL_475142 | Karolinska Universitetslaboratoriet | The Public Health Agency of Sweden | Oskar Karlsson Lindsjo, Maria Lind Karlberg, Mattias Haukland, Reza Advani, Olov Svartstrom, Anna-Malin Linde, Sandra Broddesson, Petra Edquist, Shamam Muradrasoli, Anna Risberg, Karin Tegmark-Wisell |
| EPI_ISL_475143 | Umea klinisk mikrobiologi | The Public Health Agency of Sweden | Oskar Karlsson Lindsjo, Maria Lind Karlberg, Mattias Haukland, Reza Advani, Olov Svartstrom, Anna-Malin Linde, Sandra Broddesson, Petra Edquist, Shamam Muradrasoli, Anna Risberg, Karin Tegmark-Wisell |
| EPI_ISL_475144, EPI_ISL_475145, EPI_ISL_475146, EPI_ISL_475147, EPI_ISL_475148 | Klinisk mikrobiologi Vasternorrland | The Public Health Agency of Sweden | Oskar Karlsson Lindsjo, Maria Lind Karlberg, Mattias Haukland, Reza Advani, Olov Svartstrom, Anna-Malin Linde, Sandra Broddesson, Petra Edquist, Shamam Muradrasoli, Anna Risberg, Karin Tegmark-Wisell |
| EPI_ISL_475149, EPI_ISL_475150 | Karolinska Universitetslaboratoriet | The Public Health Agency of Sweden | Oskar Karlsson Lindsjo, Maria Lind Karlberg, Mattias Haukland, Reza Advani, Olov Svartstrom, Anna-Malin Linde, Sandra Broddesson, Petra Edquist, Shamam Muradrasoli, Anna Risberg, Karin Tegmark-Wisell |
| EPI_ISL_475151 | Uppsala klinisk mikrobiologi | The Public Health Agency of Sweden | Oskar Karlsson Lindsjo, Maria Lind Karlberg, Mattias Haukland, Reza Advani, Olov Svartstrom, Anna-Malin Linde, Sandra Broddesson, Petra Edquist, Shamam Muradrasoli, Anna Risberg, Karin Tegmark-Wisell |
| EPI_ISL_475152 | Folkhalsomyndigheten | The Public Health Agency of Sweden | Oskar Karlsson Lindsjo, Maria Lind Karlberg, Mattias Haukland, Reza Advani, Olov Svartstrom, Anna-Malin Linde, Sandra Broddesson, Petra Edquist, Shamam Muradrasoli, Anna Risberg, Karin Tegmark-Wisell |

| | | | |
|---|---|---|---|
| EPI_ISL_475153, EPI_ISL_475154, EPI_ISL_475155 | Klinisk mikrobiologi Vasternorrland | The Public Health Agency of Sweden | Oskar Karlsson Lindsjo, Maria Lind Karlberg, Mattias Haukland, Reza Advani, Olov Svartstrom, Anna-Malin Linde, Sandra Broddesson, Petra Edquist, Shamam Muradrasoli, Anna Risberg, Karin Tegmark-Wisell |
| EPI_ISL_475156, EPI_ISL_475157, EPI_ISL_475158, EPI_ISL_475159 | Halmstad klinisk mikrobiologi | The Public Health Agency of Sweden | Oskar Karlsson Lindsjo, Maria Lind Karlberg, Mattias Haukland, Reza Advani, Olov Svartstrom, Anna-Malin Linde, Sandra Broddesson, Petra Edquist, Shamam Muradrasoli, Anna Risberg, Karin Tegmark-Wisell |
| EPI_ISL_475160, EPI_ISL_475161, EPI_ISL_475162, EPI_ISL_475163 | Klinisk mikrobiologi Vasternorrland | The Public Health Agency of Sweden | Oskar Karlsson Lindsjo, Maria Lind Karlberg, Mattias Haukland, Reza Advani, Olov Svartstrom, Anna-Malin Linde, Sandra Broddesson, Petra Edquist, Shamam Muradrasoli, Anna Risberg, Karin Tegmark-Wisell |
| EPI_ISL_475164 | Halmstad klinisk mikrobiologi | The Public Health Agency of Sweden | Oskar Karlsson Lindsjo, Maria Lind Karlberg, Mattias Haukland, Reza Advani, Olov Svartstrom, Anna-Malin Linde, Sandra Broddesson, Petra Edquist, Shamam Muradrasoli, Anna Risberg, Karin Tegmark-Wisell |
| EPI_ISL_475165 | National Institute of Laboratory Medicine and Referral Center | Genomic Research Lab, BCSIR | Shahina Akter, Abu Sayeed Mohammad Mahmud, Mohammad Samir Uzzaman, Eshrar Osman, Md. Ahasan Habib, Tanjina Akhter Banu, Md. Murshed Hasan Sarkar, Barna Goswami, Iffat Jahan, Md. Saddam Hossain, Tasnim Nafisa, Md. Maruf Ahmed Molla, Mahmuda Yeasmin, Asish Kumar Ghosh, Bayzid Bin Monir, A. K. M. Shamsuzzaman, Sheikh Md. Selim Al Din, Utpal Chandra Ray, Salek Ahmed Sajib, Md. Salim Khan |
| EPI_ISL_475166 | National Institute of Laboratory Medicine and Referral Center | Genomic Research Lab, BCSIR | Tanjina Akhter Banu, Abu Sayeed Mohammad Mahmud, Mohammad Samir Uzzaman, Eshrar Osman, Md. Ahasan Habib, Shahina Akter, Md. Murshed Hasan Sarkar, Barna Goswami, Iffat Jahan, Md. Saddam Hossain, Tasnim Nafisa, Md. Maruf Ahmed Molla, Mahmuda Yeasmin, Asish Kumar Ghosh, Bayzid Bin Monir, A. K. M. Shamsuzzaman, Sheikh Md. Selim Al Din, Utpal Chandra Ray, Salek Ahmed Sajib, Md. Salim Khan |
| EPI_ISL_475167 | National Institute of Laboratory Medicine and Referral Center | Genomic Research Lab, BCSIR | Barna Goswami, Abu Sayeed Mohammad Mahmud, Mohammad Samir Uzzaman, Eshrar Osman, Md. Ahasan Habib, Shahina Akter, Tanjina Akhter Banu, Md. Murshed Hasan Sarkar, Iffat Jahan, Md. Saddam Hossain, Tasnim Nafisa, Md. Maruf Ahmed Molla, Mahmuda Yeasmin, Asish Kumar Ghosh, Bayzid Bin Monir, A. K. M. Shamsuzzaman, Sheikh Md. Selim Al Din, Utpal Chandra Ray, Salek Ahmed Sajib, Md. Salim Khan |
| EPI_ISL_475168 | National Institute of Laboratory Medicine and Referral Center | Genomic Research Lab, BCSIR | Iffat Jahan, Abu Sayeed Mohammad Mahmud, Mohammad Samir Uzzaman, Eshrar Osman, Md. Ahasan Habib, Shahina Akter, Tanjina Akhter Banu, Md. Murshed Hasan Sarkar, Barna Goswami, Md. Saddam Hossain, Tasnim Nafisa, Md. Maruf Ahmed Molla, Mahmuda Yeasmin, Asish Kumar Ghosh, Bayzid Bin Monir, A. K. M. Shamsuzzaman, Sheikh Md. Selim Al Din, Utpal Chandra Ray, Salek Ahmed Sajib, Md. Salim Khan |
| EPI_ISL_475169 | National Institute of Laboratory Medicine and Referral Center | Genomic Research Lab, BCSIR | Md. Saddam Hossain, Abu Sayeed Mohammad Mahmud, Mohammad Samir Uzzaman, Eshrar Osman, Md. Ahasan Habib, Shahina Akter, Tanjina Akhter Banu, Md. Murshed Hasan Sarkar, Barna Goswami, Iffat Jahan, Tasnim Nafisa, Md. Maruf Ahmed Molla, Mahmuda Yeasmin, Asish Kumar Ghosh, Bayzid Bin Monir, A. K. M. Shamsuzzaman, Sheikh Md. Selim Al Din, Utpal Chandra Ray, Salek Ahmed Sajib, Md. Salim Khan |
| EPI_ISL_475170, EPI_ISL_475171, EPI_ISL_475172, EPI_ISL_475173 | National Institute of Laboratory Medicine and Referral Center | Genomic Research Lab, BCSIR | Abu Sayeed Mohammad Mahmud, Mohammad Samir Uzzaman, Eshrar Osman, Md. Ahasan Habib, Shahina Akter, Tanjina Akhter Banu, Md. Murshed Hasan Sarkar, Barna Goswami, Iffat Jahan, Md. Saddam Hossain, Tasnim Nafisa, Md. Maruf Ahmed Molla, Mahmuda Yeasmin, Asish Kumar Ghosh, Bayzid Bin Monir, A. K. M. Shamsuzzaman, Sheikh Md. Selim Al Din, Utpal Chandra Ray, Salek Ahmed Sajib, Md. Salim Khan |
| EPI_ISL_475174, EPI_ISL_475175, EPI_ISL_475176, EPI_ISL_475177, EPI_ISL_475178, EPI_ISL_475179, EPI_ISL_475180, EPI_ISL_475181, EPI_ISL_475182, EPI_ISL_475183, EPI_ISL_475184, EPI_ISL_475185, EPI_ISL_475186, EPI_ISL_475187, EPI_ISL_475188, EPI_ISL_475189, EPI_ISL_475190, EPI_ISL_475191, EPI_ISL_475192, EPI_ISL_475193, EPI_ISL_475194, EPI_ISL_475195, EPI_ISL_475196, EPI_ISL_475197, EPI_ISL_475198, EPI_ISL_475199, EPI_ISL_475200, EPI_ISL_475201, EPI_ISL_475202, EPI_ISL_475203, EPI_ISL_475204, EPI_ISL_475205, EPI_ISL_475206, EPI_ISL_475207, EPI_ISL_475208, EPI_ISL_475209, EPI_ISL_475210, EPI_ISL_475211, EPI_ISL_475212, EPI_ISL_475213, EPI_ISL_475214, EPI_ISL_475215, EPI_ISL_475216, EPI_ISL_475217, EPI_ISL_475218, EPI_ISL_475219, EPI_ISL_475220, EPI_ISL_475221, EPI_ISL_475222, EPI_ISL_475223, EPI_ISL_475224, EPI_ISL_475225, EPI_ISL_475226, EPI_ISL_475227, EPI_ISL_475228, EPI_ISL_475229, EPI_ISL_475230, EPI_ISL_475231, EPI_ISL_475232, EPI_ISL_475233, EPI_ISL_475234, EPI_ISL_475235, EPI_ISL_475236, EPI_ISL_475237 | | | |
| see above | Nebraska Public Health Laboratory | UNMC COVID-19 Response Team | UNMC COVID-19 Response Team |
| EPI_ISL_475238 | National Institute of Laboratory Medicine and Referral Center | Genomic Research Lab, BCSIR | Abu Sayeed Mohammad Mahmud, Mohammad Samir Uzzaman, Eshrar Osman, Md. Ahasan Habib, Shahina Akter, Tanjina Akhter Banu, Md. Murshed Hasan Sarkar, Barna Goswami, Iffat Jahan, Md. Saddam Hossain, Tasnim Nafisa, Md. Maruf Ahmed Molla, Mahmuda Yeasmin, Asish Kumar Ghosh, Bayzid Bin Monir, A. K. M. Shamsuzzaman, Sheikh Md. Selim Al Din, Utpal Chandra Ray, Salek Ahmed Sajib, Md. Salim Khan |
| EPI_ISL_475239, EPI_ISL_475240, EPI_ISL_475241, EPI_ISL_475242, EPI_ISL_475243, EPI_ISL_475244, EPI_ISL_475245, EPI_ISL_475246, EPI_ISL_475247, EPI_ISL_475248, EPI_ISL_475249, EPI_ISL_475250, EPI_ISL_475251, EPI_ISL_475252, EPI_ISL_475253, EPI_ISL_475254, EPI_ISL_475255, EPI_ISL_475256, EPI_ISL_475257, EPI_ISL_475258, EPI_ISL_475259, EPI_ISL_475260, EPI_ISL_475261, EPI_ISL_475262, EPI_ISL_475263, EPI_ISL_475264, EPI_ISL_475265, EPI_ISL_475266, EPI_ISL_475267, EPI_ISL_475268, EPI_ISL_475269, EPI_ISL_475270, EPI_ISL_475271, EPI_ISL_475272, EPI_ISL_475273, EPI_ISL_475274, EPI_ISL_475275, EPI_ISL_475276, EPI_ISL_475277, EPI_ISL_475278, EPI_ISL_475279, EPI_ISL_475280, EPI_ISL_475281, EPI_ISL_475282, EPI_ISL_475283, EPI_ISL_475284, EPI_ISL_475285, EPI_ISL_475286, EPI_ISL_475287, EPI_ISL_475288, EPI_ISL_475289, EPI_ISL_475290, EPI_ISL_475291, EPI_ISL_475292, EPI_ISL_475293, EPI_ISL_475294, EPI_ISL_475295, EPI_ISL_475296, EPI_ISL_475297, EPI_ISL_475298, EPI_ISL_475299, EPI_ISL_475300, EPI_ISL_475301, EPI_ISL_475302, EPI_ISL_475303, EPI_ISL_475304, EPI_ISL_475305, EPI_ISL_475306, EPI_ISL_475307, EPI_ISL_475308, EPI_ISL_475309, EPI_ISL_475310, EPI_ISL_475311, EPI_ISL_475312, EPI_ISL_475313, EPI_ISL_475314, EPI_ISL_475315, EPI_ISL_475316, EPI_ISL_475317, EPI_ISL_475318, EPI_ISL_475319, EPI_ISL_475320, EPI_ISL_475321, EPI_ISL_475322, EPI_ISL_475323, EPI_ISL_475324, EPI_ISL_475325, EPI_ISL_475326, EPI_ISL_475327, EPI_ISL_475328, EPI_ISL_475329, EPI_ISL_475330, EPI_ISL_475331, EPI_ISL_475332, EPI_ISL_475333, EPI_ISL_475334, EPI_ISL_475335, EPI_ISL_475336, EPI_ISL_475337, EPI_ISL_475338, EPI_ISL_475339, EPI_ISL_475340, EPI_ISL_475341 | | | |
| see above | Centre for Enzyme Innovation, University of Portsmouth / Translational Research Laboratory, Portsmouth Hospitals NHS Trust | COVID-19 Genomics UK (COG-UK) Consortium | Angela Beckett,Yann Bourgeois,Garry Scarlett,Sharon Glaysher,Scott Elliott,Kelly Bicknell,Robert Impey,Allyson Lloyd,Sarah Wyllie,Ethan Butcher,Anoop Chauhan,Samuel Robson |
| EPI_ISL_475342, EPI_ISL_475343, EPI_ISL_475344, EPI_ISL_475345, EPI_ISL_475346, EPI_ISL_475347, EPI_ISL_475348, EPI_ISL_475349, EPI_ISL_475350, EPI_ISL_475351, EPI_ISL_475352, EPI_ISL_475353, EPI_ISL_475354, EPI_ISL_475355, EPI_ISL_475356, EPI_ISL_475357, EPI_ISL_475358, EPI_ISL_475359, EPI_ISL_475360, EPI_ISL_475361, EPI_ISL_475362, EPI_ISL_475363, EPI_ISL_475364, EPI_ISL_475365, EPI_ISL_475366, EPI_ISL_475367, EPI_ISL_475368, EPI_ISL_475369, EPI_ISL_475370, EPI_ISL_475371, EPI_ISL_475372, EPI_ISL_475373, EPI_ISL_475374, EPI_ISL_475375, EPI_ISL_475376, EPI_ISL_475377, EPI_ISL_475378, EPI_ISL_475379, EPI_ISL_475380, EPI_ISL_475381, EPI_ISL_475382, EPI_ISL_475383, EPI_ISL_475384, EPI_ISL_475385, EPI_ISL_475386, EPI_ISL_475387, EPI_ISL_475388, EPI_ISL_475389, EPI_ISL_475390, EPI_ISL_475391, EPI_ISL_475392, EPI_ISL_475393, EPI_ISL_475394, EPI_ISL_475395, EPI_ISL_475396, EPI_ISL_475397, EPI_ISL_475398, EPI_ISL_475399, EPI_ISL_475400, EPI_ISL_475401, EPI_ISL_475402, EPI_ISL_475403, EPI_ISL_475404, EPI_ISL_475405, EPI_ISL_475406, EPI_ISL_475407, EPI_ISL_475408, EPI_ISL_475409, EPI_ISL_475410, EPI_ISL_475411, EPI_ISL_475412, EPI_ISL_475413, EPI_ISL_475414, EPI_ISL_475415, EPI_ISL_475416, EPI_ISL_475417, EPI_ISL_475418, EPI_ISL_475419, EPI_ISL_475420, EPI_ISL_475421, EPI_ISL_475422, EPI_ISL_475423, EPI_ISL_475424, EPI_ISL_475425, EPI_ISL_475426, EPI_ISL_475427, EPI_ISL_475428, EPI_ISL_475429, EPI_ISL_475430, EPI_ISL_475431, EPI_ISL_475432, EPI_ISL_475433, EPI_ISL_475434, EPI_ISL_475435, EPI_ISL_475436, EPI_ISL_475437, EPI_ISL_475438, EPI_ISL_475439, EPI_ISL_475440, EPI_ISL_475441, EPI_ISL_475442, EPI_ISL_475443, EPI_ISL_475444, EPI_ISL_475445, EPI_ISL_475446, EPI_ISL_475447, EPI_ISL_475448, EPI_ISL_475449, EPI_ISL_475450, EPI_ISL_475451, EPI_ISL_475452, EPI_ISL_475453, EPI_ISL_475454, EPI_ISL_475455, EPI_ISL_475456, EPI_ISL_475457, EPI_ISL_475458, EPI_ISL_475459, EPI_ISL_475460, EPI_ISL_475461, EPI_ISL_475462, EPI_ISL_475463, EPI_ISL_475464, EPI_ISL_475465, EPI_ISL_475466, EPI_ISL_475467, EPI_ISL_475468, EPI_ISL_475469, EPI_ISL_475470, EPI_ISL_475471, EPI_ISL_475472, EPI_ISL_475473, EPI_ISL_475474, EPI_ISL_475475, EPI_ISL_475476, EPI_ISL_475477, EPI_ISL_475478, EPI_ISL_475479, EPI_ISL_475480, EPI_ISL_475481, EPI_ISL_475482, EPI_ISL_475483, EPI_ISL_475484, EPI_ISL_475485, EPI_ISL_475486, EPI_ISL_475487, EPI_ISL_475488, EPI_ISL_475489, EPI_ISL_475490, EPI_ISL_475491, EPI_ISL_475492, EPI_ISL_475493, EPI_ISL_475494, EPI_ISL_475495, EPI_ISL_475496, EPI_ISL_475497, EPI_ISL_475498, EPI_ISL_475499, EPI_ISL_475500, EPI_ISL_475501, EPI_ISL_475502, EPI_ISL_475503, EPI_ISL_475504, EPI_ISL_475505, EPI_ISL_475506, EPI_ISL_475507, EPI_ISL_475508, EPI_ISL_475509, EPI_ISL_475510 | | | |
| see above | Virology Department, Sheffield Teaching Hospitals NHS Foundation Trust/Department of Infection, Immunity and Cardiovascular Disease, The Medical School, University of Sheffield | COVID-19 Genomics UK (COG-UK) Consortium | Thushan de Silva, Matthew Parker, Nikki Smith, Adri Angyal, Rebecca Brown, Luke Green, Rachel Tucker, Paul Parsons, Danielle Groves, Katie Johnson, Laura Carrilero, Alex Keeley, Dave Partridge, Matthew Wyles, Benjamin Lindsey, Mehmet Yavuz, Mohammad Raza, Cariad Evans |
| EPI_ISL_475511 | Orestadsklinikens VC | The Public Health Agency of Sweden | Oskar Karlsson Lindsjo, Maria Lind Karlberg, Mattias Haukland, Reza Advani, Olov Svartstrom, Anna-Malin Linde, Sandra Broddesson, Mia Brytting, Anna Risberg, Karin Tegmark-Wisell |
| EPI_ISL_475512 | Din Klinik | The Public Health Agency of Sweden | Oskar Karlsson Lindsjo, Maria Lind Karlberg, Mattias Haukland, Reza Advani, Olov Svartstrom, Anna-Malin Linde, Sandra Broddesson, Mia Brytting, Anna Risberg, Karin Tegmark-Wisell |
| EPI_ISL_475513 | Huddinge VC | The Public Health Agency of Sweden | Oskar Karlsson Lindsjo, Maria Lind Karlberg, Mattias Haukland, Reza Advani, Olov Svartstrom, Anna-Malin Linde, Sandra Broddesson, Mia Brytting, Anna Risberg, Karin Tegmark-Wisell |
| EPI_ISL_475514 | Uppsala Narakut Aleris | The Public Health Agency of Sweden | Oskar Karlsson Lindsjo, Maria Lind Karlberg, Mattias Haukland, Reza Advani, Olov Svartstrom, Anna-Malin Linde, Sandra Broddesson, Mia Brytting, Anna Risberg, Karin Tegmark-Wisell |
| EPI_ISL_475515 | Lakargruppen | The Public Health Agency of Sweden | Oskar Karlsson Lindsjo, Maria Lind Karlberg, Mattias Haukland, Reza Advani, Olov Svartstrom, Anna-Malin Linde, Sandra Broddesson, Mia Brytting, Anna Risberg, Karin Tegmark-Wisell |
| EPI_ISL_475516, EPI_ISL_475517 | Uppsala Narakut Aleris | The Public Health Agency of Sweden | Oskar Karlsson Lindsjo, Maria Lind Karlberg, Mattias Haukland, Reza Advani, Olov Svartstrom, Anna-Malin Linde, Sandra Broddesson, Mia Brytting, Anna Risberg, Karin Tegmark-Wisell |
| EPI_ISL_475518 | Trollbackens VC | The Public Health Agency of Sweden | Oskar Karlsson Lindsjo, Maria Lind Karlberg, Mattias Haukland, Reza Advani, Olov Svartstrom, Anna-Malin Linde, Sandra Broddesson, Mia Brytting, Anna Risberg, Karin Tegmark-Wisell |
| EPI_ISL_475519 | Orsa VC | The Public Health Agency of Sweden | Oskar Karlsson Lindsjo, Maria Lind Karlberg, Mattias Haukland, Reza Advani, Olov Svartstrom, Anna-Malin Linde, Sandra Broddesson, Mia Brytting, Anna Risberg, Karin Tegmark-Wisell |
| EPI_ISL_475520 | Vardcentralen Brinken | The Public Health Agency of Sweden | Oskar Karlsson Lindsjo, Maria Lind Karlberg, Mattias Haukland, Reza Advani, Olov Svartstrom, Anna-Malin Linde, Sandra Broddesson, Mia Brytting, Anna Risberg, Karin Tegmark-Wisell |

| | | | |
|---|---|---|---|
| EPI_ISL_475521 | Ulltuna Vardcentral | The Public Health Agency of Sweden | Oskar Karlsson Lindsjo, Maria Lind Karlberg, Mattias Haukland, Reza Advani, Olov Svartstrom, Anna-Malin Linde, Sandra Broddesson, Mia Brytting, Anna Risberg, Karin Tegmark-Wisell |
| EPI_ISL_475522, EPI_ISL_475523 | Huddinge VC | The Public Health Agency of Sweden | Oskar Karlsson Lindsjo, Maria Lind Karlberg, Mattias Haukland, Reza Advani, Olov Svartstrom, Anna-Malin Linde, Sandra Broddesson, Mia Brytting, Anna Risberg, Karin Tegmark-Wisell |
| EPI_ISL_475524 | Narhalsan Sjobo vardcentral | The Public Health Agency of Sweden | Oskar Karlsson Lindsjo, Maria Lind Karlberg, Mattias Haukland, Reza Advani, Olov Svartstrom, Anna-Malin Linde, Sandra Broddesson, Mia Brytting, Anna Risberg, Karin Tegmark-Wisell |
| EPI_ISL_475525 | Huddinge VC | The Public Health Agency of Sweden | Oskar Karlsson Lindsjo, Maria Lind Karlberg, Mattias Haukland, Reza Advani, Olov Svartstrom, Anna-Malin Linde, Sandra Broddesson, Mia Brytting, Anna Risberg, Karin Tegmark-Wisell |
| EPI_ISL_475526, EPI_ISL_475527 | Uppsala Narakut Aleris | The Public Health Agency of Sweden | Oskar Karlsson Lindsjo, Maria Lind Karlberg, Mattias Haukland, Reza Advani, Olov Svartstrom, Anna-Malin Linde, Sandra Broddesson, Mia Brytting, Anna Risberg, Karin Tegmark-Wisell |
| EPI_ISL_475528 | Omtanken Grimmered | The Public Health Agency of Sweden | Oskar Karlsson Lindsjo, Maria Lind Karlberg, Mattias Haukland, Reza Advani, Olov Svartstrom, Anna-Malin Linde, Sandra Broddesson, Mia Brytting, Anna Risberg, Karin Tegmark-Wisell |
| EPI_ISL_475529, EPI_ISL_475530, EPI_ISL_475531, EPI_ISL_475532 | Kungsors VC | The Public Health Agency of Sweden | Oskar Karlsson Lindsjo, Maria Lind Karlberg, Mattias Haukland, Reza Advani, Olov Svartstrom, Anna-Malin Linde, Sandra Broddesson, Mia Brytting, Anna Risberg, Karin Tegmark-Wisell |
| EPI_ISL_475533, EPI_ISL_475534, EPI_ISL_475535 | Omtanken Grimmered | The Public Health Agency of Sweden | Oskar Karlsson Lindsjo, Maria Lind Karlberg, Mattias Haukland, Reza Advani, Olov Svartstrom, Anna-Malin Linde, Sandra Broddesson, Mia Brytting, Anna Risberg, Karin Tegmark-Wisell |
| EPI_ISL_475536 | Follinge Halsocentral | The Public Health Agency of Sweden | Oskar Karlsson Lindsjo, Maria Lind Karlberg, Mattias Haukland, Reza Advani, Olov Svartstrom, Anna-Malin Linde, Sandra Broddesson, Mia Brytting, Anna Risberg, Karin Tegmark-Wisell |
| EPI_ISL_475537 | Narhalsan Oden VC | The Public Health Agency of Sweden | Oskar Karlsson Lindsjo, Maria Lind Karlberg, Mattias Haukland, Reza Advani, Olov Svartstrom, Anna-Malin Linde, Sandra Broddesson, Mia Brytting, Anna Risberg, Karin Tegmark-Wisell |
| EPI_ISL_475538 | Omtanken Grimmered | The Public Health Agency of Sweden | Oskar Karlsson Lindsjo, Maria Lind Karlberg, Mattias Haukland, Reza Advani, Olov Svartstrom, Anna-Malin Linde, Sandra Broddesson, Mia Brytting, Anna Risberg, Karin Tegmark-Wisell |
| EPI_ISL_475539 | Narhalsan Sjobo vardcentral | The Public Health Agency of Sweden | Oskar Karlsson Lindsjo, Maria Lind Karlberg, Mattias Haukland, Reza Advani, Olov Svartstrom, Anna-Malin Linde, Sandra Broddesson, Mia Brytting, Anna Risberg, Karin Tegmark-Wisell |
| EPI_ISL_475540 | Bla Kustens halsocentral | The Public Health Agency of Sweden | Oskar Karlsson Lindsjo, Maria Lind Karlberg, Mattias Haukland, Reza Advani, Olov Svartstrom, Anna-Malin Linde, Sandra Broddesson, Mia Brytting, Anna Risberg, Karin Tegmark-Wisell |
| EPI_ISL_475541 | Follinge Halsocentral | The Public Health Agency of Sweden | Oskar Karlsson Lindsjo, Maria Lind Karlberg, Mattias Haukland, Reza Advani, Olov Svartstrom, Anna-Malin Linde, Sandra Broddesson, Mia Brytting, Anna Risberg, Karin Tegmark-Wisell |
| EPI_ISL_475542 | Kungsholmsdoktorn | The Public Health Agency of Sweden | Oskar Karlsson Lindsjo, Maria Lind Karlberg, Mattias Haukland, Reza Advani, Olov Svartstrom, Anna-Malin Linde, Sandra Broddesson, Mia Brytting, Anna Risberg, Karin Tegmark-Wisell |
| EPI_ISL_475543 | Surbrunns VC | The Public Health Agency of Sweden | Oskar Karlsson Lindsjo, Maria Lind Karlberg, Mattias Haukland, Reza Advani, Olov Svartstrom, Anna-Malin Linde, Sandra Broddesson, Mia Brytting, Anna Risberg, Karin Tegmark-Wisell |
| EPI_ISL_475544, EPI_ISL_475545, EPI_ISL_475546, EPI_ISL_475547 | Karolinska Universitetslaboratoriet | The Public Health Agency of Sweden | Oskar Karlsson Lindsjo, Maria Lind Karlberg, Mattias Haukland, Reza Advani, Olov Svartstrom, Anna-Malin Linde, Sandra Broddesson, Shaman Muradrasoli, Anna Risberg, Karin Tegmark-Wisell |
| EPI_ISL_475548 | Halmstad klinisk mikrobiologi | The Public Health Agency of Sweden | Oskar Karlsson Lindsjo, Maria Lind Karlberg, Mattias Haukland, Reza Advani, Olov Svartstrom, Anna-Malin Linde, Sandra Broddesson, Shaman Muradrasoli, Anna Risberg, Karin Tegmark-Wisell |
| EPI_ISL_475549, EPI_ISL_475550 | Skovde/Unilabs | The Public Health Agency of Sweden | Oskar Karlsson Lindsjo, Maria Lind Karlberg, Mattias Haukland, Reza Advani, Olov Svartstrom, Anna-Malin Linde, Sandra Broddesson, Shaman Muradrasoli, Anna Risberg, Karin Tegmark-Wisell |
| EPI_ISL_475551, EPI_ISL_475552 | Karolinska Universitetslaboratoriet | The Public Health Agency of Sweden | Oskar Karlsson Lindsjo, Maria Lind Karlberg, Mattias Haukland, Reza Advani, Olov Svartstrom, Anna-Malin Linde, Sandra Broddesson, Shaman Muradrasoli, Anna Risberg, Karin Tegmark-Wisell |
| EPI_ISL_475553 | Halmstad klinisk mikrobiologi | The Public Health Agency of Sweden | Oskar Karlsson Lindsjo, Maria Lind Karlberg, Mattias Haukland, Reza Advani, Olov Svartstrom, Anna-Malin Linde, Sandra Broddesson, Shaman Muradrasoli, Anna Risberg, Karin Tegmark-Wisell |
| EPI_ISL_475554, EPI_ISL_475555 | Skovde/Unilabs | The Public Health Agency of Sweden | Oskar Karlsson Lindsjo, Maria Lind Karlberg, Mattias Haukland, Reza Advani, Olov Svartstrom, Anna-Malin Linde, Sandra Broddesson, Shaman Muradrasoli, Anna Risberg, Karin Tegmark-Wisell |
| EPI_ISL_475556, EPI_ISL_475557 | Halmstad klinisk mikrobiologi | The Public Health Agency of Sweden | Oskar Karlsson Lindsjo, Maria Lind Karlberg, Mattias Haukland, Reza Advani, Olov Svartstrom, Anna-Malin Linde, Sandra Broddesson, Shaman Muradrasoli, Anna Risberg, Karin Tegmark-Wisell |
| EPI_ISL_475558, EPI_ISL_475559, EPI_ISL_475560, EPI_ISL_475561 | Karolinska Universitetslaboratoriet | The Public Health Agency of Sweden | Oskar Karlsson Lindsjo, Maria Lind Karlberg, Mattias Haukland, Reza Advani, Olov Svartstrom, Anna-Malin Linde, Sandra Broddesson, Shaman Muradrasoli, Anna Risberg, Karin Tegmark-Wisell |
| EPI_ISL_475562, EPI_ISL_475563 | Din Klinik | The Public Health Agency of Sweden | Oskar Karlsson Lindsjo, Maria Lind Karlberg, Mattias Haukland, Reza Advani, Olov Svartstrom, Anna-Malin Linde, Sandra Broddesson, Mia Brytting, Anna Risberg, Karin Tegmark-Wisell |
| EPI_ISL_475564 | Surbrunns VC | The Public Health Agency of Sweden | Oskar Karlsson Lindsjo, Maria Lind Karlberg, Mattias Haukland, Reza Advani, Olov Svartstrom, Anna-Malin Linde, Sandra Broddesson, Mia Brytting, Anna Risberg, Karin Tegmark-Wisell |
| EPI_ISL_475565 | Narhalsan Backa vardcentral | The Public Health Agency of Sweden | Oskar Karlsson Lindsjo, Maria Lind Karlberg, Mattias Haukland, Reza Advani, Olov Svartstrom, Anna-Malin Linde, Sandra Broddesson, Mia Brytting, Anna Risberg, Karin Tegmark-Wisell |
| EPI_ISL_475566 | Vardcentralen Brinken | The Public Health Agency of Sweden | Oskar Karlsson Lindsjo, Maria Lind Karlberg, Mattias Haukland, Reza Advani, Olov Svartstrom, Anna-Malin Linde, Sandra Broddesson, Mia Brytting, Anna Risberg, Karin Tegmark-Wisell |
| EPI_ISL_475567 | Huddinge VC | The Public Health Agency of Sweden | Oskar Karlsson Lindsjo, Maria Lind Karlberg, Mattias Haukland, Reza Advani, Olov Svartstrom, Anna-Malin Linde, Sandra Broddesson, Mia Brytting, Anna Risberg, Karin Tegmark-Wisell |
| EPI_ISL_475568 | Kungsors VC | The Public Health Agency of Sweden | Oskar Karlsson Lindsjo, Maria Lind Karlberg, Mattias Haukland, Reza Advani, Olov Svartstrom, Anna-Malin Linde, Sandra Broddesson, Mia Brytting, Anna Risberg, Karin Tegmark-Wisell |
| EPI_ISL_475569 | Kungsholmsdoktorn | The Public Health Agency of Sweden | Oskar Karlsson Lindsjo, Maria Lind Karlberg, Mattias Haukland, Reza Advani, Olov Svartstrom, Anna-Malin Linde, Sandra Broddesson, Mia Brytting, Anna Risberg, Karin Tegmark-Wisell |
| EPI_ISL_475570 | Genome Center | Genome Center | A. S. M. Rubayet- Ul- Alam, Ovinu Kibria Islam, Md. Shazid Hasan, Hassan M. Al-Emran, Shireen Nigar, Selina Akter, Pravas Chandra Roy, Md. Tanvir Islam, Shovon Lal Sarkar, M. Shaminur Rahman, M. Rafiul Islam, Habiba Ibnat, Md Nur Kabidul Azam, Chakrabortty Atonu, Proshanto Kumar Das, Md. Hasan al Pramanik, Md. Zannat Ali, Shohanur Rahaman, Md. Aminul Islam, Ashok Kumar, Md. Nazmul Hasan, Md. Iqbal Kabir Jahid, Md. Anwar Hossain |
| EPI_ISL_475571 | Genome Center | Genome Center | Hassan M. Al-Emran, Md. Shazid Hasan, Ovinu Kibria Islam, A. S. M. Rubayet- Ul- Alam, Pravas Chandra Roy, Selina Akter, Shireen Nigar, Shovon Lal Sarkar, Md. Tanvir Islam, Mithun Talukder Md. Tawyabur, Md. Taijul Islam, Provakar Mondol, Md. Muzahidul Islam, Md. Iqbal Kabir Jahid Md. Anwar Hossain |
| EPI_ISL_475572 | Imperial College London | Imperial College London | Jie Zhou, Wendy Barclay |
| EPI_ISL_475573 | Genome Center | Genome Center | Md. Shazid Hasan, Hassan M. Al-Emran, Ovinu Kibria Islam, A. S. M. Rubayet- Ul- Alam, Selina Akter, Shireen Nigar, Md. Tanvir Islam, Pravas Chandra |

| | | | |
|---|---|---|---|
| | | | Roy, Shovon Lal Sarkar, Md. Nazmul Hasan, Tanay Chakrovarty, Md. Ali Ahasan Setu, Sourav Dutta, Ruhul Amin, Md. Iqbal Kabir Jahid, Md. Anwar Hossain |
| EPI_ISL_475574, EPI_ISL_475575, EPI_ISL_475576, EPI_ISL_475577, EPI_ISL_475578, EPI_ISL_475579, EPI_ISL_475580, EPI_ISL_475581, EPI_ISL_475582, EPI_ISL_475583, EPI_ISL_475584, EPI_ISL_475585, EPI_ISL_475586, EPI_ISL_475587, EPI_ISL_475588, EPI_ISL_475589, EPI_ISL_475590, EPI_ISL_475591, EPI_ISL_475592, EPI_ISL_475593, EPI_ISL_475594, EPI_ISL_475595, EPI_ISL_475596, EPI_ISL_475597, EPI_ISL_475598, EPI_ISL_475599, EPI_ISL_475600, EPI_ISL_475601, EPI_ISL_475602, EPI_ISL_475603, EPI_ISL_475604, EPI_ISL_475605, EPI_ISL_475606, EPI_ISL_475607, EPI_ISL_475608, EPI_ISL_475609, EPI_ISL_475610, EPI_ISL_475611, EPI_ISL_475612, EPI_ISL_475613, EPI_ISL_475614, EPI_ISL_475615, EPI_ISL_475616, EPI_ISL_475617, EPI_ISL_475618, EPI_ISL_475619, EPI_ISL_475620, EPI_ISL_475621, EPI_ISL_475622, EPI_ISL_475623, EPI_ISL_475624, EPI_ISL_475625, EPI_ISL_475626, EPI_ISL_475627, EPI_ISL_475628, EPI_ISL_475629, EPI_ISL_475630, EPI_ISL_475631, EPI_ISL_475632, EPI_ISL_475633, EPI_ISL_475634, EPI_ISL_475635, EPI_ISL_475636, EPI_ISL_475637, EPI_ISL_475638, EPI_ISL_475639, EPI_ISL_475640, EPI_ISL_475641, EPI_ISL_475642, EPI_ISL_475643, EPI_ISL_475644, EPI_ISL_475645, EPI_ISL_475646, EPI_ISL_475647, EPI_ISL_475648, EPI_ISL_475649, EPI_ISL_475650, EPI_ISL_475651, EPI_ISL_475652, EPI_ISL_475653, EPI_ISL_475654, EPI_ISL_475655, EPI_ISL_475656, EPI_ISL_475657, EPI_ISL_475658, EPI_ISL_475659, EPI_ISL_475660, EPI_ISL_475661, EPI_ISL_475662, EPI_ISL_475663, EPI_ISL_475664, EPI_ISL_475665, EPI_ISL_475666, EPI_ISL_475667, EPI_ISL_475668, EPI_ISL_475669, EPI_ISL_475670, EPI_ISL_475671, EPI_ISL_475672, EPI_ISL_475673, EPI_ISL_475674, EPI_ISL_475675, EPI_ISL_475676, EPI_ISL_475677, EPI_ISL_475678, EPI_ISL_475679, EPI_ISL_475680, EPI_ISL_475681, EPI_ISL_475682, EPI_ISL_475683, EPI_ISL_475684, EPI_ISL_475685, EPI_ISL_475686, EPI_ISL_475687, EPI_ISL_475688, EPI_ISL_475689, EPI_ISL_475690, EPI_ISL_475691, EPI_ISL_475692, EPI_ISL_475693, EPI_ISL_475694, EPI_ISL_475695, EPI_ISL_475696, EPI_ISL_475697, EPI_ISL_475698, EPI_ISL_475699, EPI_ISL_475700, EPI_ISL_475701, EPI_ISL_475702, EPI_ISL_475703, EPI_ISL_475704, EPI_ISL_475705, EPI_ISL_475706, EPI_ISL_475707, EPI_ISL_475708, EPI_ISL_475709, EPI_ISL_475710, EPI_ISL_475711, EPI_ISL_475712, EPI_ISL_475713, EPI_ISL_475714, EPI_ISL_475715, EPI_ISL_475716 | | | |
| see above | Cedars-Sinai Medical Center, Department of Pathology & Laboratory Medicine, Molecular Pathology Laboratory | Cedars-Sinai Medical Center, Molecular Pathology Laboratory of Department of Pathology & Laboratory Medicine and Genomic Core | Wenjuan Zhang, John Paul Govindavari, Brian Davis, Stephanie Chen, Jong Taek Kim, Jianbo Song, Jean Lopategui, Jasmine T Plummer, Eric Vail |
| EPI_ISL_475717, EPI_ISL_475718, EPI_ISL_475719, EPI_ISL_475720, EPI_ISL_475721 | unknown | Microbiology | Cilla,G., Montes,M., Pineiro,L., Marimon,J.M. |
| EPI_ISL_475723, EPI_ISL_475724 | unknown | Cancer Biology Department | Zekri,A.N., Amer,K.E., Ahmed,O.S., Soliman,H.K., Hafez,M.M., Bahnassy,A.A., Abdelhamid,W., Khattab,A., Ali,M., Hassan,W., Samir,M., Raouf,A., Hamdy,M.S., Soliman,M.S., Elsissy,M.H., Elkhateeb,S.M., Ezzelarab,M.H., Abouelhoda,M. |
| EPI_ISL_475725, EPI_ISL_475726, EPI_ISL_475727, EPI_ISL_475728, EPI_ISL_475729, EPI_ISL_475730, EPI_ISL_475731, EPI_ISL_475732, EPI_ISL_475733, EPI_ISL_475734, EPI_ISL_475735, EPI_ISL_475736, EPI_ISL_475737, EPI_ISL_475738, EPI_ISL_475739, EPI_ISL_475740, EPI_ISL_475741, EPI_ISL_475742, EPI_ISL_475743, EPI_ISL_475744 | | | |
| see above | Utah Public Health Laboratory | Utah Public Health Laboratory | Erin Young, Kelly Oakeson |
| EPI_ISL_475745, EPI_ISL_475746, EPI_ISL_475747, EPI_ISL_475748, EPI_ISL_475749, EPI_ISL_475750, EPI_ISL_475751, EPI_ISL_475752, EPI_ISL_475753 | Medical Ain Shams Research Institute (MASRI), Ain Shams University | Medical Ain Shams Research Institute (MASRI), Ain Shams University | Hesham Elghazaly , Sara Hassan Agwa, Mahmoud Elmeteini , Ahmad Moustafa , Ashraf Omar, Osama Mansour, Samia Abdo, Hala Hafez, Ghada Ismael , Shaimaa Moustafa , Aya Mohamed, Reham Mamdouh , Hoda Abd Elsatar, Manal Hamdy Elsaid, Fatma Ebied |
| EPI_ISL_475754 | National Institute of Laboratory Medicine and Referral Center | Genomic Research Lab, BCSIR | Shahina Akter, Abu Sayeed Mohammad Mahmud, Mohammad Samir Uzzaman, Eshrar Osman, Md. Ahasan Habib, Tanjina Akhter Banu, Md. Murshed Hasan Sarkar, Barna Goswami, Iffat Jahan, Md. Saddam Hossain, Tasnim Nafisa, Md. Maruf Ahmed Molla, Mahmuda Yeasmin, Asish Kumar Ghosh, Arifa Akram, A. K. M. Shamsuzzaman, Sheikh Md. Selim Al Din, Utpal Chandra Ray, Salek Ahmed Sajib, Md. Salim Khan |
| EPI_ISL_475755 | National Institute of Laboratory Medicine and Referral Center | Genomic Research Lab, BCSIR | Md. Murshed Hasan Sarkar, Abu Sayeed Mohammad Mahmud, Mohammad Samir Uzzaman, Eshrar Osman, Md. Ahasan Habib, Shahina Akter, Tanjina Akhter Banu, Barna Goswami, Iffat Jahan, Md. Saddam Hossain, Tasnim Nafisa, Md. Maruf Ahmed Molla, Mahmuda Yeasmin, Asish Kumar Ghosh, Arifa Akram, A. K. M. Shamsuzzaman, Sheikh Md. Selim Al Din, Utpal Chandra Ray, Salek Ahmed Sajib, Md. Salim Khan |
| EPI_ISL_475756 | National Institute of Laboratory Medicine and Referral Center | Genomic Research Lab, BCSIR | Tanjina Akhter Banu, Abu Sayeed Mohammad Mahmud, Mohammad Samir Uzzaman, Eshrar Osman, Md. Ahasan Habib, Shahina Akter, Md. Murshed Hasan Sarkar, Barna Goswami, Iffat Jahan, Md. Saddam Hossain, Tasnim Nafisa, Md. Maruf Ahmed Molla, Mahmuda Yeasmin, Asish Kumar Ghosh, Arifa Akram, A. K. M. Shamsuzzaman, Sheikh Md. Selim Al Din, Utpal Chandra Ray, Salek Ahmed Sajib, Md. Salim Khan |
| EPI_ISL_475757 | National Institute of Laboratory Medicine and Referral Center | Genomic Research Lab, BCSIR | Barna Goswami, Abu Sayeed Mohammad Mahmud, Mohammad Samir Uzzaman, Eshrar Osman, Md. Ahasan Habib, Shahina Akter, Tanjina Akhter Banu, Md. Murshed Hasan Sarkar, Iffat Jahan, Md. Saddam Hossain, Tasnim Nafisa, Md. Maruf Ahmed Molla, Mahmuda Yeasmin, Asish Kumar Ghosh, Arifa Akram, A. K. M. Shamsuzzaman, Sheikh Md. Selim Al Din, Utpal Chandra Ray, Salek Ahmed Sajib, Md. Salim Khan |
| EPI_ISL_475758 | National Institute of Laboratory Medicine and Referral Center | Genomic Research Lab, BCSIR | Iffat Jahan, Abu Sayeed Mohammad Mahmud, Mohammad Samir Uzzaman, Eshrar Osman, Md. Ahasan Habib, Shahina Akter, Tanjina Akhter Banu, Md. Murshed Hasan Sarkar, Barna Goswami, Md. Saddam Hossain, Tasnim Nafisa, Md. Maruf Ahmed Molla, Mahmuda Yeasmin, Asish Kumar Ghosh, Arifa Akram, A. K. M. Shamsuzzaman, Sheikh Md. Selim Al Din, Utpal Chandra Ray, Salek Ahmed Sajib, Md. Salim Khan |
| EPI_ISL_475759 | National Institute of Laboratory Medicine and Referral Center | Genomic Research Lab, BCSIR | Md. Saddam Hossain, Abu Sayeed Mohammad Mahmud, Mohammad Samir Uzzaman, Eshrar Osman, Md. Ahasan Habib, Shahina Akter, Tanjina Akhter Banu, Md. Murshed Hasan Sarkar, Barna Goswami, Iffat Jahan, Tasnim Nafisa, Md. Maruf Ahmed Molla, Mahmuda Yeasmin, Asish Kumar Ghosh, Arifa Akram, A. K. M. Shamsuzzaman, Sheikh Md. Selim Al Din, Utpal Chandra Ray, Salek Ahmed Sajib, Md. Salim Khan |
| EPI_ISL_475760, EPI_ISL_475761 | National Institute of Laboratory Medicine and Referral Center | Genomic Research Lab, BCSIR | Abu Sayeed Mohammad Mahmud, Mohammad Samir Uzzaman, Eshrar Osman, Md. Ahasan Habib, Shahina Akter, Tanjina Akhter Banu, Md. Murshed Hasan Sarkar, Barna Goswami, Iffat Jahan, Md. Saddam Hossain, Tasnim Nafisa, Md. Maruf Ahmed Molla, Mahmuda Yeasmin, Asish Kumar Ghosh, Arifa Akram, A. K. M. Shamsuzzaman, Sheikh Md. Selim Al Din, Utpal Chandra Ray, Salek Ahmed Sajib, Md. Salim Khan |
| EPI_ISL_475762 | Oklahoma State Department of Health | França Lab | Caio Martinelle B. de França, Graham Wiley, Samuel T. Dunn, and Matthew J. Miller. |
| EPI_ISL_475763 | Institut für Virologie am Department für Hygiene, Mikrobiologie und Public Health | Bergthaler laboratory, CeMM Research Center for Molecular Medicine of the Austrian Academy of Sciences | Alexandra Popa, Benedikt Agerer, Henrique Colaco, Lukas Endler, Jakob-Wendelin Genger, Alexander Lercher, Mark Smyth, Thomas Penz, Michael Schuster, Jan Laine, Martin Senekowitsch, Judith Aberle, Stephan Aberle, Peter Hufnagl, Daniela Schmid, Franz Allerberger, Elisabeth Puchhammer-Stoeckl, Manfred Nairz, Guenter Weiss, Gregor Hörmann, Kinga Rigler-Hohenwarter, Rainer Gattringer, Wegene Borena, Dorothee von Laer, Christoph Bock, Andreas Bergthaler |
| EPI_ISL_475764, EPI_ISL_475765, EPI_ISL_475766, EPI_ISL_475767, EPI_ISL_475768 | Universitaetsklinik für Innere Medizin II Innsbruck | Bergthaler laboratory, CeMM Research Center for Molecular Medicine of the Austrian Academy of Sciences | Alexandra Popa, Benedikt Agerer, Henrique Colaco, Lukas Endler, Jakob-Wendelin Genger, Alexander Lercher, Mark Smyth, Thomas Penz, Michael Schuster, Jan Laine, Martin Senekowitsch, Judith Aberle, Stephan Aberle, Peter Hufnagl, Daniela Schmid, Franz Allerberger, Elisabeth Puchhammer-Stoeckl, Manfred Nairz, Guenter Weiss, Gregor Hörmann, Kinga Rigler-Hohenwarter, Rainer Gattringer, Wegene Borena, Dorothee von Laer, Christoph Bock, Andreas Bergthaler |
| EPI_ISL_475769 | Institut für Virologie am Department für Hygiene, Mikrobiologie und Public Health | Bergthaler laboratory, CeMM Research Center for Molecular Medicine of the Austrian Academy of Sciences | Alexandra Popa, Benedikt Agerer, Henrique Colaco, Lukas Endler, Jakob-Wendelin Genger, Alexander Lercher, Mark Smyth, Thomas Penz, Michael Schuster, Jan Laine, Martin Senekowitsch, Judith Aberle, Stephan Aberle, Peter Hufnagl, Daniela Schmid, Franz Allerberger, Elisabeth Puchhammer-Stoeckl, Manfred Nairz, Guenter Weiss, Gregor Hörmann, Kinga Rigler-Hohenwarter, Rainer Gattringer, Wegene Borena, Dorothee von Laer, Christoph Bock, Andreas Bergthaler |
| EPI_ISL_475770, EPI_ISL_475771, EPI_ISL_475772, EPI_ISL_475773, EPI_ISL_475774, EPI_ISL_475775, EPI_ISL_475776, EPI_ISL_475777, EPI_ISL_475778, EPI_ISL_475779, EPI_ISL_475780, EPI_ISL_475781, EPI_ISL_475782, EPI_ISL_475783, EPI_ISL_475784, EPI_ISL_475785, EPI_ISL_475786, EPI_ISL_475787, EPI_ISL_475788, EPI_ISL_475789, EPI_ISL_475790, EPI_ISL_475791, EPI_ISL_475792, EPI_ISL_475793, EPI_ISL_475794, EPI_ISL_475795, EPI_ISL_475796, EPI_ISL_475797, EPI_ISL_475798, EPI_ISL_475799, EPI_ISL_475800, EPI_ISL_475801, EPI_ISL_475802, EPI_ISL_475803, EPI_ISL_475804, EPI_ISL_475805, EPI_ISL_475806, EPI_ISL_475807, EPI_ISL_475808, EPI_ISL_475809, EPI_ISL_475810, EPI_ISL_475811, EPI_ISL_475812 | | | |
| see above | Center for Virology, Medical University of Vienna | Bergthaler laboratory, CeMM Research Center for Molecular Medicine of the Austrian Academy of Sciences | Alexandra Popa, Benedikt Agerer, Henrique Colaco, Lukas Endler, Jakob-Wendelin Genger, Alexander Lercher, Mark Smyth, Thomas Penz, Michael Schuster, Jan Laine, Martin Senekowitsch, Judith Aberle, Stephan Aberle, Peter Hufnagl, Daniela Schmid, Franz Allerberger, Elisabeth Puchhammer-Stoeckl, Manfred Nairz, Guenter Weiss, Gregor Hörmann, Kinga Rigler-Hohenwarter, Rainer Gattringer, Wegene Borena, Dorothee von Laer, Christoph Bock, Andreas Bergthaler |
| EPI_ISL_475813, EPI_ISL_475814, EPI_ISL_475815, EPI_ISL_475816, EPI_ISL_475817, EPI_ISL_475818, EPI_ISL_475819, EPI_ISL_475820, EPI_ISL_475821, EPI_ISL_475822, EPI_ISL_475823, EPI_ISL_475824, EPI_ISL_475825, EPI_ISL_475826, EPI_ISL_475827, EPI_ISL_475828, EPI_ISL_475829 | | | |
| see above | Institut für Virologie am Department für Hygiene, Mikrobiologie und Public Health | Bergthaler laboratory, CeMM Research Center for Molecular Medicine of the Austrian Academy of Sciences | Alexandra Popa, Benedikt Agerer, Henrique Colaco, Lukas Endler, Jakob-Wendelin Genger, Alexander Lercher, Mark Smyth, Thomas Penz, Michael Schuster, Jan Laine, Martin Senekowitsch, Judith Aberle, Stephan Aberle, Peter Hufnagl, Daniela Schmid, Franz Allerberger, Elisabeth Puchhammer-Stoeckl, Manfred Nairz, Guenter Weiss, Gregor Hörmann, Kinga Rigler-Hohenwarter, Rainer Gattringer, Wegene Borena, Dorothee von Laer, Christoph Bock, Andreas Bergthaler |
| EPI_ISL_475830, EPI_ISL_475831, EPI_ISL_475832, EPI_ISL_475833, EPI_ISL_475834, EPI_ISL_475835, EPI_ISL_475836, EPI_ISL_475837, EPI_ISL_475838, EPI_ISL_475839, EPI_ISL_475840, EPI_ISL_475841, EPI_ISL_475842, EPI_ISL_475843, EPI_ISL_475844, EPI_ISL_475845, EPI_ISL_475846, EPI_ISL_475847, EPI_ISL_475848, EPI_ISL_475849, EPI_ISL_475850, EPI_ISL_475851, EPI_ISL_475852, EPI_ISL_475853, EPI_ISL_475854, EPI_ISL_475855, EPI_ISL_475856, EPI_ISL_475857, EPI_ISL_475858, EPI_ISL_475859, EPI_ISL_475860, EPI_ISL_475861, EPI_ISL_475862, EPI_ISL_475863, EPI_ISL_475864, EPI_ISL_475865, EPI_ISL_475866, EPI_ISL_475867, EPI_ISL_475868, EPI_ISL_475869, EPI_ISL_475870, EPI_ISL_475871, EPI_ISL_475872, EPI_ISL_475873, EPI_ISL_475874, EPI_ISL_475875, EPI_ISL_475876, EPI_ISL_475877, EPI_ISL_475878, EPI_ISL_475879, EPI_ISL_475880, EPI_ISL_475881, EPI_ISL_475882, EPI_ISL_475883, EPI_ISL_475884, EPI_ISL_475885, EPI_ISL_475886 | | | |
| see above | Austrian Agency for Health and Food Safety (AGES) | Bergthaler laboratory, CeMM Research Center for Molecular Medicine of the Austrian Academy of | Alexandra Popa, Benedikt Agerer, Henrique Colaco, Lukas Endler, Jakob-Wendelin Genger, Alexander Lercher, Mark Smyth, Thomas Penz, Michael Schuster, Jan Laine, Martin Senekowitsch, Judith Aberle, Stephan Aberle, Peter Hufnagl, Daniela Schmid, Franz Allerberger, Elisabeth |

| | | | |
|---|---|---|---|
| | | Sciences | Puchhammer-Stoeckl, Manfred Nairz, Guenter Weiss, Gregor Hörmann, Kinga Rigler-Hohenwarter, Rainer Gattringer, Wegene Borena, Dorothee von Laer, Christoph Bock, Andreas Bergthaler |
| EPI_ISL_475887, EPI_ISL_475888, EPI_ISL_475889, EPI_ISL_475890, EPI_ISL_475891, EPI_ISL_475892, EPI_ISL_475893, EPI_ISL_475894, EPI_ISL_475895, EPI_ISL_475896, EPI_ISL_475897, EPI_ISL_475898, EPI_ISL_475899, EPI_ISL_475900, EPI_ISL_475901, EPI_ISL_475902, EPI_ISL_475903, EPI_ISL_475904, EPI_ISL_475905, EPI_ISL_475906, EPI_ISL_475907, EPI_ISL_475908, EPI_ISL_475909 | | | |
| see above | Zentralinstitut für medizinische und chemische Labordiagnostik, Universitätskliniken Insbruck | Bergthaler laboratory, CeMM Research Center for Molecular Medicine of the Austrian Academy of Sciences | Alexandra Popa, Benedikt Agerer, Henrique Colaco, Lukas Endler, Jakob-Wendelin Genger, Alexander Lercher, Mark Smyth, Thomas Penz, Michael Schuster, Jan Laine, Martin Senekowitsch, Judith Aberle, Stephan Aberle, Peter Hufnagl, Daniela Schmid, Franz Allerberger, Elisabeth Puchhammer-Stoeckl, Manfred Nairz, Guenter Weiss, Gregor Hörmann, Kinga Rigler-Hohenwarter, Rainer Gattringer, Wegene Borena, Dorothee von Laer, Christoph Bock, Andreas Bergthaler |
| EPI_ISL_475910, EPI_ISL_475911, EPI_ISL_475912, EPI_ISL_475913, EPI_ISL_475914, EPI_ISL_475915 | Klinikum Wels-Grieskirchen | Bergthaler laboratory, CeMM Research Center for Molecular Medicine of the Austrian Academy of Sciences | Alexandra Popa, Benedikt Agerer, Henrique Colaco, Lukas Endler, Jakob-Wendelin Genger, Alexander Lercher, Mark Smyth, Thomas Penz, Michael Schuster, Jan Laine, Martin Senekowitsch, Judith Aberle, Stephan Aberle, Peter Hufnagl, Daniela Schmid, Franz Allerberger, Elisabeth Puchhammer-Stoeckl, Manfred Nairz, Guenter Weiss, Gregor Hörmann, Kinga Rigler-Hohenwarter, Rainer Gattringer, Wegene Borena, Dorothee von Laer, Christoph Bock, Andreas Bergthaler |
| EPI_ISL_475916, EPI_ISL_475917, EPI_ISL_475918, EPI_ISL_475919, EPI_ISL_475920, EPI_ISL_475921, EPI_ISL_475922, EPI_ISL_475923, EPI_ISL_475924, EPI_ISL_475925, EPI_ISL_475926, EPI_ISL_475927, EPI_ISL_475928 | | | |
| see above | Institut für Virologie am Department für Hygiene, Mikrobiologie und Public Health | Bergthaler laboratory, CeMM Research Center for Molecular Medicine of the Austrian Academy of Sciences | Alexandra Popa, Benedikt Agerer, Henrique Colaco, Lukas Endler, Jakob-Wendelin Genger, Alexander Lercher, Mark Smyth, Thomas Penz, Michael Schuster, Jan Laine, Martin Senekowitsch, Judith Aberle, Stephan Aberle, Peter Hufnagl, Daniela Schmid, Franz Allerberger, Elisabeth Puchhammer-Stoeckl, Manfred Nairz, Guenter Weiss, Gregor Hörmann, Kinga Rigler-Hohenwarter, Rainer Gattringer, Wegene Borena, Dorothee von Laer, Christoph Bock, Andreas Bergthaler |
| EPI_ISL_475929, EPI_ISL_475930, EPI_ISL_475931, EPI_ISL_475932, EPI_ISL_475933, EPI_ISL_475934, EPI_ISL_475935, EPI_ISL_475936 | Universitaetsklinik für Innere Medizin II Innsbruck | Bergthaler laboratory, CeMM Research Center for Molecular Medicine of the Austrian Academy of Sciences | Alexandra Popa, Benedikt Agerer, Henrique Colaco, Lukas Endler, Jakob-Wendelin Genger, Alexander Lercher, Mark Smyth, Thomas Penz, Michael Schuster, Jan Laine, Martin Senekowitsch, Judith Aberle, Stephan Aberle, Peter Hufnagl, Daniela Schmid, Franz Allerberger, Elisabeth Puchhammer-Stoeckl, Manfred Nairz, Guenter Weiss, Gregor Hörmann, Kinga Rigler-Hohenwarter, Rainer Gattringer, Wegene Borena, Dorothee von Laer, Christoph Bock, Andreas Bergthaler |
| EPI_ISL_475937, EPI_ISL_475938, EPI_ISL_475939, EPI_ISL_475940, EPI_ISL_475941, EPI_ISL_475942, EPI_ISL_475943, EPI_ISL_475944, EPI_ISL_475945, EPI_ISL_475946, EPI_ISL_475947, EPI_ISL_475948, EPI_ISL_475949, EPI_ISL_475950, EPI_ISL_475951, EPI_ISL_475952, EPI_ISL_475953, EPI_ISL_475954, EPI_ISL_475955, EPI_ISL_475956, EPI_ISL_475957, EPI_ISL_475958, EPI_ISL_475959, EPI_ISL_475960, EPI_ISL_475961, EPI_ISL_475962, EPI_ISL_475963, EPI_ISL_475964, EPI_ISL_475965, EPI_ISL_475966, EPI_ISL_475967, EPI_ISL_475968, EPI_ISL_475969, EPI_ISL_475970, EPI_ISL_475971, EPI_ISL_475972, EPI_ISL_475973, EPI_ISL_475974, EPI_ISL_475975, EPI_ISL_475976, EPI_ISL_475977, EPI_ISL_475978, EPI_ISL_475979, EPI_ISL_475980, EPI_ISL_475981, EPI_ISL_475982, EPI_ISL_475983, EPI_ISL_475984, EPI_ISL_475985, EPI_ISL_475986, EPI_ISL_475987, EPI_ISL_475988, EPI_ISL_475989, EPI_ISL_475990, EPI_ISL_475991, EPI_ISL_475992, EPI_ISL_475993, EPI_ISL_475994, EPI_ISL_475995, EPI_ISL_475996, EPI_ISL_475997, EPI_ISL_475998, EPI_ISL_475999 | | | |
| see above | National Public Health Laboratory, National Centre for Infectious Diseases | National Public Health Laboratory, National Centre for Infectious Diseases | Mak TM, Octavia S, Chavatte JM, Cui L, Lin RTP |
| EPI_ISL_476018, EPI_ISL_476019, EPI_ISL_476020, EPI_ISL_476021 | Washington University in St. Louis | Washington University in St. Louis | David Wang, Carey-Ann Burnham, Scott Handley, Lindsay Droit, Stephan Tahan |
| EPI_ISL_476022 | Defence Research & Development Establishment | Defence Research & Development Establishment | Shashi Sharma, Paban Kumar Dash, Jyoti S Kumar, Sushil Kumar Sharma, Ambuj Shrivastava |
| EPI_ISL_476023 | Defence Research & Development Establishment (DRDE) | Defence Research & Development Establishment (DRDE) | Shashi Sharma, Paban Kumar Dash, Sushil Kumar Sharma, Ambuj Shrivastava, Jyoti S. Kumar |
| EPI_ISL_476024 | Laboratoire de Recherche et d'Analyses Médicales de la Gendarmerie Royale | Laboratoire de Recherche et d'Analyses Médicales de la Gendarmerie Royale | Sanaâ Lemriss, Amal SOUIRI, Nabil Lemzaoui, Omar Mestoui, Mohamed Labioui, Nabil Ouaariba, Ayoub Jibjibe, Mahmoud Yartaoui, Mohamed Chahmi, Marouane El Rhouila, Samiha Sellak, Nadia Kandoussi, Saâd El Kabbaj. |
| EPI_ISL_476025 | Laboratoire de Recherche et d'Analyses Médicales de la Gendarmerie Royale | Laboratoire de Recherche et d'Analyses Médicales de la Gendarmerie Royale | Sanaâ LEMRISS, Amal Souiri, Saâd EL KABBAJ |
| EPI_ISL_476026 | Laboratoire de Recherche et d'Analyses Médicales de la Gendarmerie Royale | Laboratoire de Recherche et d'Analyses Médicales de la Gendarmerie Royale | Sanaâ Lemriss, Amal SOUIRI, Saâd EL KABBAJ |
| EPI_ISL_476027, EPI_ISL_476028, EPI_ISL_476029, EPI_ISL_476030, EPI_ISL_476031, EPI_ISL_476032, EPI_ISL_476033, EPI_ISL_476034, EPI_ISL_476035, EPI_ISL_476036, EPI_ISL_476037, EPI_ISL_476038, EPI_ISL_476039, EPI_ISL_476040, EPI_ISL_476041, EPI_ISL_476042, EPI_ISL_476043, EPI_ISL_476044, EPI_ISL_476045, EPI_ISL_476046, EPI_ISL_476047, EPI_ISL_476048, EPI_ISL_476049, EPI_ISL_476050, EPI_ISL_476051, EPI_ISL_476052, EPI_ISL_476053, EPI_ISL_476054, EPI_ISL_476055, EPI_ISL_476056, EPI_ISL_476057, EPI_ISL_476058, EPI_ISL_476059, EPI_ISL_476060, EPI_ISL_476061, EPI_ISL_476062, EPI_ISL_476063, EPI_ISL_476064, EPI_ISL_476065, EPI_ISL_476066 | | | |
| see above | Michigan Department of Health and Human Services, Bureau of Laboratories | Michigan Department of Health and Human Services, Bureau of Laboratories | Blankenship HM, Riner D, Soehnlen MK |
| EPI_ISL_476067 | The National Institute of Public Health | State Veterinary Institute Prague and The National Institute of Public Health | Nagy,A;Jirincova,H;Novakova,L;Trnka,D;Vecerova,J |
| EPI_ISL_476068, EPI_ISL_476069, EPI_ISL_476070, EPI_ISL_476071, EPI_ISL_476072, EPI_ISL_476073, EPI_ISL_476074, EPI_ISL_476075, EPI_ISL_476076, EPI_ISL_476077 | University of Debrecen, Department of Medical Microbiology | National Laboratory of Virology, Szentágothai Research Centre | Endre Gábor Tóth, Balázs Somogyi, Brigitta Zana, Eszter Csoma, Ferenc Jakab, Gábor Kemenesi |
| EPI_ISL_476078 | University of Szeged, Institute of Clinical Microbiology | National Laboratory of Virology, Szentágothai Research Centre | Endre Gábor Tóth, Balázs Somogyi, Brigitta Zana, Terhes Gabriella, Ferenc Jakab, Gábor Kemenesi |
| EPI_ISL_476079, EPI_ISL_476080, EPI_ISL_476081, EPI_ISL_476082, EPI_ISL_476083, EPI_ISL_476084, EPI_ISL_476085, EPI_ISL_476086, EPI_ISL_476087, EPI_ISL_476088, EPI_ISL_476089, EPI_ISL_476090, EPI_ISL_476091, EPI_ISL_476092, EPI_ISL_476093, EPI_ISL_476094, EPI_ISL_476095, EPI_ISL_476096, EPI_ISL_476097, EPI_ISL_476098, EPI_ISL_476099, EPI_ISL_476100, EPI_ISL_476101, EPI_ISL_476102, EPI_ISL_476103, EPI_ISL_476104, EPI_ISL_476105, EPI_ISL_476106, EPI_ISL_476107, EPI_ISL_476108, EPI_ISL_476109, EPI_ISL_476110, EPI_ISL_476111, EPI_ISL_476112, EPI_ISL_476113, EPI_ISL_476114, EPI_ISL_476115, EPI_ISL_476116, EPI_ISL_476117, EPI_ISL_476118, EPI_ISL_476119, EPI_ISL_476120, EPI_ISL_476121, EPI_ISL_476122, EPI_ISL_476123, EPI_ISL_476124, EPI_ISL_476125, EPI_ISL_476126, EPI_ISL_476127, EPI_ISL_476128, EPI_ISL_476129, EPI_ISL_476130, EPI_ISL_476131, EPI_ISL_476132, EPI_ISL_476133, EPI_ISL_476134 | | | |
| see above | Viollier AG | Department of Biosystems Science and Engineering, ETH Zürich | Christian Beisel, Sarah Nadeau, Ivan Topolsky, Pedro Ferreira, Philipp Jablonski, Susana Posada-Céspedes, Tobias Schär, Ina Nissen, Natascha Santacroce, Elodie Burcklen, Christiane Beckmann, Maurice Redondo, Olivier Kobel, Christoph Noppen, Sophie Seidel, Noemie Santamaria de Souza, Niko Beerenwinkel, Tanja Stadler |
| EPI_ISL_476135 | Achima Care Fristadens VC | The Public Health Agency of Sweden | Oskar Karlsson Lindsjo, Maria Lind Karlberg, Mattias Haukland, Reza Advani, Olov Svartstrom, Anna-Malin Linde, Sandra Broddesson, Petra Edquist, Mia Brytting, Anna Risberg, Karin Tegmark-Wisell |
| EPI_ISL_476136 | Surbrunns VC | The Public Health Agency of Sweden | Oskar Karlsson Lindsjo, Maria Lind Karlberg, Mattias Haukland, Reza Advani, Olov Svartstrom, Anna-Malin Linde, Sandra Broddesson, Petra Edquist, Mia Brytting, Anna Risberg, Karin Tegmark-Wisell |
| EPI_ISL_476137 | Wasterlakarna | The Public Health Agency of Sweden | Oskar Karlsson Lindsjo, Maria Lind Karlberg, Mattias Haukland, Reza Advani, Olov Svartstrom, Anna-Malin Linde, Sandra Broddesson, Petra Edquist, Mia Brytting, Anna Risberg, Karin Tegmark-Wisell |
| EPI_ISL_476138 | Ulltuna Vardcentral | The Public Health Agency of Sweden | Oskar Karlsson Lindsjo, Maria Lind Karlberg, Mattias Haukland, Reza Advani, Olov Svartstrom, Anna-Malin Linde, Sandra Broddesson, Petra Edquist, Mia Brytting, Anna Risberg, Karin Tegmark-Wisell |
| EPI_ISL_476139 | Folkhalsomyndigheten | The Public Health Agency of Sweden | Oskar Karlsson Lindsjo, Maria Lind Karlberg, Mattias Haukland, Reza Advani, Olov Svartstrom, Anna-Malin Linde, Sandra Broddesson, Petra Edquist, Shamam Muradrasoli, Anna Risberg, Karin Tegmark-Wisell |
| EPI_ISL_476140, EPI_ISL_476141, EPI_ISL_476142 | Klinisk Mikrobiologi | The Public Health Agency of Sweden | Oskar Karlsson Lindsjo, Maria Lind Karlberg, Mattias Haukland, Reza Advani, Olov Svartstrom, Anna-Malin Linde, Sandra Broddesson, Petra Edquist, Shamam Muradrasoli, Anna Risberg, Karin Tegmark-Wisell |
| EPI_ISL_476143, EPI_ISL_476144 | Skovde/Unilabs | The Public Health Agency of Sweden | Oskar Karlsson Lindsjo, Maria Lind Karlberg, Mattias Haukland, Reza Advani, Olov Svartstrom, Anna-Malin Linde, Sandra Broddesson, Petra Edquist, Shamam Muradrasoli, Anna Risberg, Karin Tegmark-Wisell |
| EPI_ISL_476145, EPI_ISL_476146, EPI_ISL_476147 | Ostersund klinisk mikrobiologi | The Public Health Agency of Sweden | Oskar Karlsson Lindsjo, Maria Lind Karlberg, Mattias Haukland, Reza Advani, Olov Svartstrom, Anna-Malin Linde, Sandra Broddesson, Petra Edquist, |

| | | | |
|---|---|---|---|
| | | | Shamam Muradrasoli, Anna Risberg, Karin Tegmark-Wisell |
| EPI_ISL_476148, EPI_ISL_476149 | Institut Pasteur Dakar | Institut Pasteur de Dakar | Ndongo Dia, Moussa Moise Diagne, Mamadou Diop, Ousmane Faye, Amadou Alpha Sall |
| EPI_ISL_476150 | Institut Pasteur Dakar | Institut Pasteur de Dakar | Ndongo Dia, Moussa Moise Diagne, Mamadou diop, Ousmane Faye, Amadou Alpha Sall |
| EPI_ISL_476151 | Institut Pasteur Dakar | Institut Pasteur de Dakar | Ndongo Dia, Moussa Moise Diagne, Mamadou Diop, Ousmane faye, Amadou Alpha Sall |
| EPI_ISL_476152, EPI_ISL_476153, EPI_ISL_476154, EPI_ISL_476155, EPI_ISL_476156, EPI_ISL_476157, EPI_ISL_476158, EPI_ISL_476159, EPI_ISL_476160, EPI_ISL_476161, EPI_ISL_476162, EPI_ISL_476163, EPI_ISL_476164, EPI_ISL_476165, EPI_ISL_476166, EPI_ISL_476167, EPI_ISL_476168, EPI_ISL_476169, EPI_ISL_476170 | | | |
| see above | Laboratório de Patologia Clínica - UNICAMP | Laboratório de Estudos de Vírus Emergentes - UNICAMP | José Luiz Proença-Modena, Magnun Nueldo Nunes dos Santos, Angelica Schreiber, Julia Forato,Camila Simeoni, Marcilio Jorge Fumagalli, Mariene Ribeiro Amorim, Darlan da Silva Candido, Nuno Rodrigues Faria, Julien Theze, Luiz Gonzaga,Jaqueline Goes Jesus e William Marciel de Souza |
| EPI_ISL_476171, EPI_ISL_476172, EPI_ISL_476173, EPI_ISL_476174, EPI_ISL_476175, EPI_ISL_476176, EPI_ISL_476177, EPI_ISL_476178, EPI_ISL_476179, EPI_ISL_476180, EPI_ISL_476181, EPI_ISL_476182, EPI_ISL_476183, EPI_ISL_476184, EPI_ISL_476185, EPI_ISL_476186, EPI_ISL_476187, EPI_ISL_476188, EPI_ISL_476189, EPI_ISL_476190, EPI_ISL_476191, EPI_ISL_476192, EPI_ISL_476193, EPI_ISL_476194, EPI_ISL_476195, EPI_ISL_476196, EPI_ISL_476197, EPI_ISL_476198, EPI_ISL_476199, EPI_ISL_476200, EPI_ISL_476201, EPI_ISL_476202 | | | |
| see above | DB Diagnósticos do Brasil | Instituto de Medicina Tropical da Univesidade de São Paulo | Samples: Nelson Gaburo Jr; Sequencing: Ingra Morales Claro, Jaqueline Goes de Jesus, Erika Regina Manuli, Flavia Cristina da Silva Sales, Thais de Moura Coletti, Camila Alves Maia da Silva, Mariana Severo Ramundo, Giula Magalhaes Ferreira, Darlan da Silva Candido, Julien Theze, Nuno Faria, Ester Sabino |
| EPI_ISL_476203, EPI_ISL_476204, EPI_ISL_476205, EPI_ISL_476206, EPI_ISL_476207, EPI_ISL_476208 | Hospital da Clínicas da Faculdade de Medicina da Universidade de São Paulo | Instituto de Medicina Tropical da Univesidade de São Paulo | Samples: Ingra Morales Claro, Erika Regina Manuli, Cecilia Salete Alencar, Carolina S. Lazar, Silvia F. Costa; Sequencing: Ingra Morales Claro, Jaqueline Goes de Jesus, Erika Regina Manuli, Flavia Cristina da Silva Sales, Thais de Moura Coletti, Camila Alves Maia da Silva, Mariana Severo Ramundo, Giula Magalhaes Ferreira, Darlan da Silva Candido, Julien Theze, Nuno Faria, Ester Sabino |
| EPI_ISL_476209, EPI_ISL_476210, EPI_ISL_476211, EPI_ISL_476212, EPI_ISL_476213, EPI_ISL_476214, EPI_ISL_476215, EPI_ISL_476216 | DB Diagnósticos do Brasil | Instituto de Medicina Tropical da Univesidade de São Paulo | Samples: Nelson Gaburo Jr; Sequencing: Ingra Morales Claro, Jaqueline Goes de Jesus, Erika Regina Manuli, Flavia Cristina da Silva Sales, Thais de Moura Coletti, Camila Alves Maia da Silva, Mariana Severo Ramundo, Giula Magalhaes Ferreira, Darlan da Silva Candido, Julien Theze, Nuno Faria, Ester Sabino |
| EPI_ISL_476217 | Hospital da Clínicas da Faculdade de Medicina da Universidade de São Paulo | Instituto de Medicina Tropical da Univesidade de São Paulo | Samples: Ingra Morales Claro, Erika Regina Manuli, Cecilia Salete Alencar, Carolina S. Lazar, Silvia F. Costa; Sequencing: Ingra Morales Claro, Jaqueline Goes de Jesus, Erika Regina Manuli, Flavia Cristina da Silva Sales, Thais de Moura Coletti, Camila Alves Maia da Silva, Mariana Severo Ramundo, Giula Magalhaes Ferreira, Darlan da Silva Candido, Julien Theze, Nuno Faria, Ester Sabino |
| EPI_ISL_476218, EPI_ISL_476219 | DB Diagnósticos do Brasil | Instituto de Medicina Tropical da Univesidade de São Paulo | Samples: Nelson Gaburo Jr; Sequencing: Ingra Morales Claro, Jaqueline Goes de Jesus, Erika Regina Manuli, Flavia Cristina da Silva Sales, Thais de Moura Coletti, Camila Alves Maia da Silva, Mariana Severo Ramundo, Giula Magalhaes Ferreira, Darlan da Silva Candido, Julien Theze, Nuno Faria, Ester Sabino |
| EPI_ISL_476220, EPI_ISL_476221 | Laboratory Fleury | Instituto de Medicina Tropical da Univesidade de São Paulo | Samples: Celso Granato; Sequencing: Ingra Morales Claro, Jaqueline Goes de Jesus, Erika Regina Manuli, Flavia Cristina da Silva Sales, Thais de Moura Coletti, Camila Alves Maia da Silva, Mariana Severo Ramundo, Giula Magalhaes Ferreira, Darlan da Silva Candido, Julien Theze, Nuno Faria, Ester Sabino |
| EPI_ISL_476222, EPI_ISL_476223 | Hospital da Clínicas da Faculdade de Medicina da Universidade de São Paulo | Instituto de Medicina Tropical da Univesidade de São Paulo | Samples: Ingra Morales Claro, Erika Regina Manuli, Cecilia Salete Alencar, Carolina S. Lazar, Silvia F. Costa; Sequencing: Ingra Morales Claro, Jaqueline Goes de Jesus, Erika Regina Manuli, Flavia Cristina da Silva Sales, Thais de Moura Coletti, Camila Alves Maia da Silva, Mariana Severo Ramundo, Giula Magalhaes Ferreira, Darlan da Silva Candido, Julien Theze, Nuno Faria, Ester Sabino |
| EPI_ISL_476224, EPI_ISL_476225, EPI_ISL_476226, EPI_ISL_476227, EPI_ISL_476228, EPI_ISL_476229 | DB Diagnósticos do Brasil | Instituto de Medicina Tropical da Univesidade de São Paulo | Samples: Nelson Gaburo Jr; Sequencing: Ingra Morales Claro, Jaqueline Goes de Jesus, Erika Regina Manuli, Flavia Cristina da Silva Sales, Thais de Moura Coletti, Camila Alves Maia da Silva, Mariana Severo Ramundo, Giula Magalhaes Ferreira, Darlan da Silva Candido, Julien Theze, Nuno Faria, Ester Sabino |
| EPI_ISL_476230, EPI_ISL_476231 | Laboratory Fleury | Instituto de Medicina Tropical da Univesidade de São Paulo | Samples: Celso Granato; Sequencing: Ingra Morales Claro, Jaqueline Goes de Jesus, Erika Regina Manuli, Flavia Cristina da Silva Sales, Thais de Moura Coletti, Camila Alves Maia da Silva, Mariana Severo Ramundo, Giula Magalhaes Ferreira, Darlan da Silva Candido, Julien Theze, Nuno Faria, Ester Sabino |
| EPI_ISL_476232 | DB Diagnósticos do Brasil | Instituto de Medicina Tropical da Univesidade de São Paulo | Samples: Nelson Gaburo Jr; Sequencing: Ingra Morales Claro, Jaqueline Goes de Jesus, Erika Regina Manuli, Flavia Cristina da Silva Sales, Thais de Moura Coletti, Camila Alves Maia da Silva, Mariana Severo Ramundo, Giula Magalhaes Ferreira, Darlan da Silva Candido, Julien Theze, Nuno Faria, Ester Sabino |
| EPI_ISL_476233, EPI_ISL_476234, EPI_ISL_476235, EPI_ISL_476236 | Laboratory Fleury | Instituto de Medicina Tropical da Univesidade de São Paulo | Samples: Celso Granato; Sequencing: Ingra Morales Claro, Jaqueline Goes de Jesus, Erika Regina Manuli, Flavia Cristina da Silva Sales, Thais de Moura Coletti, Camila Alves Maia da Silva, Mariana Severo Ramundo, Giula Magalhaes Ferreira, Darlan da Silva Candido, Julien Theze, Nuno Faria, Ester Sabino |
| EPI_ISL_476237, EPI_ISL_476238, EPI_ISL_476239, EPI_ISL_476240, EPI_ISL_476241, EPI_ISL_476242, EPI_ISL_476243, EPI_ISL_476244, EPI_ISL_476245, EPI_ISL_476246, EPI_ISL_476247, EPI_ISL_476248, EPI_ISL_476249, EPI_ISL_476250, EPI_ISL_476251, EPI_ISL_476252, EPI_ISL_476253, EPI_ISL_476254, EPI_ISL_476255, EPI_ISL_476256, EPI_ISL_476257, EPI_ISL_476258, EPI_ISL_476259, EPI_ISL_476260, EPI_ISL_476261, EPI_ISL_476262, EPI_ISL_476263, EPI_ISL_476264, EPI_ISL_476265, EPI_ISL_476266, EPI_ISL_476267, EPI_ISL_476268, EPI_ISL_476269, EPI_ISL_476270, EPI_ISL_476271, EPI_ISL_476272, EPI_ISL_476273, EPI_ISL_476274, EPI_ISL_476275, EPI_ISL_476276, EPI_ISL_476277 | | | |
| see above | Hospital da Clínicas da Faculdade de Medicina da Universidade de São Paulo | Instituto de Medicina Tropical da Univesidade de São Paulo | Samples: Ingra Morales Claro, Erika Regina Manuli, Cecilia Salete Alencar, Carolina S. Lazar, Silvia F. Costa; Sequencing: Ingra Morales Claro, Jaqueline Goes de Jesus, Erika Regina Manuli, Flavia Cristina da Silva Sales, Thais de Moura Coletti, Camila Alves Maia da Silva, Mariana Severo Ramundo, Giula Magalhaes Ferreira, Darlan da Silva Candido, Julien Theze, Nuno Faria, Ester Sabino |
| EPI_ISL_476278, EPI_ISL_476279, EPI_ISL_476280, EPI_ISL_476281, EPI_ISL_476282, EPI_ISL_476283, EPI_ISL_476284, EPI_ISL_476285, EPI_ISL_476286, EPI_ISL_476287, EPI_ISL_476288, EPI_ISL_476289, EPI_ISL_476290, EPI_ISL_476291, EPI_ISL_476292, EPI_ISL_476293, EPI_ISL_476294, EPI_ISL_476295, EPI_ISL_476296, EPI_ISL_476297, EPI_ISL_476298, EPI_ISL_476299, EPI_ISL_476300, EPI_ISL_476301, EPI_ISL_476302, EPI_ISL_476303, EPI_ISL_476304, EPI_ISL_476305, EPI_ISL_476306, EPI_ISL_476307, EPI_ISL_476308, EPI_ISL_476309, EPI_ISL_476310, EPI_ISL_476311, EPI_ISL_476312, EPI_ISL_476313, EPI_ISL_476314, EPI_ISL_476315, EPI_ISL_476316, EPI_ISL_476317, EPI_ISL_476318, EPI_ISL_476319, EPI_ISL_476320, EPI_ISL_476321, EPI_ISL_476322, EPI_ISL_476323, EPI_ISL_476324, EPI_ISL_476325, EPI_ISL_476326, EPI_ISL_476327, EPI_ISL_476328, EPI_ISL_476329, EPI_ISL_476330, EPI_ISL_476331, EPI_ISL_476332, EPI_ISL_476333, EPI_ISL_476334, EPI_ISL_476335, EPI_ISL_476336 | | | |
| see above | DB Diagnósticos do Brasil | Instituto de Medicina Tropical da Univesidade de São Paulo | Samples: Nelson Gaburo Jr; Sequencing: Ingra Morales Claro, Jaqueline Goes de Jesus, Erika Regina Manuli, Flavia Cristina da Silva Sales, Thais de Moura Coletti, Camila Alves Maia da Silva, Mariana Severo Ramundo, Giula Magalhaes Ferreira, Darlan da Silva Candido, Julien Theze, Nuno Faria, Ester Sabino |
| EPI_ISL_476337, EPI_ISL_476338, EPI_ISL_476339, EPI_ISL_476340, EPI_ISL_476341, EPI_ISL_476342, EPI_ISL_476343, EPI_ISL_476344, EPI_ISL_476345, EPI_ISL_476346, EPI_ISL_476347, EPI_ISL_476348, EPI_ISL_476349 | | | |
| see above | Laboratório de Patologia Clínica - UNICAMP | Laboratório de Estudos de Vírus Emergentes - UNICAMP | José Luiz Proença-Modena, Magnun Nueldo Nunes dos Santos, Angelica Schreiber, Julia Forato,Camila Simeoni, Marcilio Jorge Fumagalli, Mariene Ribeiro Amorim, Darlan da Silva Candido, Nuno Rodrigues Faria, Julien Theze, Luiz Gonzaga,Jaqueline Goes Jesus e William Marciel de Souza |
| EPI_ISL_476350, EPI_ISL_476351, EPI_ISL_476352, EPI_ISL_476353, EPI_ISL_476354, EPI_ISL_476355, EPI_ISL_476356, EPI_ISL_476357, EPI_ISL_476358, EPI_ISL_476359, EPI_ISL_476360, EPI_ISL_476361, EPI_ISL_476362, EPI_ISL_476363, EPI_ISL_476364, EPI_ISL_476365, EPI_ISL_476366, EPI_ISL_476367, EPI_ISL_476368, EPI_ISL_476369, EPI_ISL_476370, EPI_ISL_476371 | | | |
| see above | DB Diagnósticos do Brasil | Instituto de Medicina Tropical da Univesidade de São Paulo | Samples: Nelson Gaburo Jr; Sequencing: Ingra Morales Claro, Jaqueline Goes de Jesus, Erika Regina Manuli, Flavia Cristina da Silva Sales, Thais de Moura Coletti, Camila Alves Maia da Silva, Mariana Severo Ramundo, Giula Magalhaes Ferreira, Darlan da Silva Candido, Julien Theze, Nuno Faria, Ester Sabino |
| EPI_ISL_476372, EPI_ISL_476373, EPI_ISL_476374, EPI_ISL_476375, EPI_ISL_476376, EPI_ISL_476377, EPI_ISL_476378, EPI_ISL_476379, EPI_ISL_476380, EPI_ISL_476381, EPI_ISL_476382, EPI_ISL_476383, EPI_ISL_476384, EPI_ISL_476385, EPI_ISL_476386 | | | |
| see above | Hospital da Clínicas da Faculdade de Medicina da Universidade de São Paulo | Instituto de Medicina Tropical da Univesidade de São Paulo | Samples: Ingra Morales Claro, Erika Regina Manuli, Cecilia Salete Alencar, Carolina S. Lazar, Silvia F. Costa; Sequencing: Ingra Morales Claro, Jaqueline Goes de Jesus, Erika Regina Manuli, Flavia Cristina da Silva Sales, Thais de Moura Coletti, Camila Alves Maia da Silva, Mariana Severo Ramundo, Giula Magalhaes Ferreira, Darlan da Silva Candido, Julien Theze, Nuno Faria, Ester Sabino |
| EPI_ISL_476387, EPI_ISL_476388, EPI_ISL_476389, EPI_ISL_476390, EPI_ISL_476391, EPI_ISL_476392, EPI_ISL_476393, EPI_ISL_476394, EPI_ISL_476395, EPI_ISL_476396, EPI_ISL_476397, EPI_ISL_476398, EPI_ISL_476399, EPI_ISL_476400, EPI_ISL_476401, EPI_ISL_476402 | | | |
| see above | Laboratório de Patologia Clínica - UNICAMP | Laboratório de Estudos de Vírus Emergentes - UNICAMP | José Luiz Proença-Modena, Magnun Nueldo Nunes dos Santos, Angelica Schreiber, Julia Forato,Camila Simeoni, Marcilio Jorge Fumagalli, Mariene Ribeiro Amorim, Darlan da Silva Candido, Nuno Rodrigues Faria, Julien Theze, Luiz Gonzaga,Jaqueline Goes Jesus e William Marciel de Souza |
| EPI_ISL_476403, EPI_ISL_476404, EPI_ISL_476405 | Laboratory DASA | Instituto de Medicina Tropical da Univesidade de São Paulo | Samples: José Eduardo Levi; Sequencing: Ingra Morales Claro, Jaqueline Goes de Jesus, Erika Regina Manuli, Flavia Cristina da Silva Sales, Thais de Moura Coletti, Camila Alves Maia da Silva, Mariana Severo Ramundo, Giula Magalhaes Ferreira, Darlan da Silva Candido, Julien Theze, Nuno Faria, Ester Sabino |
| EPI_ISL_476406, EPI_ISL_476407 | Laboratory Fleury | Instituto de Medicina Tropical da Univesidade de São | Samples: Celso Granato; Sequencing: Ingra Morales Claro, Jaqueline Goes de Jesus, Erika Regina Manuli, Flavia Cristina da Silva Sales, Thais de Moura |

| | | | |
|---|---|---|---|
| | | Paulo | Coletti, Camila Alves Maia da Silva, Mariana Severo Ramundo, Giula Magalhaes Ferreira, Darlan da Silva Candido, Julien Theze, Nuno Faria, Ester Sabino |
| EPI_ISL_476408, EPI_ISL_476409, EPI_ISL_476410, EPI_ISL_476411, EPI_ISL_476412, EPI_ISL_476413, EPI_ISL_476414, EPI_ISL_476415, EPI_ISL_476416, EPI_ISL_476417, EPI_ISL_476418, EPI_ISL_476419, EPI_ISL_476420, EPI_ISL_476421, EPI_ISL_476422, EPI_ISL_476423, EPI_ISL_476424, EPI_ISL_476425 | | | |
| see above | Laboratório de Patologia Clínica - UNICAMP | Laboratório de Estudos de Vírus Emergentes - UNICAMP | José Luiz Proença-Modena, Magnun Nueldo Nunes dos Santos, Angelica Schreiber, Julia Forato,Camila Simeoni, Marcilio Jorge Fumagalli, Mariene Ribeiro Amorim, Darlan da Silva Candido, Nuno Rodrigues Faria, Julien Theze, Luiz Gonzaga,Jaqueline Goes Jesus e William Marciel de Souza |
| EPI_ISL_476426, EPI_ISL_476427 | Laboratory Fleury | Instituto de Medicina Tropical da Univesidade de São Paulo | Samples: Celso Granato; Sequencing: Ingra Morales Claro, Jaqueline Goes de Jesus, Erika Regina Manuli, Flavia Cristina da Silva Sales, Thais de Moura Coletti, Camila Alves Maia da Silva, Mariana Severo Ramundo, Giula Magalhaes Ferreira, Darlan da Silva Candido, Julien Theze, Nuno Faria, Ester Sabino |
| EPI_ISL_476428, EPI_ISL_476429, EPI_ISL_476430, EPI_ISL_476431, EPI_ISL_476432, EPI_ISL_476433, EPI_ISL_476434, EPI_ISL_476435, EPI_ISL_476436, EPI_ISL_476437, EPI_ISL_476438, EPI_ISL_476439, EPI_ISL_476440, EPI_ISL_476441, EPI_ISL_476442, EPI_ISL_476443, EPI_ISL_476444, EPI_ISL_476445, EPI_ISL_476446, EPI_ISL_476447, EPI_ISL_476448, EPI_ISL_476449, EPI_ISL_476450, EPI_ISL_476451, EPI_ISL_476452, EPI_ISL_476453, EPI_ISL_476454, EPI_ISL_476455, EPI_ISL_476456, EPI_ISL_476457, EPI_ISL_476458, EPI_ISL_476459, EPI_ISL_476460, EPI_ISL_476461, EPI_ISL_476462, EPI_ISL_476463, EPI_ISL_476464, EPI_ISL_476465, EPI_ISL_476466, EPI_ISL_476467, EPI_ISL_476468, EPI_ISL_476469, EPI_ISL_476470, EPI_ISL_476471, EPI_ISL_476472, EPI_ISL_476473, EPI_ISL_476474, EPI_ISL_476475, EPI_ISL_476476, EPI_ISL_476477, EPI_ISL_476478, EPI_ISL_476479, EPI_ISL_476480, EPI_ISL_476481, EPI_ISL_476482, EPI_ISL_476483, EPI_ISL_476484, EPI_ISL_476485, EPI_ISL_476486, EPI_ISL_476487, EPI_ISL_476488, EPI_ISL_476489, EPI_ISL_476490 | | | |
| see above | Hospital da Clínicas da Faculdade de Medicina da Universidade de São Paulo | Instituto de Medicina Tropical da Univesidade de São Paulo | Samples: Ingra Morales Claro, Erika Regina Manuli, Cecilia Salete Alencar, Carolina S. Lazar, Silvia F. Costa; Sequencing: Ingra Morales Claro, Jaqueline Goes de Jesus, Erika Regina Manuli, Flavia Cristina da Silva Sales, Thais de Moura Coletti, Camila Alves Maia da Silva, Mariana Severo Ramundo, Giula Magalhaes Ferreira, Darlan da Silva Candido, Julien Theze, Nuno Faria, Ester Sabino |
| EPI_ISL_476491, EPI_ISL_476492 | Institut Pasteur Dakar | Institut Pasteur de Dakar | Ndongo Dia, Moussa Moise Diagne, Mamadou Diop, Ousmane Faye, Amadou Alpha Sall |
| EPI_ISL_476493 | Institut Pasteur Dakar | Institut Pasteur de Dakar | Ndongo Dia, Moussa Moise Diagne, Mamadou Diop, Ousmane Faye, Amadou alpha Sall |
| EPI_ISL_476494 | Institut Pasteur Dakar | Institut Pasteur de Dakar | Ndongo Dia, Moussa Moise Diagne, Mamadou Diop, Ousmane Faye, Amadou Alpha Sall |
| EPI_ISL_476495 | Institut Pasteur Dakar | Institut Pasteur de Dakar | Ndongo Dia, Moussa Moise Diagne, Mamadou Diop, Ousmane Faye, Amadou Alpha Sall |
| EPI_ISL_476496 | Hospital Garrahan | Héritas | Dalmacio Pereyra, Roberta Crespo, Mauricio Grisolia, Cristian Rohr, Andrea Mangano, Maria Florencia Fernandez, Fabian Fay, Martin Vazquez |
| EPI_ISL_476497 | Institut Pasteur Dakar | Institut Pasteur de Dakar | Ndongo Dia, Moussa Moise Diagne, Mamadou Diop, Ousmane Faye, Amadou alpha Sall |
| EPI_ISL_476498, EPI_ISL_476499, EPI_ISL_476500, EPI_ISL_476501, EPI_ISL_476502, EPI_ISL_476503, EPI_ISL_476504, EPI_ISL_476505, EPI_ISL_476506, EPI_ISL_476507, EPI_ISL_476508, EPI_ISL_476509, EPI_ISL_476510, EPI_ISL_476511, EPI_ISL_476512, EPI_ISL_476513 | | | |
| see above | Laboratoire de microbiologie, Hopital de Verdun | Smith Laboratory, Centre de Recherche CHU Sainte-Justine | Martin Smith, Marieke Rozendaal, Ivan Pavlov |
| EPI_ISL_476514 | Institut Pasteur Dakar | Institut Pasteur de Dakar | Ndongo Dia, Moussa Moise Diagne, Mamadou Diop, Ousmane Faye, Amadou Alpha Sall |
| EPI_ISL_476515 | Institut Pasteur Dakar | Institut Pasteur de Dakar | Ndongo Dia, Moussa Moise Diagne, Mamadou diop, Ousmane Faye, Amadou alpha Sall |
| EPI_ISL_476516 | Institut Pasteur Dakar | Institut Pasteur de Dakar | Ndongo Dia, Moussa Moise Diagne, mamadou Diop, Ousmane Faye, Amadou Alpha Sall |
| EPI_ISL_476517, EPI_ISL_476518, EPI_ISL_476519, EPI_ISL_476520, EPI_ISL_476521, EPI_ISL_476522, EPI_ISL_476523, EPI_ISL_476524, EPI_ISL_476525, EPI_ISL_476526, EPI_ISL_476527, EPI_ISL_476528, EPI_ISL_476529, EPI_ISL_476530, EPI_ISL_476531, EPI_ISL_476532, EPI_ISL_476533, EPI_ISL_476534, EPI_ISL_476535, EPI_ISL_476536, EPI_ISL_476537, EPI_ISL_476538, EPI_ISL_476539, EPI_ISL_476540, EPI_ISL_476541, EPI_ISL_476542, EPI_ISL_476543, EPI_ISL_476544, EPI_ISL_476545, EPI_ISL_476546, EPI_ISL_476547, EPI_ISL_476548, EPI_ISL_476549, EPI_ISL_476550, EPI_ISL_476551, EPI_ISL_476552, EPI_ISL_476553, EPI_ISL_476554, EPI_ISL_476555, EPI_ISL_476556, EPI_ISL_476557 | | | |
| see above | Yale Clinical Virology Laboratory | Grubaugh Lab - Yale School of Public Health | Joseph Fauver, Tara Alpert, Anderson Brito, Anne Wyllie, Chantal Vogels, Mary Petrone, Cole Jensen, Chaney Kalinich, Isabel Ott, Arnau Casanovas, Catherine Muenker, Adam Moore, Alice Lu, Maria Tokuyama, Patrick Wong, Peiwen Lu, Saad Omer, Richard Martinello, Allison Nelson, Shelli Farhadian, Akiko Iwasaki, Charlese Dela Cruz, Albert Ko, Nathan Grubaugh |
| EPI_ISL_476558 | Institut Pasteur Dakar | Institut Pasteur de Dakar | Ndongo Dia, Moussa Moise Diagne, Mamadou Diop, Ousmane Faye, Amadou Alpha Sall |
| EPI_ISL_476559 | unknown | Laboratoire Sciences et Technologies de la Santé (STS) Institut Supérieur des Sciences de la Santé Université Hassan 1er, Settat, Morocco | Hajar Lemriss, Sanaâ Lemriss, Amal Souiri, Narjis Amar, Mustapha Mouallif, Touria Essayagh, Jawad Bouzid, Saâd EL Kabbaj, Abderraouf Hilali |
| EPI_ISL_476560 | Institut Pasteur Dakar | Institut Pasteur de Dakar | Ndongo Dia, Moussa Moise Diagne, Mamadou Diop, Ousmane Faye, Amadou Alpha Sall |
| EPI_ISL_476561 | Hospital Garrahan | Héritas | Roberta Crespo, Dalmacio Pereyra, Mauricio Grisolia, Cristian Rohr, Andrea Mangano, Maria Florencia Fernandez, Fabian Fay, Martin Vazquez |
| EPI_ISL_476562 | Institut Pasteur Dakar | Institut Pasteur de Dakar | Ndongo Dia, Moussa Moise Diagne, Mamadou Diop, Ousmane Faye, Amadou Alpha Sall |
| EPI_ISL_476563 | Hospital de Pediatría "Prof. Dr. Juan P Garrahan" | Héritas | Dalmacio Pereyra, Roberta Crespo, Mauricio Grisolia, Cristian Rohr, Andrea Mangano, Maria Florencia Fernandez, Fabian Fay, Martin Vazquez |
| EPI_ISL_476564 | Institut Pasteur Dakar | Institut Pasteur de Dakar | Ndongo Dia, Moussa Moise Diagne, Mamadou diop, Ousmane Faye, Amadou alpha Sall |
| EPI_ISL_476565 | Hospital de Pediatría "Prof. Dr. Juan P Garrahan" | Héritas | Andrea Mangano, Maria Florencia Fernandez, Dalmacio Pereyra, Roberta Crespo, Mauricio Grisolia, Cristian Rohr, Fabian Fay, Martin Vazquez |
| EPI_ISL_476566 | Institut pasteur Dakar | Institut Pasteur de Dakar | Ndongo Dia, Moussa Moise Diagne, Mamadou Diop, Ousmane Faye, Amadou Alpha Sall |
| EPI_ISL_476567 | Hospital de Pediatría "Prof. Dr. Juan P Garrahan" | Héritas | Dalmacio Pereyra, Roberta Crespo, Mauricio Grisolia, Cristian Rohr, Andrea Mangano, Maria Florencia Fernandez, Fabian Fay, Martin Vazquez |
| EPI_ISL_476568 | Hospital de Pediatría "Prof. Dr. Juan P Garrahan" | Héritas | Cristian Rohr, Andrea Mangano, Maria Florencia Fernandez, Dalmacio Pereyra, Roberta Crespo, Mauricio Grisolia, Fabian Fay, Martin Vazquez |
| EPI_ISL_476569 | Institut Pasteur Dakar | Institut Pasteur de Dakar | Ndongo Dia, Moussa Moise, Mamadou Diop, Ousmane Faye, Amadou Alpha Sall |
| EPI_ISL_476570 | Institut Pasteur Dakar | Institut Pasteur de Dakar | Ndongo Dia, Moussa Moise Diagne, Mamadou Diop, Ousmane Faye, Amadou Alpha Sall |
| EPI_ISL_476571 | Hospital de Pediatría "Prof. Dr. Juan P Garrahan" | Héritas | Dalmacio Pereyra, Roberta Crespo, Mauricio Grisolia, Cristian Rohr, Andrea Mangano, Maria Florencia Fernandez, Fabian Fay, Martin Vazquez |
| EPI_ISL_476572 | Institut Pasteur Dakar | Institut Pasteur de Dakar | Ndongo Dia, Moussa Moise Diagne, Mamadou Diop, Ousmane Faye, Amadou Alpha Sall |
| EPI_ISL_476573 | Hospital de Pediatría "Prof. Dr. Juan P Garrahan" | Héritas | Dalmacio Pereyra, Roberta Crespo, Mauricio Grisolia, Cristian Rohr, Andrea Mangano, Maria Florencia Fernandez, Fabian Fay, Martin Vazquez |
| EPI_ISL_476574 | Institut Pasteur Dakar | Institut Pasteur de Dakar | Ndongo Dia, Moussa Moise Diagne, Mamadou Diop, Ousmane Faye, Amadou Alpha Sall |
| EPI_ISL_476702 | Incubadora Venezolana de Ciencia, Venezuela | Incubadora Venezolana de Ciencia, Venezuela / Instituto Nacional de Salud, Bogotá, Colombia / Grupo de Investigaciones Microbiológicas-UR (GIMUR), Departamento de Biología, Facultad de Ciencias Naturales, Universidad del Rosario, Bogotá, Colombia / Icahn School of Medicine at Mount Sinai, New York, USA | Alberto Paniz-Mondolfi, Marina Muñoz, Luis Perez-Garcia, Lourdes Delgado, Carolina Florez, Sergio Gomez, Angelica Rico, Lisseth Pardo, Esther C. Barros, Carolina Hernández, Jesús E. Jaimes, Aníbal A. Teherán, Ana S. Gonzalez-Reiche, Matthew M. Hernandez, Emilia Mia Sordillo, Viviana Simon, Harm van Bakel, Juan David Ramírez |
| EPI_ISL_476703, EPI_ISL_476704 | Incubadora Venezolana de Ciencia, Venezuela | Incubadora Venezolana de Ciencia, Venezuela / Instituto Nacional de Salud, Bogotá, Colombia / Grupo de Investigaciones Microbiológicas-UR (GIMUR), Departamento de Biología, Facultad de Ciencias Naturales, Universidad del Rosario, Bogotá, Colombia / Icahn School of Medicine at Mount Sinai, New York, USA | Alberto Paniz-Mondolfi, Marina Muñoz, Luis Perez-Garcia, Lourdes Delgado, Carolina Florez, Sergio Gomez, Angelica Rico, Lisseth Pardo, Esther C. Barros, Carolina Hernández, Jesús E. Jaimes, Aníbal A. Teherán, Ana S. Gonzalez-Reiche, Matthew M. Hernandez, Emilia Mia Sordillo, Viviana Simon, Harm van Bakel, Juan David Ramírez |
| EPI_ISL_476705 | Labor Kneißler GmbH & Co. KG | Heinrich Pette Institute, Leibniz Institute for Experimental Virology | Günther, Thomas; Grundhoff, Adam; Czech-Sioli, Manya; Fischer, Nicole; Ottinger, Matthias; Brinkmann, Melanie M. |
| EPI_ISL_476706, EPI_ISL_476707, EPI_ISL_476708, EPI_ISL_476709, EPI_ISL_476710, EPI_ISL_476711, EPI_ISL_476712, EPI_ISL_476713, EPI_ISL_476714, EPI_ISL_476715, EPI_ISL_476716, EPI_ISL_476717, EPI_ISL_476718, EPI_ISL_476719, EPI_ISL_476720, EPI_ISL_476721, EPI_ISL_476722, EPI_ISL_476723, EPI_ISL_476724, EPI_ISL_476725, EPI_ISL_476726, EPI_ISL_476727, EPI_ISL_476728, EPI_ISL_476729, EPI_ISL_476730, EPI_ISL_476731, EPI_ISL_476732, EPI_ISL_476733, EPI_ISL_476734, EPI_ISL_476735, EPI_ISL_476736, EPI_ISL_476737, EPI_ISL_476738, EPI_ISL_476739, EPI_ISL_476740, EPI_ISL_476741, | | | |

| | | | |
|---|---|---|---|
| EPI_ISL_476742, EPI_ISL_476743, EPI_ISL_476744, EPI_ISL_476745, EPI_ISL_476746, EPI_ISL_476747, EPI_ISL_476748, EPI_ISL_476749, EPI_ISL_476750, EPI_ISL_476751, EPI_ISL_476752, EPI_ISL_476753, EPI_ISL_476754, EPI_ISL_476755, EPI_ISL_476756, EPI_ISL_476757, EPI_ISL_476758, EPI_ISL_476759, EPI_ISL_476760, EPI_ISL_476761, EPI_ISL_476762, EPI_ISL_476763, EPI_ISL_476764, EPI_ISL_476765, EPI_ISL_476766 | | | |
| see above | Minnesota Department of Health, Public Health Laboratory | Minnesota Department of Health, Public Health Laboratory | Matt Plumb, Jacob Garfin, and Xiong Wang |
| EPI_ISL_476767, EPI_ISL_476768, EPI_ISL_476769, EPI_ISL_476770, EPI_ISL_476771, EPI_ISL_476772, EPI_ISL_476773, EPI_ISL_476774, EPI_ISL_476775, EPI_ISL_476776, EPI_ISL_476777, EPI_ISL_476778, EPI_ISL_476779, EPI_ISL_476780, EPI_ISL_476781, EPI_ISL_476782, EPI_ISL_476783, EPI_ISL_476784, EPI_ISL_476785, EPI_ISL_476786, EPI_ISL_476787, EPI_ISL_476788, EPI_ISL_476789, EPI_ISL_476790, EPI_ISL_476791, EPI_ISL_476792, EPI_ISL_476793, EPI_ISL_476794 | | | |
| see above | Stanford clinical virology lab | Chan-Zuckerberg Biohub | Benjamin Pinksy, Katharine Walter, Victoria N. Parikh, John Gorzynski, Hannah N. DeJong, Matthew T. Wheeler, Jason Andrews, Manuel Rivas, Carlos Bustamante, Euan Ashley, with CZB Cliahub Consortium |
| EPI_ISL_476795 | Department of Laboratory Medicine Tan Tock Seng Hospital | Department of Laboratory Medicine Tan Tock Seng Hospital | Chen YYC, Zair X, Li C, Tang WY, Maurer-Stroh S, Barkham TMS, Nagarajan N, Sessions OM |
| EPI_ISL_476796, EPI_ISL_476797 | Department of Laboratory Medicine Tan Tock Seng Hospital | Department of Laboratory Medicine Tan Tock Seng Hospital | Chen YYC, Zair X, Li C, Tang WY, Maurer-Stroh S, Barkham TMS, Nagarajan N, Sessions OM |
| EPI_ISL_476801, EPI_ISL_476802, EPI_ISL_476803, EPI_ISL_476804 | Hong Kong Department of Health | School of Public Health, The University of Hong Kong | Dominic N.C. Tsang, Daniel K.W. Chu, Leo L.M. Poon, Malik Peiris |
| EPI_ISL_476805, EPI_ISL_476806, EPI_ISL_476807, EPI_ISL_476808, EPI_ISL_476809, EPI_ISL_476810, EPI_ISL_476811, EPI_ISL_476812, EPI_ISL_476813, EPI_ISL_476815, EPI_ISL_476816, EPI_ISL_476817, EPI_ISL_476818, EPI_ISL_476819 | | | |
| see above | Department of Laboratory Medicine Tan Tock Seng Hospital | Department of Laboratory Medicine Tan Tock Seng Hospital | Chen YYC, Zair X, Li C, Tang WY, Maurer-Stroh S, Barkham TMS, Nagarajan N, Sessions OM |
| EPI_ISL_476820 | Department of Laboratory Medicine Tan Tock Seng Hospital | Department of Laboratory Medicine Tan Tock Seng Hospital | Chen YYC, Zair X, Li C, Tang WY, Maurer-Stroh S, Barkham TMS, Nagarajan N, Sessions OM |
| EPI_ISL_476821 | Department of Laboratory Medicine Tan Tock Seng Hospital | Department of Laboratory Medicine Tan Tock Seng Hospital | Chen YYC, Zair X, Li C, Tang WY, Maurer-Stroh S, Barkham TMS, Nagarajan N, Sessions OM |
| EPI_ISL_476822, EPI_ISL_476823, EPI_ISL_476824, EPI_ISL_476825, EPI_ISL_476826, EPI_ISL_476827, EPI_ISL_476828, EPI_ISL_476829, EPI_ISL_476830, EPI_ISL_476831 | Laboratoire des Fièvres Hémorragiques Virales du Benin | Charité-Universitätsmedizin Berlin | Yadouleton, Anges; Sander Anna-Lena; Moreira-Soto Andres; Drexler, Jan Felix |
| EPI_ISL_476832 | Medical Biology Department, Kocaeli University | Medical Genetics Department, Kocaeli University | Savli H, Cine N, Sunnetci-Akkoyunlu D, Eren-Keskin S, Ilgazli A, Akhan S, Karadenizli A, Kasap M, Sayan M, Akpinar G, Canturk NZ. |
| EPI_ISL_476833, EPI_ISL_476834 | Laboratoire des Fièvres Hémorragiques Virales du Benin | Charité-Universitätsmedizin Berlin | Yadouleton, Anges; Sander Anna-Lena; Moreira-Soto Andres; Drexler, Jan Felix |
| EPI_ISL_476835 | National Influenza Centre for Northern Greece | National Influenza Centre for Northern Greece | Maria Christoforidi |
| EPI_ISL_476836 | National Influenza Centre for Nothern Greece | National Influenza Centre for Nothern Greece | Maria Christoforidi |
| EPI_ISL_476837, EPI_ISL_476838, EPI_ISL_476839 | National Influenza Centre for Northern Greece | National Influenza Centre for Northern Greece | Maria Christoforidi |
| EPI_ISL_476840 | Defence Research & Development Establishment (DRDE) | Defence Research & Development Establishment (DRDE) | Shashi Sharma, Paban Kumar Dash, Sushil Kumar Sharma, Ambuj Shrivastava, Jyoti S. Kumar |
| EPI_ISL_476841 | National Influenza Centre for Northern Greece | National Influenza Centre for Northern Greece | Maria Christoforidi |
| EPI_ISL_476842 | Defence Research & Development Establishment (DRDE) | Defence Research & Development Establishment (DRDE) | Shashi Sharma, Paban Kumar Dash, Sushil Kumar Sharma, Ambuj Shrivastava, Jyoti S. Kumar |
| EPI_ISL_476843 | National Influenza Centre for Northern Greece | National Influenza Centre for Northern Greece | Maria Christoforidi |
| EPI_ISL_476844 | Defence Research & Development Establishment (DRDE) | Defence Research & Development Establishment (DRDE) | Shashi Sharma, Paban Kumar Dash, Sushil Kumar Sharma, Ambuj Shrivastava, Jyoti S. Kumar |
| EPI_ISL_476845 | National Influenza Centre for Northern Greece | National Influenza Centre for Northern Greece | Maria Christoforidi |
| EPI_ISL_476846 | Defence Research & Development Establishment (DRDE) | Defence Research & Development Establishment (DRDE) | Shashi Sharma, Paban Kumar Dash, Sushil Kumar Sharma, Ambuj Shrivastava, Jyoti S. Kumar |
| EPI_ISL_476847 | National Influenza Centre for Northern Greece | National Influenza Centre for Northern Greece | Maria Christoforidi |
| EPI_ISL_476848, EPI_ISL_476849, EPI_ISL_476850 | Defence Research & Development Establishment (DRDE) | Defence Research & Development Establishment (DRDE) | Shashi Sharma, Paban Kumar Dash, Sushil Kumar Sharma, Ambuj Shrivastava, Jyoti S. Kumar |
| EPI_ISL_476851 | National Influenza Centre for Northern Greece | National Influenza Centre for Northern Greece | Maria Christoforidi |
| EPI_ISL_476852, EPI_ISL_476853, EPI_ISL_476854 | Defence Research & Development Establishment (DRDE) | Defence Research & Development Establishment (DRDE) | Shashi Sharma, Paban Kumar Dash, Sushil Kumar Sharma, Ambuj Shrivastava, Jyoti S. Kumar |
| EPI_ISL_476855 | GMERS Medical College & Hospital, Gotri, Vadodara | Gujarat Biotechnology Research Centre | Apurvasinh Puvar, Janvi Raval, Zarna Patel, Monika Gandhi, Pinal Trivedi, Maharshi Pandya, Nidhi Patel, Nitin Savaliya, Raghawendra Kumar, Dinesh Kumar, Zuber Saiyed, Komal Patel, Labdhi Pandya, Afzal Ansari, Nikha Trivedi, Meenakshi Shah, Neena Doshi, Varsha Godbole, R D Dixit, A M Kadri, Harsh Bakshi, Chaitanya Joshi, Madhvi Joshi |
| EPI_ISL_476856 | GMERS Medical College & Hospital, Gotri, Vadodara | Gujarat Biotechnology Research Centre | Janvi Raval, Zarna Patel, Monika Gandhi, Pinal Trivedi, Maharshi Pandya, Nidhi Patel, Nitin Savaliya, Raghawendra Kumar, Dinesh Kumar, Zuber Saiyed, Komal Patel, Labdhi Pandya, Afzal Ansari, Nikha Trivedi, Meenakshi Shah, Neena Doshi, Varsha Godbole, Apurvasinh Puvar, R D Dixit, A M Kadri, Harsh Bakshi, Chaitanya Joshi, Madhvi Joshi |
| EPI_ISL_476857 | GMERS Medical College & Hospital, Gotri, Vadodara | Gujarat Biotechnology Research Centre | Zarna Patel, Monika Gandhi, Pinal Trivedi, Maharshi Pandya, Nidhi Patel, Nitin Savaliya, Raghawendra Kumar, Dinesh Kumar, Zuber Saiyed, Komal Patel, Labdhi Pandya, Afzal Ansari, Nikha Trivedi, Meenakshi Shah, Neena Doshi, Varsha Godbole, Apurvasinh Puvar, Janvi Raval, R D Dixit, A M Kadri, Harsh Bakshi, Chaitanya Joshi, Madhvi Joshi |
| EPI_ISL_476858 | GMERS Medical College & Hospital, Gotri, Vadodara | Gujarat Biotechnology Research Centre | Monika Gandhi, Pinal Trivedi, Maharshi Pandya, Nidhi Patel, Nitin Savaliya, Raghawendra Kumar, Dinesh Kumar, Zuber Saiyed, Komal Patel, Labdhi Pandya, Afzal Ansari, Nikha Trivedi, Meenakshi Shah, Neena Doshi, Varsha Godbole, Apurvasinh Puvar, Janvi Raval, Zarna Patel, R D Dixit, A M Kadri, Harsh Bakshi, Chaitanya Joshi, Madhvi Joshi |
| EPI_ISL_476859 | GMERS Medical College & Hospital, Gotri, Vadodara | Gujarat Biotechnology Research Centre | Pinal Trivedi, Maharshi Pandya, Nidhi Patel, Nitin Savaliya, Raghawendra Kumar, Dinesh Kumar, Zuber Saiyed, Komal Patel, Labdhi Pandya, Afzal Ansari, Nikha Trivedi, Meenakshi Shah, Neena Doshi, Varsha Godbole, Apurvasinh Puvar, Janvi Raval, Zarna Patel, Monika Gandhi, R D Dixit, A M Kadri, Harsh Bakshi, Chaitanya Joshi, Madhvi Joshi |
| EPI_ISL_476860 | GMERS Medical College & Hospital, Gotri, Vadodara | Gujarat Biotechnology Research Centre | Maharshi Pandya, Nidhi Patel, Nitin Savaliya, Raghawendra Kumar, Dinesh Kumar, Zuber Saiyed, Komal Patel, Labdhi Pandya, Afzal Ansari, Nikha Trivedi, Meenakshi Shah, Neena Doshi, Varsha Godbole, Apurvasinh Puvar, Janvi Raval, Zarna Patel, Monika Gandhi, Pinal Trivedi, R D Dixit, A M Kadri, Harsh Bakshi, Chaitanya Joshi, Madhvi Joshi |
| EPI_ISL_476861 | GMERS Medical College & Hospital, Gotri, Vadodara | Gujarat Biotechnology Research Centre | Nidhi Patel, Nitin Savaliya, Raghawendra Kumar, Dinesh Kumar, Zuber Saiyed, Komal Patel, Labdhi Pandya, Afzal Ansari, Nikha Trivedi, Meenakshi Shah, Neena Doshi, Varsha Godbole, Apurvasinh Puvar, Janvi Raval, Zarna Patel, Monika Gandhi, Pinal Trivedi, Maharshi Pandya, R D Dixit, A M Kadri, Harsh Bakshi, Chaitanya Joshi, Madhvi Joshi |
| EPI_ISL_476862 | GMERS Medical College & Hospital, Gotri, Vadodara | Gujarat Biotechnology Research Centre | Nitin Savaliya, Raghawendra Kumar, Dinesh Kumar, Zuber Saiyed, Komal Patel, Labdhi Pandya, Afzal Ansari, Nikha Trivedi, Meenakshi Shah, Neena |

| | | | |
|---|---|---|---|
| | | | Doshi, Varsha Godbole, Apurvasinh Puvar, Janvi Raval, Zarna Patel, Monika Gandhi, Pinal Trivedi, Maharshi Pandya, Nidhi Patel, R D Dixit, A M Kadri, Harsh Bakshi, Chaitanya Joshi, Madhvi Joshi |
| EPI_ISL_476863 | GMERS Medical College and Hospital, Gandhinagar | Gujarat Biotechnology Research Centre | Raghawendra Kumar, Dinesh Kumar, Zuber Saiyed, Komal Patel, Labdhi Pandya, Afzal Ansari, Nikha Trivedi, Seema Bhatt, Gaurishankar Shrimali, Bhavesh Modi, Bharti Rajani, Apurvasinh Puvar, Janvi Raval, Zarna Patel, Monika Gandhi, Pinal Trivedi, Maharshi Pandya, Nidhi Patel, Nitin Savaliya, R D Dixit, A M Kadri, Harsh Bakshi, Chaitanya Joshi, Madhvi Joshi |
| EPI_ISL_476864 | GMERS Medical College and Hospital, Gandhinagar | Gujarat Biotechnology Research Centre | Dinesh Kumar, Zuber Saiyed, Komal Patel, Labdhi Pandya, Afzal Ansari, Nikha Trivedi, Seema Bhatt, Gaurishankar Shrimali, Bhavesh Modi, Bharti Rajani, Apurvasinh Puvar, Janvi Raval, Zarna Patel, Monika Gandhi, Pinal Trivedi, Maharshi Pandya, Nidhi Patel, Nitin Savaliya, Raghawendra Kumar, R D Dixit, A M Kadri, Harsh Bakshi, Chaitanya Joshi, Madhvi Joshi |
| EPI_ISL_476865 | GMERS Medical College and Hospital, Gandhinagar | Gujarat Biotechnology Research Centre | Zuber Saiyed, Komal Patel, Labdhi Pandya, Afzal Ansari, Nikha Trivedi, Seema Bhatt, Gaurishankar Shrimali, Bhavesh Modi, Bharti Rajani, Apurvasinh Puvar, Janvi Raval, Zarna Patel, Monika Gandhi, Pinal Trivedi, Maharshi Pandya, Nidhi Patel, Nitin Savaliya, Raghawendra Kumar, Dinesh Kumar, R D Dixit, A M Kadri, Harsh Bakshi, Chaitanya Joshi, Madhvi Joshi |
| EPI_ISL_476866 | GMERS Medical College and Hospital, Gandhinagar | Gujarat Biotechnology Research Centre | Komal Patel, Labdhi Pandya, Afzal Ansari, Nikha Trivedi, Seema Bhatt, Gaurishankar Shrimali, Bhavesh Modi, Bharti Rajani, Apurvasinh Puvar, Janvi Raval, Zarna Patel, Monika Gandhi, Pinal Trivedi, Maharshi Pandya, Nidhi Patel, Nitin Savaliya, Raghawendra Kumar, Dinesh Kumar, Zuber Saiyed, R D Dixit, A M Kadri, Harsh Bakshi, Chaitanya Joshi, Madhvi Joshi |
| EPI_ISL_476867 | Banas Medical College and Research Institute | Gujarat Biotechnology Research Centre | Labdhi Pandya, Afzal Ansari, Nikha Trivedi, Radhika Khara, Sunil R Joshi, Viren s Doshi, Apurvasinh Puvar, Janvi Raval, Zarna Patel, Monika Gandhi, Pinal Trivedi, Maharshi Pandya, Nidhi Patel, Nitin Savaliya, Raghawendra Kumar, Dinesh Kumar, Zuber Saiyed, Komal Patel, R D Dixit, A M Kadri, Harsh Bakshi, Chaitanya Joshi, Madhvi Joshi |
| EPI_ISL_476868 | Banas Medical College and Research Institute | Gujarat Biotechnology Research Centre | Afzal Ansari, Nikha Trivedi, Radhika Khara, Sunil R Joshi, Viren s Doshi, Apurvasinh Puvar, Janvi Raval, Zarna Patel, Monika Gandhi, Pinal Trivedi, Maharshi Pandya, Nidhi Patel, Nitin Savaliya, Raghawendra Kumar, Dinesh Kumar, Zuber Saiyed, Komal Patel, Labdhi Pandya, R D Dixit, A M Kadri, Harsh Bakshi, Chaitanya Joshi, Madhvi Joshi |
| EPI_ISL_476869 | Department of MicroBiology, Government Medical College, Surat | Gujarat Biotechnology Research Centre | Nikha Trivedi, Naresh Chauhan, Summaiya Mullan, Amit gamit, Apurvasinh Puvar, Janvi Raval, Zarna Patel, Monika Gandhi, Pinal Trivedi, Maharshi Pandya, Nidhi Patel, Nitin Savaliya, Raghawendra Kumar, Dinesh Kumar, Zuber Saiyed, Komal Patel, Labdhi Pandya, Afzal Ansari, R D Dixit, A M Kadri, Harsh Bakshi, Chaitanya Joshi, Madhvi Joshi |
| EPI_ISL_476870 | Department of MicroBiology, Government Medical College, Surat | Gujarat Biotechnology Research Centre | Naresh Chauhan, Summaiya Mullan, Amit gamit, Apurvasinh Puvar, Janvi Raval, Zarna Patel, Monika Gandhi, Pinal Trivedi, Maharshi Pandya, Nidhi Patel, Nitin Savaliya, Raghawendra Kumar, Dinesh Kumar, Zuber Saiyed, Komal Patel, Labdhi Pandya, Afzal Ansari, Nikha Trivedi, R D Dixit, A M Kadri, Harsh Bakshi, Chaitanya Joshi, Madhvi Joshi |
| EPI_ISL_476871 | Department of MicroBiology, Government Medical College, Surat | Gujarat Biotechnology Research Centre | Summaiya Mullan, Amit gamit, Apurvasinh Puvar, Janvi Raval, Zarna Patel, Monika Gandhi, Pinal Trivedi, Maharshi Pandya, Nidhi Patel, Nitin Savaliya, Raghawendra Kumar, Dinesh Kumar, Zuber Saiyed, Komal Patel, Labdhi Pandya, Afzal Ansari, Nikha Trivedi, Naresh Chauhan, R D Dixit, A M Kadri, Harsh Bakshi, Chaitanya Joshi, Madhvi Joshi |
| EPI_ISL_476872 | Department of MicroBiology, Government Medical College, Surat | Gujarat Biotechnology Research Centre | Amit gamit, Apurvasinh Puvar, Janvi Raval, Zarna Patel, Monika Gandhi, Pinal Trivedi, Maharshi Pandya, Nidhi Patel, Nitin Savaliya, Raghawendra Kumar, Dinesh Kumar, Zuber Saiyed, Komal Patel, Labdhi Pandya, Afzal Ansari, Nikha Trivedi, Naresh Chauhan, Summaiya Mullan, R D Dixit, A M Kadri, Harsh Bakshi, Chaitanya Joshi, Madhvi Joshi |
| EPI_ISL_476873 | Department of MicroBiology, Government Medical College, Surat | Gujarat Biotechnology Research Centre | Apurvasinh Puvar, Janvi Raval, Zarna Patel, Monika Gandhi, Pinal Trivedi, Maharshi Pandya, Nidhi Patel, Nitin Savaliya, Raghawendra Kumar, Dinesh Kumar, Zuber Saiyed, Komal Patel, Labdhi Pandya, Afzal Ansari, Nikha Trivedi, Naresh Chauhan, Summaiya Mullan, Amit gamit, R D Dixit, A M Kadri, Harsh Bakshi, Chaitanya Joshi, Madhvi Joshi |
| EPI_ISL_476874 | Department of MicroBiology, Government Medical College, Surat | Gujarat Biotechnology Research Centre | Janvi Raval, Zarna Patel, Monika Gandhi, Pinal Trivedi, Maharshi Pandya, Nidhi Patel, Nitin Savaliya, Raghawendra Kumar, Dinesh Kumar, Zuber Saiyed, Komal Patel, Labdhi Pandya, Afzal Ansari, Nikha Trivedi, Naresh Chauhan, Summaiya Mullan, Amit gamit, Apurvasinh Puvar, R D Dixit, A M Kadri, Harsh Bakshi, Chaitanya Joshi, Madhvi Joshi |
| EPI_ISL_476875 | Department of MicroBiology, Government Medical College, Surat | Gujarat Biotechnology Research Centre | Zarna Patel, Monika Gandhi, Pinal Trivedi, Maharshi Pandya, Nidhi Patel, Nitin Savaliya, Raghawendra Kumar, Dinesh Kumar, Zuber Saiyed, Komal Patel, Labdhi Pandya, Afzal Ansari, Nikha Trivedi, Naresh Chauhan, Summaiya Mullan, Amit gamit, Apurvasinh Puvar, Janvi Raval, R D Dixit, A M Kadri, Harsh Bakshi, Chaitanya Joshi, Madhvi Joshi |
| EPI_ISL_476876 | Department of MicroBiology, Government Medical College, Surat | Gujarat Biotechnology Research Centre | Pinal Trivedi, Maharshi Pandya, Nidhi Patel, Nitin Savaliya, Raghawendra Kumar, Dinesh Kumar, Zuber Saiyed, Komal Patel, Labdhi Pandya, Afzal Ansari, Nikha Trivedi, Naresh Chauhan, Summaiya Mullan, Amit gamit, Apurvasinh Puvar, Janvi Raval, Zarna Patel, Monika Gandhi, R D Dixit, A M Kadri, Harsh Bakshi, Chaitanya Joshi, Madhvi Joshi |
| EPI_ISL_476877 | Department of MicroBiology, Government Medical College, Surat | Gujarat Biotechnology Research Centre | Maharshi Pandya, Nidhi Patel, Nitin Savaliya, Raghawendra Kumar, Dinesh Kumar, Zuber Saiyed, Komal Patel, Labdhi Pandya, Afzal Ansari, Nikha Trivedi, Naresh Chauhan, Summaiya Mullan, Amit gamit, Apurvasinh Puvar, Janvi Raval, Zarna Patel, Monika Gandhi, Pinal Trivedi, R D Dixit, A M Kadri, Harsh Bakshi, Chaitanya Joshi, Madhvi Joshi |
| EPI_ISL_476878 | Department of MicroBiology, Government Medical College, Surat | Gujarat Biotechnology Research Centre | Nidhi Patel, Nitin Savaliya, Raghawendra Kumar, Dinesh Kumar, Zuber Saiyed, Komal Patel, Labdhi Pandya, Afzal Ansari, Nikha Trivedi, Naresh Chauhan, Summaiya Mullan, Amit gamit, Apurvasinh Puvar, Janvi Raval, Zarna Patel, Monika Gandhi, Pinal Trivedi, Maharshi Pandya, R D Dixit, A M Kadri, Harsh Bakshi, Chaitanya Joshi, Madhvi Joshi |
| EPI_ISL_476879 | Department of MicroBiology, Government Medical College, Surat | Gujarat Biotechnology Research Centre | Nitin Savaliya, Raghawendra Kumar, Dinesh Kumar, Zuber Saiyed, Komal Patel, Labdhi Pandya, Afzal Ansari, Nikha Trivedi, Naresh Chauhan, Summaiya Mullan, Amit gamit, Apurvasinh Puvar, Janvi Raval, Zarna Patel, Monika Gandhi, Pinal Trivedi, Maharshi Pandya, Nidhi Patel, R D Dixit, A M Kadri, Harsh Bakshi, Chaitanya Joshi, Madhvi Joshi |
| EPI_ISL_476880 | Department of MicroBiology, Government Medical College, Surat | Gujarat Biotechnology Research Centre | Raghawendra Kumar, Dinesh Kumar, Zuber Saiyed, Komal Patel, Labdhi Pandya, Afzal Ansari, Nikha Trivedi, Naresh Chauhan, Summaiya Mullan, Amit gamit, Apurvasinh Puvar, Janvi Raval, Zarna Patel, Monika Gandhi, Pinal Trivedi, Maharshi Pandya, Nidhi Patel, Nitin Savaliya, R D Dixit, A M Kadri, Harsh Bakshi, Chaitanya Joshi, Madhvi Joshi |
| EPI_ISL_476881 | Department of MicroBiology, Government Medical College, Surat | Gujarat Biotechnology Research Centre | Dinesh Kumar, Zuber Saiyed, Komal Patel, Labdhi Pandya, Afzal Ansari, Nikha Trivedi, Naresh Chauhan, Summaiya Mullan, Amit gamit, Apurvasinh Puvar, Janvi Raval, Zarna Patel, Monika Gandhi, Pinal Trivedi, Maharshi Pandya, Nidhi Patel, Nitin Savaliya, Raghawendra Kumar, R D Dixit, A M Kadri, Harsh Bakshi, Chaitanya Joshi, Madhvi Joshi |
| EPI_ISL_476882 | Department of MicroBiology, Government Medical College, Surat | Gujarat Biotechnology Research Centre | Zuber Saiyed, Komal Patel, Labdhi Pandya, Afzal Ansari, Nikha Trivedi, Naresh Chauhan, Summaiya Mullan, Amit gamit, Apurvasinh Puvar, Janvi Raval, Zarna Patel, Monika Gandhi, Pinal Trivedi, Maharshi Pandya, Nidhi Patel, Nitin Savaliya, Raghawendra Kumar, Dinesh Kumar, R D Dixit, A M Kadri, Harsh Bakshi, Chaitanya Joshi, Madhvi Joshi |
| EPI_ISL_476883, EPI_ISL_476884, EPI_ISL_476885, EPI_ISL_476886, EPI_ISL_476887, EPI_ISL_476888, EPI_ISL_476889, EPI_ISL_476890, EPI_ISL_476891, EPI_ISL_476892, EPI_ISL_476893, EPI_ISL_476894, EPI_ISL_476895, EPI_ISL_476896 | | | |
| see above | Defence Research & Development Establishment (DRDE) | Defence Research & Development Establishment (DRDE) | Shashi Sharma, Paban Kumar Dash, Sushil Kumar Sharma, Ambuj Shrivastava, Jyoti S. Kumar |
| EPI_ISL_476897 | University of South Carolina Functional Genomics Core | University of South Carolina Functional Genomics Core | Michael Shtutman |
| EPI_ISL_476898, EPI_ISL_476899 | Alaska State Virology Laboratory | Alaska State Virology Laboratory | Jack Chen, Ph.D. |
| EPI_ISL_476900, EPI_ISL_476901, EPI_ISL_476902, EPI_ISL_476903, EPI_ISL_476904, EPI_ISL_476905, EPI_ISL_476906, EPI_ISL_476907, EPI_ISL_476908, EPI_ISL_476909, EPI_ISL_476910, EPI_ISL_476911, EPI_ISL_476912, EPI_ISL_476913, EPI_ISL_476914, EPI_ISL_476915, EPI_ISL_476916, EPI_ISL_476917, EPI_ISL_476918, EPI_ISL_476919, EPI_ISL_476920, EPI_ISL_476921, EPI_ISL_476922, EPI_ISL_476923, EPI_ISL_476924, EPI_ISL_476925, EPI_ISL_476926, EPI_ISL_476927, EPI_ISL_476928, EPI_ISL_476929, EPI_ISL_476930, EPI_ISL_476931, EPI_ISL_476932, EPI_ISL_476933, EPI_ISL_476934, EPI_ISL_476935, EPI_ISL_476936, EPI_ISL_476937, EPI_ISL_476938, EPI_ISL_476939, EPI_ISL_476940 | | | |
| see above | UW Virology Lab | UW Virology Lab | Pavitra Roychoudhury, Hong Xie, Lasata Shrestha, Amin Addetia, Truong Nguyen, Victoria M Rachleff, Meei-Li Huang, Keith R Jerome, Alexander Greninger |
| EPI_ISL_476941, EPI_ISL_476942, EPI_ISL_476943, EPI_ISL_476944, EPI_ISL_476945, EPI_ISL_476946, EPI_ISL_476947, EPI_ISL_476948, EPI_ISL_476949, EPI_ISL_476950, EPI_ISL_476951, EPI_ISL_476952, EPI_ISL_476953, EPI_ISL_476954, EPI_ISL_476955, EPI_ISL_476956, EPI_ISL_476957, EPI_ISL_476958, EPI_ISL_476959, EPI_ISL_476960, EPI_ISL_476961, EPI_ISL_476962, EPI_ISL_476963, EPI_ISL_476964, EPI_ISL_476965, EPI_ISL_476966, EPI_ISL_476967, EPI_ISL_476968, EPI_ISL_476969, EPI_ISL_476970, EPI_ISL_476971, EPI_ISL_476972, EPI_ISL_476973, EPI_ISL_476974, EPI_ISL_476975, EPI_ISL_476976, | | | |

| | | | |
|---|---|---|---|
| EPI_ISL_476977, EPI_ISL_476978, EPI_ISL_476979, EPI_ISL_476980, EPI_ISL_476981, EPI_ISL_476982, EPI_ISL_476983, EPI_ISL_476984, EPI_ISL_476985, EPI_ISL_476986, EPI_ISL_476987, EPI_ISL_476988, EPI_ISL_476989, EPI_ISL_476990, EPI_ISL_476991, EPI_ISL_476992, EPI_ISL_476993, EPI_ISL_476994, EPI_ISL_476995, EPI_ISL_476996, EPI_ISL_476997, EPI_ISL_476998, EPI_ISL_476999, EPI_ISL_477000, EPI_ISL_477001, EPI_ISL_477002, EPI_ISL_477003, EPI_ISL_477004, EPI_ISL_477005, EPI_ISL_477006, EPI_ISL_477007 | | | |
| see above | KU Leuven, Rega Institute, Clinical and Epidemiological Virology | KU Leuven, Rega Institute, Clinical and Epidemiological Virology | Tony Wawina-Bokalanga, Joan Marti-Carerras, Bert Vanmechelen, Piet Maes |
| EPI_ISL_477008, EPI_ISL_477009, EPI_ISL_477010, EPI_ISL_477011, EPI_ISL_477012, EPI_ISL_477013 | University of Debrecen, Department of Medical Microbiology | National Laboratory of Virology, Szentágothai Research Centre | Endre Gábor Tóth, Balázs Somogyi, Brigitta Zana, Eszter Csoma, Ferenc Jakab, Gábor Kemenesi |
| EPI_ISL_477014 | Institute of Microbiology, Universidad San Francisco de Quito | Institute of Microbiology, Universidad San Francisco de Quito | Belen Prado-Vivar, Sully Marquez, Juan Jose Guadalupe, Monica Becerra-Wong, Carla Torres, Bernardo Gutierrez, Francisco Mora, Juan Gaviria, Alejandra Ramones, Franklin Espinoza, Edison Ligña, Jorge Reyes, Patricio Rojas-Silva, Veronica Barragan, Gabriel Trueba, Michelle Grunauer, Paul Cardenas |
| EPI_ISL_477015 | Institute of Microbiology, Universidad San Francisco de Quito | Institute of Microbiology, Universidad San Francisco de Quito | Sully Márquez, Belén Prado-Vivar, Juan José Guadalupe, Monica Becerra-Wong, Carla Torres, Bernardo Gutiérrez, Jorge Luis Velez, Verónica Barragán, Patricio Rojas-Silva, Gabriel Trueba, Michelle Grunauer, Paúl Cárdenas |
| EPI_ISL_477016 | Institute of Microbiology, Universidad San Francisco de Quito | Institute of Microbiology, Universidad San Francisco de Quito | Juan José Guadalupe, Sully Márquez, Belén Prado-Vivar, Monica Becerra-Wong, Carla Torres, Bernardo Gutiérrez, Jorge Luis Velez, Verónica Barragán, Patricio Rojas-Silva, Gabriel Trueba, Michelle Grunauer, Paúl Cárdenas |
| EPI_ISL_477017, EPI_ISL_477018, EPI_ISL_477019, EPI_ISL_477020, EPI_ISL_477021, EPI_ISL_477022, EPI_ISL_477023, EPI_ISL_477024, EPI_ISL_477025, EPI_ISL_477026, EPI_ISL_477027, EPI_ISL_477028, EPI_ISL_477029, EPI_ISL_477030, EPI_ISL_477031, EPI_ISL_477032, EPI_ISL_477033, EPI_ISL_477034, EPI_ISL_477035, EPI_ISL_477036, EPI_ISL_477037, EPI_ISL_477038, EPI_ISL_477039, EPI_ISL_477040, EPI_ISL_477041, EPI_ISL_477042, EPI_ISL_477043, EPI_ISL_477044, EPI_ISL_477045, EPI_ISL_477046, EPI_ISL_477047, EPI_ISL_477048, EPI_ISL_477049, EPI_ISL_477050, EPI_ISL_477051, EPI_ISL_477052, EPI_ISL_477053, EPI_ISL_477054, EPI_ISL_477055, EPI_ISL_477056, EPI_ISL_477057, EPI_ISL_477058, EPI_ISL_477059, EPI_ISL_477060, EPI_ISL_477061, EPI_ISL_477062, EPI_ISL_477063, EPI_ISL_477064, EPI_ISL_477065, EPI_ISL_477066, EPI_ISL_477067, EPI_ISL_477068, EPI_ISL_477069, EPI_ISL_477070, EPI_ISL_477071, EPI_ISL_477072, EPI_ISL_477073, EPI_ISL_477074, EPI_ISL_477075, EPI_ISL_477076, EPI_ISL_477077, EPI_ISL_477078, EPI_ISL_477079, EPI_ISL_477080, EPI_ISL_477081, EPI_ISL_477082, EPI_ISL_477083, EPI_ISL_477084, EPI_ISL_477085, EPI_ISL_477086, EPI_ISL_477087, EPI_ISL_477088, EPI_ISL_477089, EPI_ISL_477090, EPI_ISL_477091, EPI_ISL_477092, EPI_ISL_477093, EPI_ISL_477094, EPI_ISL_477095, EPI_ISL_477096, EPI_ISL_477097, EPI_ISL_477098, EPI_ISL_477099, EPI_ISL_477100, EPI_ISL_477101, EPI_ISL_477102, EPI_ISL_477103, EPI_ISL_477104, EPI_ISL_477105, EPI_ISL_477106, EPI_ISL_477107, EPI_ISL_477108, EPI_ISL_477109, EPI_ISL_477110, EPI_ISL_477111, EPI_ISL_477112, EPI_ISL_477113, EPI_ISL_477114, EPI_ISL_477115, EPI_ISL_477116, EPI_ISL_477117, EPI_ISL_477118, EPI_ISL_477119 | | | |
| see above | BCCDC Public Health Laboratory | BCCDC Public Health Laboratory | Richard Harrigan, Hope Lapointe, Jinny Choi, Kimia Kamelian, John Tyson,Terry Snutch, Linda Hoang, Inna Sekirov, Paul Levett, Mel Krajden, Natalie Prystajecky |
| EPI_ISL_477125, EPI_ISL_477126, EPI_ISL_477127, EPI_ISL_477128, EPI_ISL_477129, EPI_ISL_477130, EPI_ISL_477131, EPI_ISL_477132, EPI_ISL_477133, EPI_ISL_477134, EPI_ISL_477135, EPI_ISL_477136, EPI_ISL_477137, EPI_ISL_477138, EPI_ISL_477139, EPI_ISL_477140 | | | |
| see above | Child Health Research Foundation | Child Health Research Foundation | Senjuti Saha, Md Saiful Islam Sajib, Roly Malaker, Md Hafizur Rahman, Afroza Akter Tanni, Syed Muktadir Al Sium, Maksuda Islam, Samir K Saha |
| EPI_ISL_477141, EPI_ISL_477142, EPI_ISL_477143, EPI_ISL_477144, EPI_ISL_477145, EPI_ISL_477146, EPI_ISL_477147, EPI_ISL_477148, EPI_ISL_477149, EPI_ISL_477150, EPI_ISL_477151, EPI_ISL_477152, EPI_ISL_477153, EPI_ISL_477154, EPI_ISL_477155, EPI_ISL_477156, EPI_ISL_477157, EPI_ISL_477158, EPI_ISL_477159 | | | |
| see above | Institut Pasteur Dakar | Institut Pasteur de Dakar | Ndongo Dia, Moussa Moise Diagne, Mamadou Diop, Mamadou Malado Jallow, Marie Henriette Dior Ndione, Safietou Sankhe, Ousmane Faye, Amadou Alpha Sall. |
| EPI_ISL_477160 | Laboratory of Dr. John Lednicky | University of Florida | John A. Lednicky, Chang-Yu Wu, and John Glenn Morris, Jr. |
| EPI_ISL_477161 | unknown | Cancer Biology Department | Zekri,A.N., Amer,K.E., Ahmed,O.S., Soliman,H.K., Hafez,M.M., Bahnassy,A.A., Abdelhamid,W., Khattab,A., Ali,M., Hassan,W., Samir,M., Raouf,A., Hamdy,M.S., Soliman,M.S., Elsissy,M.H., Elkhateeb,S.M., Ezzelarab,M.H. and Abouelhoda,M. |
| EPI_ISL_477163 | Laboratory of Dr. John Lednicky | University of Florida | John A. Lednicky, Maha A. Elbadry, Kuttichantran Subramaniam, Thomas B. Waltzek, John Glenn Morris, Jr. |
| EPI_ISL_477164 | Department of Virology, Public Health Laboratories Division, National Institute of Health | Department of Virology, Public Health Laboratories Division, National Institute of Health | Nazish Badar, Aamer Ikram, Muhammad Salman, Hamza Ahmed Mirza, Abdul Ahad, Yasir Arshad, Massab Umair |
| EPI_ISL_477165, EPI_ISL_477166, EPI_ISL_477167 | Department of Virology, Public Health Laboratories Division, National Institute of Health | Department of Virology, Public Health Laboratories Division, National Institute of Health | Nazish Badar,Aamer Ikram, Muhammad Salman, Massab Umair, Hamza Ahmed Mirza, Abdul Ahad, Yasir Arshad |
| EPI_ISL_477168 | Institute for Stem Cell Science and Regenerative Medicine | National Centre for Biological Sciences | Farhan Ali, Vanessa Molin Paynter, Srikar Krishna, Mohak Sharda, Shah-e-Jahan Gulzar, Awadhesh Pandit, Varadha Sundarmurthy, Uma Ramakrishnan, Dasaradhi Palakodeti, Aswin Seshasayee |
| EPI_ISL_477170, EPI_ISL_477171, EPI_ISL_477172, EPI_ISL_477174, EPI_ISL_477175, EPI_ISL_477177, EPI_ISL_477178, EPI_ISL_477180, EPI_ISL_477182 | Department of Laboratory Medicine Tan Tock Seng Hospital | Department of Laboratory Medicine Tan Tock Seng Hospital | Chen YYC, Zair X, Li C, Tang WY, Maurer-Stroh S, Barkham TMS, Nagarajan N, Sessions OM |
| EPI_ISL_477183 | Department of MicroBiology, Government Medical College, Surat | Gujarat Biotechnology Research Centre | Monika Gandhi, Pinal Trivedi, Maharshi Pandya, Nidhi Patel, Nitin Savaliya, Raghawendra Kumar, Dinesh Kumar, Zuber Saiyed, Komal Patel, Labdhi Pandya, Afzal Ansari, Nikha Trivedi, Naresh Chauhan, Summaiya Mullan, Amit gamit, Apurvasinh Puvar, Janvi Raval, Zarna Patel, R D Dixit, A M Kadri, Harsh Bakshi, Chaitanya Joshi, Madhvi Joshi |
| EPI_ISL_477184, EPI_ISL_477187, EPI_ISL_477188, EPI_ISL_477189, EPI_ISL_477190, EPI_ISL_477191, EPI_ISL_477192 | Department of Laboratory Medicine Tan Tock Seng Hospital | Department of Laboratory Medicine Tan Tock Seng Hospital | Chen YYC, Zair X, Li C, Tang WY, Maurer-Stroh S, Barkham TMS, Nagarajan N, Sessions OM |
| EPI_ISL_477193, EPI_ISL_477194, EPI_ISL_477195, EPI_ISL_477196, EPI_ISL_477197, EPI_ISL_477198, EPI_ISL_477199, EPI_ISL_477200, EPI_ISL_477201, EPI_ISL_477202, EPI_ISL_477203 | | | |
| see above | Istituto Zooprofilattico Sperimentale Puglia e Basilicata; | Beaconlab (Bioinformatics, Evolution and Comparative Genomics lab), Dept of Biosciences, University on Mila | Parisi A.,Pesole G., Manzari C., Chiara M. |
| EPI_ISL_477204 | Prof. Massimo Zollo CEINGE TASK-FORCE COVID19 - Regione Campania | Prof. Massimo Zollo CEINGE TASK-FORCE COVID19 - Regione Campania | Veronica Ferrucci1,2, Dae young Kong8, Fatemeh asadzadeh1,2, Laura Marrone1,2, Roberto Siciliano1,2, Rino Cerino3, Giovanna Fusco3, Marika Comegna1,2, Angelo Boccia2, Maurizio Viscardi3, Giorgia Borriello3, Sergio Brandi3, Claudia Tiberio4, Luigi Atripaldi4, Giovanni Paolella1,2, Giuseppe Castaldo1,2, Stefano Pascarella4, Martina Bianchi4, Lorenzo Chiariotti1,2, Jae Myun Lee5, Jae Ho Jung6, Kyong Seop Yun7, Hong Yeoul Kim 7,8* and Massimo Zollo1,2* 1 CEINGE Biotecnologie Avanzate, Naples, Italia 2 Dipartimento di Medicina Molecolare e Biotecnologie Mediche DMMBM University of Naples Federico II, Italia 3 Istituto Zooprofilattico Sperimentale del Mezzogiorno, Naples, Italia 4 -U.O.C. di Patologia Clinica Ospedale D. Cotugno, Azienda Sanitaria Ospedali dei Colli, Naples, Italy. 5 Università La Sapienza di Roma, Italia 6 Department of Microbiology, Yonsei University College of Medicine, Seoul, Korea 7 Department of Surgery, Yonsei University College of Medicine, Seoul, Korea 8 Haim bio co., Ltd, , Indust |
| EPI_ISL_477205, EPI_ISL_477206, EPI_ISL_477207, EPI_ISL_477208, EPI_ISL_477209, EPI_ISL_477210, EPI_ISL_477211, EPI_ISL_477212, EPI_ISL_477213, EPI_ISL_477214, EPI_ISL_477215, EPI_ISL_477216, EPI_ISL_477217, EPI_ISL_477218, EPI_ISL_477219, EPI_ISL_477220, EPI_ISL_477221, EPI_ISL_477222, EPI_ISL_477223, EPI_ISL_477224, EPI_ISL_477225, EPI_ISL_477226, EPI_ISL_477227, EPI_ISL_477228, EPI_ISL_477229, EPI_ISL_477230, EPI_ISL_477231, EPI_ISL_477232, EPI_ISL_477233, EPI_ISL_477234, EPI_ISL_477235, EPI_ISL_477236, EPI_ISL_477237, EPI_ISL_477238, EPI_ISL_477239, EPI_ISL_477240, EPI_ISL_477241, EPI_ISL_477242, EPI_ISL_477243, EPI_ISL_477244, EPI_ISL_477245, EPI_ISL_477246, EPI_ISL_477247, EPI_ISL_477248, EPI_ISL_477249, EPI_ISL_477250, EPI_ISL_477251, EPI_ISL_477252, EPI_ISL_477253, EPI_ISL_477254, EPI_ISL_477255, EPI_ISL_477256, EPI_ISL_477257, EPI_ISL_477258, EPI_ISL_477259, EPI_ISL_477260, EPI_ISL_477261, EPI_ISL_477262, EPI_ISL_477263 | | | |
| see above | Institute for Stem Cell Science and Regenerative Medicine | National Centre for Biological Sciences | Farhan Ali, Vanessa Molin Paynter, Srikar Krishna, Mohak Sharda, Shah-e-Jahan Gulzar, Awadhesh Pandit, Varadha Sundarmurthy, Uma Ramakrishnan, Dasaradhi Palakodeti, Aswin Seshasayee |
| EPI_ISL_477272, EPI_ISL_477273, EPI_ISL_477274, EPI_ISL_477275, EPI_ISL_477276 | Mayo Clinic & Mayo Clinic Laboratories | Minnesota Department of Health, Public Health Laboratory | Matt Plumb, Jacob Garfin, Kelly Pung, and Xiong Wang |
| EPI_ISL_477277, EPI_ISL_477278, EPI_ISL_477279, EPI_ISL_477280, EPI_ISL_477281, EPI_ISL_477282, EPI_ISL_477283, EPI_ISL_477284, EPI_ISL_477285, EPI_ISL_477286, EPI_ISL_477287, EPI_ISL_477288, EPI_ISL_477289, EPI_ISL_477290 | | | |
| see above | M Health Fairview | Minnesota Department of Health, Public Health Laboratory | Matt Plumb, Jacob Garfin, Kelly Pung, and Xiong Wang |
| EPI_ISL_477291, EPI_ISL_477292, EPI_ISL_477293, EPI_ISL_477294, EPI_ISL_477295, EPI_ISL_477296, EPI_ISL_477297, EPI_ISL_477298, EPI_ISL_477299, EPI_ISL_477300, EPI_ISL_477301, EPI_ISL_477302, EPI_ISL_477303, EPI_ISL_477304, EPI_ISL_477305, EPI_ISL_477306, EPI_ISL_477307, EPI_ISL_477308, EPI_ISL_477309, EPI_ISL_477310, EPI_ISL_477311, EPI_ISL_477312 | | | |
| see above | Mayo Clinic & Mayo Clinic Laboratories | Minnesota Department of Health, Public Health Laboratory | Matt Plumb, Jacob Garfin, Kelly Pung, and Xiong Wang |
| EPI_ISL_477615, EPI_ISL_477616, EPI_ISL_477617, EPI_ISL_477618, EPI_ISL_477619, EPI_ISL_477620, EPI_ISL_477621, EPI_ISL_477622, EPI_ISL_477623, EPI_ISL_477624, EPI_ISL_477625 | | | |

| | | | |
|---|---|---|---|
| see above | University of Szeged, Institute of Clinical Microbiology | National Laboratory of Virology, Szentágothai Research Centre | Endre Gábor Tóth, Balázs Somogyi, Brigitta Zana, Terhes Gabriella, Ferenc Jakab, Gábor Kemenesi |

EPI_ISL_477631, EPI_ISL_477632, EPI_ISL_477633, EPI_ISL_477634, EPI_ISL_477635, EPI_ISL_477636, EPI_ISL_477637, EPI_ISL_477638, EPI_ISL_477639, EPI_ISL_477640, EPI_ISL_477641, EPI_ISL_477642, EPI_ISL_477643, EPI_ISL_477644, EPI_ISL_477645, EPI_ISL_477646, EPI_ISL_477647, EPI_ISL_477648, EPI_ISL_477649, EPI_ISL_477650, EPI_ISL_477651, EPI_ISL_477652, EPI_ISL_477653, EPI_ISL_477654, EPI_ISL_477655, EPI_ISL_477656, EPI_ISL_477657, EPI_ISL_477658, EPI_ISL_477659, EPI_ISL_477660, EPI_ISL_477661, EPI_ISL_477662, EPI_ISL_477663, EPI_ISL_477664, EPI_ISL_477665, EPI_ISL_477666, EPI_ISL_477667, EPI_ISL_477668, EPI_ISL_477669, EPI_ISL_477670, EPI_ISL_477671, EPI_ISL_477672

| | | | |
|---|---|---|---|
| see above | Virginia DCLS | Virginia DCLS | Virginia DCLS |

EPI_ISL_477673, EPI_ISL_477674, EPI_ISL_477675, EPI_ISL_477676, EPI_ISL_477677, EPI_ISL_477678, EPI_ISL_477679, EPI_ISL_477680, EPI_ISL_477681, EPI_ISL_477682, EPI_ISL_477683, EPI_ISL_477684, EPI_ISL_477685, EPI_ISL_477686, EPI_ISL_477687, EPI_ISL_477688, EPI_ISL_477689, EPI_ISL_477690, EPI_ISL_477691, EPI_ISL_477692, EPI_ISL_477693, EPI_ISL_477694, EPI_ISL_477695, EPI_ISL_477696, EPI_ISL_477697, EPI_ISL_477698, EPI_ISL_477699, EPI_ISL_477700, EPI_ISL_477701, EPI_ISL_477702, EPI_ISL_477703, EPI_ISL_477704, EPI_ISL_477705, EPI_ISL_477706, EPI_ISL_477707, EPI_ISL_477708, EPI_ISL_477709, EPI_ISL_477710, EPI_ISL_477711, EPI_ISL_477712, EPI_ISL_477713, EPI_ISL_477714, EPI_ISL_477715, EPI_ISL_477716, EPI_ISL_477717, EPI_ISL_477718, EPI_ISL_477719, EPI_ISL_477720, EPI_ISL_477721, EPI_ISL_477722, EPI_ISL_477723, EPI_ISL_477724

| | | | |
|---|---|---|---|
| see above | UW Virology Lab | UW Virology Lab | Pavitra Roychoudhury, Hong Xie, Lasata Shrestha, Amin Addetia, Truong Nguyen, Victoria M Rachleff, Meei-Li Huang, Keith R Jerome, Alexander Greninger |

EPI_ISL_477727, EPI_ISL_477728, EPI_ISL_477729, EPI_ISL_477730, EPI_ISL_477731, EPI_ISL_477732, EPI_ISL_477733, EPI_ISL_477734, EPI_ISL_477735, EPI_ISL_477736, EPI_ISL_477737, EPI_ISL_477738, EPI_ISL_477739, EPI_ISL_477740, EPI_ISL_477741, EPI_ISL_477742, EPI_ISL_477743, EPI_ISL_477744, EPI_ISL_477745, EPI_ISL_477746, EPI_ISL_477747, EPI_ISL_477748, EPI_ISL_477749, EPI_ISL_477750, EPI_ISL_477751, EPI_ISL_477752, EPI_ISL_477753, EPI_ISL_477754, EPI_ISL_477755, EPI_ISL_477756, EPI_ISL_477757, EPI_ISL_477758, EPI_ISL_477759, EPI_ISL_477760, EPI_ISL_477761, EPI_ISL_477762, EPI_ISL_477763, EPI_ISL_477764, EPI_ISL_477765, EPI_ISL_477766, EPI_ISL_477767, EPI_ISL_477768, EPI_ISL_477769, EPI_ISL_477770, EPI_ISL_477771, EPI_ISL_477772, EPI_ISL_477773, EPI_ISL_477774, EPI_ISL_477775, EPI_ISL_477776, EPI_ISL_477777, EPI_ISL_477778, EPI_ISL_477779, EPI_ISL_477780, EPI_ISL_477781, EPI_ISL_477782

| | | | |
|---|---|---|---|
| see above | University of Birmingham | COVID-19 Genomics UK (COG-UK) Consortium | Institute of Microbiology, University of Birmingham: Claire McMurray, Joanne Stockton, Samuel Nicholls, Radoslaw Poplawski, Will Rowe, Josh Quick, Nicholas Loman. University of Birmingham Testing Laboratory: Celina M Whalley, Andrew Bosworth, Charlotte Poxon, Kasun Wanigasooriya, Oliver Pickles, Mike Kidd, Alex Richter, Andrew D Beggs PHE Heartlands Lab: Husam Osman, Andrew Bosworth. Queen Elizabeth Hospital: Anna Casey |

EPI_ISL_477783, EPI_ISL_477784, EPI_ISL_477785, EPI_ISL_477786, EPI_ISL_477787, EPI_ISL_477788, EPI_ISL_477789, EPI_ISL_477790, EPI_ISL_477791, EPI_ISL_477792, EPI_ISL_477793, EPI_ISL_477794, EPI_ISL_477795, EPI_ISL_477796, EPI_ISL_477797, EPI_ISL_477798, EPI_ISL_477799, EPI_ISL_477800, EPI_ISL_477801, EPI_ISL_477802, EPI_ISL_477803, EPI_ISL_477804, EPI_ISL_477805, EPI_ISL_477806, EPI_ISL_477807, EPI_ISL_477808, EPI_ISL_477809, EPI_ISL_477810, EPI_ISL_477811, EPI_ISL_477812, EPI_ISL_477813, EPI_ISL_477814, EPI_ISL_477815, EPI_ISL_477816, EPI_ISL_477817, EPI_ISL_477818, EPI_ISL_477819

| | | | |
|---|---|---|---|
| see above | Department of Pathology, University of Cambridge | COVID-19 Genomics UK (COG-UK) Consortium | Luke W Meredith, M. Estée Török, Myra Hosmillo, William L. Hamilton, Martin D. Curran, Theresa Feltwell, Grant Hall, Anna Yakovleva, Fahad A Khokhar, Charlotte J. Houldcroft, Laura G Caller, Aminu S. Jahun, Sarah L. Caddy, Yasmin Chaudhry, Malte Pinckert, Ian Goodfellow |

EPI_ISL_477820, EPI_ISL_477821, EPI_ISL_477822, EPI_ISL_477823, EPI_ISL_477824, EPI_ISL_477825, EPI_ISL_477826, EPI_ISL_477827, EPI_ISL_477828, EPI_ISL_477829, EPI_ISL_477830, EPI_ISL_477831, EPI_ISL_477832, EPI_ISL_477833, EPI_ISL_477834, EPI_ISL_477835, EPI_ISL_477836, EPI_ISL_477837, EPI_ISL_477838, EPI_ISL_477839, EPI_ISL_477840, EPI_ISL_477841, EPI_ISL_477842, EPI_ISL_477843, EPI_ISL_477844, EPI_ISL_477845, EPI_ISL_477846, EPI_ISL_477847, EPI_ISL_477848, EPI_ISL_477849, EPI_ISL_477850, EPI_ISL_477851, EPI_ISL_477852, EPI_ISL_477853, EPI_ISL_477854, EPI_ISL_477855, EPI_ISL_477856, EPI_ISL_477857, EPI_ISL_477858, EPI_ISL_477859, EPI_ISL_477860, EPI_ISL_477861, EPI_ISL_477862, EPI_ISL_477863, EPI_ISL_477864, EPI_ISL_477865, EPI_ISL_477866, EPI_ISL_477867, EPI_ISL_477868, EPI_ISL_477869, EPI_ISL_477870, EPI_ISL_477871, EPI_ISL_477872, EPI_ISL_477873, EPI_ISL_477874, EPI_ISL_477875, EPI_ISL_477876, EPI_ISL_477877, EPI_ISL_477878, EPI_ISL_477879, EPI_ISL_477880, EPI_ISL_477881, EPI_ISL_477882, EPI_ISL_477883, EPI_ISL_477884, EPI_ISL_477885, EPI_ISL_477886, EPI_ISL_477887, EPI_ISL_477888, EPI_ISL_477889, EPI_ISL_477890, EPI_ISL_477891, EPI_ISL_477892, EPI_ISL_477893, EPI_ISL_477894, EPI_ISL_477895, EPI_ISL_477896, EPI_ISL_477897, EPI_ISL_477898, EPI_ISL_477899, EPI_ISL_477900, EPI_ISL_477901, EPI_ISL_477902, EPI_ISL_477903, EPI_ISL_477904, EPI_ISL_477905, EPI_ISL_477906, EPI_ISL_477907, EPI_ISL_477908, EPI_ISL_477909, EPI_ISL_477910, EPI_ISL_477911, EPI_ISL_477912, EPI_ISL_477913, EPI_ISL_477914, EPI_ISL_477915, EPI_ISL_477916, EPI_ISL_477917, EPI_ISL_477918, EPI_ISL_477919, EPI_ISL_477920, EPI_ISL_477921, EPI_ISL_477922, EPI_ISL_477923, EPI_ISL_477924, EPI_ISL_477925, EPI_ISL_477926, EPI_ISL_477927, EPI_ISL_477928, EPI_ISL_477929, EPI_ISL_477930, EPI_ISL_477931, EPI_ISL_477932, EPI_ISL_477933, EPI_ISL_477934, EPI_ISL_477935, EPI_ISL_477936, EPI_ISL_477937, EPI_ISL_477938, EPI_ISL_477939, EPI_ISL_477940, EPI_ISL_477941, EPI_ISL_477942, EPI_ISL_477943, EPI_ISL_477944, EPI_ISL_477945, EPI_ISL_477946, EPI_ISL_477947, EPI_ISL_477948, EPI_ISL_477949, EPI_ISL_477950, EPI_ISL_477951, EPI_ISL_477952, EPI_ISL_477953, EPI_ISL_477954, EPI_ISL_477955, EPI_ISL_477956, EPI_ISL_477957, EPI_ISL_477958, EPI_ISL_477959, EPI_ISL_477960, EPI_ISL_477961, EPI_ISL_477962, EPI_ISL_477963, EPI_ISL_477964, EPI_ISL_477965, EPI_ISL_477966, EPI_ISL_477967, EPI_ISL_477968, EPI_ISL_477969, EPI_ISL_477970, EPI_ISL_477971, EPI_ISL_477972, EPI_ISL_477973, EPI_ISL_477974, EPI_ISL_477975, EPI_ISL_477976, EPI_ISL_477977, EPI_ISL_477978, EPI_ISL_477979, EPI_ISL_477980, EPI_ISL_477981, EPI_ISL_477982, EPI_ISL_477983, EPI_ISL_477984, EPI_ISL_477985, EPI_ISL_477986, EPI_ISL_477987, EPI_ISL_477988, EPI_ISL_477989, EPI_ISL_477990, EPI_ISL_477991, EPI_ISL_477992, EPI_ISL_477993, EPI_ISL_477994, EPI_ISL_477995, EPI_ISL_477996, EPI_ISL_477997, EPI_ISL_477998, EPI_ISL_477999, EPI_ISL_478000, EPI_ISL_478001, EPI_ISL_478002, EPI_ISL_478003, EPI_ISL_478004, EPI_ISL_478005, EPI_ISL_478006, EPI_ISL_478007, EPI_ISL_478008, EPI_ISL_478009, EPI_ISL_478010, EPI_ISL_478011, EPI_ISL_478012, EPI_ISL_478013, EPI_ISL_478014, EPI_ISL_478015, EPI_ISL_478016, EPI_ISL_478017, EPI_ISL_478018, EPI_ISL_478019, EPI_ISL_478020, EPI_ISL_478021, EPI_ISL_478022, EPI_ISL_478023, EPI_ISL_478024, EPI_ISL_478025, EPI_ISL_478026, EPI_ISL_478027, EPI_ISL_478028, EPI_ISL_478029, EPI_ISL_478030, EPI_ISL_478031, EPI_ISL_478032, EPI_ISL_478033, EPI_ISL_478034, EPI_ISL_478035, EPI_ISL_478036, EPI_ISL_478037, EPI_ISL_478038, EPI_ISL_478039, EPI_ISL_478040, EPI_ISL_478041, EPI_ISL_478042, EPI_ISL_478043, EPI_ISL_478044, EPI_ISL_478045, EPI_ISL_478046, EPI_ISL_478047, EPI_ISL_478048, EPI_ISL_478049, EPI_ISL_478050, EPI_ISL_478051, EPI_ISL_478052, EPI_ISL_478053, EPI_ISL_478054, EPI_ISL_478055, EPI_ISL_478056, EPI_ISL_478057, EPI_ISL_478058, EPI_ISL_478059, EPI_ISL_478060, EPI_ISL_478061, EPI_ISL_478062, EPI_ISL_478063, EPI_ISL_478064, EPI_ISL_478065, EPI_ISL_478066, EPI_ISL_478067, EPI_ISL_478068, EPI_ISL_478069, EPI_ISL_478070, EPI_ISL_478071, EPI_ISL_478072, EPI_ISL_478073, EPI_ISL_478074, EPI_ISL_478075, EPI_ISL_478076, EPI_ISL_478077, EPI_ISL_478078, EPI_ISL_478079, EPI_ISL_478080, EPI_ISL_478081, EPI_ISL_478082, EPI_ISL_478083, EPI_ISL_478084, EPI_ISL_478085, EPI_ISL_478086, EPI_ISL_478087, EPI_ISL_478088, EPI_ISL_478089, EPI_ISL_478090, EPI_ISL_478091, EPI_ISL_478092, EPI_ISL_478093, EPI_ISL_478094, EPI_ISL_478095, EPI_ISL_478096, EPI_ISL_478097, EPI_ISL_478098, EPI_ISL_478099, EPI_ISL_478100, EPI_ISL_478101, EPI_ISL_478102, EPI_ISL_478103, EPI_ISL_478104, EPI_ISL_478105, EPI_ISL_478106, EPI_ISL_478107, EPI_ISL_478108, EPI_ISL_478109, EPI_ISL_478110, EPI_ISL_478111, EPI_ISL_478112, EPI_ISL_478113, EPI_ISL_478114, EPI_ISL_478115, EPI_ISL_478116, EPI_ISL_478117, EPI_ISL_478118, EPI_ISL_478119, EPI_ISL_478120, EPI_ISL_478121, EPI_ISL_478122, EPI_ISL_478123, EPI_ISL_478124, EPI_ISL_478125, EPI_ISL_478126, EPI_ISL_478127, EPI_ISL_478128, EPI_ISL_478129, EPI_ISL_478130, EPI_ISL_478131, EPI_ISL_478132, EPI_ISL_478133, EPI_ISL_478134, EPI_ISL_478135, EPI_ISL_478136, EPI_ISL_478137, EPI_ISL_478138, EPI_ISL_478139, EPI_ISL_478140, EPI_ISL_478141, EPI_ISL_478142, EPI_ISL_478143, EPI_ISL_478144, EPI_ISL_478145, EPI_ISL_478146, EPI_ISL_478147, EPI_ISL_478148, EPI_ISL_478149, EPI_ISL_478150, EPI_ISL_478151, EPI_ISL_478152, EPI_ISL_478153, EPI_ISL_478154, EPI_ISL_478155, EPI_ISL_478156, EPI_ISL_478157, EPI_ISL_478158, EPI_ISL_478159, EPI_ISL_478160, EPI_ISL_478161, EPI_ISL_478162

| | | | |
|---|---|---|---|
| see above | West of Scotland Specialist Virology Centre, NHSGGC / MRC-University of Glasgow Centre for Virus Research | COVID-19 Genomics UK (COG-UK) Consortium | Ana da Silva Filipe, Natasha Johnson, Kathy Smollett, Daniel Mair, Stephen Carmichael, Lily Tong, Jenna Nichols, Elihu Aranday-Cortes, Kirstyn Brunker, Yasmin Parr, Alice Broos, Kyriaki Nomikou; Sarah McDonald, Marc Niebel, Patawee Asamaphan; Richard Orton, Joseph Hughes, Sreenu Vattipally, David L Robertson; Alasdair MacLean, Rory Gunson; Kathy Li, Natasha Jesudason, Rajiv Shah, James Shepherd, Antonia Ho, Emma Thomson |

EPI_ISL_478163, EPI_ISL_478164, EPI_ISL_478165, EPI_ISL_478166, EPI_ISL_478167, EPI_ISL_478168, EPI_ISL_478169, EPI_ISL_478170, EPI_ISL_478171, EPI_ISL_478172, EPI_ISL_478173, EPI_ISL_478174, EPI_ISL_478175, EPI_ISL_478176, EPI_ISL_478177, EPI_ISL_478178, EPI_ISL_478179, EPI_ISL_478180, EPI_ISL_478181, EPI_ISL_478182, EPI_ISL_478183, EPI_ISL_478184, EPI_ISL_478185, EPI_ISL_478186, EPI_ISL_478187, EPI_ISL_478188, EPI_ISL_478189, EPI_ISL_478190, EPI_ISL_478191, EPI_ISL_478192, EPI_ISL_478193, EPI_ISL_478194, EPI_ISL_478195, EPI_ISL_478196, EPI_ISL_478197, EPI_ISL_478198, EPI_ISL_478199, EPI_ISL_478200, EPI_ISL_478201, EPI_ISL_478202, EPI_ISL_478203, EPI_ISL_478204, EPI_ISL_478205, EPI_ISL_478206, EPI_ISL_478207, EPI_ISL_478208, EPI_ISL_478209, EPI_ISL_478210, EPI_ISL_478211, EPI_ISL_478212, EPI_ISL_478213, EPI_ISL_478214, EPI_ISL_478215, EPI_ISL_478216, EPI_ISL_478217, EPI_ISL_478218, EPI_ISL_478219, EPI_ISL_478220, EPI_ISL_478221, EPI_ISL_478222, EPI_ISL_478223, EPI_ISL_478224, EPI_ISL_478225, EPI_ISL_478226, EPI_ISL_478227, EPI_ISL_478228, EPI_ISL_478229, EPI_ISL_478230, EPI_ISL_478231, EPI_ISL_478232, EPI_ISL_478233, EPI_ISL_478234, EPI_ISL_478235, EPI_ISL_478236, EPI_ISL_478237, EPI_ISL_478238, EPI_ISL_478239, EPI_ISL_478240, EPI_ISL_478241, EPI_ISL_478242, EPI_ISL_478243, EPI_ISL_478244, EPI_ISL_478245, EPI_ISL_478246, EPI_ISL_478247, EPI_ISL_478248, EPI_ISL_478249, EPI_ISL_478250, EPI_ISL_478251, EPI_ISL_478252, EPI_ISL_478253, EPI_ISL_478254, EPI_ISL_478255, EPI_ISL_478256, EPI_ISL_478257, EPI_ISL_478258, EPI_ISL_478259, EPI_ISL_478260, EPI_ISL_478261, EPI_ISL_478262, EPI_ISL_478263, EPI_ISL_478264, EPI_ISL_478265, EPI_ISL_478266, EPI_ISL_478267, EPI_ISL_478268, EPI_ISL_478269, EPI_ISL_478270, EPI_ISL_478271, EPI_ISL_478272, EPI_ISL_478273

| | | | |
|---|---|---|---|
| see above | Virology Department, Royal Infirmary of Edinburgh, NHS Lothian / School of Biological Sciences, University of Edinburgh / Institute of Genetics and Molecular Medicine, University of Edinburgh | COVID-19 Genomics UK (COG-UK) Consortium | McHugh M, Dewar R, Rooke S, Gallagher M, Balcaza C, O'Toole Á, Scher E, Hill V, McCrone JT, Colquhoun R, Yu X, Jackson B, Rambaut A, Williams TC, Templeton K |

EPI_ISL_478274, EPI_ISL_478275, EPI_ISL_478276, EPI_ISL_478277, EPI_ISL_478278, EPI_ISL_478279, EPI_ISL_478280, EPI_ISL_478281, EPI_ISL_478282, EPI_ISL_478283, EPI_ISL_478284, EPI_ISL_478285, EPI_ISL_478286, EPI_ISL_478287, EPI_ISL_478288

| | | | |
|---|---|---|---|
| see above | University of Exeter | COVID-19 Genomics UK (COG-UK) Consortium | Ben Temperton,Aaron Jeffries,Michelle Michelsen,Joanna Warwick-Dugdale,Audrey Farbos,Robyn Manley,Stephen Michell,Jane Masoli |

EPI_ISL_478289, EPI_ISL_478290, EPI_ISL_478291, EPI_ISL_478292, EPI_ISL_478293, EPI_ISL_478294, EPI_ISL_478295, EPI_ISL_478296, EPI_ISL_478297, EPI_ISL_478298, EPI_ISL_478299, EPI_ISL_478300, EPI_ISL_478301, EPI_ISL_478302, EPI_ISL_478303, EPI_ISL_478304, EPI_ISL_478305, EPI_ISL_478306, EPI_ISL_478307, EPI_ISL_478308, EPI_ISL_478309, EPI_ISL_478310, EPI_ISL_478311, EPI_ISL_478312, EPI_ISL_478313, EPI_ISL_478314, EPI_ISL_478315, EPI_ISL_478316, EPI_ISL_478317, EPI_ISL_478318, EPI_ISL_478319, EPI_ISL_478320, EPI_ISL_478321, EPI_ISL_478322, EPI_ISL_478323, EPI_ISL_478324, EPI_ISL_478325, EPI_ISL_478326, EPI_ISL_478327, EPI_ISL_478328, EPI_ISL_478329, EPI_ISL_478330, EPI_ISL_478331, EPI_ISL_478332, EPI_ISL_478333, EPI_ISL_478334, EPI_ISL_478335, EPI_ISL_478336, EPI_ISL_478337, EPI_ISL_478338, EPI_ISL_478339, EPI_ISL_478340, EPI_ISL_478341, EPI_ISL_478342, EPI_ISL_478343, EPI_ISL_478344, EPI_ISL_478345, EPI_ISL_478346, EPI_ISL_478347, EPI_ISL_478348, EPI_ISL_478349, EPI_ISL_478350, EPI_ISL_478351, EPI_ISL_478352, EPI_ISL_478353, EPI_ISL_478354, EPI_ISL_478355, EPI_ISL_478356, EPI_ISL_478357, EPI_ISL_478358, EPI_ISL_478359, EPI_ISL_478360, EPI_ISL_478361, EPI_ISL_478362, EPI_ISL_478363, EPI_ISL_478364, EPI_ISL_478365, EPI_ISL_478366, EPI_ISL_478367, EPI_ISL_478368, EPI_ISL_478369, EPI_ISL_478370, EPI_ISL_478371

| | | | |
|---|---|---|---|
| see above | University Hospitals Of Leicester NHS Trust and DeepSeq Nottingham | COVID-19 Genomics UK (COG-UK) Consortium | Christopher Holmes, Paul Bird, Thomas Helmer, Karlie Fallon, Julian Tang, Jonathan Ball, Patrick McClure, Joeseph Chappell, Nadine Holmes, Matthew Carlisle, Christopher Moore, Fei Sang, Johnny Debebe, Victoria Wright, Matthew Loose |

EPI_ISL_478372, EPI_ISL_478373, EPI_ISL_478374, EPI_ISL_478375, EPI_ISL_478376, EPI_ISL_478377, EPI_ISL_478378, EPI_ISL_478379, EPI_ISL_478380, EPI_ISL_478381, EPI_ISL_478382, EPI_ISL_478383, EPI_ISL_478384, EPI_ISL_478385, EPI_ISL_478386, EPI_ISL_478387, EPI_ISL_478388, EPI_ISL_478389, EPI_ISL_478390, EPI_ISL_478391, EPI_ISL_478392, EPI_ISL_478393, EPI_ISL_478394, EPI_ISL_478395, EPI_ISL_478396, EPI_ISL_478397, EPI_ISL_478398, EPI_ISL_478399, EPI_ISL_478400, EPI_ISL_478401, EPI_ISL_478402, EPI_ISL_478403

| | | | |
|---|---|---|---|
| see above | Liverpool Clinical Laboratories | COVID-19 Genomics UK (COG-UK) Consortium | Sam Haldenby, Anita Lucaci, Steve Paterson, Julian Hiscox, Alistair Darby, M Almsaud, A Alrezaihi, Muhannad Alruwaili, Stuart D Armstrong, Jones Benjamin, Eleanor G Bentley, Anu Chawla, Jordan J Clark, Angela Cowell, Richard Eccles, Isabel García-Dorival, Matthew Gemmell, Alessandro Gerada, PKF Gilmore, Richard Gregory, Ximeng Han, Catherine Hartley, Margaret Hughes, Miren Iturriza-Gomara, James Johnson, L Luu, Jenifer Manson, Charlotte Nelson, Elaine O'Toole, Cassie Olateju, Rebekah Penrice-Randal , Lucille Rainbow, N.P Randle, Trevor Ian Robinson, Parul Sharma, Ghada T Shawli, James P Stewart, Neil Swainston, Ecaterina Vamos, Joanne Watts, Mark Whitehead |

EPI_ISL_478404, EPI_ISL_478405, EPI_ISL_478406, EPI_ISL_478407, EPI_ISL_478408, EPI_ISL_478409, EPI_ISL_478410, EPI_ISL_478411, EPI_ISL_478412, EPI_ISL_478413, EPI_ISL_478414, EPI_ISL_478415, EPI_ISL_478416, EPI_ISL_478417, EPI_ISL_478418, EPI_ISL_478419, EPI_ISL_478420, EPI_ISL_478421, EPI_ISL_478422, EPI_ISL_478423, EPI_ISL_478424, EPI_ISL_478425, EPI_ISL_478426, EPI_ISL_478427, EPI_ISL_478428, EPI_ISL_478429, EPI_ISL_478430, EPI_ISL_478431, EPI_ISL_478432, EPI_ISL_478433, EPI_ISL_478434, EPI_ISL_478435, EPI_ISL_478436, EPI_ISL_478437, EPI_ISL_478438, EPI_ISL_478439, EPI_ISL_478440, EPI_ISL_478441, EPI_ISL_478442, EPI_ISL_478443, EPI_ISL_478444, EPI_ISL_478445, EPI_ISL_478446, EPI_ISL_478447, EPI_ISL_478448, EPI_ISL_478449, EPI_ISL_478450, EPI_ISL_478451, EPI_ISL_478452, EPI_ISL_478453, EPI_ISL_478454, EPI_ISL_478455, EPI_ISL_478456, EPI_ISL_478457, EPI_ISL_478458, EPI_ISL_478459, EPI_ISL_478460, EPI_ISL_478461, EPI_ISL_478462, EPI_ISL_478463, EPI_ISL_478464, EPI_ISL_478465, EPI_ISL_478466, EPI_ISL_478467, EPI_ISL_478468, EPI_ISL_478469, EPI_ISL_478470, EPI_ISL_478471, EPI_ISL_478472

| | | | |
|---|---|---|---|
| see above | University College London, Great Ormond Street Hospital for Children NHS Foundation Trust, Imperial College Healthcare NHS Trust | COVID-19 Genomics UK (COG-UK) Consortium | Sergi Castellano, Rachel Williams, Mark Kristiansen, Paola Resende Silva, Sunando Roy, Tony Brooks, Helena Tutill, Paola Niola, Patricia Dyal, Charlotte Williams, Leysa Forrest, Yasmin Panchbhaya, Jacqueline Findlay, Samuel Weeks, Julianne Brown, Kathryn Harris, Paul Randell, James Price, Alison Holmes, Judith Breuer |
| EPI_ISL_478473, EPI_ISL_478474, EPI_ISL_478475, EPI_ISL_478476, EPI_ISL_478477, EPI_ISL_478478, EPI_ISL_478479, EPI_ISL_478480, EPI_ISL_478481, EPI_ISL_478482, EPI_ISL_478483, EPI_ISL_478484, EPI_ISL_478485, EPI_ISL_478486, EPI_ISL_478487, EPI_ISL_478488, EPI_ISL_478489, EPI_ISL_478490, EPI_ISL_478491, EPI_ISL_478492, EPI_ISL_478493, EPI_ISL_478494, EPI_ISL_478495, EPI_ISL_478496, EPI_ISL_478497, EPI_ISL_478498, EPI_ISL_478499, EPI_ISL_478500, EPI_ISL_478501, EPI_ISL_478502, EPI_ISL_478503, EPI_ISL_478504, EPI_ISL_478505, EPI_ISL_478506, EPI_ISL_478507, EPI_ISL_478508, EPI_ISL_478509, EPI_ISL_478510, EPI_ISL_478511, EPI_ISL_478512, EPI_ISL_478513, EPI_ISL_478514, EPI_ISL_478515, EPI_ISL_478516, EPI_ISL_478517, EPI_ISL_478518, EPI_ISL_478519, EPI_ISL_478520, EPI_ISL_478521, EPI_ISL_478522, EPI_ISL_478523, EPI_ISL_478524, EPI_ISL_478525, EPI_ISL_478526, EPI_ISL_478527, EPI_ISL_478528, EPI_ISL_478529, EPI_ISL_478530, EPI_ISL_478531, EPI_ISL_478532, EPI_ISL_478533, EPI_ISL_478534, EPI_ISL_478535, EPI_ISL_478536, EPI_ISL_478537, EPI_ISL_478538, EPI_ISL_478539, EPI_ISL_478540, EPI_ISL_478541, EPI_ISL_478542, EPI_ISL_478543, EPI_ISL_478544, EPI_ISL_478545, EPI_ISL_478546, EPI_ISL_478547, EPI_ISL_478548, EPI_ISL_478549, EPI_ISL_478550, EPI_ISL_478551, EPI_ISL_478552, EPI_ISL_478553, EPI_ISL_478554, EPI_ISL_478555, EPI_ISL_478556, EPI_ISL_478557, EPI_ISL_478558, EPI_ISL_478559, EPI_ISL_478560, EPI_ISL_478561, EPI_ISL_478562, EPI_ISL_478563, EPI_ISL_478564, EPI_ISL_478565, EPI_ISL_478566, EPI_ISL_478567, EPI_ISL_478568, EPI_ISL_478569, EPI_ISL_478570, EPI_ISL_478571, EPI_ISL_478572, EPI_ISL_478573, EPI_ISL_478574, EPI_ISL_478575, EPI_ISL_478576, EPI_ISL_478577, EPI_ISL_478578, EPI_ISL_478579, EPI_ISL_478580, EPI_ISL_478581, EPI_ISL_478582, EPI_ISL_478583, EPI_ISL_478584, EPI_ISL_478585, EPI_ISL_478586, EPI_ISL_478587, EPI_ISL_478588, EPI_ISL_478589, EPI_ISL_478590, EPI_ISL_478591, EPI_ISL_478592, EPI_ISL_478593, EPI_ISL_478594, EPI_ISL_478595, EPI_ISL_478596, EPI_ISL_478597, EPI_ISL_478598, EPI_ISL_478599, EPI_ISL_478600, EPI_ISL_478601, EPI_ISL_478602, EPI_ISL_478603, EPI_ISL_478604, EPI_ISL_478605, EPI_ISL_478606, EPI_ISL_478607, EPI_ISL_478608, EPI_ISL_478609, EPI_ISL_478610, EPI_ISL_478611, EPI_ISL_478612, EPI_ISL_478613, EPI_ISL_478614, EPI_ISL_478615, EPI_ISL_478616, EPI_ISL_478617, EPI_ISL_478618, EPI_ISL_478619, EPI_ISL_478620, EPI_ISL_478621, EPI_ISL_478622, EPI_ISL_478623, EPI_ISL_478624, EPI_ISL_478625, EPI_ISL_478626, EPI_ISL_478627, EPI_ISL_478628, EPI_ISL_478629, EPI_ISL_478630, EPI_ISL_478631, EPI_ISL_478632, EPI_ISL_478633, EPI_ISL_478634, EPI_ISL_478635, EPI_ISL_478636, EPI_ISL_478637, EPI_ISL_478638, EPI_ISL_478639, EPI_ISL_478640, EPI_ISL_478641, EPI_ISL_478642, EPI_ISL_478643, EPI_ISL_478644, EPI_ISL_478645, EPI_ISL_478646, EPI_ISL_478647, EPI_ISL_478648, EPI_ISL_478649, EPI_ISL_478650, EPI_ISL_478651, EPI_ISL_478652, EPI_ISL_478653, EPI_ISL_478654, EPI_ISL_478655, EPI_ISL_478656, EPI_ISL_478657, EPI_ISL_478658, EPI_ISL_478659, EPI_ISL_478660, EPI_ISL_478661, EPI_ISL_478662, EPI_ISL_478663, EPI_ISL_478664, EPI_ISL_478665, EPI_ISL_478666, EPI_ISL_478667, EPI_ISL_478668 | | | |
| see above | Northumbria University / South Tees Hospitals NHS Foundation Trust / North Cumbria Integrated Care NHS Foundation Trust / North Tees and Hartlepool NHS Foundation Trust / Newcastle Hospitals NHS Foundation Trust | COVID-19 Genomics UK (COG-UK) Consortium | Darren L Smith,Andrew Nelson,Matthew Bashton,Greg R Young,Joshua Loh,John Allan,Mohammad A Tariq,Giles S Holt,Gary Black,Wen C Yew,Lynn Dover,Paul Baker,Steve Liggett,Sarah Essex,Jane Greenaway,Debra Padgett,Clive Graham,Garren Scott,Edward Barton,Emma Swindells,Brendan Payne,Jennifer Collins,Yusri Taha,Gary Eltringham |
| EPI_ISL_478669, EPI_ISL_478670 | unknown | Microbiology, Koc University | Can,F., Ozer,B., Nurtop,E., Dogan,O. |
| EPI_ISL_478671 | unknown | Molecular and Cell Biology | Baray,J.C., Mahmud,A., Khan,M.R., Nag,K., Sultana,N. |
| EPI_ISL_478673, EPI_ISL_478674 | Pathology North - Royal North Shore Hospital - NSW Health Pathology | NSW Health Pathology - Institute of Clinical Pathology and Medical Research; Westmead Hospital; University of Sydney | CIDM-PH et al. |
| EPI_ISL_478675, EPI_ISL_478676, EPI_ISL_478677, EPI_ISL_478678, EPI_ISL_478679, EPI_ISL_478680, EPI_ISL_478681, EPI_ISL_478682 | Sydney South West Pathology Service (SSWPS) - Liverpool Hospital - NSW Health Pathology | NSW Health Pathology - Institute of Clinical Pathology and Medical Research; Westmead Hospital; University of Sydney | CIDM-PH et al. |
| EPI_ISL_478683, EPI_ISL_478684, EPI_ISL_478685, EPI_ISL_478686, EPI_ISL_478687, EPI_ISL_478688, EPI_ISL_478689, EPI_ISL_478690, EPI_ISL_478691, EPI_ISL_478692, EPI_ISL_478693, EPI_ISL_478694, EPI_ISL_478695, EPI_ISL_478696, EPI_ISL_478697, EPI_ISL_478698, EPI_ISL_478699, EPI_ISL_478700, EPI_ISL_478701, EPI_ISL_478702, EPI_ISL_478703, EPI_ISL_478704, EPI_ISL_478705 | | | |
| see above | South Eastern Area Laboratory Services (SEALS) | NSW Health Pathology - Institute of Clinical Pathology and Medical Research; Westmead Hospital; University of Sydney | CIDM-PH et al. |
| EPI_ISL_478706, EPI_ISL_478707 | Sydney South West Pathology Service (SSWPS) - Liverpool Hospital - NSW Health Pathology | NSW Health Pathology - Institute of Clinical Pathology and Medical Research; Westmead Hospital; University of Sydney | CIDM-PH et al. |
| EPI_ISL_478708, EPI_ISL_478709, EPI_ISL_478710, EPI_ISL_478711 | South Eastern Area Laboratory Services (SEALS) | NSW Health Pathology - Institute of Clinical Pathology and Medical Research; Westmead Hospital; University of Sydney | CIDM-PH et al. |
| EPI_ISL_478712, EPI_ISL_478713, EPI_ISL_478714, EPI_ISL_478715, EPI_ISL_478716 | Sydney South West Pathology Service (SSWPS) - Liverpool Hospital - NSW Health Pathology | NSW Health Pathology - Institute of Clinical Pathology and Medical Research; Westmead Hospital; University of Sydney | CIDM-PH et al. |
| EPI_ISL_478717, EPI_ISL_478718 | Sydney South West Pathology Service (SSWPS) - Concord Repatriation General Hospital - NSW Health Pathology | NSW Health Pathology - Institute of Clinical Pathology and Medical Research; Westmead Hospital; University of Sydney | CIDM-PH et al. |
| EPI_ISL_478719, EPI_ISL_478720 | Quadram Institute Bioscience | COVID-19 Genomics UK (COG-UK) Consortium | Dave J. Baker, Gemma L. Kay, Alp Aydin, Thanh Le-Viet, Steven Rudder, Ana P. Tedim, Anastasia Kolyva, Maria Diaz, Leonardo de Oliveira Martins, Nabil-Fareed Alikhan, Lizzie Meadows, Rachael Stanley, Ngozi Elumogo, Muhammed Yasir, Nicholas M. Thomson, Alexander J Trotter, Rachel Gilroy, Samuel Bloomfield, Claire Stuart, Andrew Bell, Reenesh Prakash, Samir Dervisevic, Alison E. Mather, John Wain, Mark Webber, Andrew J. Page, Justin O'Grady |
| EPI_ISL_478721, EPI_ISL_478722, EPI_ISL_478723, EPI_ISL_478724, EPI_ISL_478725, EPI_ISL_478726, EPI_ISL_478727, EPI_ISL_478728 | Queens Medical Centre, Clinical Microbiology Department / DeepSeq Nottingham | COVID-19 Genomics UK (COG-UK) Consortium | Gemma Clark, Wendy Smith, Manjinder Khakh, Vicki M Fleming, Michelle M Lister, Hannah Howson-Wells, Jonathan Ball, Patrick McClure, Joseph Chappell, Theocharis Tsoleridis, Nadine Holmes, Matthew Carlisle, Christopher Moore, Fei Sang, Johnny Debebe, Victoria Wright, Matthew Loose |
| EPI_ISL_478729, EPI_ISL_478730, EPI_ISL_478731, EPI_ISL_478732, EPI_ISL_478733, EPI_ISL_478734, EPI_ISL_478735, EPI_ISL_478736, EPI_ISL_478737, EPI_ISL_478738, EPI_ISL_478739, EPI_ISL_478740, EPI_ISL_478741, EPI_ISL_478742, EPI_ISL_478743, EPI_ISL_478744, EPI_ISL_478745, EPI_ISL_478746, EPI_ISL_478747, EPI_ISL_478748, EPI_ISL_478749, EPI_ISL_478750, EPI_ISL_478751, EPI_ISL_478752, EPI_ISL_478753, EPI_ISL_478754, EPI_ISL_478755, EPI_ISL_478756, EPI_ISL_478757, EPI_ISL_478758, EPI_ISL_478759, EPI_ISL_478760, EPI_ISL_478761, EPI_ISL_478762, EPI_ISL_478763, EPI_ISL_478764, EPI_ISL_478765, EPI_ISL_478766, EPI_ISL_478767, EPI_ISL_478768, EPI_ISL_478769, EPI_ISL_478770, EPI_ISL_478771, EPI_ISL_478772, EPI_ISL_478773, EPI_ISL_478774, EPI_ISL_478775, EPI_ISL_478776, EPI_ISL_478777, EPI_ISL_478778, EPI_ISL_478779, EPI_ISL_478780, EPI_ISL_478781, EPI_ISL_478782, EPI_ISL_478783, EPI_ISL_478784, EPI_ISL_478785, EPI_ISL_478786, EPI_ISL_478787, EPI_ISL_478788, EPI_ISL_478789, EPI_ISL_478790, EPI_ISL_478791, EPI_ISL_478792, EPI_ISL_478793, EPI_ISL_478794, EPI_ISL_478795, EPI_ISL_478796, EPI_ISL_478797, EPI_ISL_478798, EPI_ISL_478799, EPI_ISL_478800, EPI_ISL_478801, EPI_ISL_478802, EPI_ISL_478803, EPI_ISL_478804, EPI_ISL_478805, EPI_ISL_478806, EPI_ISL_478807, EPI_ISL_478808, EPI_ISL_478809, EPI_ISL_478810, EPI_ISL_478811, EPI_ISL_478812, EPI_ISL_478813, EPI_ISL_478814, EPI_ISL_478815, EPI_ISL_478816, EPI_ISL_478817, EPI_ISL_478818, EPI_ISL_478819, EPI_ISL_478820, EPI_ISL_478821, EPI_ISL_478822, EPI_ISL_478823, EPI_ISL_478824, EPI_ISL_478825, EPI_ISL_478826, EPI_ISL_478827, EPI_ISL_478828, EPI_ISL_478829, EPI_ISL_478830, EPI_ISL_478831, EPI_ISL_478832, EPI_ISL_478833, EPI_ISL_478834, EPI_ISL_478835, EPI_ISL_478836, EPI_ISL_478837, EPI_ISL_478838, EPI_ISL_478839, EPI_ISL_478840, EPI_ISL_478841, EPI_ISL_478842, EPI_ISL_478843, EPI_ISL_478844, EPI_ISL_478845, EPI_ISL_478846, EPI_ISL_478847, EPI_ISL_478848, EPI_ISL_478849, EPI_ISL_478850, EPI_ISL_478851, EPI_ISL_478852, EPI_ISL_478853, EPI_ISL_478854, EPI_ISL_478855, EPI_ISL_478856, EPI_ISL_478857, EPI_ISL_478858, EPI_ISL_478859, EPI_ISL_478860, EPI_ISL_478861, EPI_ISL_478862, EPI_ISL_478863, EPI_ISL_478864, EPI_ISL_478865, EPI_ISL_478866, EPI_ISL_478867, EPI_ISL_478868, EPI_ISL_478869, EPI_ISL_478870, EPI_ISL_478871, EPI_ISL_478872, EPI_ISL_478873, EPI_ISL_478874, EPI_ISL_478875, EPI_ISL_478876, EPI_ISL_478877, EPI_ISL_478878, EPI_ISL_478879, EPI_ISL_478880, EPI_ISL_478881, EPI_ISL_478882, EPI_ISL_478883, EPI_ISL_478884, EPI_ISL_478885, EPI_ISL_478886, EPI_ISL_478887, EPI_ISL_478888, EPI_ISL_478889, EPI_ISL_478890, EPI_ISL_478891, EPI_ISL_478892, EPI_ISL_478893, EPI_ISL_478894, EPI_ISL_478895, EPI_ISL_478896, EPI_ISL_478897, EPI_ISL_478898, EPI_ISL_478899, EPI_ISL_478900, EPI_ISL_478901, EPI_ISL_478902, EPI_ISL_478903, EPI_ISL_478904, EPI_ISL_478905, EPI_ISL_478906, EPI_ISL_478907, EPI_ISL_478908, EPI_ISL_478909, EPI_ISL_478910, EPI_ISL_478911, EPI_ISL_478912, EPI_ISL_478913, EPI_ISL_478914, EPI_ISL_478915, EPI_ISL_478916, EPI_ISL_478917, EPI_ISL_478918, EPI_ISL_478919, EPI_ISL_478920, EPI_ISL_478921, EPI_ISL_478922, EPI_ISL_478923, EPI_ISL_478924, EPI_ISL_478925, EPI_ISL_478926, EPI_ISL_478927, EPI_ISL_478928, EPI_ISL_478929, EPI_ISL_478930, EPI_ISL_478931, EPI_ISL_478932, EPI_ISL_478933, EPI_ISL_478934, EPI_ISL_478935, EPI_ISL_478936, EPI_ISL_478937, EPI_ISL_478938, EPI_ISL_478939, EPI_ISL_478940, EPI_ISL_478941, EPI_ISL_478942, EPI_ISL_478943, EPI_ISL_478944, EPI_ISL_478945, EPI_ISL_478946, EPI_ISL_478947, EPI_ISL_478948, EPI_ISL_478949, EPI_ISL_478950, EPI_ISL_478951, EPI_ISL_478952, EPI_ISL_478953, EPI_ISL_478954, EPI_ISL_478955, EPI_ISL_478956, EPI_ISL_478957, EPI_ISL_478958, EPI_ISL_478959, EPI_ISL_478960, EPI_ISL_478961, EPI_ISL_478962, EPI_ISL_478963, EPI_ISL_478964, EPI_ISL_478965, EPI_ISL_478966, EPI_ISL_478967, EPI_ISL_478968, EPI_ISL_478969, EPI_ISL_478970, EPI_ISL_478971, EPI_ISL_478972, EPI_ISL_478973, EPI_ISL_478974, EPI_ISL_478975, EPI_ISL_478976, EPI_ISL_478977, EPI_ISL_478978, EPI_ISL_478979, EPI_ISL_478980, EPI_ISL_478981, EPI_ISL_478982, EPI_ISL_478983, EPI_ISL_478984, EPI_ISL_478985, EPI_ISL_478986, EPI_ISL_478987, EPI_ISL_478988, EPI_ISL_478989, EPI_ISL_478990, EPI_ISL_478991, EPI_ISL_478992, EPI_ISL_478993, EPI_ISL_478994, EPI_ISL_478995, EPI_ISL_478996, EPI_ISL_478997, EPI_ISL_478998, EPI_ISL_478999, EPI_ISL_479000, EPI_ISL_479001, EPI_ISL_479002, EPI_ISL_479003, EPI_ISL_479004, EPI_ISL_479005, EPI_ISL_479006, EPI_ISL_479007, EPI_ISL_479008, EPI_ISL_479009, EPI_ISL_479010, EPI_ISL_479011, EPI_ISL_479012, EPI_ISL_479013, EPI_ISL_479014, EPI_ISL_479015, EPI_ISL_479016, EPI_ISL_479017, EPI_ISL_479018, EPI_ISL_479019, EPI_ISL_479020, EPI_ISL_479021, EPI_ISL_479022, EPI_ISL_479023, EPI_ISL_479024, EPI_ISL_479025, EPI_ISL_479026, EPI_ISL_479027, EPI_ISL_479028, EPI_ISL_479029, EPI_ISL_479030, EPI_ISL_479031, EPI_ISL_479032, EPI_ISL_479033, EPI_ISL_479034, EPI_ISL_479035, EPI_ISL_479036, EPI_ISL_479037, EPI_ISL_479038, EPI_ISL_479039, EPI_ISL_479040, EPI_ISL_479041, EPI_ISL_479042, EPI_ISL_479043, EPI_ISL_479044, EPI_ISL_479045, EPI_ISL_479046, EPI_ISL_479047, EPI_ISL_479048, EPI_ISL_479049, EPI_ISL_479050, EPI_ISL_479051, EPI_ISL_479052, EPI_ISL_479053, EPI_ISL_479054, EPI_ISL_479055, EPI_ISL_479056, EPI_ISL_479057, EPI_ISL_479058, EPI_ISL_479059, EPI_ISL_479060, EPI_ISL_479061, EPI_ISL_479062, EPI_ISL_479063, EPI_ISL_479064, EPI_ISL_479065, EPI_ISL_479066, EPI_ISL_479067, EPI_ISL_479068, EPI_ISL_479069, EPI_ISL_479070, EPI_ISL_479071, EPI_ISL_479072, EPI_ISL_479073, EPI_ISL_479074, EPI_ISL_479075, EPI_ISL_479076, EPI_ISL_479077, EPI_ISL_479078, EPI_ISL_479079, EPI_ISL_479080, EPI_ISL_479081, EPI_ISL_479082, EPI_ISL_479083, EPI_ISL_479084, EPI_ISL_479085, EPI_ISL_479086, EPI_ISL_479087, EPI_ISL_479088, EPI_ISL_479089, EPI_ISL_479090, EPI_ISL_479091, EPI_ISL_479092, EPI_ISL_479093, EPI_ISL_479094, EPI_ISL_479095, EPI_ISL_479096, EPI_ISL_479097, EPI_ISL_479098, EPI_ISL_479099, EPI_ISL_479100, EPI_ISL_479101, EPI_ISL_479102, EPI_ISL_479103, EPI_ISL_479104, EPI_ISL_479105, EPI_ISL_479106, EPI_ISL_479107, EPI_ISL_479108, EPI_ISL_479109, EPI_ISL_479110, EPI_ISL_479111, EPI_ISL_479112, EPI_ISL_479113, EPI_ISL_479114, EPI_ISL_479115, EPI_ISL_479116, EPI_ISL_479117, EPI_ISL_479118, EPI_ISL_479119, EPI_ISL_479120, EPI_ISL_479121, EPI_ISL_479122, EPI_ISL_479123, EPI_ISL_479124, EPI_ISL_479125, EPI_ISL_479126, EPI_ISL_479127, EPI_ISL_479128, EPI_ISL_479129, EPI_ISL_479130, EPI_ISL_479131, EPI_ISL_479132, EPI_ISL_479133, EPI_ISL_479134, EPI_ISL_479135, EPI_ISL_479136, EPI_ISL_479137, EPI_ISL_479138, EPI_ISL_479139, EPI_ISL_479140, EPI_ISL_479141, EPI_ISL_479142, EPI_ISL_479143, EPI_ISL_479144, EPI_ISL_479145, EPI_ISL_479146, EPI_ISL_479147, EPI_ISL_479148, EPI_ISL_479149, EPI_ISL_479150, EPI_ISL_479151, EPI_ISL_479152, EPI_ISL_479153, EPI_ISL_479154, EPI_ISL_479155, EPI_ISL_479156, EPI_ISL_479157, EPI_ISL_479158, EPI_ISL_479159, EPI_ISL_479160, EPI_ISL_479161, EPI_ISL_479162, EPI_ISL_479163, EPI_ISL_479164, EPI_ISL_479165, EPI_ISL_479166, EPI_ISL_479167, EPI_ISL_479168, EPI_ISL_479169, EPI_ISL_479170, EPI_ISL_479171, EPI_ISL_479172, EPI_ISL_479173, EPI_ISL_479174 | | | |
| see above | Oxford Viromics, NDM, University of Oxford; Oxford University Hospitals; Basingstoke and North Hampshire Hospital | COVID-19 Genomics UK (COG-UK) Consortium | Tanya Golubchik, David Bonsall, George Macintyre, Amy Trebes, Mariateresa de Cesare, Catrin Moore, Alex Mobbs, Anita Justice, Robert Shaw, Monique Andersson, Timothy Peto, Emma Wise, Nathan Moore, Jessica Lynch, Nick Cortes, Matilde Mori, Stephen Kidd, David Buck, John Todd, Christophe Fraser |

| | | | |
|---|---|---|---|
| EPI_ISL_479175, EPI_ISL_479176, EPI_ISL_479177, EPI_ISL_479178, EPI_ISL_479179, EPI_ISL_479180, EPI_ISL_479181, EPI_ISL_479182, EPI_ISL_479183, EPI_ISL_479184, EPI_ISL_479185, EPI_ISL_479186, EPI_ISL_479187, EPI_ISL_479188, EPI_ISL_479189, EPI_ISL_479190, EPI_ISL_479191, EPI_ISL_479192, EPI_ISL_479193, EPI_ISL_479194 | | | |
| see above | Centre for Enzyme Innovation, University of Portsmouth / Translational Research Laboratory, Portsmouth Hospitals NHS Trust | COVID-19 Genomics UK (COG-UK) Consortium | Angela Beckett,Yann Bourgeois,Garry Scarlett,Sharon Glaysher,Scott Elliott,Kelly Bicknell,Robert Impey,Allyson Lloyd,Sarah Wyllie,Ethan Butcher,Anoop Chauhan,Samuel Robson |
| EPI_ISL_479195, EPI_ISL_479196, EPI_ISL_479197, EPI_ISL_479198, EPI_ISL_479199, EPI_ISL_479200, EPI_ISL_479201, EPI_ISL_479202, EPI_ISL_479203, EPI_ISL_479204, EPI_ISL_479205, EPI_ISL_479206, EPI_ISL_479207, EPI_ISL_479208, EPI_ISL_479209, EPI_ISL_479210, EPI_ISL_479211, EPI_ISL_479212, EPI_ISL_479213, EPI_ISL_479214, EPI_ISL_479215, EPI_ISL_479216, EPI_ISL_479217, EPI_ISL_479218, EPI_ISL_479219, EPI_ISL_479220, EPI_ISL_479221, EPI_ISL_479222, EPI_ISL_479223, EPI_ISL_479224, EPI_ISL_479225, EPI_ISL_479226, EPI_ISL_479227, EPI_ISL_479228, EPI_ISL_479229, EPI_ISL_479230, EPI_ISL_479231, EPI_ISL_479232, EPI_ISL_479233, EPI_ISL_479234, EPI_ISL_479235, EPI_ISL_479236, EPI_ISL_479237, EPI_ISL_479238, EPI_ISL_479239, EPI_ISL_479240, EPI_ISL_479241, EPI_ISL_479242, EPI_ISL_479243, EPI_ISL_479244, EPI_ISL_479245, EPI_ISL_479246, EPI_ISL_479247, EPI_ISL_479248, EPI_ISL_479249, EPI_ISL_479250, EPI_ISL_479251, EPI_ISL_479252, EPI_ISL_479253, EPI_ISL_479254, EPI_ISL_479255, EPI_ISL_479256, EPI_ISL_479257, EPI_ISL_479258, EPI_ISL_479259, EPI_ISL_479260, EPI_ISL_479261, EPI_ISL_479262, EPI_ISL_479263, EPI_ISL_479264, EPI_ISL_479265, EPI_ISL_479266, EPI_ISL_479267, EPI_ISL_479268, EPI_ISL_479269, EPI_ISL_479270, EPI_ISL_479271, EPI_ISL_479272, EPI_ISL_479273, EPI_ISL_479274, EPI_ISL_479275, EPI_ISL_479276, EPI_ISL_479277, EPI_ISL_479278, EPI_ISL_479279, EPI_ISL_479280, EPI_ISL_479281, EPI_ISL_479282, EPI_ISL_479283 | | | |
| see above | Virology Department, Sheffield Teaching Hospitals NHS Foundation Trust/Department of Infection, Immunity and Cardiovascular Disease, The Medical School, University of Sheffield | COVID-19 Genomics UK (COG-UK) Consortium | Thushan de Silva, Matthew Parker, Nikki Smith, Adri Angyal, Rebecca Brown, Luke Green, Rachel Tucker, Paul Parsons, Danielle Groves, Katie Johnson, Laura Carrilero, Alex Keeley, Dave Partridge, Matthew Wyles, Benjamin Lindsey, Mehmet Yavuz, Mohammad Raza, Cariad Evans |
| EPI_ISL_479284, EPI_ISL_479285, EPI_ISL_479286, EPI_ISL_479287, EPI_ISL_479288, EPI_ISL_479289, EPI_ISL_479290, EPI_ISL_479291, EPI_ISL_479292, EPI_ISL_479293, EPI_ISL_479294, EPI_ISL_479295, EPI_ISL_479296, EPI_ISL_479297, EPI_ISL_479298, EPI_ISL_479299, EPI_ISL_479300, EPI_ISL_479301, EPI_ISL_479302, EPI_ISL_479303, EPI_ISL_479304, EPI_ISL_479305, EPI_ISL_479306, EPI_ISL_479307, EPI_ISL_479308, EPI_ISL_479309, EPI_ISL_479310, EPI_ISL_479311, EPI_ISL_479312, EPI_ISL_479313, EPI_ISL_479314, EPI_ISL_479315, EPI_ISL_479316, EPI_ISL_479317, EPI_ISL_479318, EPI_ISL_479319, EPI_ISL_479320, EPI_ISL_479321, EPI_ISL_479322, EPI_ISL_479323, EPI_ISL_479324, EPI_ISL_479325, EPI_ISL_479326, EPI_ISL_479327, EPI_ISL_479328, EPI_ISL_479329, EPI_ISL_479330, EPI_ISL_479331, EPI_ISL_479332, EPI_ISL_479333, EPI_ISL_479334, EPI_ISL_479335, EPI_ISL_479336, EPI_ISL_479337, EPI_ISL_479338, EPI_ISL_479339, EPI_ISL_479340, EPI_ISL_479341, EPI_ISL_479342, EPI_ISL_479343, EPI_ISL_479344, EPI_ISL_479345, EPI_ISL_479346, EPI_ISL_479347, EPI_ISL_479348, EPI_ISL_479349, EPI_ISL_479350, EPI_ISL_479351, EPI_ISL_479352, EPI_ISL_479353, EPI_ISL_479354, EPI_ISL_479355, EPI_ISL_479356, EPI_ISL_479357, EPI_ISL_479358, EPI_ISL_479359, EPI_ISL_479360, EPI_ISL_479361, EPI_ISL_479362, EPI_ISL_479363, EPI_ISL_479364, EPI_ISL_479365, EPI_ISL_479366, EPI_ISL_479367, EPI_ISL_479368, EPI_ISL_479369, EPI_ISL_479370, EPI_ISL_479371, EPI_ISL_479372, EPI_ISL_479373, EPI_ISL_479374, EPI_ISL_479375, EPI_ISL_479376, EPI_ISL_479377, EPI_ISL_479378, EPI_ISL_479379, EPI_ISL_479380, EPI_ISL_479381, EPI_ISL_479382, EPI_ISL_479383, EPI_ISL_479384, EPI_ISL_479385, EPI_ISL_479386, EPI_ISL_479387, EPI_ISL_479388, EPI_ISL_479389, EPI_ISL_479390, EPI_ISL_479391, EPI_ISL_479392, EPI_ISL_479393, EPI_ISL_479394, EPI_ISL_479395, EPI_ISL_479396, EPI_ISL_479397, EPI_ISL_479398, EPI_ISL_479399, EPI_ISL_479400, EPI_ISL_479401, EPI_ISL_479402, EPI_ISL_479403, EPI_ISL_479404, EPI_ISL_479405, EPI_ISL_479406, EPI_ISL_479407, EPI_ISL_479408, EPI_ISL_479409, EPI_ISL_479410, EPI_ISL_479411, EPI_ISL_479412, EPI_ISL_479413, EPI_ISL_479414, EPI_ISL_479415, EPI_ISL_479416, EPI_ISL_479417, EPI_ISL_479418, EPI_ISL_479419, EPI_ISL_479420, EPI_ISL_479421, EPI_ISL_479422, EPI_ISL_479423, EPI_ISL_479424, EPI_ISL_479425, EPI_ISL_479426, EPI_ISL_479427, EPI_ISL_479428, EPI_ISL_479429, EPI_ISL_479430, EPI_ISL_479431, EPI_ISL_479432, EPI_ISL_479433, EPI_ISL_479434, EPI_ISL_479435, EPI_ISL_479436, EPI_ISL_479437, EPI_ISL_479438, EPI_ISL_479439, EPI_ISL_479440, EPI_ISL_479441, EPI_ISL_479442, EPI_ISL_479443, EPI_ISL_479444, EPI_ISL_479445, EPI_ISL_479446, EPI_ISL_479447, EPI_ISL_479448, EPI_ISL_479449, EPI_ISL_479450, EPI_ISL_479451, EPI_ISL_479452, EPI_ISL_479453, EPI_ISL_479454, EPI_ISL_479455, EPI_ISL_479456, EPI_ISL_479457, EPI_ISL_479458, EPI_ISL_479459, EPI_ISL_479460, EPI_ISL_479461, EPI_ISL_479462, EPI_ISL_479463, EPI_ISL_479464, EPI_ISL_479465, EPI_ISL_479466, EPI_ISL_479467, EPI_ISL_479468, EPI_ISL_479469, EPI_ISL_479470, EPI_ISL_479471, EPI_ISL_479472, EPI_ISL_479473, EPI_ISL_479474, EPI_ISL_479475, EPI_ISL_479476, EPI_ISL_479477, EPI_ISL_479478, EPI_ISL_479479, EPI_ISL_479480, EPI_ISL_479481 | | | |
| see above | Wales Specialist Virology Centre Sequencing lab: Pathogen Genomics Unit | COVID-19 Genomics UK (COG-UK) Consortium | Catherine Moore, Johnathan Evans, Laura Gifford, Malorie Perry, Simon Cottrell, Angela Marchbank, Alec Birchley, Alexander Adams, Amy Gaskin, Bree Gatica-Wilcox, Jason Coombes, Joel Southgate, Lauren Gilbert, Lee Graham, Nicole Pacchiarini, Sara Kumziene-Summerhayes, Sarah Taylor, Sophie Jones, Sara Rey, Matthew Bull, Joanne Watkins, Sally Corden, Tom Connor |
| EPI_ISL_479482, EPI_ISL_479483, EPI_ISL_479484, EPI_ISL_479485, EPI_ISL_479486, EPI_ISL_479487, EPI_ISL_479488, EPI_ISL_479489, EPI_ISL_479490, EPI_ISL_479491, EPI_ISL_479492 | | | |
| see above | Department of Laboratory Medicine Tan Tock Seng Hospital | Department of Laboratory Medicine Tan Tock Seng Hospital | Chen YYC, Zair X, Li C, Tang WY, Maurer-Stroh S, Barkham TMS, Nagarajan N, Sessions OM |
| EPI_ISL_479493, EPI_ISL_479494, EPI_ISL_479495, EPI_ISL_479496, EPI_ISL_479497, EPI_ISL_479498, EPI_ISL_479499, EPI_ISL_479500, EPI_ISL_479501, EPI_ISL_479502, EPI_ISL_479503, EPI_ISL_479504, EPI_ISL_479505, EPI_ISL_479506, EPI_ISL_479507, EPI_ISL_479508, EPI_ISL_479509, EPI_ISL_479510, EPI_ISL_479511, EPI_ISL_479512, EPI_ISL_479513, EPI_ISL_479514, EPI_ISL_479515, EPI_ISL_479516, EPI_ISL_479517, EPI_ISL_479518, EPI_ISL_479519, EPI_ISL_479520, EPI_ISL_479521, EPI_ISL_479522, EPI_ISL_479523, EPI_ISL_479524, EPI_ISL_479525, EPI_ISL_479526, EPI_ISL_479527, EPI_ISL_479528, EPI_ISL_479529, EPI_ISL_479530, EPI_ISL_479531, EPI_ISL_479532, EPI_ISL_479533, EPI_ISL_479534, EPI_ISL_479535, EPI_ISL_479536, EPI_ISL_479537, EPI_ISL_479538, EPI_ISL_479539, EPI_ISL_479540, EPI_ISL_479541, EPI_ISL_479542, EPI_ISL_479543, EPI_ISL_479544, EPI_ISL_479545, EPI_ISL_479546, EPI_ISL_479547, EPI_ISL_479548, EPI_ISL_479549, EPI_ISL_479550, EPI_ISL_479551, EPI_ISL_479552, EPI_ISL_479553, EPI_ISL_479554, EPI_ISL_479555, EPI_ISL_479556, EPI_ISL_479557, EPI_ISL_479558, EPI_ISL_479559, EPI_ISL_479560, EPI_ISL_479561, EPI_ISL_479562, EPI_ISL_479563, EPI_ISL_479564, EPI_ISL_479565, EPI_ISL_479566, EPI_ISL_479567, EPI_ISL_479568, EPI_ISL_479569, EPI_ISL_479570, EPI_ISL_479571, EPI_ISL_479572, EPI_ISL_479573 | | | |
| see above | NIV Influenza | NIV Influenza | Potdar V |
| EPI_ISL_479574, EPI_ISL_479575, EPI_ISL_479576, EPI_ISL_479577, EPI_ISL_479578, EPI_ISL_479579, EPI_ISL_479580, EPI_ISL_479581, EPI_ISL_479582, EPI_ISL_479583, EPI_ISL_479584, EPI_ISL_479585, EPI_ISL_479586, EPI_ISL_479587, EPI_ISL_479588, EPI_ISL_479589, EPI_ISL_479590, EPI_ISL_479591, EPI_ISL_479592, EPI_ISL_479593, EPI_ISL_479594, EPI_ISL_479595, EPI_ISL_479596, EPI_ISL_479597, EPI_ISL_479598, EPI_ISL_479599, EPI_ISL_479600, EPI_ISL_479601, EPI_ISL_479602, EPI_ISL_479603 | | | |
| see above | National Public Health Laboratory, National Centre for Infectious Diseases | National Public Health Laboratory, National Centre for Infectious Diseases | Mak TM, Octavia S, Zhou Z, Chavatte JM, Cui L, Lin RTP |
| EPI_ISL_479616, EPI_ISL_479617 | Laboratory of Molecular Virology of the International Centre for Genetic Engineering and Biotechnology (ICGEB) | ARGO Open Lab Platform for Genome Sequencing | Licastro, D, Rajasekharan S, Dal Monego S, Segat L, D'Agaro P, Salton F, Confalonieri P, Confalonieri M Marcello A |
| EPI_ISL_479618 | Laboratory of Molecular Virology of the International Centre for Genetic Engineering and Biotechnology (ICGEB) | ARGO Open Lab Platform for Genome Sequencing | Licastro, D, Rajasekharan S, Dal Monego S, Segat L, D'Agaro P, Salton F, Confalonieri P, Confalonieri M, Marcello A |
| EPI_ISL_479619 | Laboratory of Molecular Virology of the International Centre for Genetic Engineering and Biotechnology (ICGEB) | ARGO Open Lab Platform for Genome Sequencing | Licastro, D, Rajasekharan S, Dal Monego S, Segat L, D'Agaro P, Salton F, Confalonieri P, Confalonieri M, Marcello A |
| EPI_ISL_479620, EPI_ISL_479621, EPI_ISL_479622, EPI_ISL_479623, EPI_ISL_479624 | Molecular diagnostic laboratory of Federal Budget Institution of Science "Central Research Institute of Epidemiology" of The Federal Service on Customers' Rights Protection and Human Well-being Surveillance | Group of Genomics and Postgenomic Technologies of Central Research Institute of Epidemilology | Speranskaya AS, Kaptelova VV,Valdokhina AV, Bulanenko VP, Samoilov AE, Korneenko EV, Sizova TV, Tivanova EV, Shipulina OY, Akimkin VG |
| EPI_ISL_479625, EPI_ISL_479626, EPI_ISL_479627, EPI_ISL_479628, EPI_ISL_479629, EPI_ISL_479630, EPI_ISL_479631, EPI_ISL_479632, EPI_ISL_479633, EPI_ISL_479634, EPI_ISL_479635, EPI_ISL_479636, EPI_ISL_479637, EPI_ISL_479638, EPI_ISL_479639, EPI_ISL_479640, EPI_ISL_479641, EPI_ISL_479642, EPI_ISL_479643, EPI_ISL_479644, EPI_ISL_479645, EPI_ISL_479646, EPI_ISL_479647, EPI_ISL_479648, EPI_ISL_479649, EPI_ISL_479650, EPI_ISL_479651 | | | |
| see above | Dr. Georges-L.-Dumont University Hospital Centre | National Microbiology Laboratory | Anna Majer, Shari Tyson, Grace Seo, Kristyn Burak, Philip Mabon, Elsie Grudeski, Rhiannon Huzarewich, Russell Mandes, Jennifer Tanner, Natalie Knox, Morag Graham, Gary Van Domselaar, Richard Garceau, Guillaume Desnoyers, Nathalie Bastien, Yan Li, Timothy Booth |
| EPI_ISL_479657, EPI_ISL_479658, EPI_ISL_479659, EPI_ISL_479660, EPI_ISL_479661 | NIV Influenza | NIV Influenza | Potdar V |
| EPI_ISL_479662 | unknown | College of Veterinary Medicine, Chungnam National University | Seo,S. |
| EPI_ISL_479663, EPI_ISL_479664, EPI_ISL_479665, EPI_ISL_479666, EPI_ISL_479667, EPI_ISL_479668, EPI_ISL_479669, EPI_ISL_479670, EPI_ISL_479671, EPI_ISL_479672, EPI_ISL_479673, EPI_ISL_479674, EPI_ISL_479675 | | | |
| see above | unknown | Center for Genomics and System Biology, New York University | Roder,A., Banakis,S., Johnson,K., Khalfan,M., Borenstein,E.S., Samanovic,M., Cornelius,A., Herati,R., Ulrich,R., Fleming,A., Kottkamp,A., Raabe,V., Mulligan,M.J., Gresham,D., Ghedin,E. |
| EPI_ISL_479728 | unknown | Cancer Biology Department, National Cancer Institute | Zekri,A.N., Amer,K.E., Ahmed,O.S., Soliman,H.K., Bahnassy,A.A., Ali,M., Abdelhamid,W., Gad,A., Hassan,W., Samir,M., Raouf,A., Hamdy,M.S., Soliman,M.S., Elsissy,M.H., Elkhateeb,S.M., Ezzelarab,M.H., Abouelhoda,M. |
| EPI_ISL_479729, EPI_ISL_479730, EPI_ISL_479731, EPI_ISL_479732, EPI_ISL_479733, EPI_ISL_479734, EPI_ISL_479735 | unknown | Cancer Biology Department, National Cancer Institute | Zekri,A.N., Amer,K.E., Ahmed,O.S., Soliman,H.K., Hafez,M.M., Bahnassy,A.A., Abdelhamid,W., Gad,A., Ali,M., Hassan,W., Samir,M., Raouf,A., Hamdy,M.S., Soliman,M.S., Elsissy,M.H., Elkhateeb,S.M., Ezzelarab,M.H., Abouelhoda,M. |
| EPI_ISL_479736, EPI_ISL_479737, EPI_ISL_479738, EPI_ISL_479739, EPI_ISL_479740, EPI_ISL_479741, EPI_ISL_479742, EPI_ISL_479743, EPI_ISL_479744, EPI_ISL_479745, EPI_ISL_479746, EPI_ISL_479747, EPI_ISL_479748, EPI_ISL_479749, EPI_ISL_479750, EPI_ISL_479751, EPI_ISL_479752, EPI_ISL_479753, EPI_ISL_479754, EPI_ISL_479755 | | | |
| see above | Institute for Stem Cell Science and Regenerative Medicine | National Centre for Biological Sciences | Farhan Ali, Vanessa Molin Paynter, Srikar Krishna, Mohak Sharda, Shah-e-Jahan Gulzar, Awadhesh Pandit, Varadha Sundarmurthy, Uma Ramakrishnan, Dasaradhi Palakodeti, Aswin Seshasayee |

| | | | |
|---|---|---|---|
| EPI_ISL_479756, EPI_ISL_479757, EPI_ISL_479758 | National Institute of Hygiene and Epidemiology (NIHE) | National Key Laboratory of Gene Technology, Institute of Biotechnology (IBT) | Le Tung Lam, Nguyen Hong Trang, Ho Thi Thuong, Tran Huyen Linh, Ung Thi Hong Trang, Le Thi Thanh, Nguyen Vu Son, Vuong Duc Cuong, Tran Thu Huong, Pham Thi Hien, Nguyen Phuong Anh, Nguyen Le Khanh Hang, Hoang Vu Mai Phuong, Hoang Ha, Taichiro Takemura, Futoshi Hasebe, Chu Hoang Ha, Le Quynh Mai, Dang Duc Anh, Truong Nam Hai |
| EPI_ISL_479759, EPI_ISL_479760, EPI_ISL_479761, EPI_ISL_479762, EPI_ISL_479763, EPI_ISL_479764, EPI_ISL_479765, EPI_ISL_479766, EPI_ISL_479767, EPI_ISL_479768, EPI_ISL_479769, EPI_ISL_479770, EPI_ISL_479771, EPI_ISL_479772, EPI_ISL_479773, EPI_ISL_479774, EPI_ISL_479775 | | | |
| see above | University of Miami Immunology and Histocompatibility Laboratory | University of Miami Immunology and Histocompatibility Laboratory | Emilio Margolles-Clark, PhD and Phillip Ruiz, MD, PhD |
| EPI_ISL_479776 | NIV Influenza | NIV Influenza | Potdar V |
| EPI_ISL_479777, EPI_ISL_479778, EPI_ISL_479779, EPI_ISL_479780, EPI_ISL_479781, EPI_ISL_479782, EPI_ISL_479783, EPI_ISL_479784, EPI_ISL_479785, EPI_ISL_479786, EPI_ISL_479787, EPI_ISL_479788, EPI_ISL_479789 | | | |
| see above | Breuer Lab, UCL | Breuer Lab, UCL | Breuer Lab |
| EPI_ISL_479790 | Laboratory of Molecular Virology of the International Centre for Genetic Engineering and Biotechnology (ICGEB) | ARGO Open Lab Platform for Genome Sequencing | Licastro, D, Rajasekharan S, Dal Monego S, Segat L, D'Agaro P, Salton F, Confalonieri P, Confalonieri M, Marcello A |
| EPI_ISL_479791 | Laboratory of Molecular Virology of the International Centre for Genetic Engineering and Biotechnology (ICGEB) | ARGO Open Lab Platform for Genome Sequencing | Licastro, D, Rajasekharan S, Dal Monego S, Segat L, D'Agaro P, Salton F, Confalonieri P, Confalonieri M, Marcello A |
| EPI_ISL_479792, EPI_ISL_479793, EPI_ISL_479794, EPI_ISL_479795 | Hokkaido Institute of Public Health | Pathogen Genomics Center, National Institute of Infectious Diseases | Tsuyoshi Sekizuka, Rika Komagome, Kentaro Itokawa, Rina Tanaka, Masanori Hashino, Hajime Kamiya, Motoi Suzuki, Makoto Kuroda |
| EPI_ISL_479796 | Ishikawa Prefectural Institute of Public Health and Environmental Science | Pathogen Genomics Center, National Institute of Infectious Diseases | Tsuyoshi Sekizuka, Sanae Kuramoto, Eri Nariai, Kentaro Itokawa, Rina Tanaka, Masanori Hashino, Hajime Kamiya, Motoi Suzuki, Makoto Kuroda |
| EPI_ISL_479797, EPI_ISL_479798 | Sagamihara City Public Health Research Institute | Pathogen Genomics Center, National Institute of Infectious Diseases | Tsuyoshi Sekizuka, Hiroshi Nakamura, Kentaro Itokawa, Rina Tanaka, Masanori Hashino, Hajime Kamiya, Motoi Suzuki, Makoto Kuroda |
| EPI_ISL_479799, EPI_ISL_479800 | Sapporo City Institute of Public Health | Pathogen Genomics Center, National Institute of Infectious Diseases | Tsuyoshi Sekizuka, Asami Ohnishi, Kentaro Itokawa, Rina Tanaka, Masanori Hashino, Hajime Kamiya, Motoi Suzuki, Makoto Kuroda |
| EPI_ISL_479801 | Hokkaido Institute of Public Health | Pathogen Genomics Center, National Institute of Infectious Diseases | Tsuyoshi Sekizuka, Rika Komagome, Kentaro Itokawa, Rina Tanaka, Masanori Hashino, Hajime Kamiya, Motoi Suzuki, Makoto Kuroda |
| EPI_ISL_479802, EPI_ISL_479803, EPI_ISL_479804 | Sagamihara City Public Health Research Institute | Pathogen Genomics Center, National Institute of Infectious Diseases | Tsuyoshi Sekizuka, Hiroshi Nakamura, Kentaro Itokawa, Rina Tanaka, Masanori Hashino, Hajime Kamiya, Motoi Suzuki, Makoto Kuroda |
| EPI_ISL_479805, EPI_ISL_479806, EPI_ISL_479807, EPI_ISL_479808 | Saitama Prefectural Institute of Public Health | Pathogen Genomics Center, National Institute of Infectious Diseases | Tsuyoshi Sekizuka, Hayato Ehara, Kentaro Itokawa, Rina Tanaka, Masanori Hashino, Hajime Kamiya, Motoi Suzuki, Makoto Kuroda |
| EPI_ISL_479809, EPI_ISL_479810, EPI_ISL_479811 | Chiba Prefectural Institute of Public Health | Pathogen Genomics Center, National Institute of Infectious Diseases | Tsuyoshi Sekizuka, Masakatsu Taira, Kentaro Itokawa, Rina Tanaka, Masanori Hashino, Hajime Kamiya, Motoi Suzuki, Makoto Kuroda |
| EPI_ISL_479812, EPI_ISL_479813, EPI_ISL_479814, EPI_ISL_479815, EPI_ISL_479816, EPI_ISL_479817, EPI_ISL_479818, EPI_ISL_479819, EPI_ISL_479820 | Hokkaido Institute of Public Health | Pathogen Genomics Center, National Institute of Infectious Diseases | Tsuyoshi Sekizuka, Rika Komagome, Kentaro Itokawa, Rina Tanaka, Masanori Hashino, Hajime Kamiya, Motoi Suzuki, Makoto Kuroda |
| EPI_ISL_479821, EPI_ISL_479822 | Department of Infectious Diseases, Kobe Institute of Health | Pathogen Genomics Center, National Institute of Infectious Diseases | Tsuyoshi Sekizuka, Ryohei Nomoto, Kentaro Itokawa, Rina Tanaka, Masanori Hashino, Hajime Kamiya, Motoi Suzuki, Makoto Kuroda |
| EPI_ISL_479823 | Kochi Prefectural Institute of Public Health | Pathogen Genomics Center, National Institute of Infectious Diseases | Tsuyoshi Sekizuka, Akihiko Tokaji, Kentaro Itokawa, Rina Tanaka, Masanori Hashino, Hajime Kamiya, Motoi Suzuki, Makoto Kuroda |
| EPI_ISL_479824 | Kumamoto PrefecturalInstituteof Public-Health and Environmental Science | Pathogen Genomics Center, National Institute of Infectious Diseases | Tsuyoshi Sekizuka, Shunsuke Yahiro, Kentaro Itokawa, Rina Tanaka, Masanori Hashino, Hajime Kamiya, Motoi Suzuki, Makoto Kuroda |
| EPI_ISL_479825 | Tokyo Metropolitan Institute of Public Health | Pathogen Genomics Center, National Institute of Infectious Diseases | Tsuyoshi Sekizuka, Kenji Sadamasu, Takashi Chiba, Mami Nagashima, Kentaro Itokawa, Rina Tanaka, Masanori Hashino, Hajime Kamiya, Motoi Suzuki, Makoto Kuroda |
| EPI_ISL_479826, EPI_ISL_479827, EPI_ISL_479828, EPI_ISL_479829, EPI_ISL_479830, EPI_ISL_479831, EPI_ISL_479832, EPI_ISL_479833, EPI_ISL_479834, EPI_ISL_479835, EPI_ISL_479836, EPI_ISL_479837, EPI_ISL_479838, EPI_ISL_479839, EPI_ISL_479840, EPI_ISL_479841, EPI_ISL_479842, EPI_ISL_479843, EPI_ISL_479844, EPI_ISL_479845, EPI_ISL_479846, EPI_ISL_479847, EPI_ISL_479848, EPI_ISL_479849 | | | |
| see above | Sapporo City Institute of Public Health | Pathogen Genomics Center, National Institute of Infectious Diseases | Tsuyoshi Sekizuka, Asami Ohnishi, Kentaro Itokawa, Rina Tanaka, Masanori Hashino, Hajime Kamiya, Motoi Suzuki, Makoto Kuroda |
| EPI_ISL_479850, EPI_ISL_479851, EPI_ISL_479852, EPI_ISL_479853, EPI_ISL_479854 | Gunma Prefectural Institute of Public Health and Environmental Sciences | Pathogen Genomics Center, National Institute of Infectious Diseases | Tsuyoshi Sekizuka, Hiroyuki Tsukagoshi, Kentaro Itokawa, Rina Tanaka, Masanori Hashino, Hajime Kamiya, Motoi Suzuki, Makoto Kuroda |
| EPI_ISL_479855, EPI_ISL_479856, EPI_ISL_479857, EPI_ISL_479858, EPI_ISL_479859, EPI_ISL_479860, EPI_ISL_479861 | Department of Infectious Diseases, Kobe Institute of Health | Pathogen Genomics Center, National Institute of Infectious Diseases | Tsuyoshi Sekizuka, Ryohei Nomoto, Kentaro Itokawa, Rina Tanaka, Masanori Hashino, Hajime Kamiya, Motoi Suzuki, Makoto Kuroda |
| EPI_ISL_479862, EPI_ISL_479863, EPI_ISL_479864, EPI_ISL_479865, EPI_ISL_479866, EPI_ISL_479867 | Wakayama Prefectural Research Center of Environment and Public Health | Pathogen Genomics Center, National Institute of Infectious Diseases | Tsuyoshi Sekizuka, Fumio Terasoma, Yosuke Hamajima, Kentaro Itokawa, Rina Tanaka, Masanori Hashino, Hajime Kamiya, Motoi Suzuki, Makoto Kuroda |
| EPI_ISL_479868 | Department of Infectious Diseases, Kobe Institute of Health | Pathogen Genomics Center, National Institute of Infectious Diseases | Tsuyoshi Sekizuka, Ryohei Nomoto, Kentaro Itokawa, Rina Tanaka, Masanori Hashino, Hajime Kamiya, Motoi Suzuki, Makoto Kuroda |
| EPI_ISL_479869 | Niigata Prefectural Institute of Public Health and Environmental Sciences | Pathogen Genomics Center, National Institute of Infectious Diseases | Tsuyoshi Sekizuka, Reiko Arai, Kentaro Itokawa, Rina Tanaka, Masanori Hashino, Hajime Kamiya, Motoi Suzuki, Makoto Kuroda |
| EPI_ISL_479870, EPI_ISL_479871 | Sagamihara City Public Health Research Institute | Pathogen Genomics Center, National Institute of Infectious Diseases | Tsuyoshi Sekizuka, Hiroshi Nakamura, Kentaro Itokawa, Rina Tanaka, Masanori Hashino, Hajime Kamiya, Motoi Suzuki, Makoto Kuroda |
| EPI_ISL_479872, EPI_ISL_479873, EPI_ISL_479874, EPI_ISL_479875, EPI_ISL_479876, EPI_ISL_479877, EPI_ISL_479878, EPI_ISL_479879, EPI_ISL_479880, EPI_ISL_479881, EPI_ISL_479882, EPI_ISL_479883, EPI_ISL_479884, EPI_ISL_479885 | | | |
| see above | Sapporo City Institute of Public Health | Pathogen Genomics Center, National Institute of Infectious Diseases | Tsuyoshi Sekizuka, Asami Ohnishi, Kentaro Itokawa, Rina Tanaka, Masanori Hashino, Hajime Kamiya, Motoi Suzuki, Makoto Kuroda |
| EPI_ISL_479886, EPI_ISL_479887, EPI_ISL_479888, EPI_ISL_479889, EPI_ISL_479890, EPI_ISL_479891, EPI_ISL_479892, EPI_ISL_479893, EPI_ISL_479894, EPI_ISL_479895 | Tokyo Metropolitan Institute of Public Health | Pathogen Genomics Center, National Institute of Infectious Diseases | Tsuyoshi Sekizuka, Kenji Sadamasu, Takashi Chiba, Mami Nagashima, Kentaro Itokawa, Rina Tanaka, Masanori Hashino, Hajime Kamiya, Motoi Suzuki, Makoto Kuroda |
| EPI_ISL_479896, EPI_ISL_479897, EPI_ISL_479898, EPI_ISL_479899, EPI_ISL_479900, EPI_ISL_479901 | Gunma Prefectural Institute of Public Health and Environmental Sciences | Pathogen Genomics Center, National Institute of Infectious Diseases | Tsuyoshi Sekizuka, Hiroyuki Tsukagoshi, Kentaro Itokawa, Rina Tanaka, Masanori Hashino, Hajime Kamiya, Motoi Suzuki, Makoto Kuroda |
| EPI_ISL_479902 | Niigata Prefectural Institute of Public Health and Environmental Sciences | Pathogen Genomics Center, National Institute of Infectious Diseases | Tsuyoshi Sekizuka, Reiko Arai, Kentaro Itokawa, Rina Tanaka, Masanori Hashino, Hajime Kamiya, Motoi Suzuki, Makoto Kuroda |
| EPI_ISL_479903, EPI_ISL_479904, EPI_ISL_479905, | Himeji City Institute of Environment and Health | Pathogen Genomics Center, National Institute of | Tsuyoshi Sekizuka, Kentaro Itokawa, Rina Tanaka, Masanori Hashino, Hajime Kamiya, Motoi Suzuki, Makoto Kuroda |

| | | | |
|---|---|---|---|
| EPI_ISL_479906, EPI_ISL_479907, EPI_ISL_479908, EPI_ISL_479909, EPI_ISL_479910, EPI_ISL_479911, EPI_ISL_479912 | | Infectious Diseases | |
| EPI_ISL_479913, EPI_ISL_479914, EPI_ISL_479915, EPI_ISL_479916, EPI_ISL_479917, EPI_ISL_479918, EPI_ISL_479919, EPI_ISL_479920, EPI_ISL_479921, EPI_ISL_479922, EPI_ISL_479923, EPI_ISL_479924 | | | |
| see above | Niigata City Public Health Research Institute | Pathogen Genomics Center, National Institute of Infectious Diseases | Tsuyoshi Sekizuka, Yurie Takahashi, Kentaro Itokawa, Rina Tanaka, Masanori Hashino, Hajime Kamiya, Motoi Suzuki, Makoto Kuroda |
| EPI_ISL_479925, EPI_ISL_479926, EPI_ISL_479927 | Sakai City Institute of Public Health | Pathogen Genomics Center, National Institute of Infectious Diseases | Tsuyoshi Sekizuka, Tatsuya Miyoshi, Kentaro Itokawa, Rina Tanaka, Masanori Hashino, Hajime Kamiya, Motoi Suzuki, Makoto Kuroda |
| EPI_ISL_479928, EPI_ISL_479929, EPI_ISL_479930, EPI_ISL_479931, EPI_ISL_479932, EPI_ISL_479933, EPI_ISL_479934, EPI_ISL_479935 | Saitama Prefectural Institute of Public Health | Pathogen Genomics Center, National Institute of Infectious Diseases | Tsuyoshi Sekizuka, Hayato Ehara, Kentaro Itokawa, Rina Tanaka, Masanori Hashino, Hajime Kamiya, Motoi Suzuki, Makoto Kuroda |
| EPI_ISL_479936, EPI_ISL_479937, EPI_ISL_479938, EPI_ISL_479939, EPI_ISL_479940, EPI_ISL_479941, EPI_ISL_479942, EPI_ISL_479943 | Ibaraki Prefectural Institute of Public Health | Pathogen Genomics Center, National Institute of Infectious Diseases | Tsuyoshi Sekizuka, Keiko Goto, Kentaro Itokawa, Rina Tanaka, Masanori Hashino, Hajime Kamiya, Motoi Suzuki, Makoto Kuroda |
| EPI_ISL_479944, EPI_ISL_479945, EPI_ISL_479946, EPI_ISL_479947, EPI_ISL_479948, EPI_ISL_479949, EPI_ISL_479950, EPI_ISL_479951, EPI_ISL_479952, EPI_ISL_479953, EPI_ISL_479954, EPI_ISL_479955, EPI_ISL_479956, EPI_ISL_479957, EPI_ISL_479958 | | | |
| see above | Osaka Institute of Public Health | Pathogen Genomics Center, National Institute of Infectious Diseases | Tsuyoshi Sekizuka, Satoshi Hiroi, Saeko Morikawa, Kazushi Motomura, Kentaro Itokawa, Rina Tanaka, Masanori Hashino, Hajime Kamiya, Motoi Suzuki, Makoto Kuroda |
| EPI_ISL_479959, EPI_ISL_479960, EPI_ISL_479961, EPI_ISL_479962, EPI_ISL_479963, EPI_ISL_479964, EPI_ISL_479965 | Tokyo Metropolitan Institute of Public Health | Pathogen Genomics Center, National Institute of Infectious Diseases | Tsuyoshi Sekizuka, Kenji Sadamasu, Takashi Chiba, Mami Nagashima, Kentaro Itokawa, Rina Tanaka, Masanori Hashino, Hajime Kamiya, Motoi Suzuki, Makoto Kuroda |
| EPI_ISL_479966 | Osaka Institute of Public Health | Pathogen Genomics Center, National Institute of Infectious Diseases | Tsuyoshi Sekizuka, Satoshi Hiroi, Saeko Morikawa, Kazushi Motomura, Kentaro Itokawa, Rina Tanaka, Masanori Hashino, Hajime Kamiya, Motoi Suzuki, Makoto Kuroda |
| EPI_ISL_479967, EPI_ISL_479968, EPI_ISL_479969, EPI_ISL_479970, EPI_ISL_479971, EPI_ISL_479972, EPI_ISL_479973, EPI_ISL_479974, EPI_ISL_479975, EPI_ISL_479976, EPI_ISL_479977, EPI_ISL_479978 | | | |
| see above | Fukui Prefectural Institute of Public Health and Environmental Science | Pathogen Genomics Center, National Institute of Infectious Diseases | Tsuyoshi Sekizuka, Miho Toho, Kentaro Itokawa, Rina Tanaka, Masanori Hashino, Hajime Kamiya, Motoi Suzuki, Makoto Kuroda |
| EPI_ISL_479979, EPI_ISL_479980, EPI_ISL_479981, EPI_ISL_479982, EPI_ISL_479983, EPI_ISL_479984 | Oita Prefectural Institute of Public Health and Environmental Science | Pathogen Genomics Center, National Institute of Infectious Diseases | Tsuyoshi Sekizuka, Mari Sasaki, Kentaro Itokawa, Rina Tanaka, Masanori Hashino, Hajime Kamiya, Motoi Suzuki, Makoto Kuroda |
| EPI_ISL_479985 | Ibaraki Prefectural Institute of Public Health | Pathogen Genomics Center, National Institute of Infectious Diseases | Tsuyoshi Sekizuka, Keiko Goto, Kentaro Itokawa, Rina Tanaka, Masanori Hashino, Hajime Kamiya, Motoi Suzuki, Makoto Kuroda |
| EPI_ISL_479986, EPI_ISL_479987, EPI_ISL_479988, EPI_ISL_479989 | Department of Infectious Diseases, Kobe Institute of Health | Pathogen Genomics Center, National Institute of Infectious Diseases | Tsuyoshi Sekizuka, Ryohei Nomoto, Kentaro Itokawa, Rina Tanaka, Masanori Hashino, Hajime Kamiya, Motoi Suzuki, Makoto Kuroda |
| EPI_ISL_479990 | Kitakyushu City Institute of Health and Environmental Sciences | Pathogen Genomics Center, National Institute of Infectious Diseases | Tsuyoshi Sekizuka, Katsuya Obata, Asuka Kikuchi Kentaro Itokawa, Rina Tanaka, Masanori Hashino, Hajime Kamiya, Motoi Suzuki, Makoto Kuroda |
| EPI_ISL_479991, EPI_ISL_479992, EPI_ISL_479993, EPI_ISL_479994, EPI_ISL_479995, EPI_ISL_479996 | Kumamoto City Public Health Research Institute | Pathogen Genomics Center, National Institute of Infectious Diseases | Tsuyoshi Sekizuka, Kaori Tashiro, Kentaro Itokawa, Rina Tanaka, Masanori Hashino, Hajime Kamiya, Motoi Suzuki, Makoto Kuroda |
| EPI_ISL_479997, EPI_ISL_479998, EPI_ISL_479999, EPI_ISL_480000, EPI_ISL_480001 | Nagano Environmental Conservation Research Institute | Pathogen Genomics Center, National Institute of Infectious Diseases | Tsuyoshi Sekizuka, Naoko Shimodaira, Kentaro Itokawa, Rina Tanaka, Masanori Hashino, Hajime Kamiya, Motoi Suzuki, Makoto Kuroda |
| EPI_ISL_480002, EPI_ISL_480003 | Nagasaki Prefectural Institute for Environmental Research and Public Health | Pathogen Genomics Center, National Institute of Infectious Diseases | Tsuyoshi Sekizuka, Fumiaki Matsumoto, Kentaro Itokawa, Rina Tanaka, Masanori Hashino, Hajime Kamiya, Motoi Suzuki, Makoto Kuroda |
| EPI_ISL_480004, EPI_ISL_480005, EPI_ISL_480006, EPI_ISL_480007, EPI_ISL_480008, EPI_ISL_480009, EPI_ISL_480010, EPI_ISL_480011, EPI_ISL_480012, EPI_ISL_480013, EPI_ISL_480014 | | | |
| see above | Chiba Prefectural Institute of Public Health | Pathogen Genomics Center, National Institute of Infectious Diseases | Tsuyoshi Sekizuka, Masakatsu Taira, Kentaro Itokawa, Rina Tanaka, Masanori Hashino, Hajime Kamiya, Motoi Suzuki, Makoto Kuroda |
| EPI_ISL_480015, EPI_ISL_480016, EPI_ISL_480017, EPI_ISL_480018, EPI_ISL_480019, EPI_ISL_480020 | Gunma Prefectural Institute of Public Health and Environmental Sciences | Pathogen Genomics Center, National Institute of Infectious Diseases | Tsuyoshi Sekizuka, Hiroyuki Tsukagoshi, Kentaro Itokawa, Rina Tanaka, Masanori Hashino, Hajime Kamiya, Motoi Suzuki, Makoto Kuroda |
| EPI_ISL_480021, EPI_ISL_480022, EPI_ISL_480023, EPI_ISL_480024, EPI_ISL_480025, EPI_ISL_480026, EPI_ISL_480027, EPI_ISL_480028, EPI_ISL_480029 | Ibaraki Prefectural Institute of Public Health | Pathogen Genomics Center, National Institute of Infectious Diseases | Tsuyoshi Sekizuka, Keiko Goto, Kentaro Itokawa, Rina Tanaka, Masanori Hashino, Hajime Kamiya, Motoi Suzuki, Makoto Kuroda |
| EPI_ISL_480030, EPI_ISL_480031, EPI_ISL_480032, EPI_ISL_480033, EPI_ISL_480034, EPI_ISL_480035, EPI_ISL_480036, EPI_ISL_480037, EPI_ISL_480038, EPI_ISL_480039, EPI_ISL_480040, EPI_ISL_480041 | | | |
| see above | Tochigi Prefectural Institute of Public Health and Environmental Science | Pathogen Genomics Center, National Institute of Infectious Diseases | Tsuyoshi Sekizuka, Ako Nakajima, Kentaro Itokawa, Rina Tanaka, Masanori Hashino, Hajime Kamiya, Motoi Suzuki, Makoto Kuroda |
| EPI_ISL_480042, EPI_ISL_480043, EPI_ISL_480044, EPI_ISL_480045, EPI_ISL_480046, EPI_ISL_480047, EPI_ISL_480048, EPI_ISL_480049, EPI_ISL_480050, EPI_ISL_480051, EPI_ISL_480052, EPI_ISL_480053, EPI_ISL_480054, EPI_ISL_480055, EPI_ISL_480056, EPI_ISL_480057, EPI_ISL_480058, EPI_ISL_480059, EPI_ISL_480060, EPI_ISL_480061, EPI_ISL_480062, EPI_ISL_480063, EPI_ISL_480064 | | | |
| see above | Nagoya City Public Health Research Institute | Pathogen Genomics Center, National Institute of Infectious Diseases | Tsuyoshi Sekizuka, Takuya Miki, Shinichiro Shibata, Kentaro Itokawa, Rina Tanaka, Masanori Hashino, Hajime Kamiya, Motoi Suzuki, Makoto Kuroda |
| EPI_ISL_480065, EPI_ISL_480066, EPI_ISL_480067, EPI_ISL_480068, EPI_ISL_480069, EPI_ISL_480070, EPI_ISL_480071, EPI_ISL_480072 | Sakai City Institute of Public Health | Pathogen Genomics Center, National Institute of Infectious Diseases | Tsuyoshi Sekizuka, Tatsuya Miyoshi, Kentaro Itokawa, Rina Tanaka, Masanori Hashino, Hajime Kamiya, Motoi Suzuki, Makoto Kuroda |
| EPI_ISL_480073 | Tochigi Prefectural Institute of Public Health and Environmental Science | Pathogen Genomics Center, National Institute of Infectious Diseases | Tsuyoshi Sekizuka, Ako Nakajima, Kentaro Itokawa, Rina Tanaka, Masanori Hashino, Hajime Kamiya, Motoi Suzuki, Makoto Kuroda |
| EPI_ISL_480074, EPI_ISL_480075, EPI_ISL_480076, EPI_ISL_480077, EPI_ISL_480078, EPI_ISL_480079, EPI_ISL_480080, EPI_ISL_480081, EPI_ISL_480082 | Shizuoka City Institute of Environmental Sciences and Public Health | Pathogen Genomics Center, National Institute of Infectious Diseases | Tsuyoshi Sekizuka, Takaharu Maehata,Sou Okamura,Yuji Kanazawa,Kenji Yagi, Kentaro Itokawa, Rina Tanaka, Masanori Hashino, Hajime Kamiya, Motoi Suzuki, Makoto Kuroda |
| EPI_ISL_480083, EPI_ISL_480084, EPI_ISL_480085, EPI_ISL_480086, EPI_ISL_480087, EPI_ISL_480088, EPI_ISL_480089 | Gifu Prefectural Institute of Public Health and Environmental Sciences | Pathogen Genomics Center, National Institute of Infectious Diseases | Tsuyoshi Sekizuka, Yoshihiko Kameyama, Kentaro Itokawa, Rina Tanaka, Masanori Hashino, Hajime Kamiya, Motoi Suzuki, Makoto Kuroda |
| EPI_ISL_480090, EPI_ISL_480091, EPI_ISL_480092, EPI_ISL_480093, EPI_ISL_480094, EPI_ISL_480095, EPI_ISL_480096, EPI_ISL_480097, EPI_ISL_480098, EPI_ISL_480099, EPI_ISL_480100, EPI_ISL_480101, EPI_ISL_480102 | | | |
| see above | Department of Infectious Diseases, Kobe Institute of Health | Pathogen Genomics Center, National Institute of Infectious Diseases | Tsuyoshi Sekizuka, Ryohei Nomoto, Kentaro Itokawa, Rina Tanaka, Masanori Hashino, Hajime Kamiya, Motoi Suzuki, Makoto Kuroda |
| EPI_ISL_480103, EPI_ISL_480104, EPI_ISL_480105, EPI_ISL_480106, EPI_ISL_480107, EPI_ISL_480108 | Koshigaya City Public Health Center | Pathogen Genomics Center, National Institute of Infectious Diseases | Tsuyoshi Sekizuka, Yuka Furui, Aya Tamura, Kyohei Sakata, Takumi Daimon, Yoko Togawa, Yoshiko Hamada, Kentaro Itokawa, Rina Tanaka, Masanori Hashino, Hajime Kamiya, Motoi Suzuki, Makoto Kuroda |
| EPI_ISL_480109, EPI_ISL_480110, EPI_ISL_480111, EPI_ISL_480112, EPI_ISL_480113, EPI_ISL_480114, EPI_ISL_480115, EPI_ISL_480116, EPI_ISL_480117, EPI_ISL_480118, EPI_ISL_480119 | | | |

| | | | |
|---|---|---|---|
| see above | Oita Prefectural Institute of Public Health and Environmental Science | Pathogen Genomics Center, National Institute of Infectious Diseases | Tsuyoshi Sekizuka, Mari Sasaki, Kentaro Itokawa, Rina Tanaka, Masanori Hashino, Hajime Kamiya, Motoi Suzuki, Makoto Kuroda |
| EPI_ISL_480120, EPI_ISL_480121, EPI_ISL_480122, EPI_ISL_480123, EPI_ISL_480124, EPI_ISL_480125, EPI_ISL_480126, EPI_ISL_480127, EPI_ISL_480128, EPI_ISL_480129, EPI_ISL_480130, EPI_ISL_480131, EPI_ISL_480132, EPI_ISL_480133, EPI_ISL_480134, EPI_ISL_480135, EPI_ISL_480136, EPI_ISL_480137, EPI_ISL_480138, EPI_ISL_480139, EPI_ISL_480140, EPI_ISL_480141, EPI_ISL_480142, EPI_ISL_480143, EPI_ISL_480144, EPI_ISL_480145, EPI_ISL_480146, EPI_ISL_480147, EPI_ISL_480148, EPI_ISL_480149, EPI_ISL_480150, EPI_ISL_480151, EPI_ISL_480152, EPI_ISL_480153, EPI_ISL_480154, EPI_ISL_480155, EPI_ISL_480156, EPI_ISL_480157, EPI_ISL_480158, EPI_ISL_480159, EPI_ISL_480160, EPI_ISL_480161, EPI_ISL_480162, EPI_ISL_480163, EPI_ISL_480164, EPI_ISL_480165, EPI_ISL_480166, EPI_ISL_480167, EPI_ISL_480168 | | | |
| see above | Fukui Prefectural Institute of Public Health and Environmental Science | Pathogen Genomics Center, National Institute of Infectious Diseases | Tsuyoshi Sekizuka, Miho Toho, Kentaro Itokawa, Rina Tanaka, Masanori Hashino, Hajime Kamiya, Motoi Suzuki, Makoto Kuroda |
| EPI_ISL_480169, EPI_ISL_480170, EPI_ISL_480171, EPI_ISL_480172, EPI_ISL_480173, EPI_ISL_480174, EPI_ISL_480175, EPI_ISL_480176, EPI_ISL_480177 | Gunma Prefectural Institute of Public Health and Environmental Sciences | Pathogen Genomics Center, National Institute of Infectious Diseases | Tsuyoshi Sekizuka, Hiroyuki Tsukagoshi, Kentaro Itokawa, Rina Tanaka, Masanori Hashino, Hajime Kamiya, Motoi Suzuki, Makoto Kuroda |
| EPI_ISL_480178, EPI_ISL_480179 | Hiroshima City Institute of Public Health | Pathogen Genomics Center, National Institute of Infectious Diseases | Tsuyoshi Sekizuka, Kota Noritsune, Kentaro Itokawa, Rina Tanaka, Masanori Hashino, Hajime Kamiya, Motoi Suzuki, Makoto Kuroda |
| EPI_ISL_480180, EPI_ISL_480181, EPI_ISL_480182, EPI_ISL_480183, EPI_ISL_480184, EPI_ISL_480185, EPI_ISL_480186, EPI_ISL_480187, EPI_ISL_480188, EPI_ISL_480189 | Ibaraki Prefectural Institute of Public Health | Pathogen Genomics Center, National Institute of Infectious Diseases | Tsuyoshi Sekizuka, Keiko Goto, Kentaro Itokawa, Rina Tanaka, Masanori Hashino, Hajime Kamiya, Motoi Suzuki, Makoto Kuroda |
| EPI_ISL_480190, EPI_ISL_480191, EPI_ISL_480192, EPI_ISL_480193, EPI_ISL_480194, EPI_ISL_480195 | Ota Health Center Welfare Section | Pathogen Genomics Center, National Institute of Infectious Diseases | Tsuyoshi Sekizuka, Chika Takahashi, Kentaro Itokawa, Rina Tanaka, Masanori Hashino, Hajime Kamiya, Motoi Suzuki, Makoto Kuroda |
| EPI_ISL_480196, EPI_ISL_480197, EPI_ISL_480198, EPI_ISL_480199, EPI_ISL_480200, EPI_ISL_480201, EPI_ISL_480202, EPI_ISL_480203 | Toyama Institute of Health | Pathogen Genomics Center, National Institute of Infectious Diseases | Tsuyoshi Sekizuka, Masae Itamochi, Kazunori Oishi, Kentaro Itokawa, Rina Tanaka, Masanori Hashino, Hajime Kamiya, Motoi Suzuki, Makoto Kuroda |
| EPI_ISL_480204 | Akita City Public Health Center | Pathogen Genomics Center, National Institute of Infectious Diseases | Tsuyoshi Sekizuka, Koichi Ito, Kentaro Itokawa, Rina Tanaka, Masanori Hashino, Hajime Kamiya, Motoi Suzuki, Makoto Kuroda |
| EPI_ISL_480205, EPI_ISL_480206, EPI_ISL_480207, EPI_ISL_480208, EPI_ISL_480209, EPI_ISL_480210, EPI_ISL_480211, EPI_ISL_480212, EPI_ISL_480213, EPI_ISL_480214, EPI_ISL_480215, EPI_ISL_480216, EPI_ISL_480217, EPI_ISL_480218, EPI_ISL_480219, EPI_ISL_480220 | | | |
| see above | Department of Infectious Diseases, Kobe Institute of Health | Pathogen Genomics Center, National Institute of Infectious Diseases | Tsuyoshi Sekizuka, Ryohei Nomoto, Kentaro Itokawa, Rina Tanaka, Masanori Hashino, Hajime Kamiya, Motoi Suzuki, Makoto Kuroda |
| EPI_ISL_480221, EPI_ISL_480222, EPI_ISL_480223 | Koshigaya City Public Health Center | Pathogen Genomics Center, National Institute of Infectious Diseases | Tsuyoshi Sekizuka, Yuka Furui, Aya Tamura, Kyohei Sakata, Takumi Daimon, Yoko Togawa, Yoshiko Hamada, Kentaro Itokawa, Rina Tanaka, Masanori Hashino, Hajime Kamiya, Motoi Suzuki, Makoto Kuroda |
| EPI_ISL_480224 | National Reference Laboratory "Influenza and acute respiratory diseases" | NRL-HIV | Ivan Ivanov, Ivailo Alexiev, Ivva Philipova |
| EPI_ISL_480225 | Fukui Prefectural Institute of Public Health and Environmental Science | Pathogen Genomics Center, National Institute of Infectious Diseases | Tsuyoshi Sekizuka, Miho Toho, Kentaro Itokawa, Rina Tanaka, Masanori Hashino, Hajime Kamiya, Motoi Suzuki, Makoto Kuroda |
| EPI_ISL_480226 | Niigata Prefectural Institute of Public Health and Environmental Sciences | Pathogen Genomics Center, National Institute of Infectious Diseases | Tsuyoshi Sekizuka, Reiko Arai, Kentaro Itokawa, Rina Tanaka, Masanori Hashino, Hajime Kamiya, Motoi Suzuki, Makoto Kuroda |
| EPI_ISL_480227 | Tokyo Metropolitan Institute of Public Health | Pathogen Genomics Center, National Institute of Infectious Diseases | Tsuyoshi Sekizuka, Kenji Sadamasu, Takashi Chiba, Mami Nagashima, Kentaro Itokawa, Rina Tanaka, Masanori Hashino, Hajime Kamiya, Motoi Suzuki, Makoto Kuroda |
| EPI_ISL_480228, EPI_ISL_480229, EPI_ISL_480230, EPI_ISL_480231, EPI_ISL_480232, EPI_ISL_480233, EPI_ISL_480234, EPI_ISL_480235, EPI_ISL_480236, EPI_ISL_480237, EPI_ISL_480238, EPI_ISL_480239, EPI_ISL_480240, EPI_ISL_480241, EPI_ISL_480242, EPI_ISL_480243, EPI_ISL_480244, EPI_ISL_480245, EPI_ISL_480246, EPI_ISL_480247, EPI_ISL_480248, EPI_ISL_480249, EPI_ISL_480250, EPI_ISL_480251, EPI_ISL_480252, EPI_ISL_480253, EPI_ISL_480254, EPI_ISL_480255, EPI_ISL_480256, EPI_ISL_480257, EPI_ISL_480258, EPI_ISL_480259, EPI_ISL_480260, EPI_ISL_480261, EPI_ISL_480262, EPI_ISL_480263, EPI_ISL_480264, EPI_ISL_480265, EPI_ISL_480266, EPI_ISL_480267, EPI_ISL_480268, EPI_ISL_480269, EPI_ISL_480270, EPI_ISL_480271, EPI_ISL_480272, EPI_ISL_480273, EPI_ISL_480274, EPI_ISL_480275, EPI_ISL_480276, EPI_ISL_480277, EPI_ISL_480278, EPI_ISL_480279, EPI_ISL_480280, EPI_ISL_480281, EPI_ISL_480282, EPI_ISL_480283, EPI_ISL_480284, EPI_ISL_480285, EPI_ISL_480286, EPI_ISL_480287, EPI_ISL_480288, EPI_ISL_480289, EPI_ISL_480290, EPI_ISL_480291, EPI_ISL_480292 | | | |
| see above | Genomic Laboratory (GLAB) (Conjoint lab of Health Directorate of Istanbul and Istanbul Technical University) | Genomic Laboratory (GLAB), Istanbul Technical University | Ilker Karacan, Tugba Kizilboga Akgun, Bugra Agaoglu, Gizem Alkurt, Jale Yildiz, Betsi Köse, Elifnaz Çelik, Arzu Irvem, Yasemin Kendir Demirkol, Ozlem Akgun Dogan, Mehtap Aydn, Levent Doganay, Gizem Dinler Doganay |
| EPI_ISL_480293, EPI_ISL_480294, EPI_ISL_480295, EPI_ISL_480296 | Institute for Stem Cell Science and Regenerative Medicine | National Centre for Biological Sciences | Farhan Ali, Vanessa Molin Paynter, Srikar Krishna, Mohak Sharda, Shah-e-Jahan Gulzar, Awadhesh Pandit, Varadha Sundarmurthy, Uma Ramakrishnan, Dasaradhi Palakodeti, Aswin Seshasayee |
| EPI_ISL_480297, EPI_ISL_480298, EPI_ISL_480299, EPI_ISL_480300, EPI_ISL_480301, EPI_ISL_480302, EPI_ISL_480303, EPI_ISL_480304, EPI_ISL_480305, EPI_ISL_480306, EPI_ISL_480307, EPI_ISL_480308, EPI_ISL_480309, EPI_ISL_480310 | | | |
| see above | National Reference Laboratory "Influenza and acute respiratory diseases" | NRL-HIV | Ivan Ivanov, Ivailo Alexiev, Ivva Philipova |
| EPI_ISL_480312, EPI_ISL_480313, EPI_ISL_480314 | Hospital Mexico | Charité Virology-University of Costa Rica | Andres Moreira-Soto, Eugenia Corrales-Aguilar, Ignacio Postigo-Hidalgo, Teresita Somogyi, Jan Felix Drexler |
| EPI_ISL_480315, EPI_ISL_480316, EPI_ISL_480317, EPI_ISL_480318, EPI_ISL_480319, EPI_ISL_480320 | Hospital Clínica Bíblica | Charité Virology-University of Costa Rica | Andres Moreira-Soto, Eugenia Corrales-Aguilar, Ignacio Postigo-Hidalgo, Karla Sofía Gutiérrez, Jan Felix Drexler |
| EPI_ISL_480321 | Laboratorio Clínico San José | Charité Virology-University of Costa Rica | Andres Moreira-Soto, Eugenia Corrales-Aguilar, Ignacio Postigo-Hidalgo, Hugo Núñez Navas, Jan Felix Drexler |
| EPI_ISL_480322, EPI_ISL_480323, EPI_ISL_480324, EPI_ISL_480325, EPI_ISL_480326, EPI_ISL_480327 | Hospital Nacional de Niños | Charité Virology-University of Costa Rica | Andres Moreira-Soto, Eugenia Corrales-Aguilar, Ignacio Postigo-Hidalgo, Cristian Pérez Corrales, Andrei Montero Bonilla, Jan Felix Drexler |
| EPI_ISL_480328 | Laboratorio LABIN | Charité Virology-University of Costa Rica | Andres Moreira-Soto, Eugenia Corrales-Aguilar, Ignacio Postigo-Hidalgo, Ignacio Soto Pacheco, Jan Felix Drexler |
| EPI_ISL_480329, EPI_ISL_480330 | Quadram Institute Bioscience | COVID-19 Genomics UK (COG-UK) Consortium | Dave J. Baker, Gemma L. Kay, Alp Aydin, Thanh Le-Viet, Steven Rudder, Ana P. Tedim, Anastasia Kolyva, Maria Diaz, Leonardo de Oliveira Martins, Nabil-Fareed Alikhan, Lizzie Meadows, Rachael Stanley, Ngozi Elumogo, Muhammed Yasir, Nicholas M. Thomson, Alexander J Trotter, Rachel Gilroy, Samuel Bloomfield, Claire Stuart, Andrew Bell, Reenesh Prakash, Samir Dervisevic, Alison E. Mather, John Wain, Mark Webber, Andrew J. Page, Justin O'Grady |
| EPI_ISL_480331, EPI_ISL_480332, EPI_ISL_480333, EPI_ISL_480334, EPI_ISL_480335, EPI_ISL_480336, EPI_ISL_480337, EPI_ISL_480338, EPI_ISL_480339, EPI_ISL_480340, EPI_ISL_480341, EPI_ISL_480342, EPI_ISL_480343, EPI_ISL_480344, EPI_ISL_480345, EPI_ISL_480346, EPI_ISL_480347, EPI_ISL_480348 | | | |
| see above | Microbial Genomics Laboratory, Institut Pasteur de Montevideo | Microbial Genomics Laboratory, Institut Pasteur de Montevideo | Cecilia Salazar, Marianoel Pereira, Ignacio Ferrés, Gonzalo Moratorio, Pilar Moreno, Gregorio Iraola |
| EPI_ISL_480351, EPI_ISL_480352, EPI_ISL_480353, EPI_ISL_480354, EPI_ISL_480355, EPI_ISL_480356, EPI_ISL_480357, EPI_ISL_480358, EPI_ISL_480359, EPI_ISL_480360, EPI_ISL_480361, EPI_ISL_480362, EPI_ISL_480363, EPI_ISL_480364, EPI_ISL_480365, EPI_ISL_480366, EPI_ISL_480367, EPI_ISL_480368, EPI_ISL_480369, EPI_ISL_480370, EPI_ISL_480371, EPI_ISL_480372, EPI_ISL_480373, EPI_ISL_480375, EPI_ISL_480376, EPI_ISL_480378, EPI_ISL_480379, EPI_ISL_480380, EPI_ISL_480381, EPI_ISL_480382, EPI_ISL_480383, EPI_ISL_480384, EPI_ISL_480385, EPI_ISL_480386, EPI_ISL_480387, EPI_ISL_480388, EPI_ISL_480389, EPI_ISL_480390, EPI_ISL_480391, EPI_ISL_480392, EPI_ISL_480393, EPI_ISL_480394, EPI_ISL_480395, EPI_ISL_480396, EPI_ISL_480397, EPI_ISL_480398, EPI_ISL_480399, EPI_ISL_480400, EPI_ISL_480401, EPI_ISL_480402, EPI_ISL_480403, EPI_ISL_480404, EPI_ISL_480405, EPI_ISL_480406, EPI_ISL_480407, EPI_ISL_480408, EPI_ISL_480409, EPI_ISL_480410, EPI_ISL_480411, EPI_ISL_480412, EPI_ISL_480413 | | | |
| see above | University of Wisconsin-Madison AIDS Vaccine Research Laboratories | University of Wisconsin-Madison AIDS Vaccine Research Laboratories | Gage Moreno, Katarina Braun, et al. AIDS Vaccine Research Laboratories |
| EPI_ISL_480414, EPI_ISL_480415, EPI_ISL_480416, EPI_ISL_480417, EPI_ISL_480418 | National Institute of Laboratory Medicine and Referral Center | Bangladesh Council of Scientific and Industrial Research | Md. Saddam Hossain, Abu Sayeed Mohammad Mahmud, Mohammad Samir Uzzaman, Eshrar Osman, Md. Ahasan Habib, Shahina Akter, Tanjina Akhter Banu, Md. Murshed Hasan Sarkar, Barna Goswami, Iffat Jahan, Tasnim Nafisa, Md. Maruf Ahmed Molla, Mahmuda Yeasmin, Asish Kumar Ghosh, Shahjahan Siddike, A. K. M. Shamsuzzaman, Sheikh Md. Selim Al Din, Utpal Chandra Ray, Salek Ahmed Sajib, Md. Salim Khan |
